# A human commons cell atlas reveals cell type specificity for OAS1 isoforms

**DOI:** 10.1101/2024.03.23.586412

**Authors:** Ángel Galvez-Merchán, A. Sina Booeshaghi, Lior Pachter

**Affiliations:** Cellarity, Somerville, MA, USA; Department of Bioengineering, University of California Berkeley, Berkeley, CA, USA; Division of Biology and Biological Engineering, California Institute of Technology, Pasadena, CA, USA; Department of Computing & Mathematical Sciences, California Institute of Technology, Pasadena, CA, USA

## Abstract

We describe an open source Human Commons Cell Atlas comprising 2.9 million cells across 27 tissues that can be easily updated and that is structured to facilitate custom analyses. To showcase the flexibility of the atlas, we demonstrate that it can be used to study isoforms of genes at cell resolution. In particular, we study cell type specificity of isoforms of OAS1, which has been shown to offer SARS-CoV-2 protection in certain individuals that display higher expression of the p46 isoform. Using our commons cell atlas we localize the OAS1 p44b isoform to the testis, and find that it is specific to round and elongating spermatids. By virtue of enabling customized analyses via a modular and dynamic atlas structure, the commons cell atlas should be useful for exploratory analyses that are intractable within the rigid framework of current gene-centric cell atlases.

## Introduction

The innate immune system plays a crucial role in defending the body against viruses (Takeuchi and Akira 2009; Koyama et al. 2008; Carty, Guy, and Bowie 2021). One of the key components of this system is Oligoadenylate synthetase 1 (OAS1), a protein that gets activated during viral infection through its binding to double-stranded RNA (Melchjorsen et al. 2009). Activated OAS1 produces 2′,5′-oligoadenylates, promoting the activity of RNase L and triggering the degradation of cellular and viral RNAs to halt viral replication (Melchjorsen et al. 2009; Hovanessian and Justesen 2007). Importantly, the last exon of the human OAS1 gene undergoes alternative splicing, leading to the production of isoforms with unique C-terminal sequences (Di, Elbahesh, and Brinton 2020). Because of their distinct antiviral activities (Soveg et al. 2021), the differential expression of OAS1 isoforms correlates with susceptibility to certain viruses (Li et al. 2017). For example, the expression of OAS1-p46 has been shown to provide protection against viral infections such as West Nile virus (Lim et al. 2009), Dengue virus (Lin et al. 2009), Hepatitis C virus, and most recently, SARS-CoV-2 (Zhou et al. 2021). Given the clinical relevance, several studies have investigated the regulation of the expression of OAS1 isoforms, with a particular focus on the impact of various SNPs on the relative isoform abundance (Li et al. 2017). However, no study has explored whether this regulation is organ or cell-type specific. This question is significant: an OAS1 isoform can only protect against a virus if it is expressed in the cell-type the virus infects, and therefore an understanding of cell type specificity of OAS1 isoforms must accompany an analysis of viral tropism.

A single cell atlas is the ideal tool to study cell-type specific OAS1 isoform regulation. There are a number of available cell atlases that together contain data from most human organs, such as the Human Cell Atlas (Li et al. 2017; Rozenblatt-Rosen et al. 2017), the adult human cell atlas (S. He et al. 2020), Tabula Sapiens (Consortium* et al. 2022), the Human Cell Landscape (Han et al. 2020), Descartes (Cao et al. 2020), and Azimuth (Hao et al. 2021). However, our attempts to use these atlases for our research question uncovered three important limitations. First, current atlases exclusively provide gene-level expression data, and lack information on isoforms. This deficiency is not minor, as evidenced by the growing body of literature supporting the critical role of isoform expression in major biological processes (E. T. Wang et al. 2008; Chaponnier and Gabbiani 2004; Warren et al. 2003; Mincarelli et al. 2023), as well as in the identification and definition of cell-types (Booeshaghi et al. 2021). Second, current cell atlas projects depend on assay-specific preprocessing tools that can introduce computationally-induced batch effects that can be challenging to identify and correct. Finally, cell atlases are static objects that cannot be easily updated or re-processed to facilitate interpretation of the stream of new findings emerging from single-cell studies. Our atlas and the infrastructure it is built on address these limitations, and we demonstrate its potential by showing how it can shed light on OAS1 isoform expression at single cell resolution.

## Results

### Building the Humans Commons Cell Atlas

To study the cell-type specificity of OAS1 isoforms in humans, we created a human cell atlas using our recently developed Commons Cell Atlas (CCA) infrastructure. The Human CCA comprises over 2.9 million cells from 525 publicly available scRNA-seq datasets across 27 organs (**Figure 1A**).

**Figure 1.**
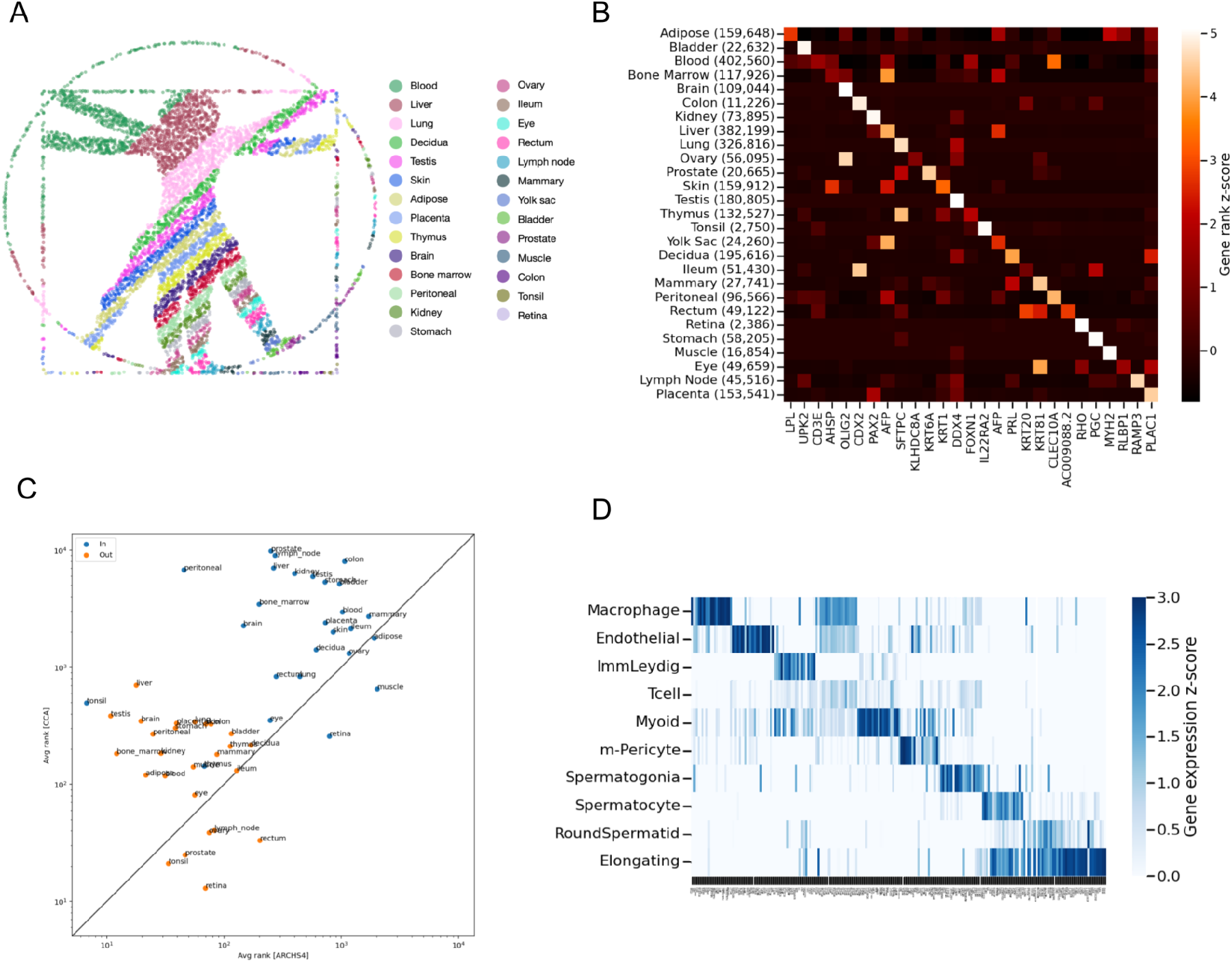
The Human Commons Cell Atlas. **(A)** 2D representation of the atlas. Cells were downsampled to match the number of coordinates of the Vitruvian man maintaining the original proportions by organ. **(B)** Heatmap displaying the z-score of the average rank of select organ marker genes calculated across organs. Elements along the diagonal indicate genes (columns) that mark major cell-types in the organ (rows). **(C)** Correlation between ranks of marker genes in the CCA and ARCHS4 computed on organ-specific marker genes (in, blue) and non-organ specific marker genes (out, red). **(D)** Z-score of average expression of testis marker genes by cell-type.

We started by compiling a list of publicly available single-cell RNA-seq datasets deposited across GEO/SRA/ENA/DDBJ (Barrett et al. 2011; Leinonen, Sugawara, et al. 2011; Leinonen, Akhtar, et al. 2011; Ogasawara et al. 2020). This collection of data consisted of 147 billion sequencing reads (20.74 TB of FASTQs), **Supplementary Table 1**. We chose to start with raw FASTQs, instead of the gene count matrices, which is crucial for 1. ensuring a uniform read alignment strategy and barcode error correction, and 2. enabling isoform quantification, which is lost when counts are aggregated at the gene level. Data and metadata were downloaded and organized by “observation” which maps to GEO sample accessions (GSM) that group a set of FASTQ files with metadata.

Atlas building requires uniform processing and sequencing read quantification to minimize computational variability and apply concordant read alignment, barcode error correction, and read counting. To that end, we leveraged the recently developed kallisto bustools (kb-python) programs (Melsted et al. 2021; Sullivan et al. 2023) to generate all 525 gene count matrices. From the raw sequencing data, the Human Commons Cell atlas was built in about two weeks (305 hours) (**Figure S1**) with less than 8GB of memory usage. Reads were pseudoaligned to the human transcriptome, cell barcodes were corrected within hamming-1 distance of a barcode “onlist”, and naïve UMI collapsing was performed to generate gene counts (see Methods, **Supplementary Knee**).

### Gene-level Human CCA

In order to study the expression of OAS1 we first sought to assess the suitability and robustness of the Human CCA for computationally identifying tissue-level marker genes. We first performed differential expression between 28 tissues on the rank of all genes and identified tissue-level markers from the list of DE genes. We identified genes that are highly and specifically expressed in tissues, most of them representing bona-fide markers for each organ. The list of genes include the LPL gene (encoding Lipoprotein Lipase) in adipose tissue (Zechner et al. 2000), the RLBP1 gene (encoding Retinaldehyde Binding Protein 1) in eye (Eichers et al. 2002), and the SFPTC gene (encoding pulmonary-associated surfactant protein C) in lung (Tredano et al. 2004) (**Figure 1B**). These findings serve as a positive control on the atlas’s ability to join tissue-level groupings and prior known marker genes.

We then assessed the concordance of gene-level pseudobulk quantifications produced with CCA to bulk gene-level bulk quantifications in ARCHS4 (Lachmann et al. 2018) across 27 organs. We found a higher concordance of ranked-pseudobulk profiles for those genes that mark organs, than those that do not (Methods). This demonstrates that pseudobulk qualifications for organ-specific marker genes computed from the CCA are consistent with those found in ARCHS4 (**Figure 1C**).

We then sought to identify if the OAS1 gene exhibits tissue-level specificity. As expected given its crucial function, OAS1 was detected in most tissues, with no significant enrichment in any of them (**Figure 2A**). Notably, OAS1 was upregulated in lung samples from COVID-19 infected individuals, which is consistent with OAS1 being a type I interferon (IFN)-induced gene (Melchjorsen et al. 2009) (Figure **2B**).

**Figure 2.**
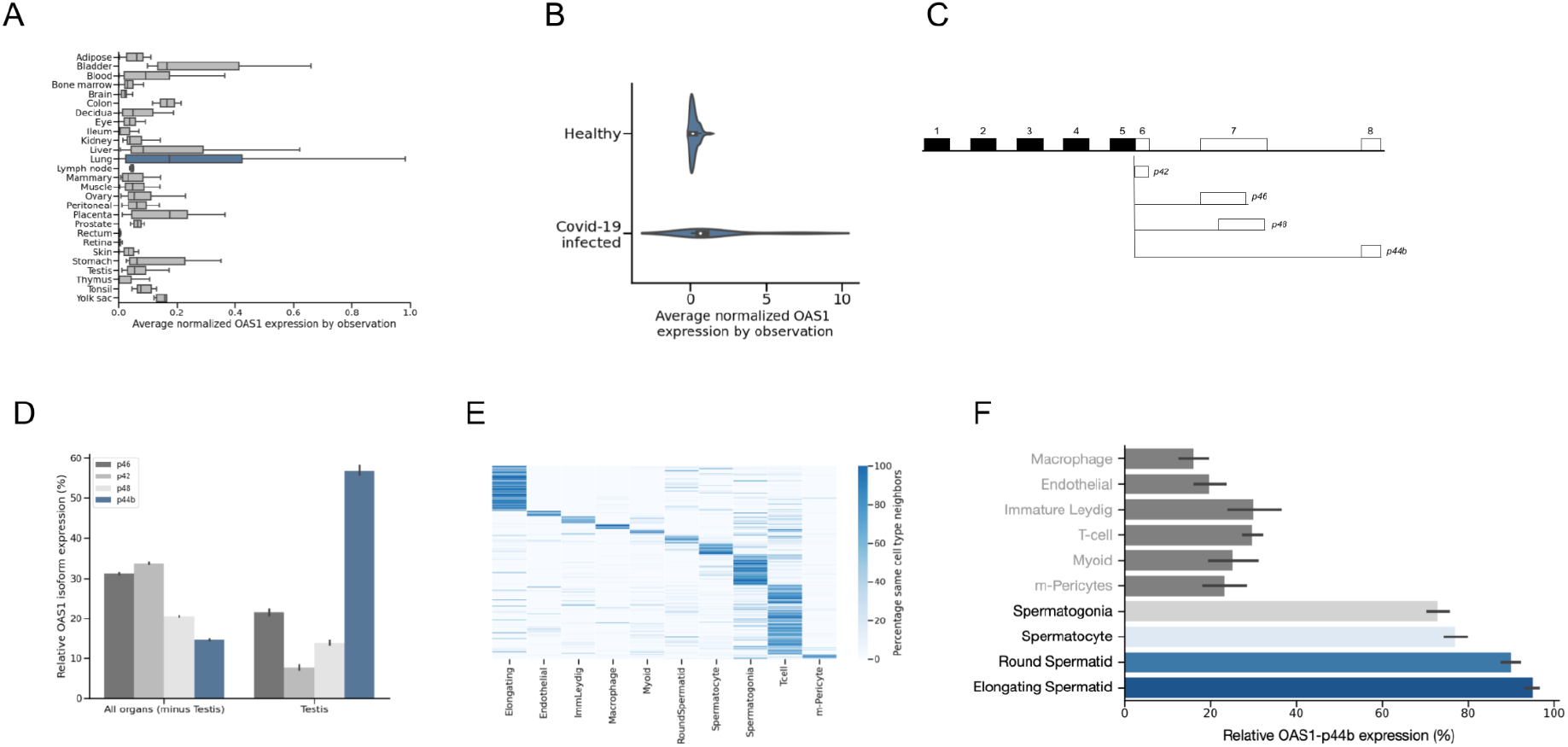
Cell-type specificity of OAS1 isoforms. (A) Normalized OAS1 gene expression by organ. Results for the lung, the organ with highest expression, are highlighted in blue. (B) Normalized OAS1 expression in Lung datasets by health status of the individual. (C) Diagram of the 4 main OAS1 isoforms we found in our data. (D) Relative OAS1 isoform expression in all organs except testis, and testis. OAS1-p44b is highly and specifically expressed in testis. (E) Validation of testis cell-type assignments. The clusters indicate that cells from the same cell-type are frequently neighbors in the K-Nearest Neighbor Graph. (F) OAS1-p44b is the main OAS1 isoform in spermatogenesis cells.

### Isoform-level Human CCA

The expression of isoforms within genes can vary greatly, even when the overall gene expression remains unchanged (Booeshaghi et al. 2021). This information is lost in currently published atlases, which fail to quantify transcript isoforms. We hypothesized that OAS1 isoforms exhibited tissue-level specificity. To test this hypothesis, we rebuilt our atlas at the isoform level leveraging transcript compatibility counts (See Methods) (Booeshaghi et al. 2021; Ntranos et al. 2019). Since most of the publicly available single-cell data derives from 3’ technologies, our quantification was limited to isoforms with distinct 3’ ends.

### OAS1 isoform cell-type specificity

We leveraged the unique 3’UTR of OAS1 isoforms to study their differential expression (**Figure 2C**). We found that, out of the 11 OAS1 isoforms annotated in the human transcriptome, only 4 of them were significantly expressed across the atlas: p46, p42, p48 and p44-b (**Figure S2**). We failed at detecting cell-type specificity of the protective isoform p46 (Zhou et al. 2021), which exhibited broad expression across all organs and was among the most highly expressed isoforms alongside p42.The least expressed isoform, p44-b, was responsible for only around 10% of the total OAS1 expression. This is in line with numerous studies consistently showing that amplifying the p44-b isoform by qPCR is difficult due to its very low or undetectable expression level (Noguchi et al. 2013; Iida et al. 2021). Interestingly, we found that this trend was reversed in testis, with p44b accounting for almost 60% of total OAS1 expression and being the predominant isoform (**Figure 2D**).

To investigate if OAS1-p44b expression was cell-type specific, we assigned cell-types using ‘mx assign’. We validated the output of ‘mx assign’ by calculating, for each cell, the percentage of cells belonging to the same cell-type within their 20-nearest neighbors (Z. Zhang 2016). Cells from the same cell-type clustered together, validating our assignment results (**Figure 2E**). We observed that the high OAS1-p44b expression was specific to germ cells undergoing spermatogenesis, where p44b represented over 80% of total OAS1 expression (**Figure 2F**). To discard any possible artifacts caused by pseudoalignment, we visualized the alignments of one of the testis samples (GSM3302525) and observed high density of reads mapping to OAS1’s exon 8, which is unique to the isoform p44b (**Figure S3**). This result was not sample or paper-dependent, with Round Spermatids across all testis samples expressing high levels of OAS1-p44b (**Figure S4**).

### Screen for cell-type specific isoform switching

To confirm that the atlas had accurate isoform information, compared isoforms quantified with ARCHS4 to those quantified with CCA that: i) derived from genes with more than one isoform, ii) had reads in testis samples that mapped uniquely to it and iii) had a minimum average normalized expression of 0.002 per cell. 20,768 isoforms passed these filters in testis, and were therefore amenable to quantification. We performed a bulk isoform-to-isoform comparison between ARCHS4 and CCA (Methods) for cells assayed from the testis and found a Spearman correlation of 0.71 pointing to concordance of pseudobulk single-cell profiles to bulk profiles (**Figure 3A**).

**Figure 3.**
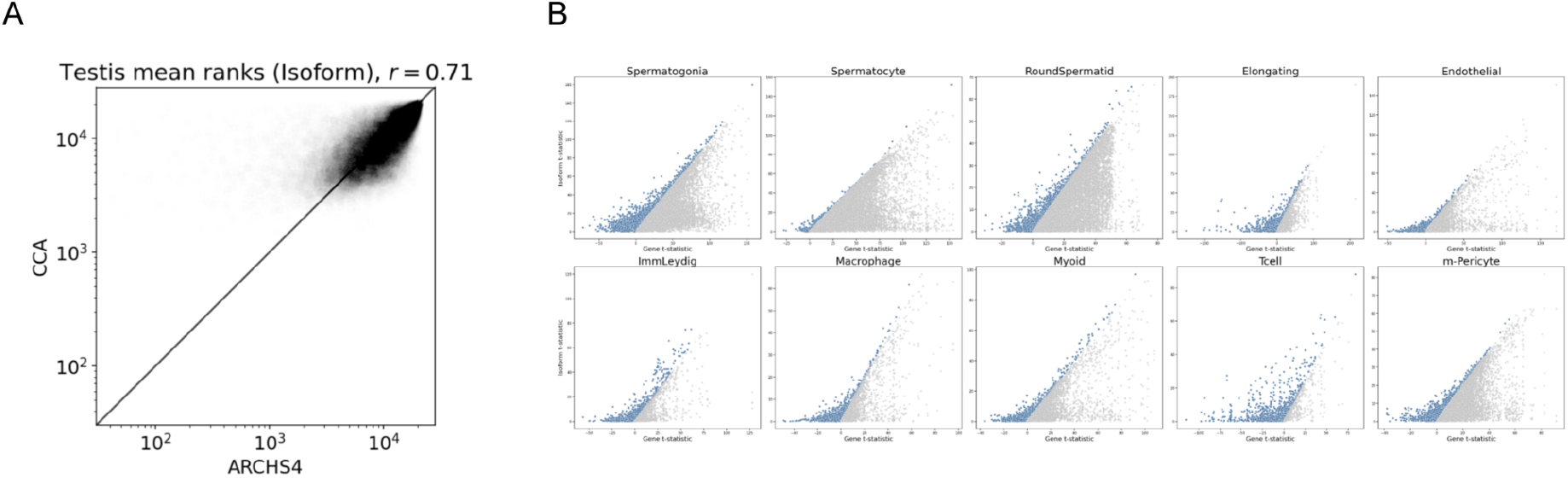
Quantifying isoforms with distinct 3’ UTRs. (A) Scatterplot of isoform ranks in the testis as measured according to CCA and ARCHS4. (B) Examples of cell-type specific isoform switches.

After confirming that the human CCA can accurately quantify isoforms with distinct 3’ UTRs, we decided to test how many isoforms were amenable to this analysis in our data. Approximately 27% of genes with more than one isoform had at least 80% of their counts in single equivalence classes (**Figure S5**), suggesting that for a relatively large fraction of genes, isoform quantifications can be meaningfully utilized. Indeed, performing differential expression of both isoforms and genes in the cell-types of the testis revealed a number of isoforms whose DE signal was stronger than that of the gene it belonged to (**Figure 3B**). We decided to leverage this additional layer of information and screen for cell-type specific isoform switching within testis. We found genes with differential isoform usage in all cell-types except for T-cells. Some of the hits, such as IFT27 in Spermatocytes (Y. Zhang et al. 2017), have been shown to have essential roles within the testis (**Figure S6**). Our results suggest that some of these roles may be carried out by specific isoforms, and that isoform switching may be an important regulatory mechanism in a number biological processes.

## Discussion

The question of how to organize a single-cell atlas is complex, and there is little agreement on even the most basic questions, such as what constitutes a “cell-type”. In addition to conceptual problems, engineering challenges abound. Single-cell genomics datasets are growing in both number and size, and it is non-trivial to engineer atlases so that they can be updated when new datasets or biology are discovered. As we have demonstrated, the Commons Cell Atlas concept offers a solution to many of these problems by virtue of reframing single-cell atlases as a dynamic collection of data and, crucially, tools for processing, querying and interpreting the data. Our isoform analysis was possible thanks to this design principle; in order to obtain isoform quantifications we needed to reprocess the data several times. Other questions may demand alternative atlas processing that, with the Commons Cell Atlas architecture, should be tractable to implement. Most importantly, the Commons Cell Atlas principle dictates that there is no definitive Commons Cell Atlas, but rather numerous Commons Cell Atlases that are customized and specific to the questions being explored.

One of the main aims of cell atlases is to provide a comprehensive characterization and classification of cells (Rozenblatt-Rosen et al. 2017), which relies heavily on identifying their cell-type. The cell-type assignments within the Commons Cell Atlas can be constantly updated, thereby enabling continual refinement of the derived results as new information emerges. To show that this is a feasible strategy, we obtained all marker genes from single cell publications whose data was not used to build the atlas. Despite data and marker genes coming from different sources, we were able to successfully assign cells from most tissues in the atlas (**Figure S7**), as measured by the percentage of same-cell-type neighbors (as shown in **Figure 2** for testis). We expect that these assignments will change as we learn more about each tissue, enabled by the dynamism and efficiency of the Common Cell Atlas infrastructure.

The pivotal function of OAS1 in innate immunity has been well-established, and recent studies have demonstrated that differential expression of its isoforms can affect susceptibility to viruses, including SARS-CoV-2 (Zhou et al. 2021). The work described here enables the study of this differential expression across organs and cell-types, providing a valuable resource for our understanding of viral immunity. We have used our Human Commons Cell Atlas to discover previously unknown cell-type specificity for an OAS1 isoform. Interestingly, this isoform was deemed undetectable across many studies, and our atlas provides an explanation for these results. Our finding is preliminary, and a comprehensive assessment of p44b function and activity in the testis is beyond the scope of our paper. But our discovery highlights the utility of a Commons Cell Atlas, and more generally, points towards the importance of carefully assessing isoform cell-type specificity.

The Human Commons Cell Atlas is composed of 27 tissues, 526 cell-types and 3,554 marker genes. The whole atlas is hosted on GitHub, and it is therefore readily available to download, inspect, modify and use. We envision that the Human Commons Cell Atlas will evolve as we increase our knowledge on tissues and cell-types, moving away from the idea of achieving a “final” or “complete” atlas. For as long as we continue discovering new cell-types, developing new single cell technologies, and gathering new single cell data, the Human Commons Cell Atlas and all its derived results can continue to grow and improve.

## Supporting information

Supplementary Knee

## Acknowledgements

We thank Tara Chari for assistance in producing **Figure 1A**. The authors acknowledge the Howard Hughes Medical Institute for funding A.S.B. through the Hanna H. Gray Fellows program. Thanks to the Caltech Bioinformatics Resource Center for assisting with pre-processing the data.

## Methods

### Downloading data

Accession ids for scRNA-seq datasets were obtained using (Svensson, da Veiga Beltrame, and Pachter 2020). Metadata and links to raw data were collected using the ffq program (Gálvez-Merchán et al. 2022) version 0.2.1 (available at https://github.com/pachterlab/ffq) by running ‘ffq DATASETID’ where DATASETID is the sample ID (GSM, SRS, etc.).

### Preprocessing data

Reads from each dataset were pseudoaligned to the human transcriptome, which was obtained by running ‘*kb ref -i index.idx -g t2g.txt -d human*’. Reads were uniformly processed using the *kallisto* | *bustools* python wrapper *kb-python*, running the command ‘*kb count -i index.idx -g t2g.txt -x [technology]*’. The datasets used in the atlas are listed below:

GSE68596 (Kind et al. 2015),

GSE130636 (Voigt et al. 2019),

GSE96583 (Kang et al. 2018)

GSE112013 (Guo et al. 2022)

GSE130228 (Xu et al. 2022)

GSE130973 (Solé-Boldo et al. 2020)

GSE129363 (Vijay et al. 2020)

SRP126175 (Sivakamasundari et al. 2017)

GSE131391 (Duclos et al. 2019)

GSE125527 (Boland et al. 2020)

GSE165860 (King et al. 2021)

GSE143704 (Fitzgerald et al. 2023)

GSE114297 (Sui et al. 2021)

GSE148963 (Leir et al. 2020)

GSE129363 (Vijay et al. 2020)

GSE131685 (Su et al. 2021)

GSE157526 (Fiege et al. 2021)

PRJCA002413 (Wen et al. 2020)

GSE128889 (Merrick et al. 2019)

GSE129845 (Yu et al. 2019)

GSE104600 (Das et al. 2018)

GSE107747 (D’Avola et al. 2018)

GSE111014 (Rendeiro et al. 2020)

GSE115189 (Freytag et al. 2018)

GSE119594 (Schiroli et al. 2019)

GSE124334 (Galinato et al. 2019)

GSE124898 (Borcherding et al. 2023)

GSE128066 (Sun et al. 2019)

GSE130117 (Richer et al. 2020)

GSE130430 (Yang et al. 2019)

GSE117824 (Nam et al. 2019)

PRJNA610059 (Turner et al. 2020)

GSE115149 (Jitschin et al. 2019)

GSE120446 (Oetjen et al. 2018)

GSE130430 (Yang et al. 2019)

GSE108571: No associated publication

GSE119212 (Zhong et al. 2020)

GSE130238 (Trujillo et al. 2019)

EMTAB6701 (Vento-Tormo et al. 2018)

GSE145926 (Liao et al. 2020)

GSE146188 (van Zyl et al. 2020)

GSE112570 (Lindström et al. 2018)

GSE114530 (Mircea et al. 2022)

GSE117211 (Kumar et al. 2019)

GSE119561 (Howden et al. 2019)

GSE130073 (Ouchi et al. 2019)

GSE103918 (McCauley et al. 2018)

GSE102592 (C. Wang et al. 10 2018)

GSE121600 (Ruiz García et al. 2019)

GSE135893 (Habermann et al. 2020)

GSE134174 (Goldfarbmuren et al. 2020)

GSE124494 (Takeda et al. 2019)

GSE11472 (Azizi et al. 2018)

GSE123926 (Merino et al. 2019)

GSE118127 (Fan et al. 2019)

GSE130888 (Kind et al. 2015)

PRJNA492324: No associated publication

GSE130318: (McCray et al. 2019)

GSE128066 (Sun et al. 2019)

GSE109037 (Hermann et al. 2018)

GSE124263 (Sohni et al. 2019)

GSE130151 (Laurentino et al. 2019)

GSE119506 (Durand et al. 2019)

GSE139522 (Durante et al. 2020)

EMTAB7407(Popescu et al. 2019)

GSE120508 (Guo et al. 2022)

GSE115469 (Sigiel et al. 1978)

EMTAB8581 (Park et al. 2020)

GSE125970 (Y. Wang et al. 2020)

### Filtering matrices

To filter the gene count matrices, barcodes with low UMIs were filtered with ‘*mx filter*’, which uses a derivation of the knee plot approach. Barcodes with more than 40% mitochondrial genes were discarded.

### Normalizing matrices

Gene count matrices were normalized using ‘*mx norm*’, which uses the log1pPF method (Sina Booeshaghi et al. 2022).

### Marker genes curation

A list of marker genes for each organ was generated from supplementary tables of single cell publications containing differential expression information. Markers were selected applying filters to the corrected p-value, log fold change, and percentage of cells expressing the gene. A Google Colab notebook that downloads the supplementary table, filters it, and generates a markers file is available at the Human Cell Atlas repository for each organ. We curated markers for the following tissues:

Adipose: (Emont et al. 2022)

Bladder: (Yu et al. 2019)

Blood:(Jiang et al. 2023)

Bone: (J. He et al. 2021)

Bone marrow: (Jiang et al. 2023)

Brain: (Jiang et al. 2023)

Colon:(Jiang et al. 2023)

Decidua: (Jiang et al. 2023)

Eye: (Gautam et al. 2021)

Heart: (Tucker et al. 2020)

Ileum: (Tucker et al. 2020)

Kidney: (Wu et al. 2018)

Liver: (Guilliams et al. 2022)

Lung: (Adams et al. 2020)

Lymph node: (Jiang et al. 2023)

Mammary: (Jiang et al. 2023)

Muscle: (Jiang et al. 2023)

Ovary: (Wagner et al. 2020)

Pancreas: (Segerstolpe et al. 2016)

Peritoneal: (Jiang et al. 2023)

Placenta: (Liu et al. 2018)

Prostate: (Jiang et al. 2023)

Rectum: (Jiang et al. 2023)

Retina: (Jiang et al. 2023)

Skin: (Jiang et al. 2023)

Stomach: (Jiang et al. 2023)

Testis: (Shami et al. 2020)

Thymus: (Jiang et al. 2023)

Tonsil: (Jiang et al. 2023)

Uterus: (Legetth et al. 2021)

Yolk sac: (Jiang et al. 2023)

### Cell-type assignment and validation

Cell-types were assigned running ‘*mx assign*’ on each individual dataset using the marker gene file generated above. Assignments were validated by calculating, for each cell, the percentage of cells that belong to the same cell-type in the k-nearest neighbors (KNN) graph, with k=20, within each dataset. The KNN graph was calculated using the normalized expression values of the union of marker genes of the corresponding organ.

### 2D representation of the atlas

To create a 2D latent space in which cells from the same organ are neighbors (**Figure 1A**), we used MCML (Chari and Pachter 2023) with the fracNCA parameter set to 1 (this is, optimizing only the Neighborhood Component Analysis (NCA) loss). We then calculated the pairwise distances of each cell’s 2D coordinates to the Vitruvian man 2D coordinates, using the L_1 norm or Manhattan distance. The distances were used as input to the *scipy* ‘*linear_sum_assignment*’ function to map the 2D latent space to the 2D shape coordinates, assigning each cell coordinate to a shape coordinate while minimizing the total cost or distance (as per the distance matrix).

### Organ-level markers

For each dataset, we calculated the average expression of each gene across all cells, and used that value to rank each gene in the dataset. The gene ranks of datasets from the same organ were averaged, resulting in an organ x gene matrix. Genes with high value in one organ and low in the others were selected, and the Z-scores across organs were plotted using a heatmap.

### OAS1 gene expression by tissue

The average normalized OAS1 expression for each dataset was calculated across all cells. Each data point in **Figure 2A**,**B** corresponds to a different dataset.

### Isoform quantification

A transcript Compatibility Counts (TCC) matrix for each sample was obtained by running ‘*bustools count*’ without the ‘--genecount’ option on the bus files generated after pseudoaligning the raw reads. Transcript abundances were quantified using the EM algorithm by running ‘*kallisto quant-tcc*’ on the TCC matrices. The transcript abundance matrix of each sample was normalized within each cell-type using log1pPF. The normalized matrix was then subsetted to isoforms that i) derived from genes with more than one isoform, ii) had reads in the samples that mapped uniquely to it and iii) had a minimum average normalized expression of 0.002 per cell.

### Isoforms screen

We performed t-tests of the expression of isoforms and genes in each cell-type vs all other cells. Isoforms with higher t-statistics than its corresponding gene were interpreted as containing isoform-specific cell-type signal.

### Percentage of genes with at least 80% counts in single equivalence classes

The raw TCC matrix of the testis sample GSM2928378 was filtered to counts that mapped to a single equivalence class, which were averaged across all cells. A isoform-to-gene ratio was calculated by dividing the result of each isoform by the averaged raw counts of the corresponding gene expression. The number of isoforms with a ratio higher than 0.8 was calculated.

### CONCORDEX ratio

The metric was calculated as previously shown (Jackson et al. 2023) for each sample of the human CCA where gene markers were available.

### ARCHS4 Gene Comparison

ARCHS4 quantifications were ranked across genes for each observation and the mean rank was computed across all observations. CCA pseudobulk profiles were computed across all cells in an observation for each organ and genes were ranked for each observation. The mean rank was computed across all observations for each organ. Ranks for marker genes and non-marker genes in CCA and ARCHS4 were plotted for each organ.

### ARCHS4 Isoform Comparison

We first took ARCHS4 quantifications in testis across 695 samples and filtered out samples with less than 1e7 counts. We then ranked all of the isoforms for each sample across the remaining 18 samples and took the average rank across the samples. We performed a similar computation with all 25 samples from the testis for the CCA data but first generated pseudobulk isoform profiles for each sample from single cells. Then the isoforms were ranked for each sample and the mean taken across all samples. A Spearman correlation was computed.

### Data and code availability

The code and data needed to reprocess the results of this manuscript can be found here https://github.com/pachterlab/GBP_2024/. The CCA GitHub can be found here https://github.com/cellatlas/human/. A summary of the datasets in the CCA can be found here https://cellatlas.github.io/human/.

## Author contributions

The CCA atlas concept emerged from an initiative by ASB to uniformly preprocess the datasets in (Svensson, da Veiga Beltrame, and Pachter 2020). ÁGM conceived of the idea of examining the OAS1 isoforms at single-cell resolution across human tissues after the publication of (Zhou et al. 2021). ASB conceived the CCA structure and associated mx toolkit. ÁGM pre-processed the CCA datasets, and ASB wrote mx. ÁGM and ASB developed the CCA quality control. ÁGM led the OAS1 analysis, with help from ASB and LP. ASB developed the ‘mx assign’ cell assignment approach, and ÁGM and ASB benchmarked it. ÁGM drafted the initial version of the manuscript, which was edited and reviewed by all authors.

