## Supplementary Information for "A human commons cell atlas reveals cell type specificity for OAS1 isoforms"

**Supplementary Figure 1:**

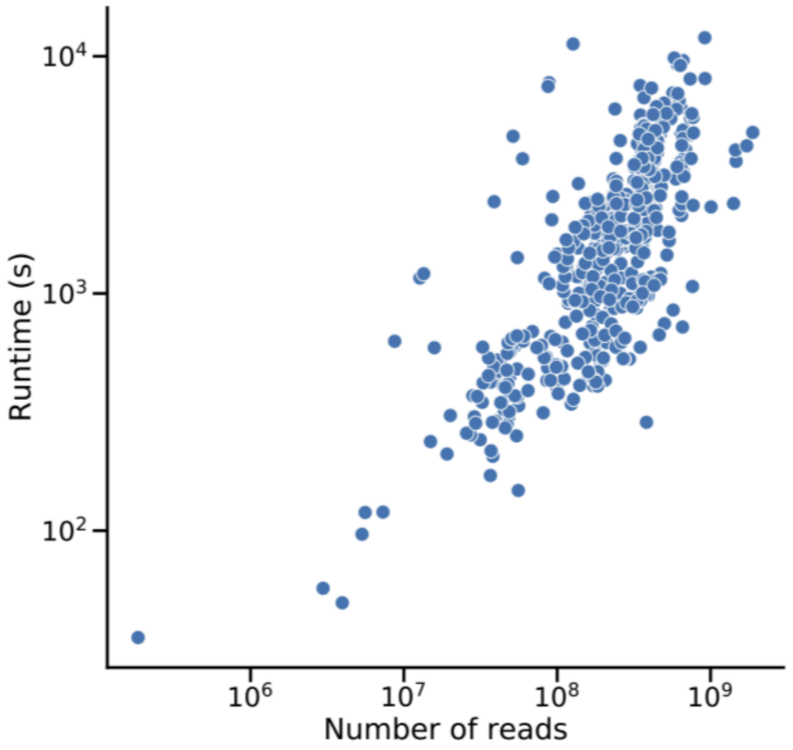


**Figure S1:** Scatterplot showing the relationship between the number of reads in the raw FASTQ and the pre-processing runtime in seconds for each sample of the human CCA. The runtime refers to the '*kb count*' command, which calls the following commands in the following order: '*kallisto bus*', '*bustools inspect*', '*bustools sort*', '*bustools inspect*', '*bustools inspect*', '*bustools correct*', '*bustools inspect*', '*bustools sort*', '*bustools inspect*', '*bustools count*', '*bustools whitelist*', '*bustools correct*', '*bustools inspect*', '*bustools sort*', '*bustools inspect*', '*bustools count*'.

**Supplementary Figure 2:**

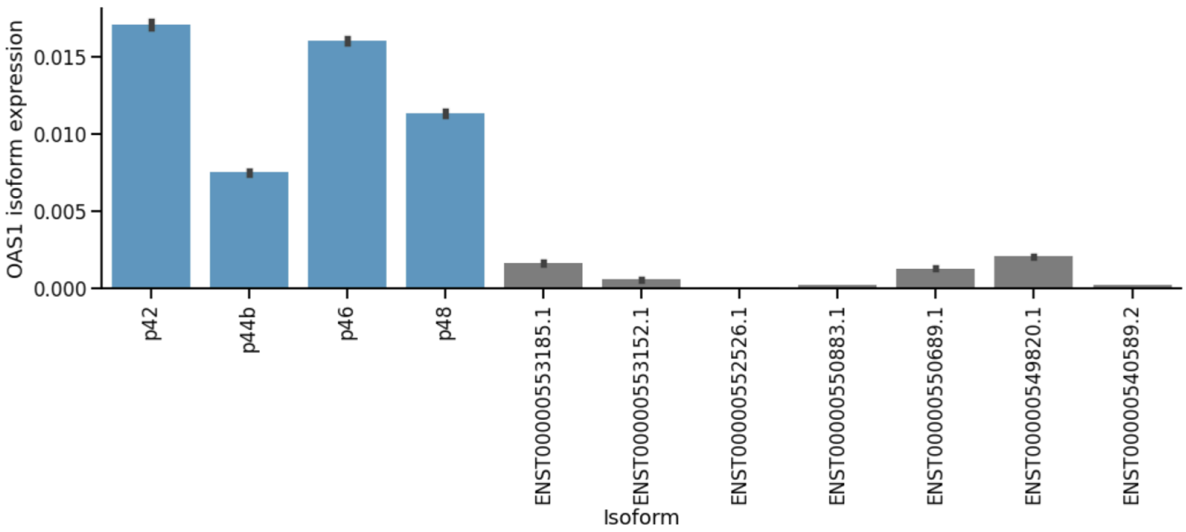


**Figure S2**: Normalised expression of OAS1 isoforms across all cells in the human CCA. The most widely expressed isoforms (p42, p44b, p56 and p48) are highlighted in blue.

**Supplementary Figure 3:**

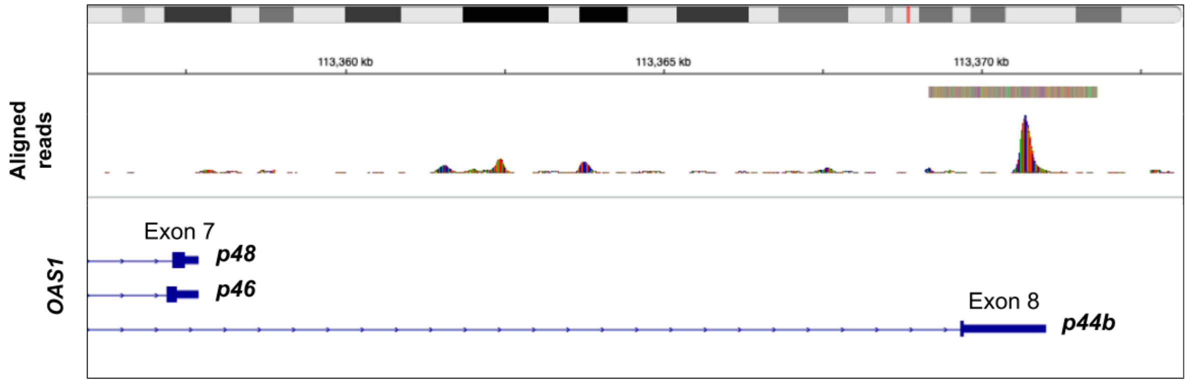


**Figure S3**: The BAM file for one of the testis observations (GSM3302525) was downloaded from GEO and visualized using the Integrative Genomics Viewer (IGV). Over 500 reads mapping to exon 8 and the p44b-specific intron were detected.

**Supplementary Figure 4:**

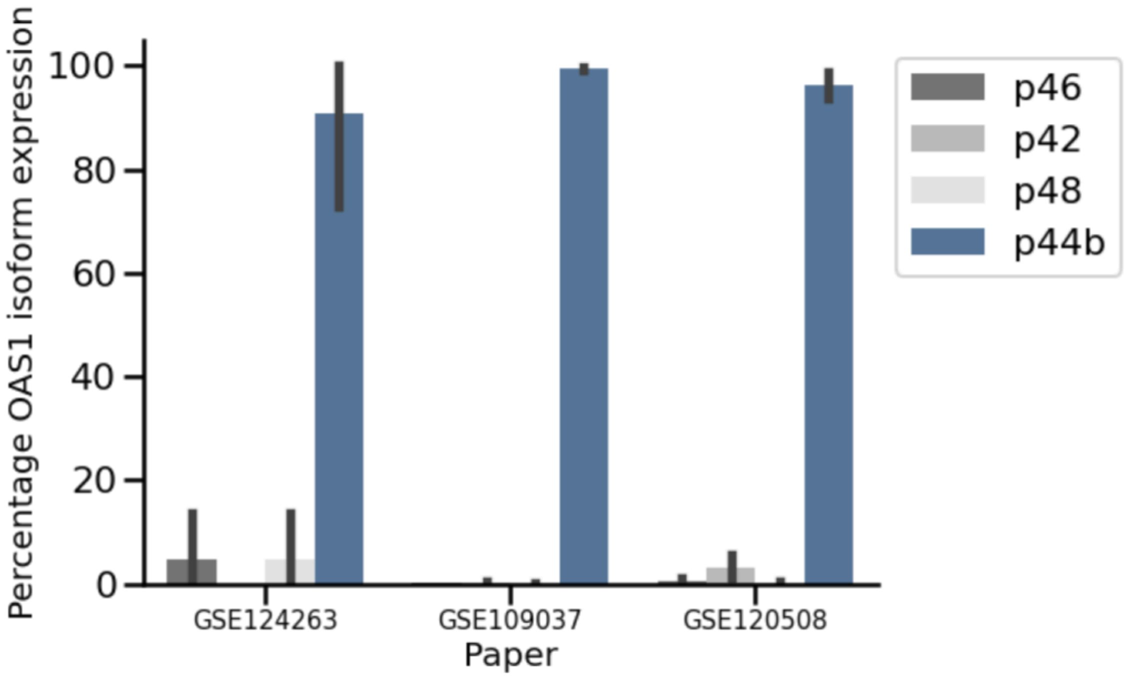


**Figure S4**: Isoform expression of the 4 main OAS1 isoforms in Round Spermatids split by paper, showing that the expression specificity of the isoform p44b reproduces across independent studies and datasets.

**Supplementary Figure 5:**

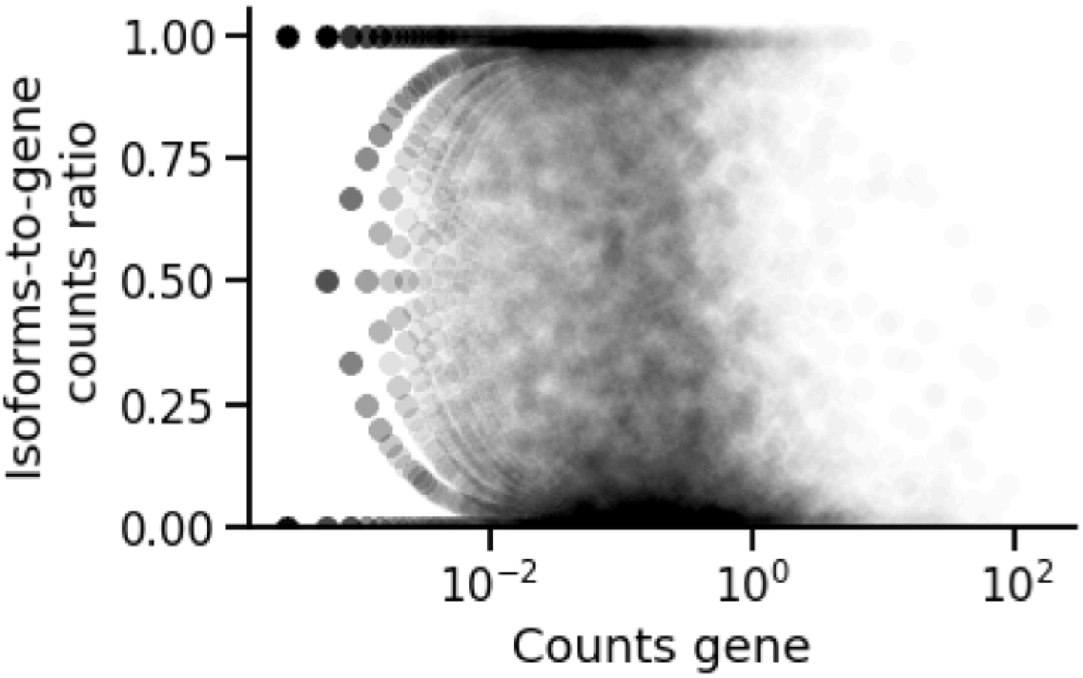


**Figure S5**: The raw TCC matrix of the testis sample GSM2928378 was filtered to counts that mapped to a single equivalence class, which were averaged across all cells. A isoform-to-gene ratio was calculated by dividing the result of each isoform by the averaged raw counts of the corresponding gene expression. For each isoform, this ratio was plotted against the averaged raw counts of the corresponding gene.

**Supplementary Figure 6:**

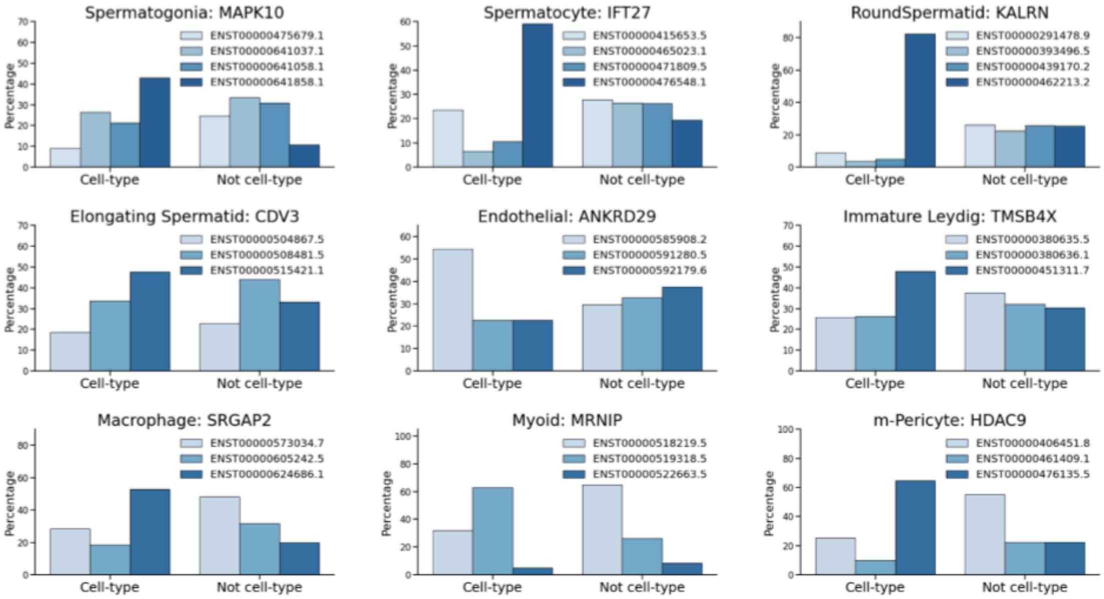


**Figure S6**: Genes that experience cell-type-specific isoform switching were manually selected, and the mean isoform expression for cells belonging or not to the cell-type was plotted.

**Supplementary Figure 7:**

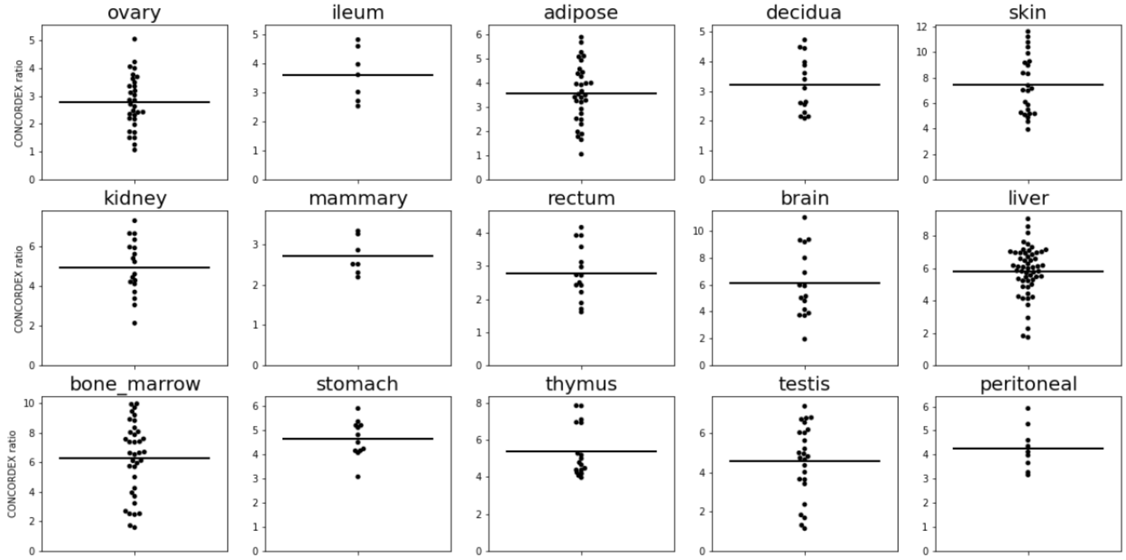


**Figure S7**: The CONCORDEX ratio (citation) of each sample was calculated using the assignments derived from ‘mx assign’. The gene markers for each organ were validated by calculating the average CONCORDEX ratio per organ. Organs with high CONCORDEX ratio are shown in the plot.
