## Supplementary Knee for "A human commons cell atlas reveals cell type specificity for OAS1 isoforms"

### 1 Algorithms for a Commons Cell Atlas

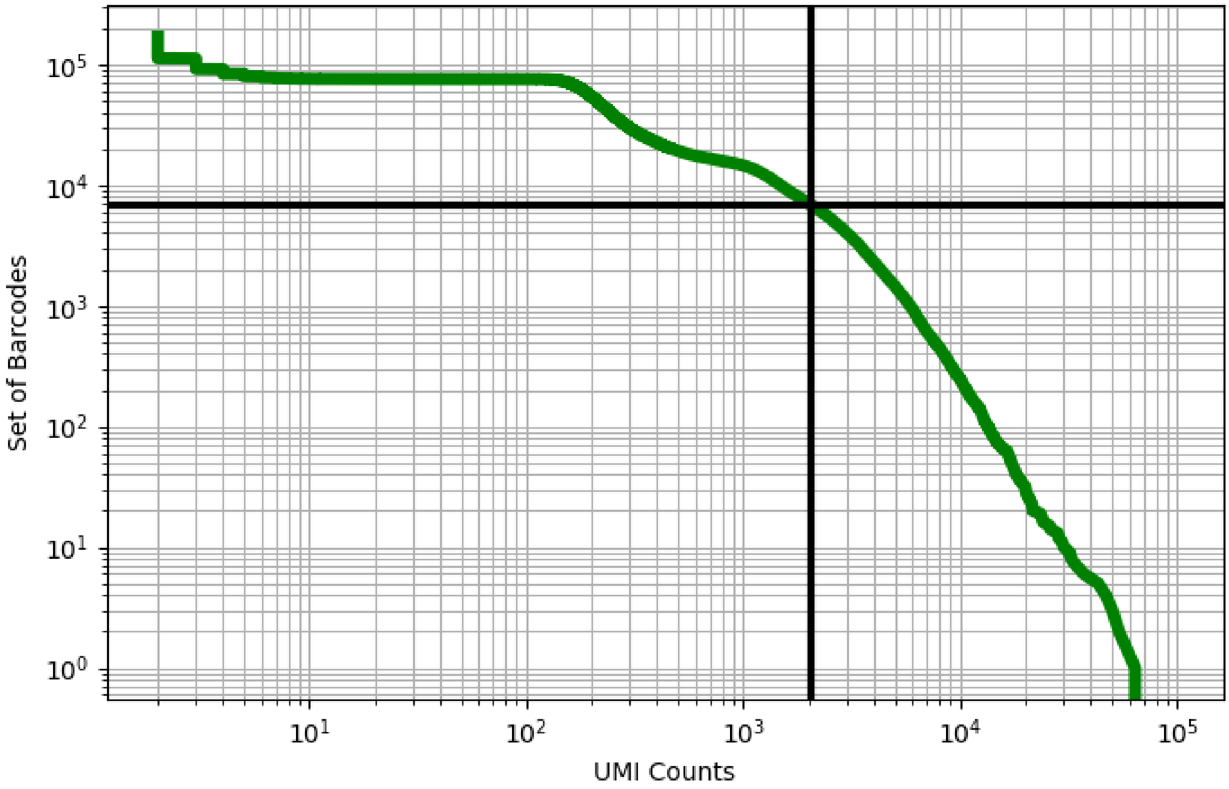


Figure 1: adipose/GSM3711757

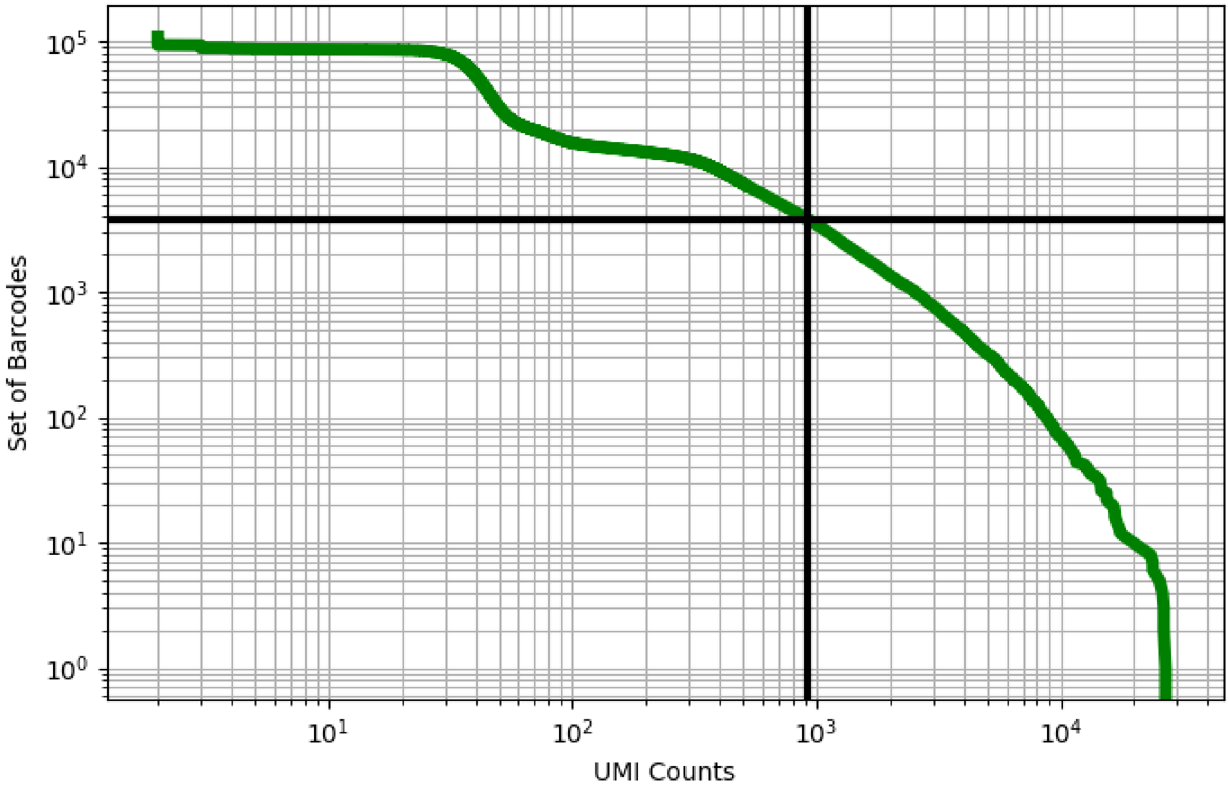


Figure 2: adipose/GSM3711758

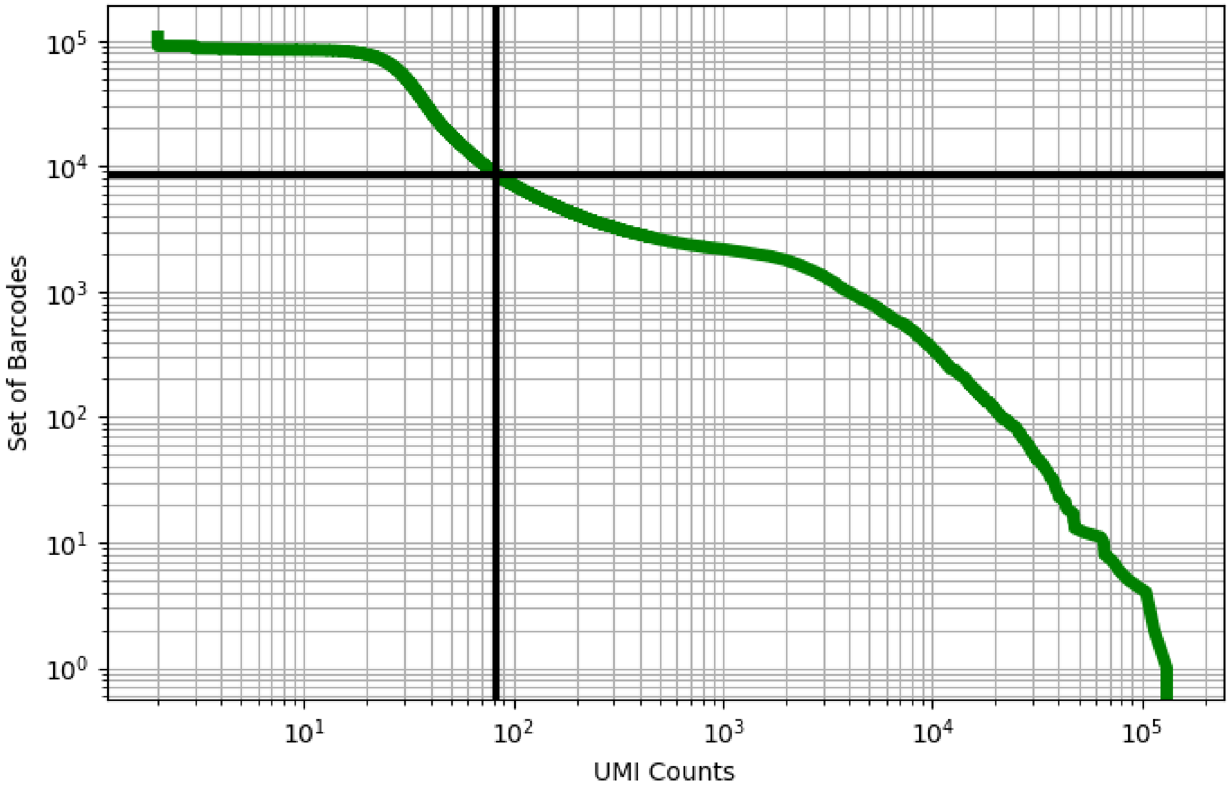


Figure 3: adipose/GSM3711759

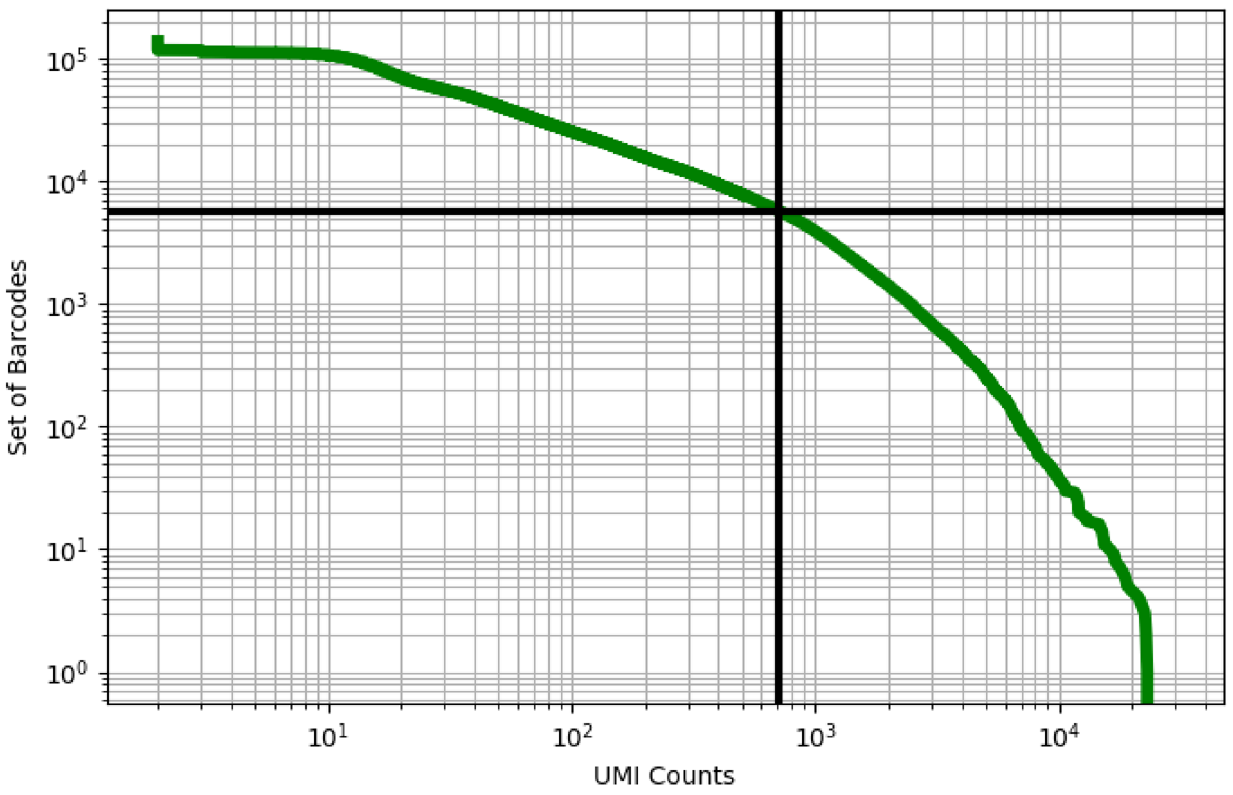


Figure 4: adipose/GSM3711760

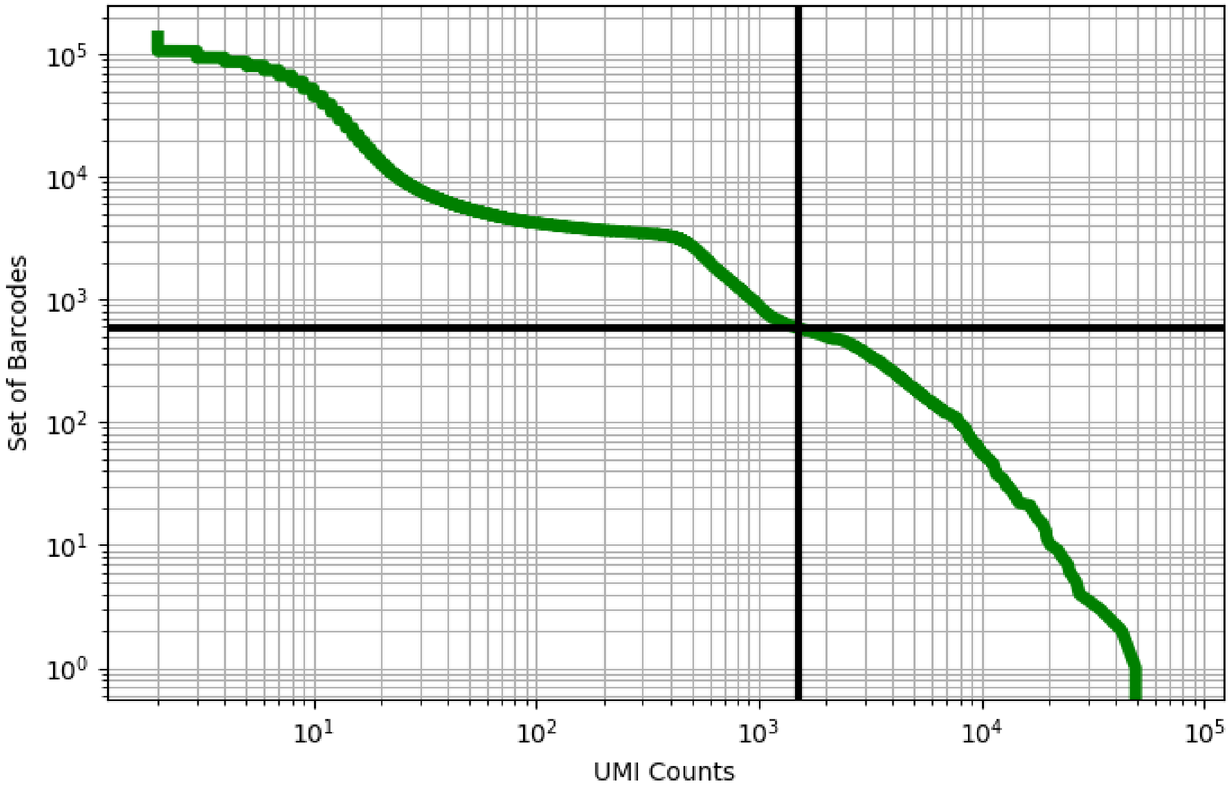


Figure 5: adipose/GSM3711761

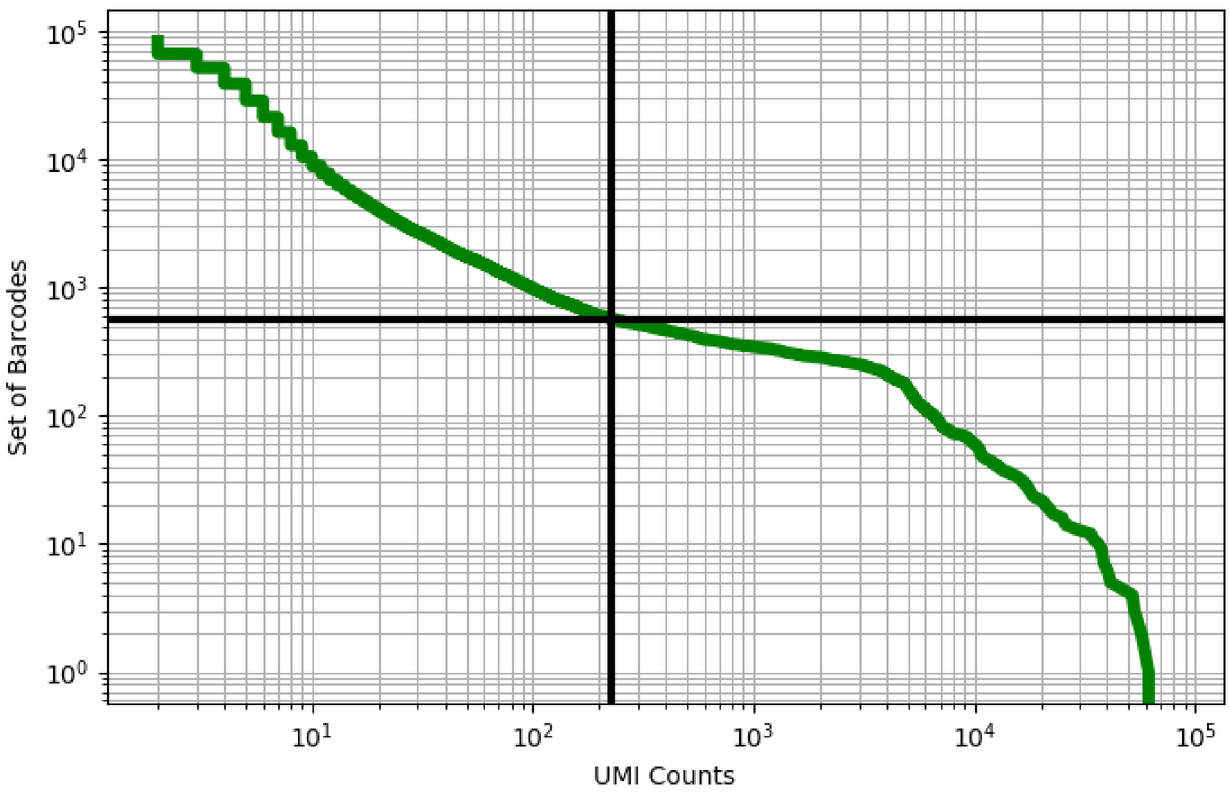


Figure 6: adipose/GSM3711762

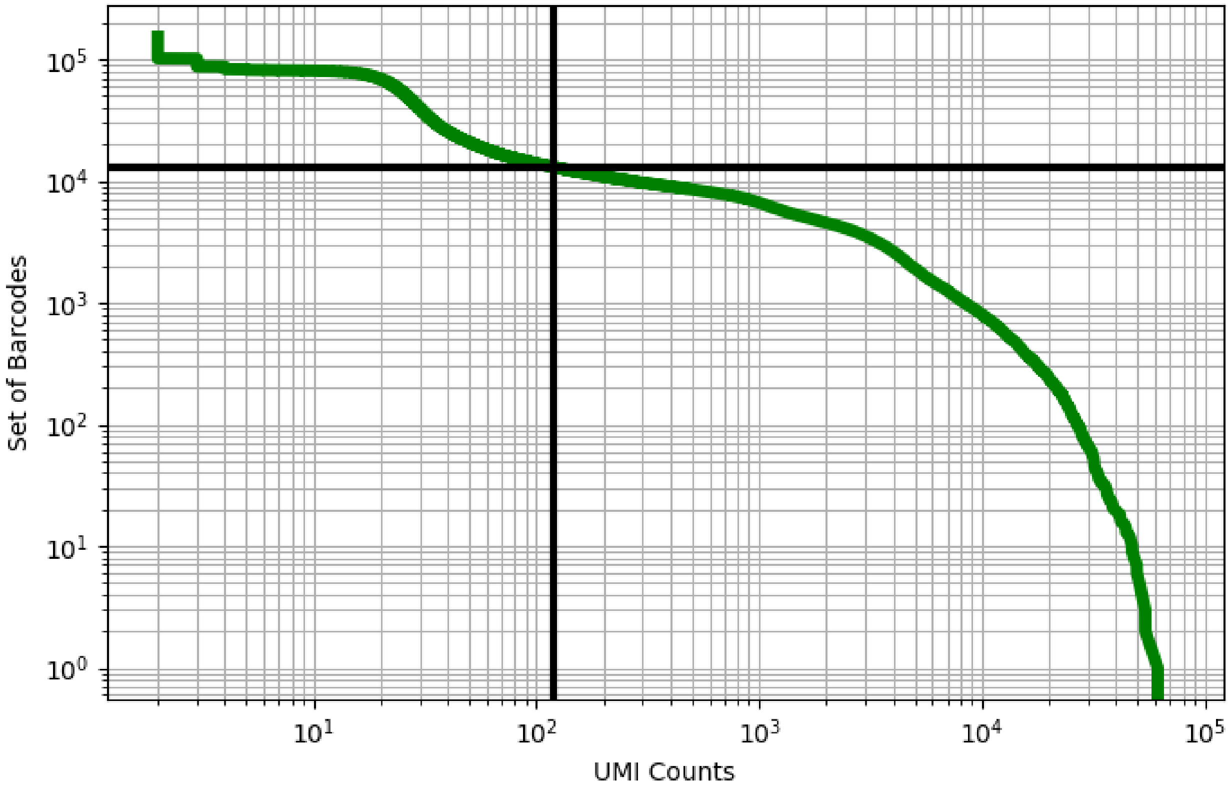


Figure 7: adipose/GSM3711763

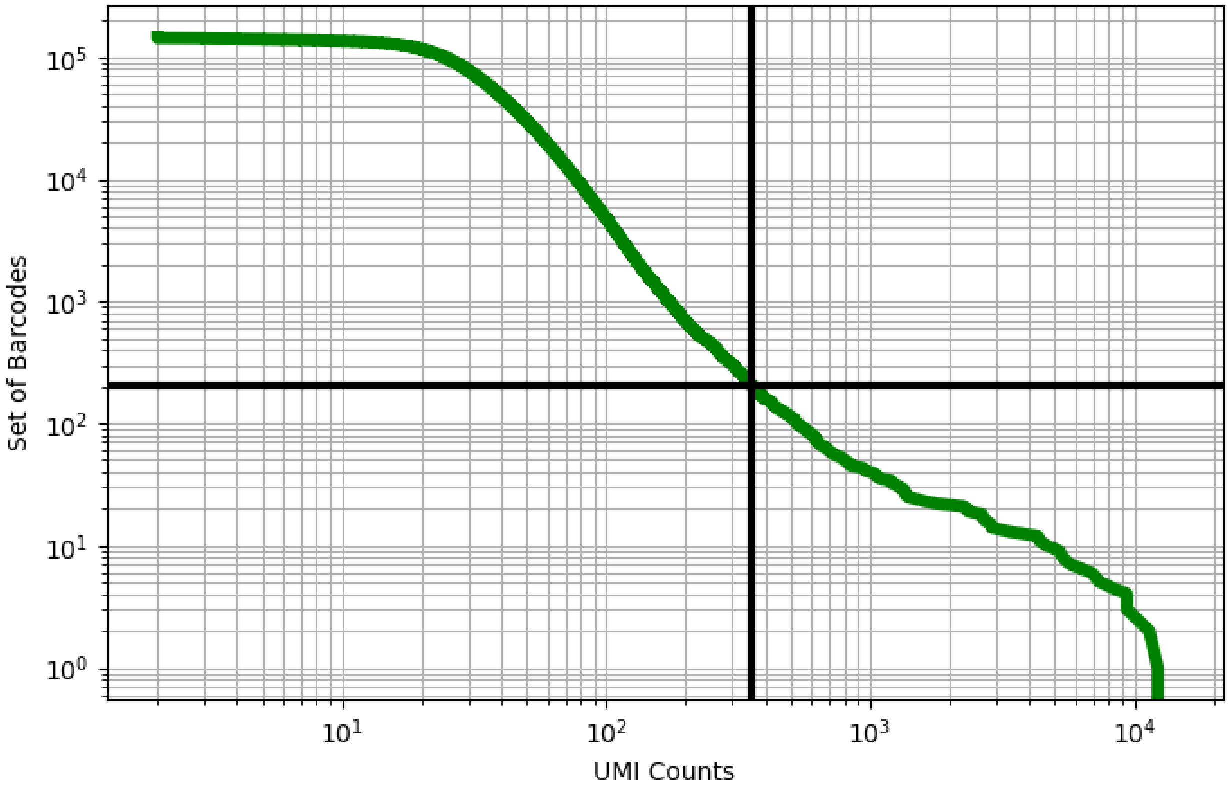


Figure 8: adipose/GSM3711764

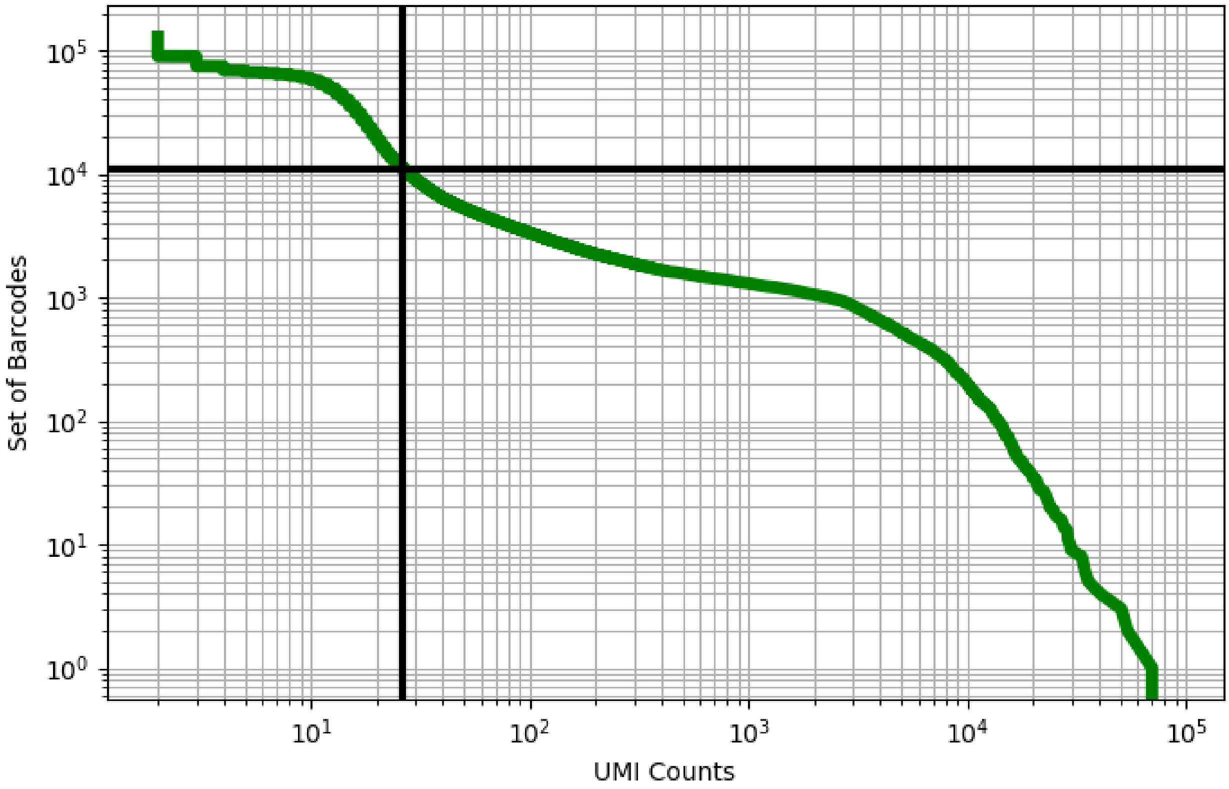


Figure 9: adipose/GSM3711765

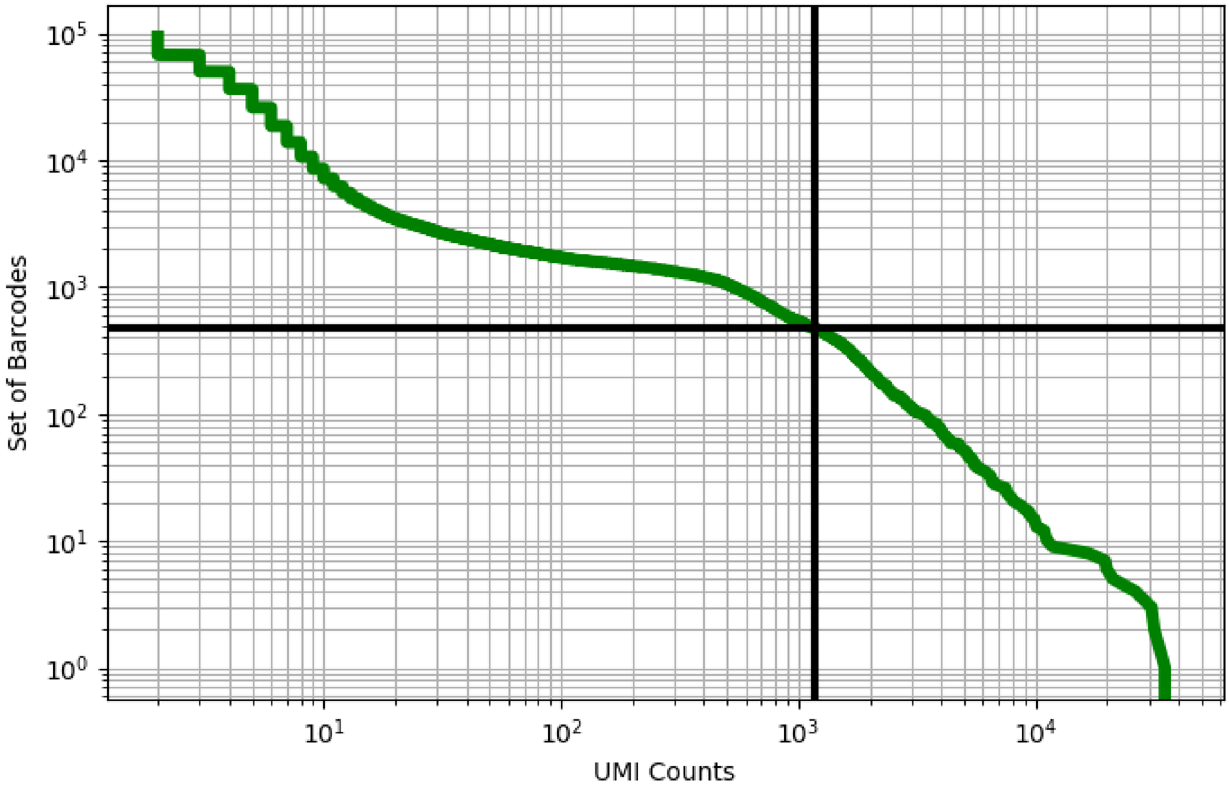


Figure 10: adipose/GSM3711766

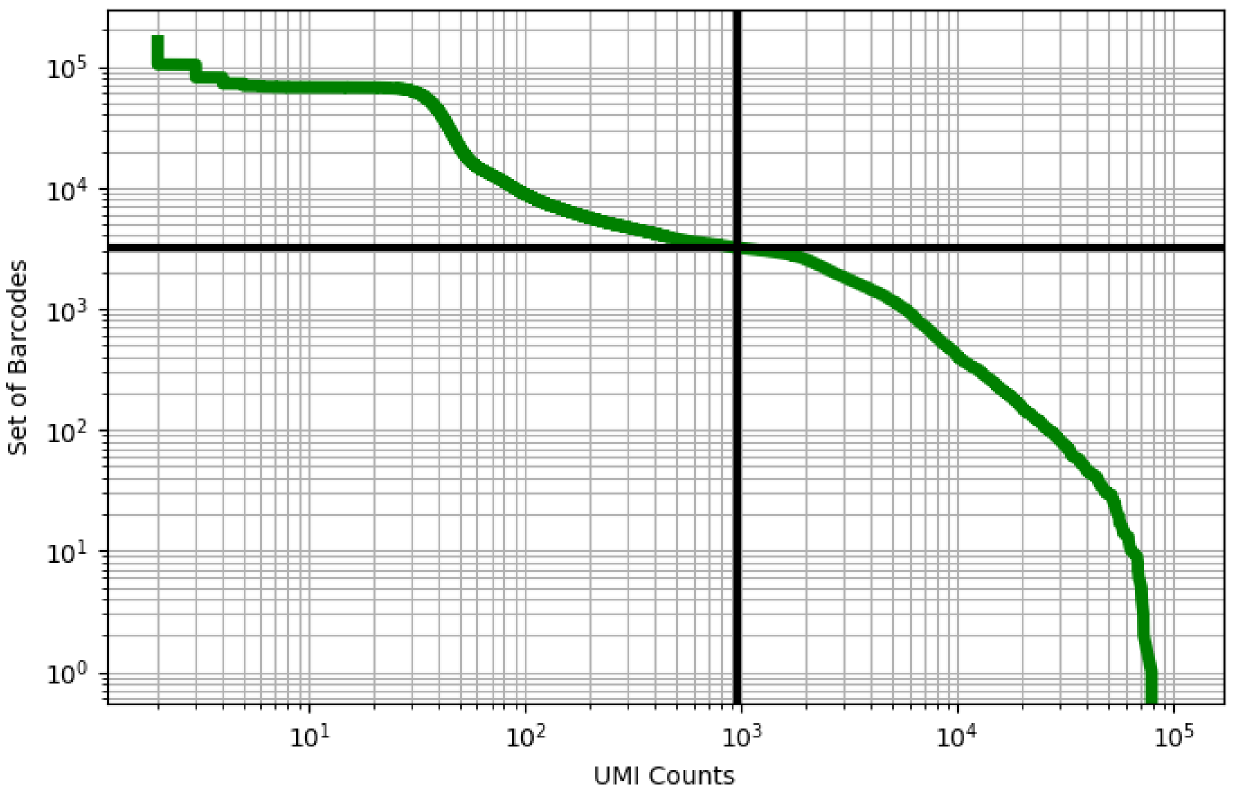


Figure 11: adipose/GSM3711767

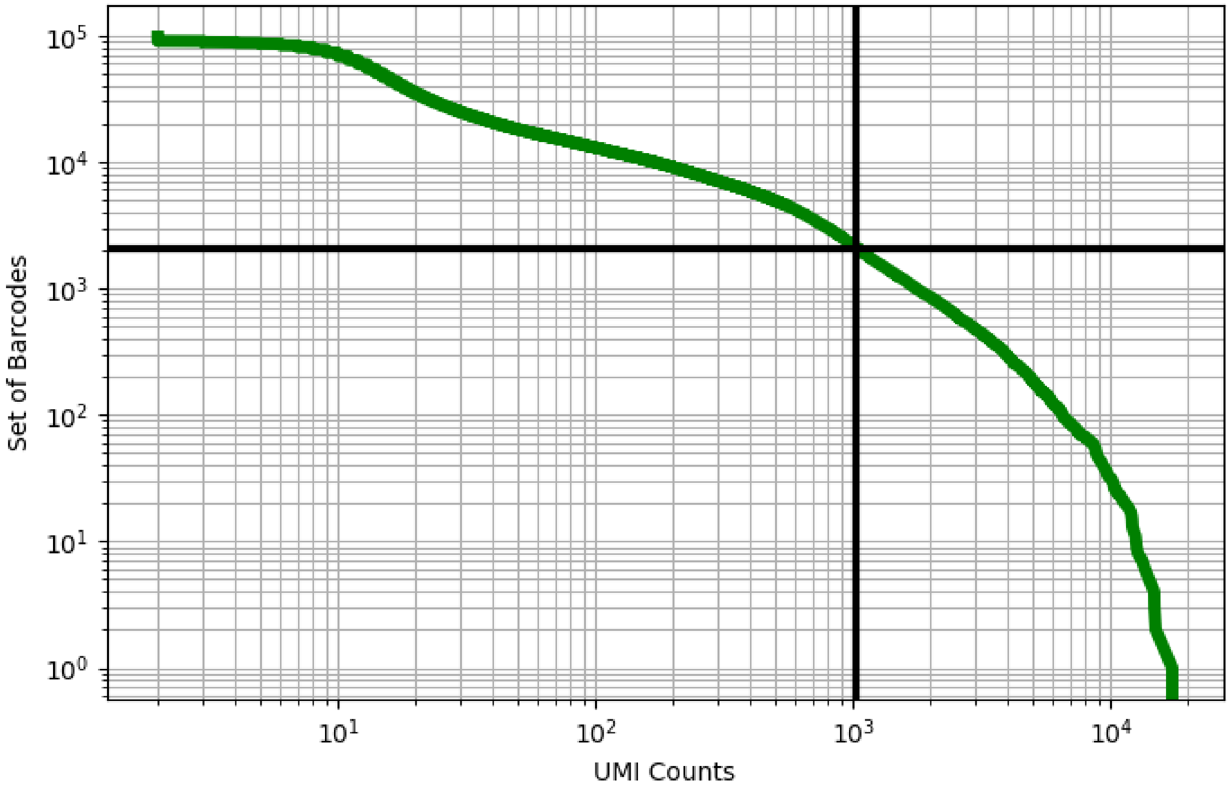


Figure 12: adipose/GSM3711768

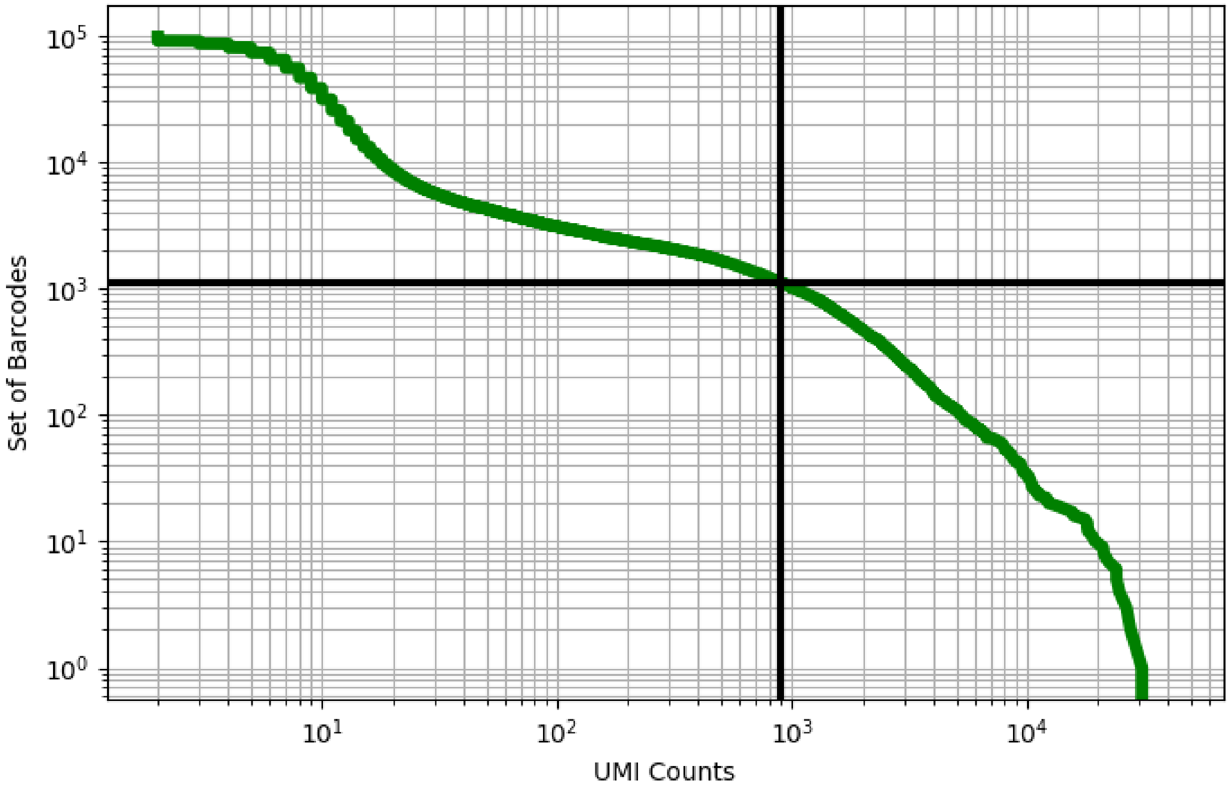


Figure 13: adipose/GSM3711769

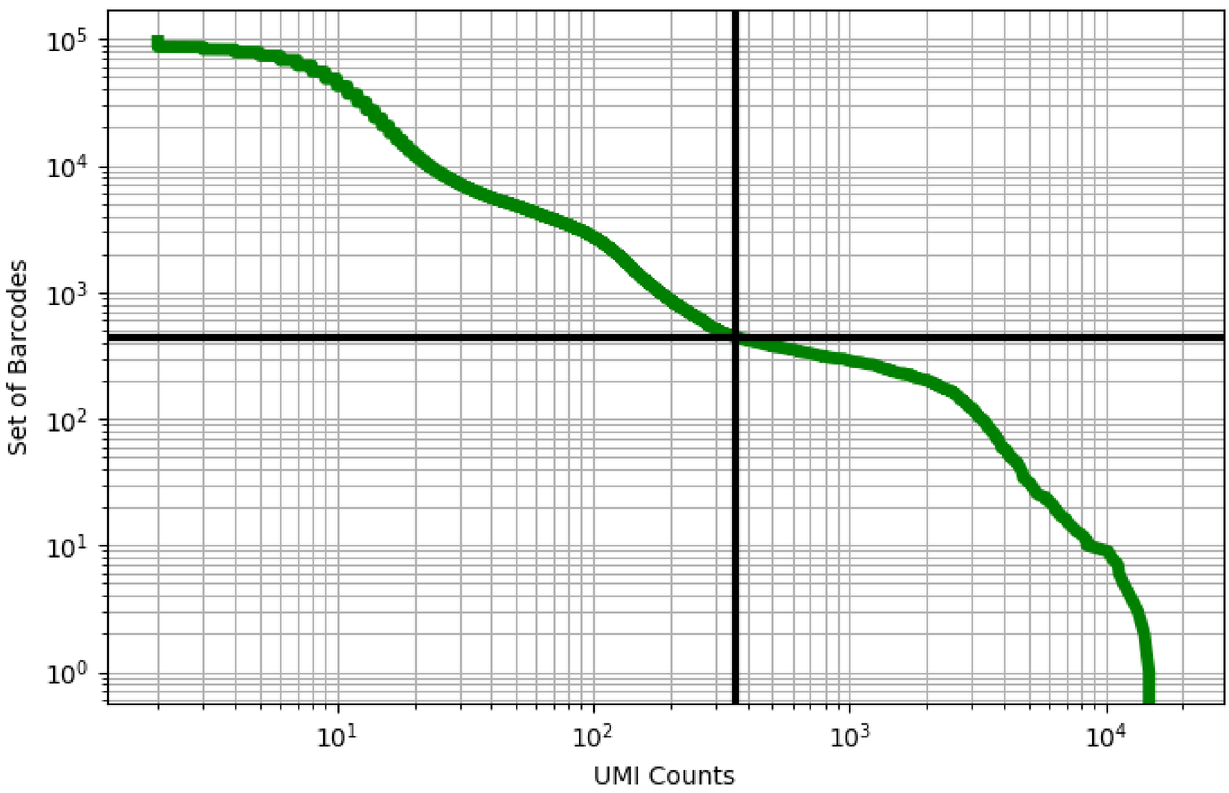


Figure 14: adipose/GSM3711770

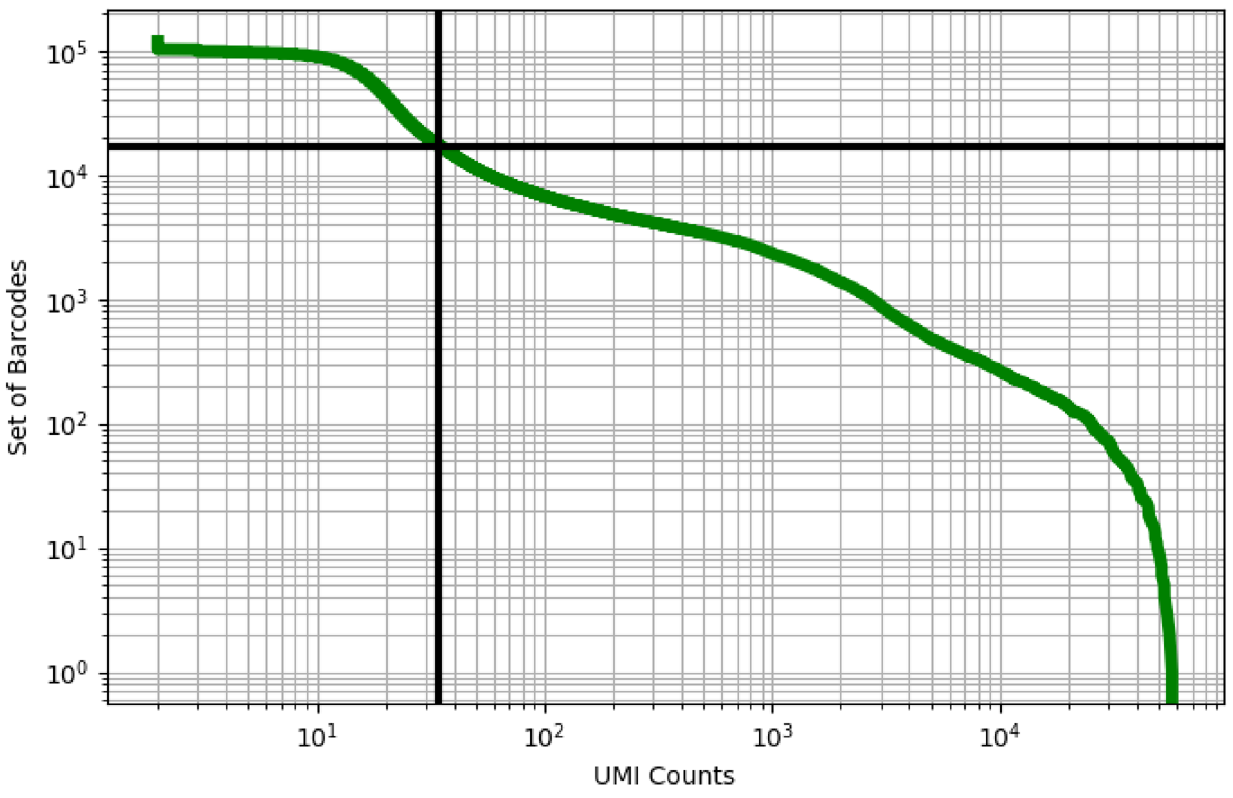


Figure 15: adipose/GSM3711771

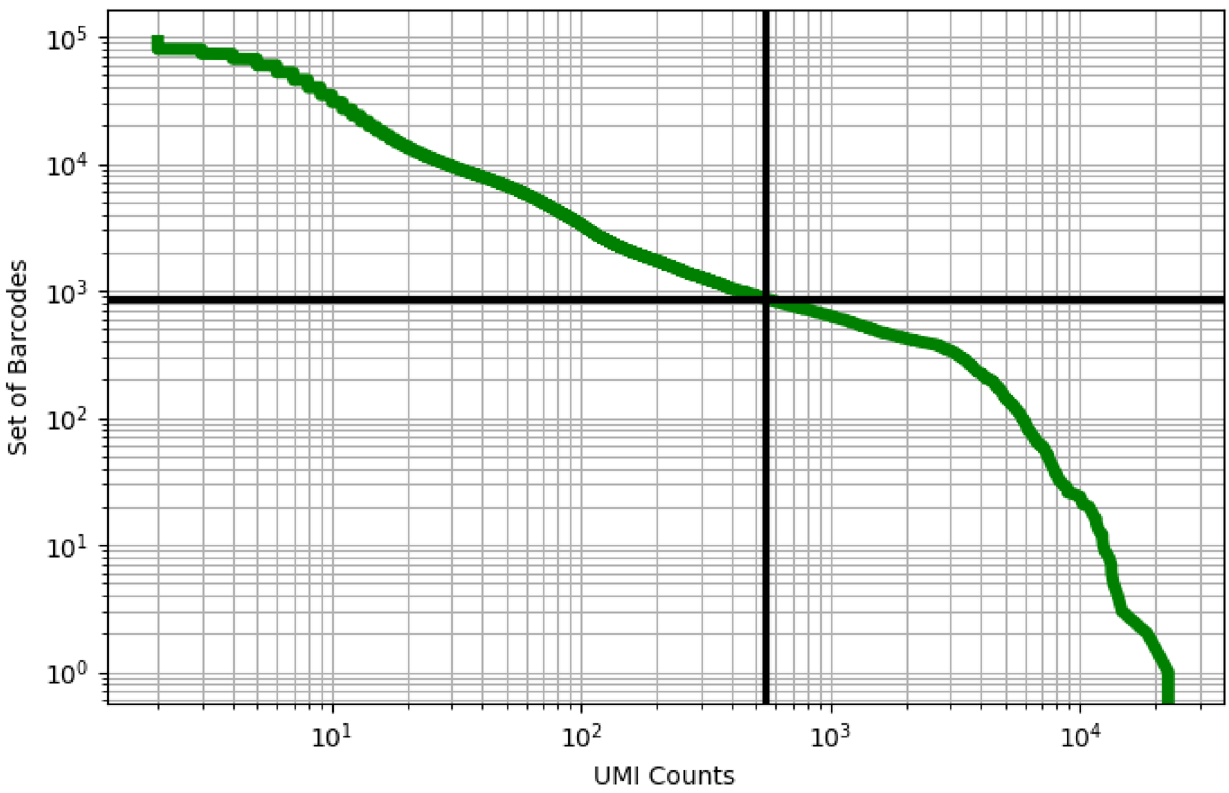


Figure 16: adipose/GSM3711772

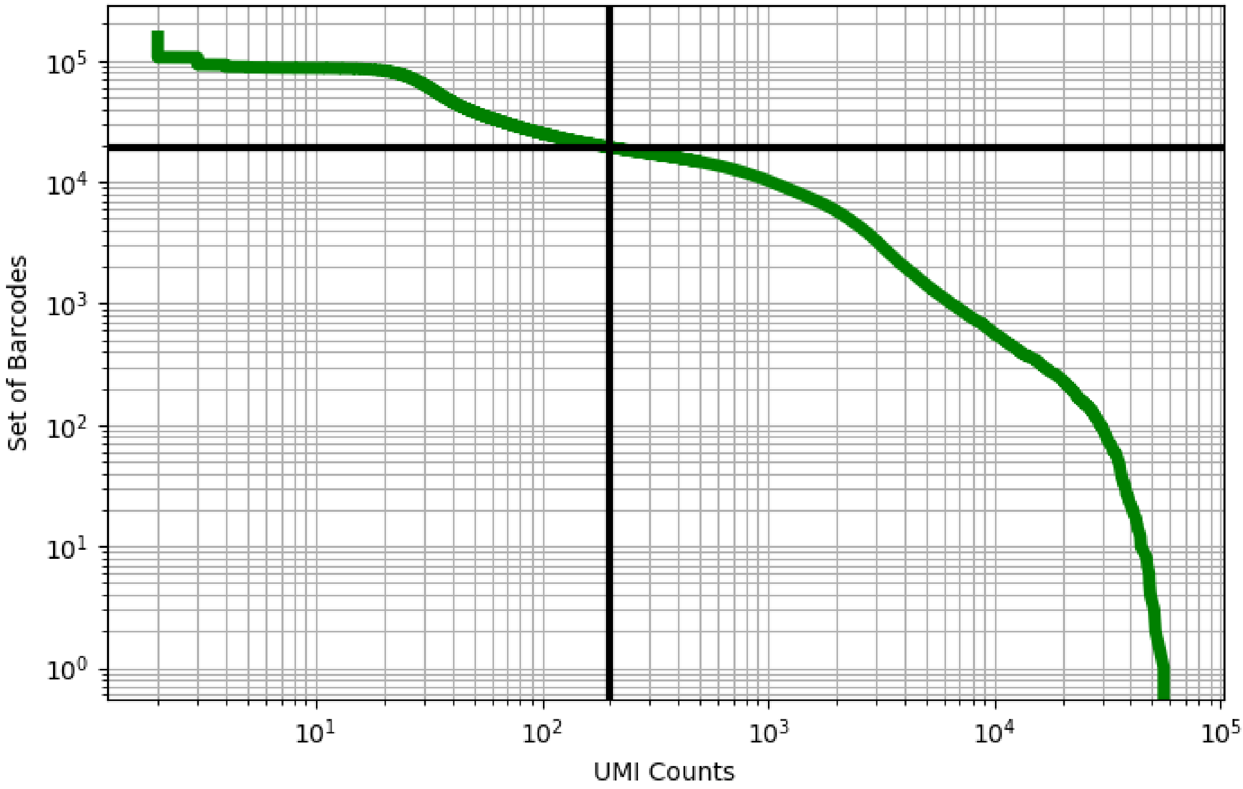


Figure 17: adipose/GSM3711773

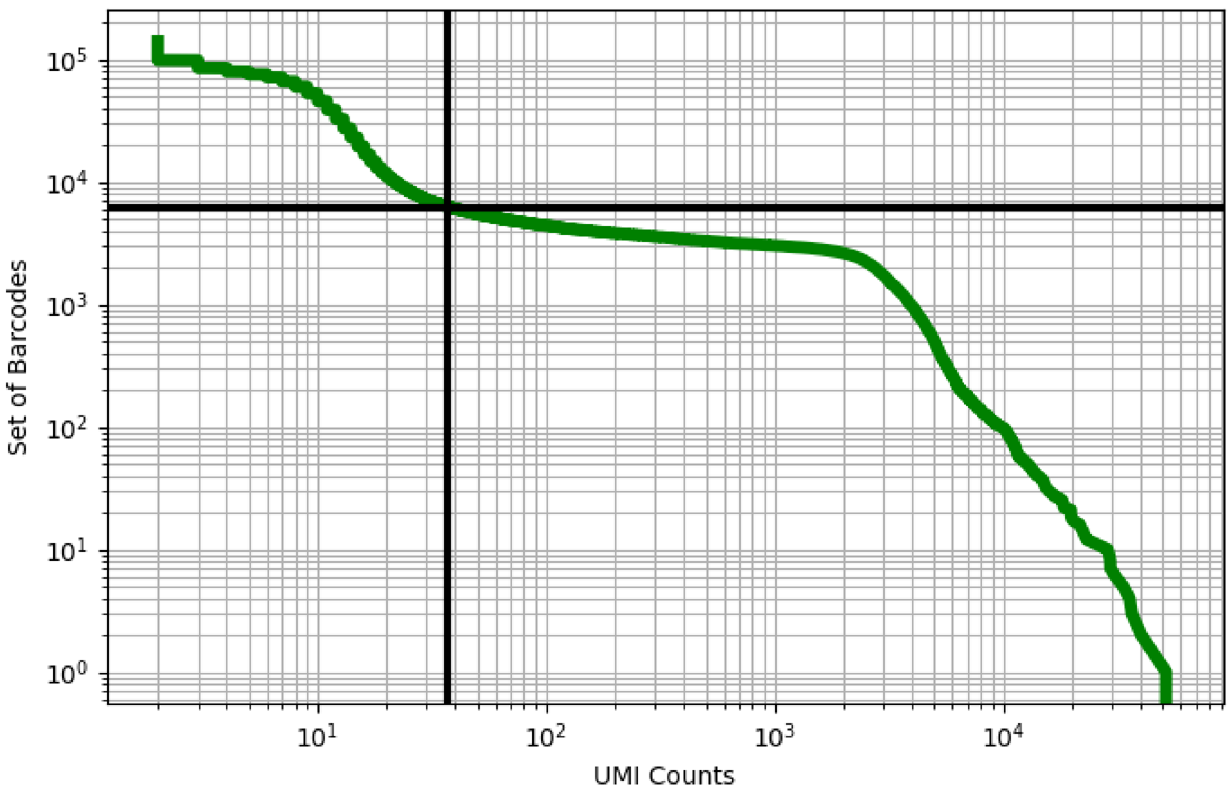


Figure 18: adipose/GSM3711774

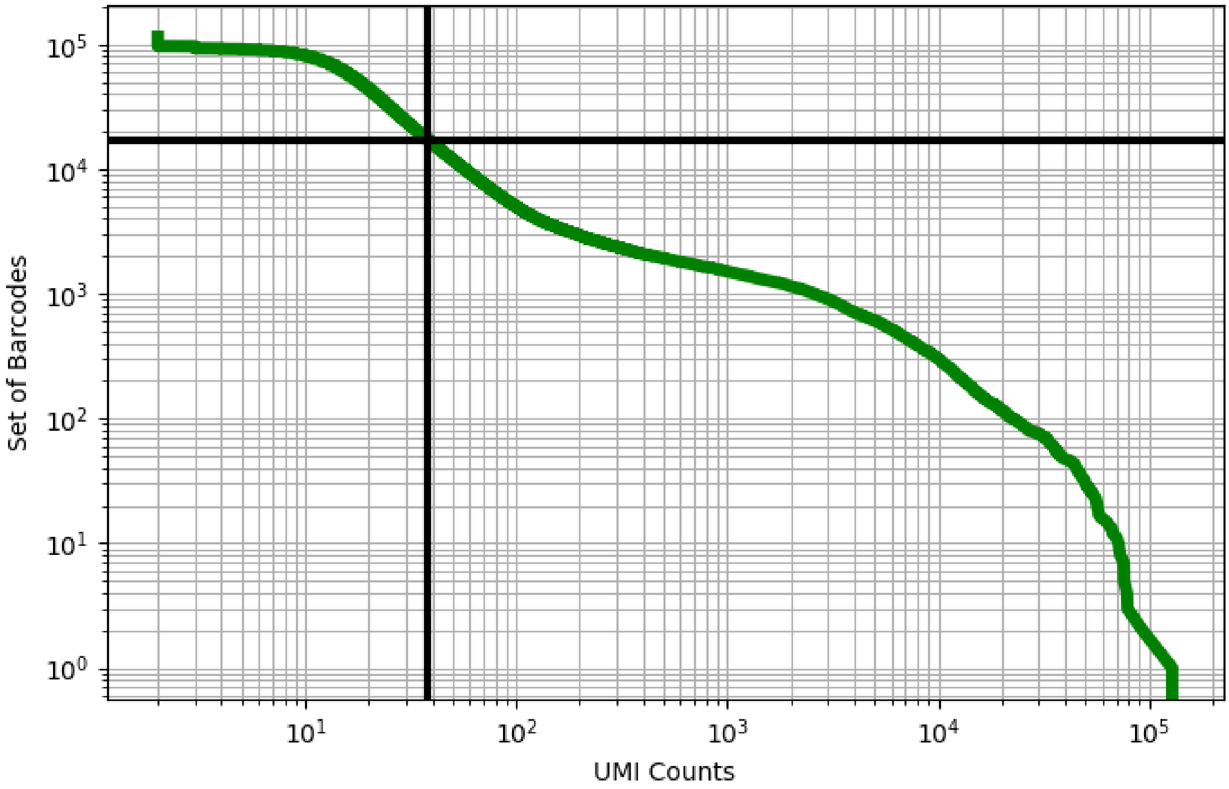


Figure 19: adipose/GSM3711775

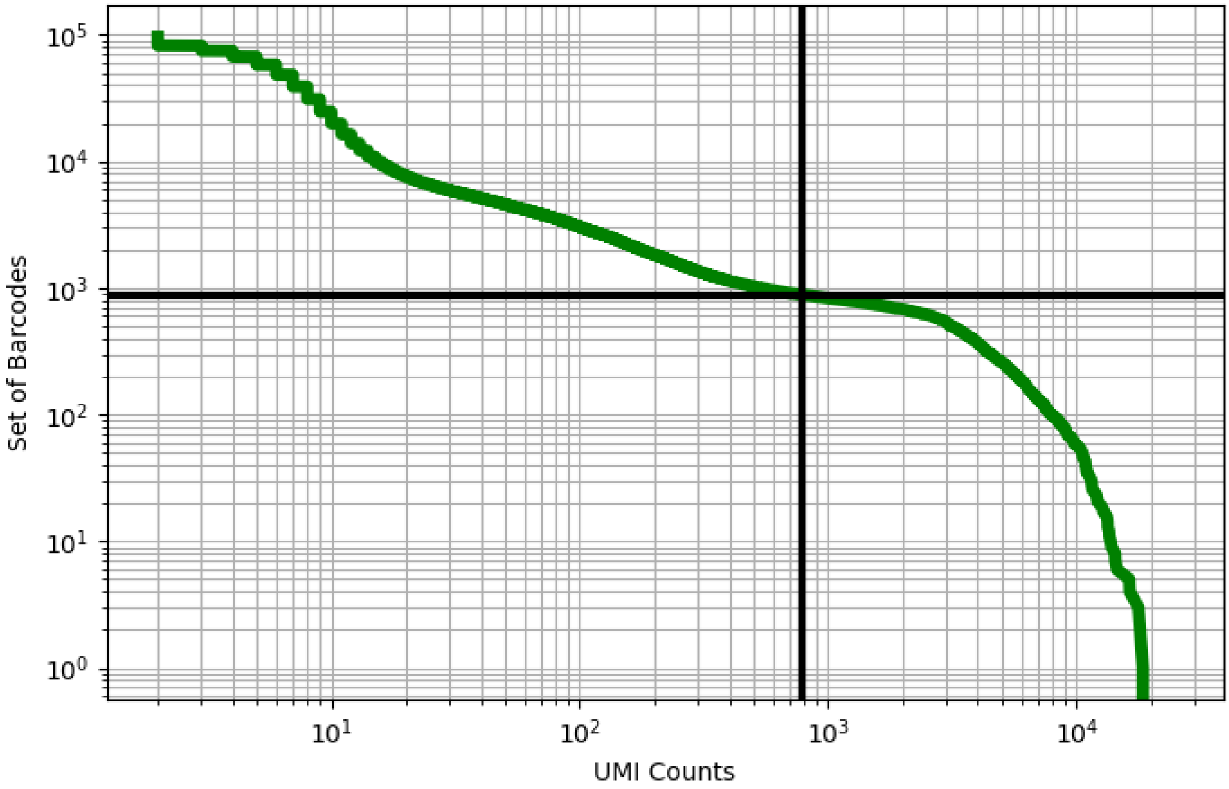


Figure 20: adipose/GSM3711776

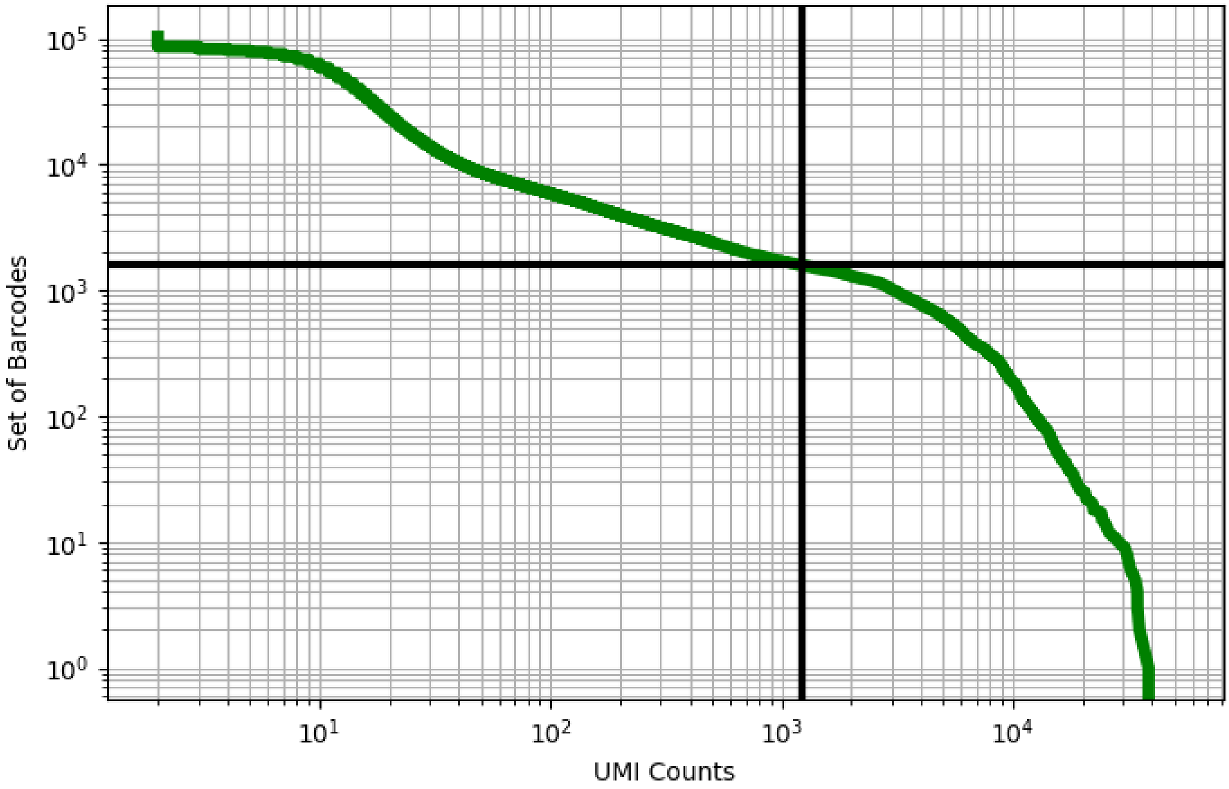


Figure 21: adipose/GSM3711777

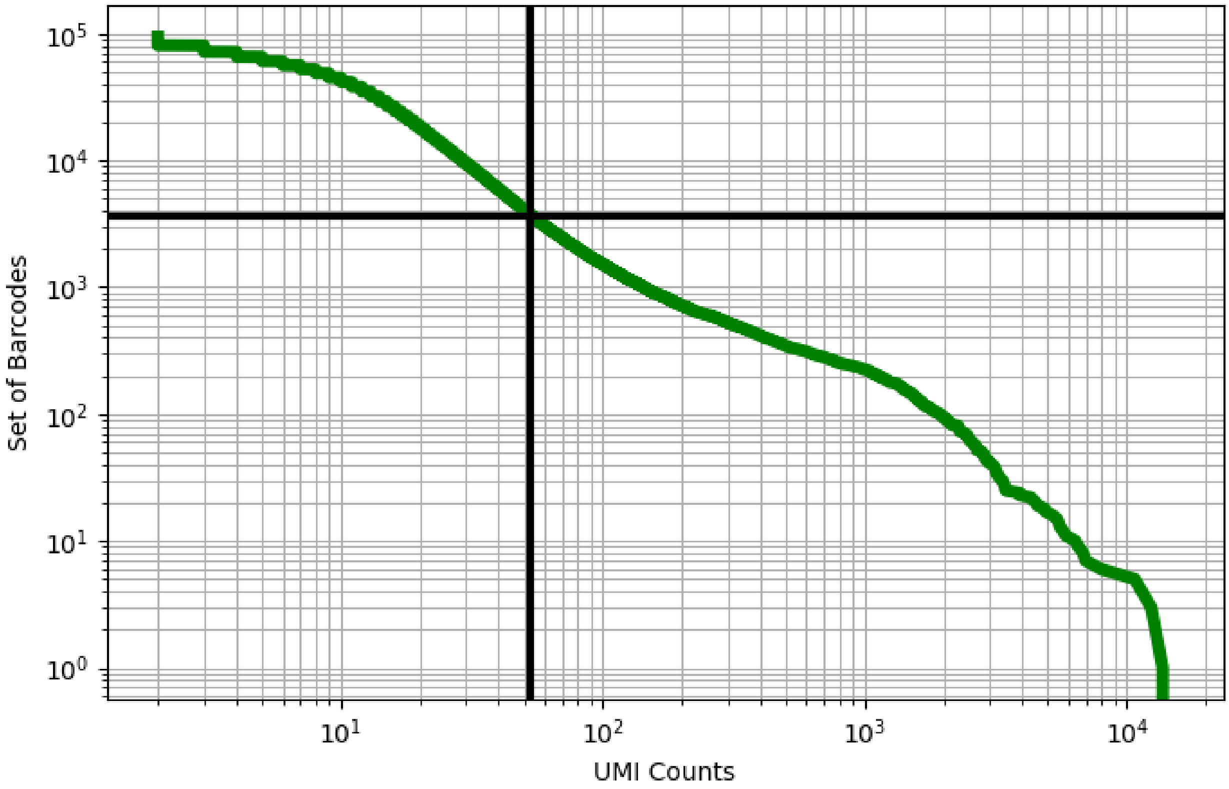


Figure 22: adipose/GSM3711778

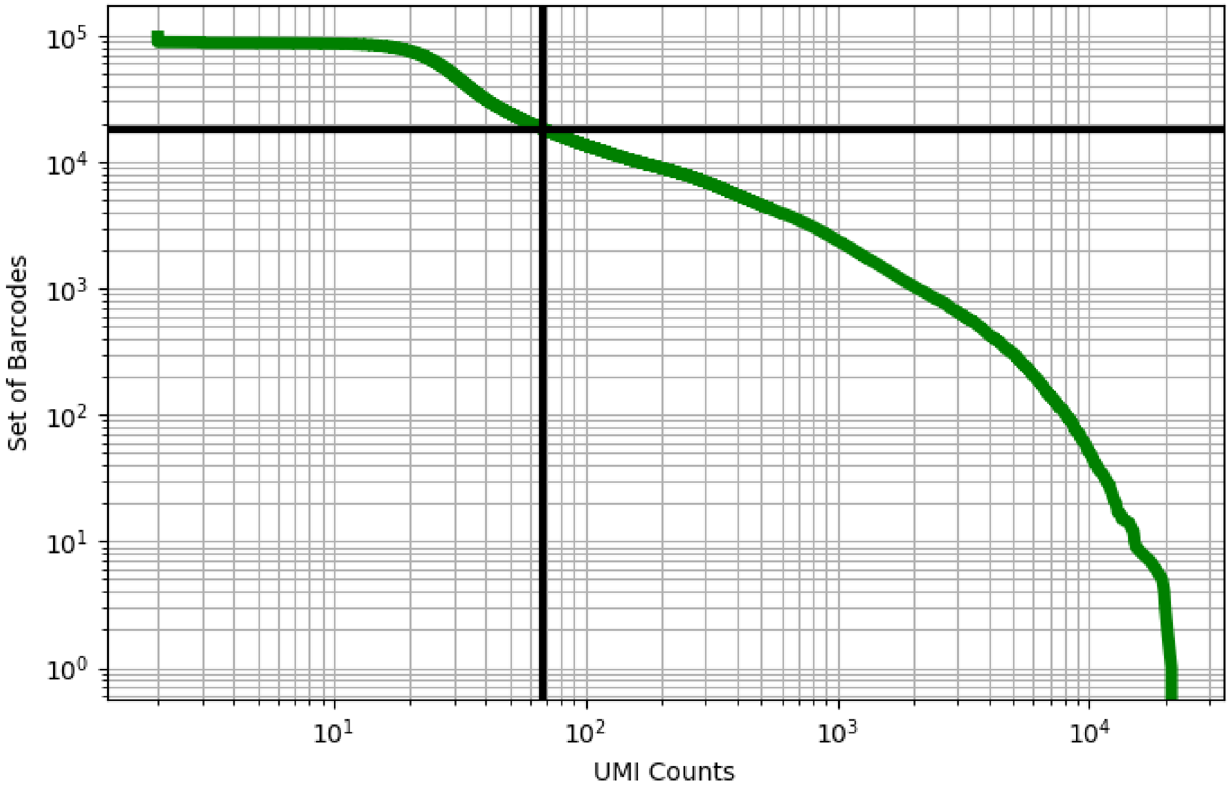


Figure 23: adipose/GSM3711779

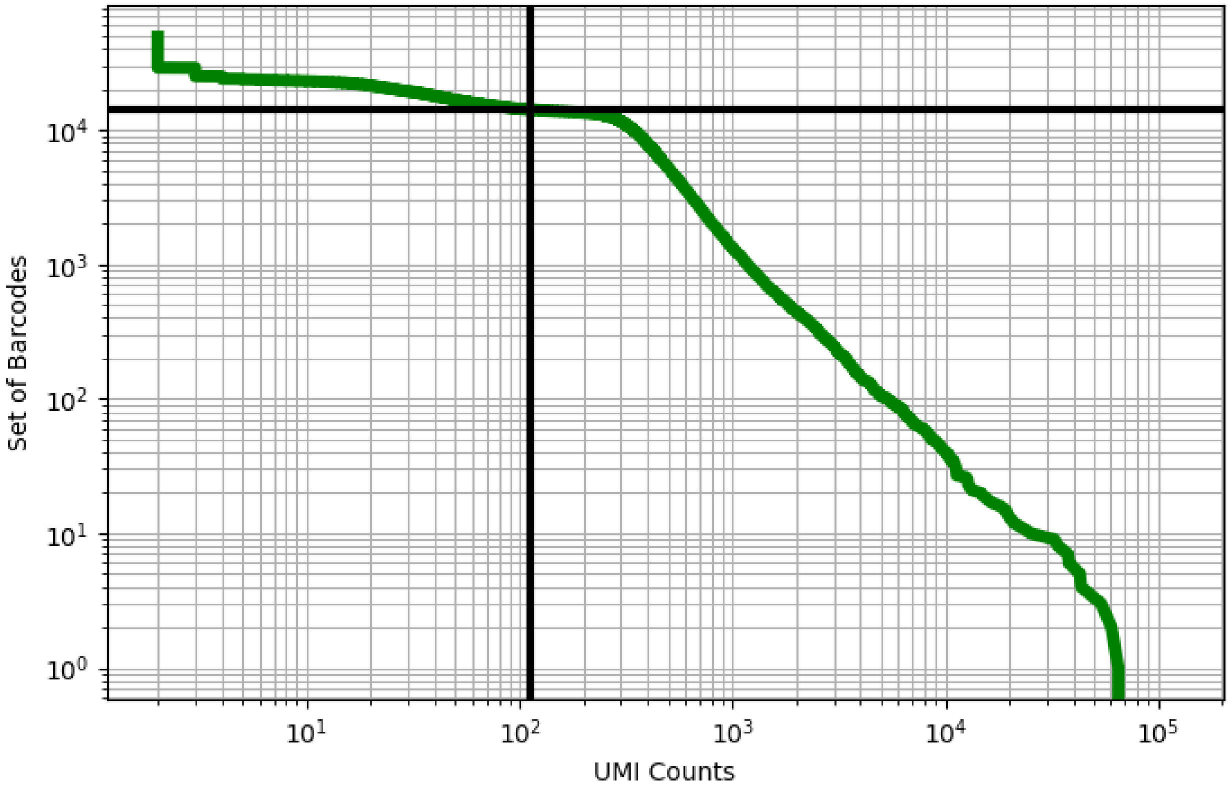


Figure 24: adipose/GSM3711780

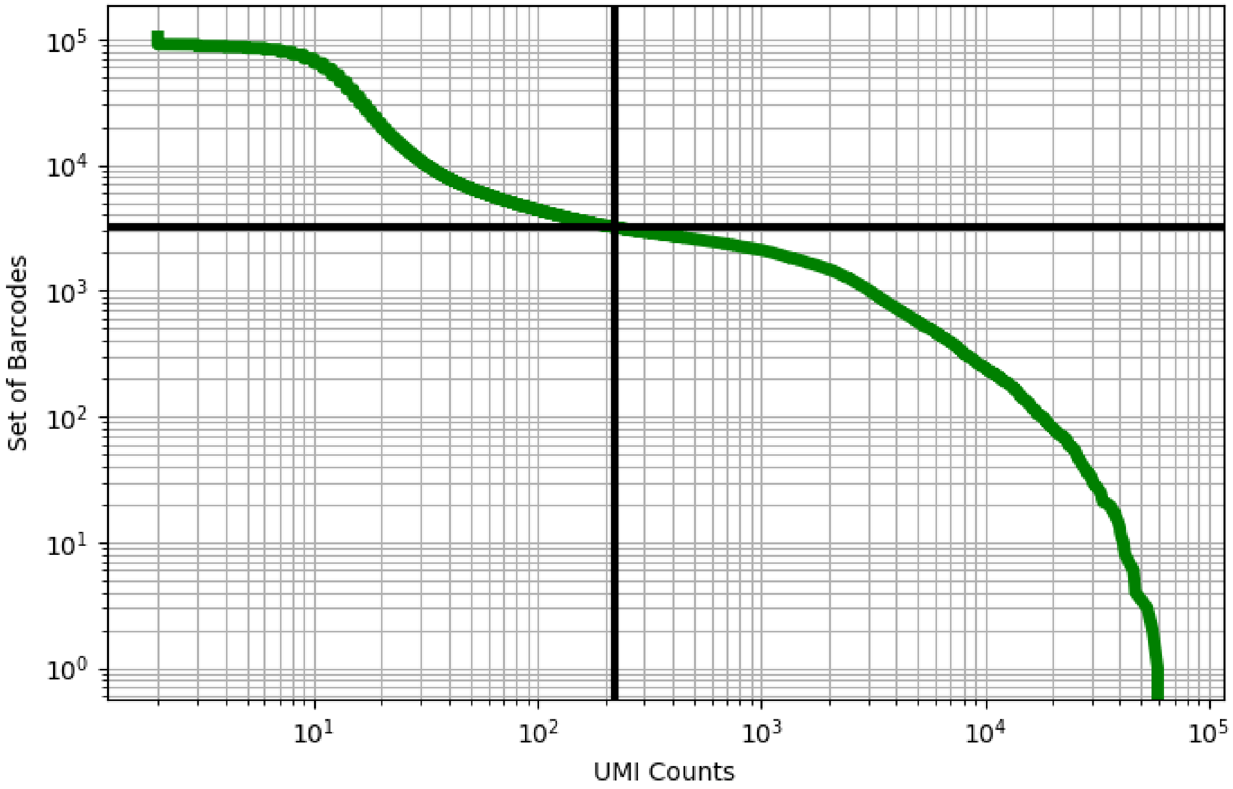


Figure 25: adipose/GSM3711781

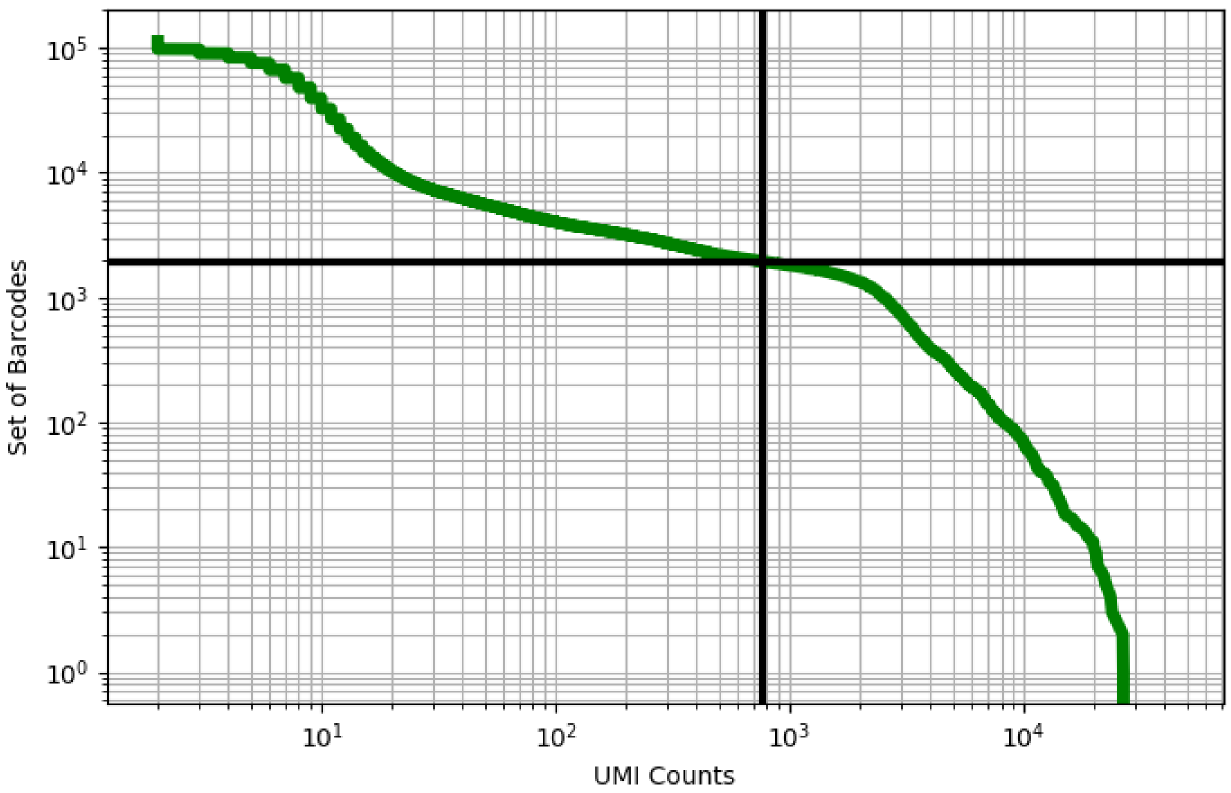


Figure 26: adipose/GSM3711782

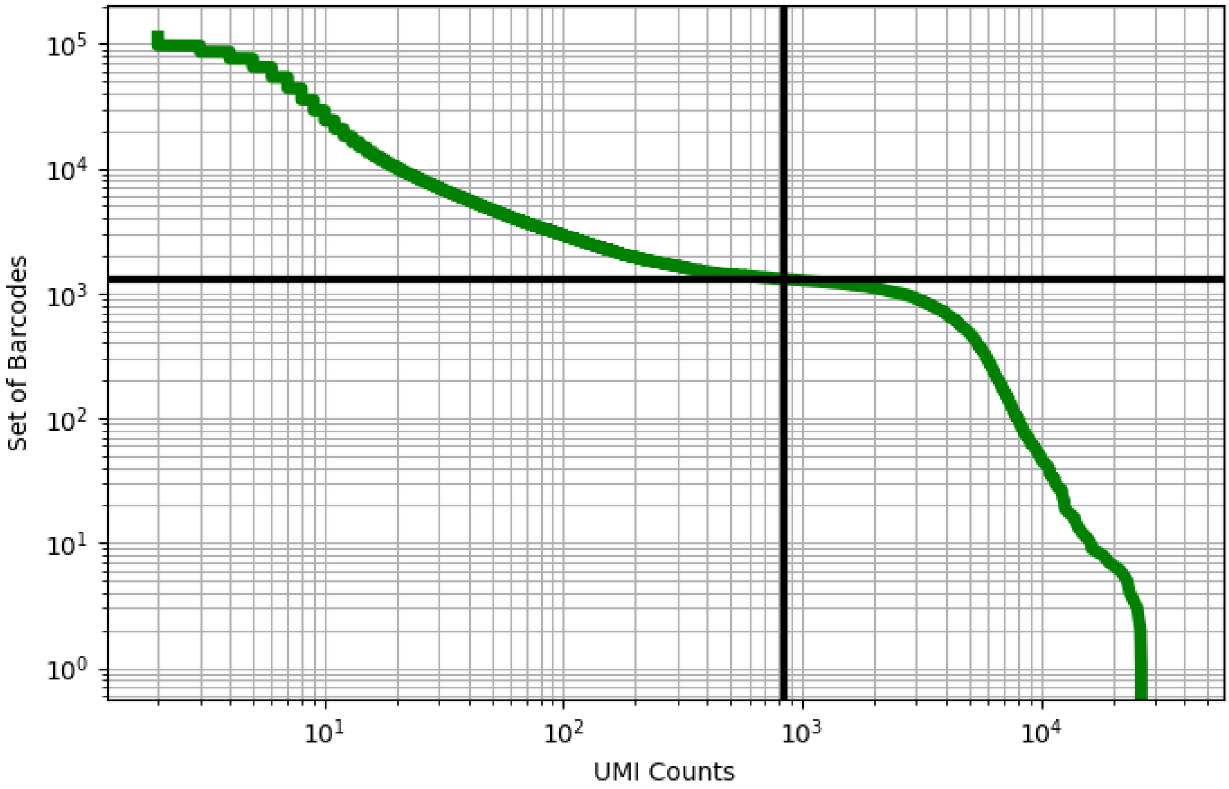


Figure 27: adipose/GSM3711783

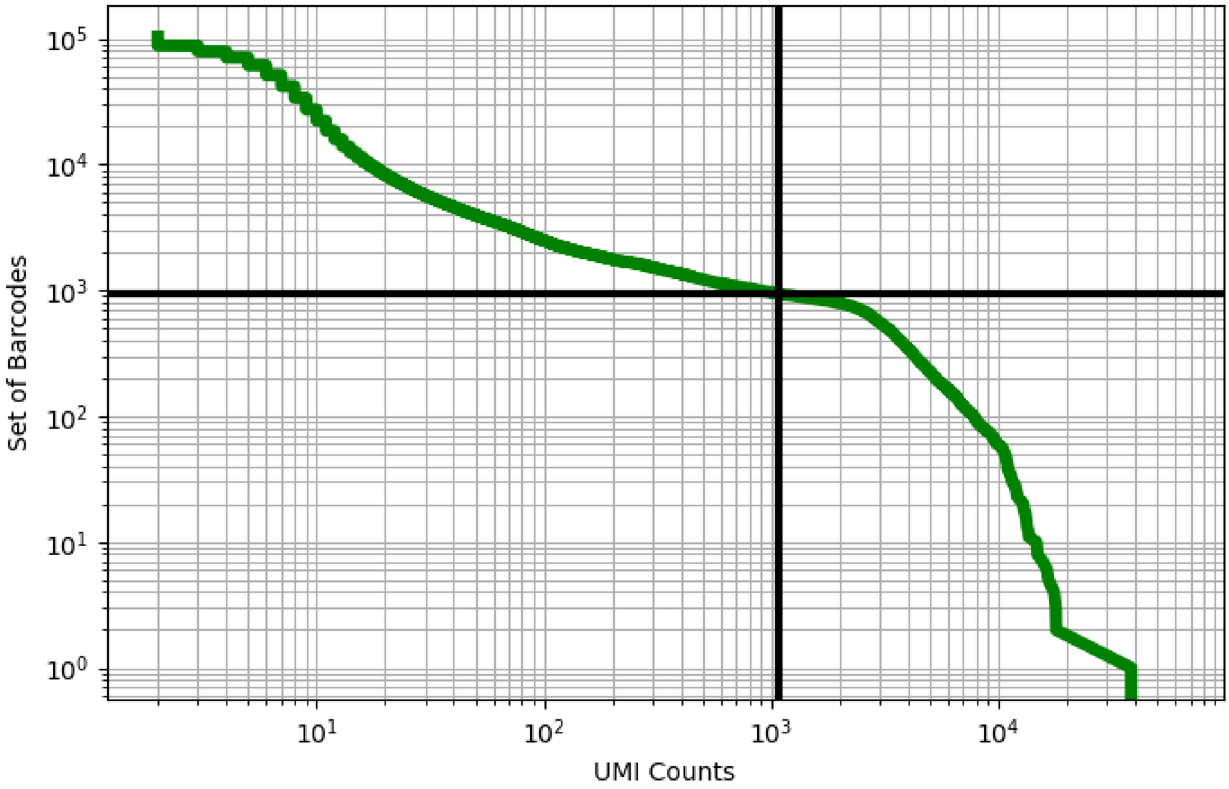


Figure 28: adipose/GSM3711784

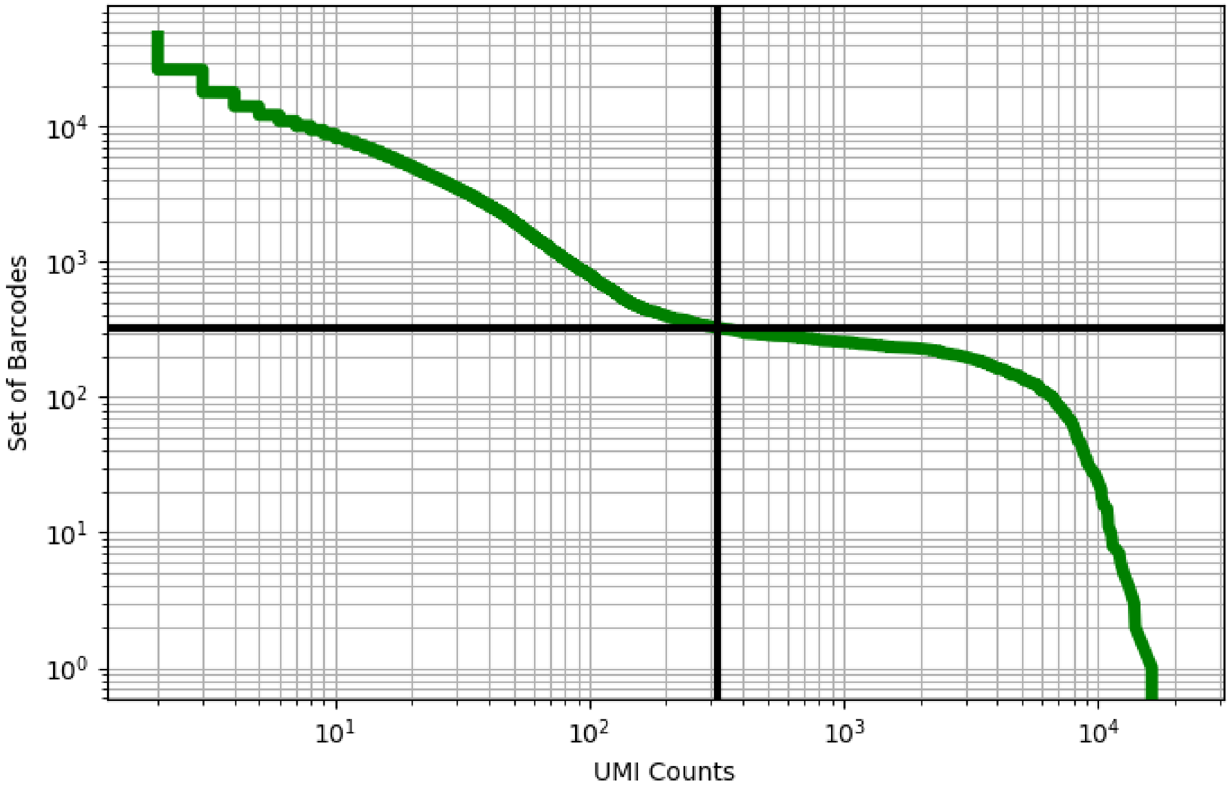


Figure 29: adipose/GSM3711785

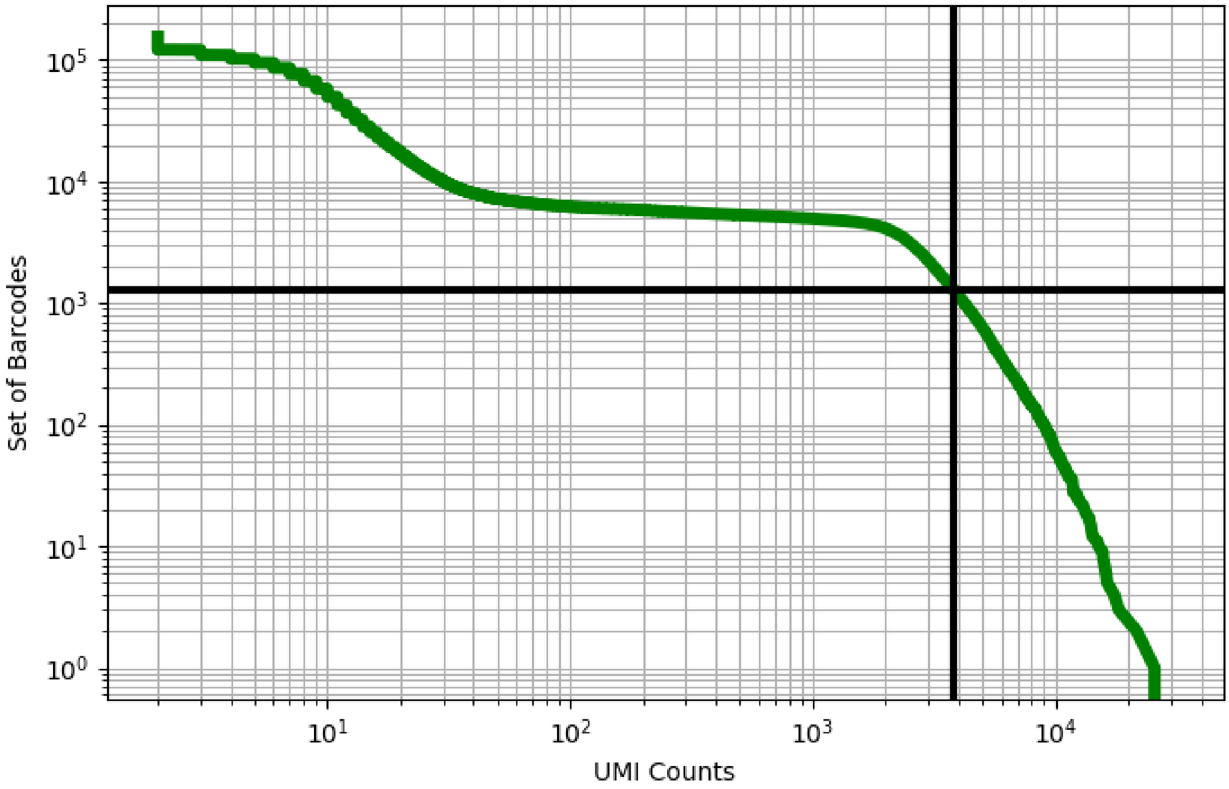


Figure 30: adipose/GSM3711786

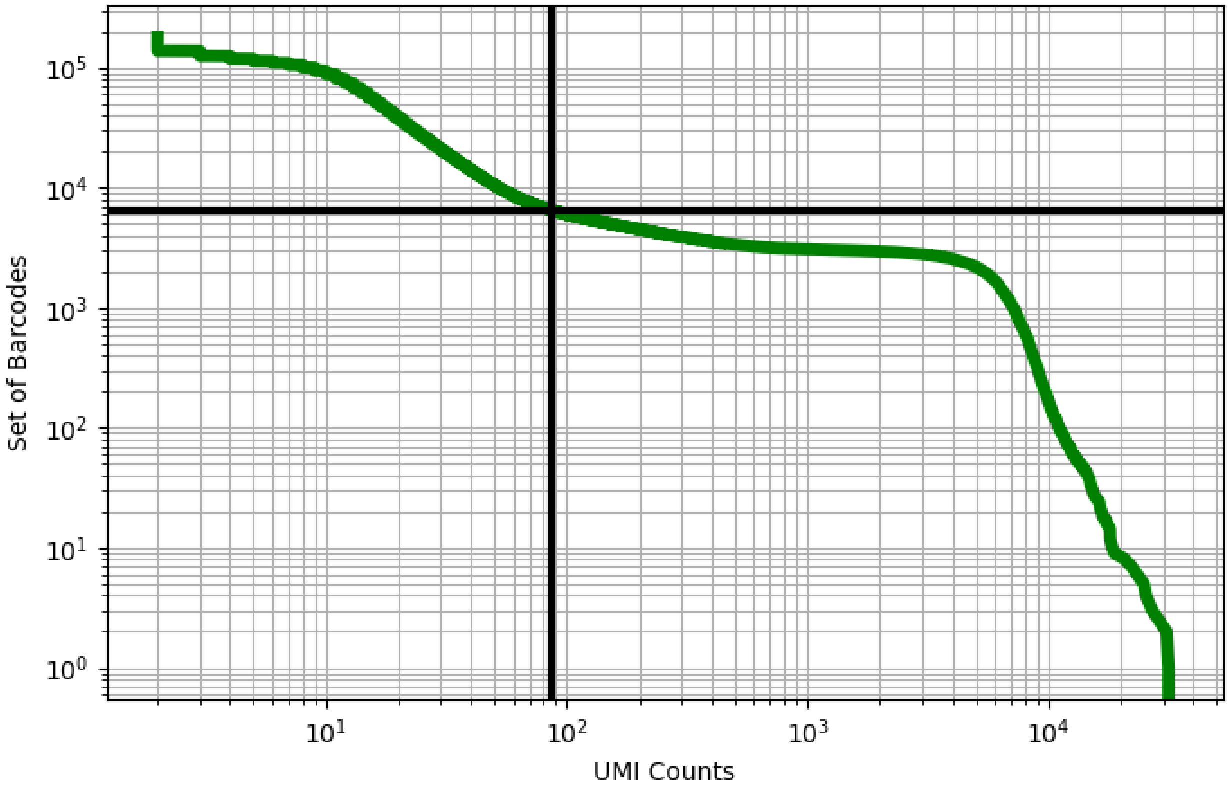


Figure 31: adipose/GSM3711787

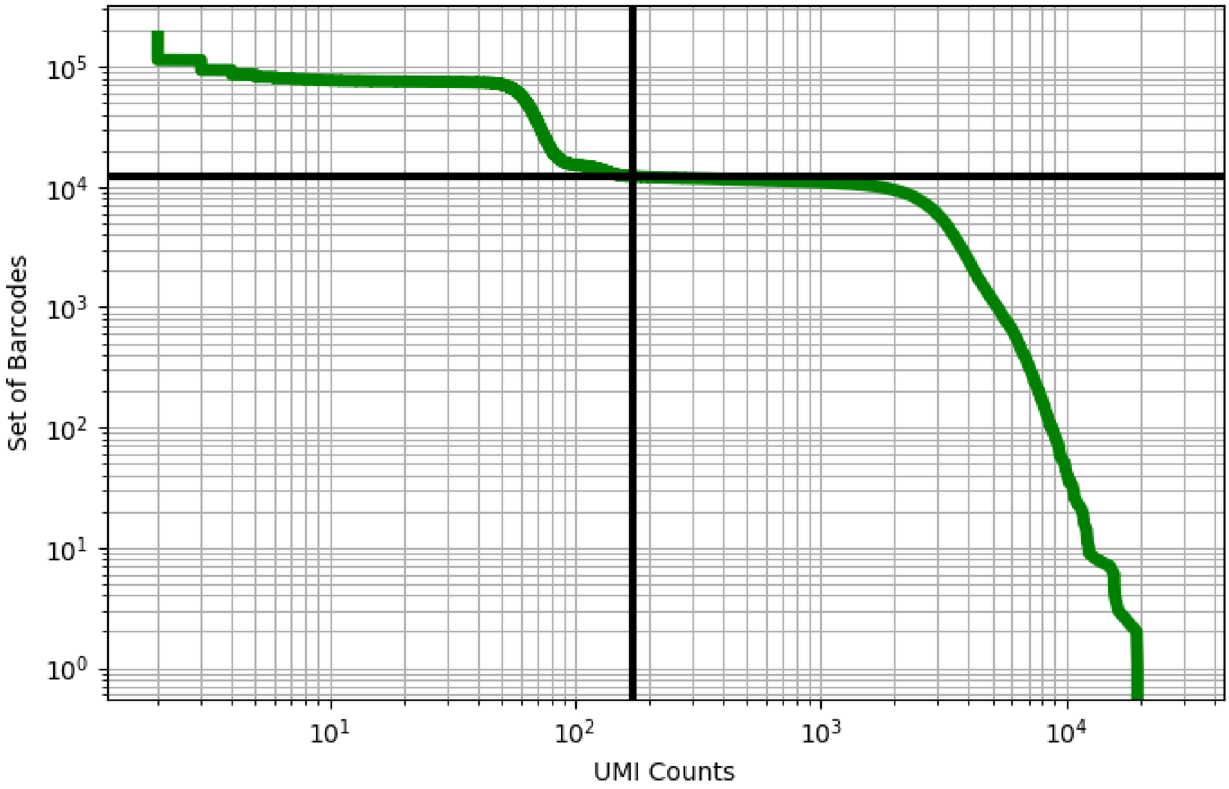


Figure 32: adipose/GSM3717979

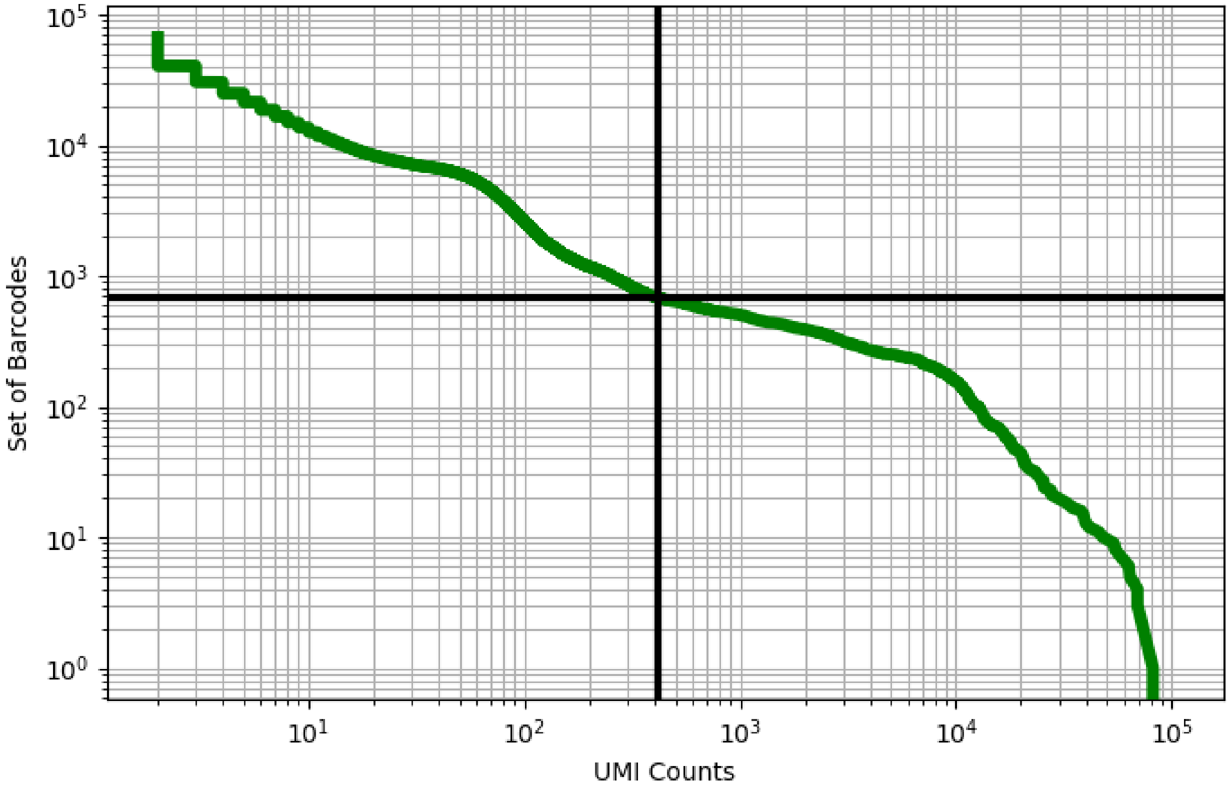


Figure 33: bladder/GSM3723357

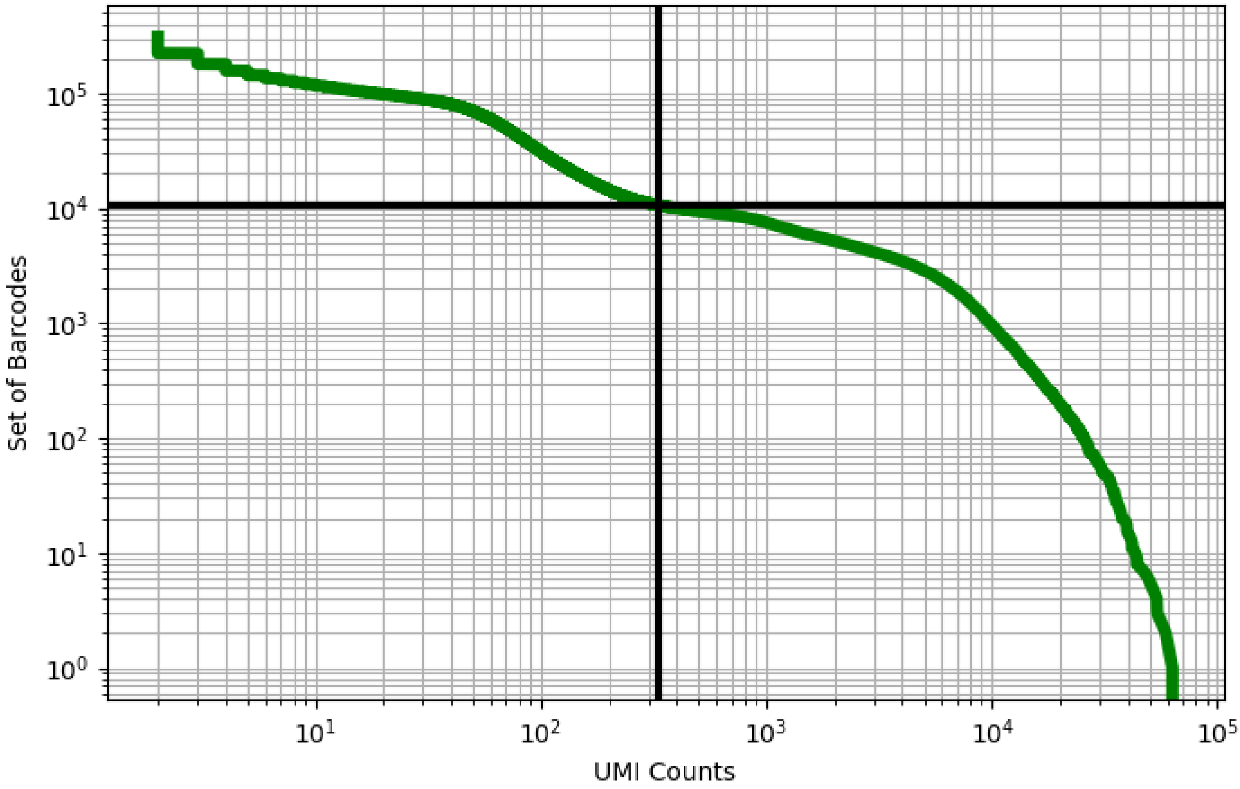


Figure 34: bladder/GSM3723358

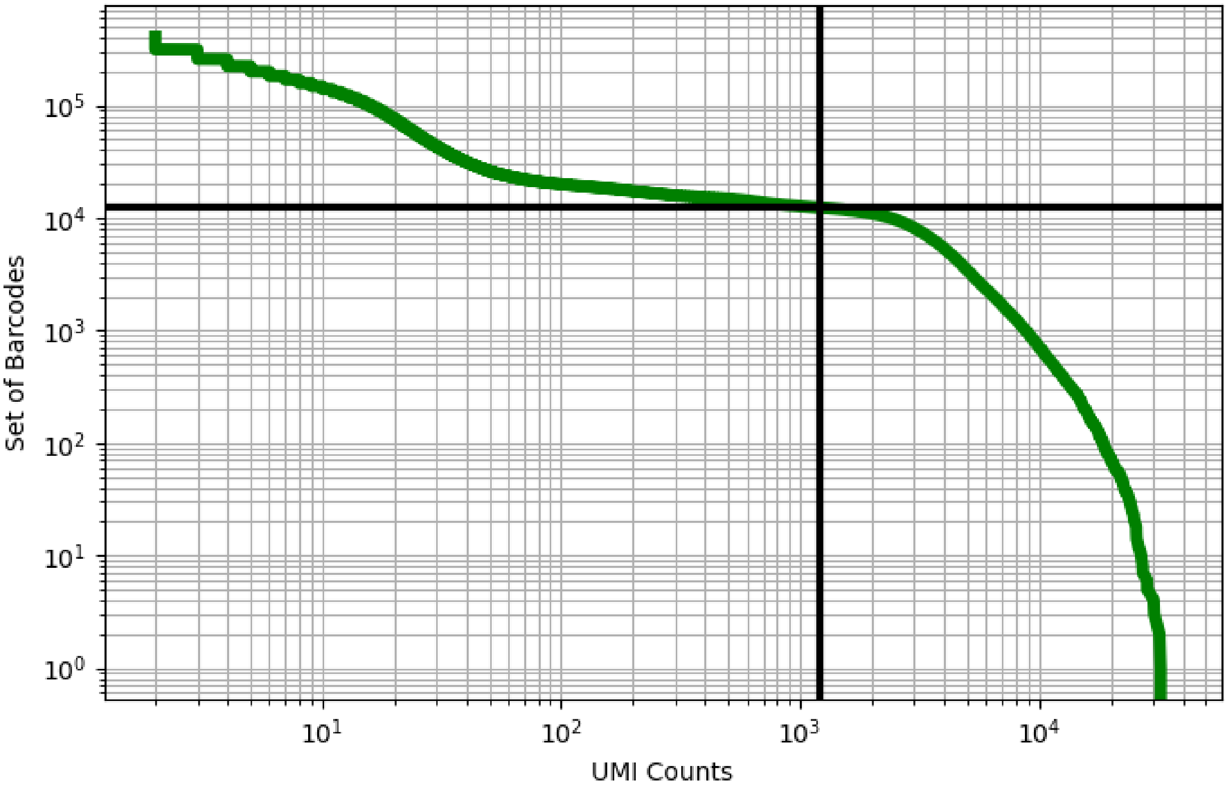


Figure 35: bladder/GSM3723359

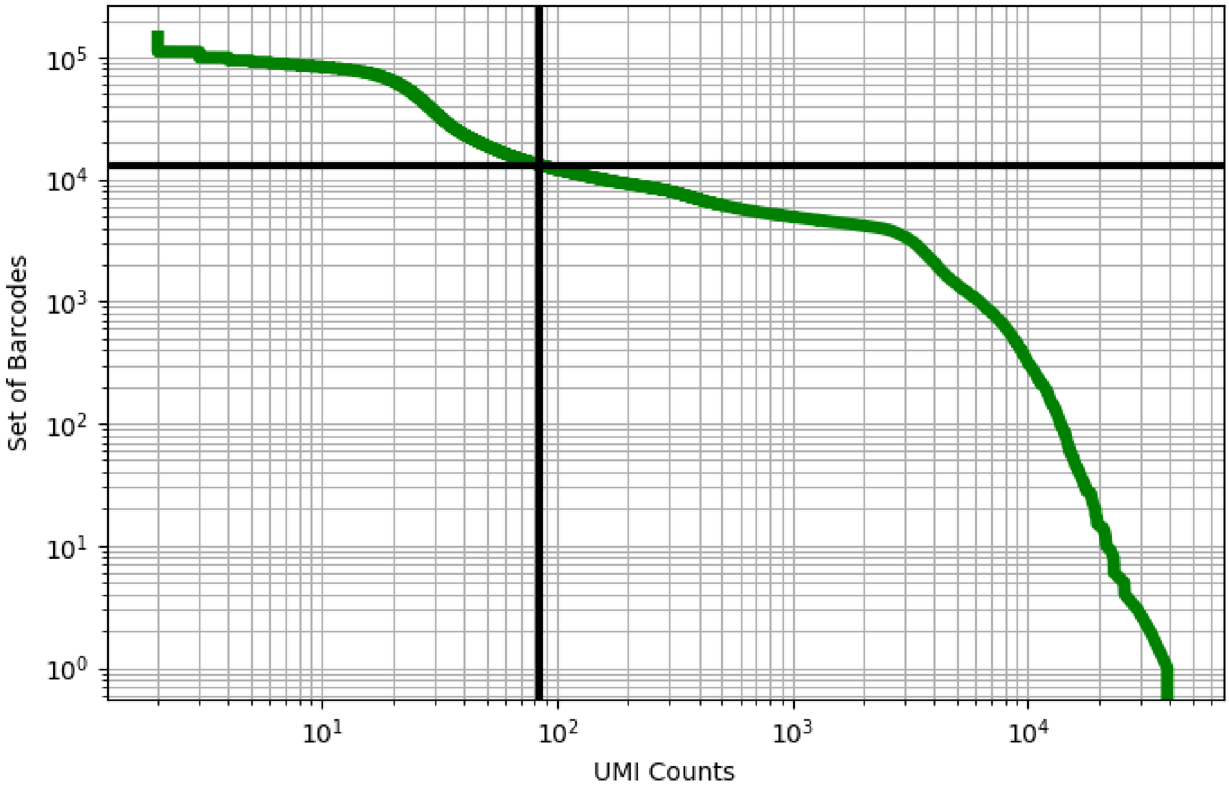


Figure 36: blood/CRX102285

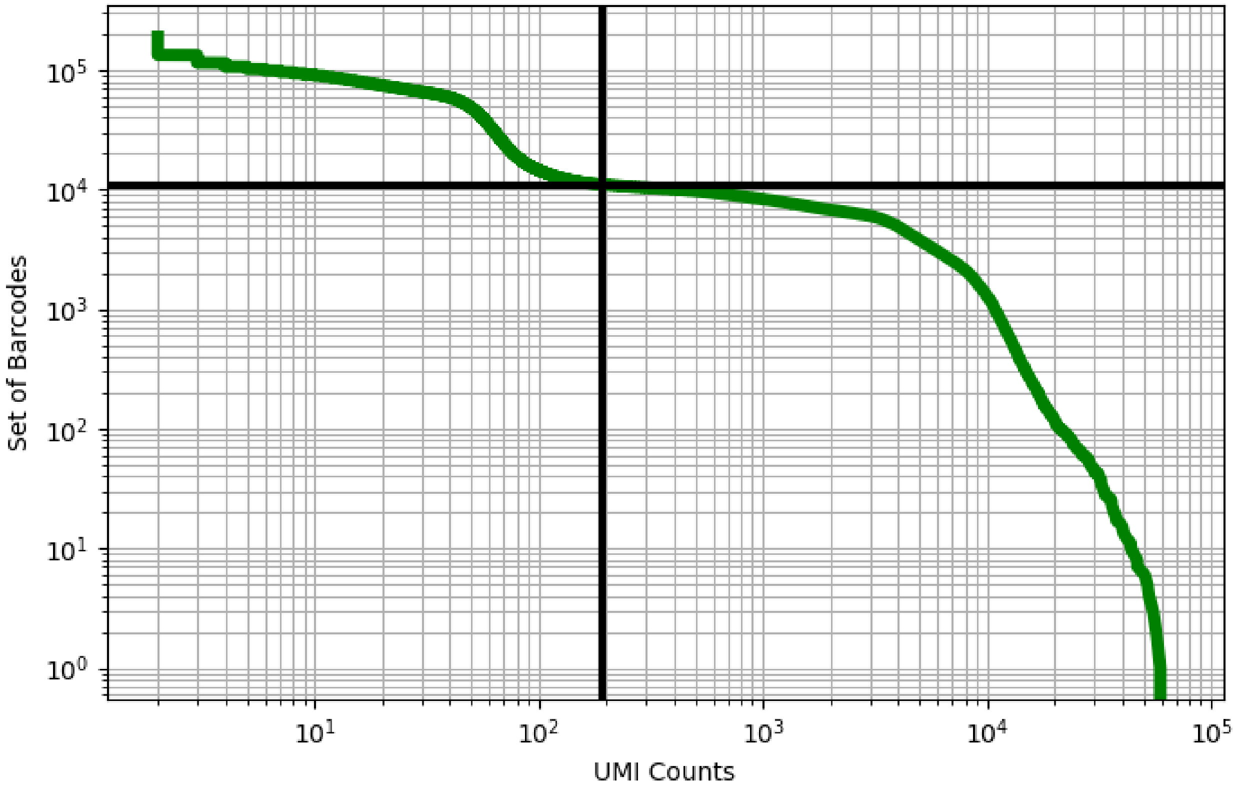


Figure 37: blood/CRX102286

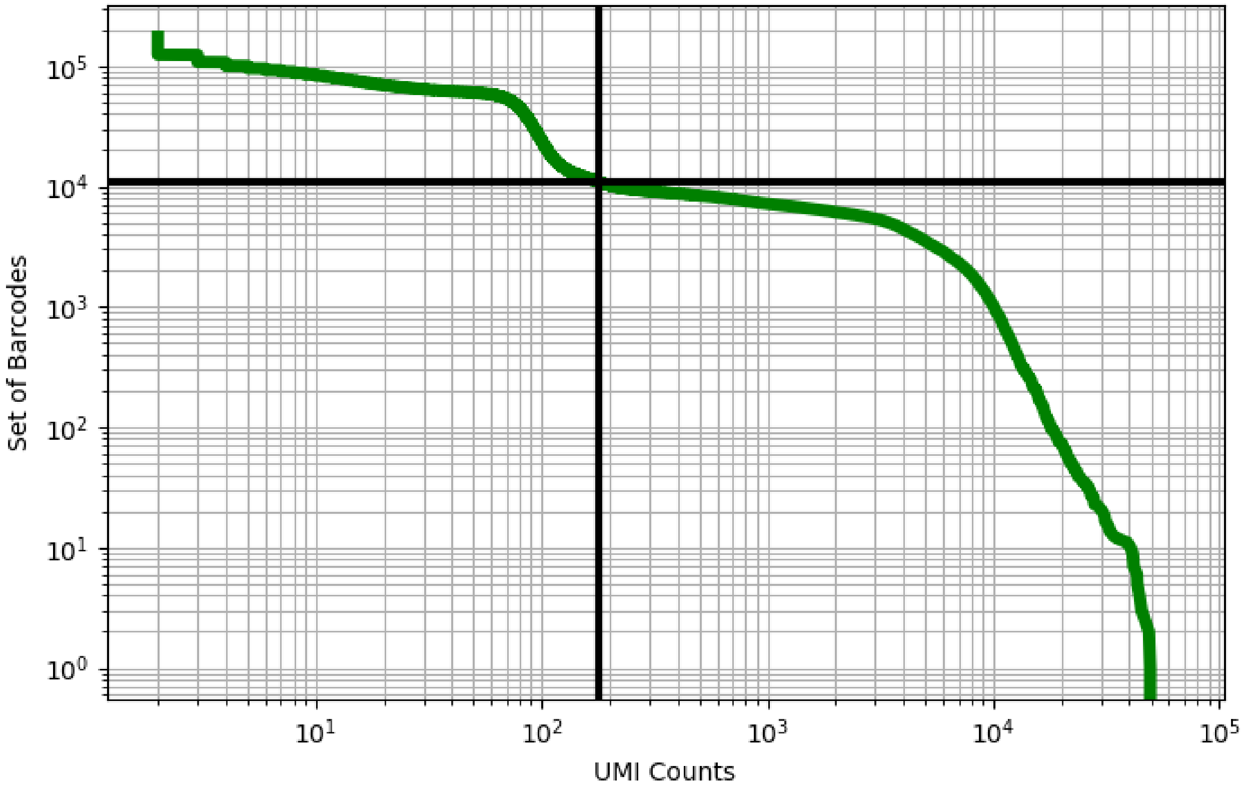


Figure 38: blood/CRX102287

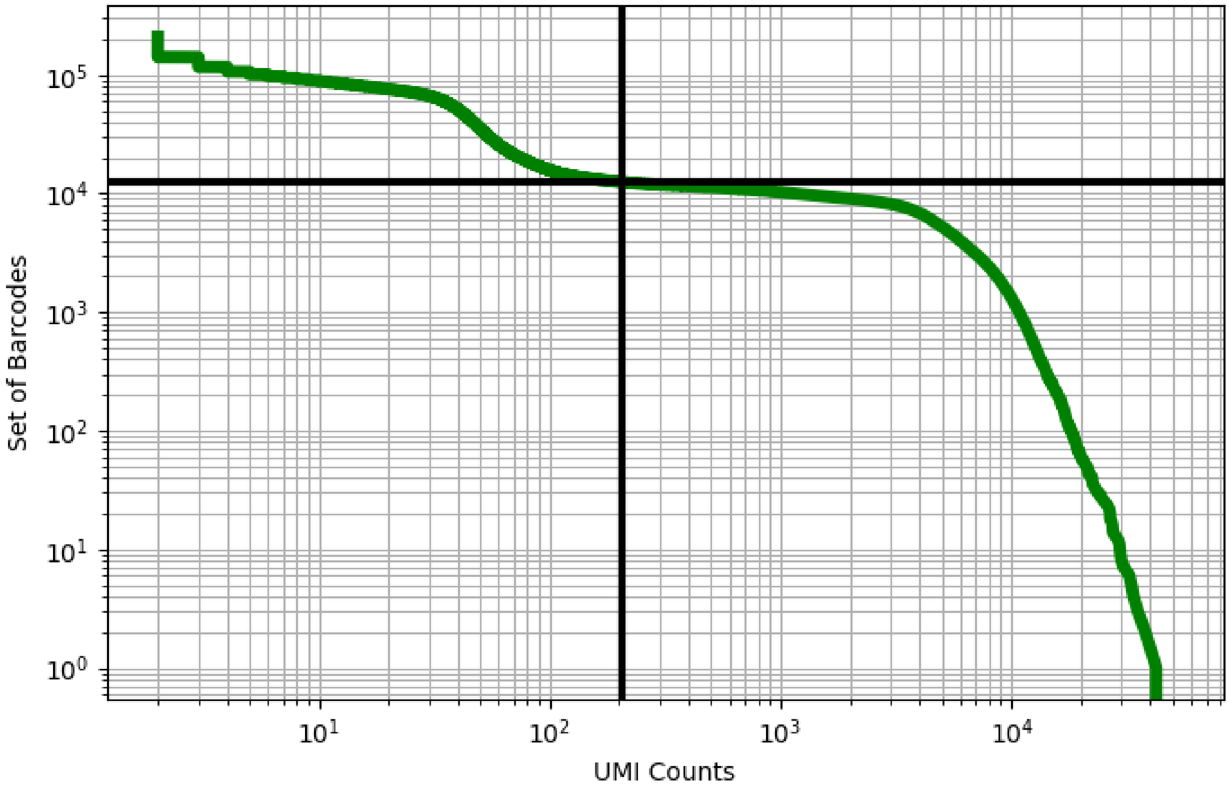


Figure 39: blood/CRX102288

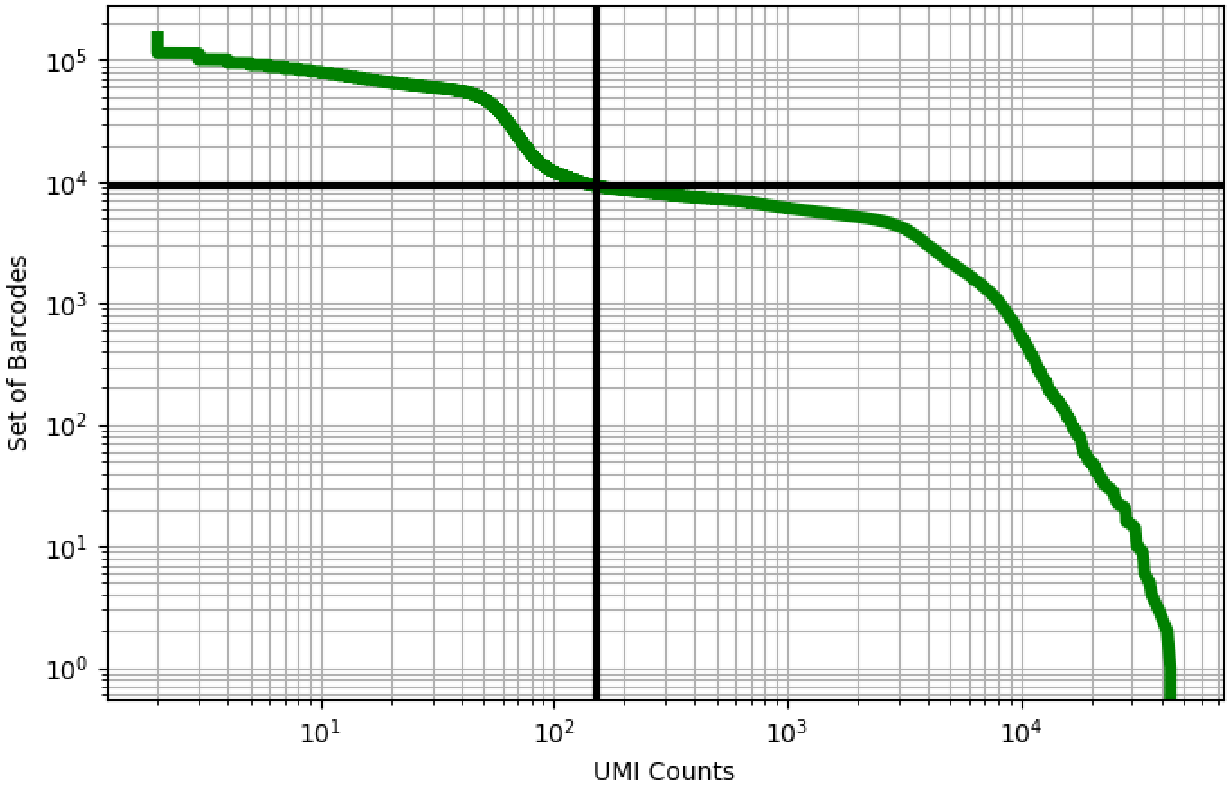


Figure 40: blood/CRX102289

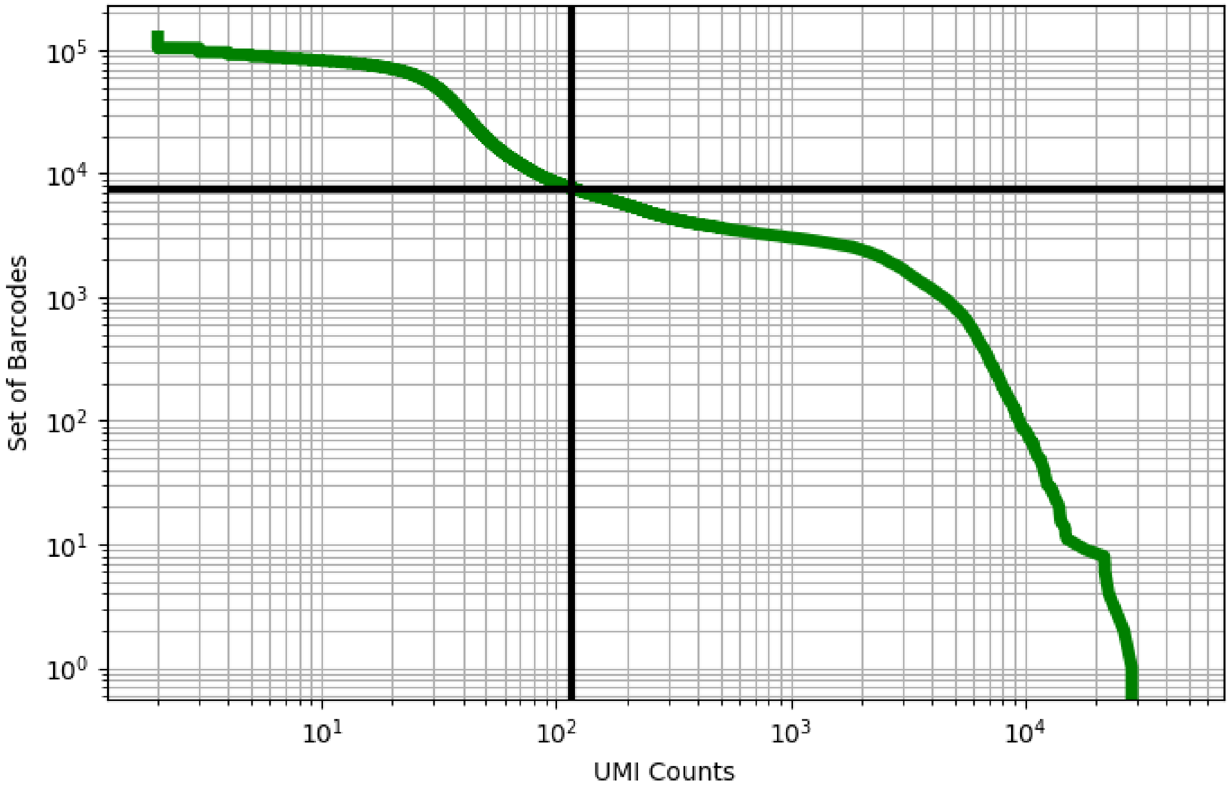


Figure 41: blood/CRX102290

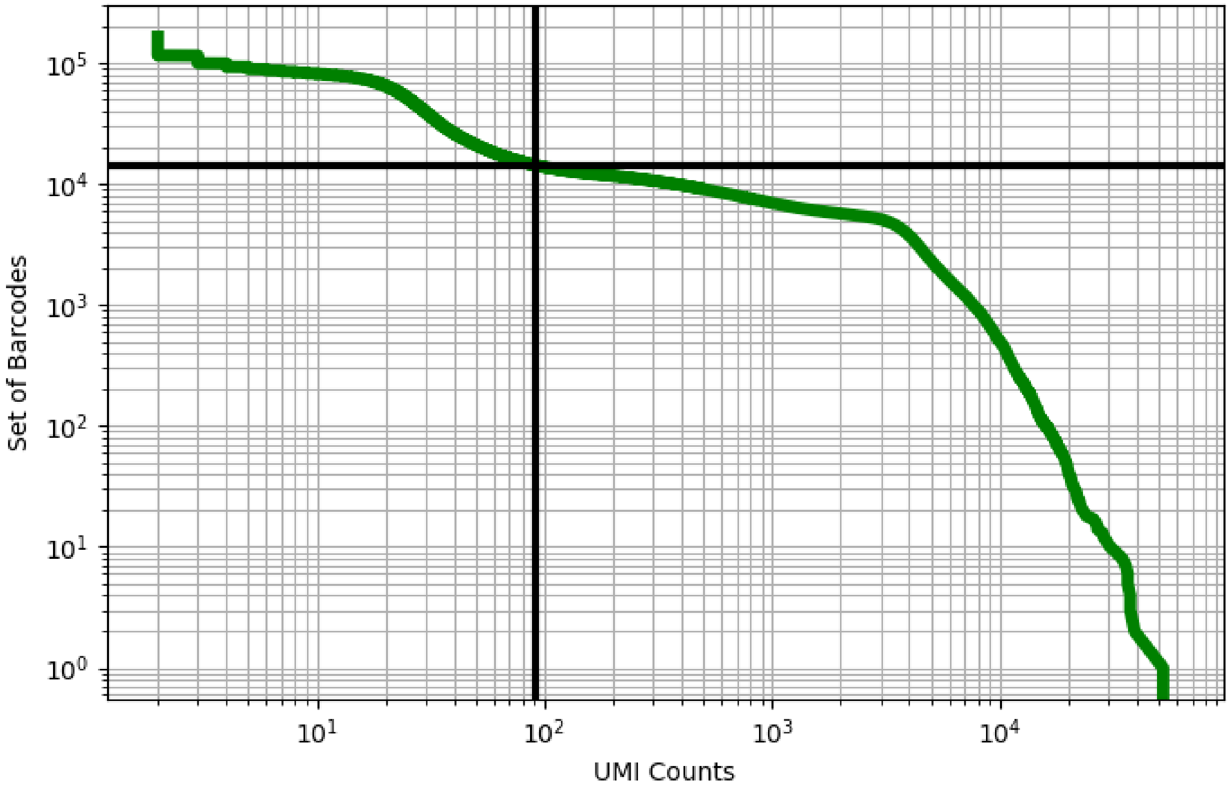


Figure 42: blood/CRX102292

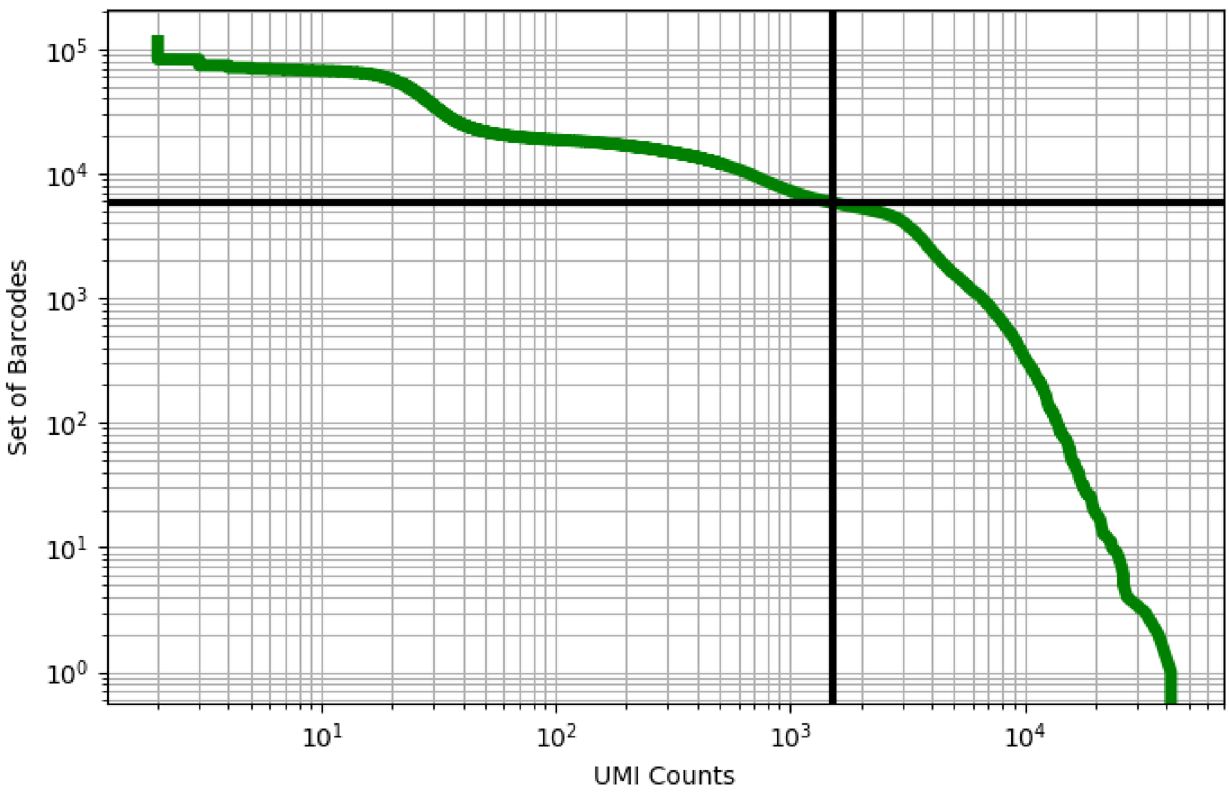


Figure 43: blood/CRX102293

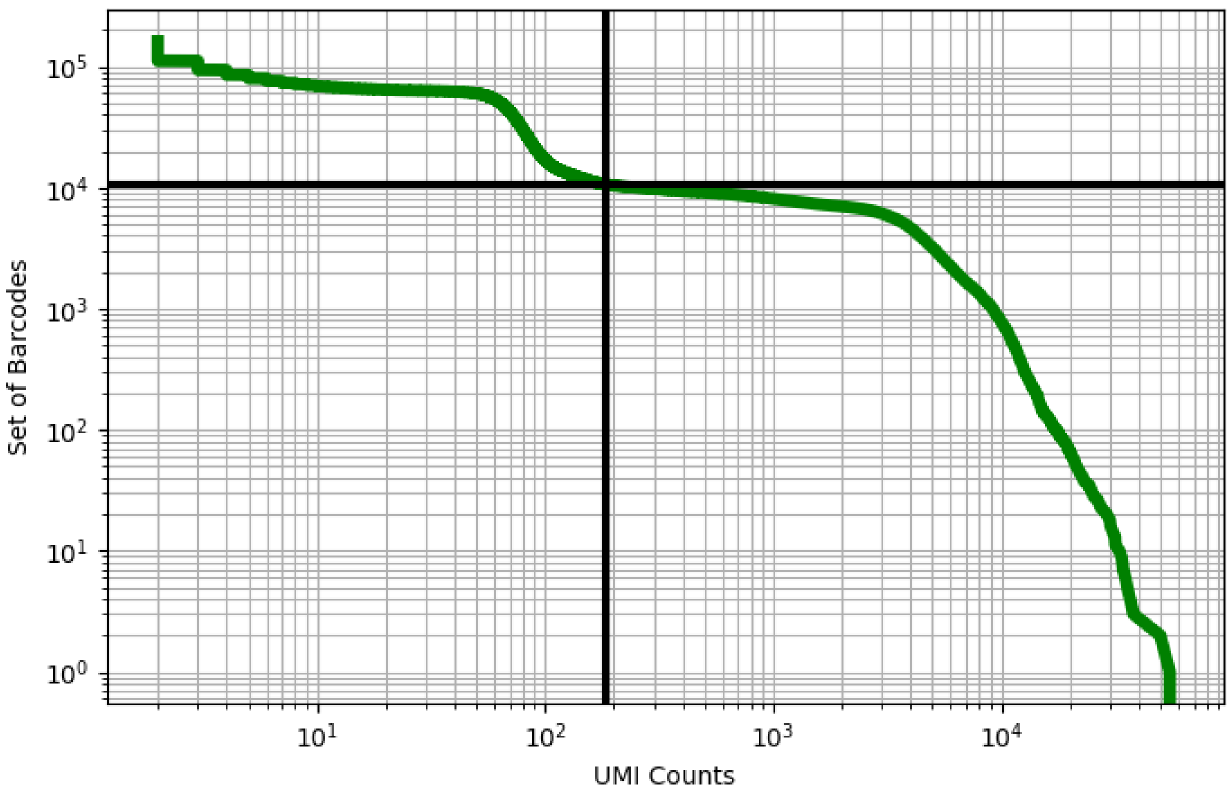


Figure 44: blood/CRX102294

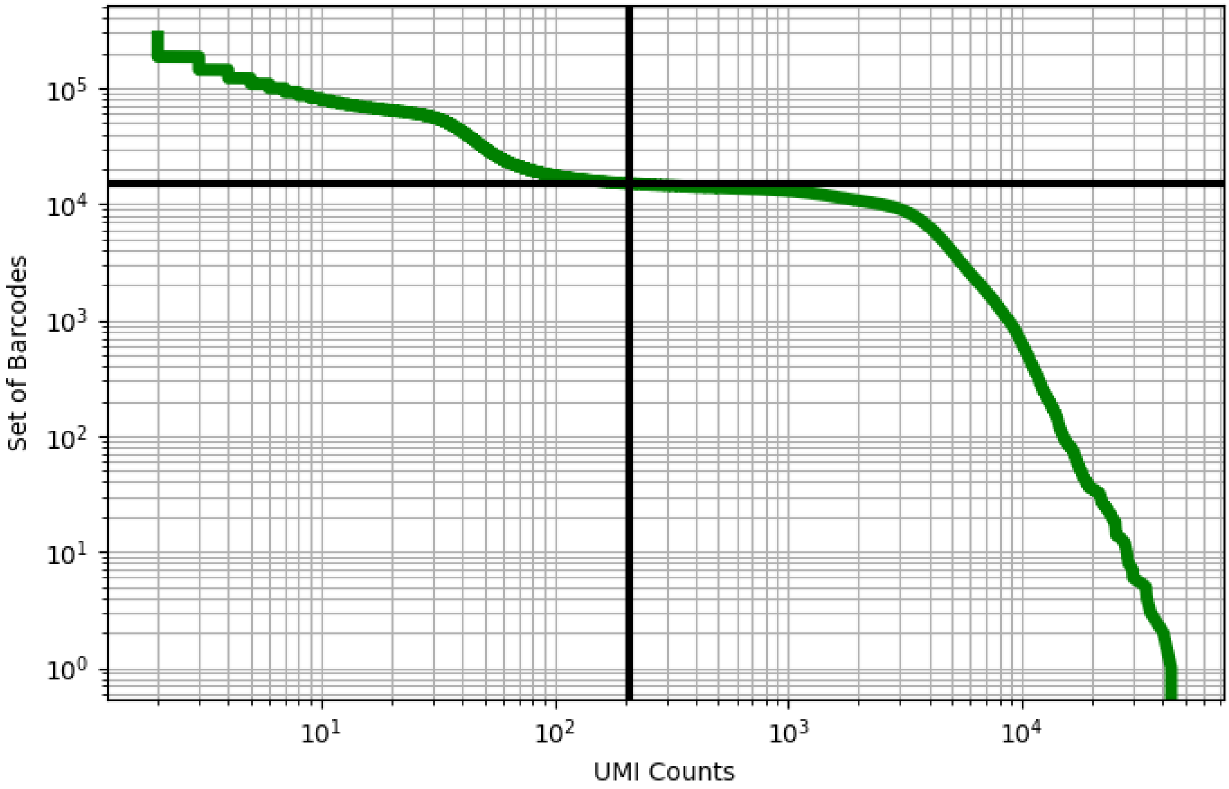


Figure 45: blood/CRX118012

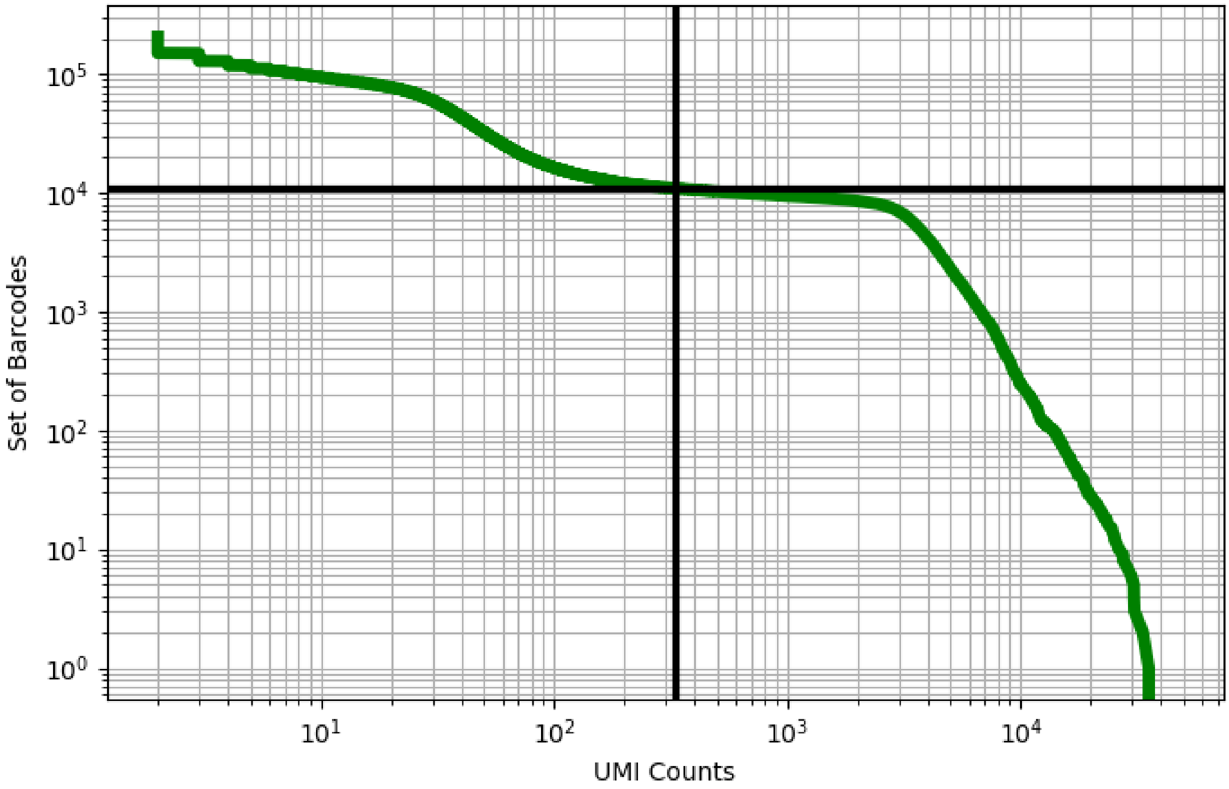


Figure 46: blood/CRX118013

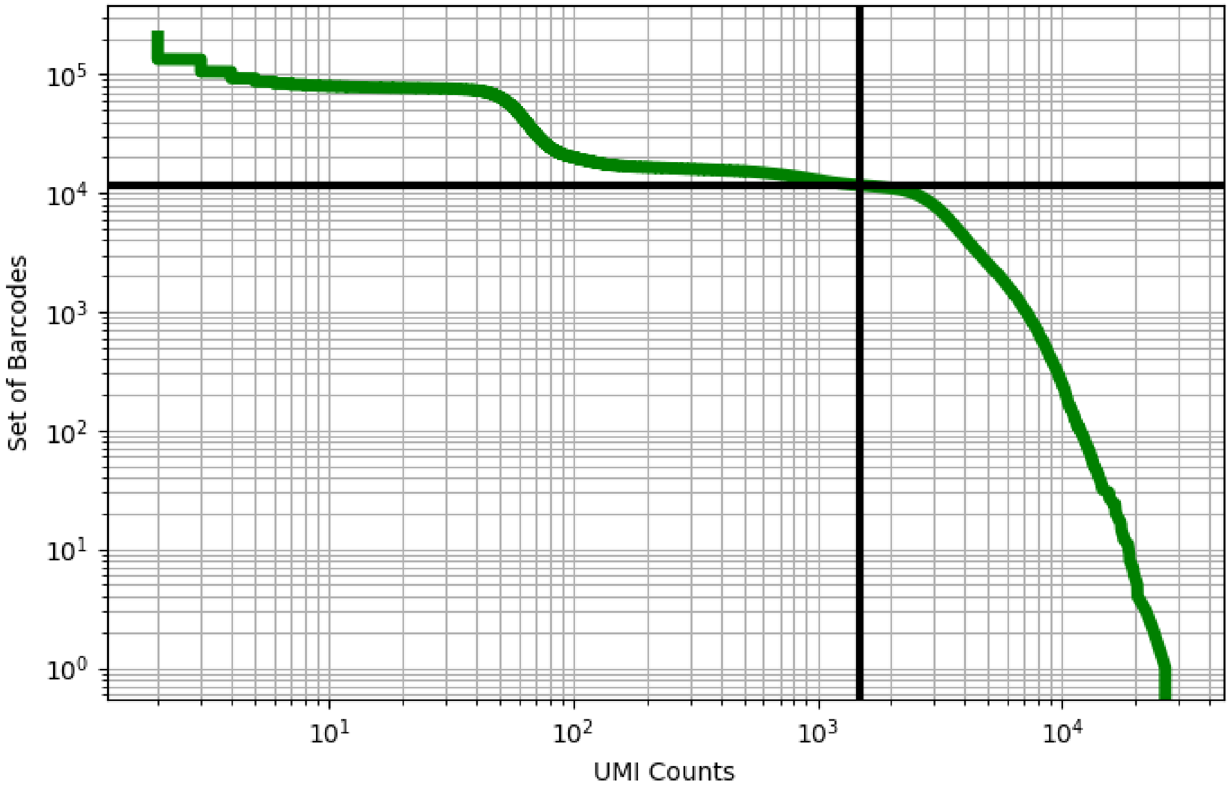


Figure 47: blood/CRX118014

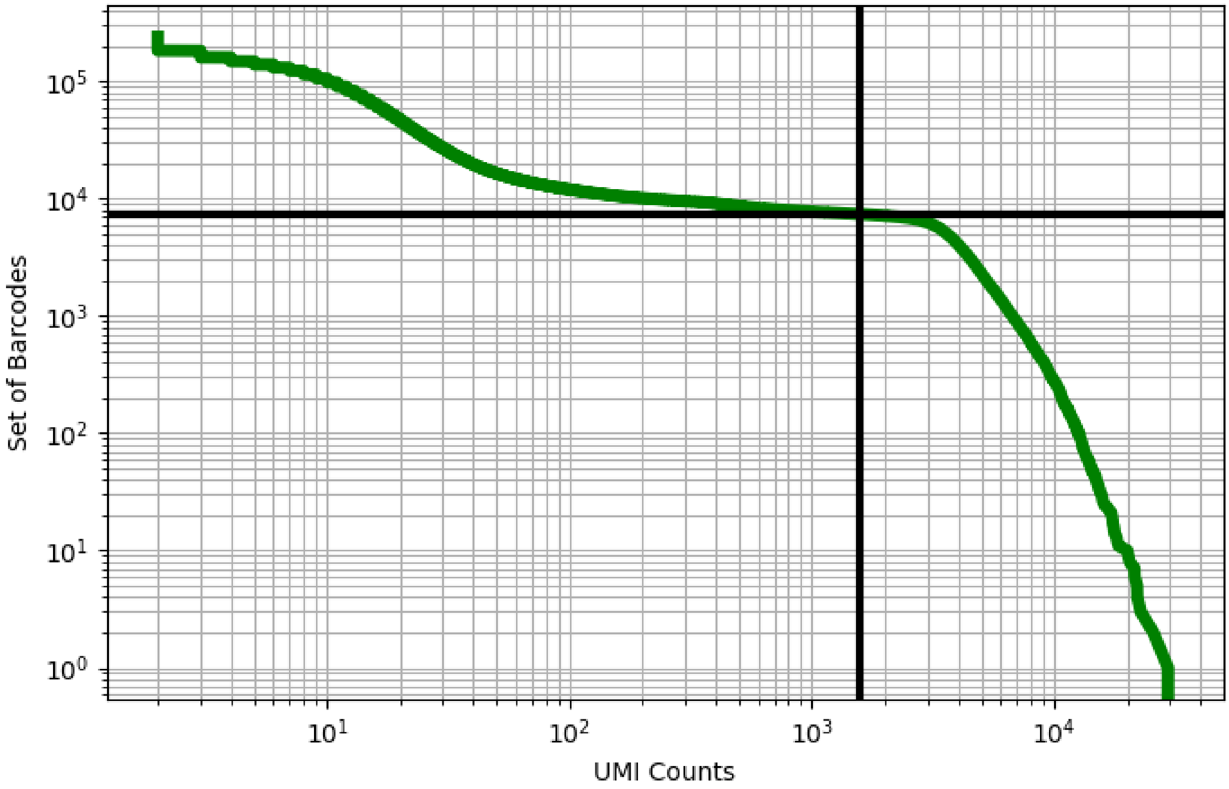


Figure 48: blood/CRX118015

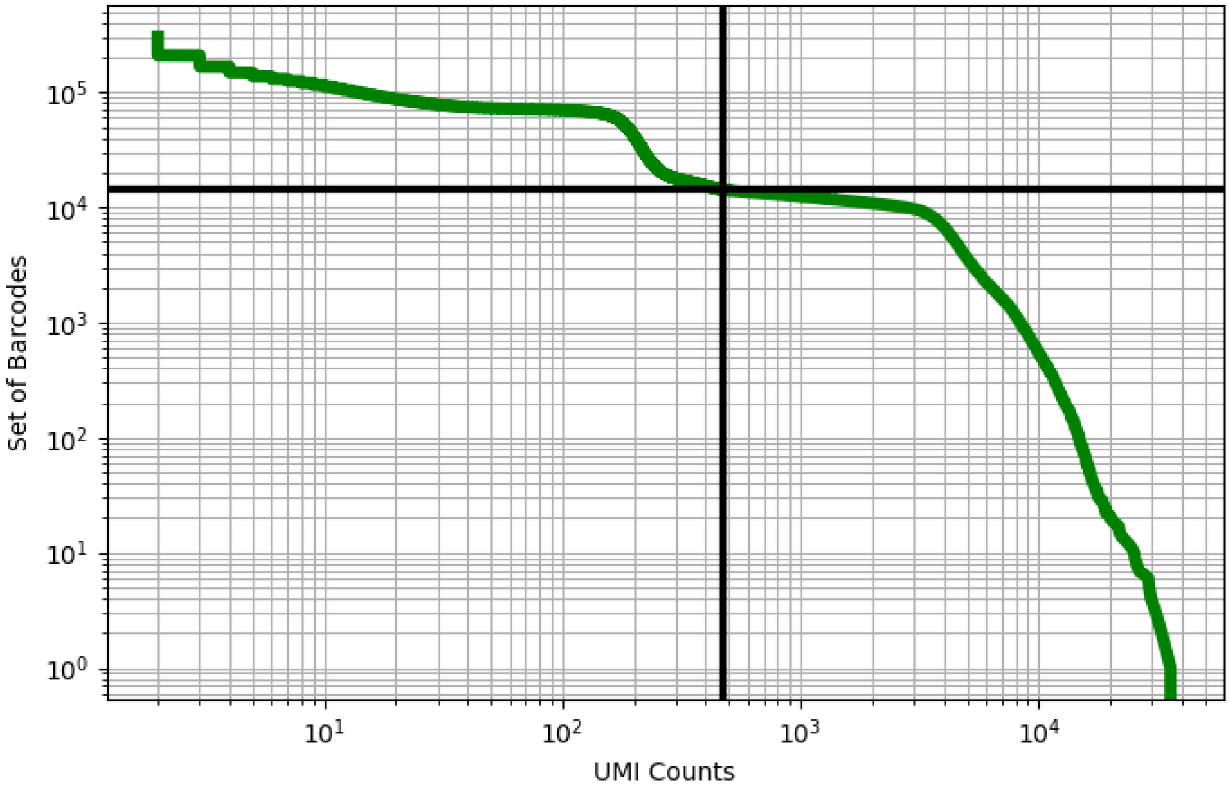


Figure 49: blood/CRX118016

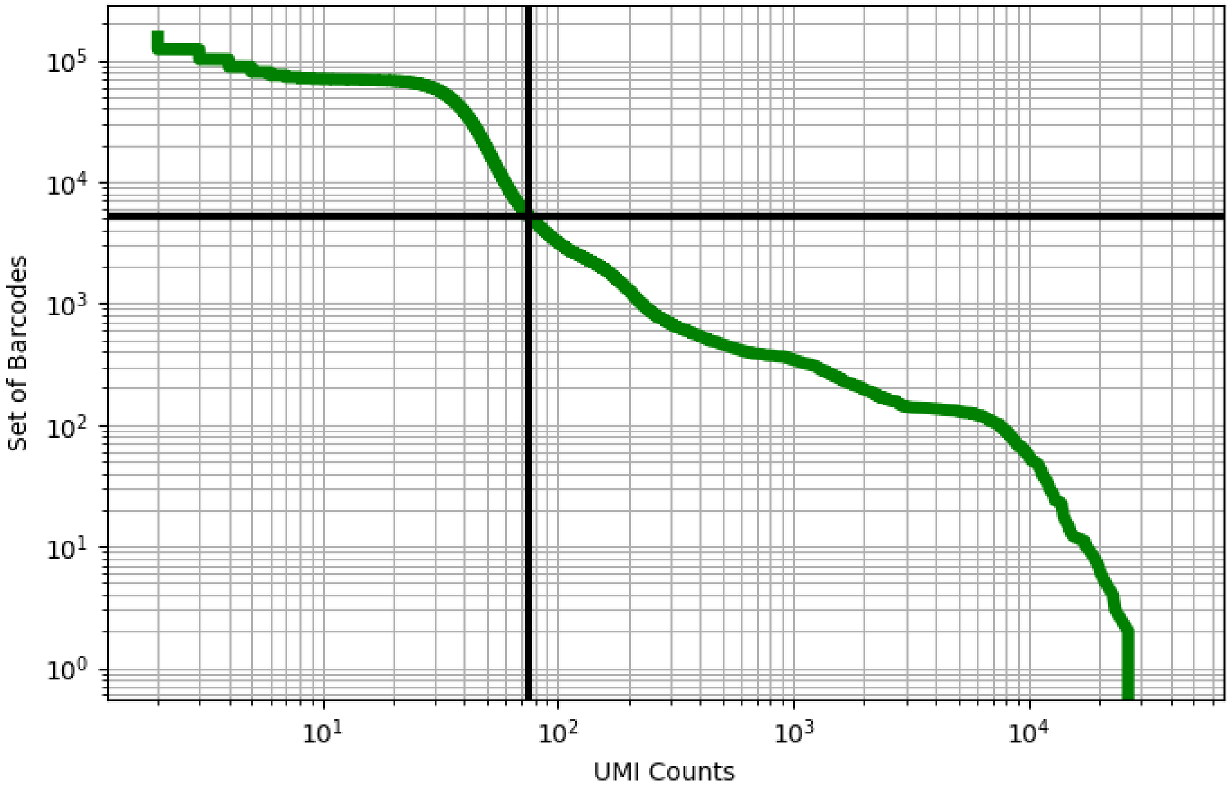


Figure 50: blood/ERX2756718

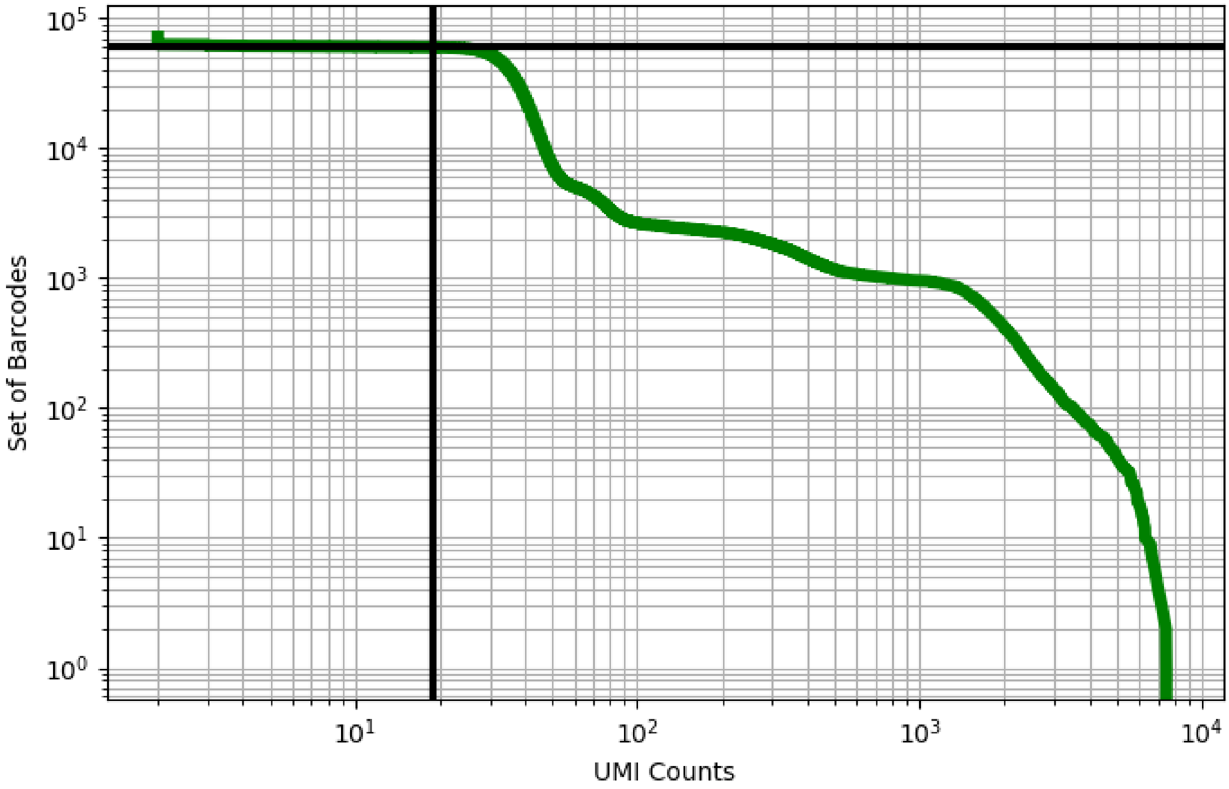


Figure 51: blood/ERX2756720

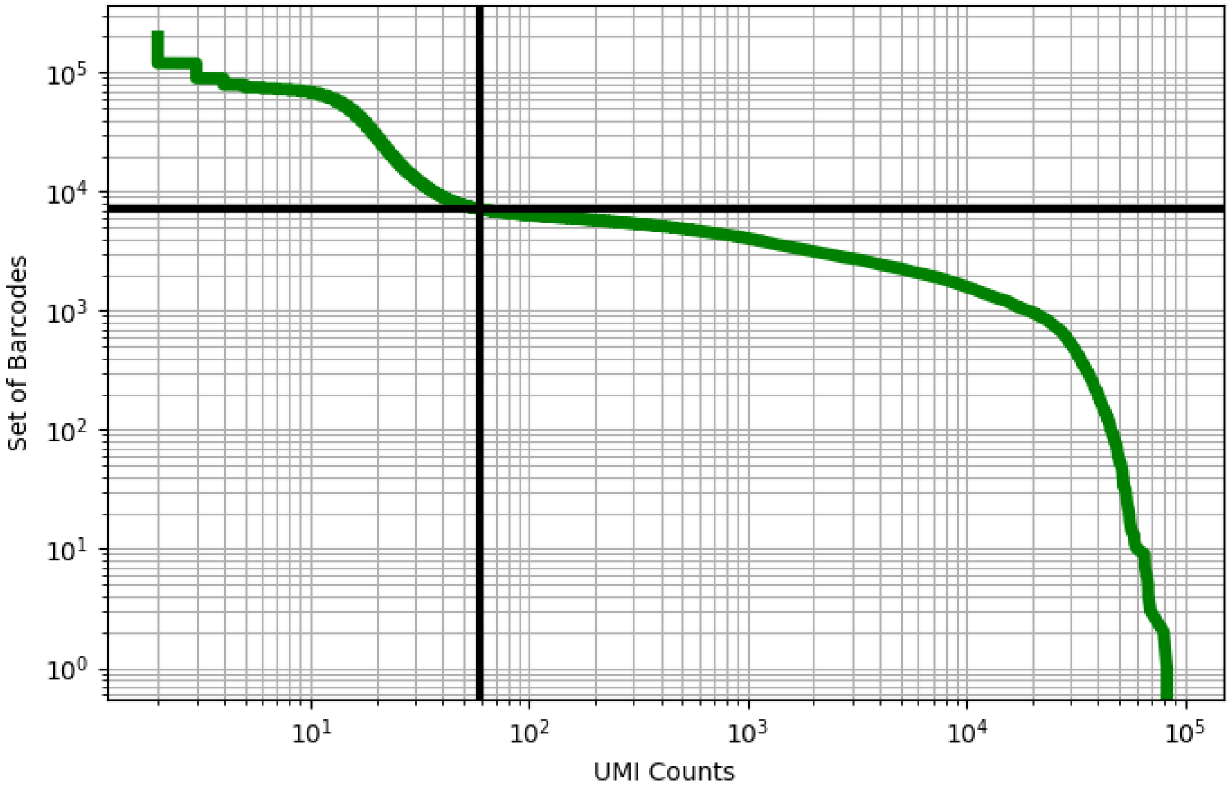


Figure 52: blood/ERX2756727

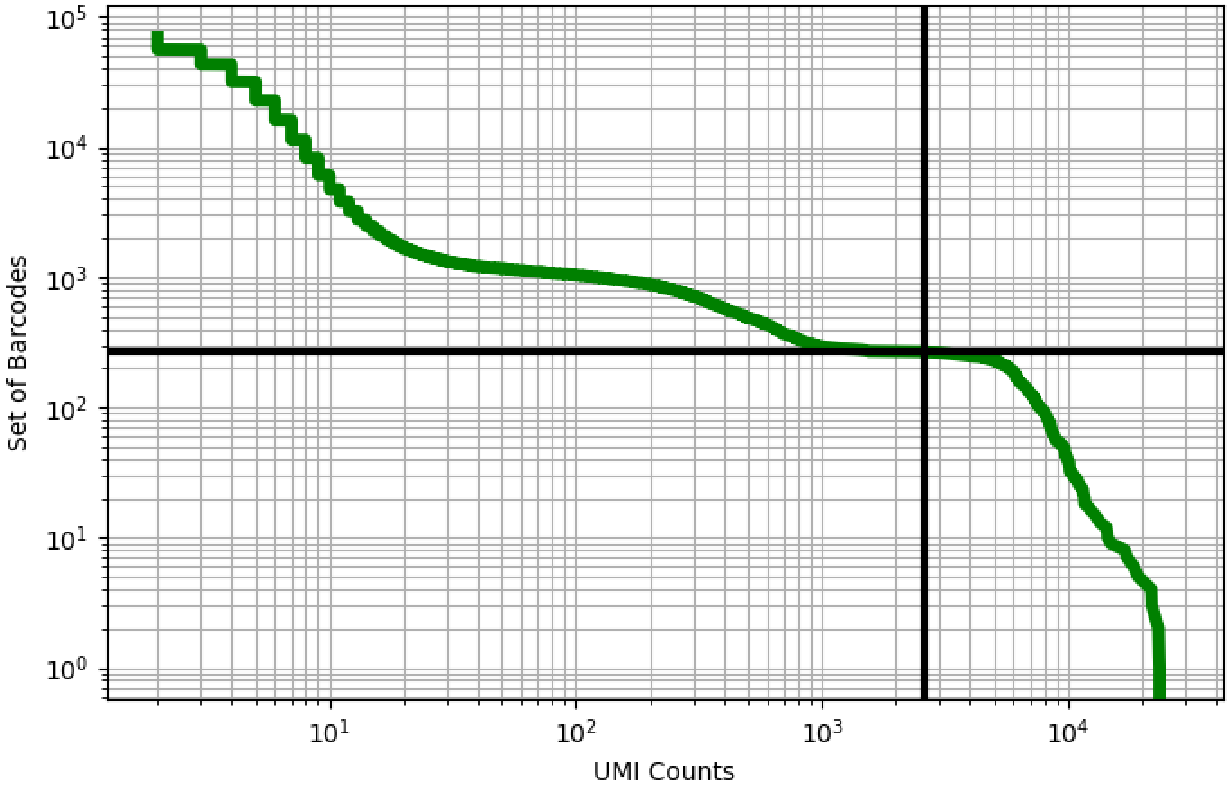


Figure 53: blood/GSM2803967

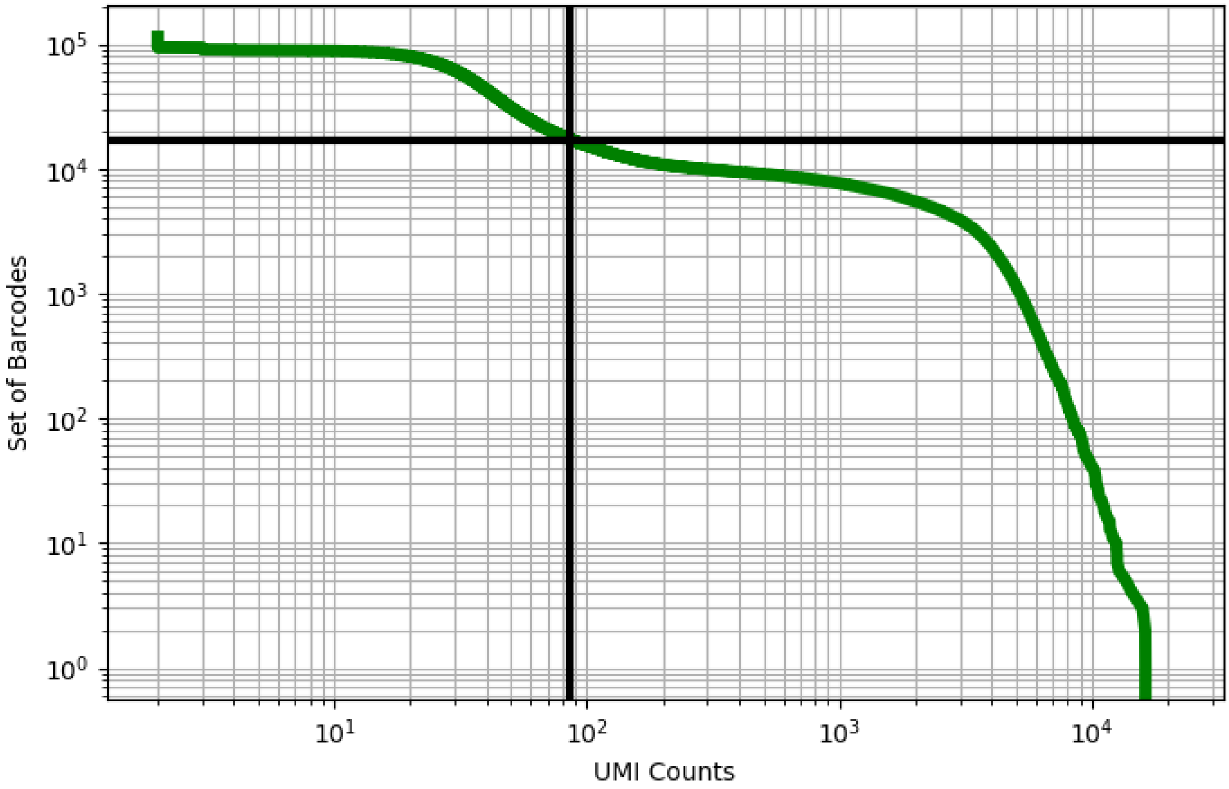


Figure 54: blood/GSM2877959

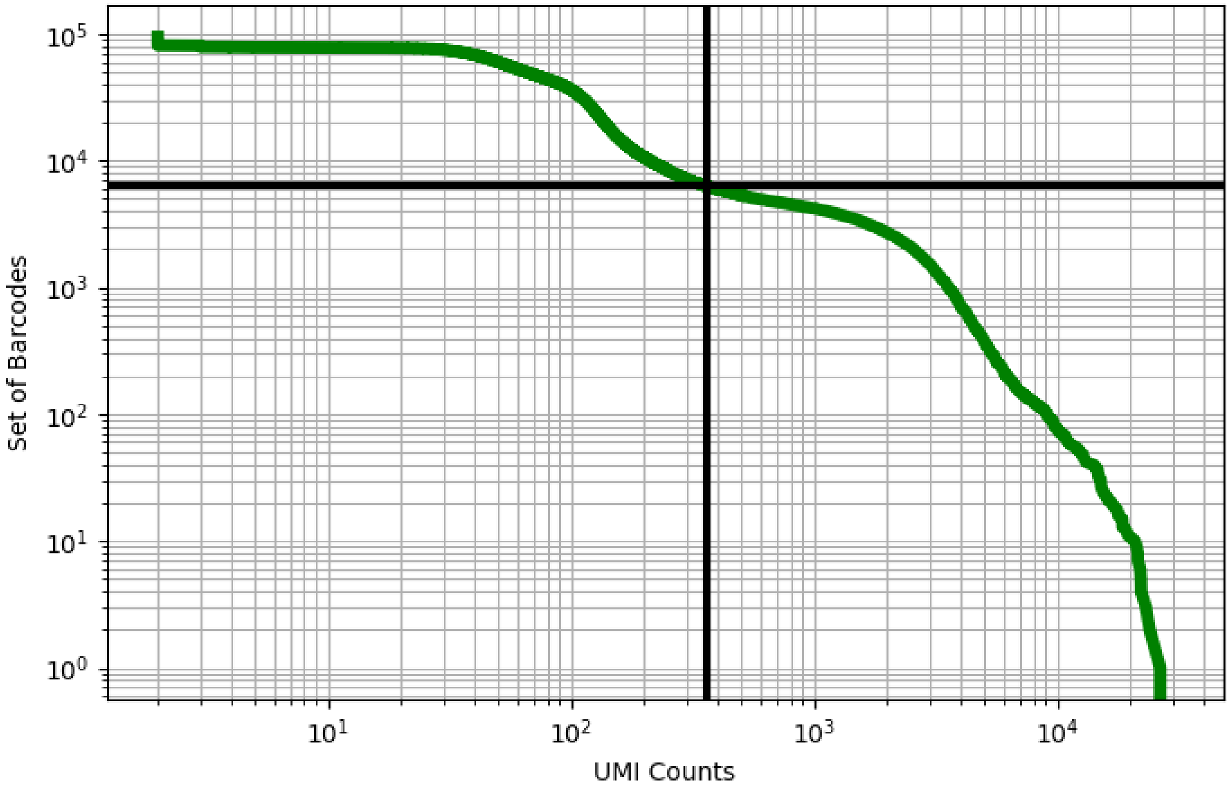


Figure 55: blood/GSM2877960

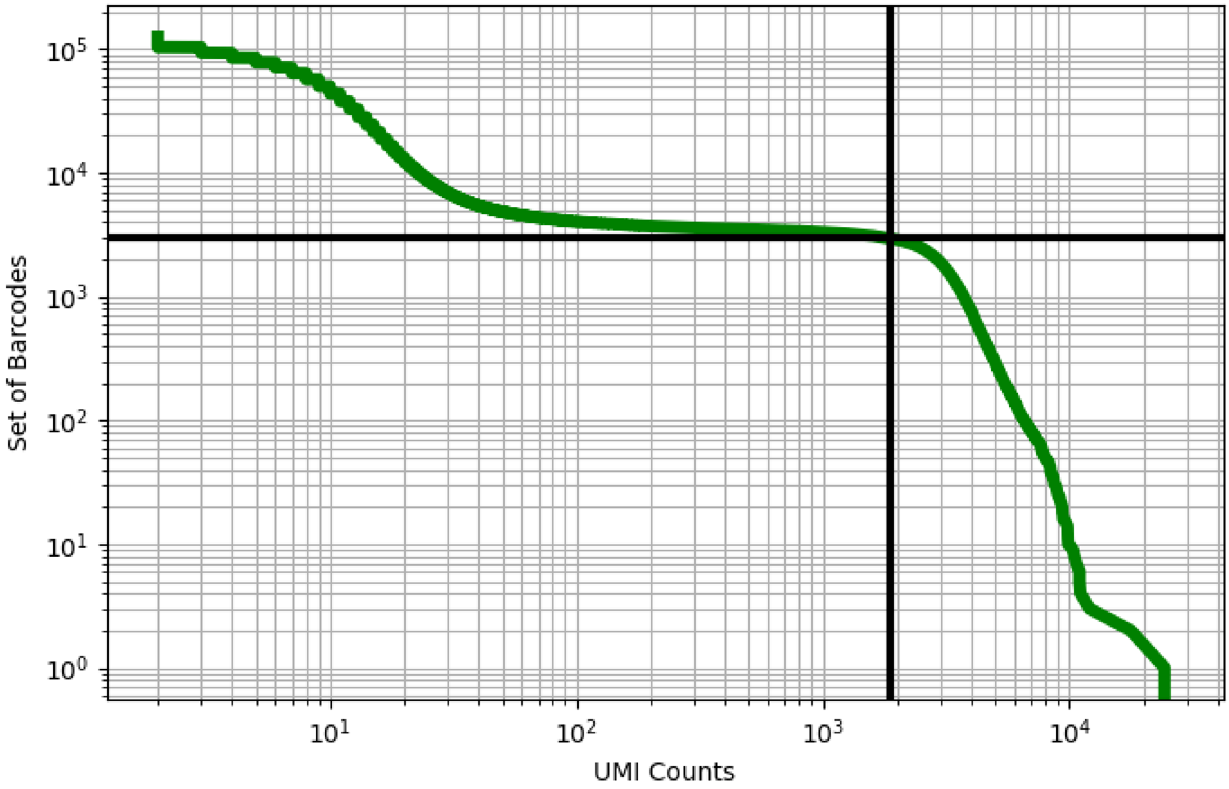


Figure 56: blood/GSM3169075

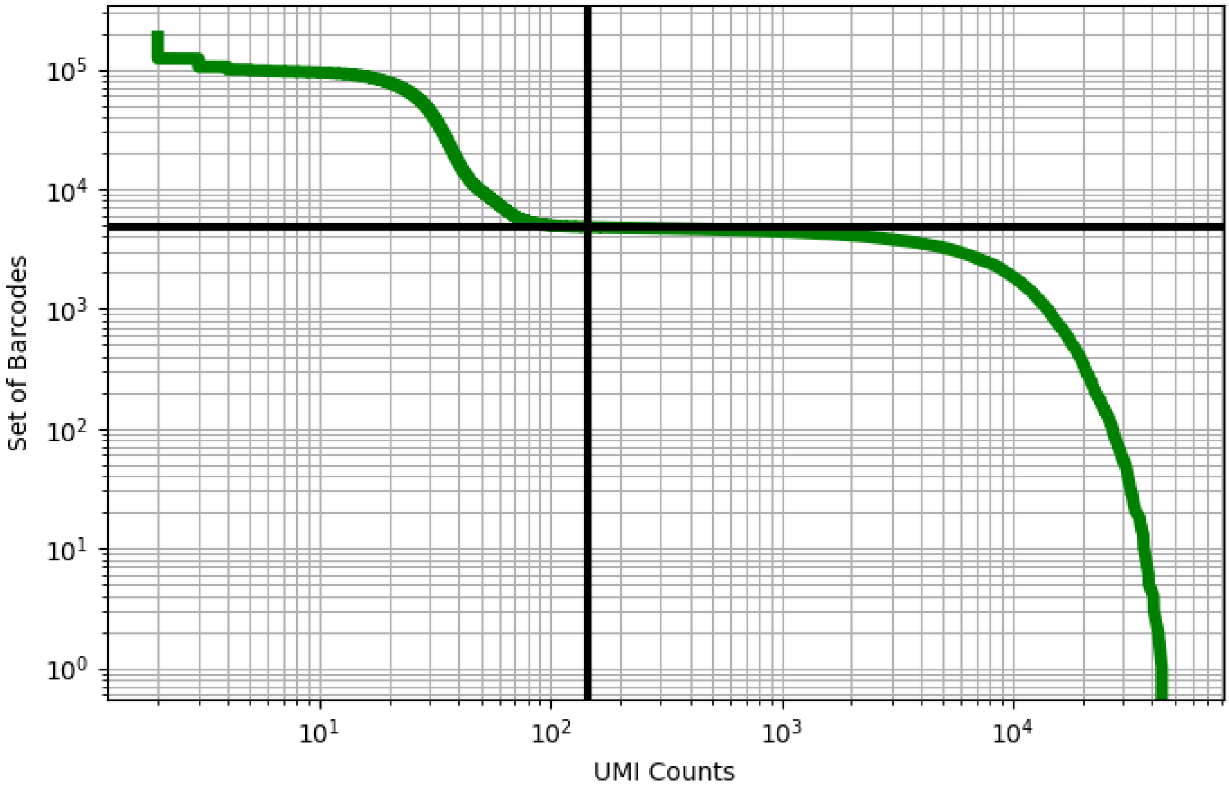


Figure 57: blood/GSM3309834

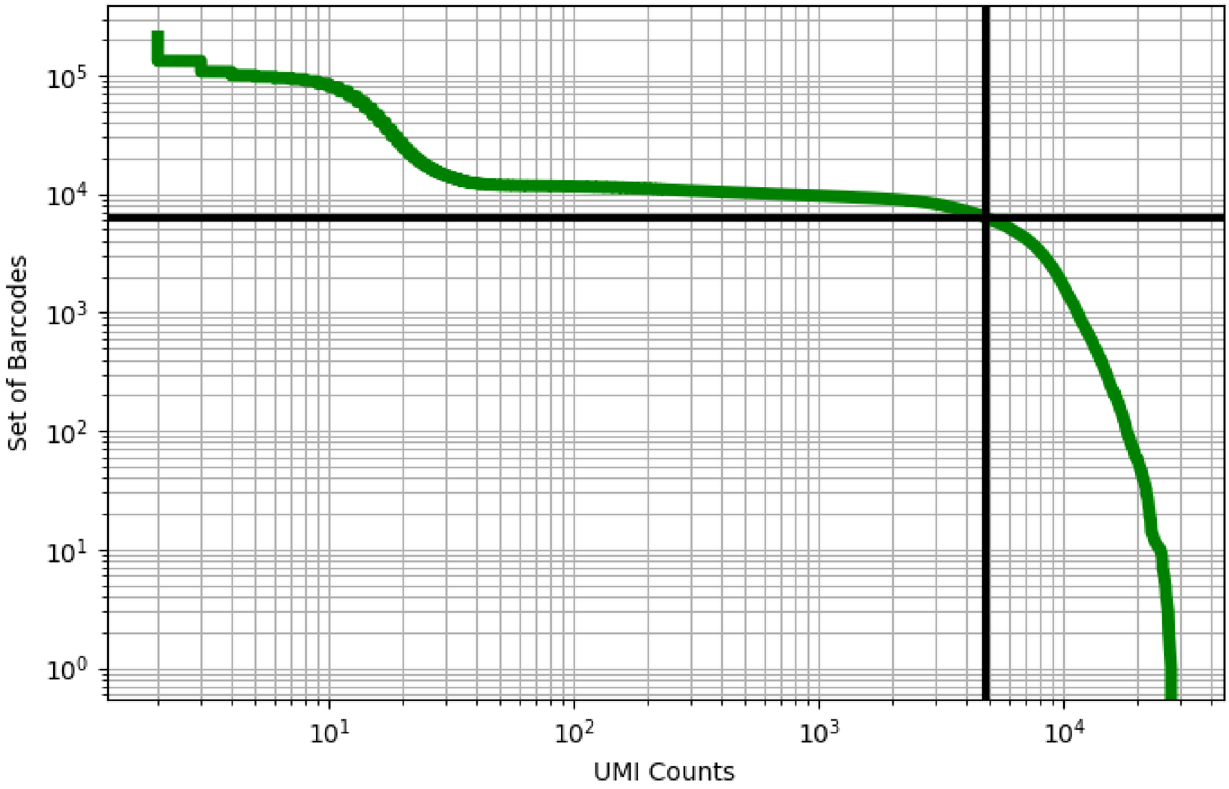


Figure 58: blood/GSM3309836

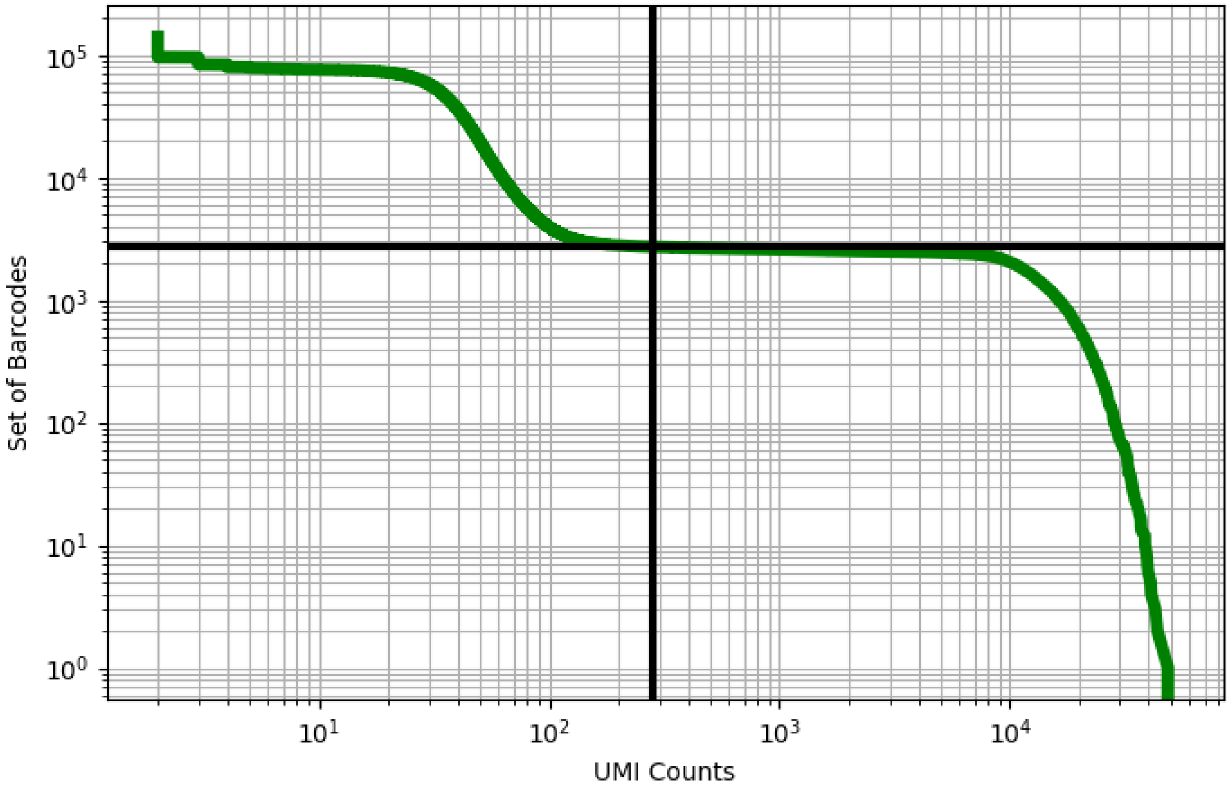


Figure 59: blood/GSM3378054

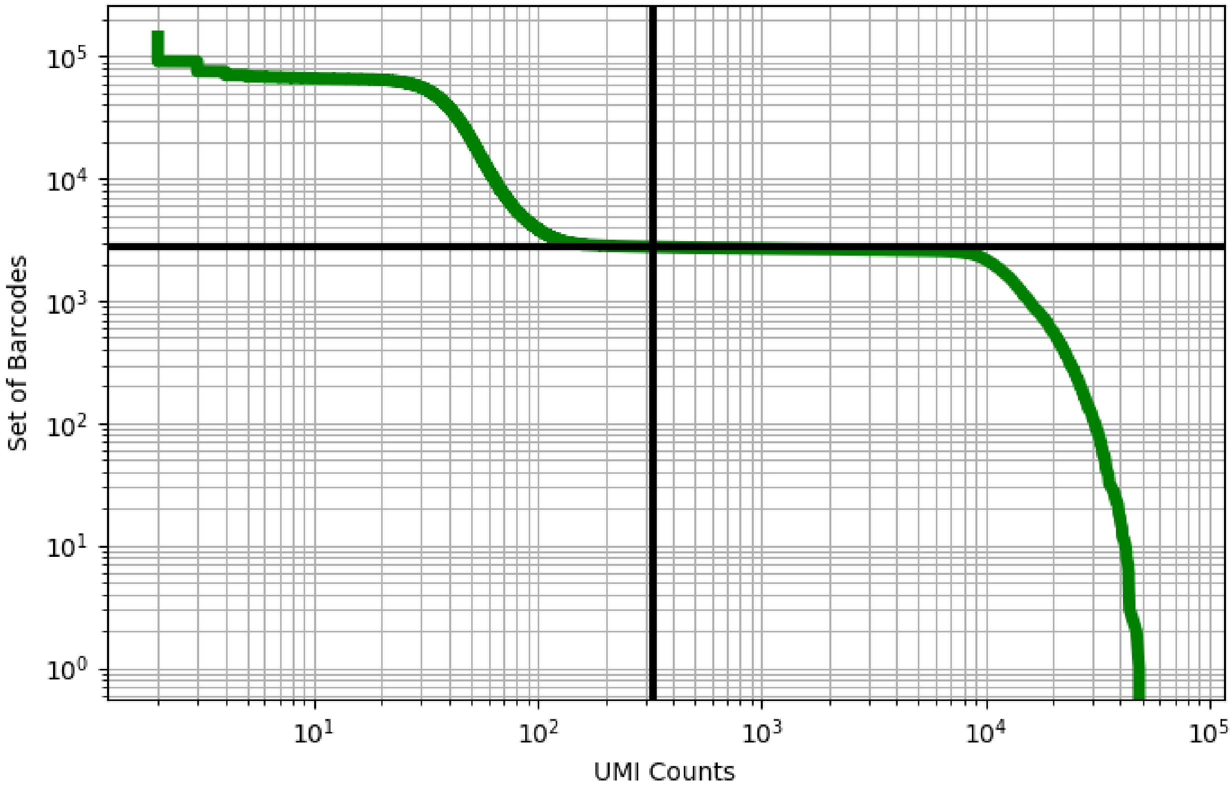


Figure 60: blood/GSM3378055

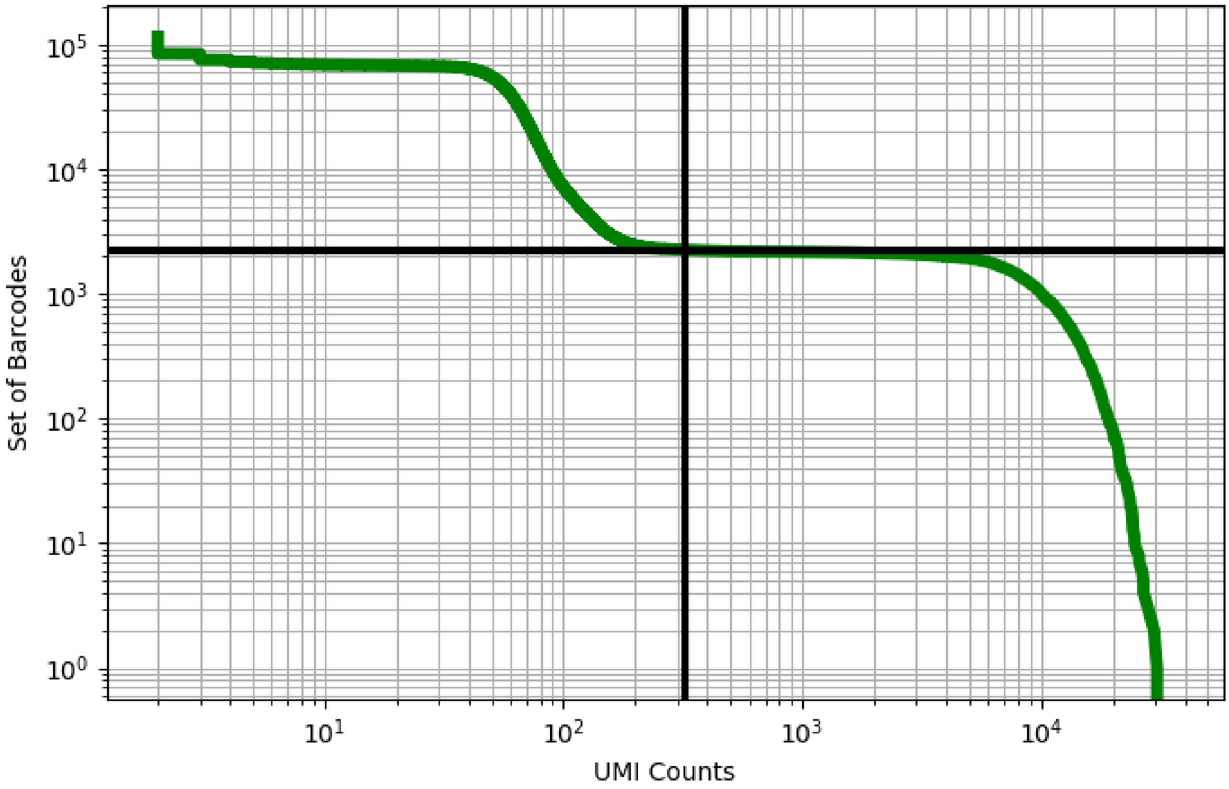


Figure 61: blood/GSM3378056

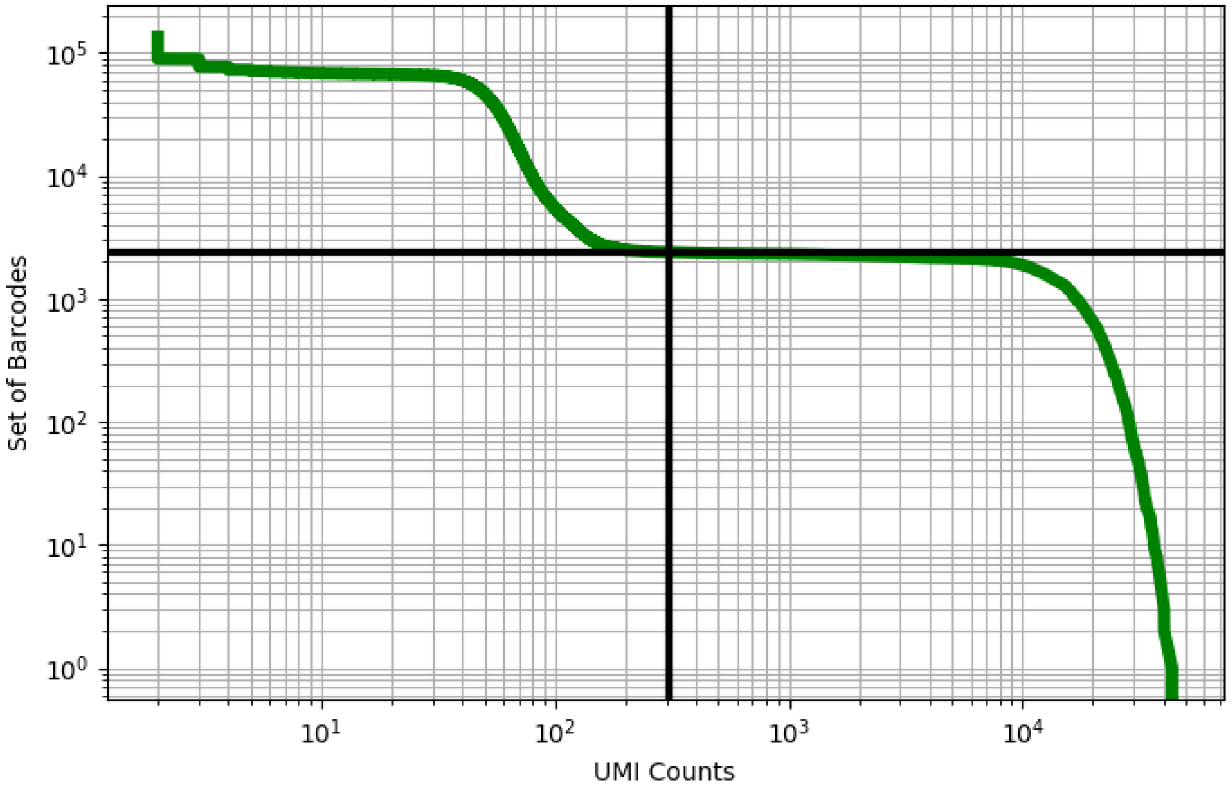


Figure 62: blood/GSM3378057

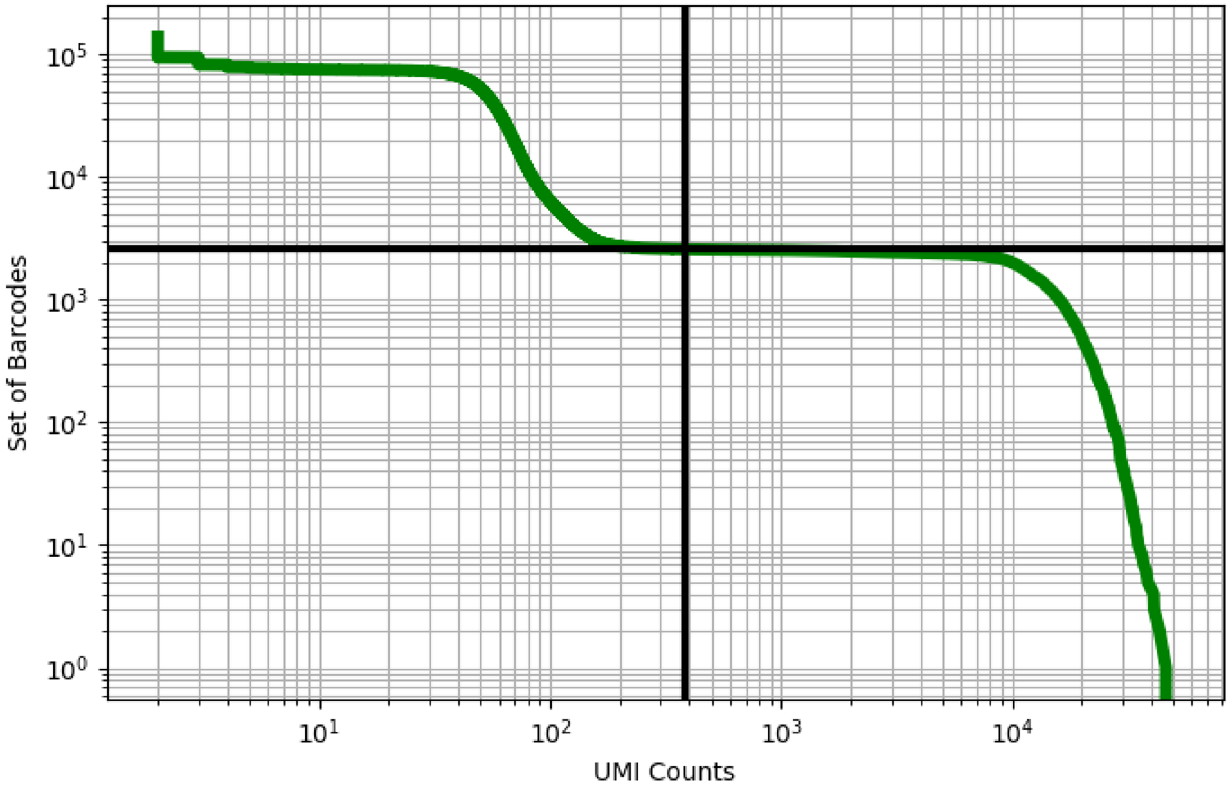


Figure 63: blood/GSM3378058

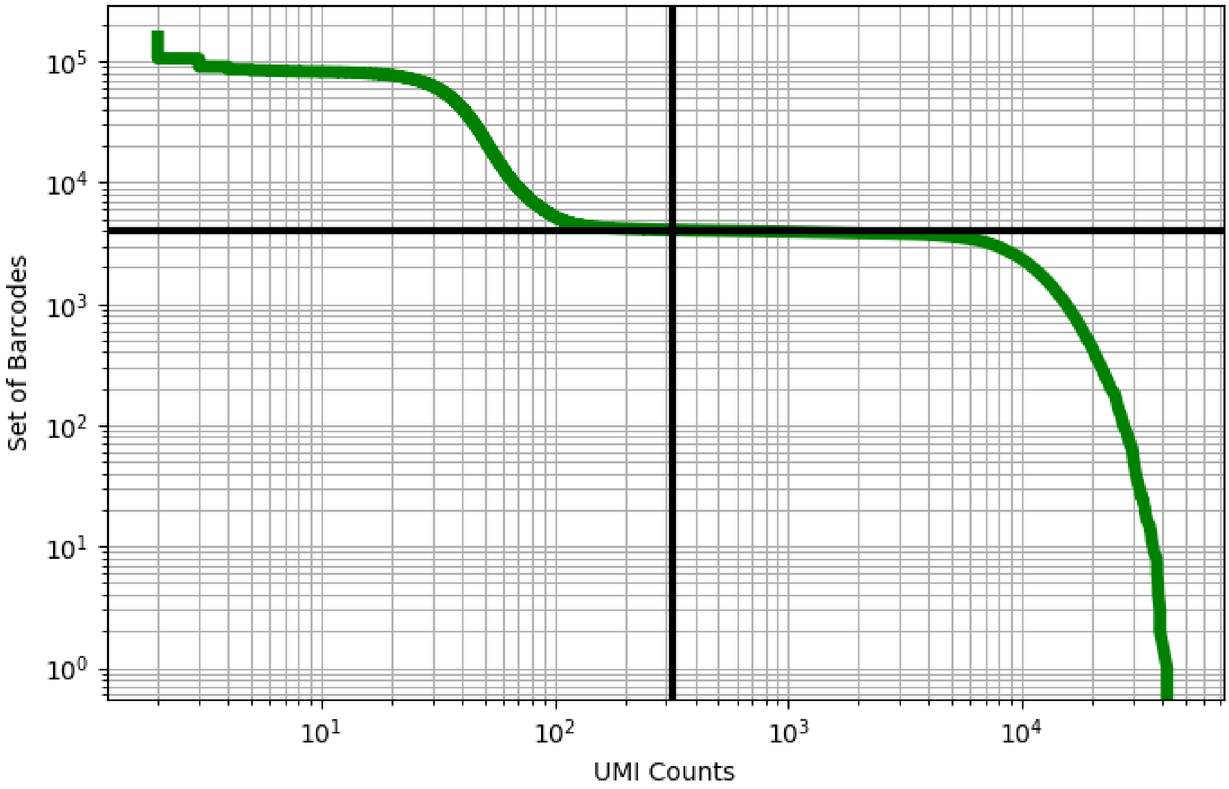


Figure 64: blood/GSM3378059

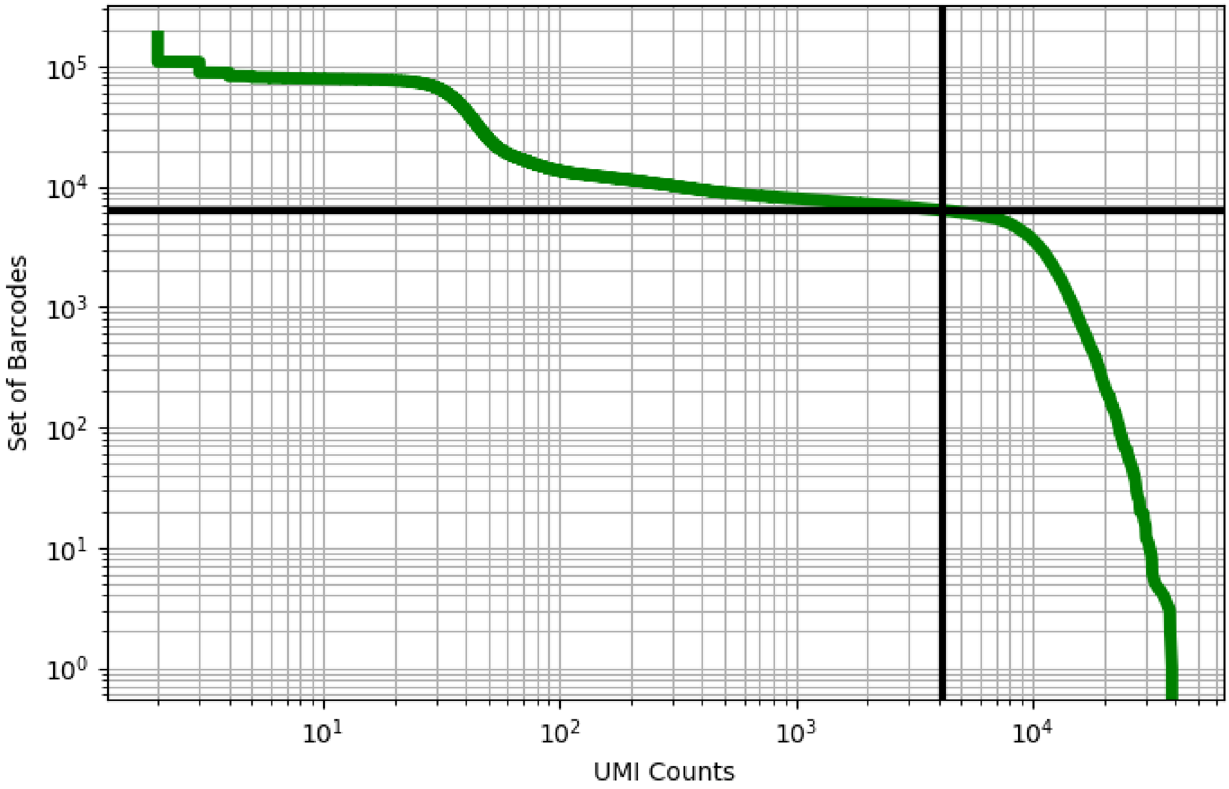


Figure 65: blood/GSM3529821

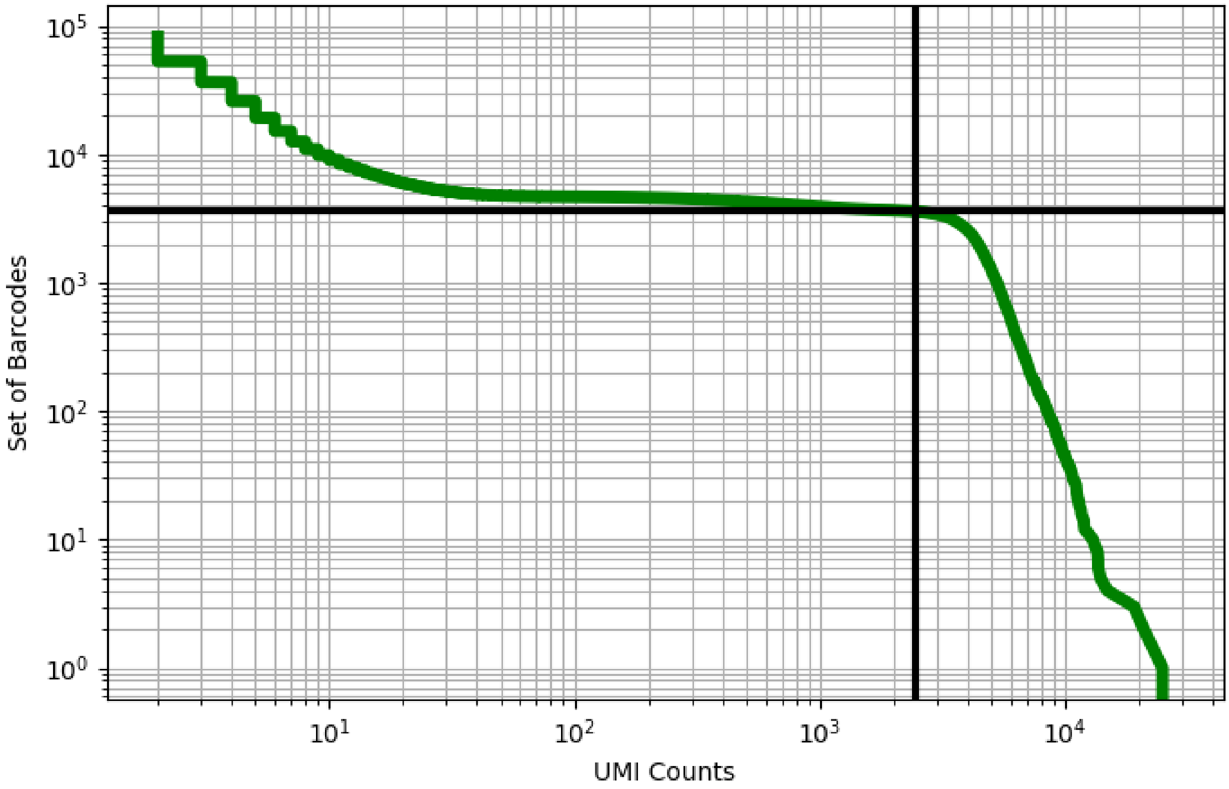


Figure 66: blood/GSM3558027

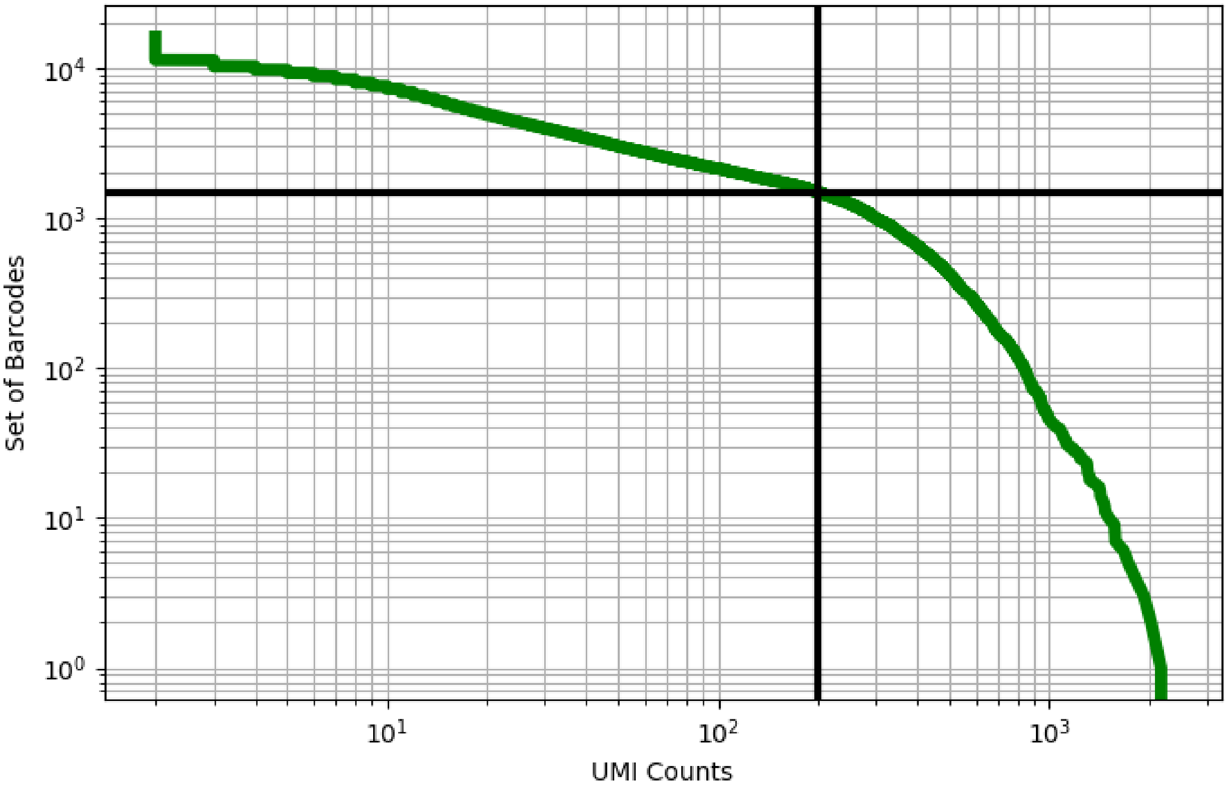


Figure 67: blood/GSM3576411

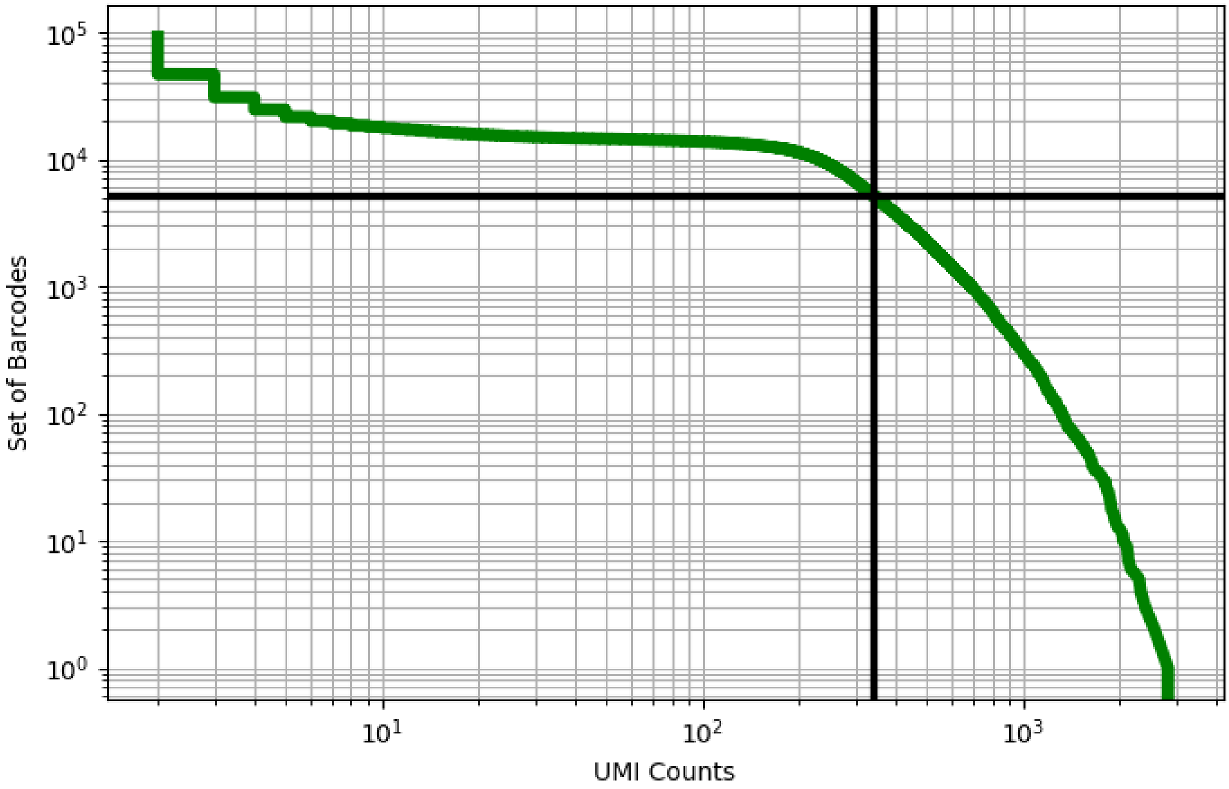


Figure 68: blood/GSM3576412

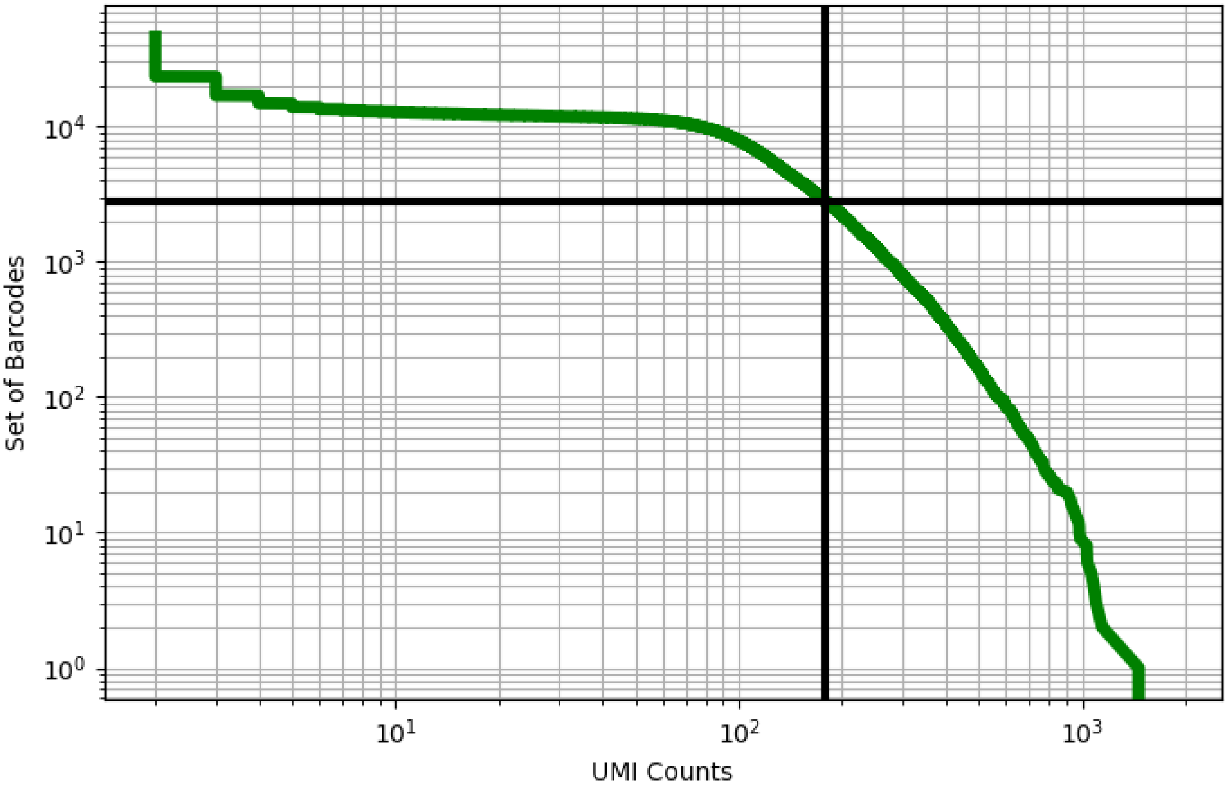


Figure 69: blood/GSM3576413

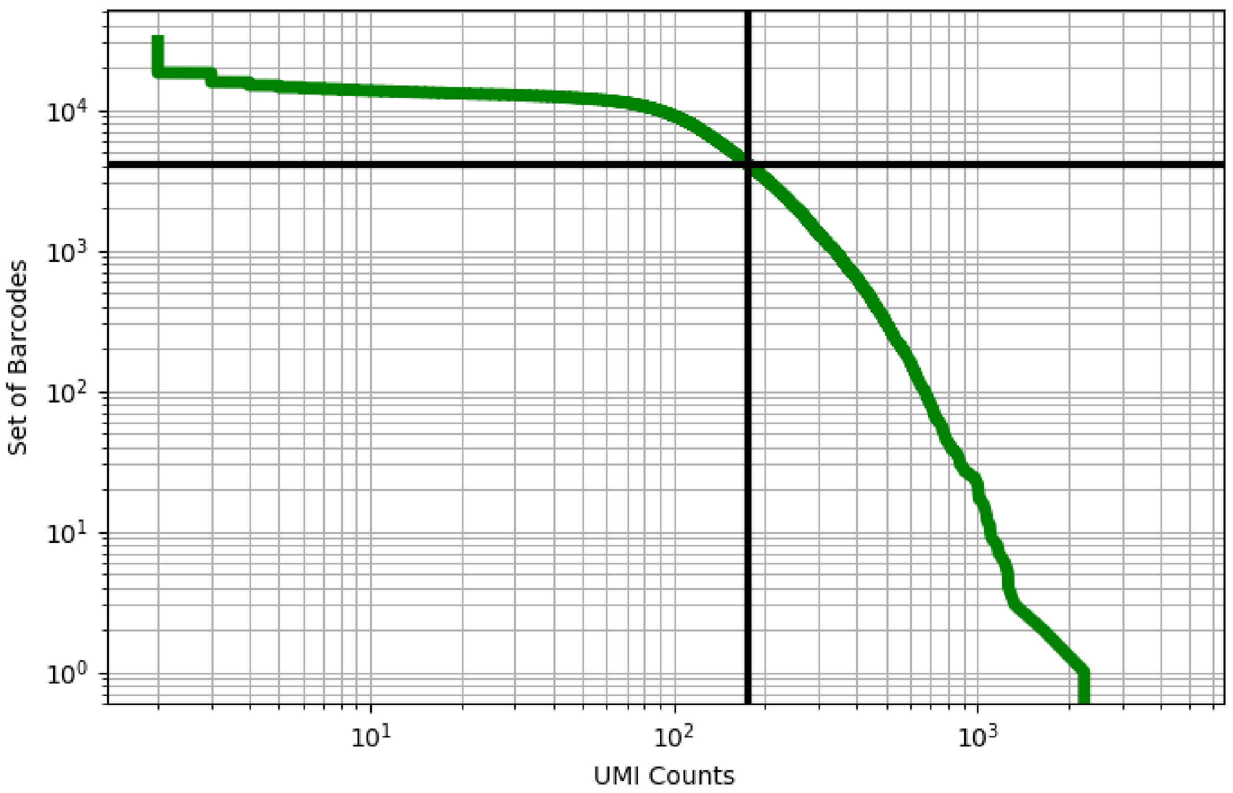


Figure 70: blood/GSM3576414

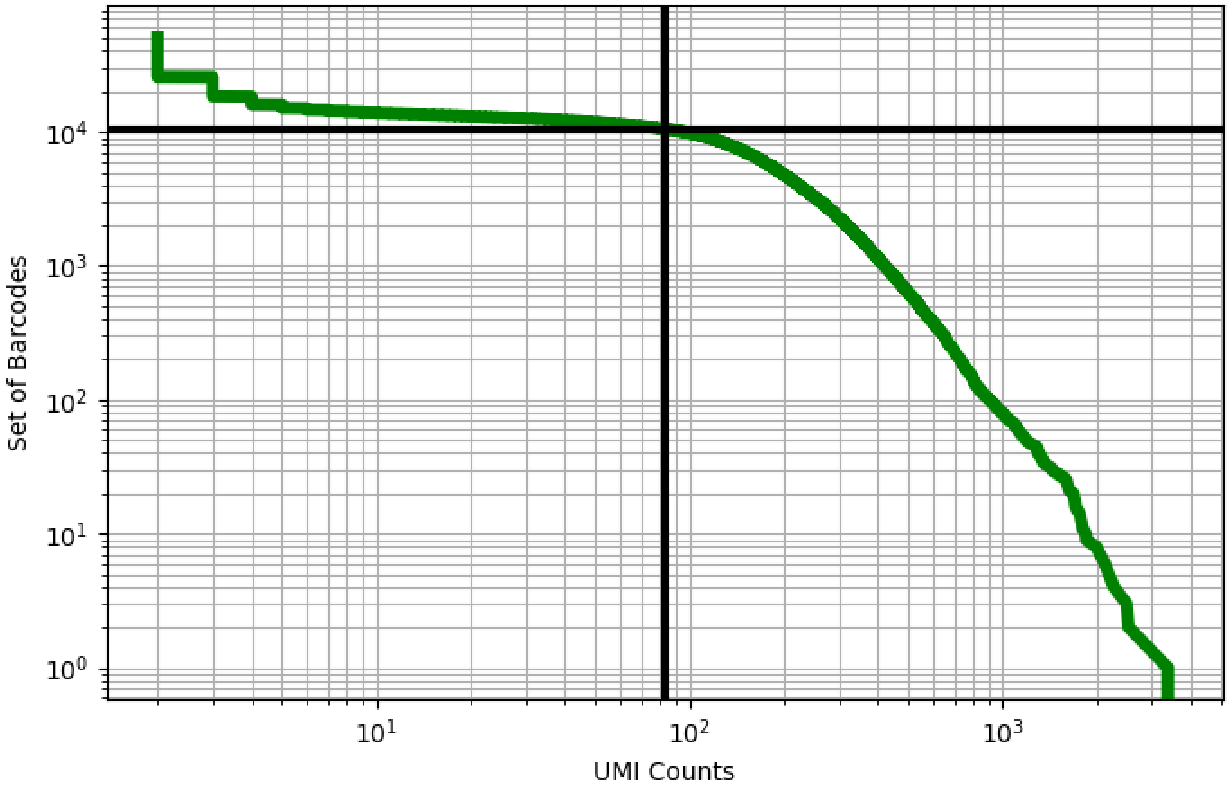


Figure 71: blood/GSM3576415

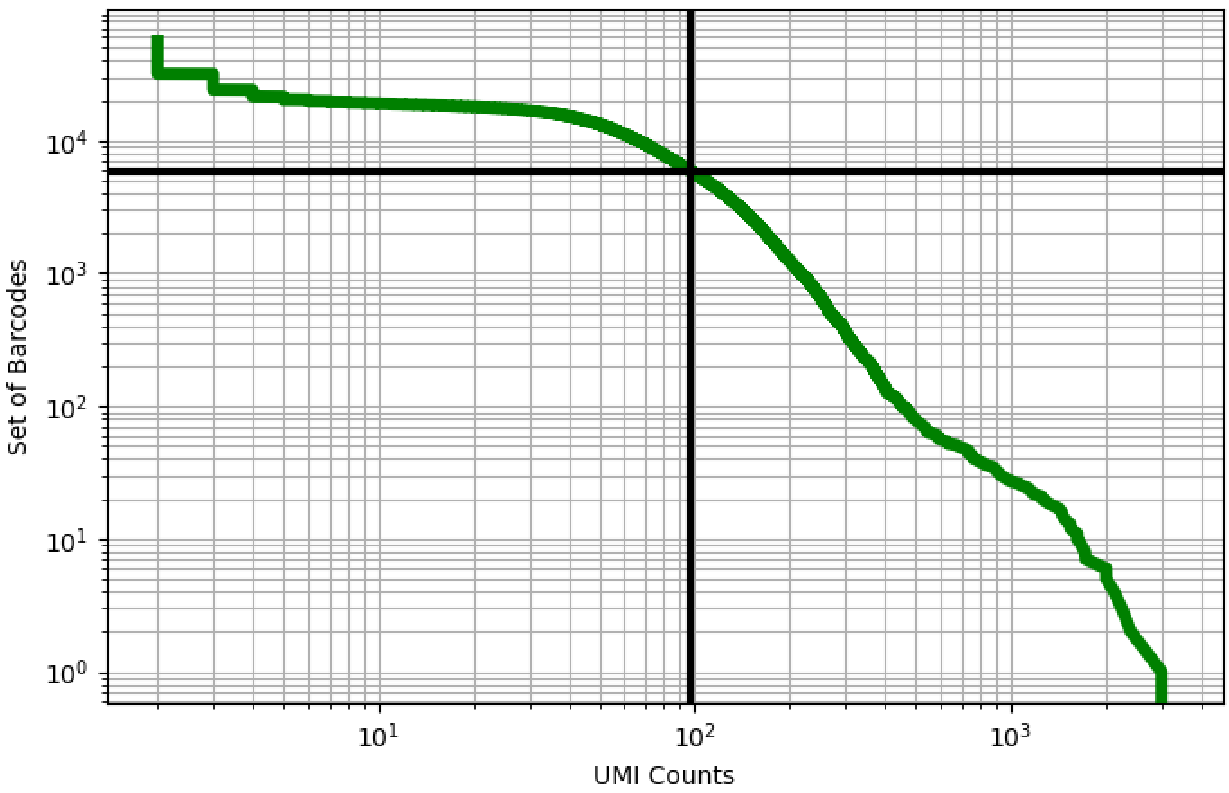


Figure 72: blood/GSM3576416

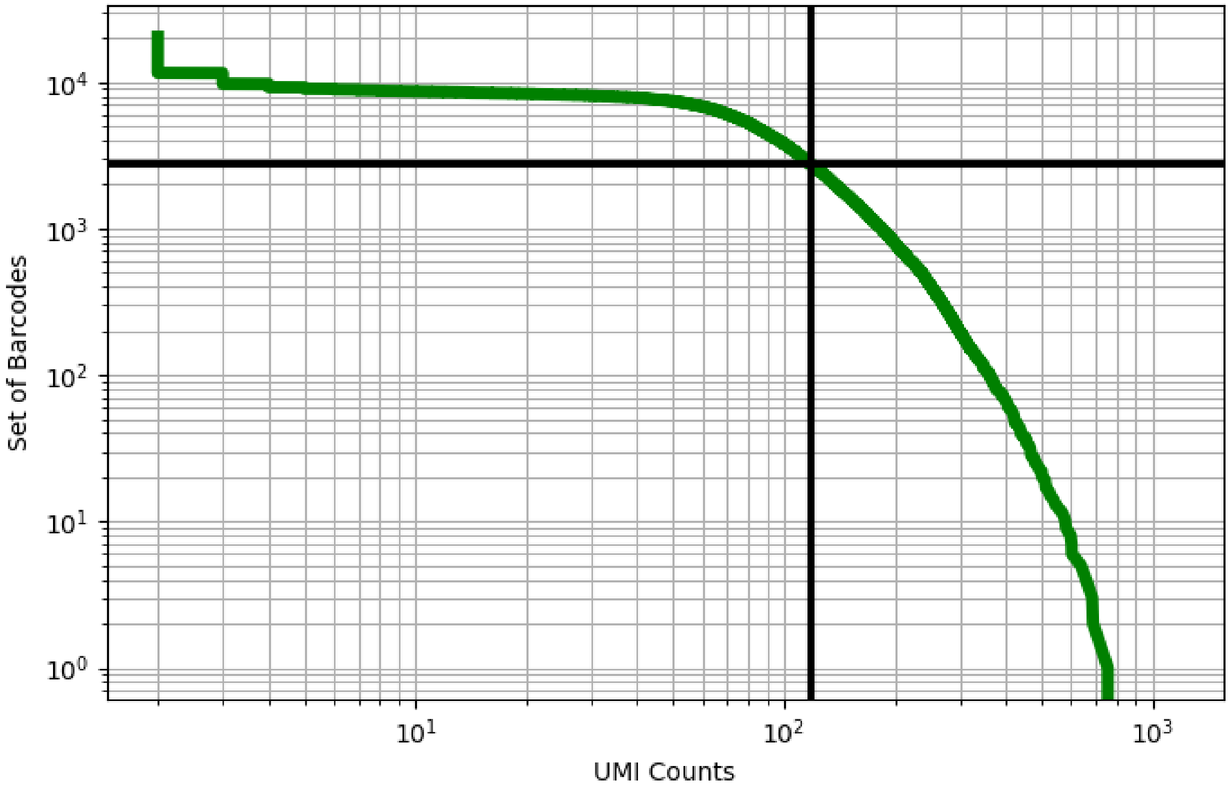


Figure 73: blood/GSM3576417

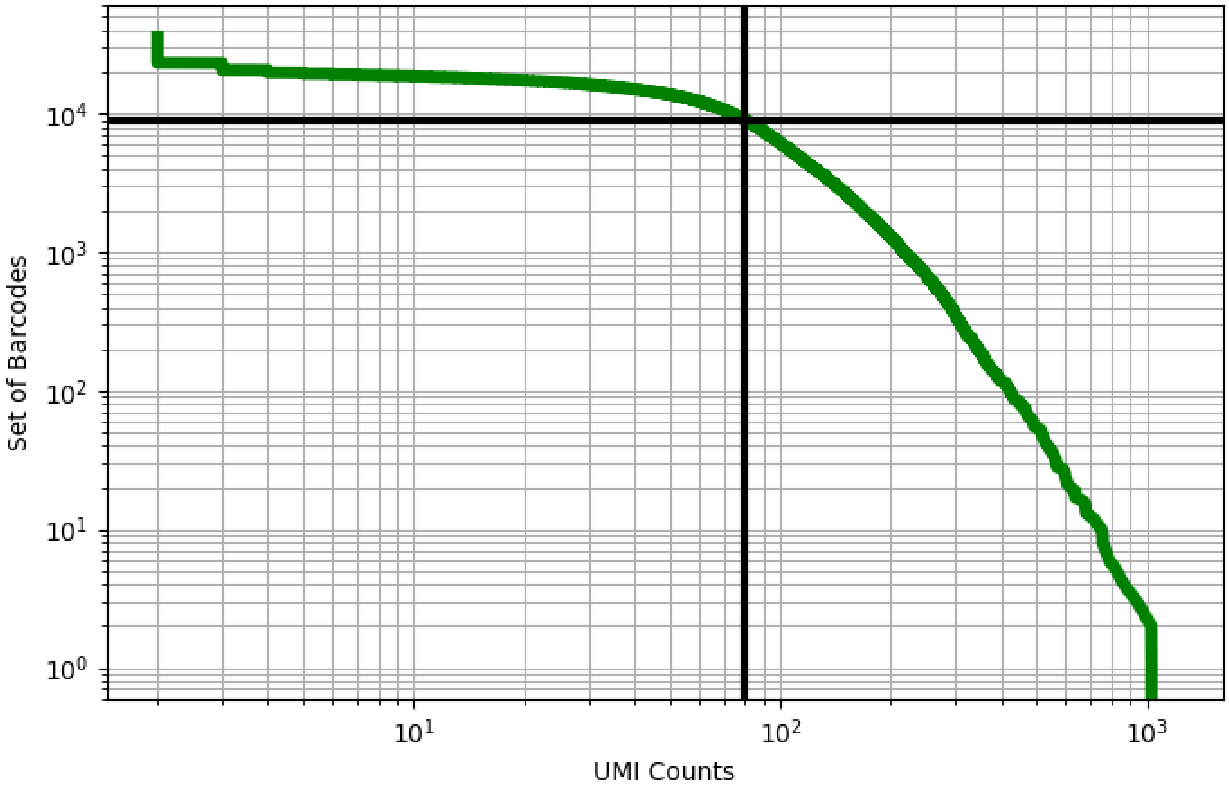


Figure 74: blood/GSM3576418

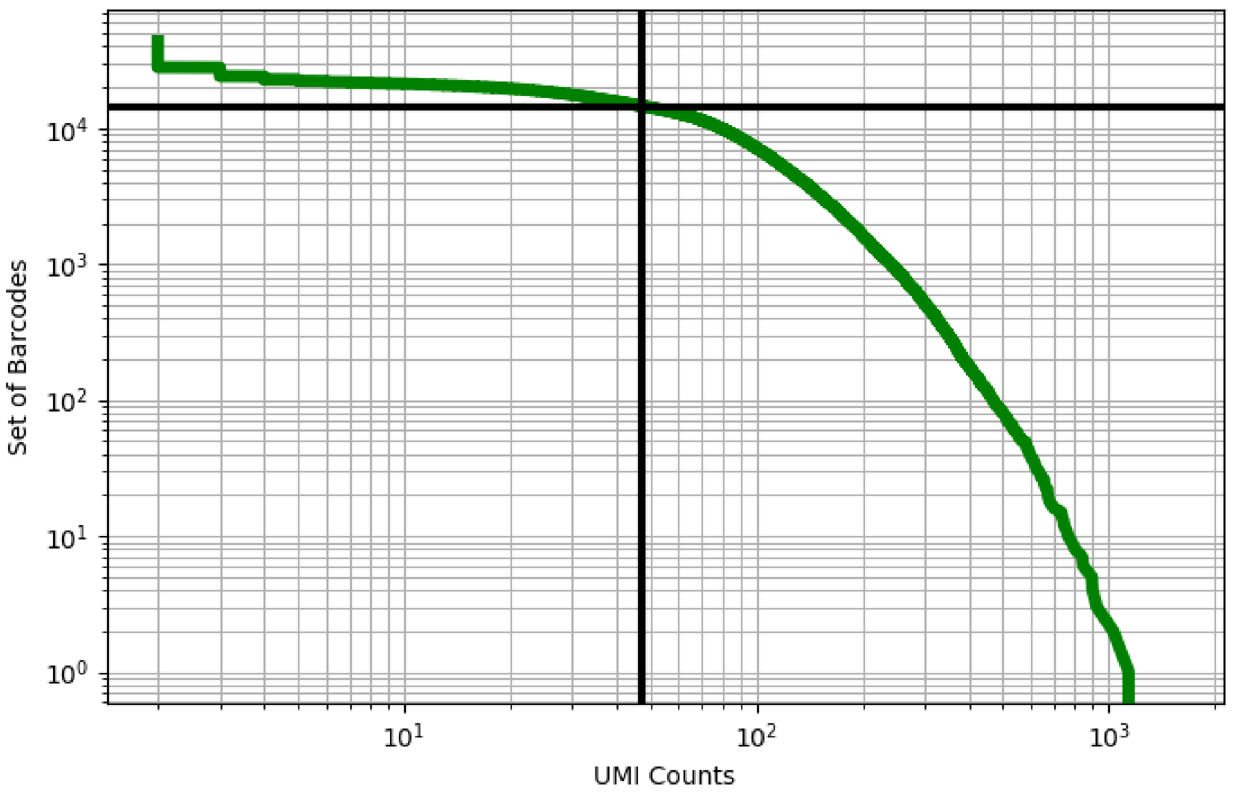


Figure 75: blood/GSM3576419

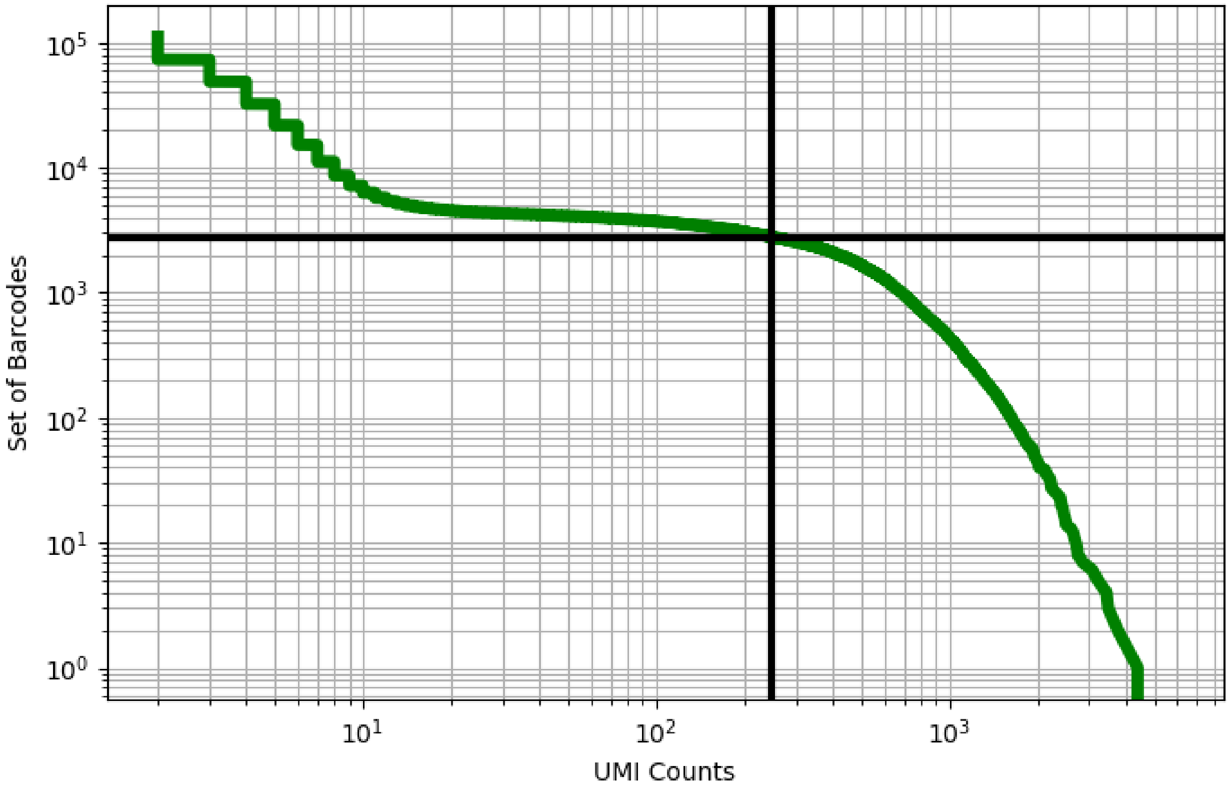


Figure 76: blood/GSM3576420

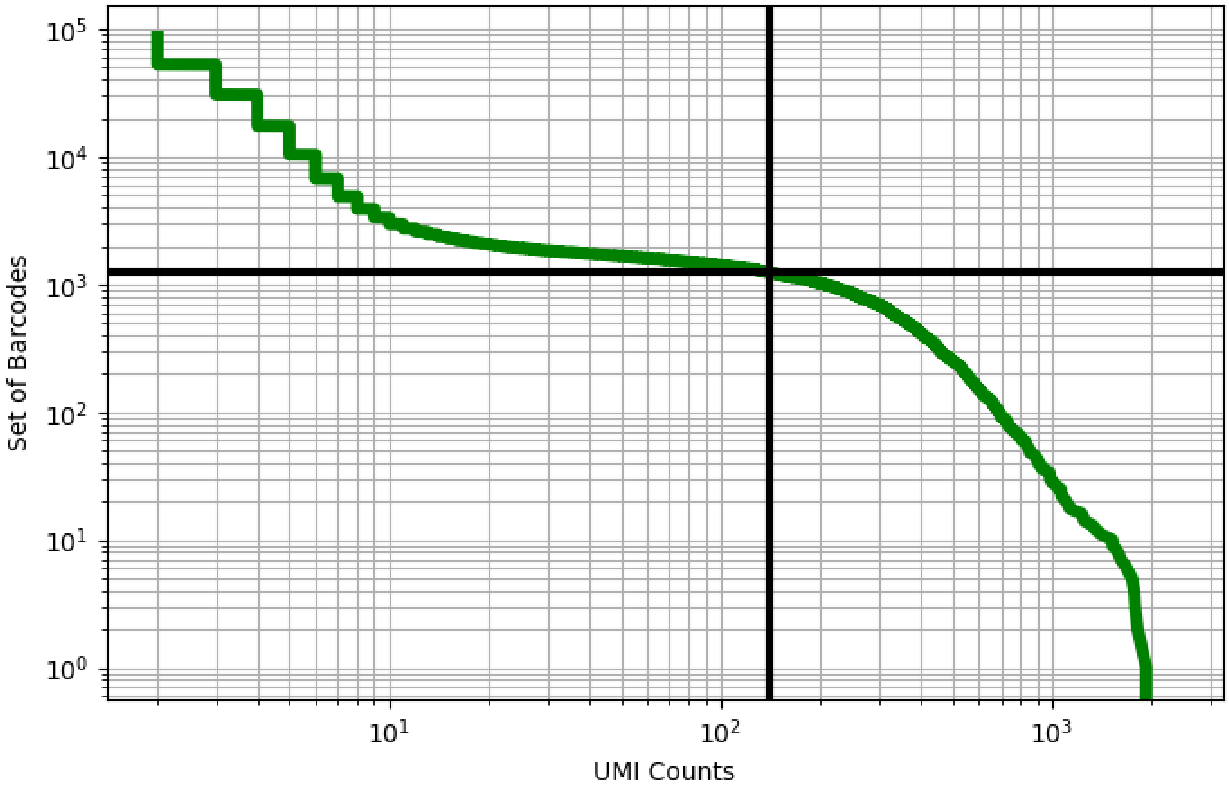


Figure 77: blood/GSM3576421

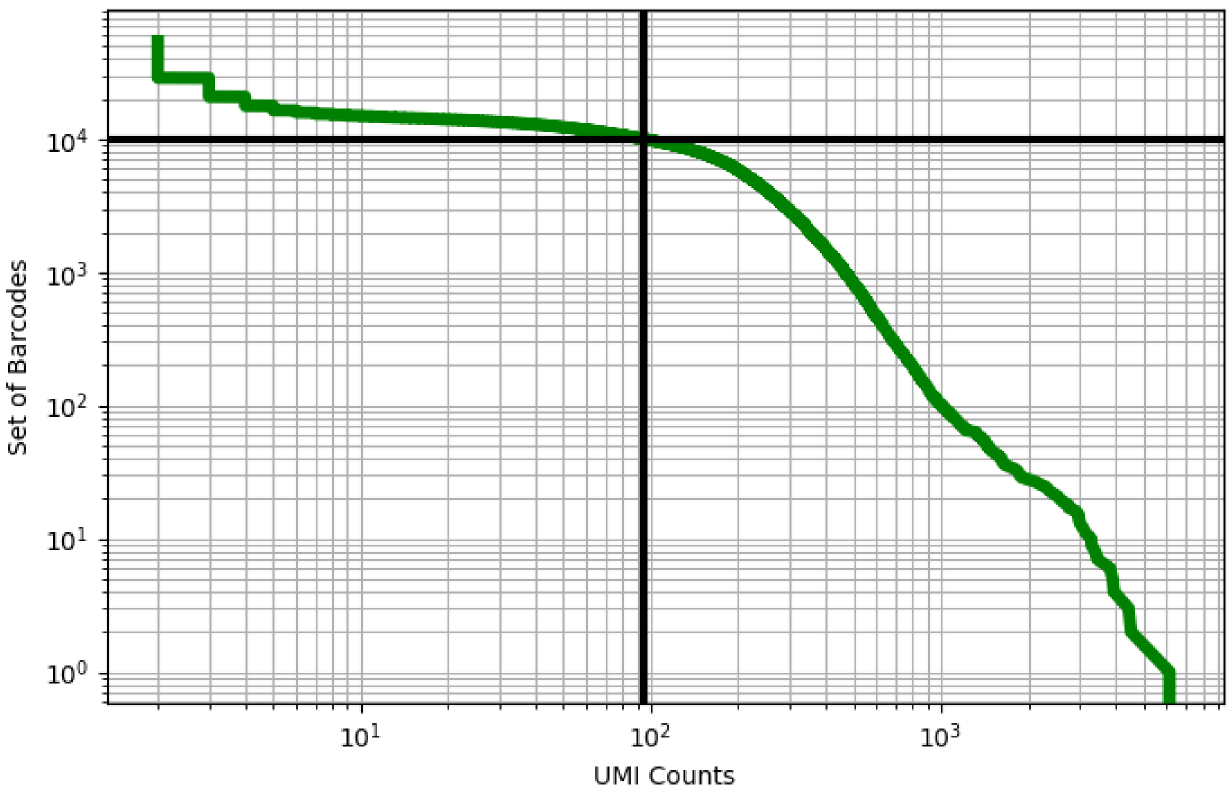


Figure 78: blood/GSM3576422

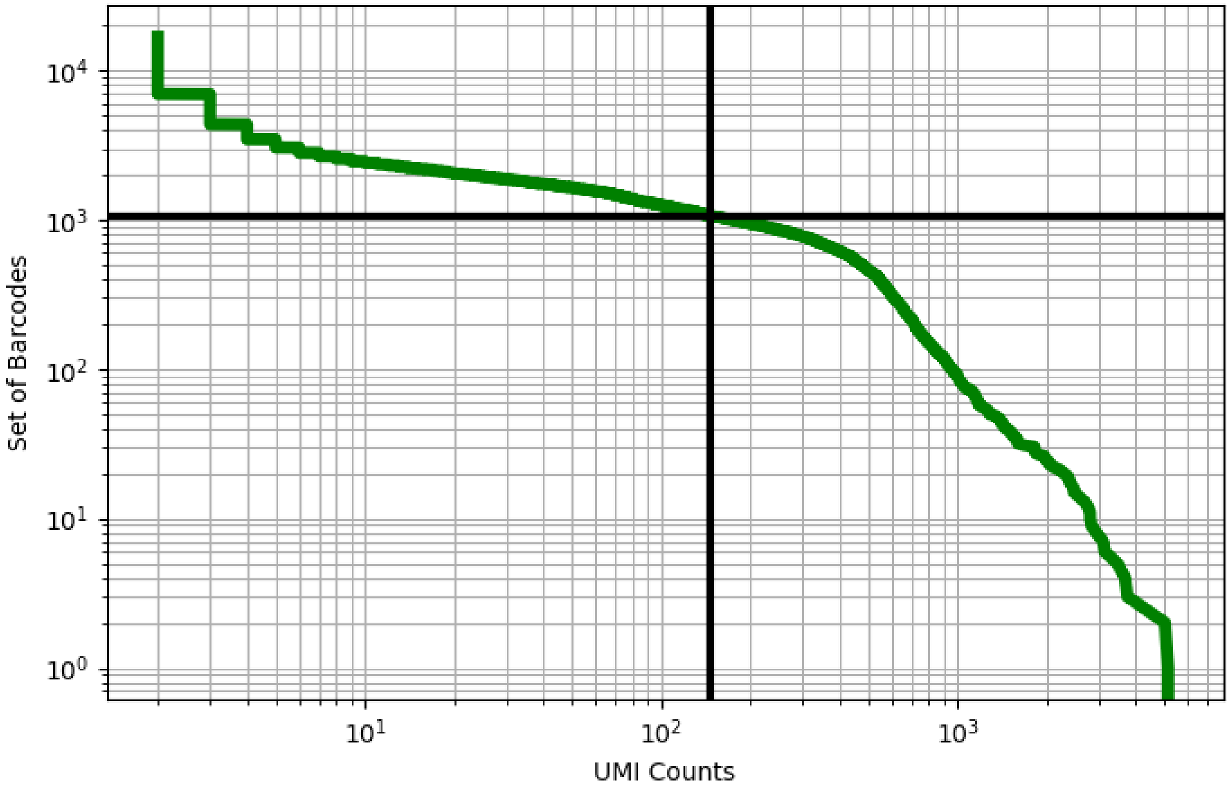


Figure 79: blood/GSM3576423

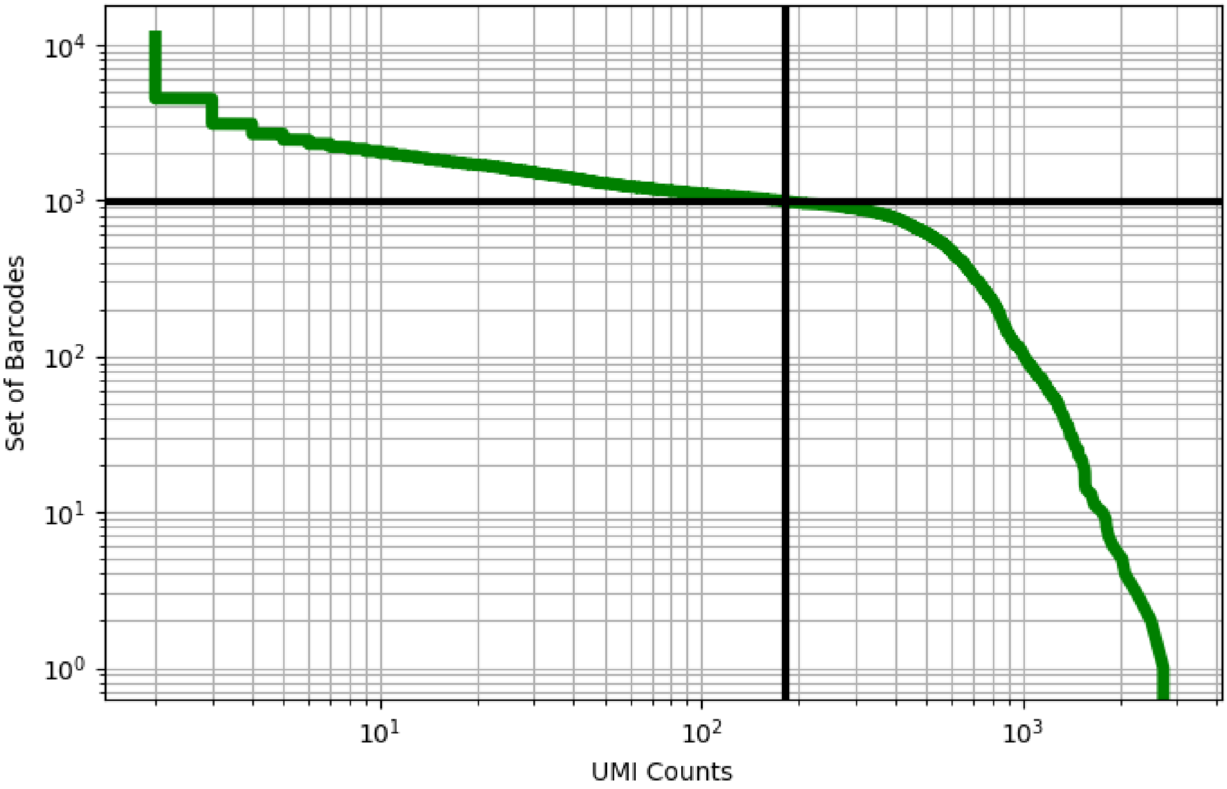


Figure 80: blood/GSM3576424

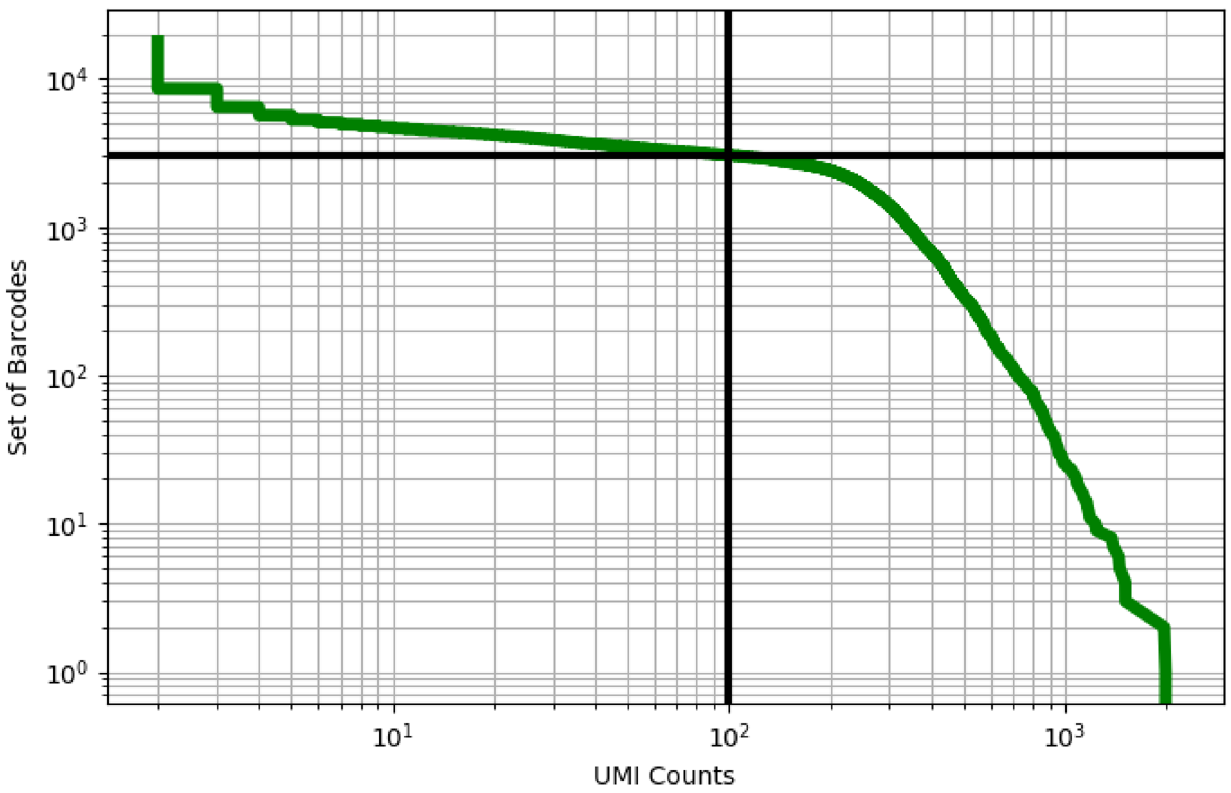


Figure 81: blood/GSM3576425

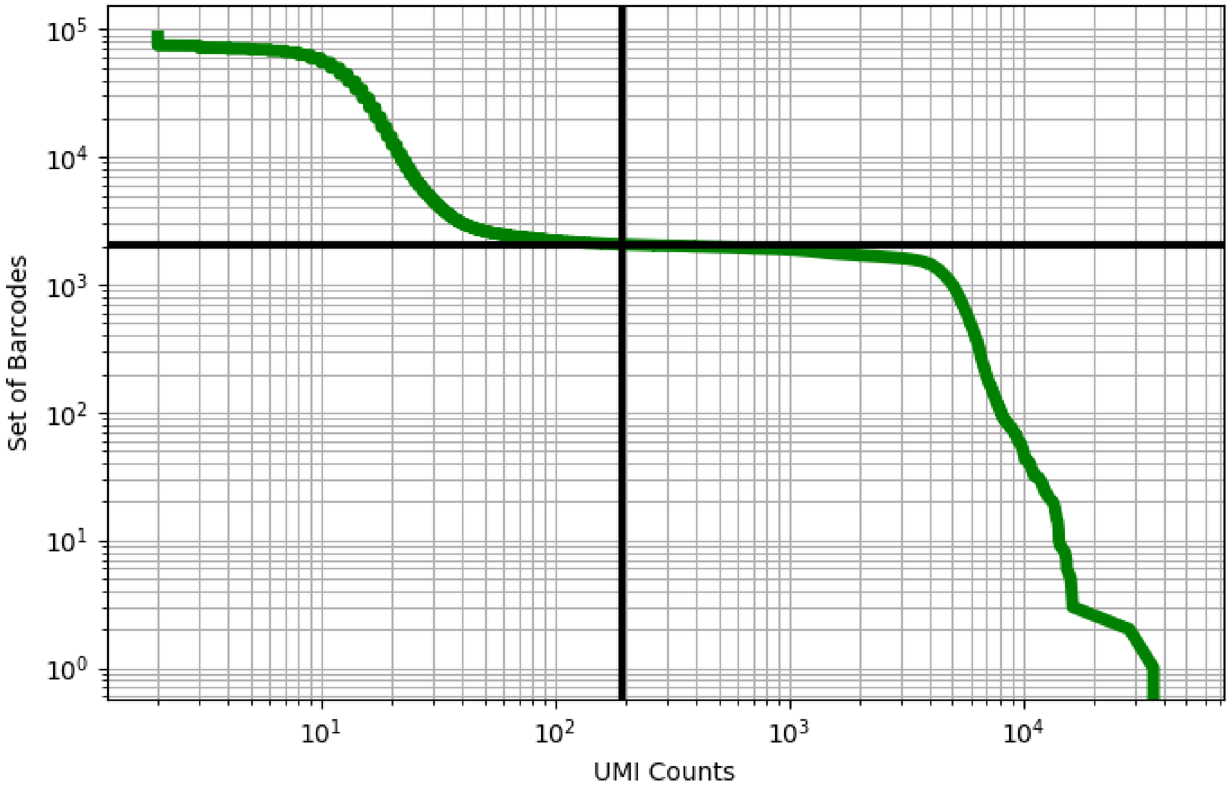


Figure 82: blood/GSM3665016

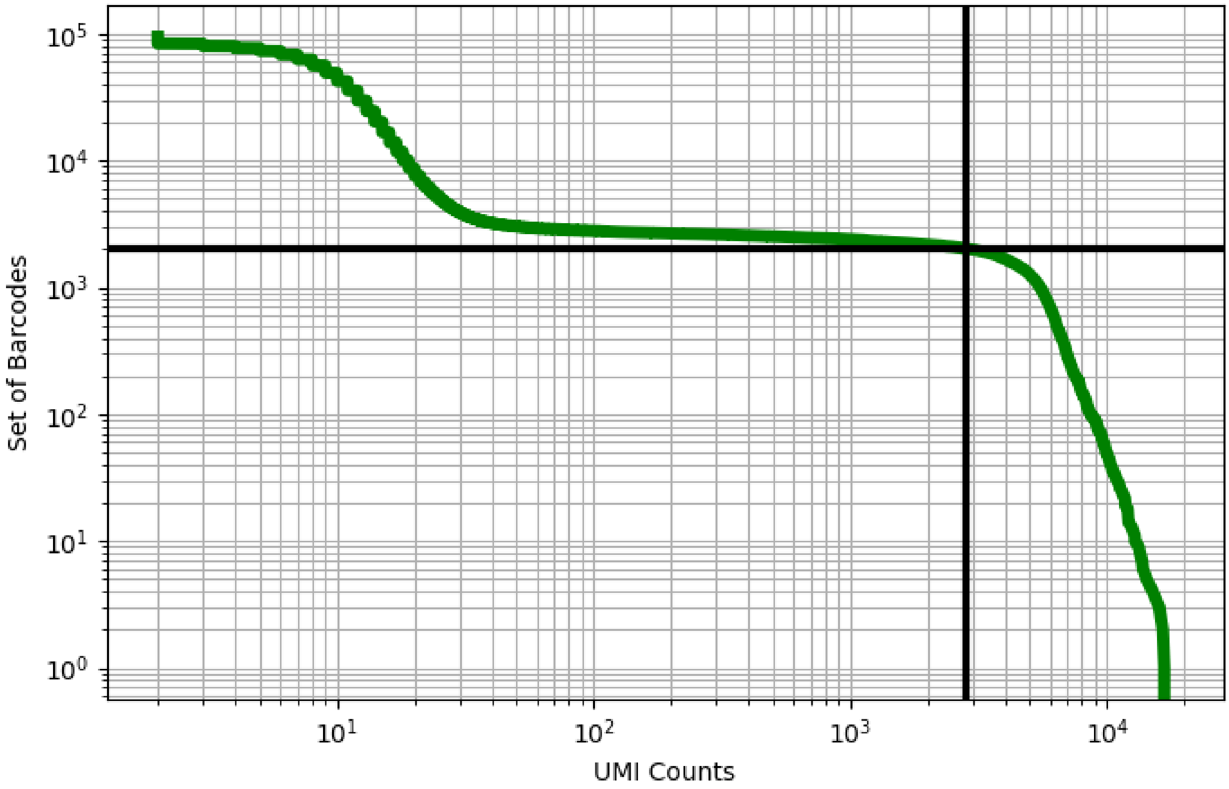


Figure 83: blood/GSM3665017

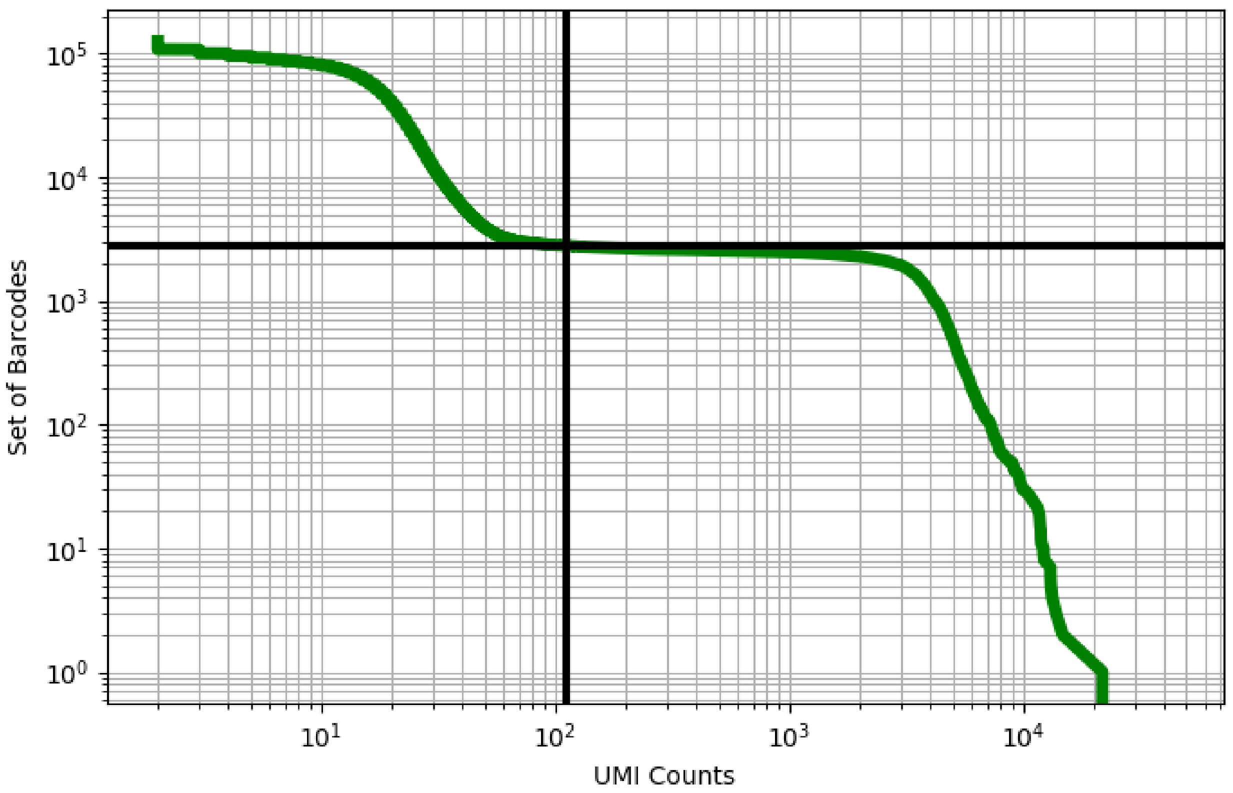


Figure 84: blood/GSM3665018

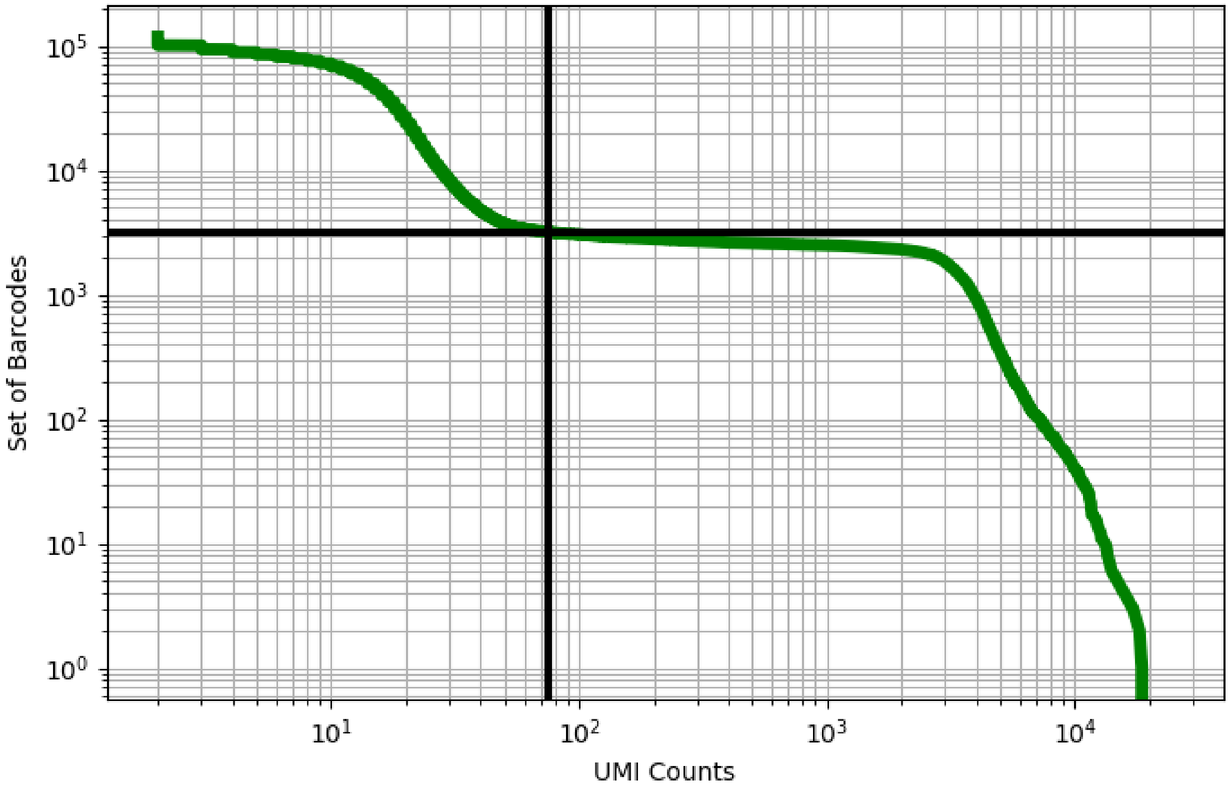


Figure 85: blood/GSM3665019

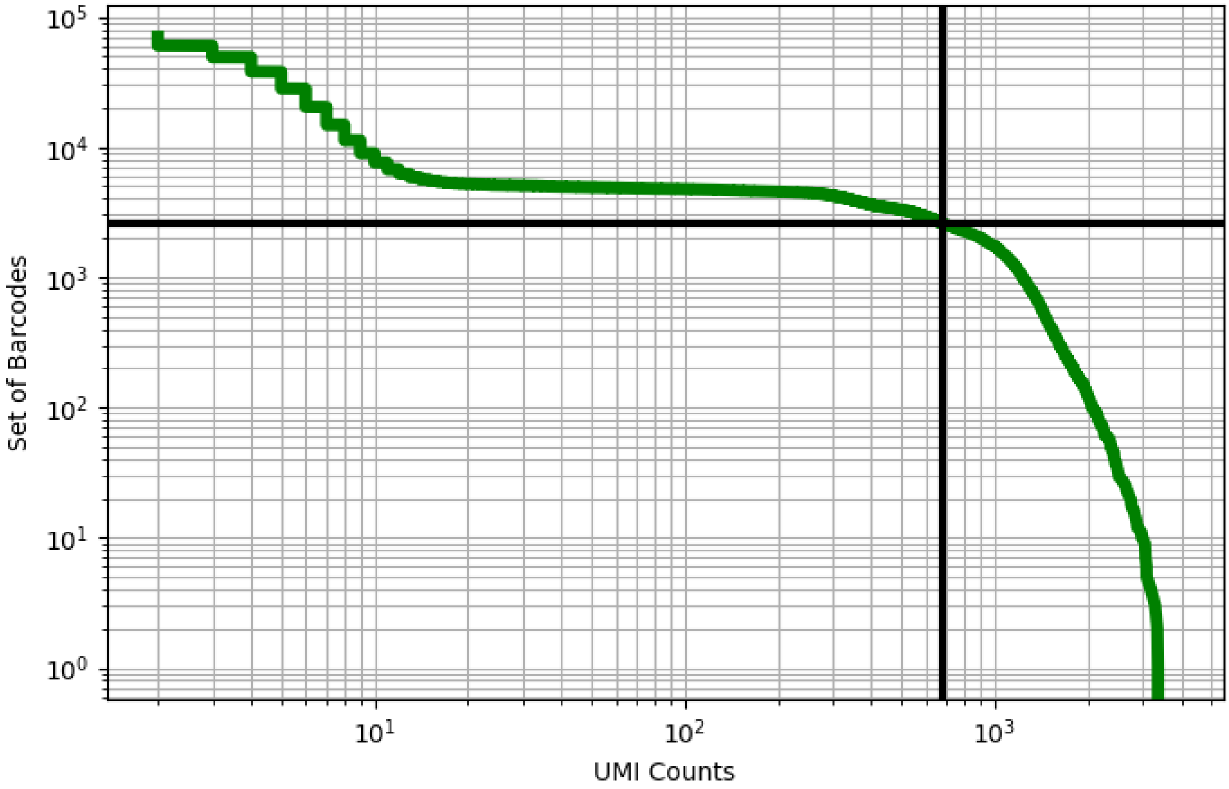


Figure 86: blood/GSM3732362

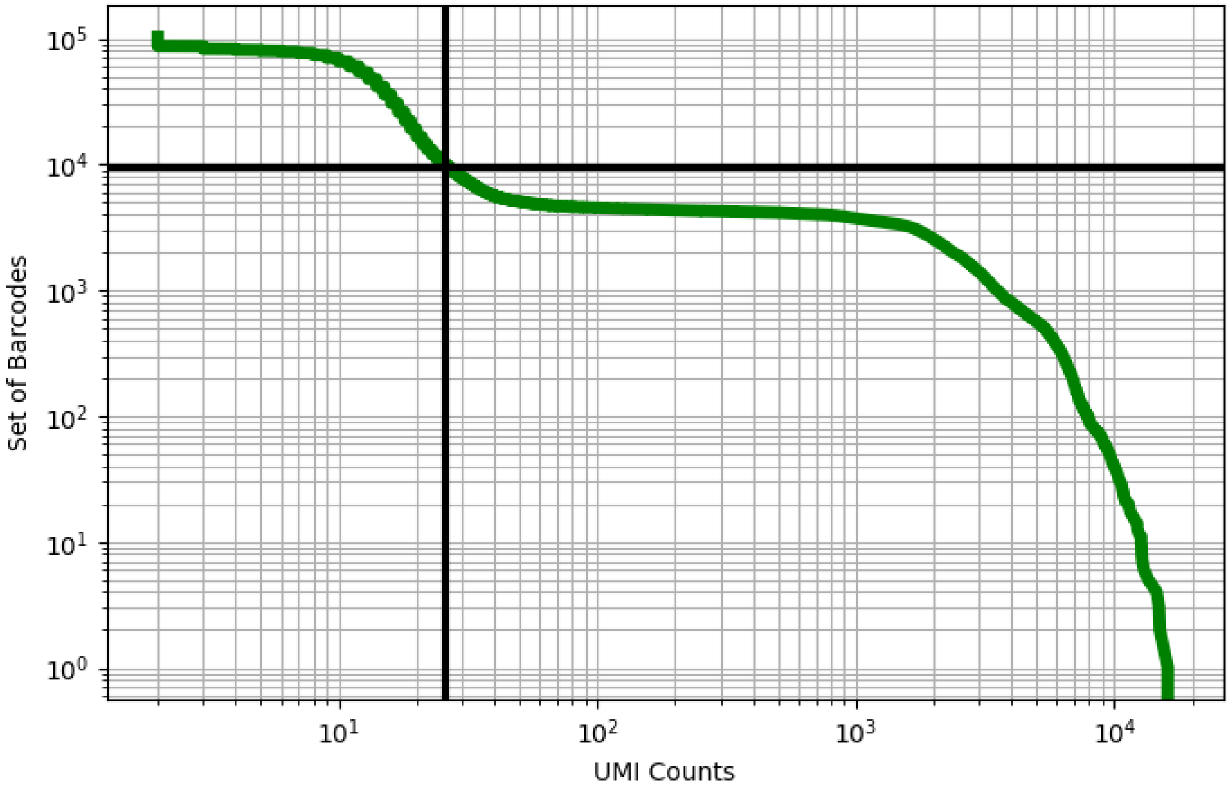


Figure 87: blood/GSM3732364

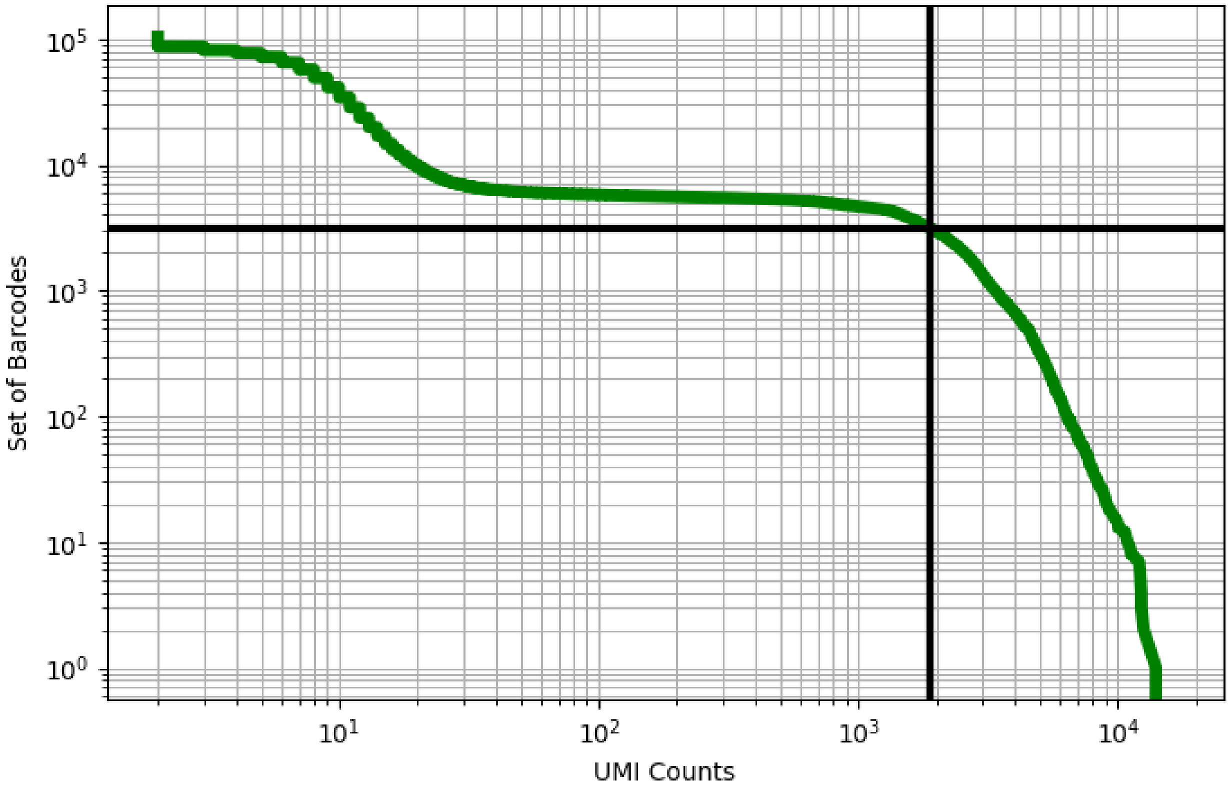


Figure 88: blood/GSM3732366

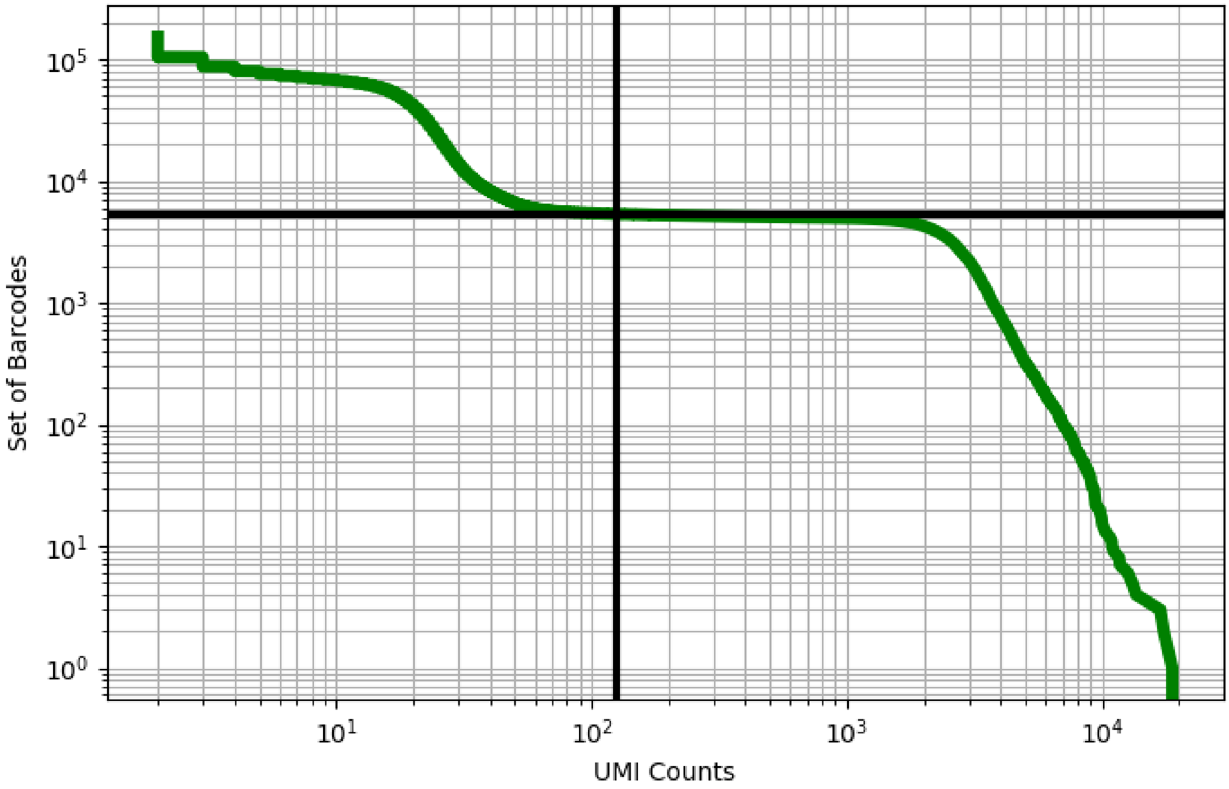


Figure 89: blood/GSM3734178

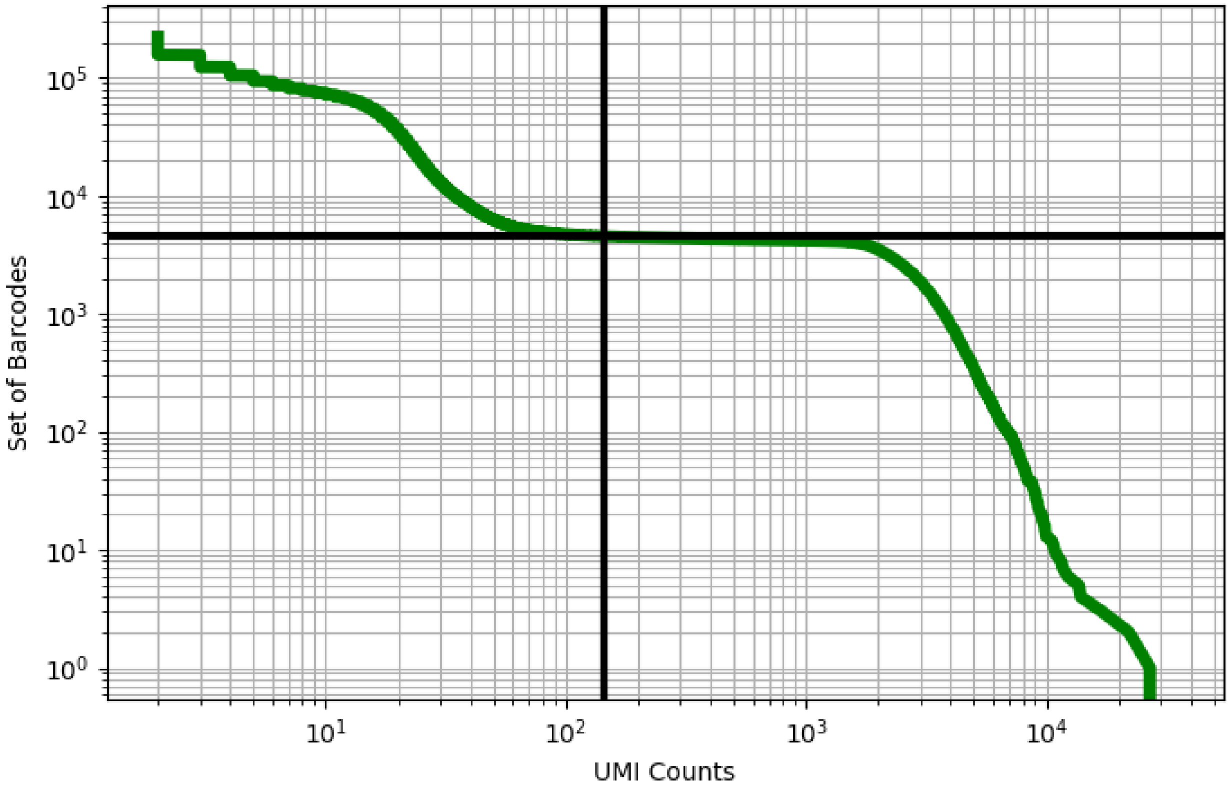


Figure 90: blood/GSM3734179

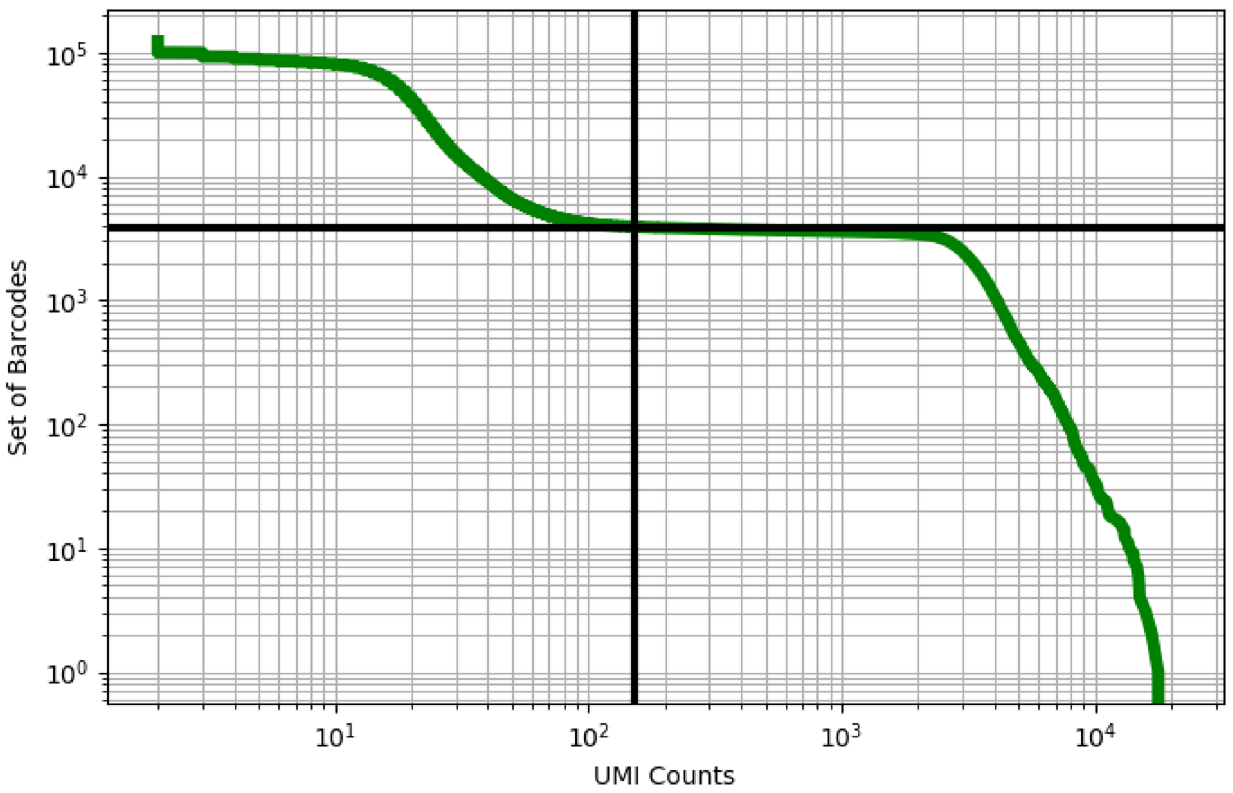


Figure 91: blood/GSM3734180

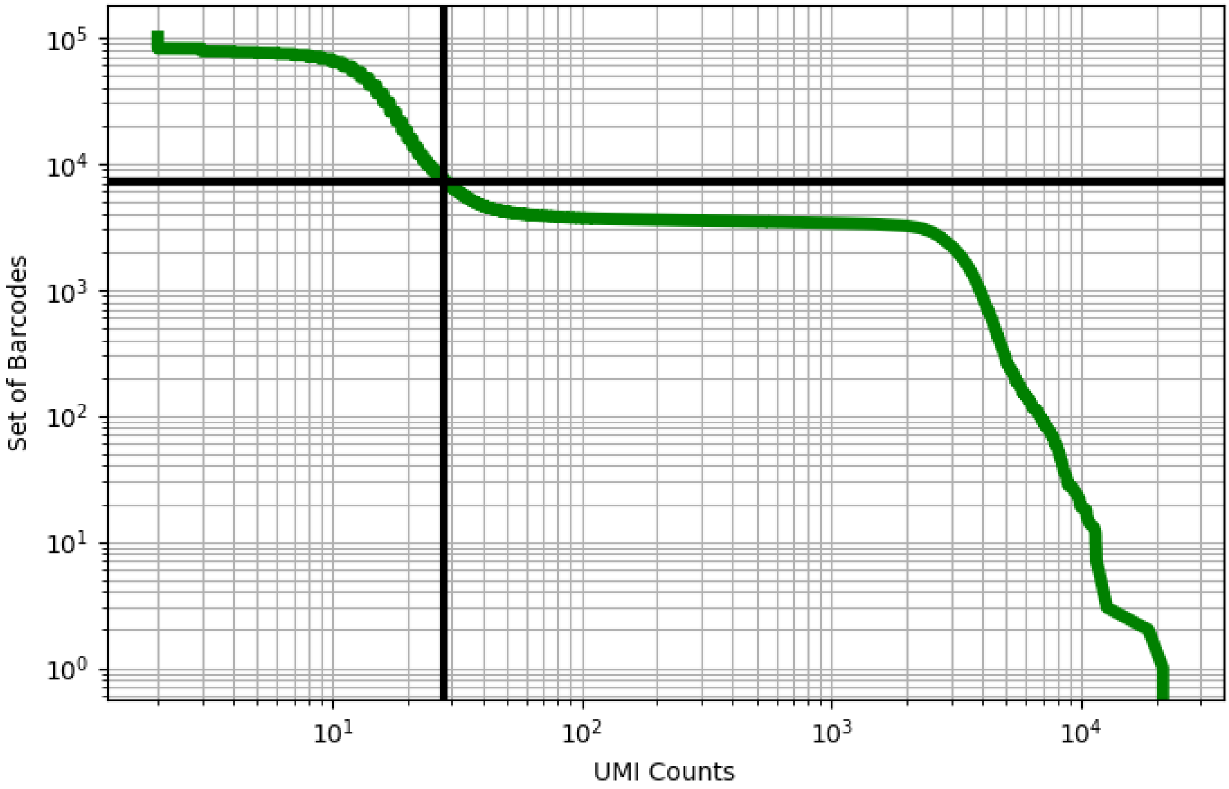


Figure 92: blood/GSM3734181

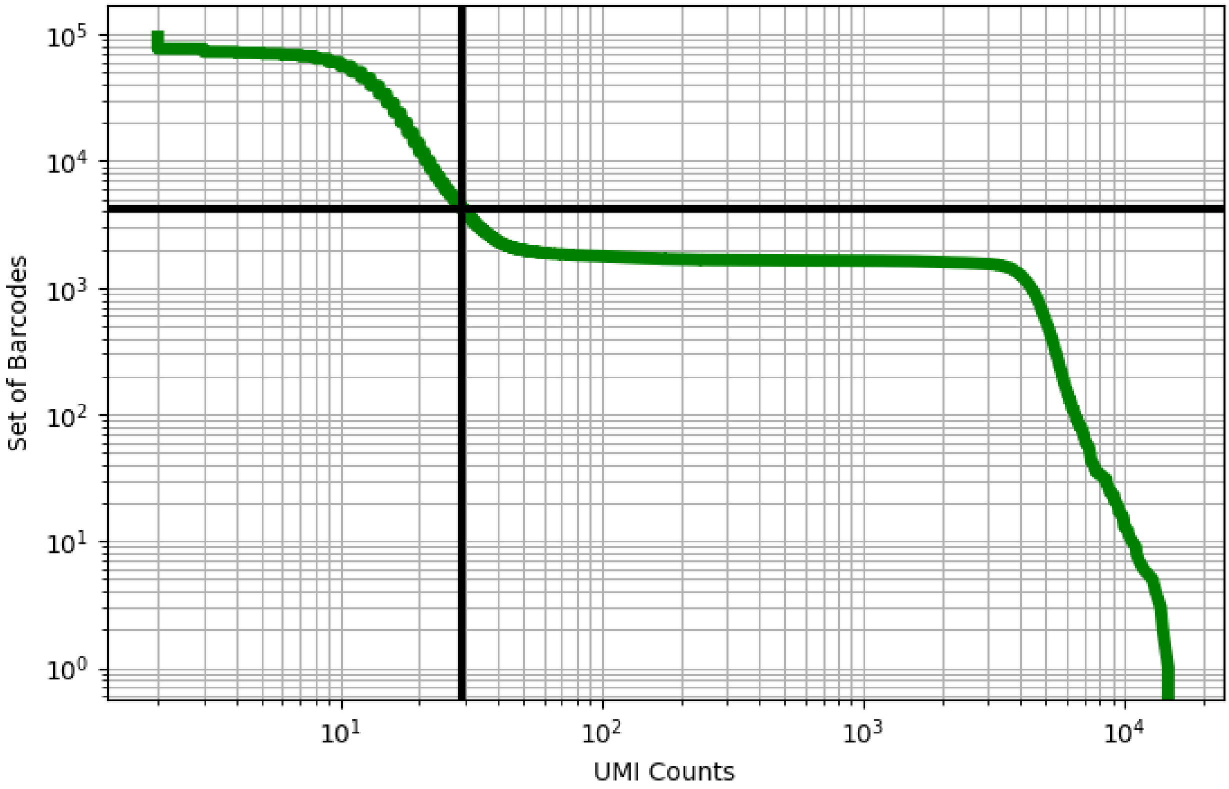


Figure 93: blood/GSM3738542

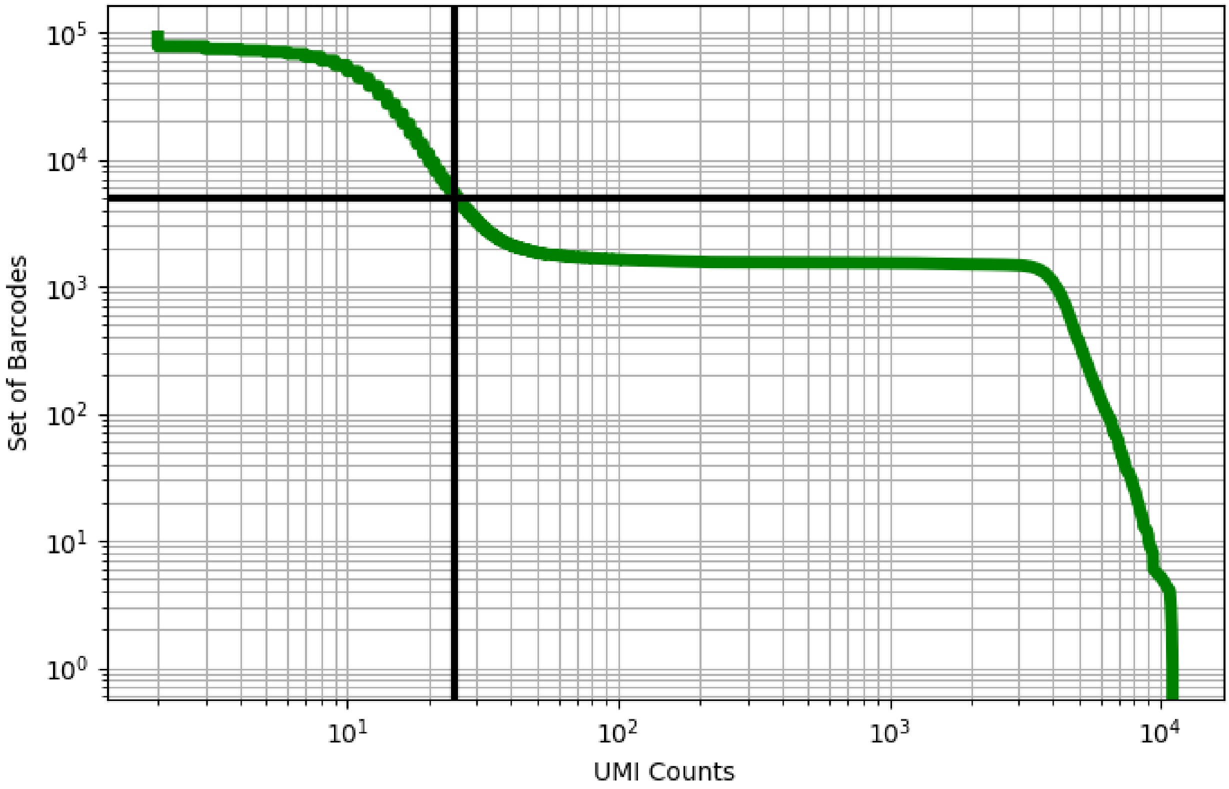


Figure 94: blood/GSM3738543

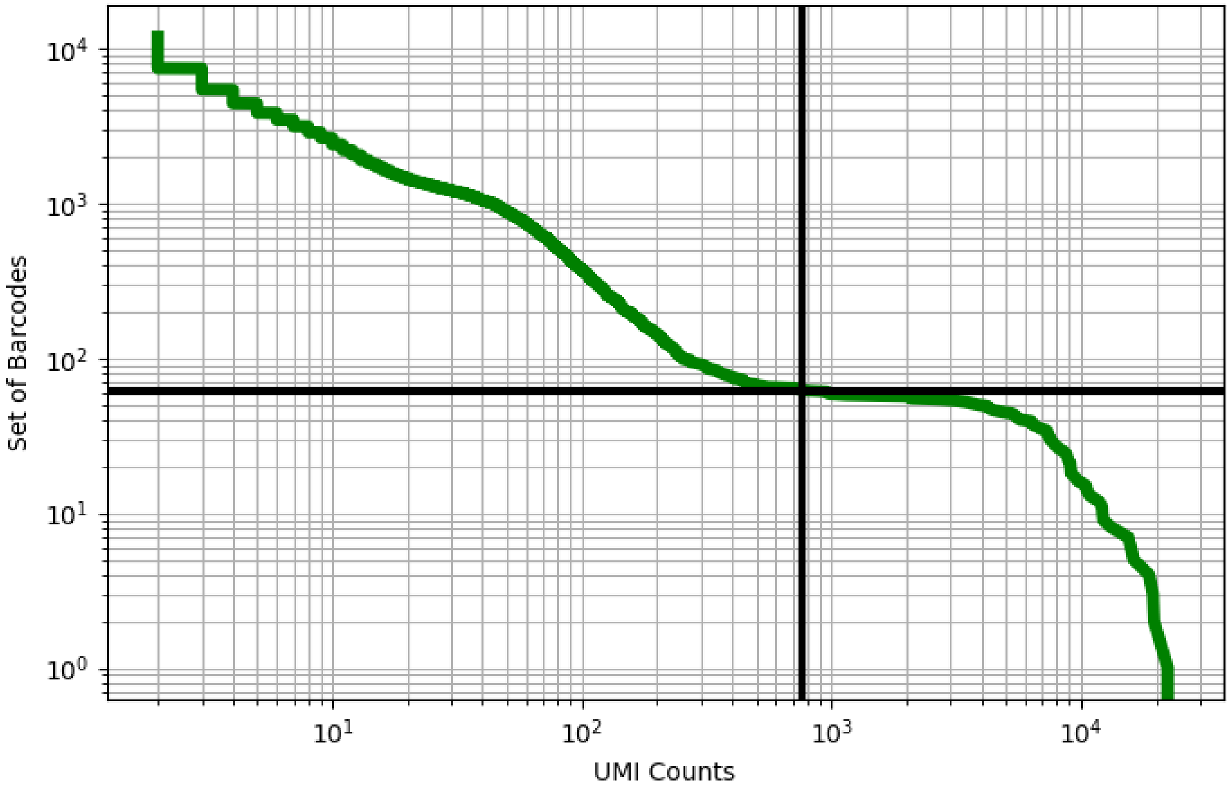


Figure 95: blood/GSM3739219

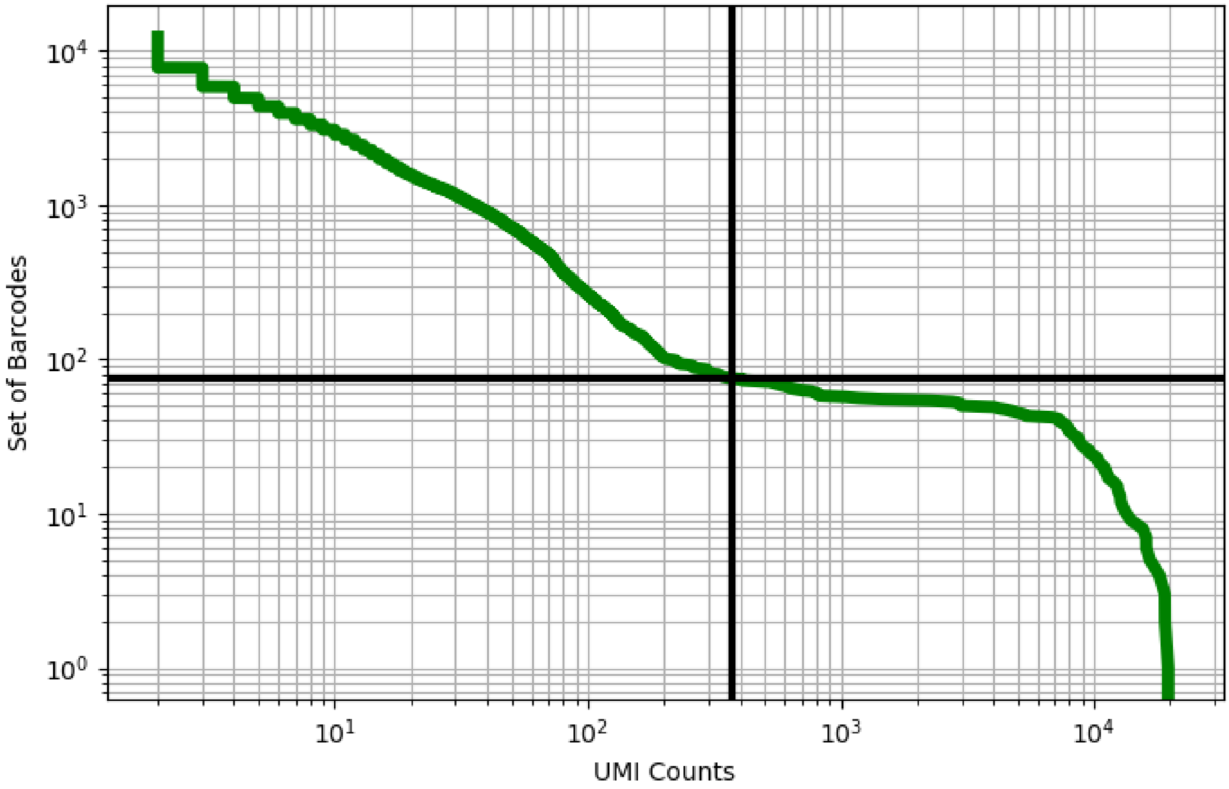


Figure 96: blood/GSM3739220

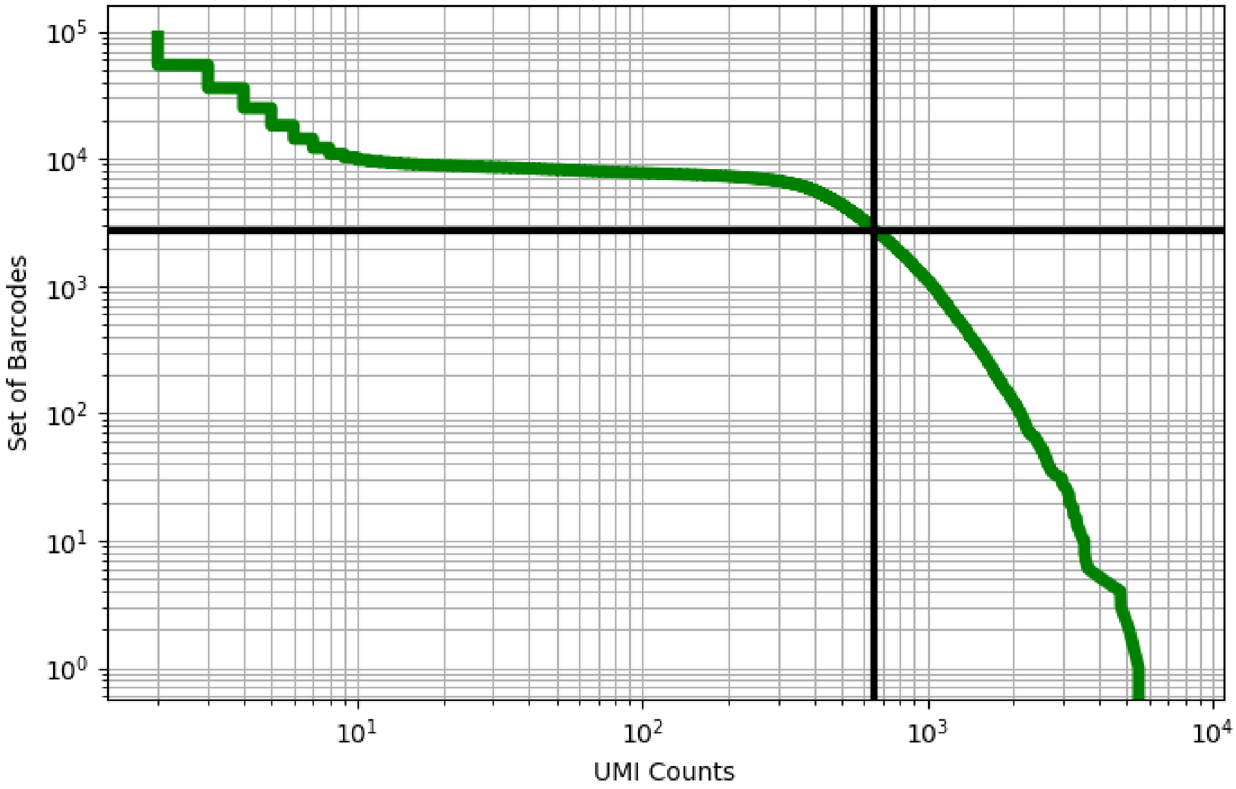


Figure 97: blood/SAMN14275543

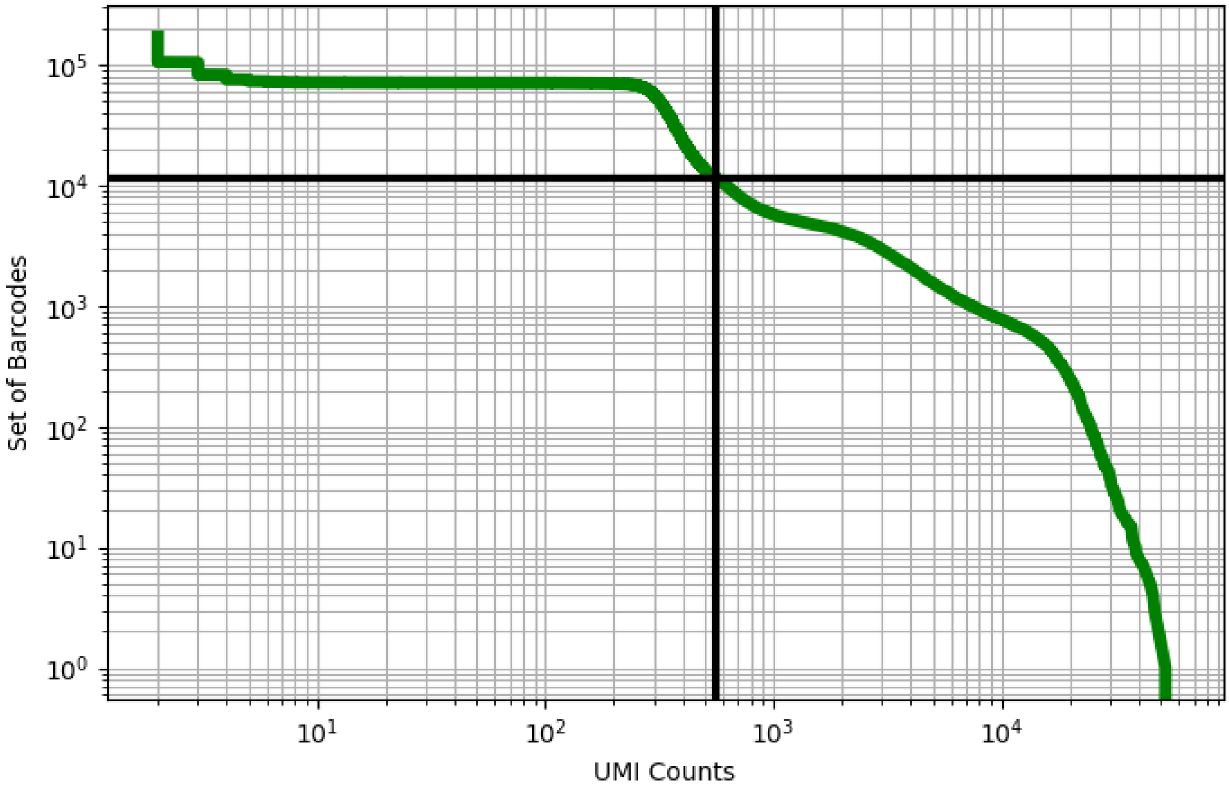


Figure 98: bone_marrow/GSM3168248

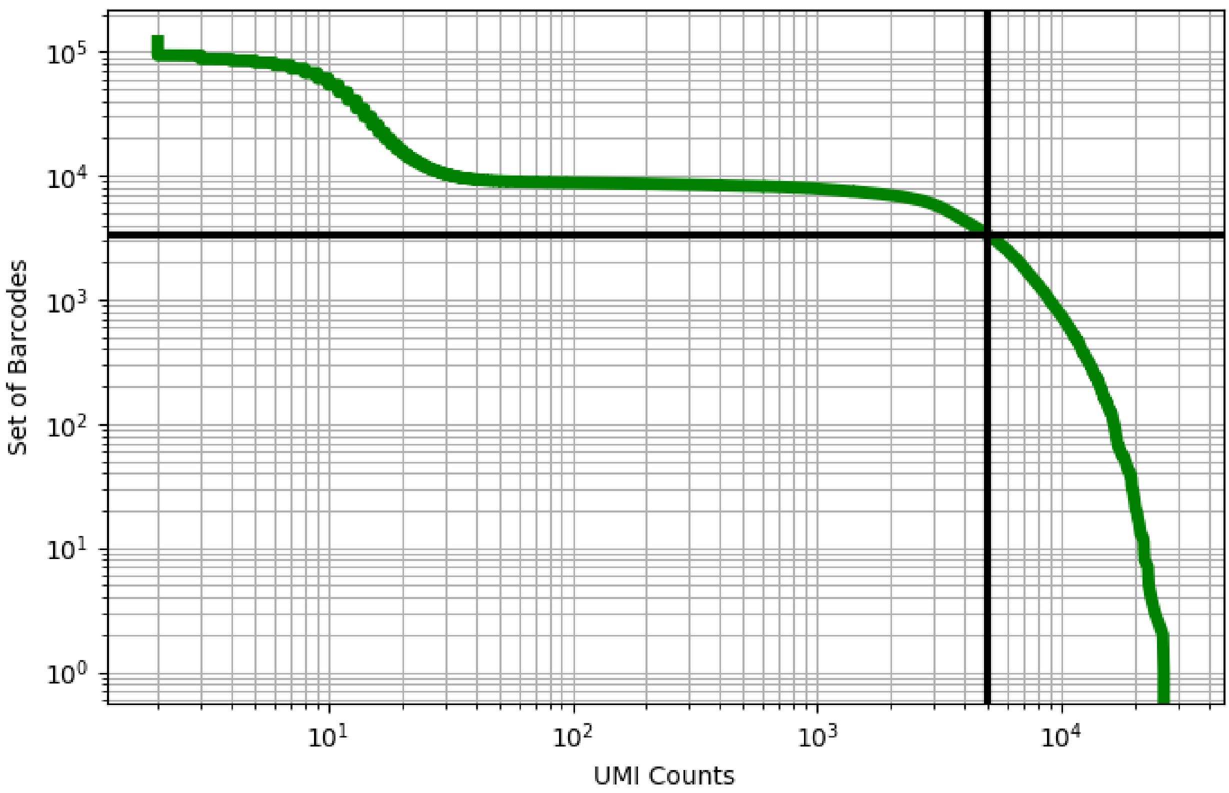


Figure 99: bone_marrow/GSM3309832

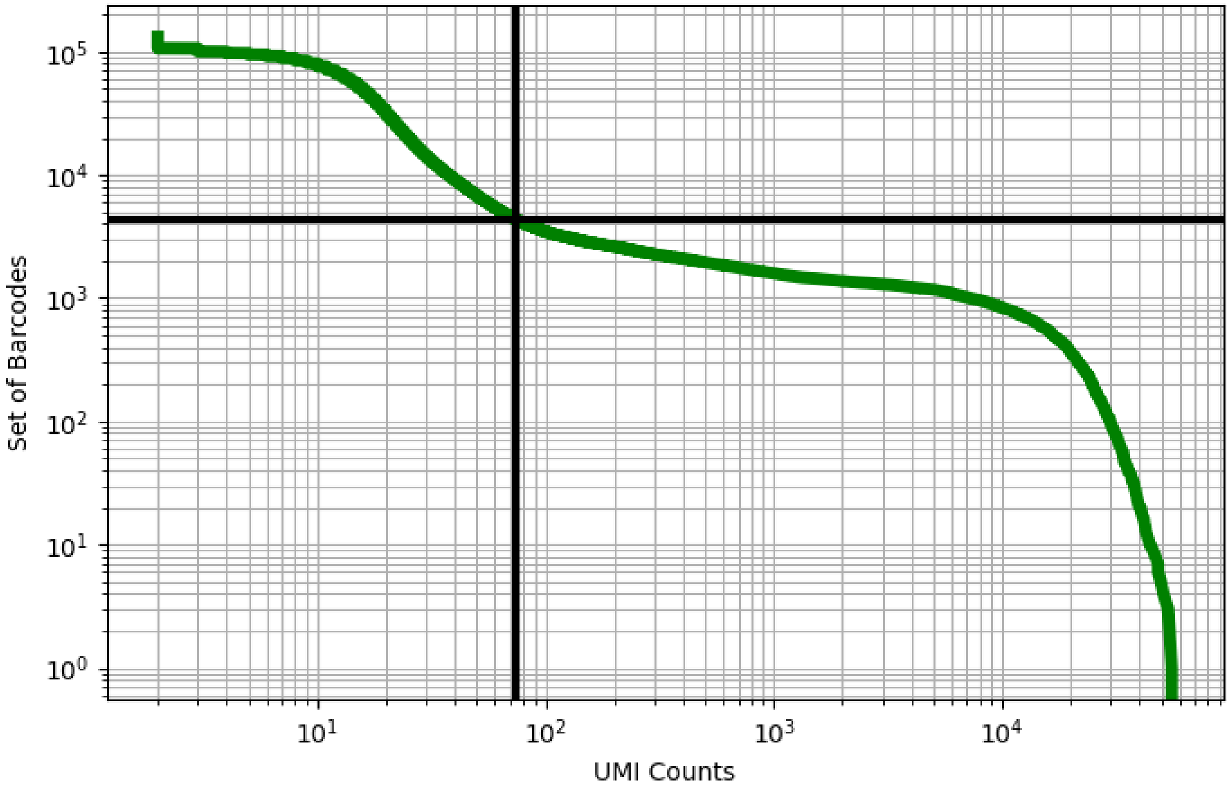


Figure 100: bone_marrow/GSM3309833

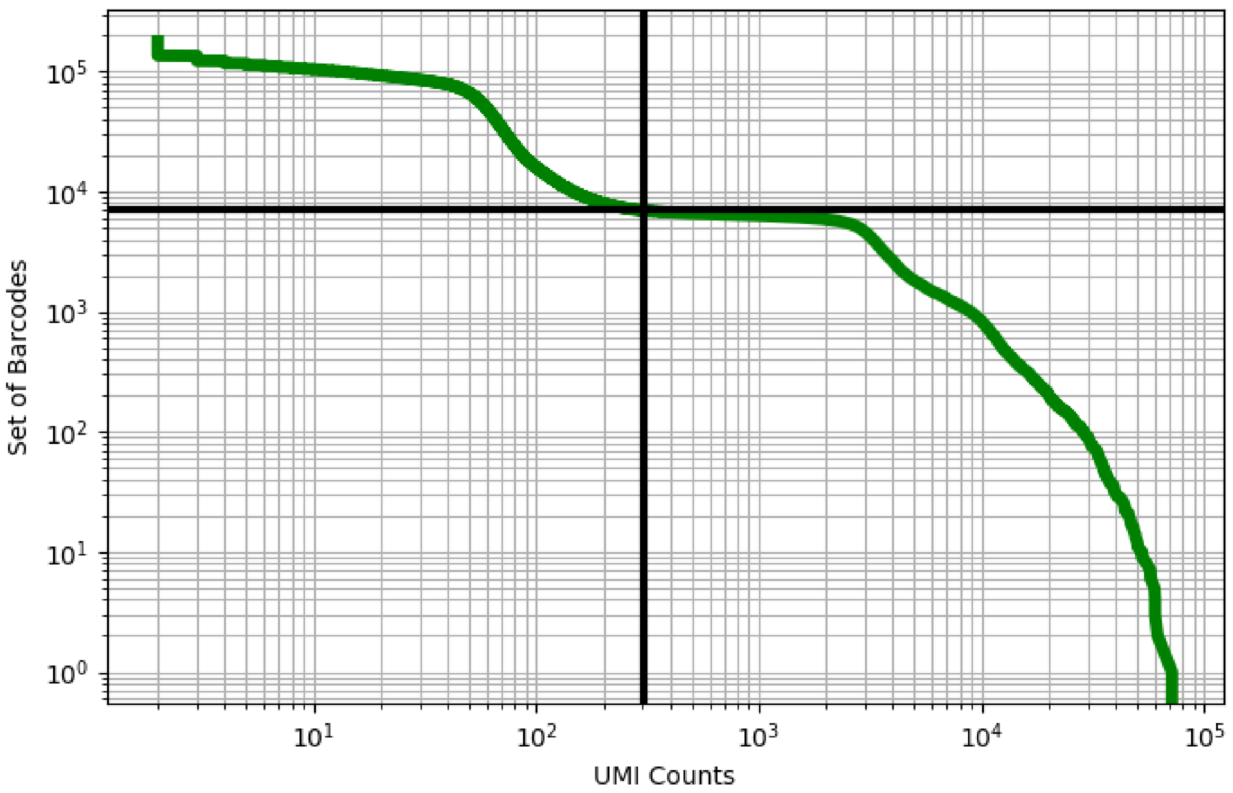


Figure 101: bone_marrow/GSM3396161

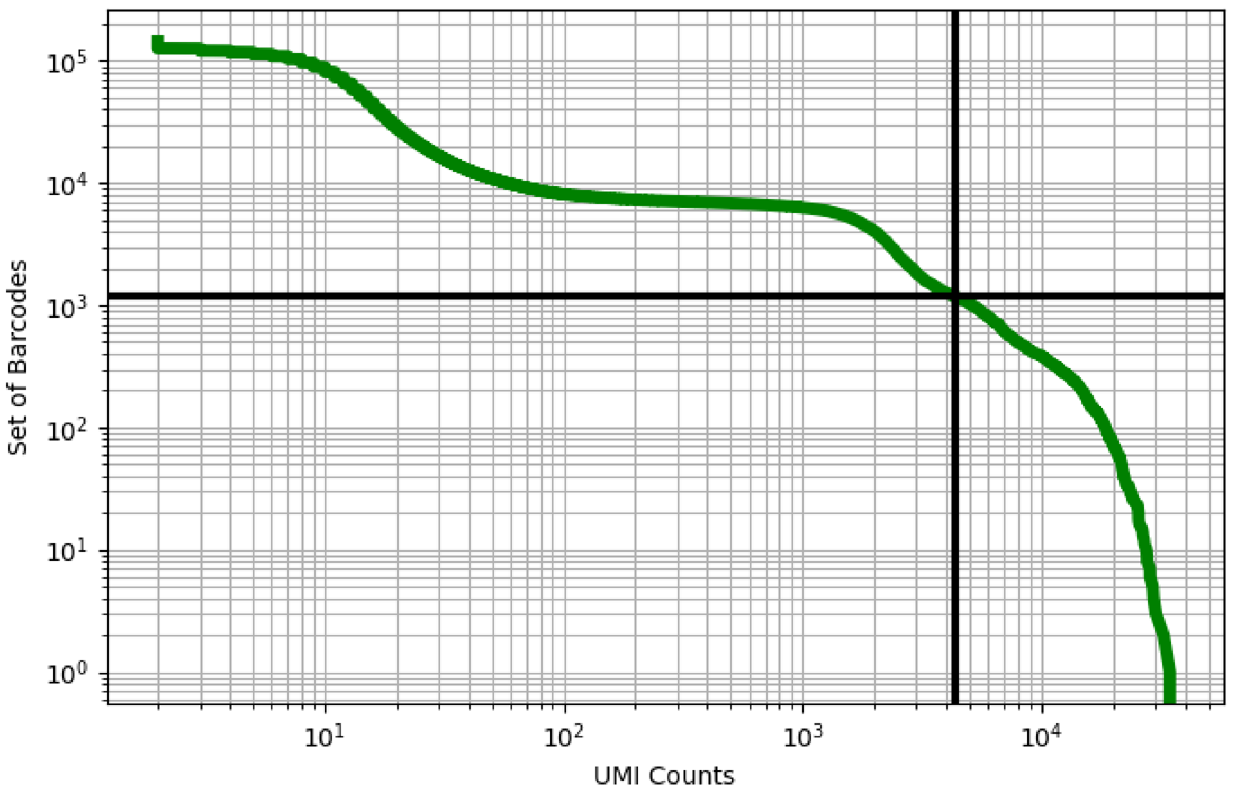


Figure 102: bone_marrow/GSM3396162

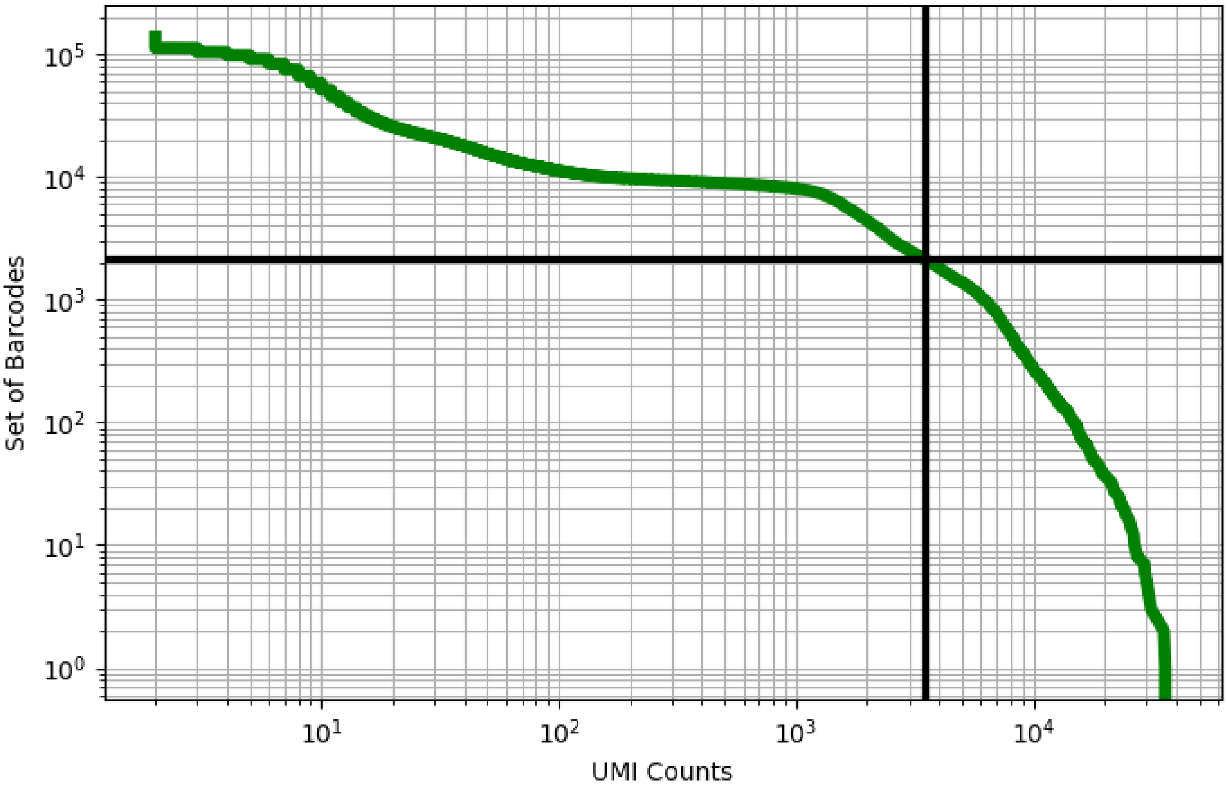


Figure 103: bone_marrow/GSM3396163

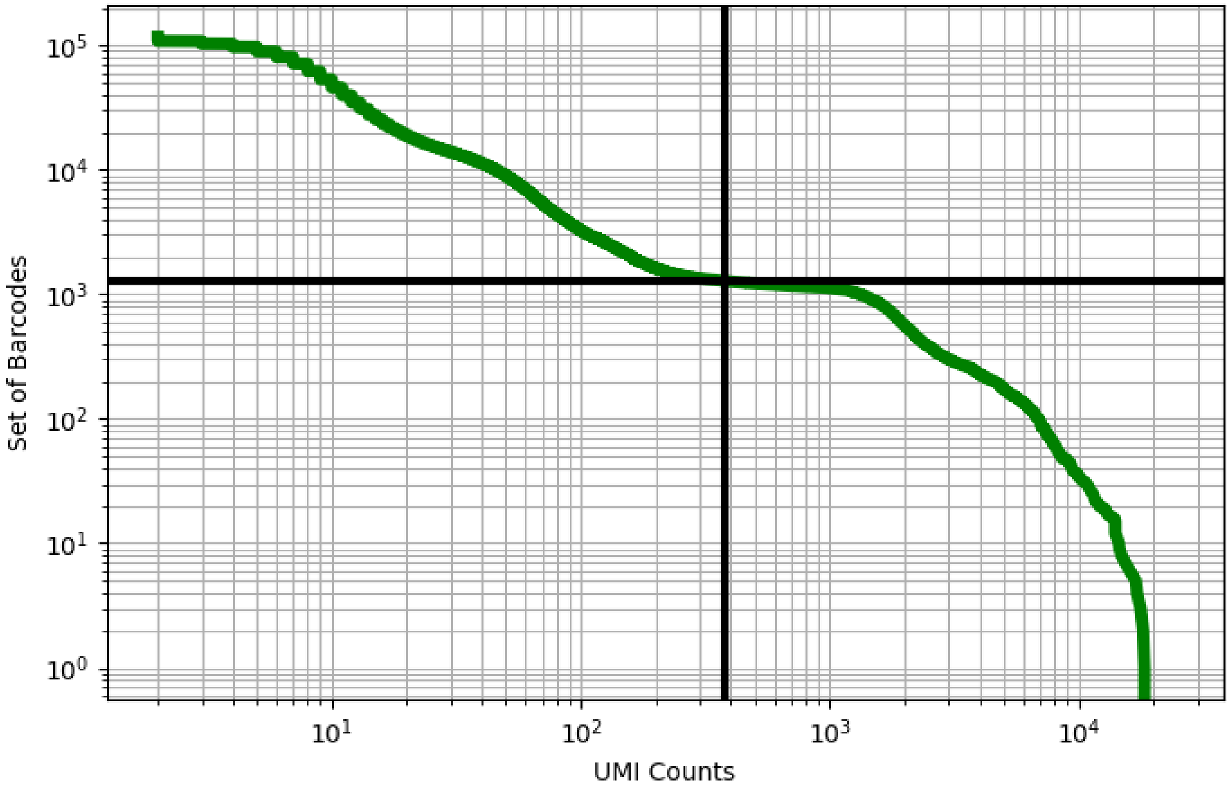


Figure 104: bone_marrow/GSM3396164

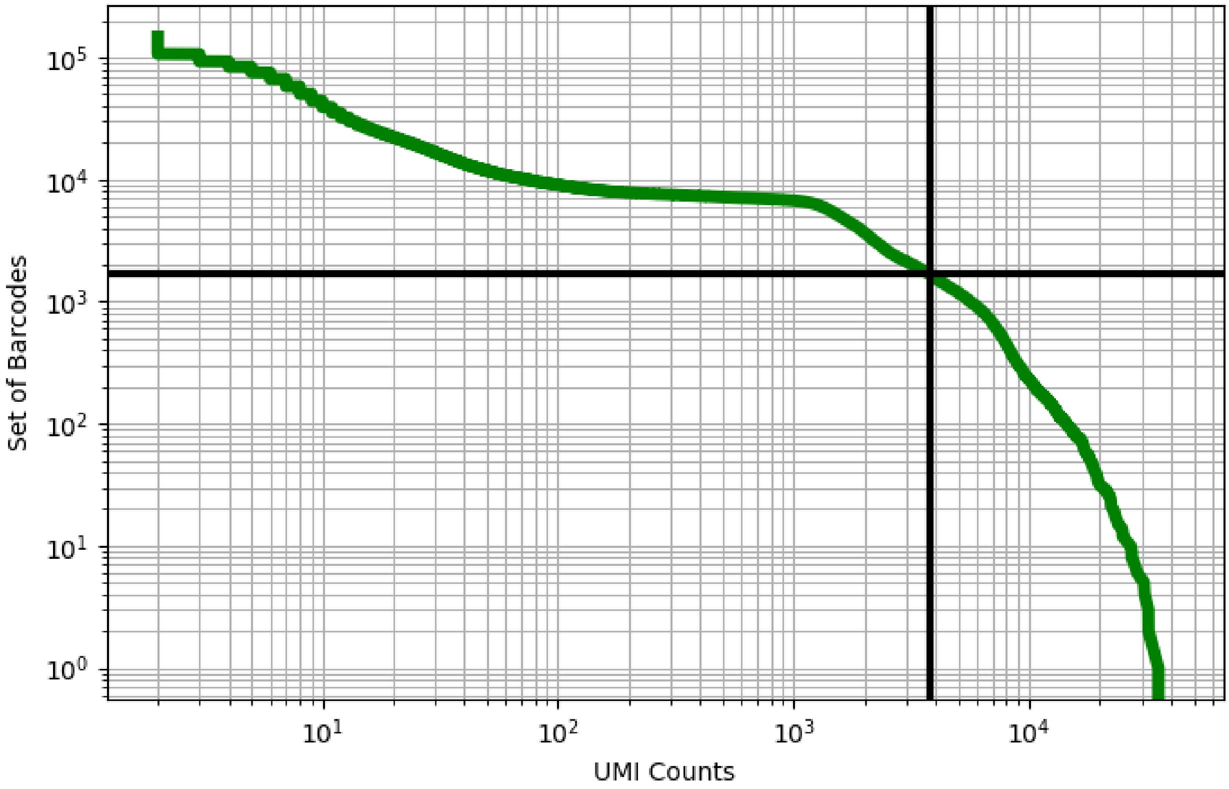


Figure 105: bone_marrow/GSM3396165

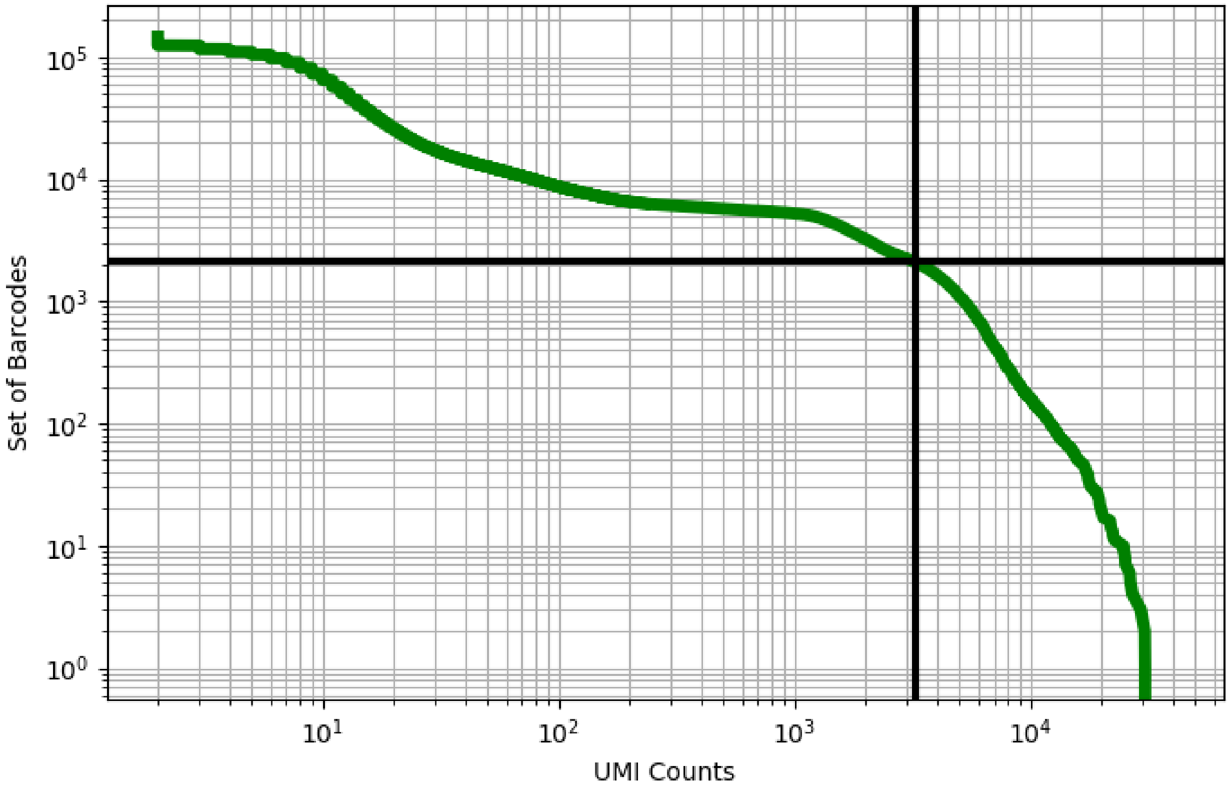


Figure 106: bone_marrow/GSM3396166

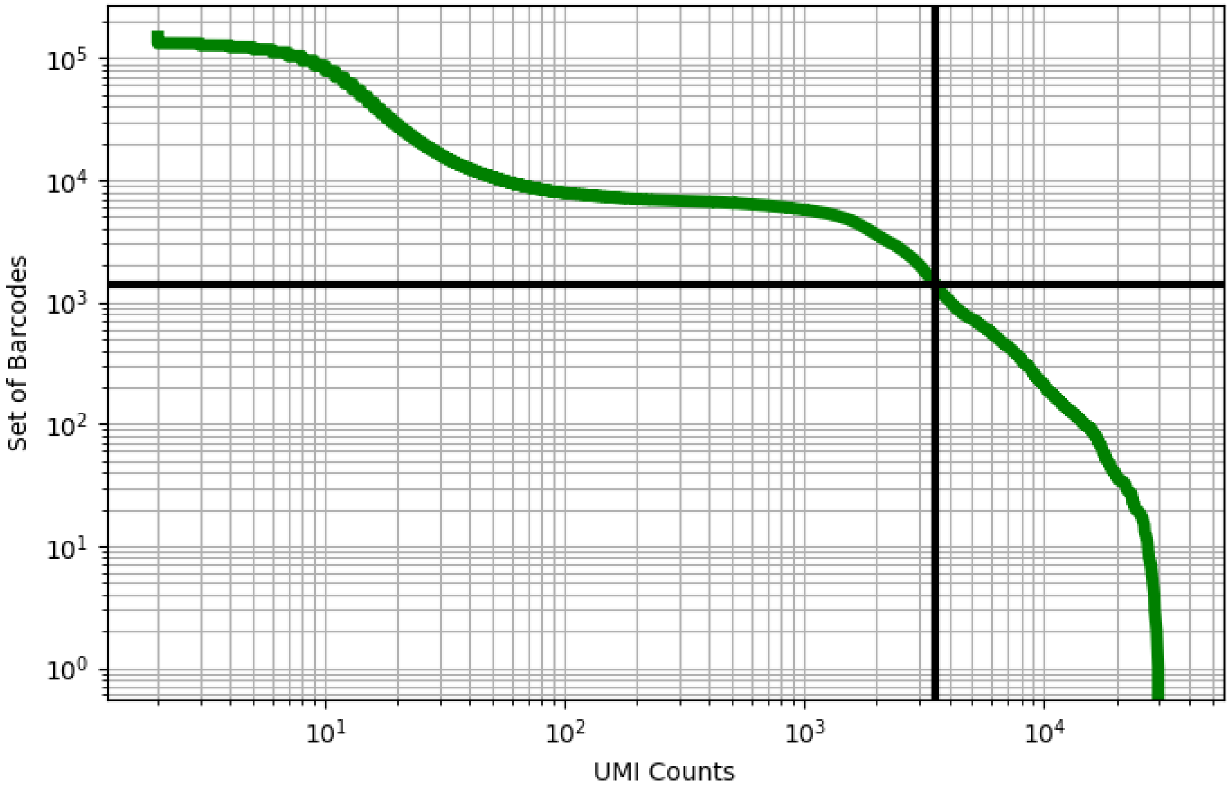


Figure 107: bone_marrow/GSM3396167

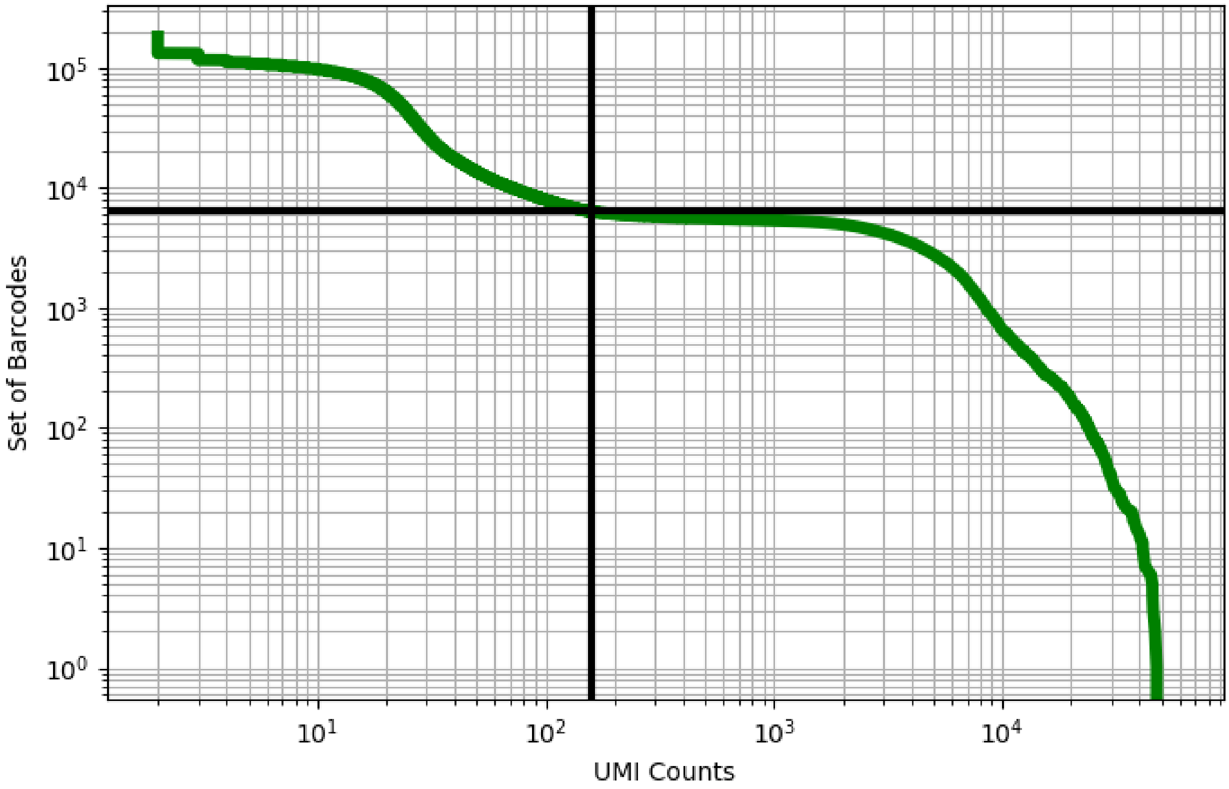


Figure 108: bone_marrow/GSM3396168

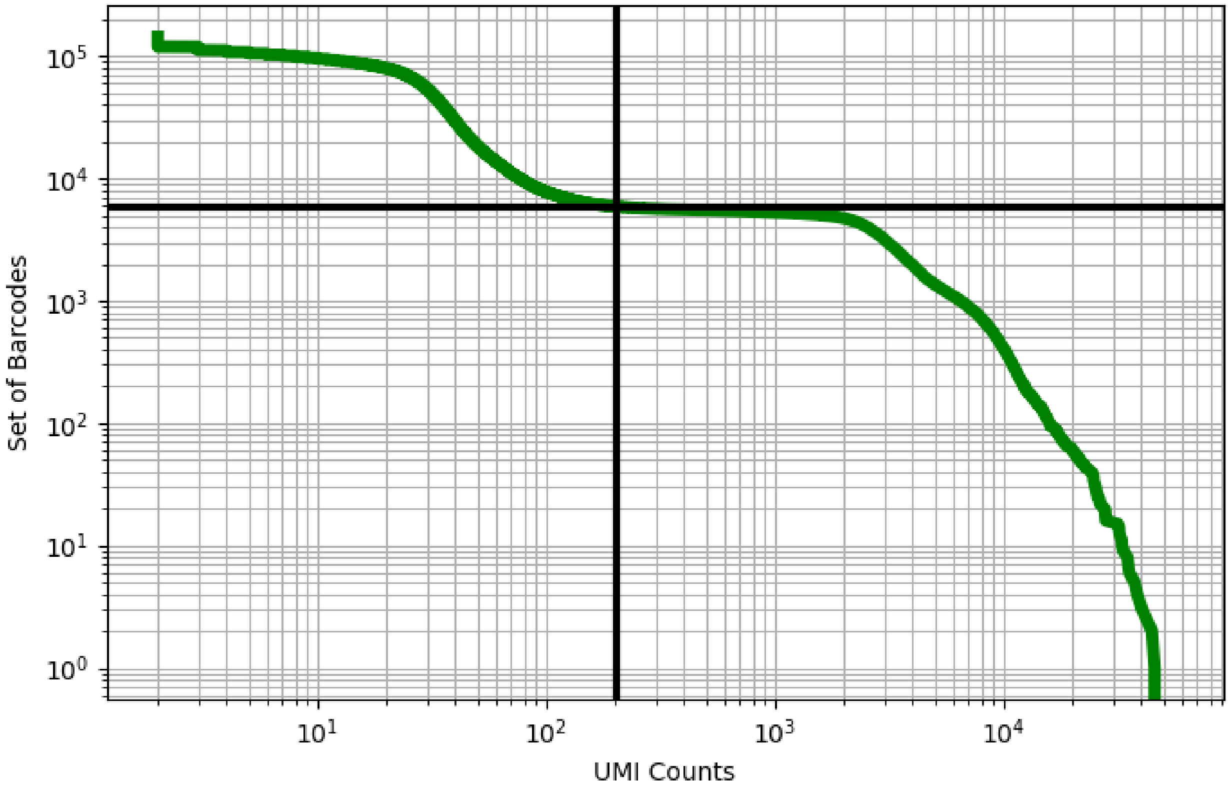


Figure 109: bone_marrow/GSM3396169

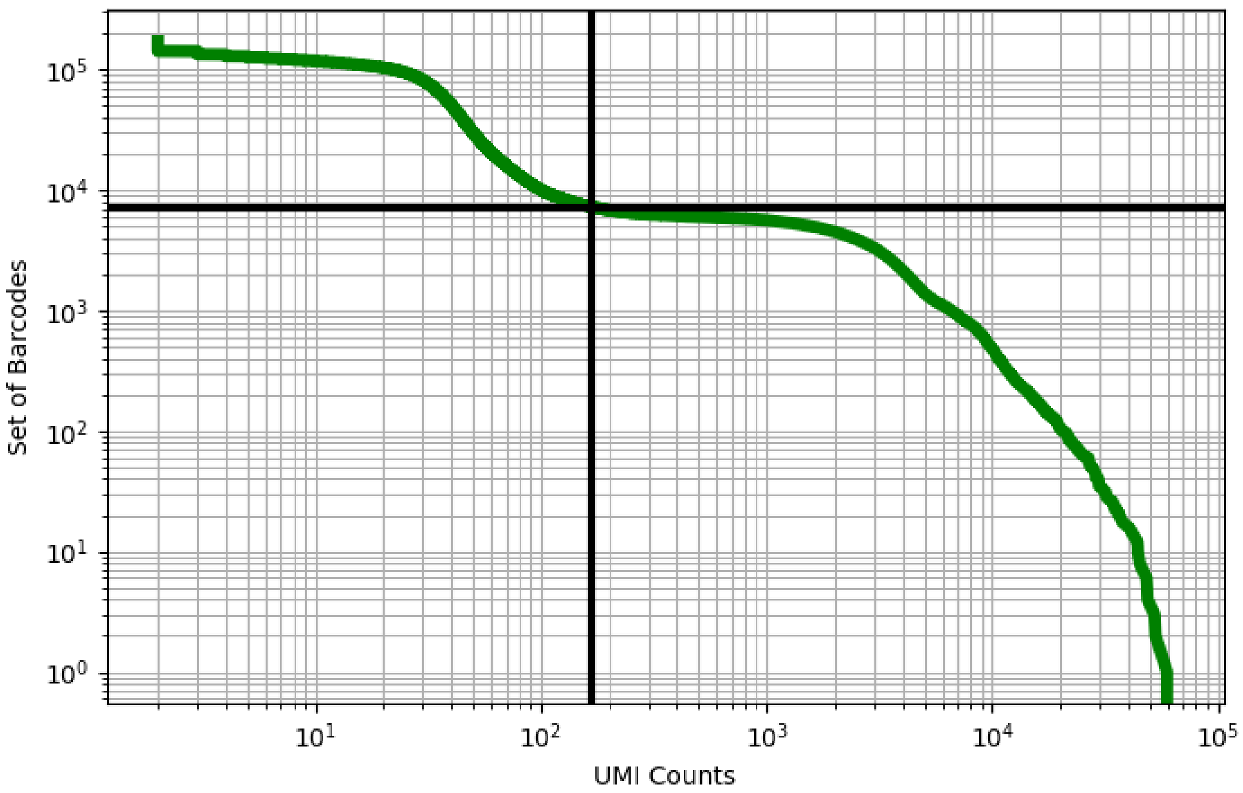


Figure 110: bone_marrow/GSM3396170

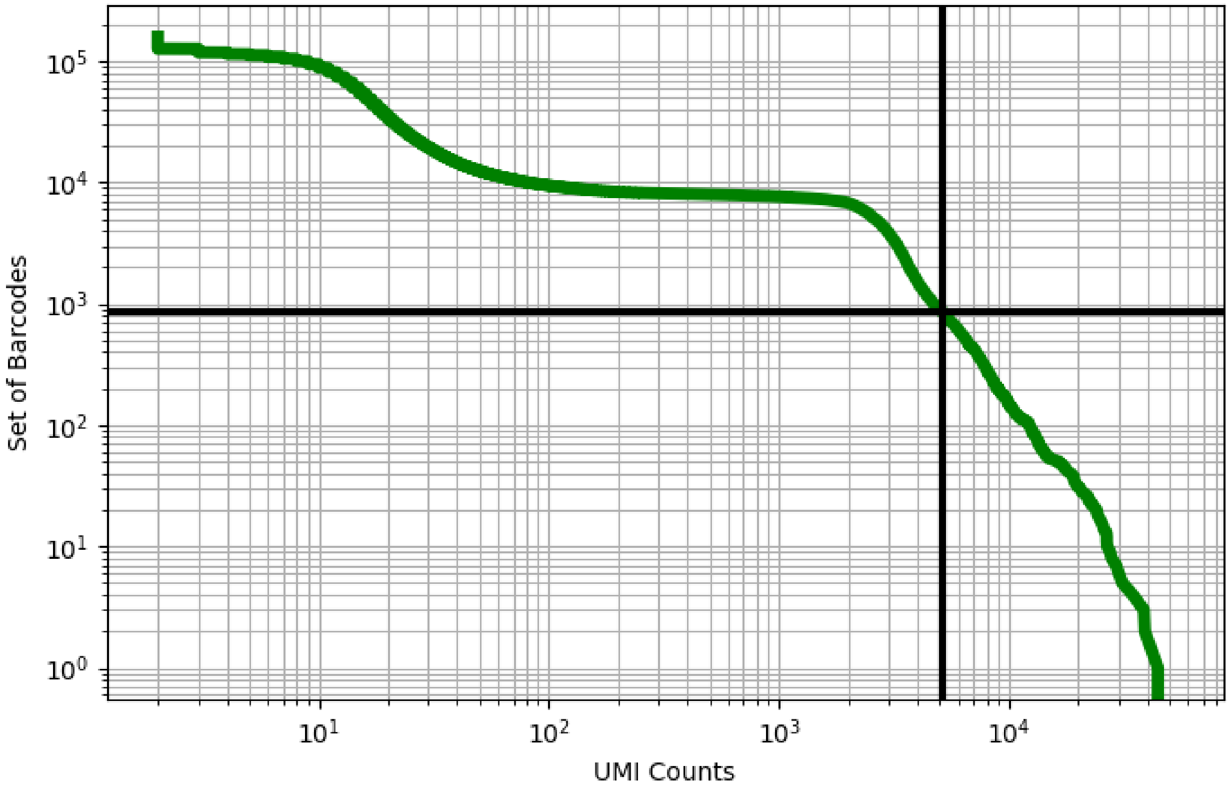


Figure 111: bone_marrow/GSM3396171

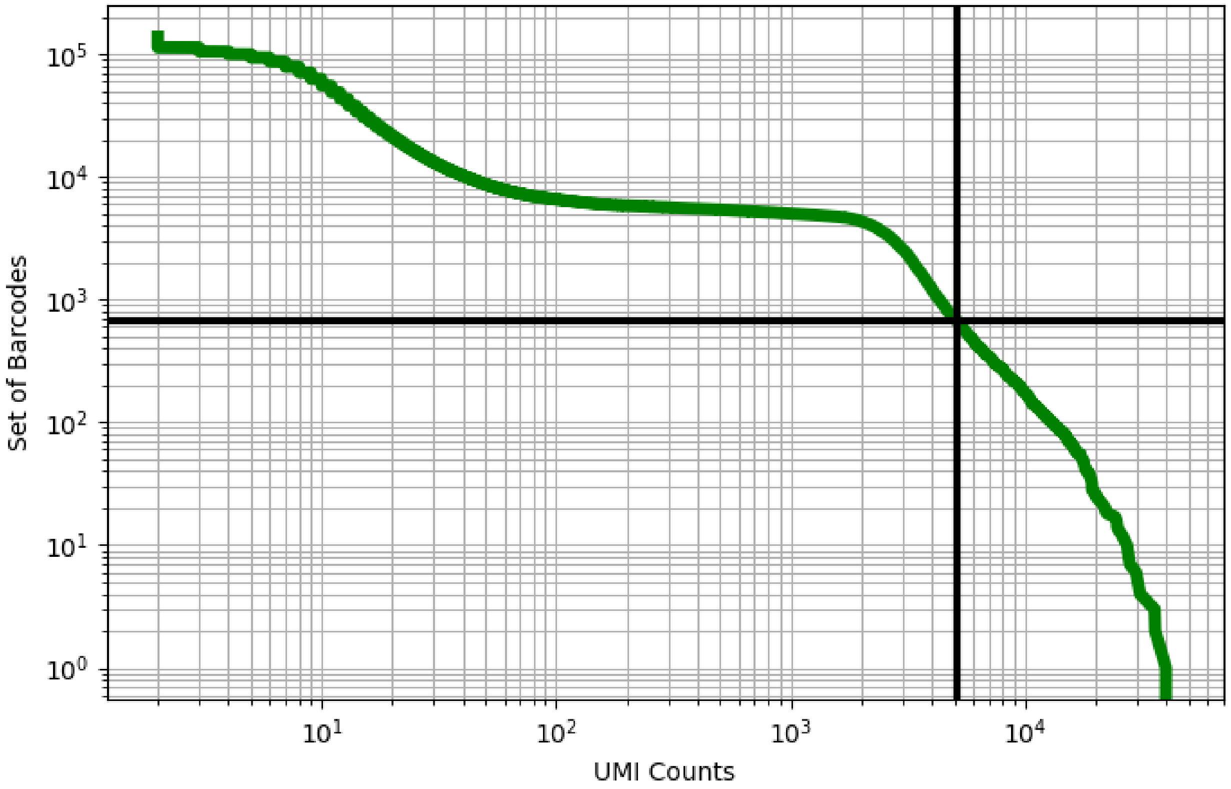


Figure 112: bone_marrow/GSM3396172

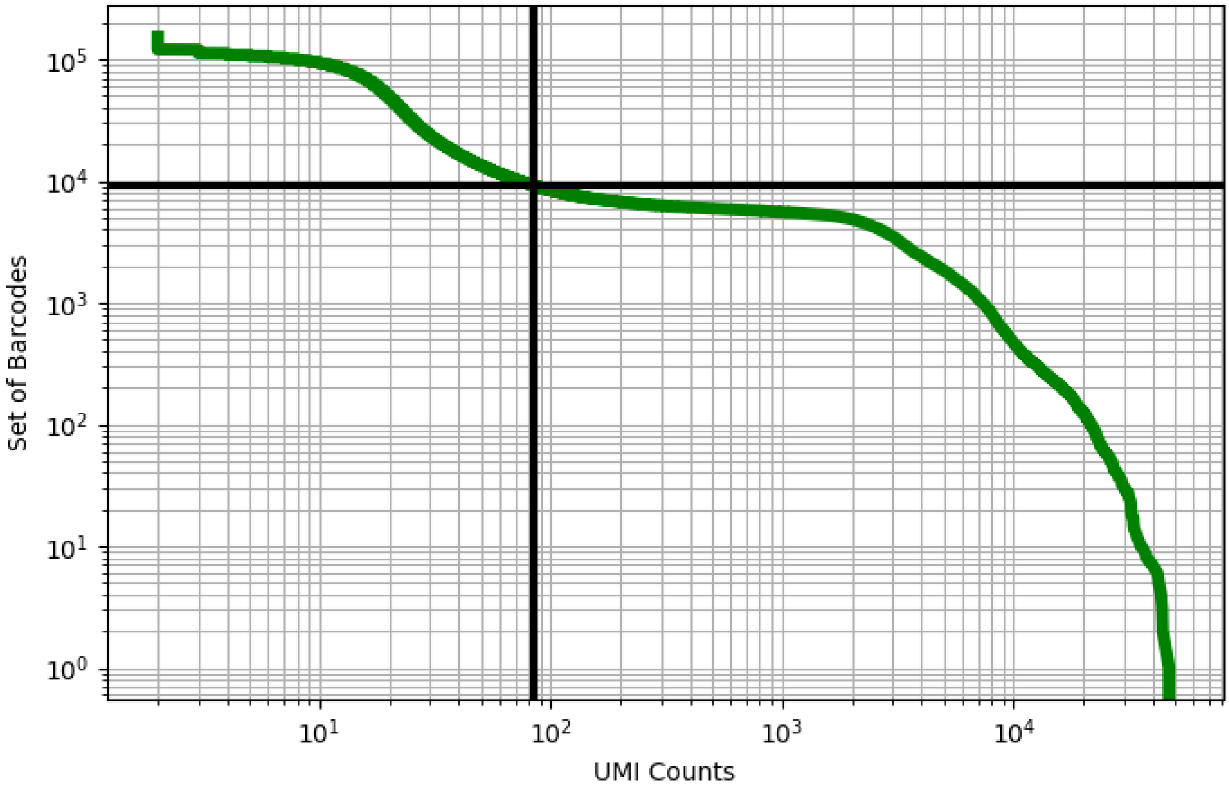


Figure 113: bone_marrow/GSM3396173

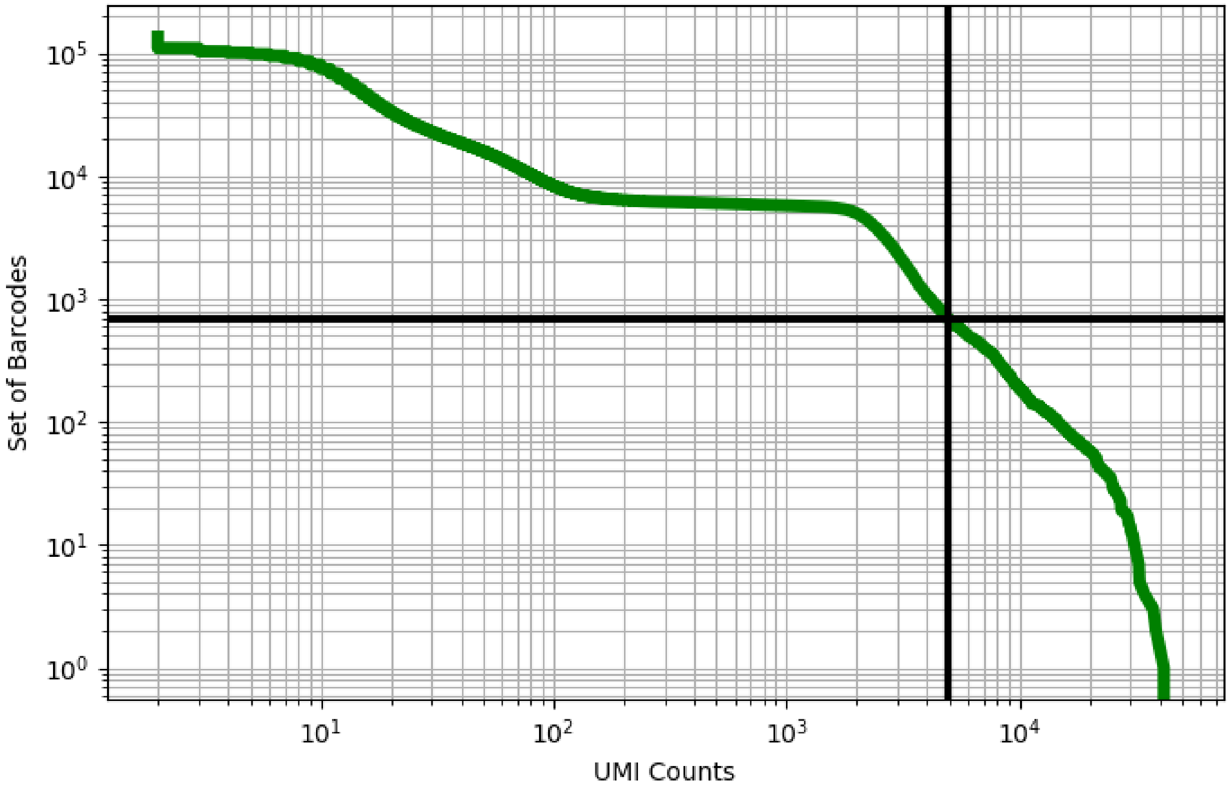


Figure 114: bone_marrow/GSM3396174

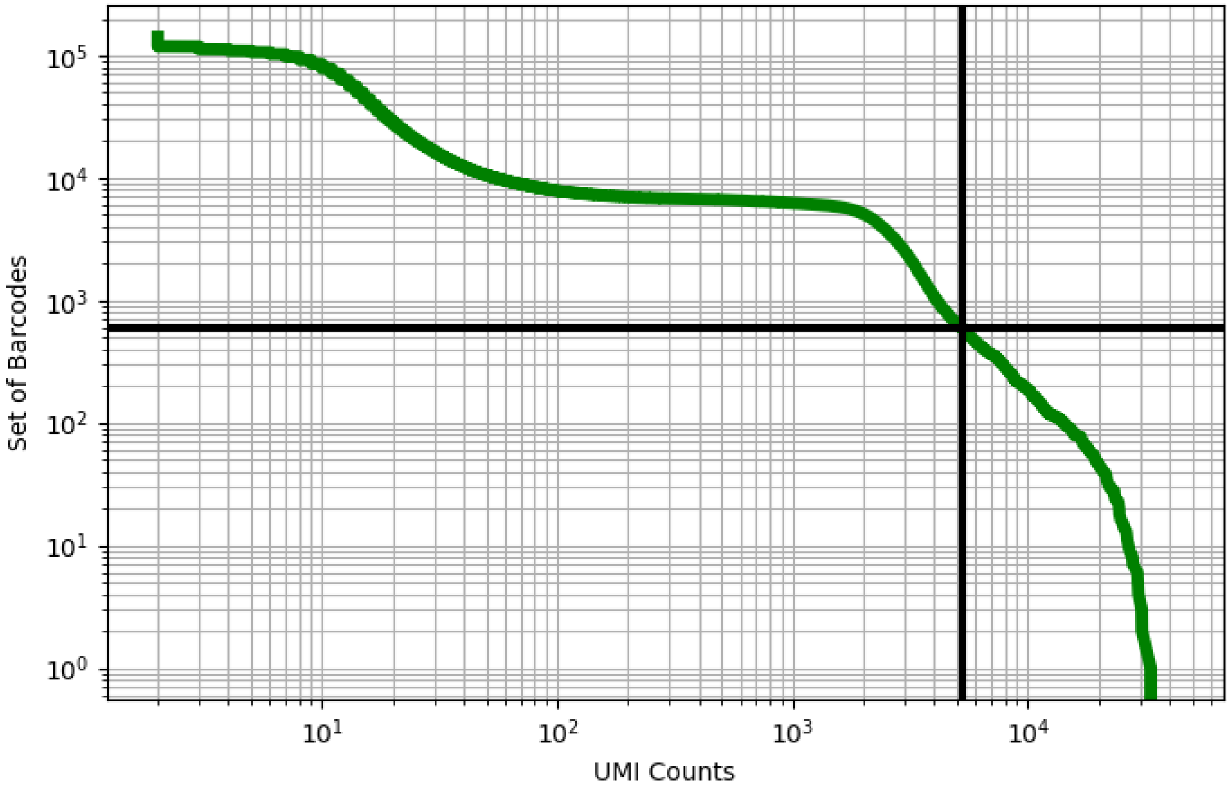


Figure 115: bone_marrow/GSM3396175

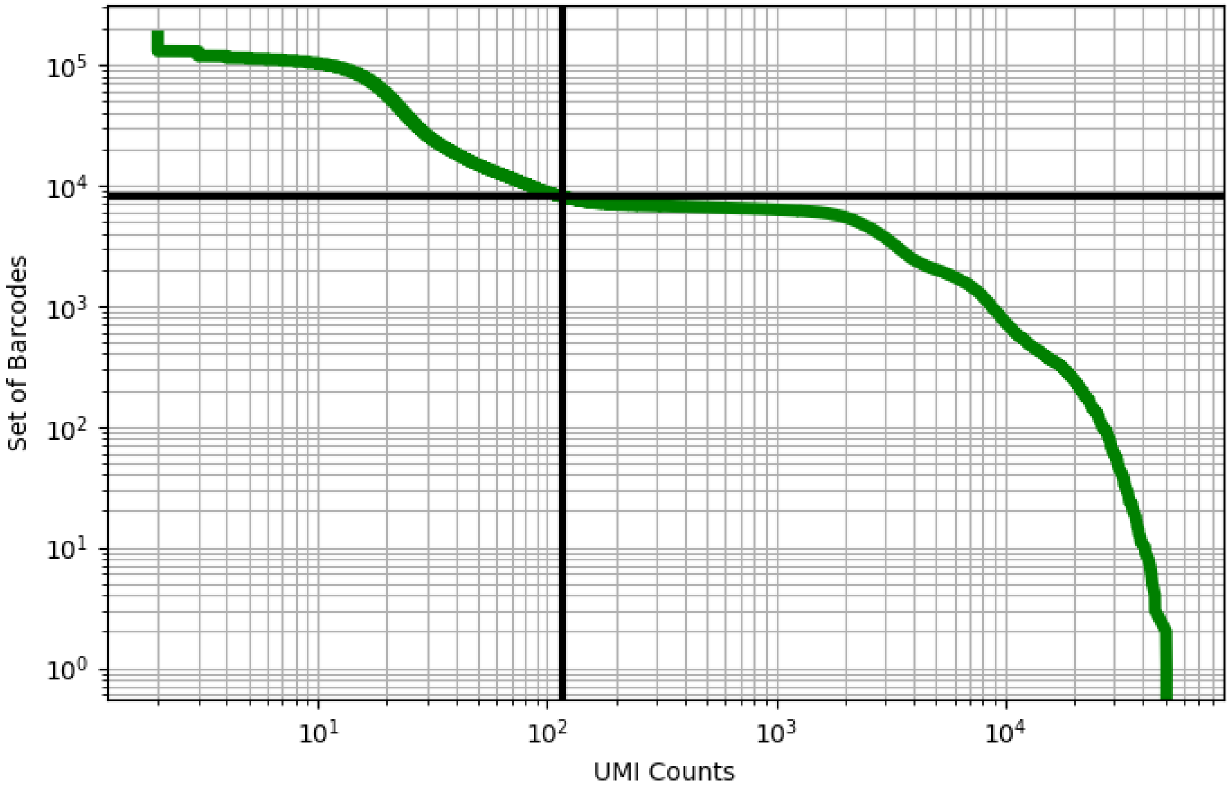


Figure 116: bone_marrow/GSM3396176

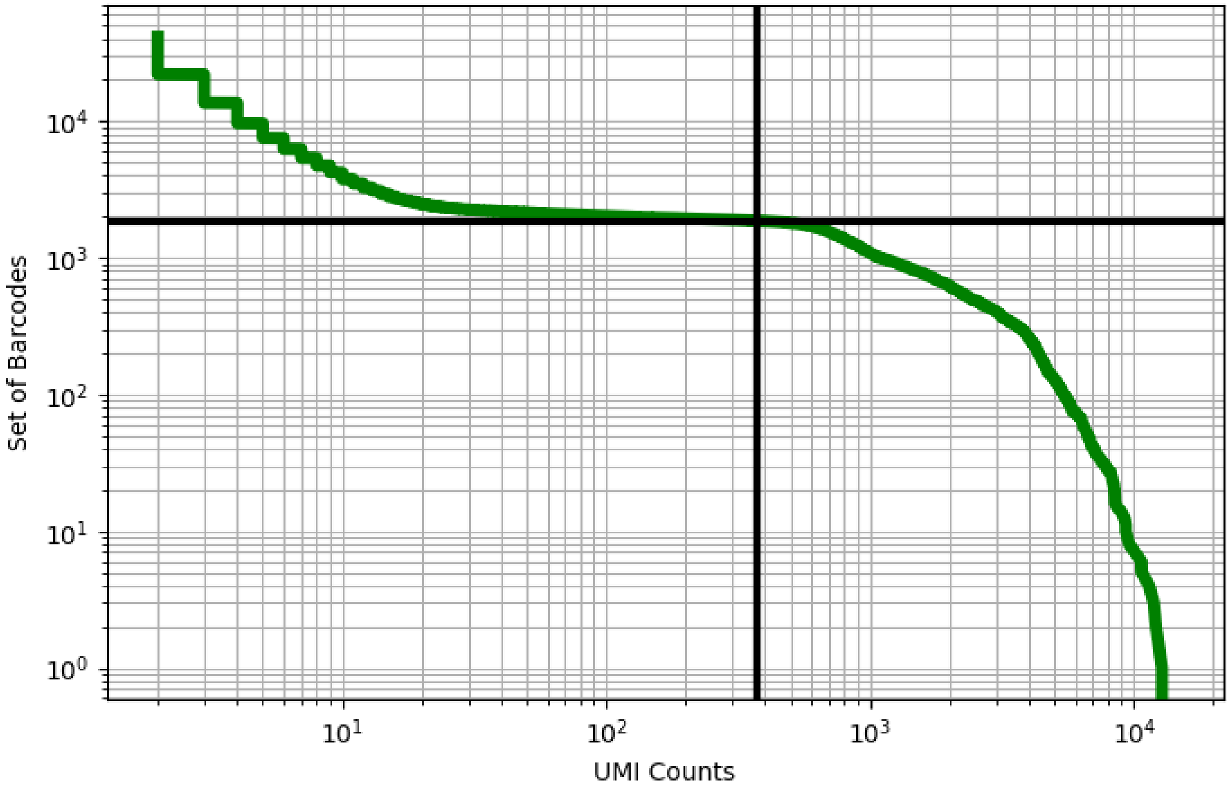


Figure 117: bone_marrow/GSM3396177

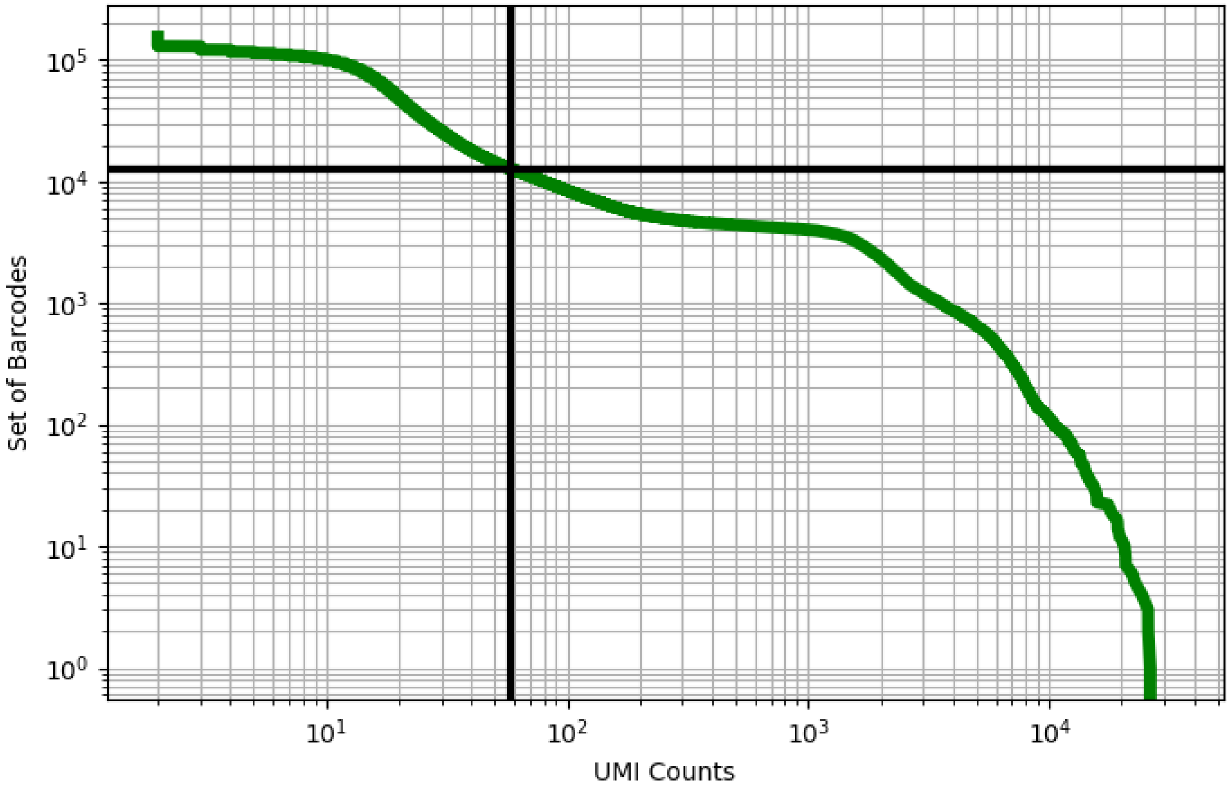


Figure 118: bone_marrow/GSM3396178

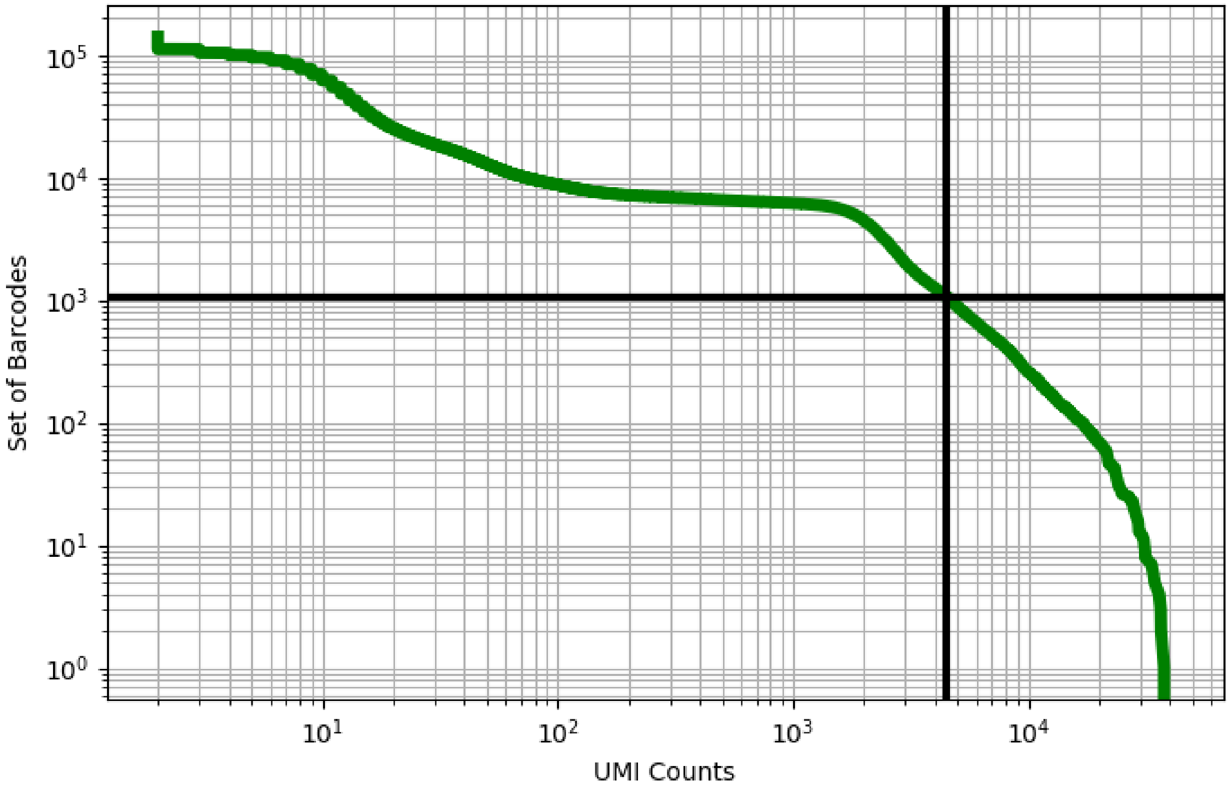


Figure 119: bone_marrow/GSM3396179

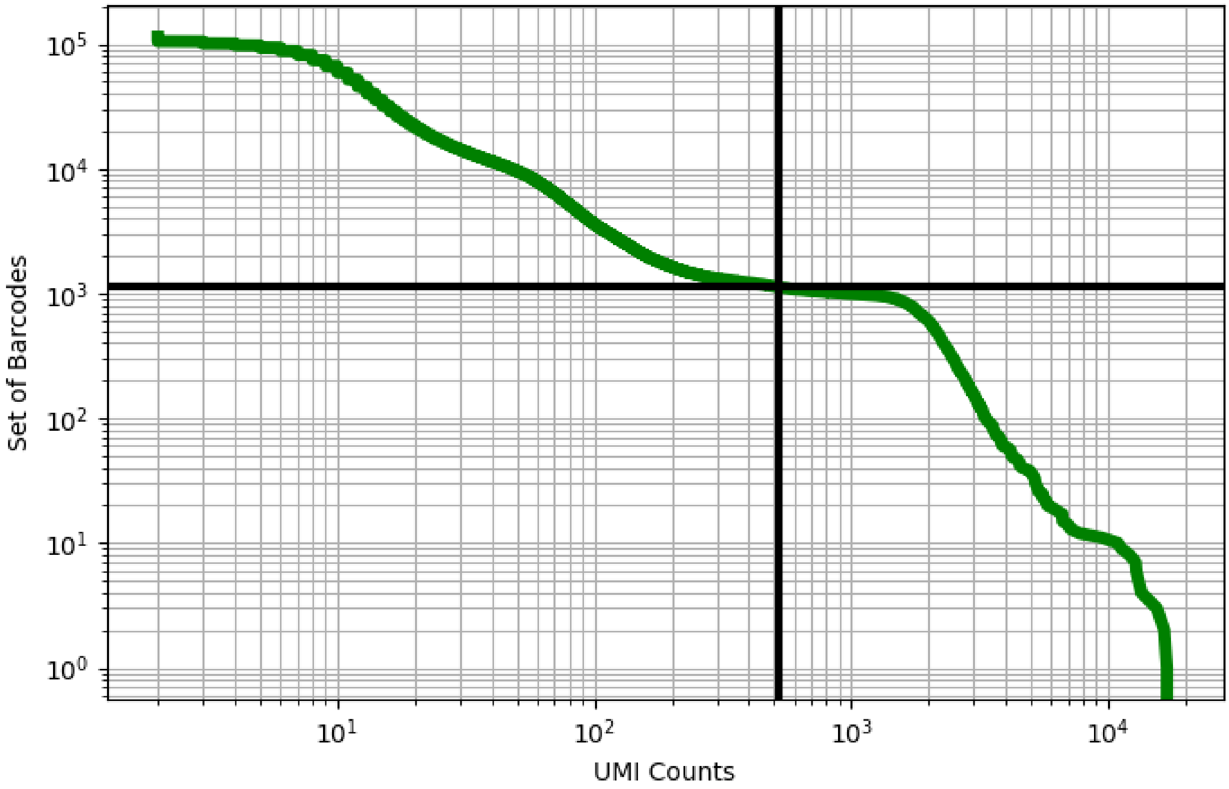


Figure 120: bone_marrow/GSM3396180

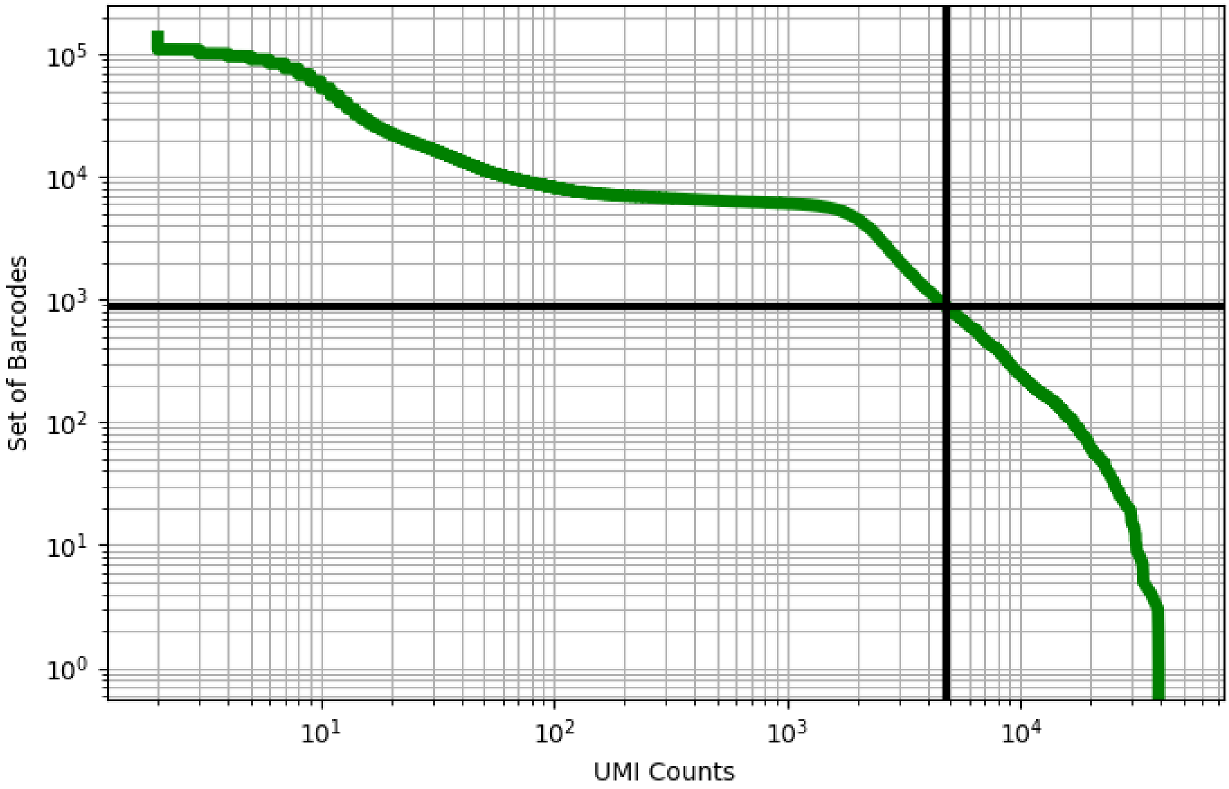


Figure 121: bone_marrow/GSM3396181

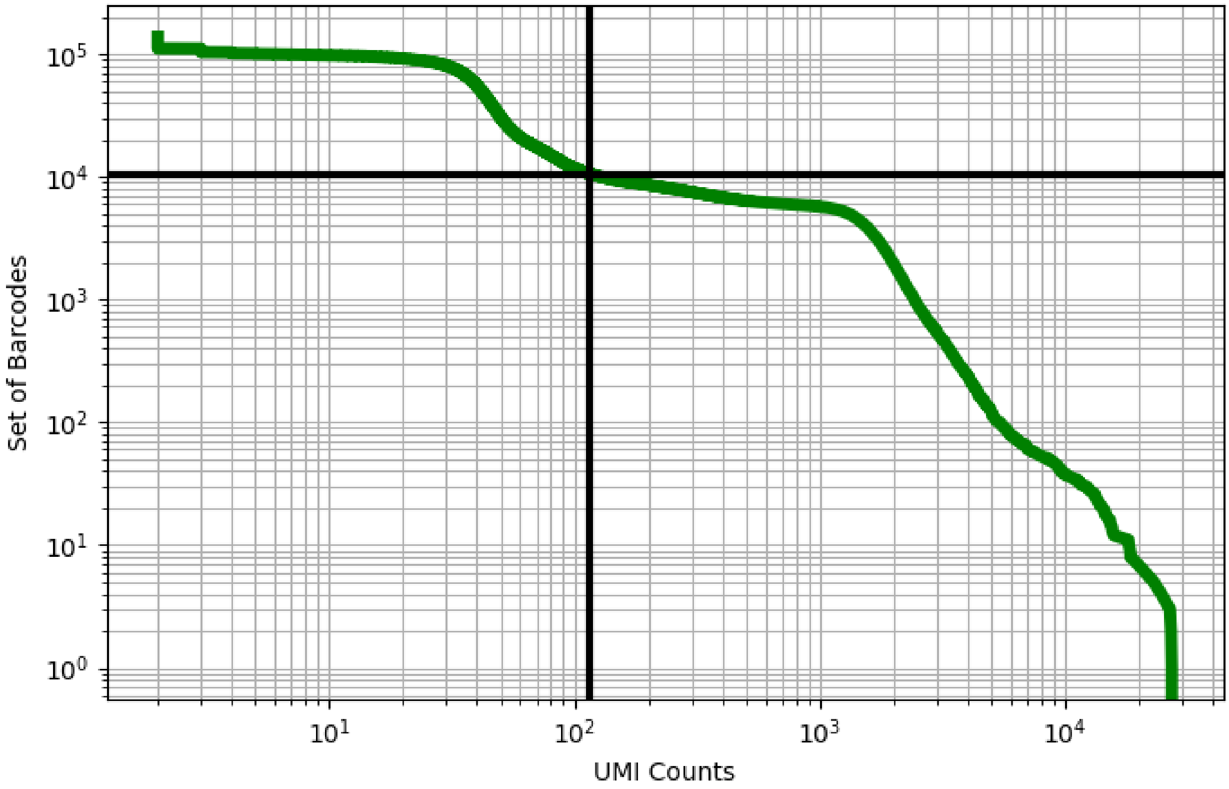


Figure 122: bone_marrow/GSM3396182

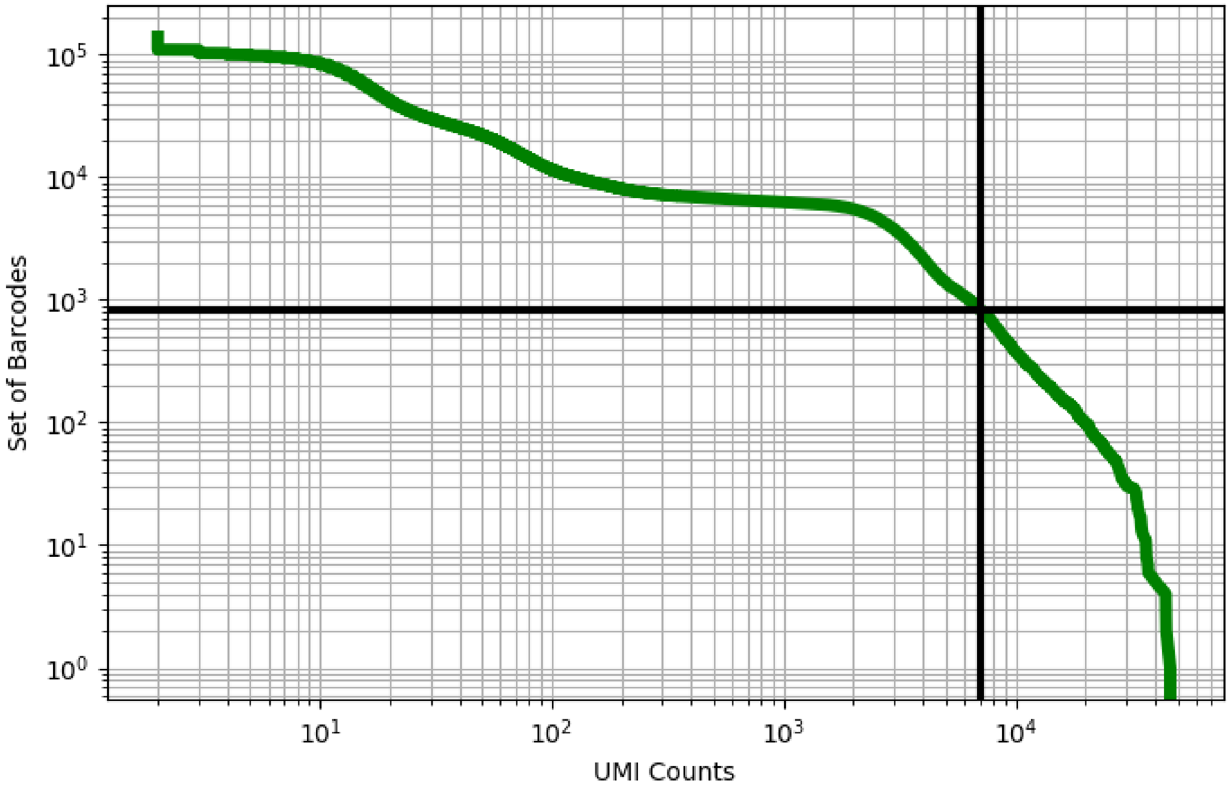


Figure 123: bone_marrow/GSM3396183

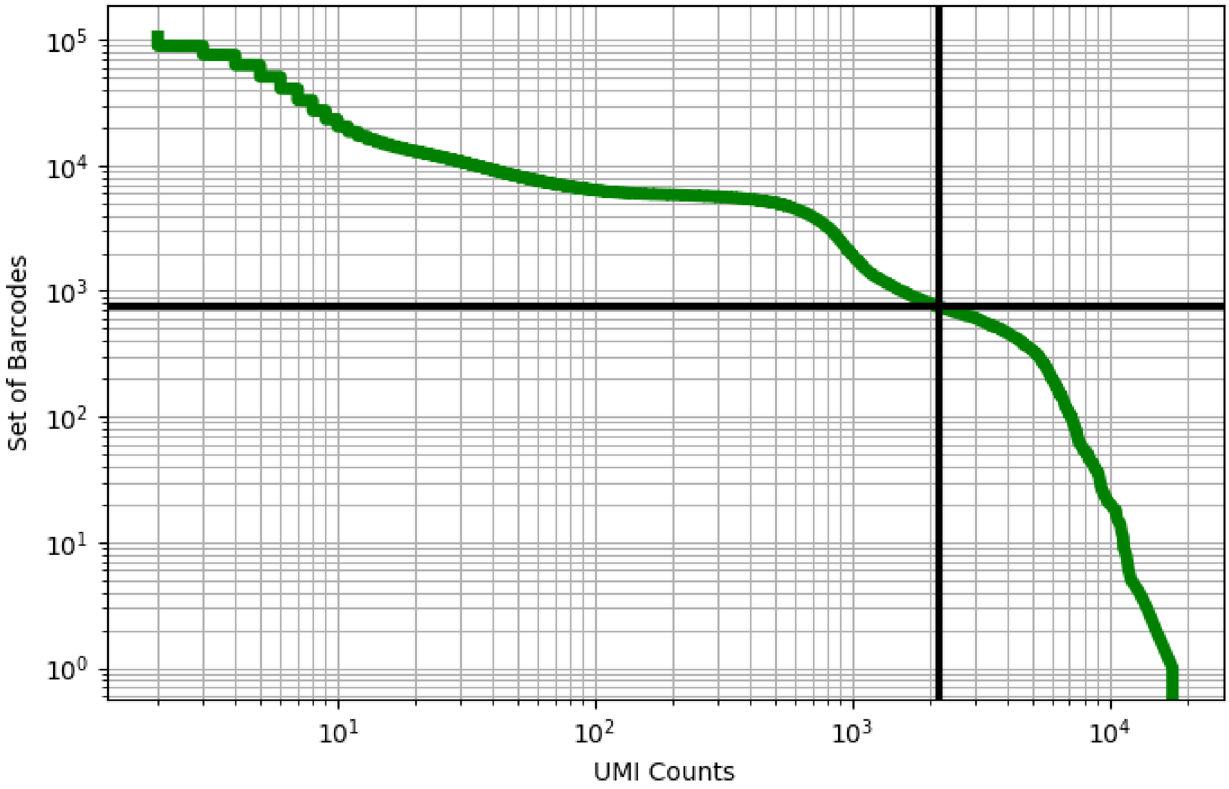


Figure 124: bone_marrow/GSM3396184

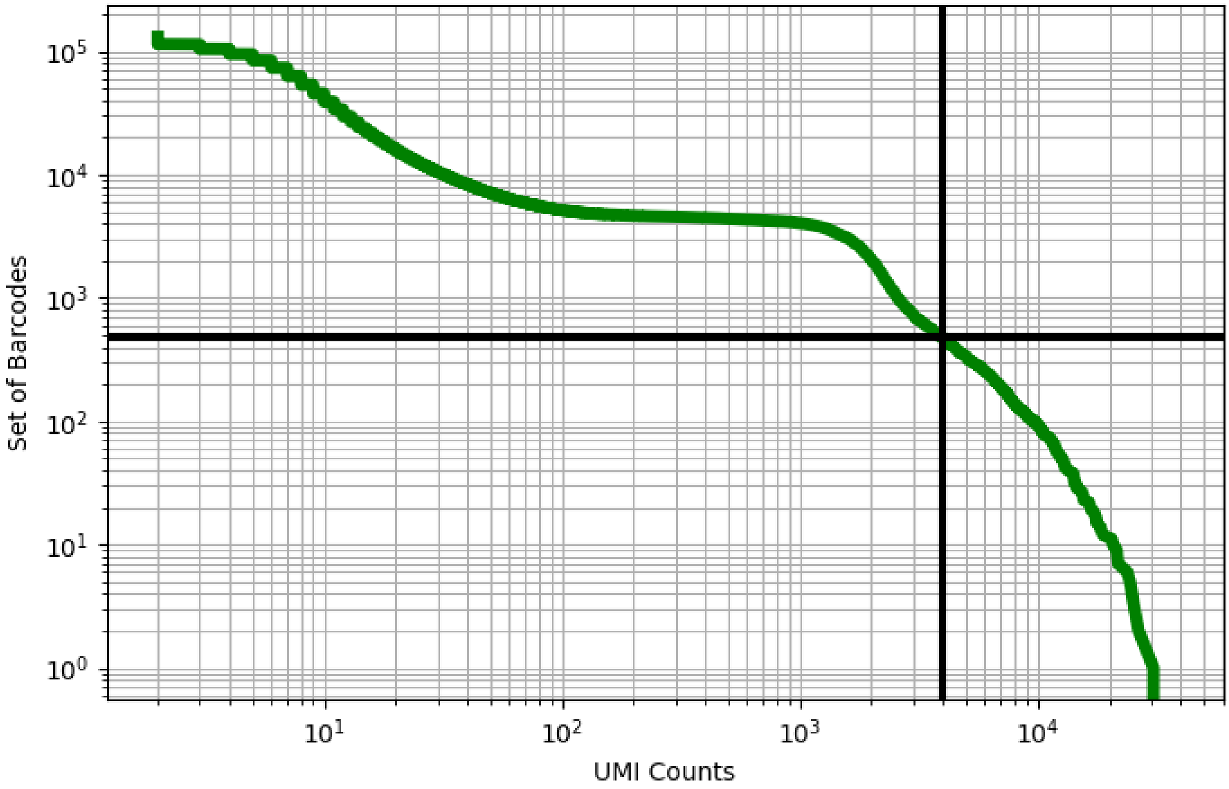


Figure 125: bone_marrow/GSM3396185

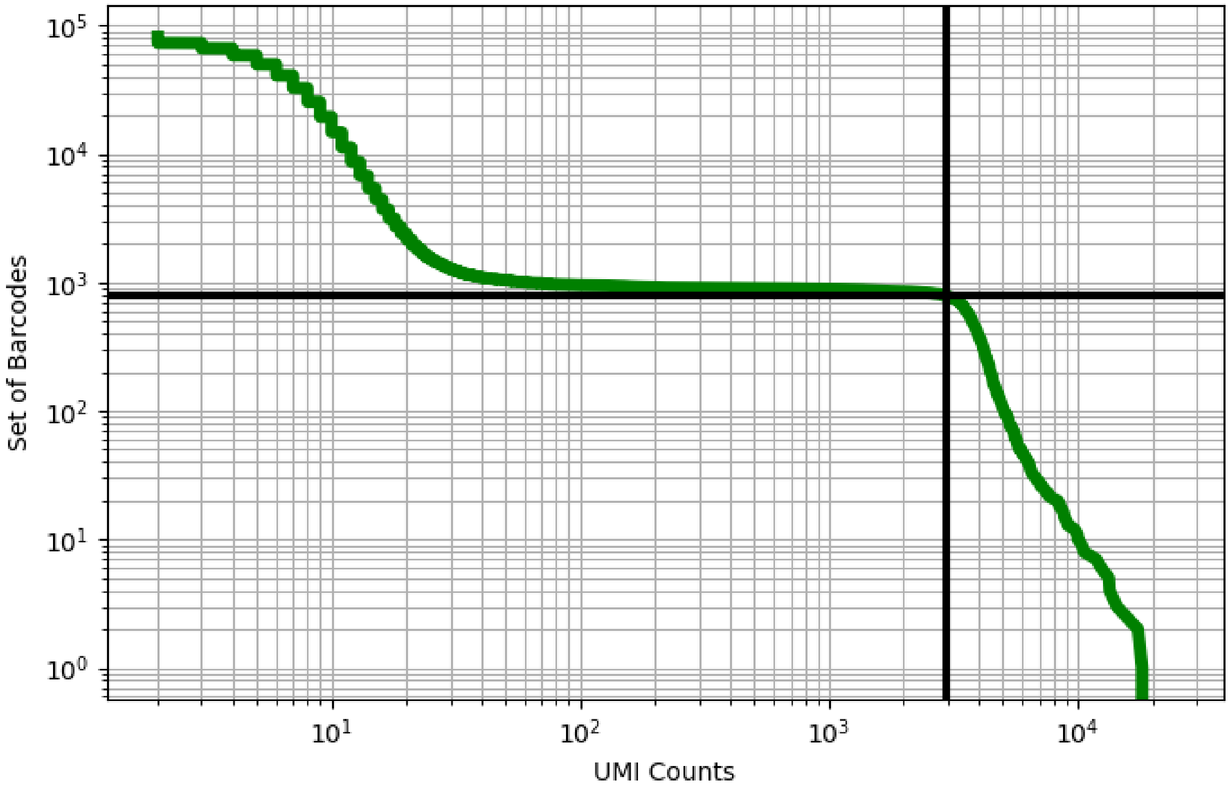


Figure 126: bone_marrow/GSM3738536

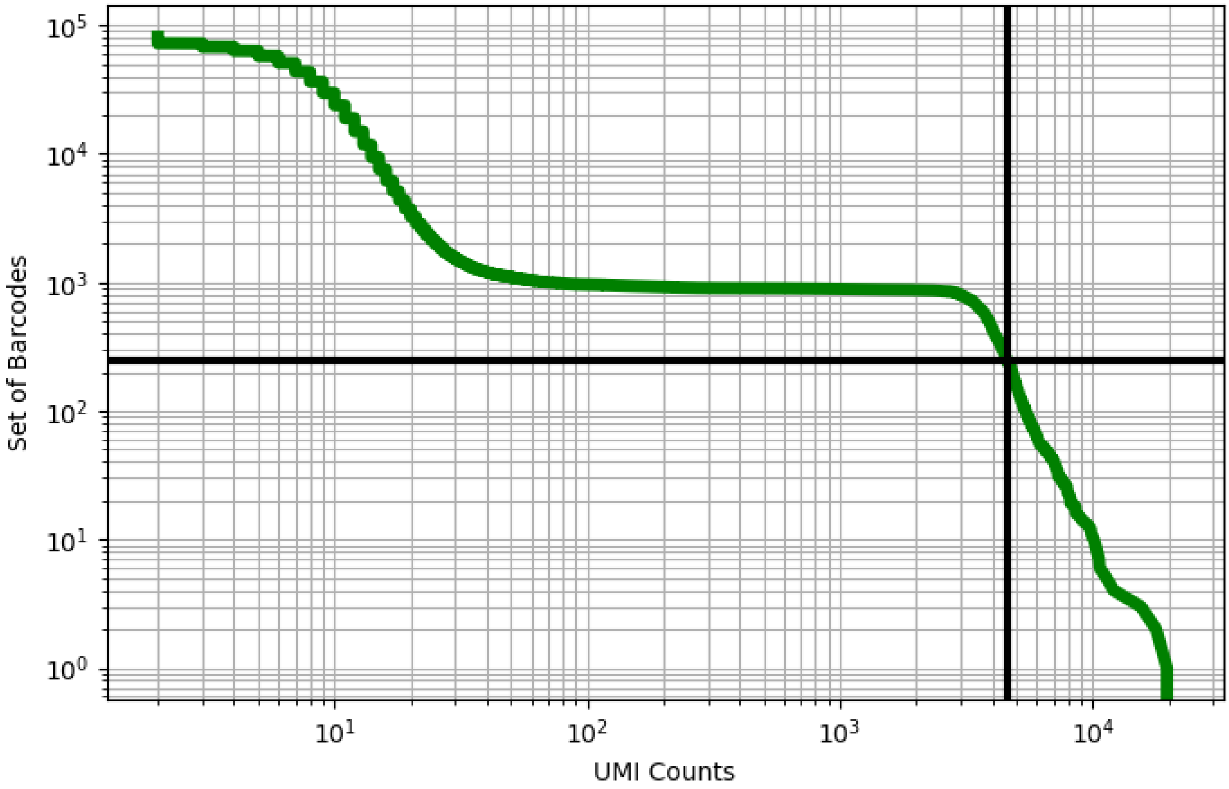


Figure 127: bone_marrow/GSM3738537

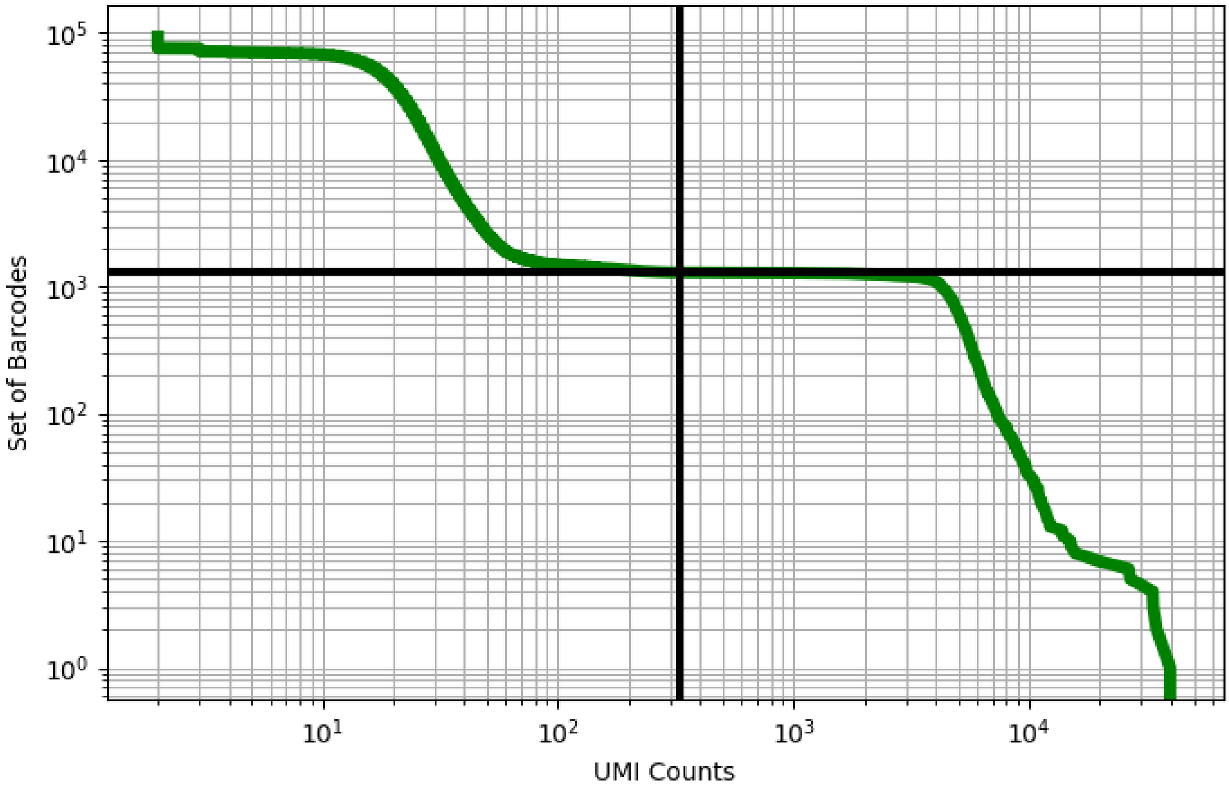


Figure 128: bone_marrow/GSM3738538

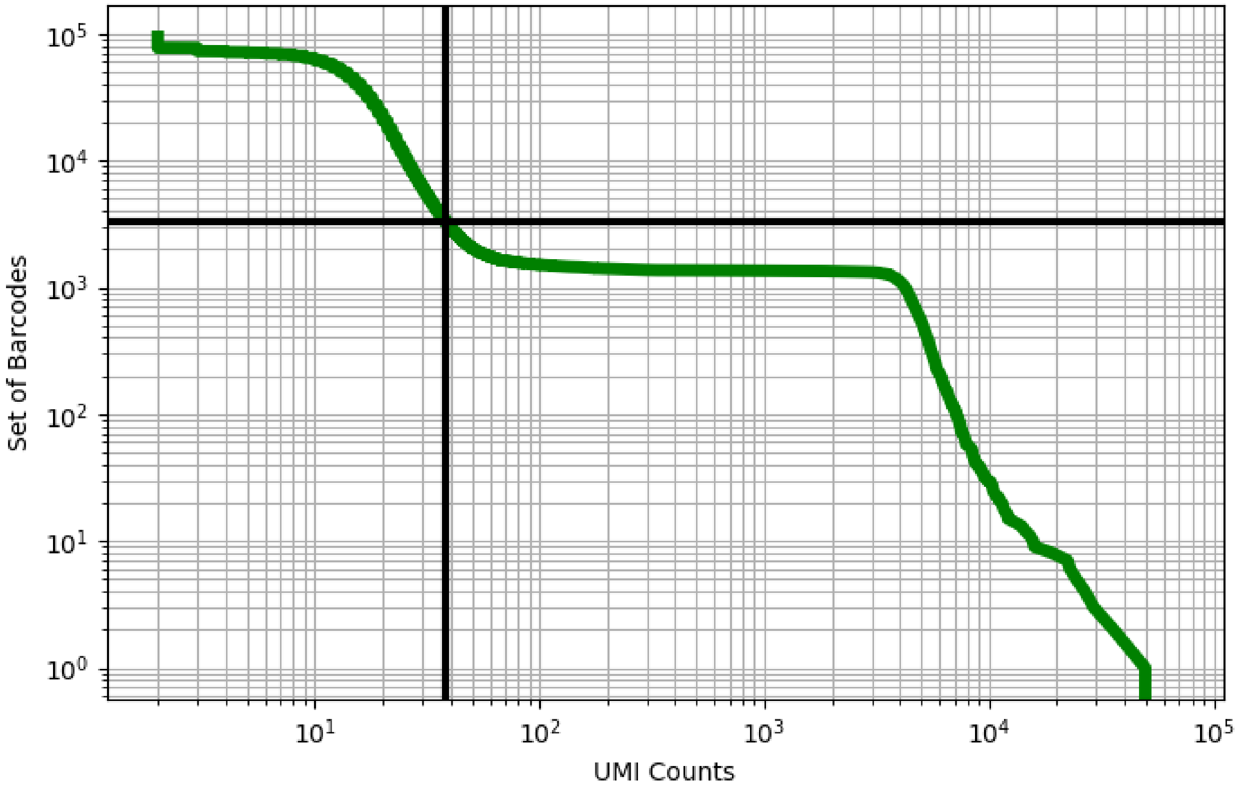


Figure 129: bone_marrow/GSM3738539

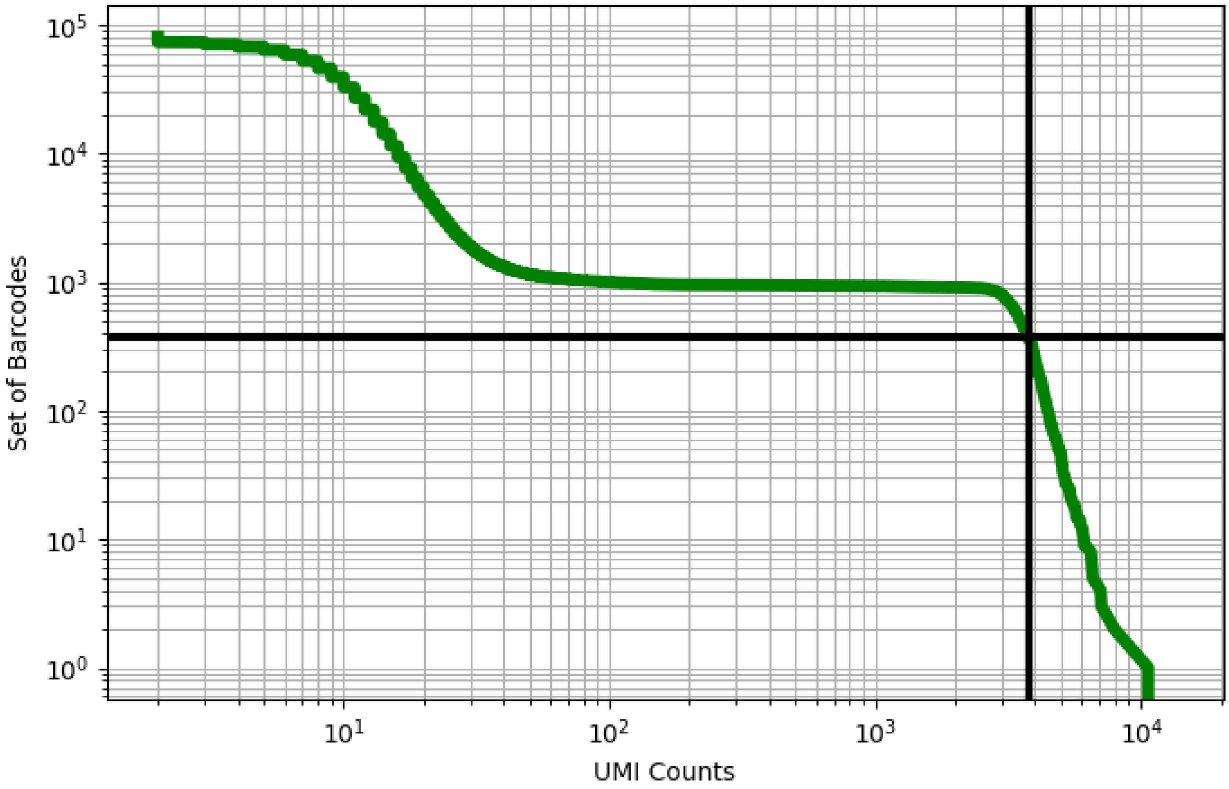


Figure 130: bone_marrow/GSM3738540

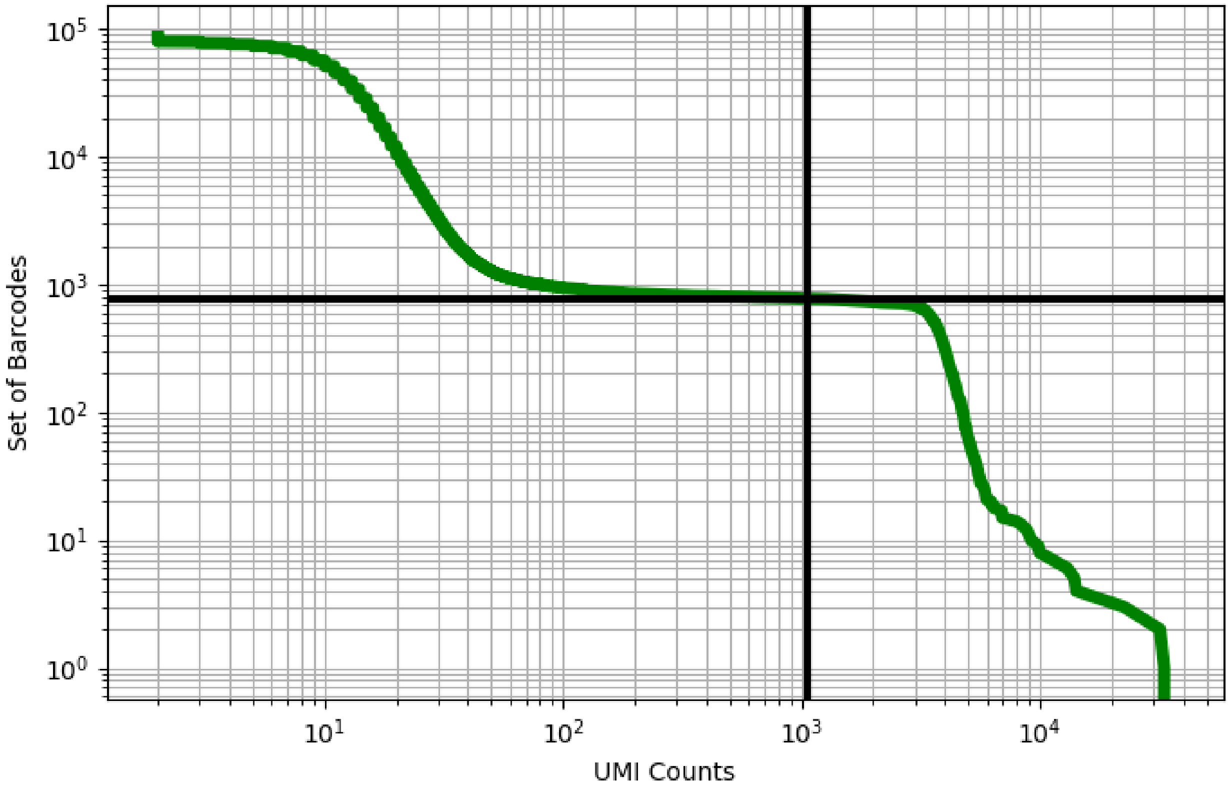


Figure 131: bone_marrow/GSM3738541

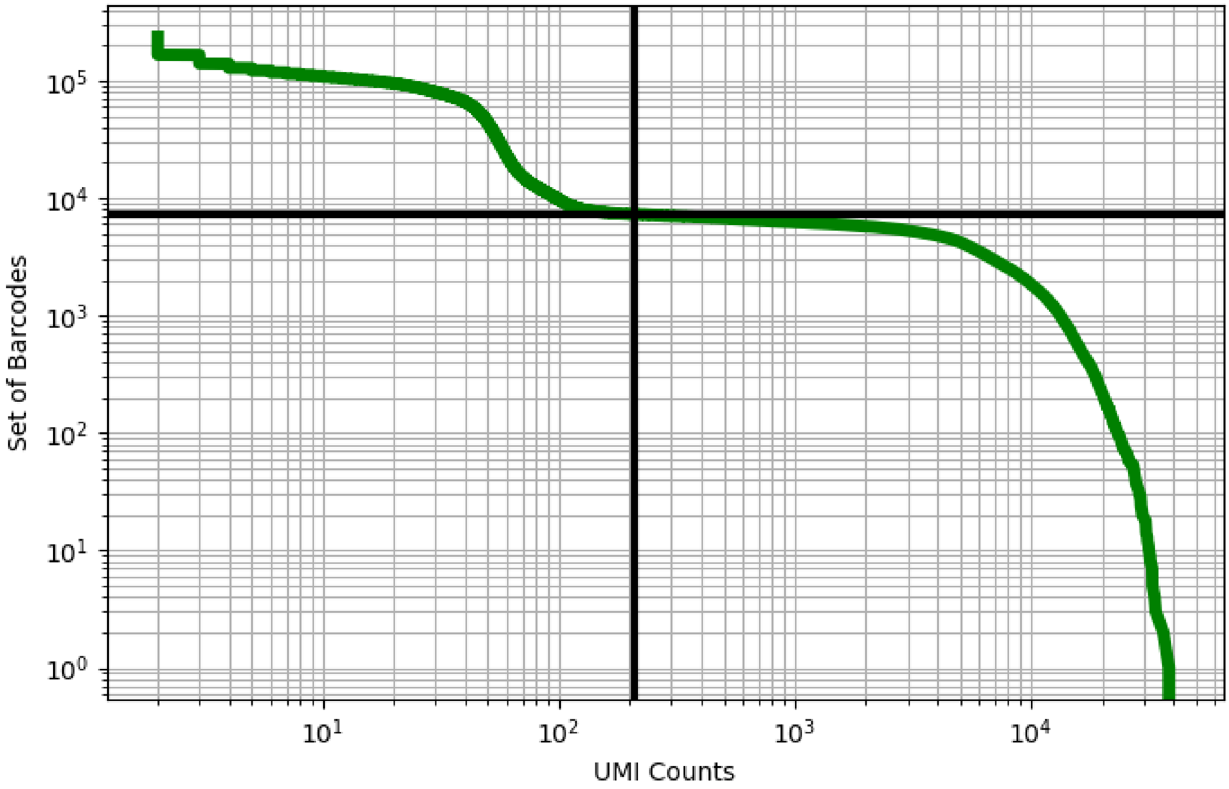


Figure 132: bone_marrow/GSM3739217

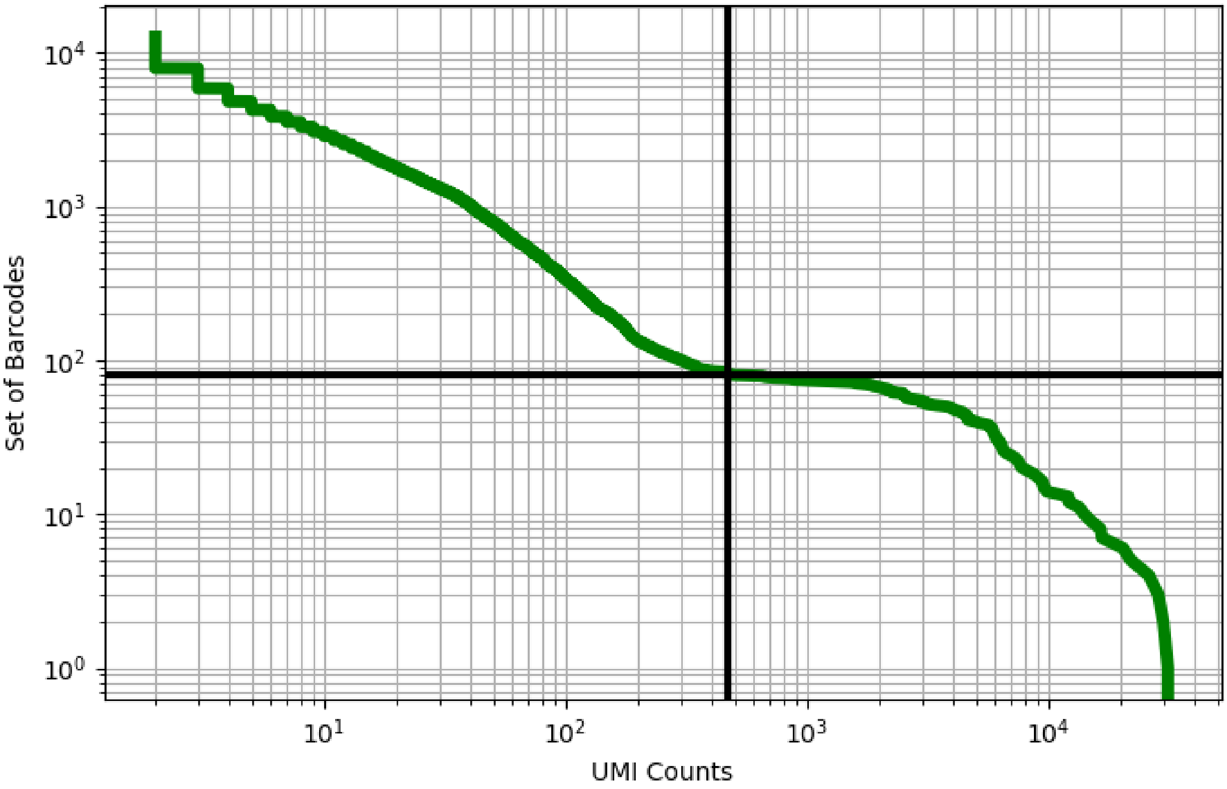


Figure 133: bone_marrow/GSM3739218

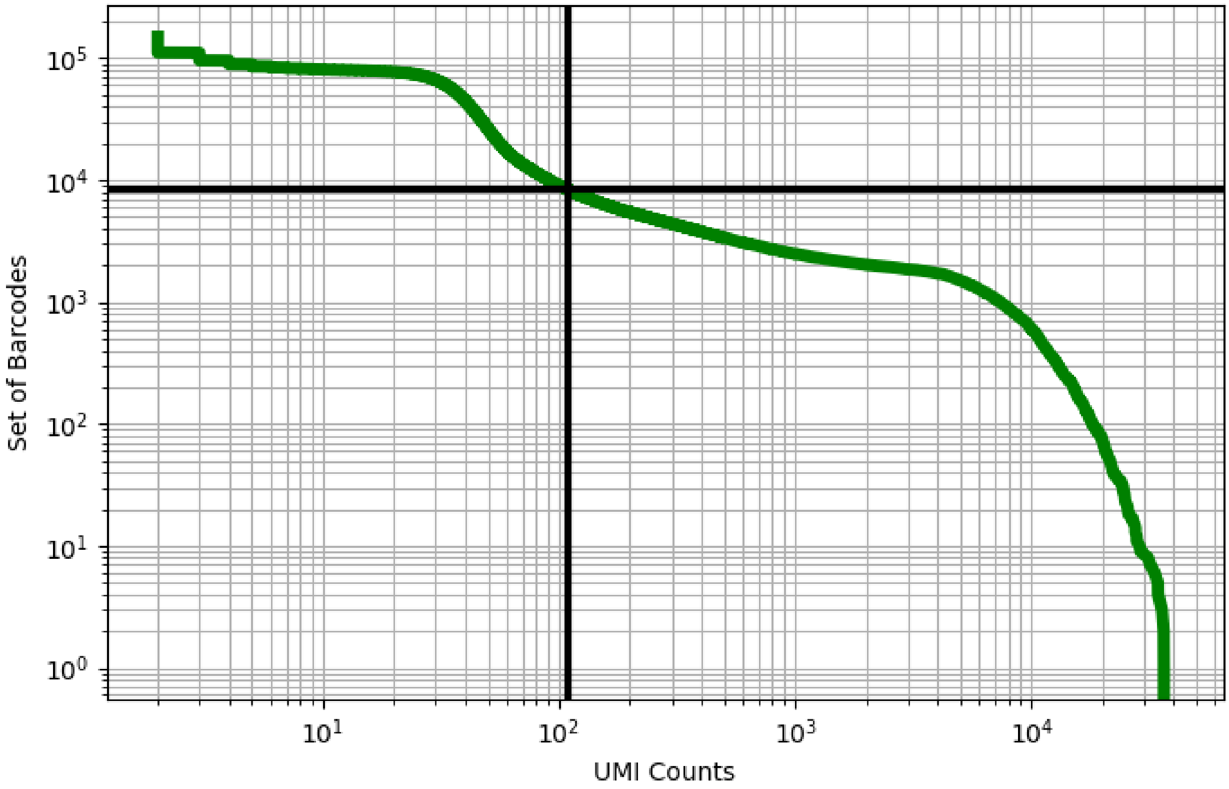


Figure 134: brain/GSM2905290

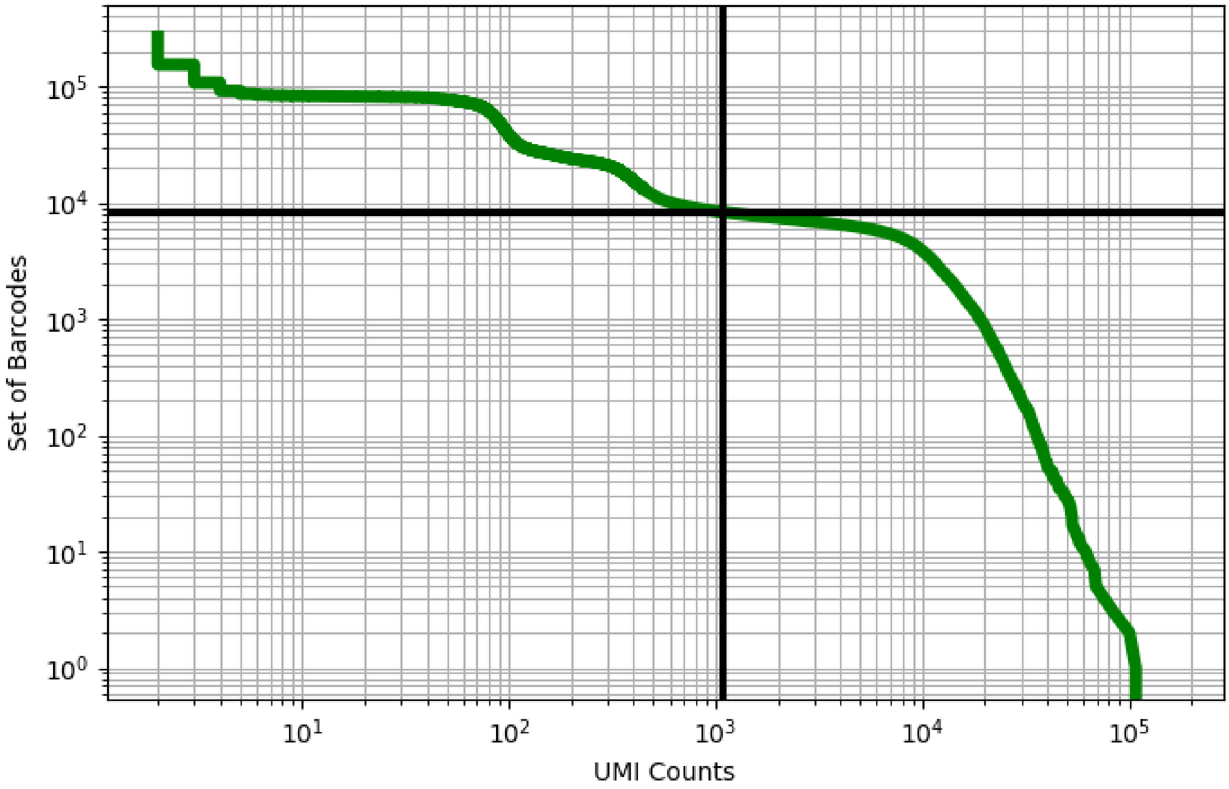


Figure 135: brain/GSM2905291

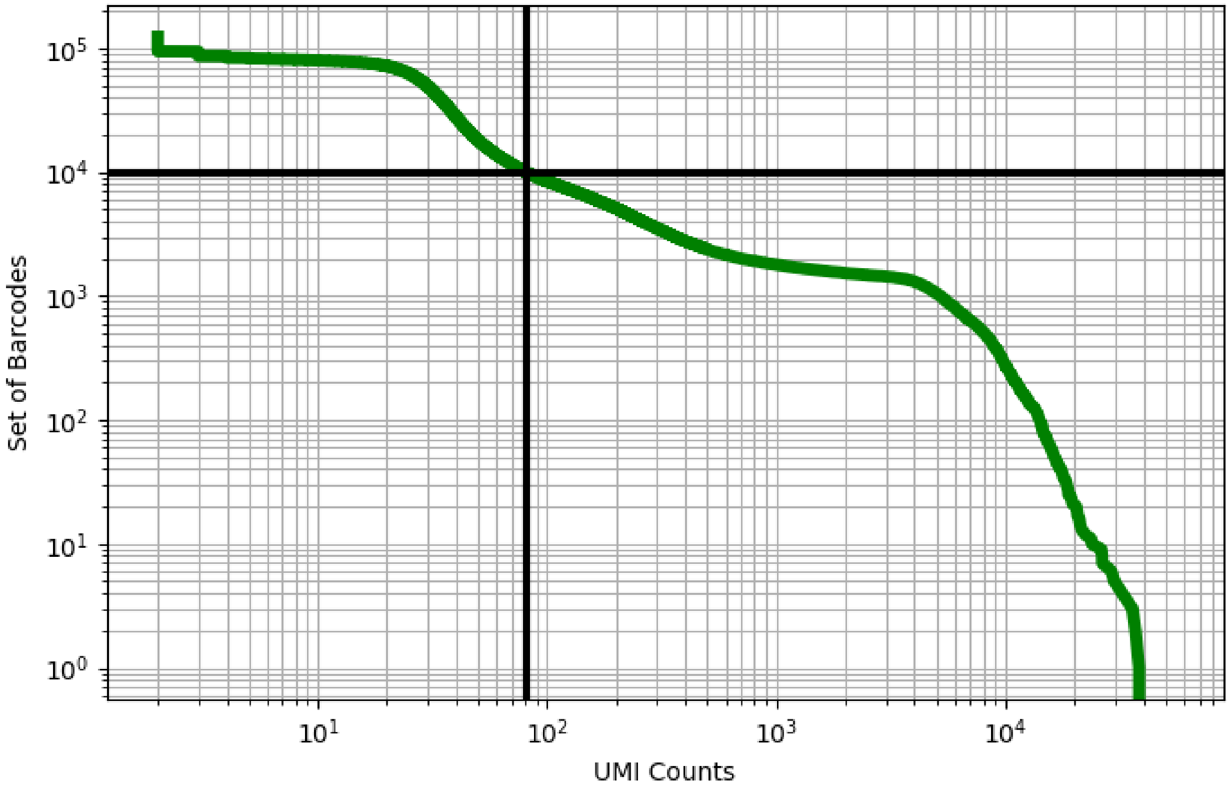


Figure 136: brain/GSM2905292

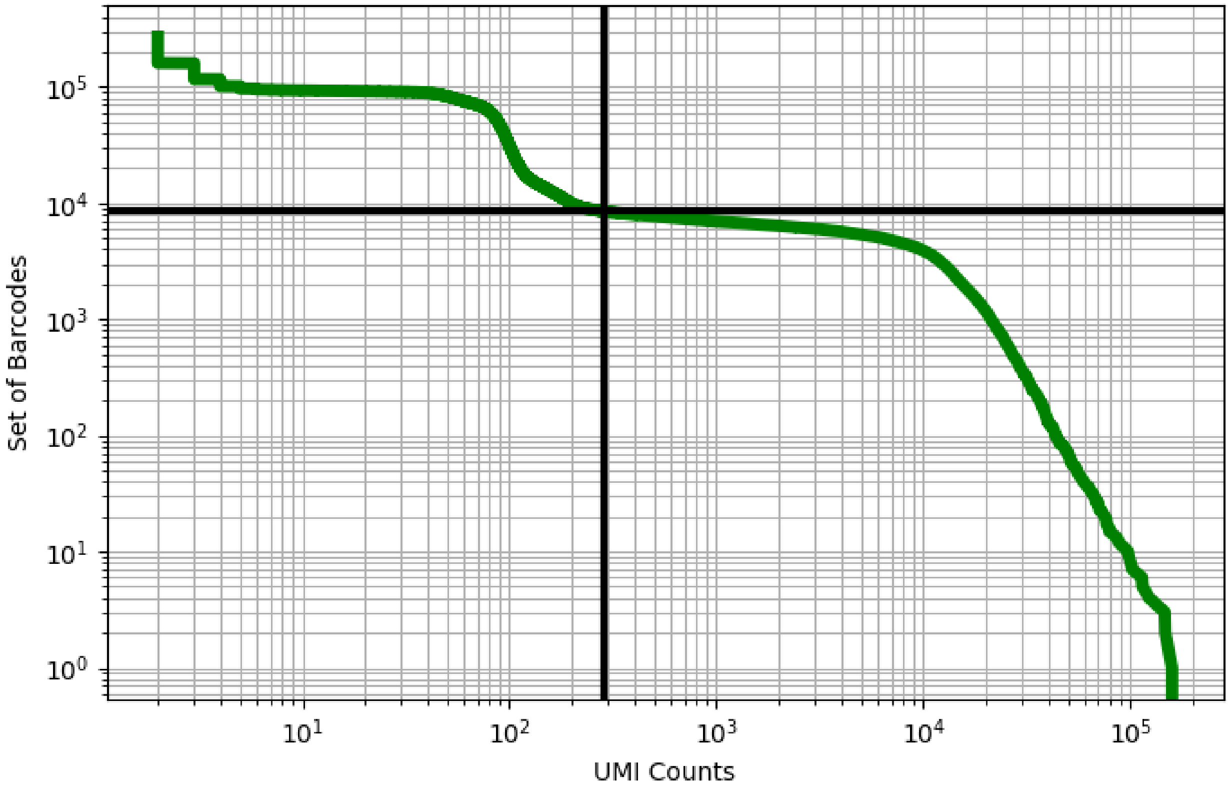


Figure 137: brain/GSM2905293

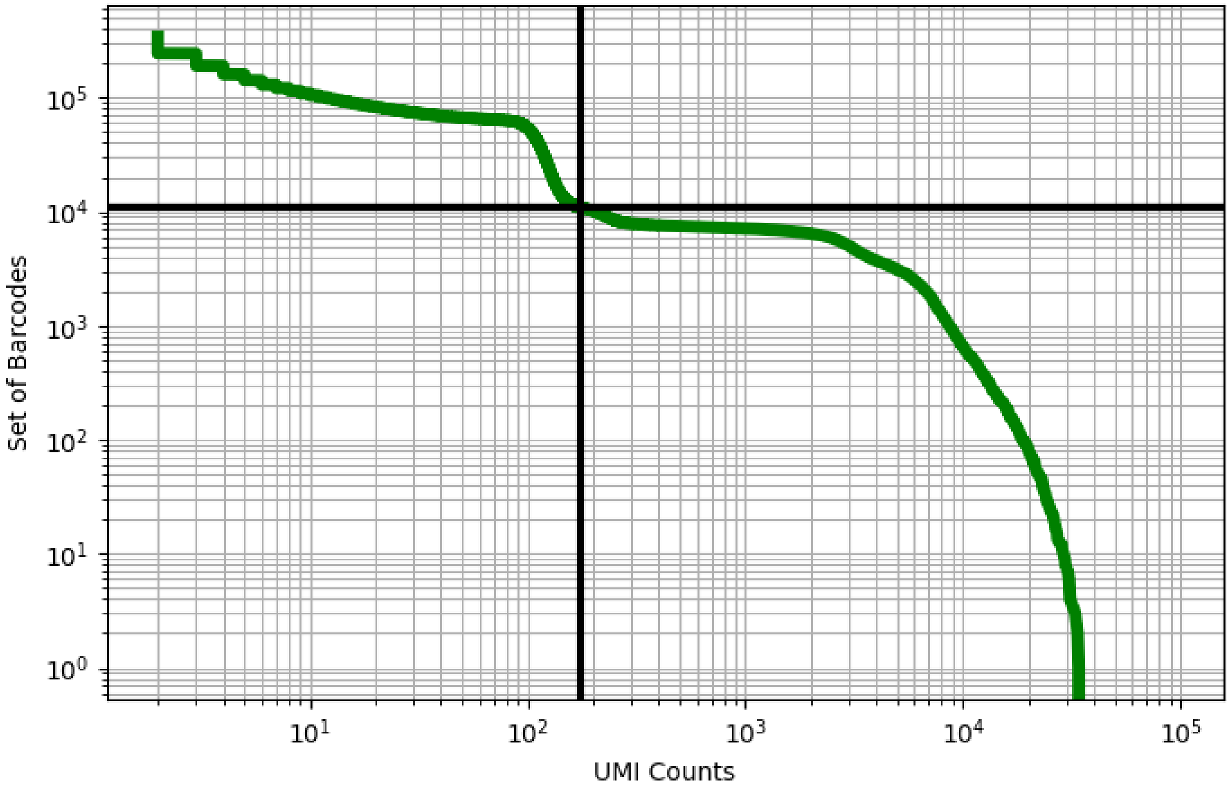


Figure 138: brain/GSM3360833

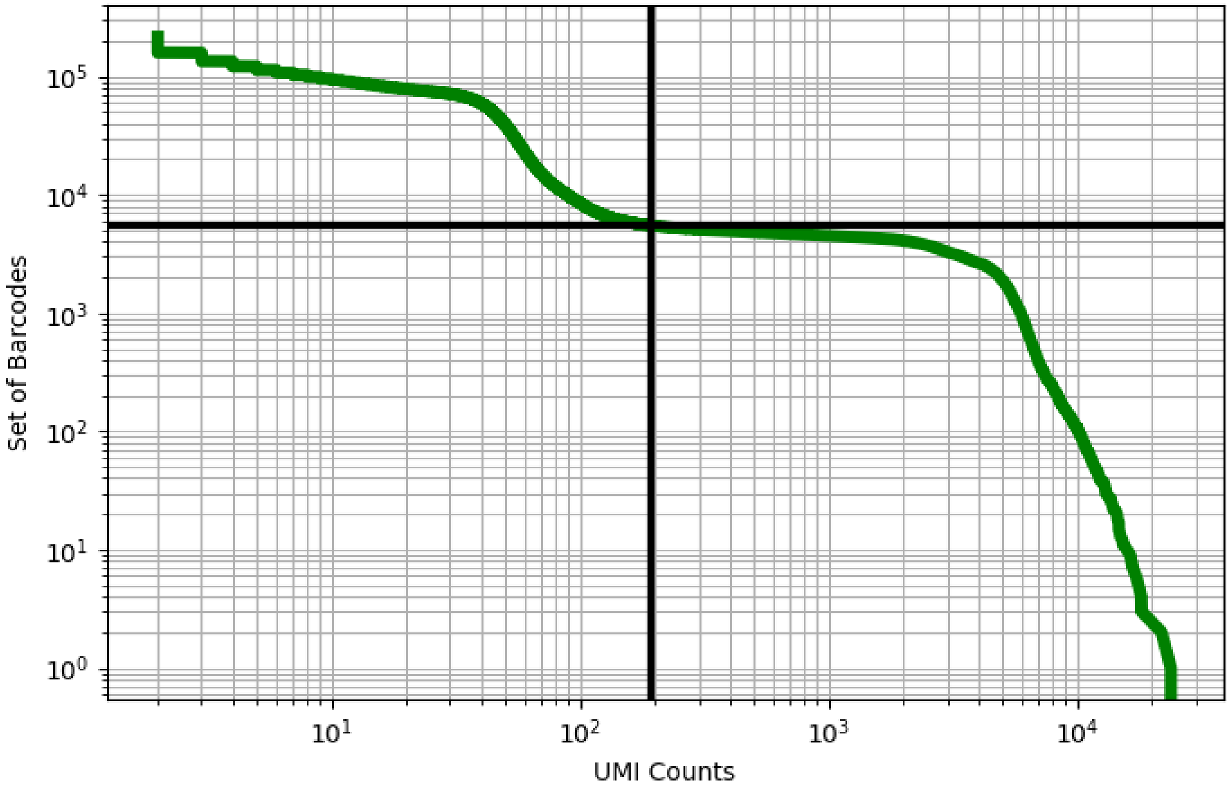


Figure 139: brain/GSM3360834

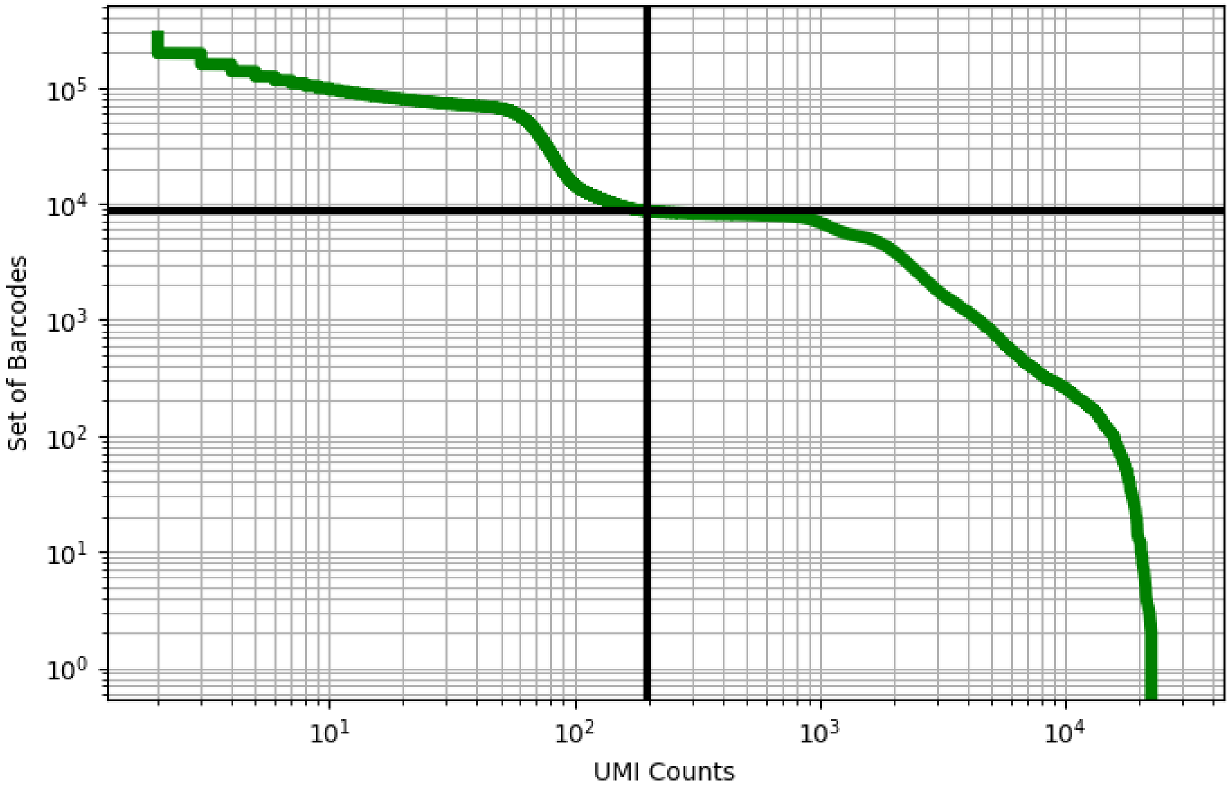


Figure 140: brain/GSM3360835

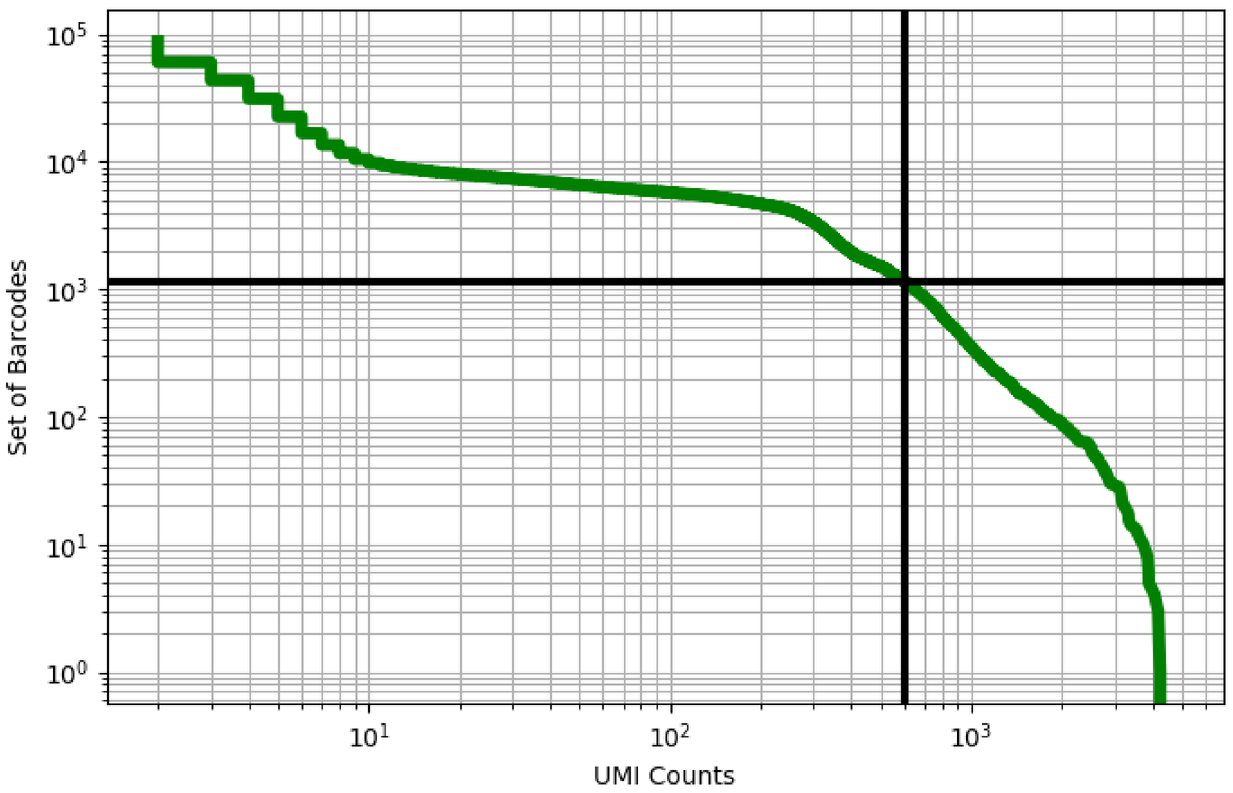


Figure 141: brain/GSM3360836

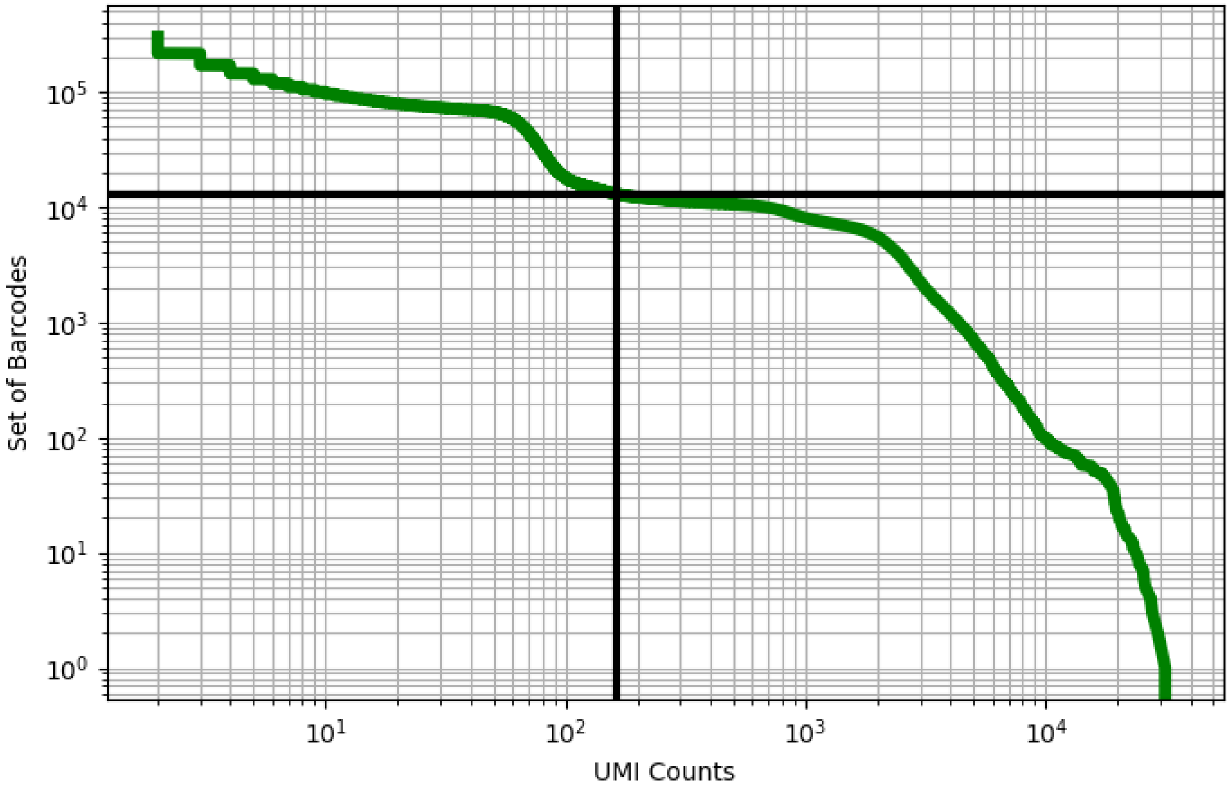


Figure 142: brain/GSM3360837

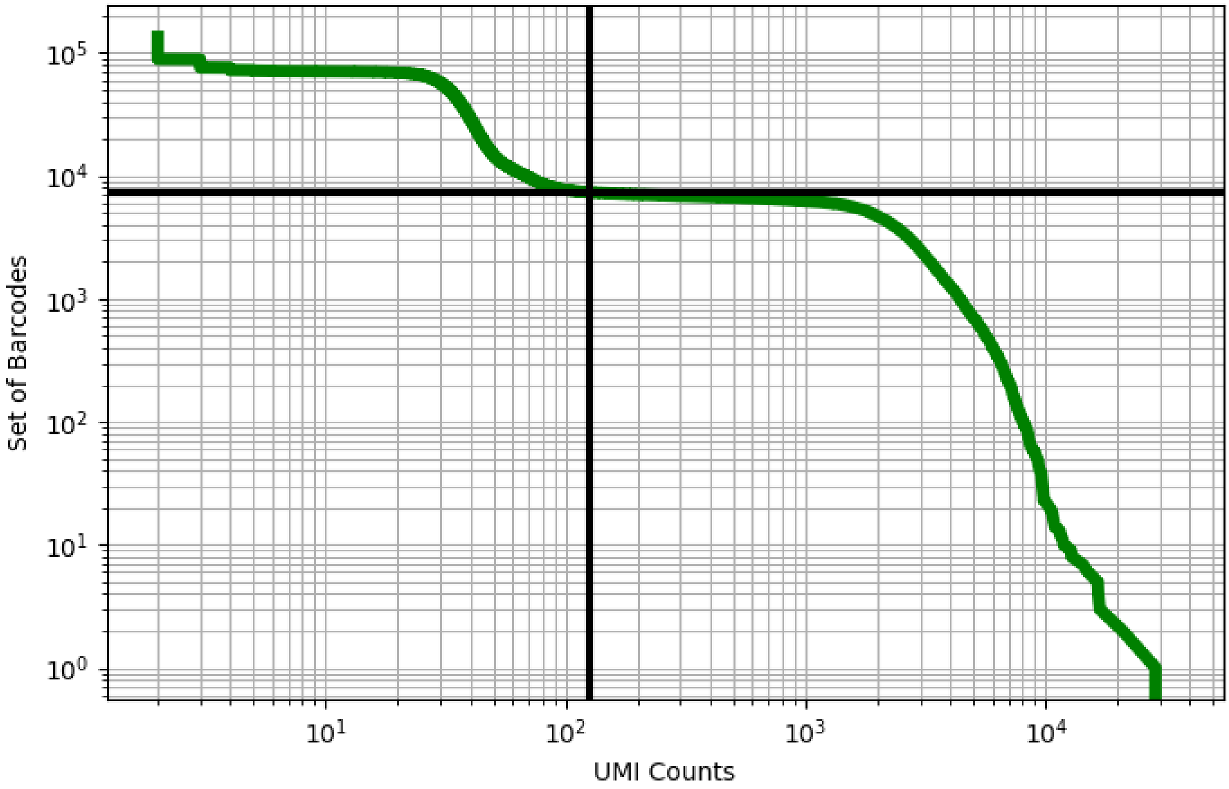


Figure 143: brain/GSM3360838

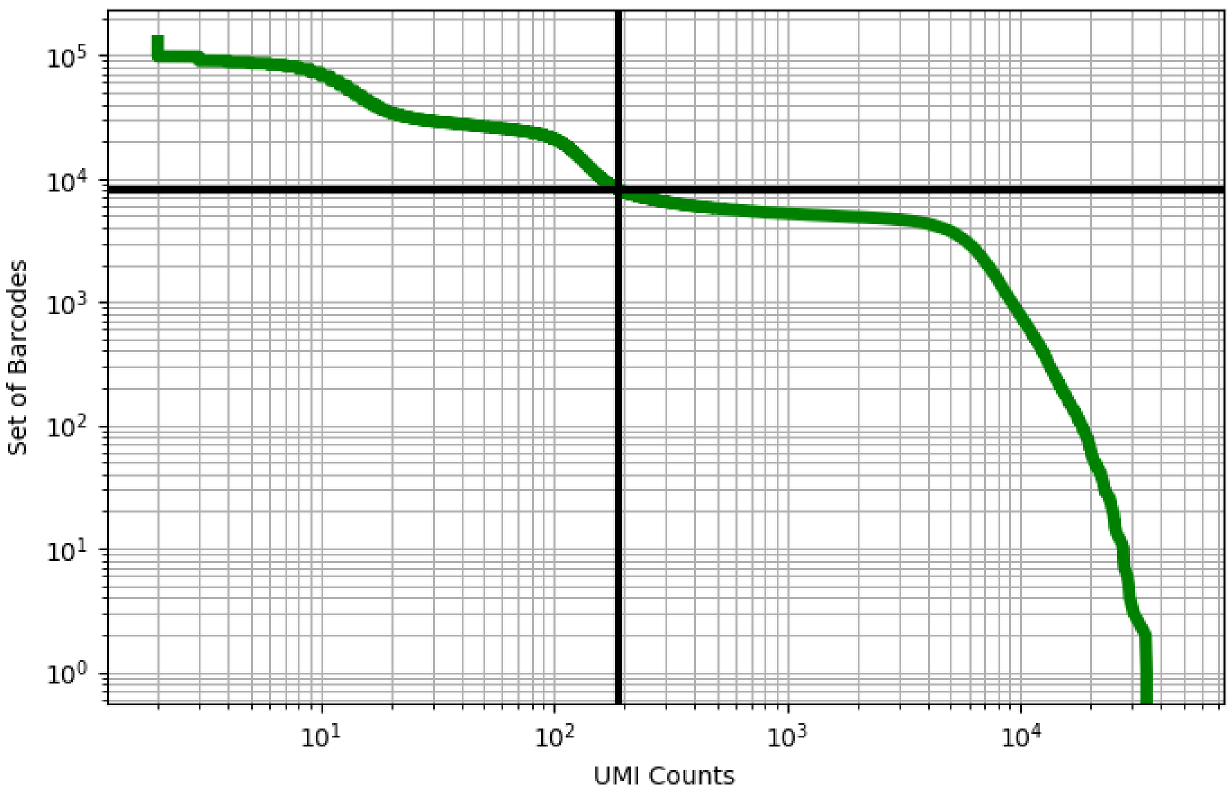


Figure 144: brain/GSM3734294

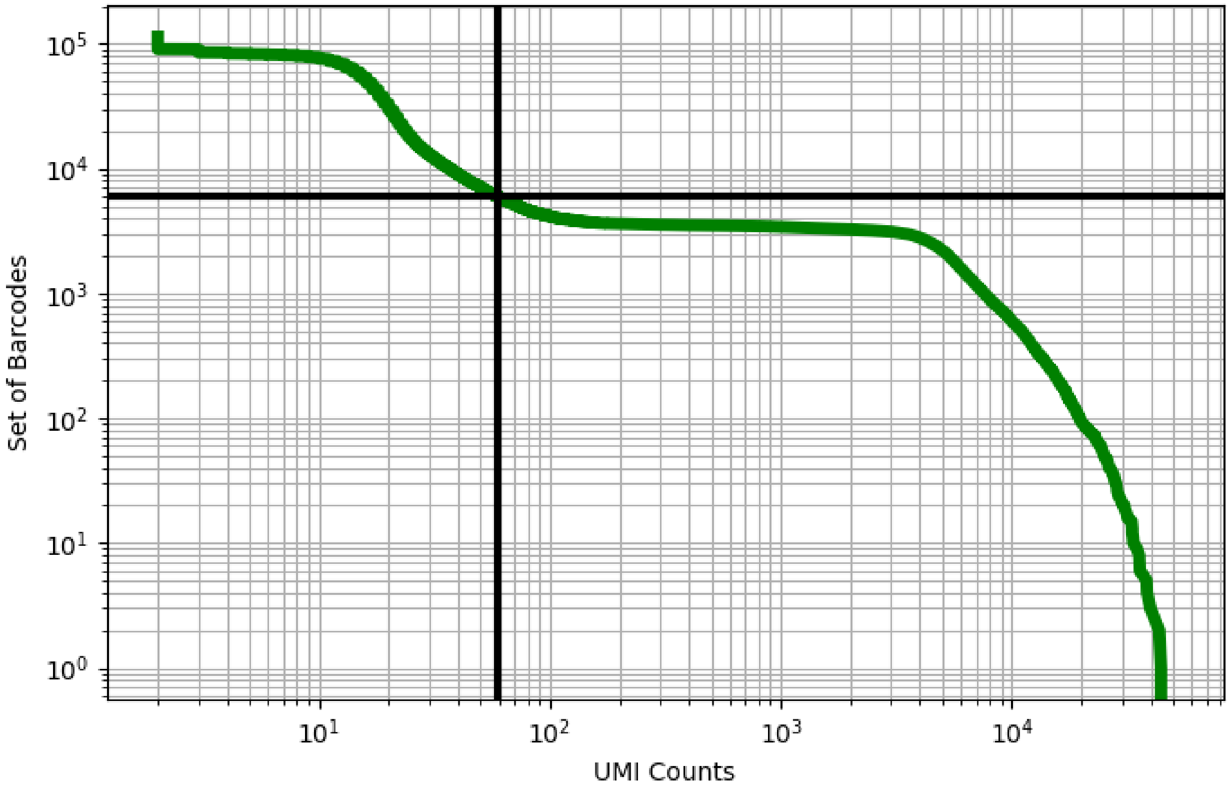


Figure 145: brain/GSM3734295

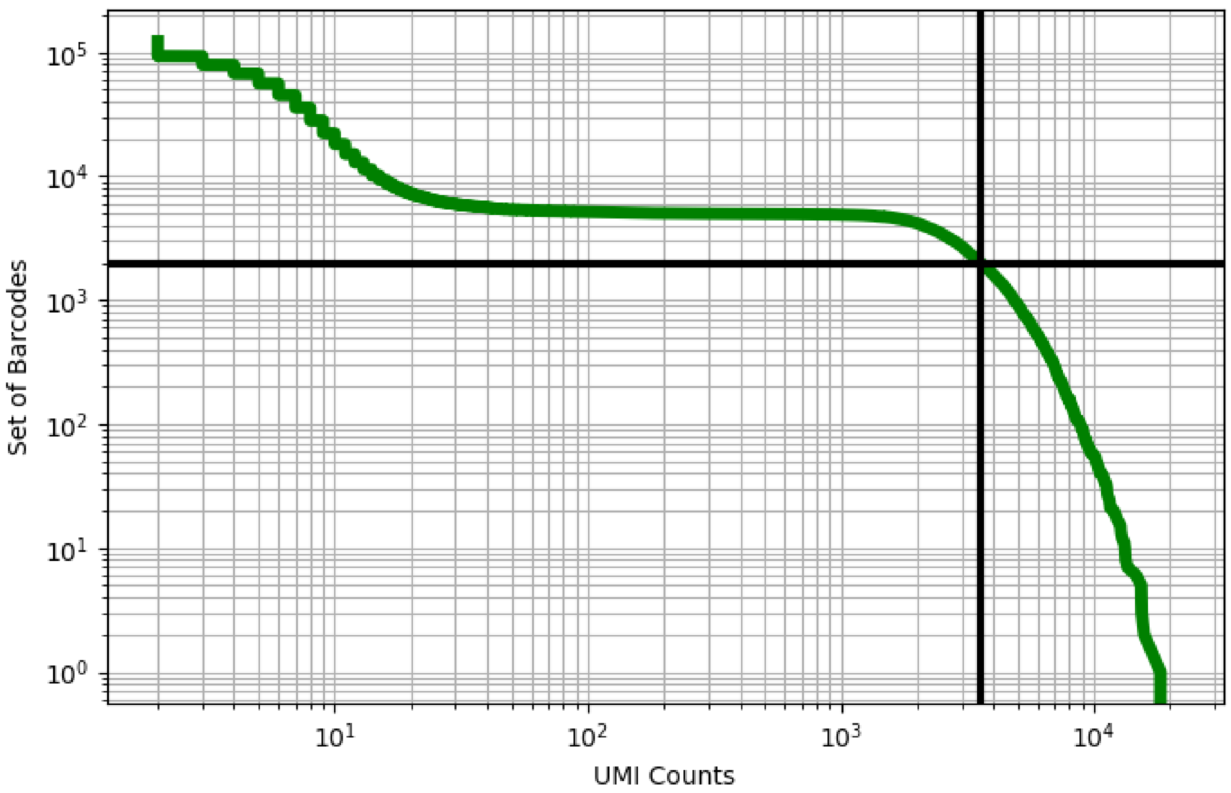


Figure 146: brain/GSM3734296

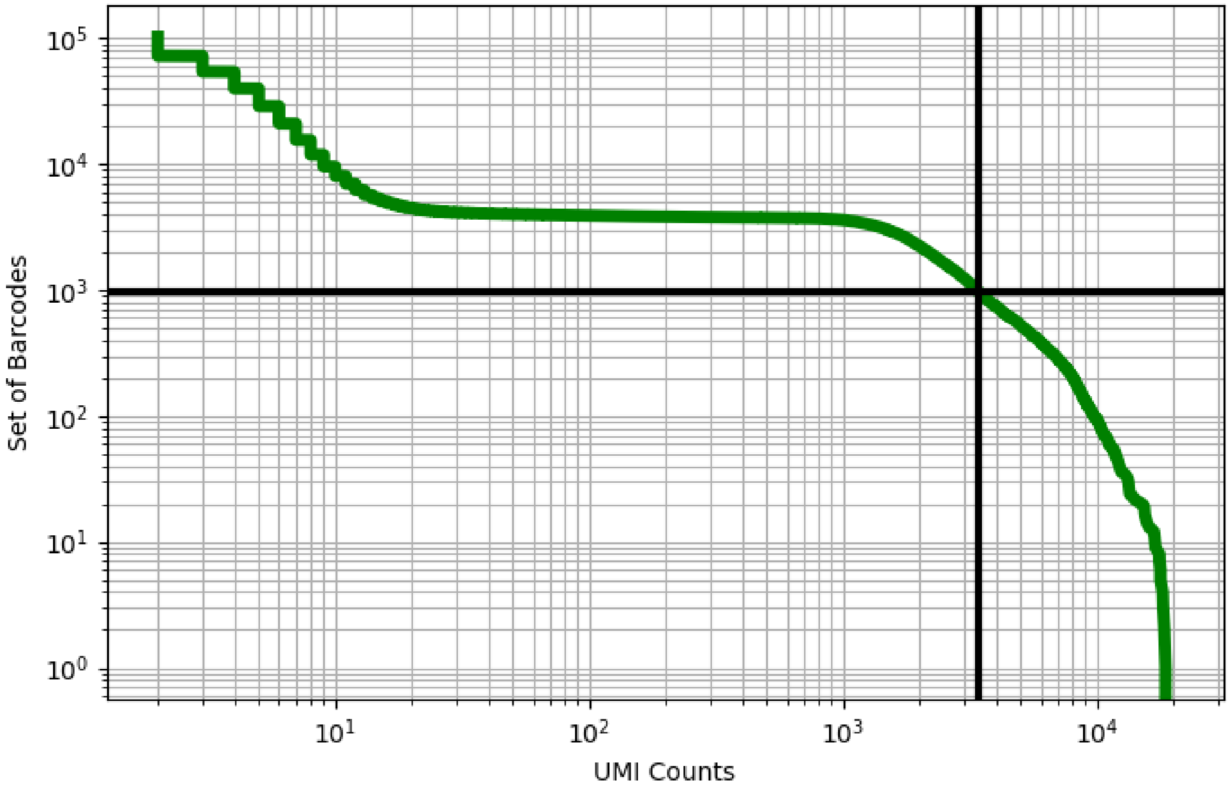


Figure 147: brain/GSM3734297

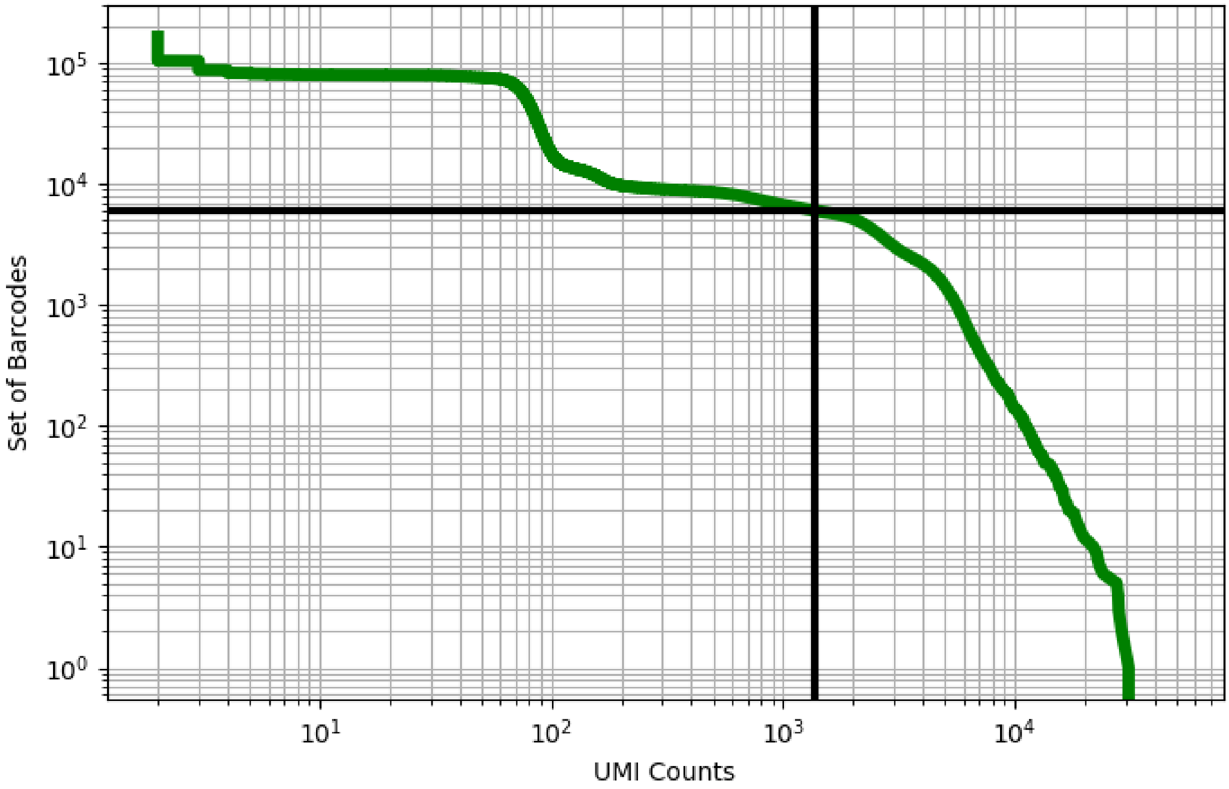


Figure 148: brain/GSM3770749

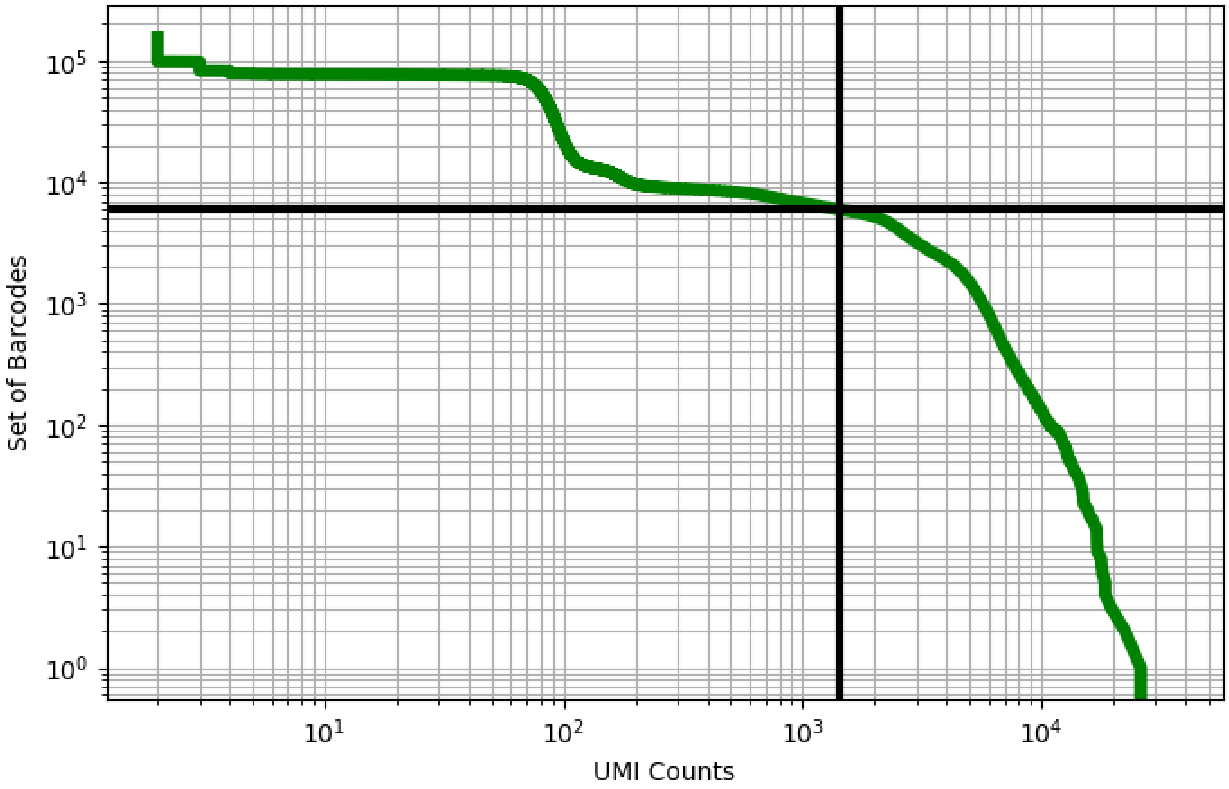


Figure 149: brain/GSM3770750

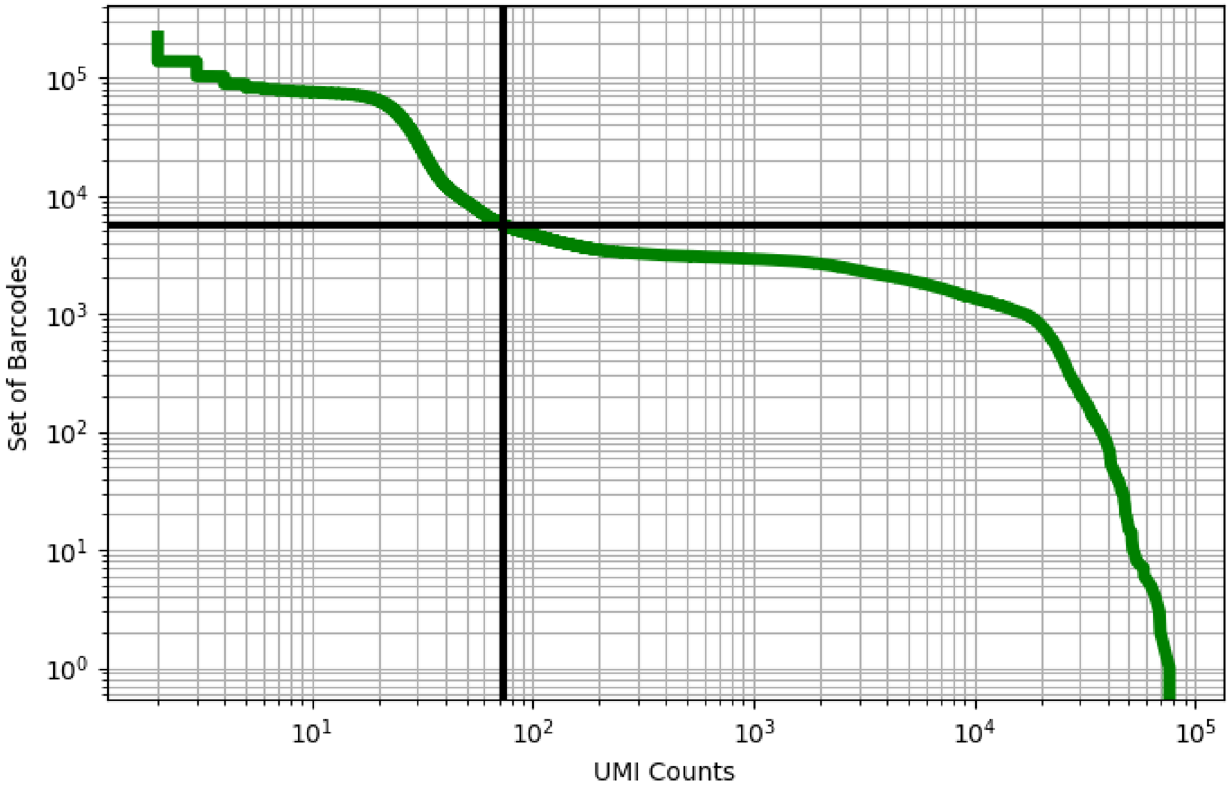


Figure 150: colon/GSM3587010

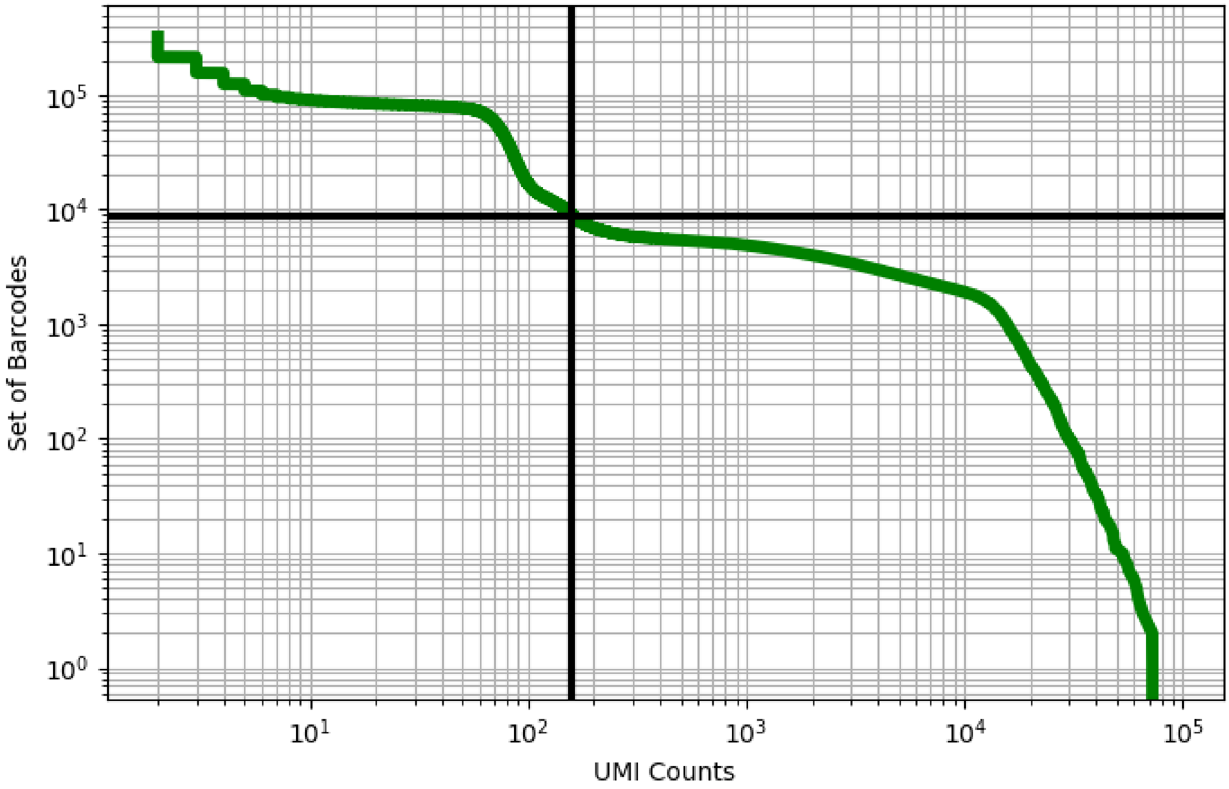


Figure 151: colon/GSM3587011

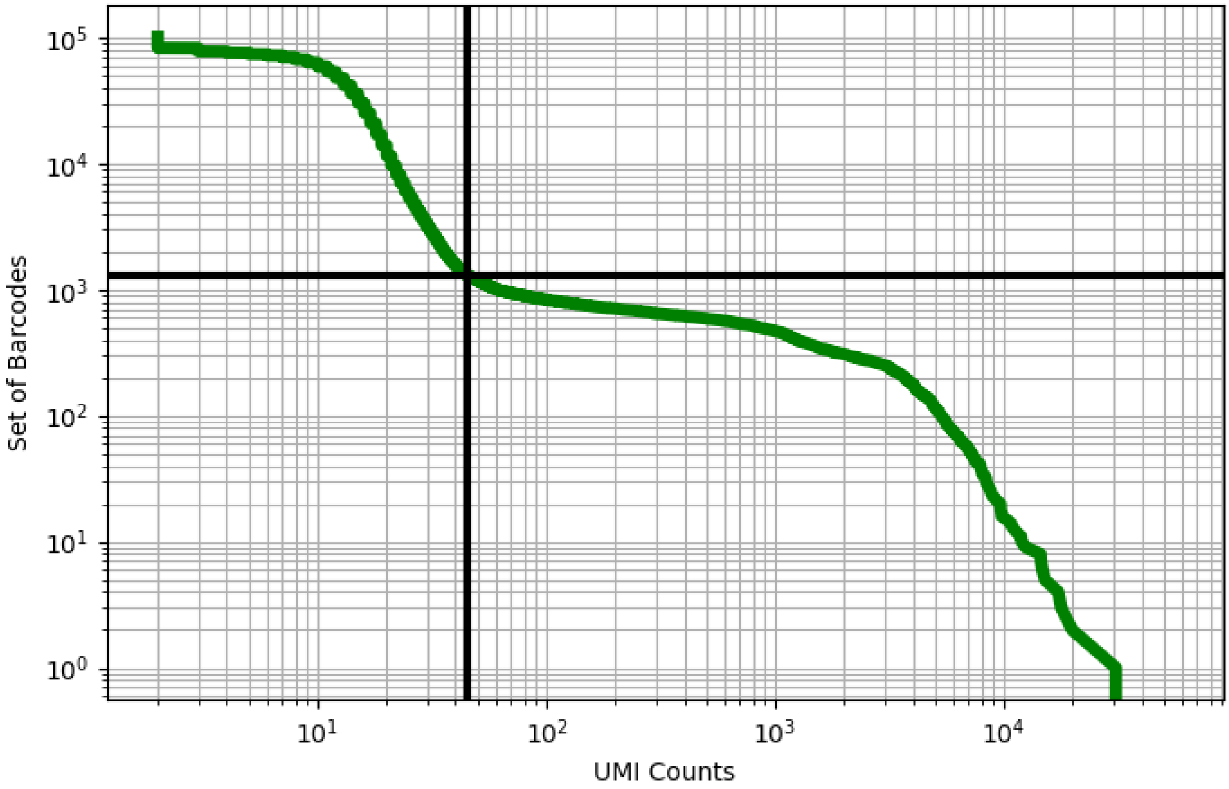


Figure 152: decidua/ERX2756708

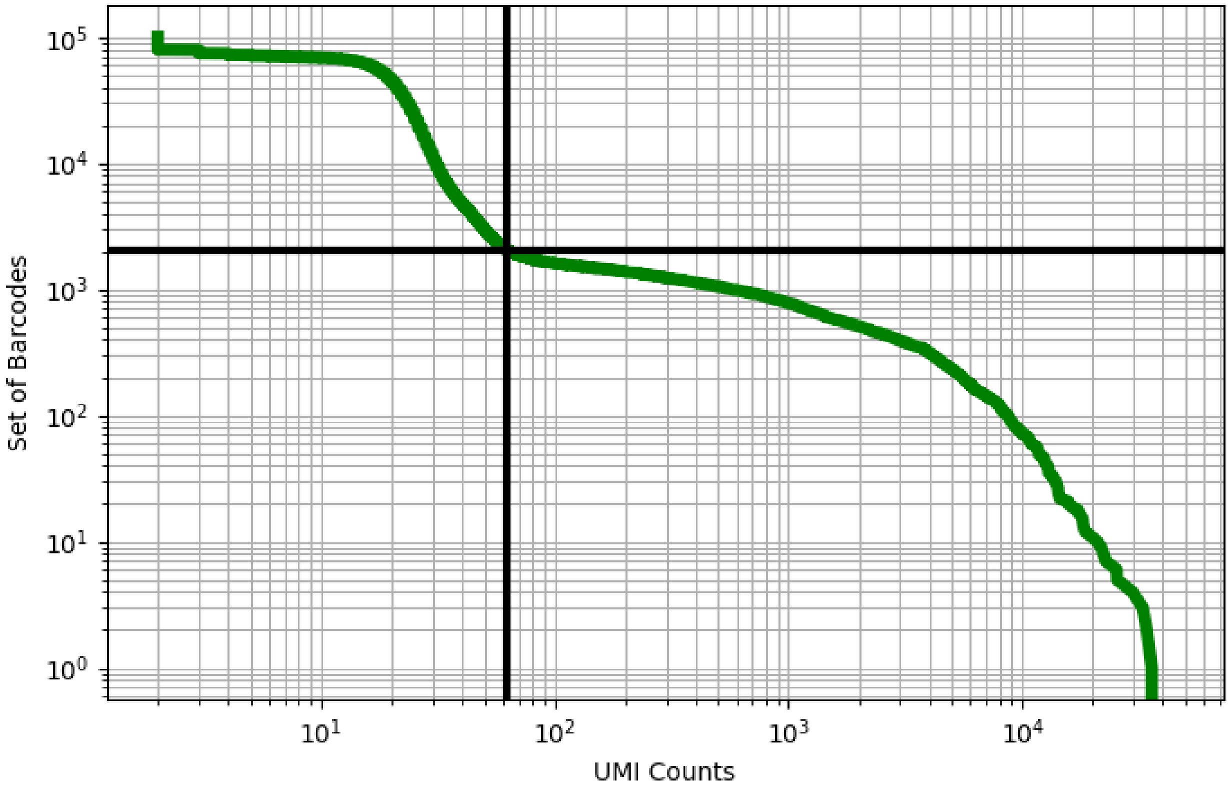


Figure 153: decidua/ERX2756710

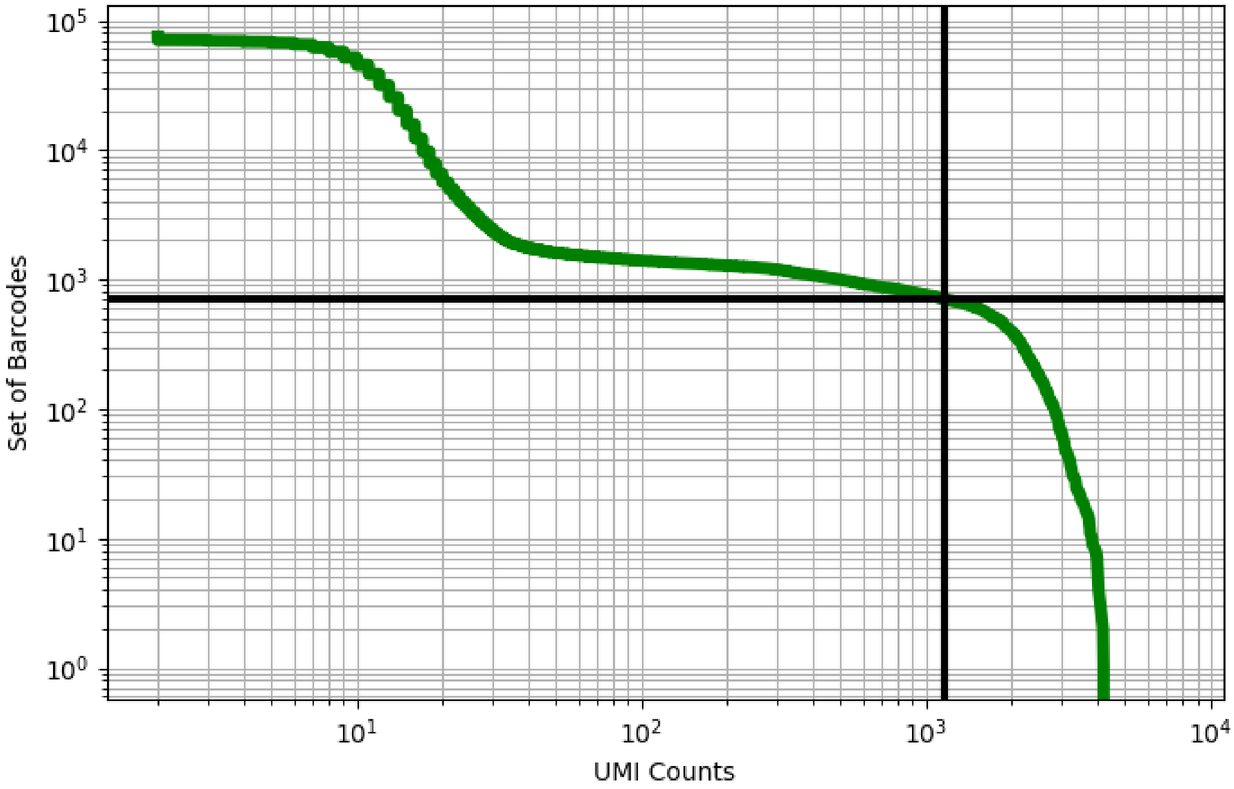


Figure 154: decidua/ERX2756711

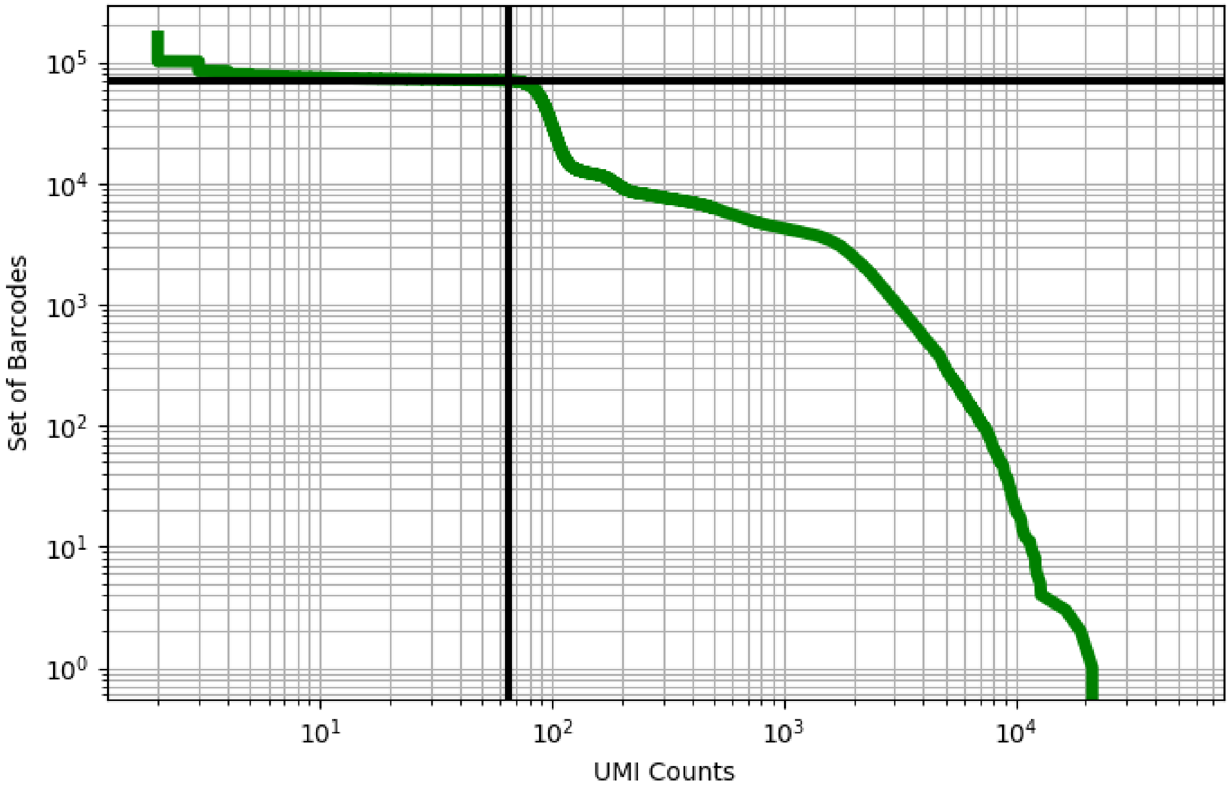


Figure 155: decidua/ERX2756712

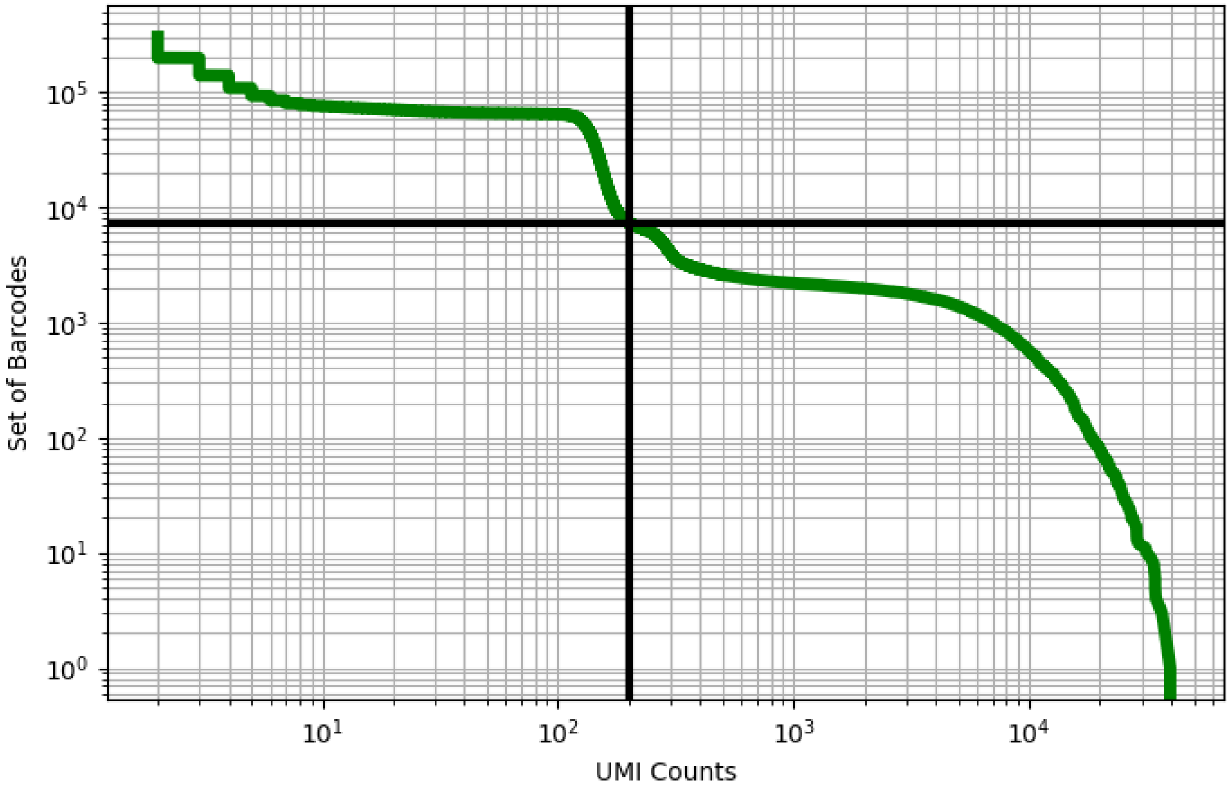


Figure 156: decidua/ERX2756713

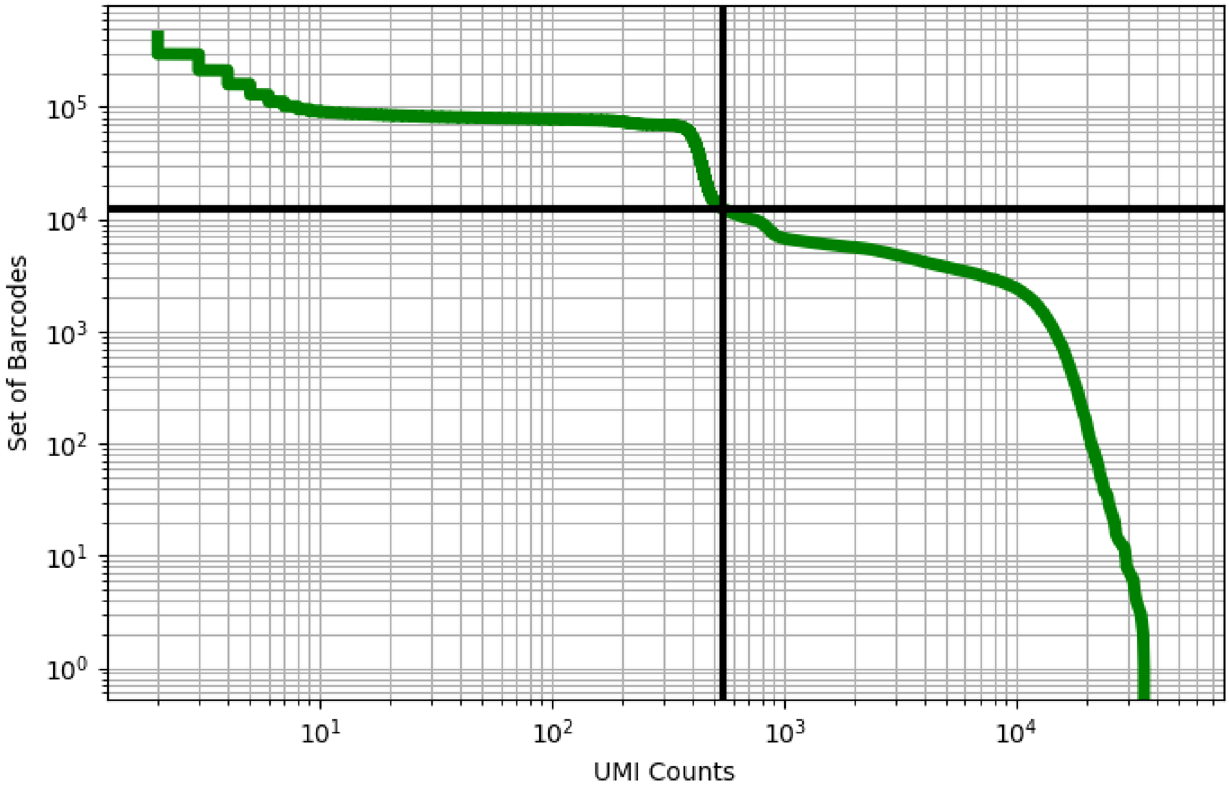


Figure 157: decidua/ERX2756714

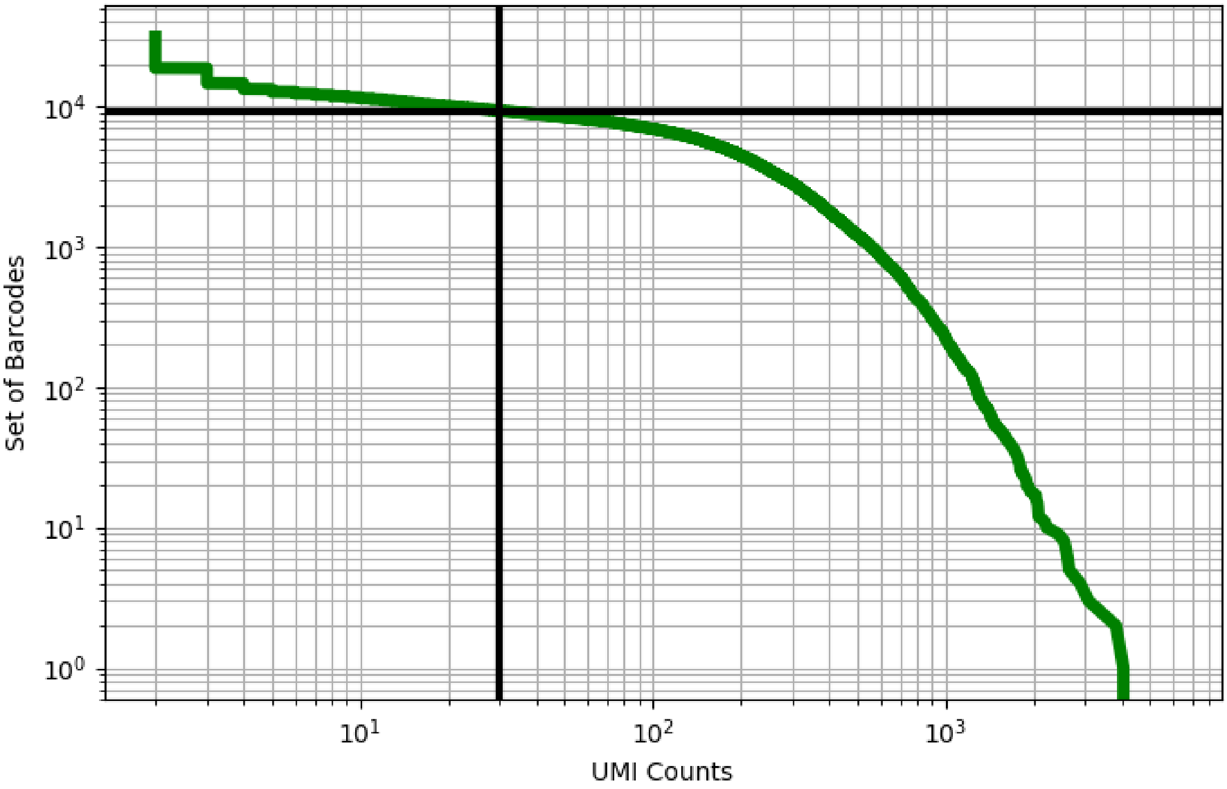


Figure 158: decidua/ERX2756721

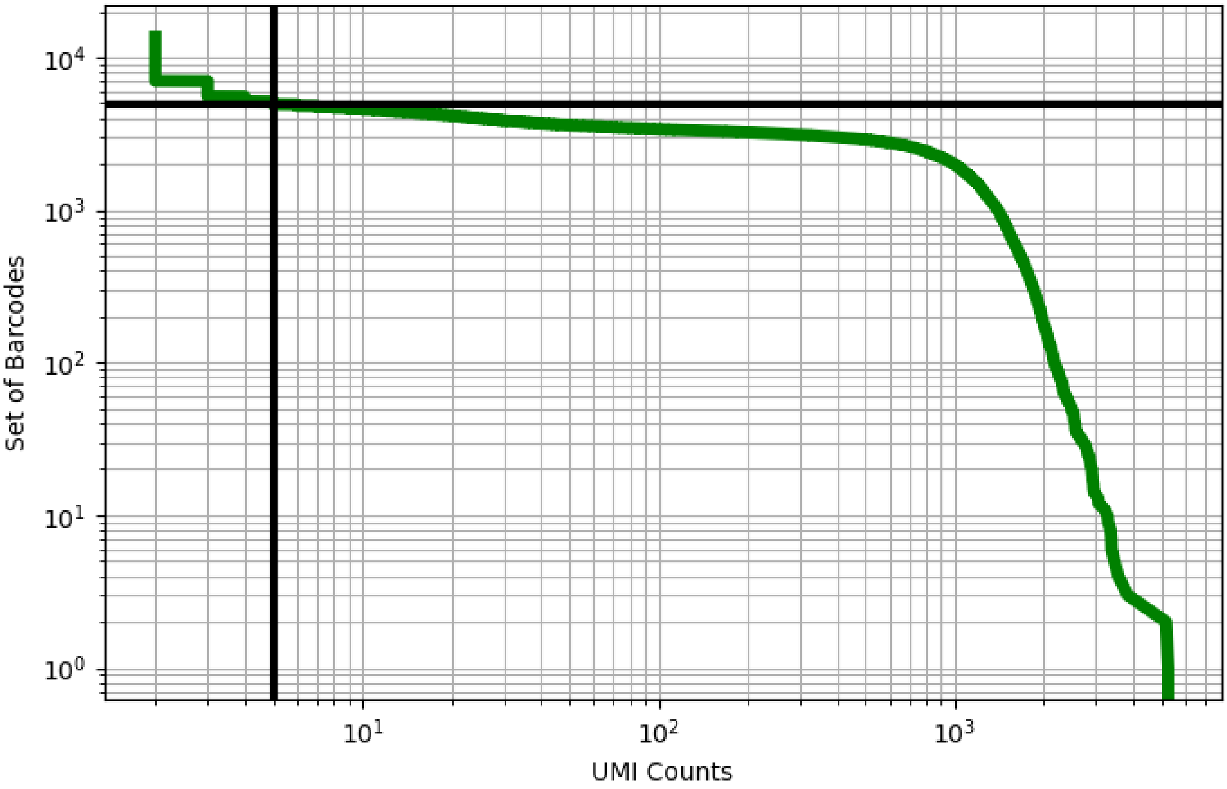


Figure 159: decidua/ERX2756722

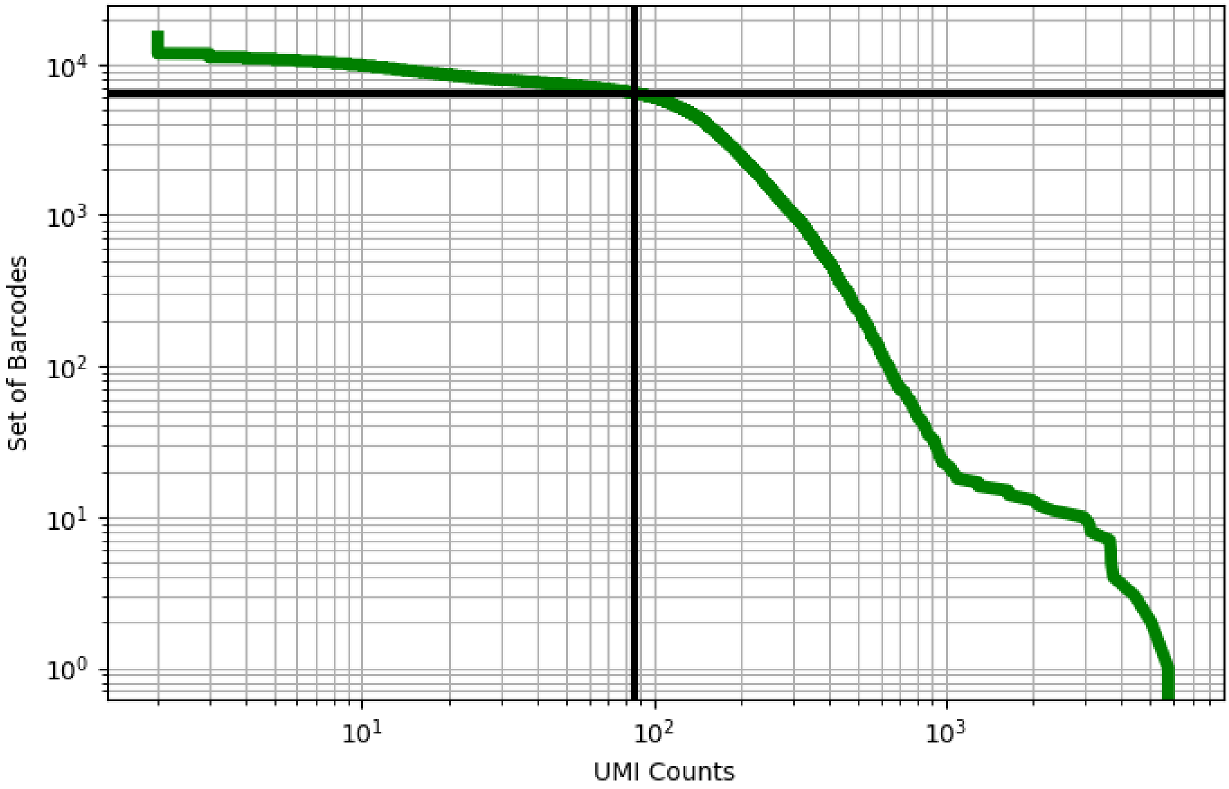


Figure 160: decidua/ERX2756724

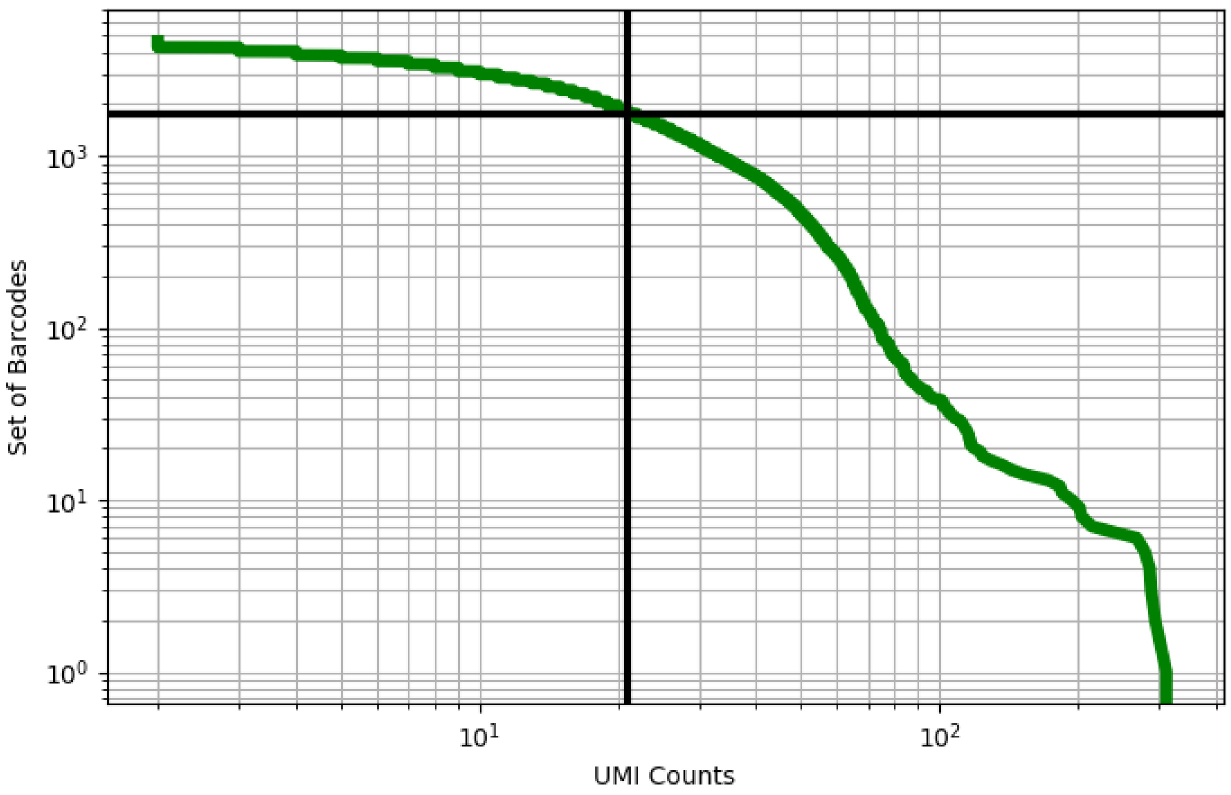


Figure 161: decidua/ERX2756725

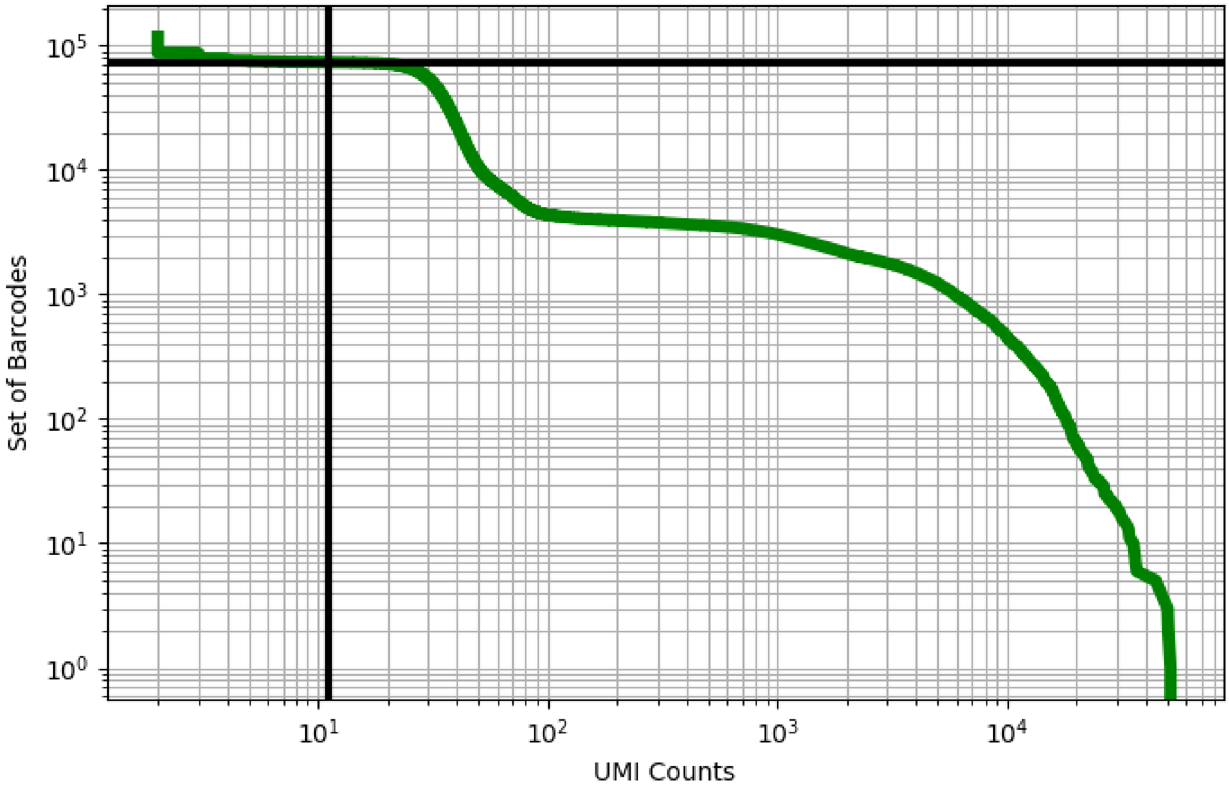


Figure 162: decidua/ERX2756729

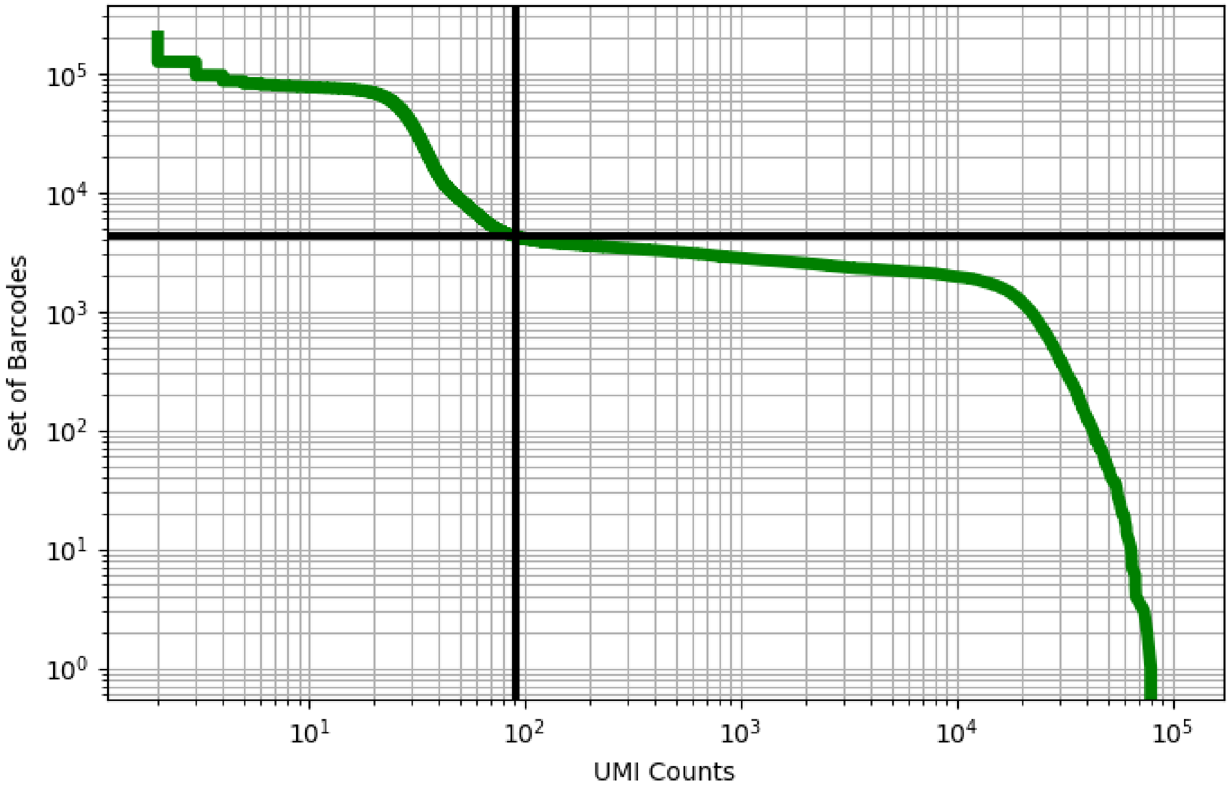


Figure 163: decidua/ERX2756730

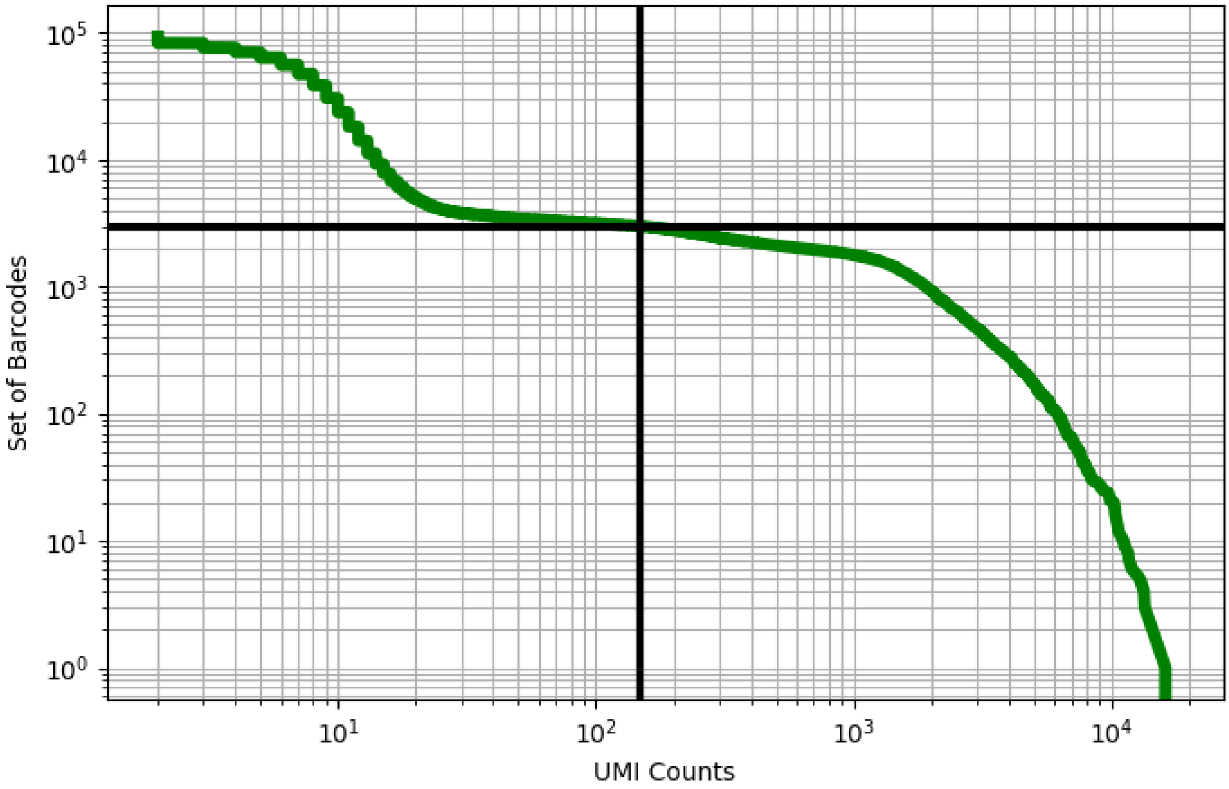


Figure 164: decidua/ERX2756737

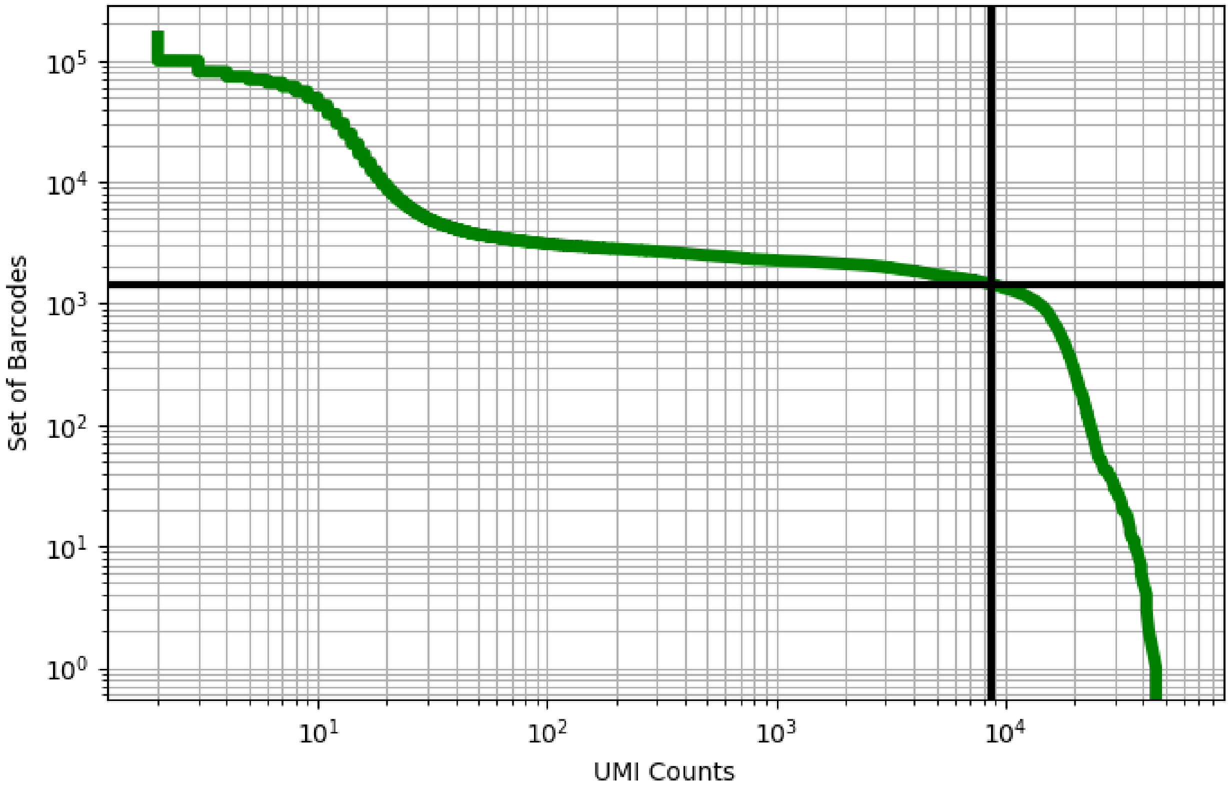


Figure 165: decidua/ERX2756738

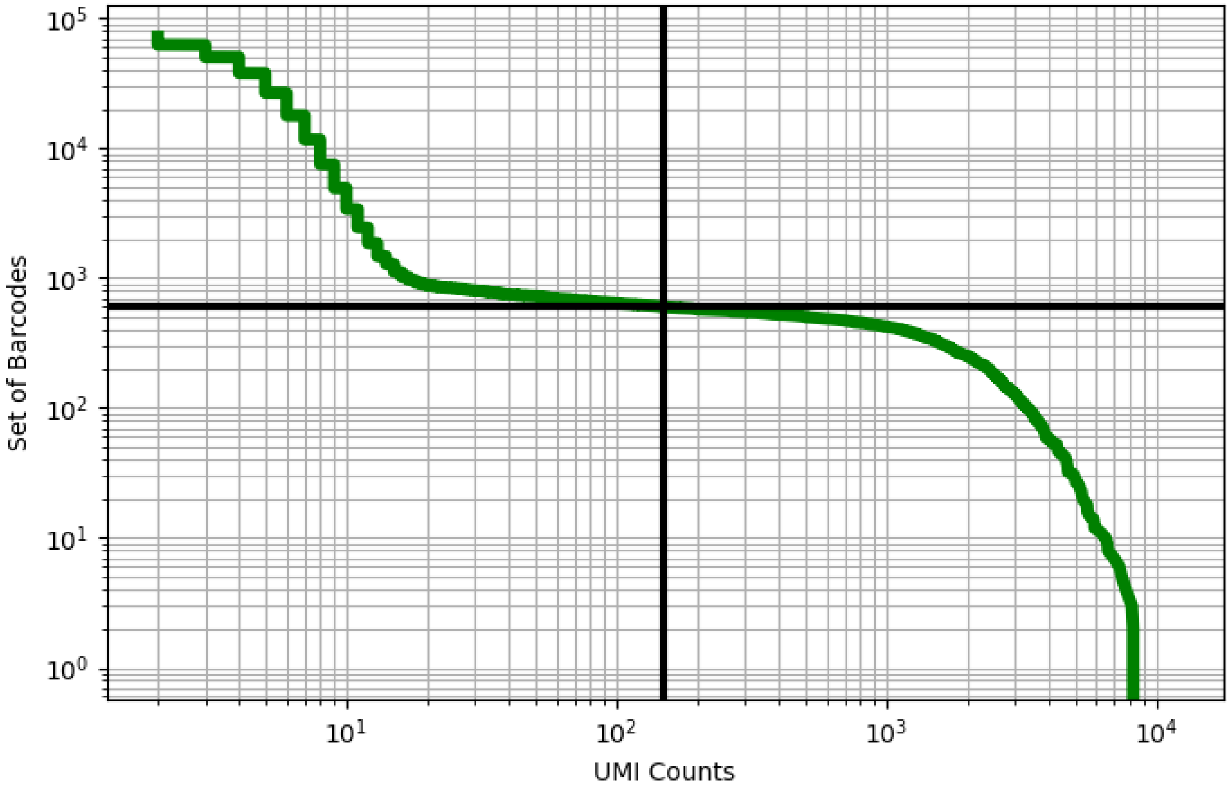


Figure 166: decidua/ERX2756739

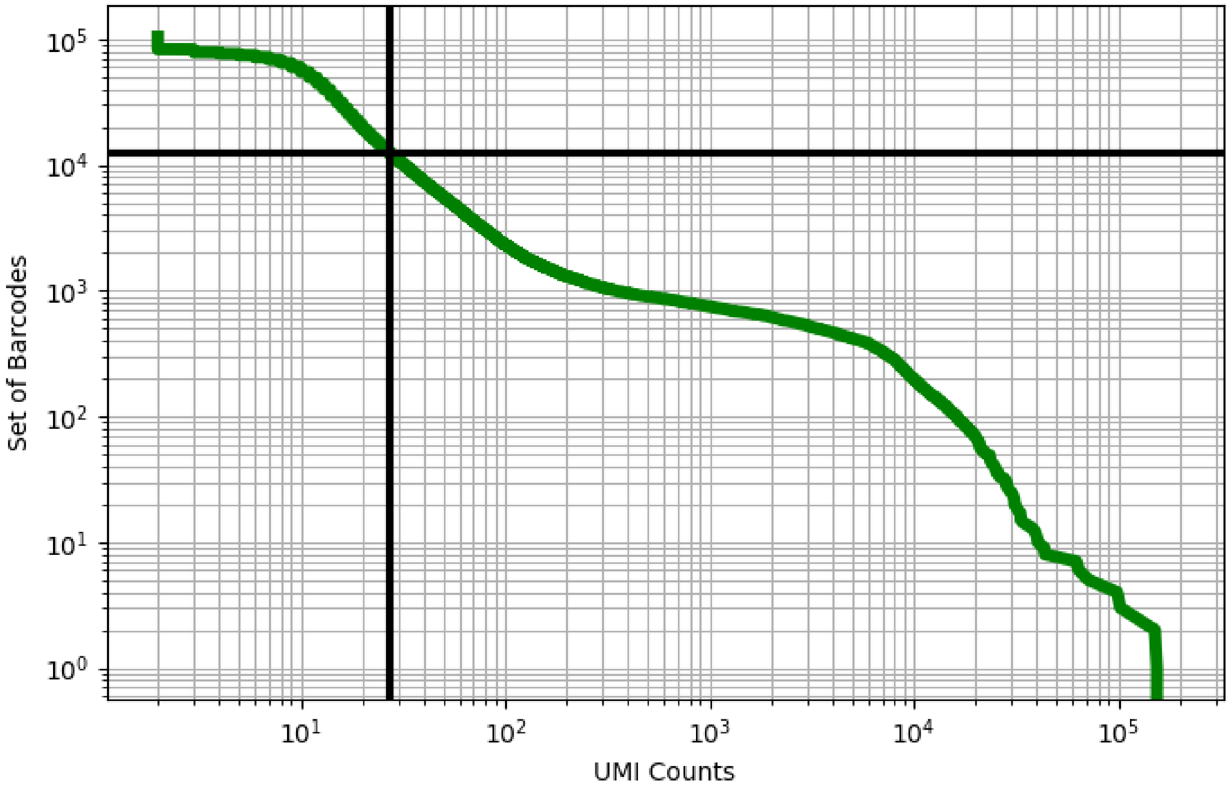


Figure 167: eye/GSM4037981

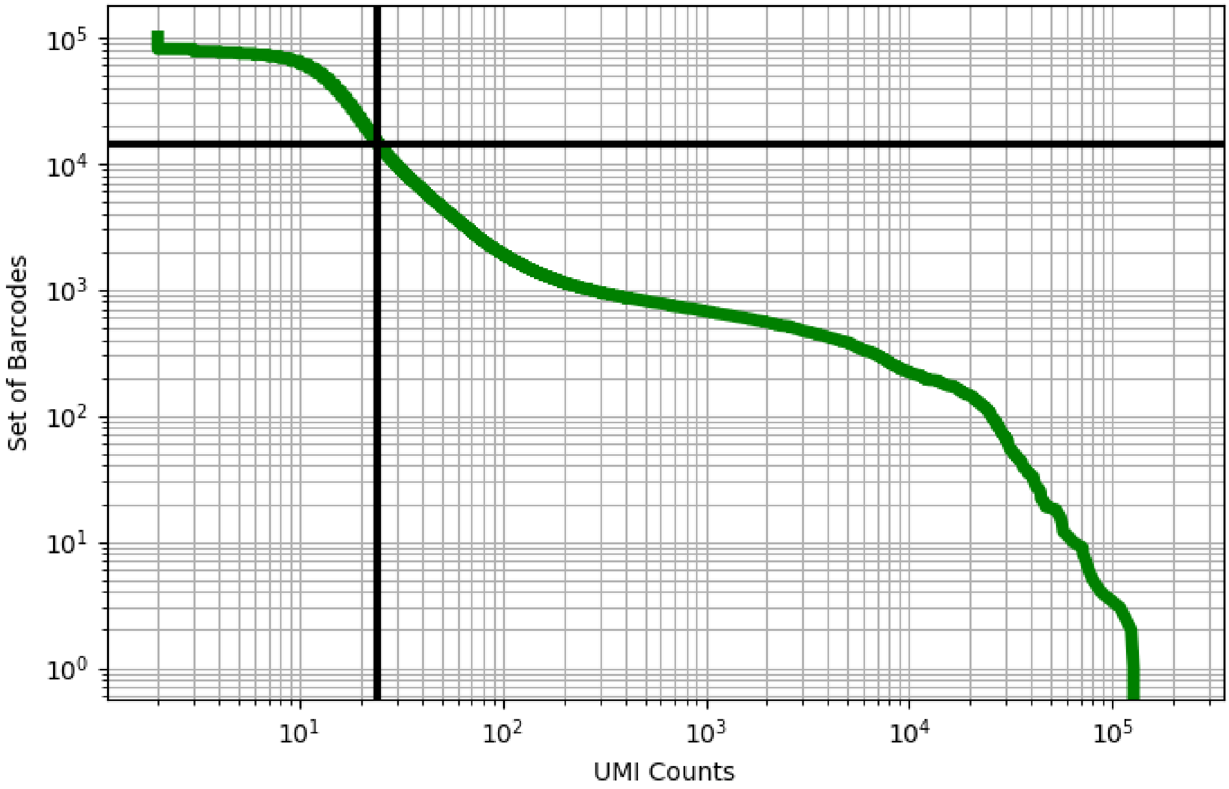


Figure 168: eye/GSM4037982

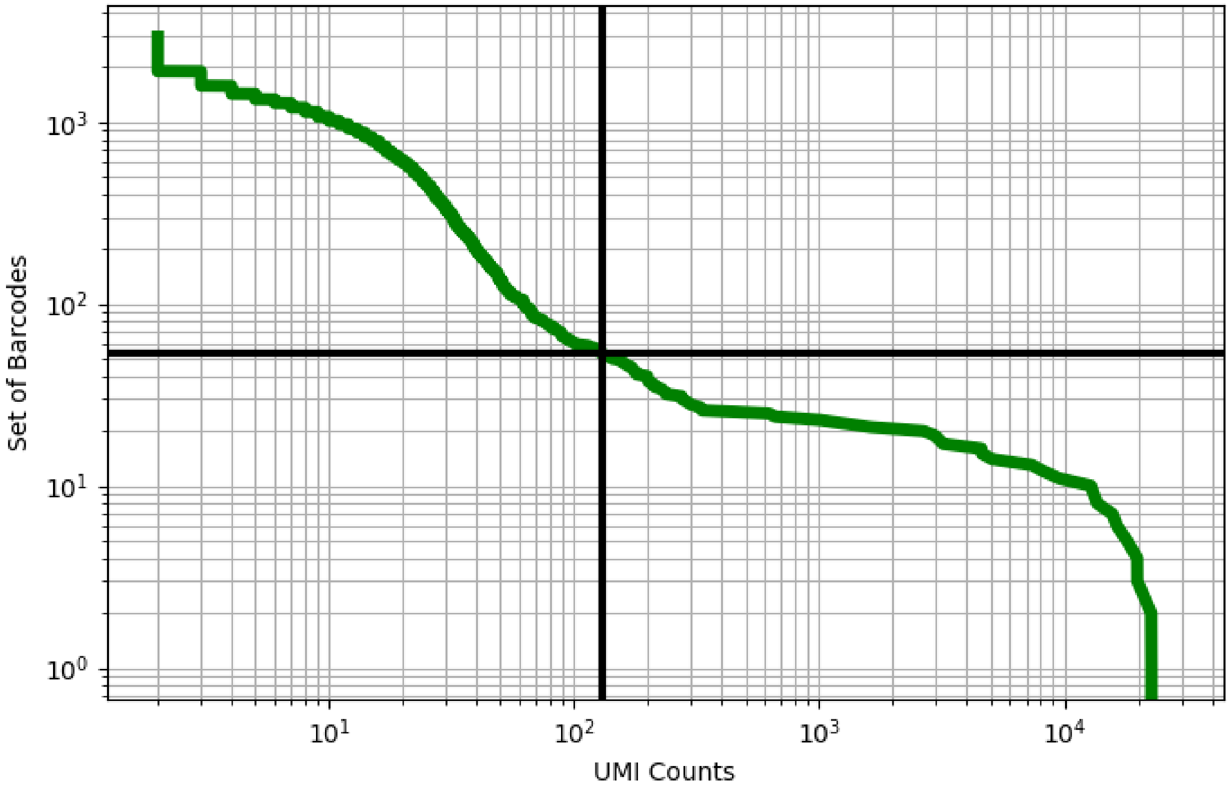


Figure 169: eye/GSM4037984

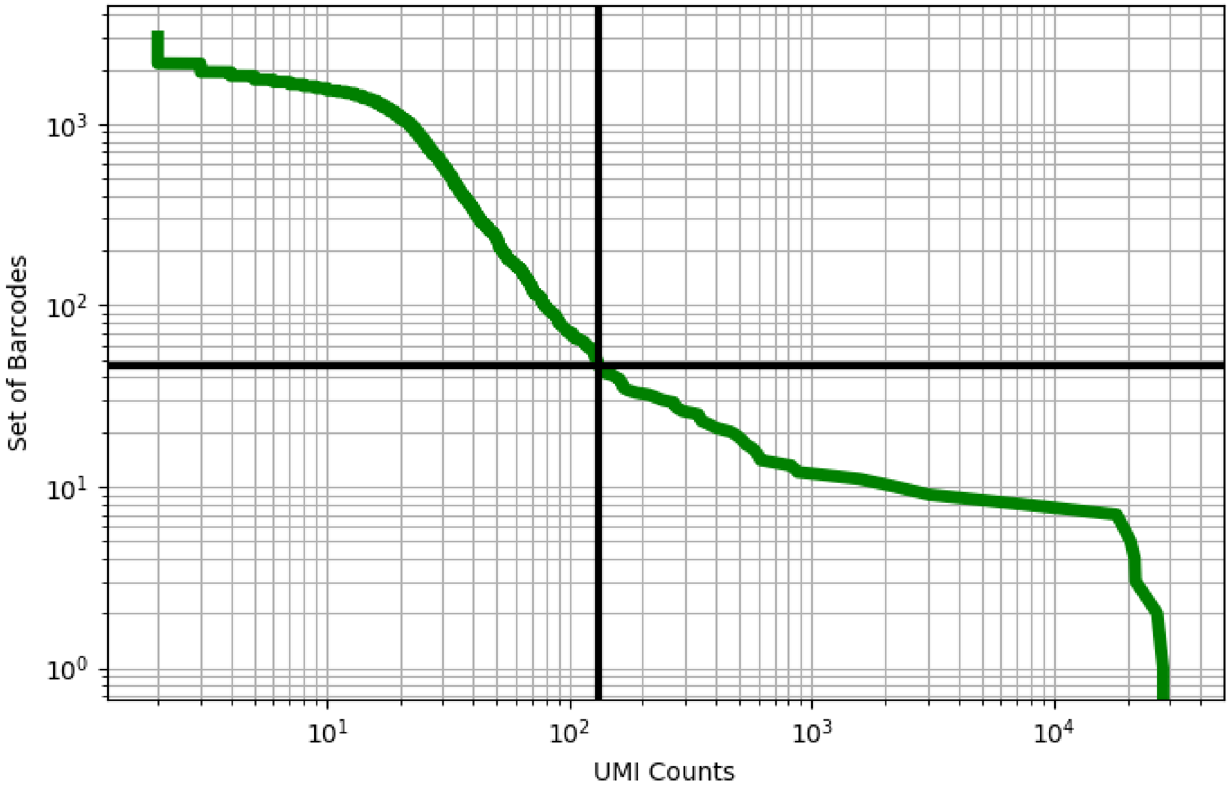


Figure 170: eye/GSM4037985

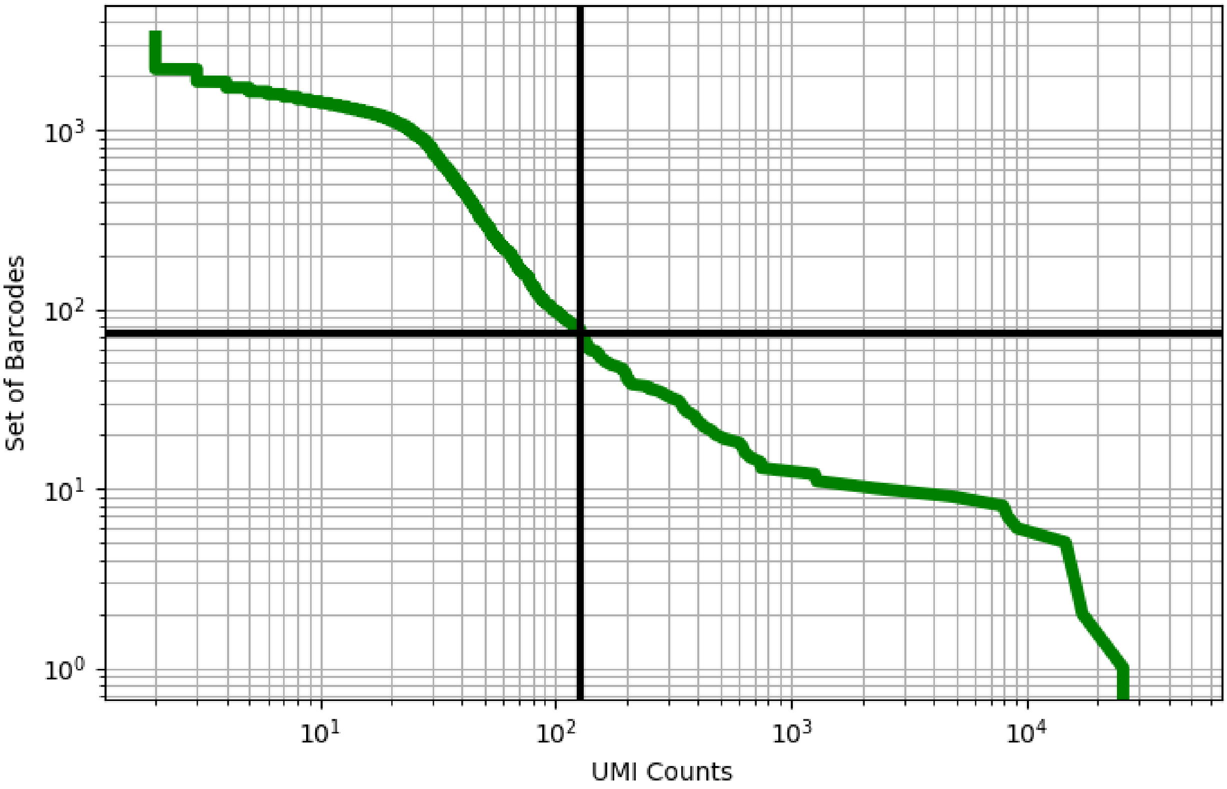


Figure 171: eye/GSM4037986

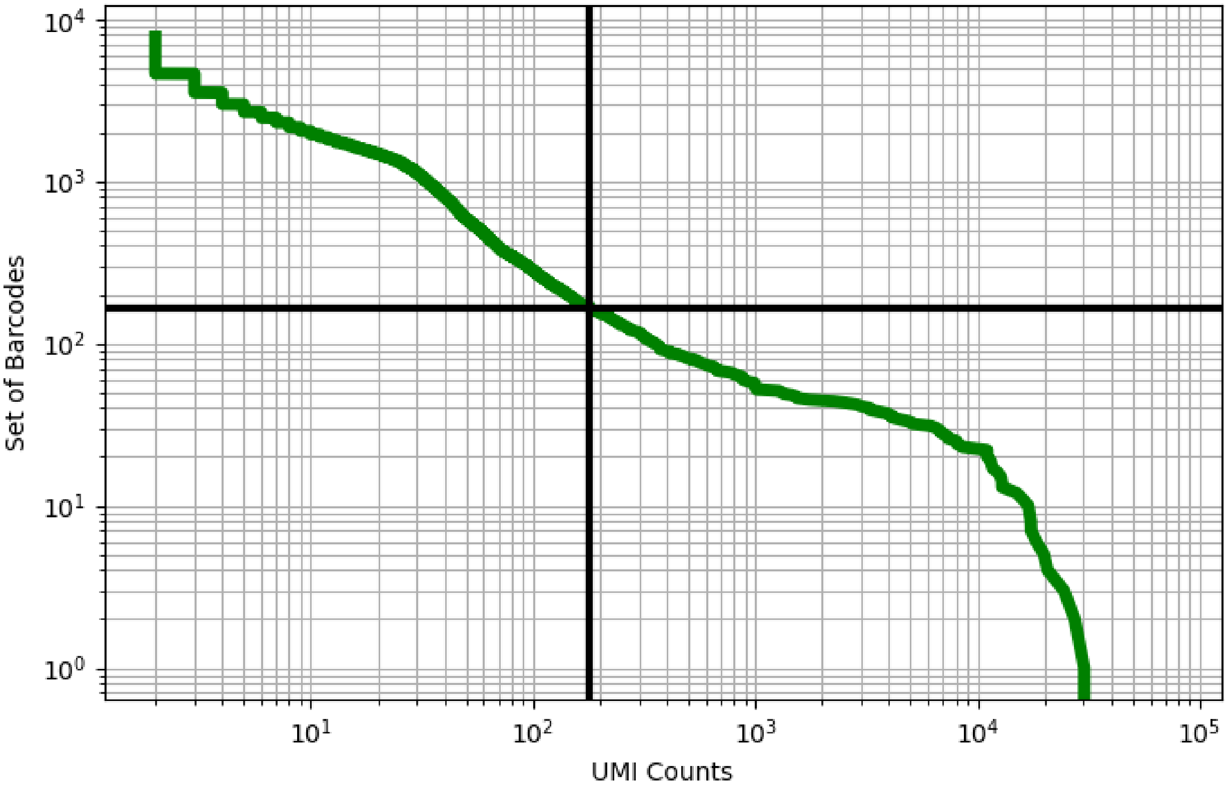


Figure 172: eye/GSM4037987

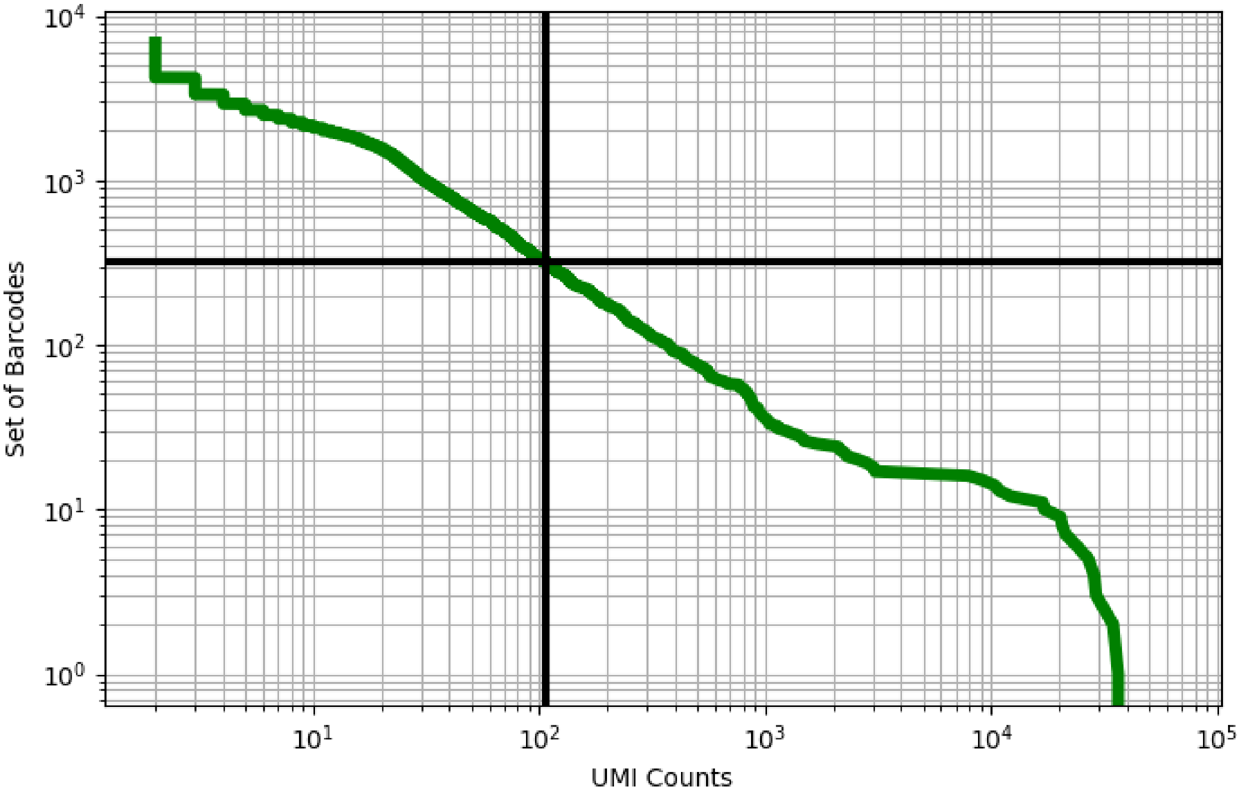


Figure 173: eye/GSM4037988

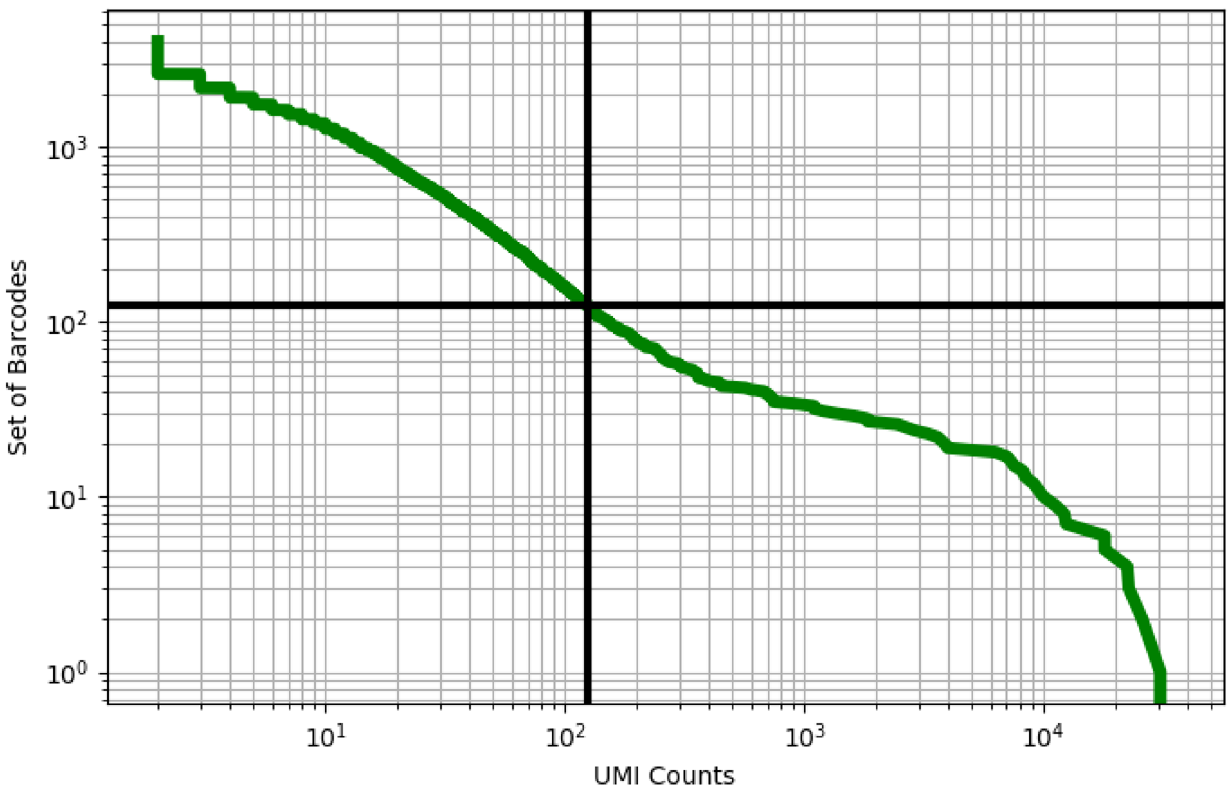


Figure 174: eye/GSM4037989

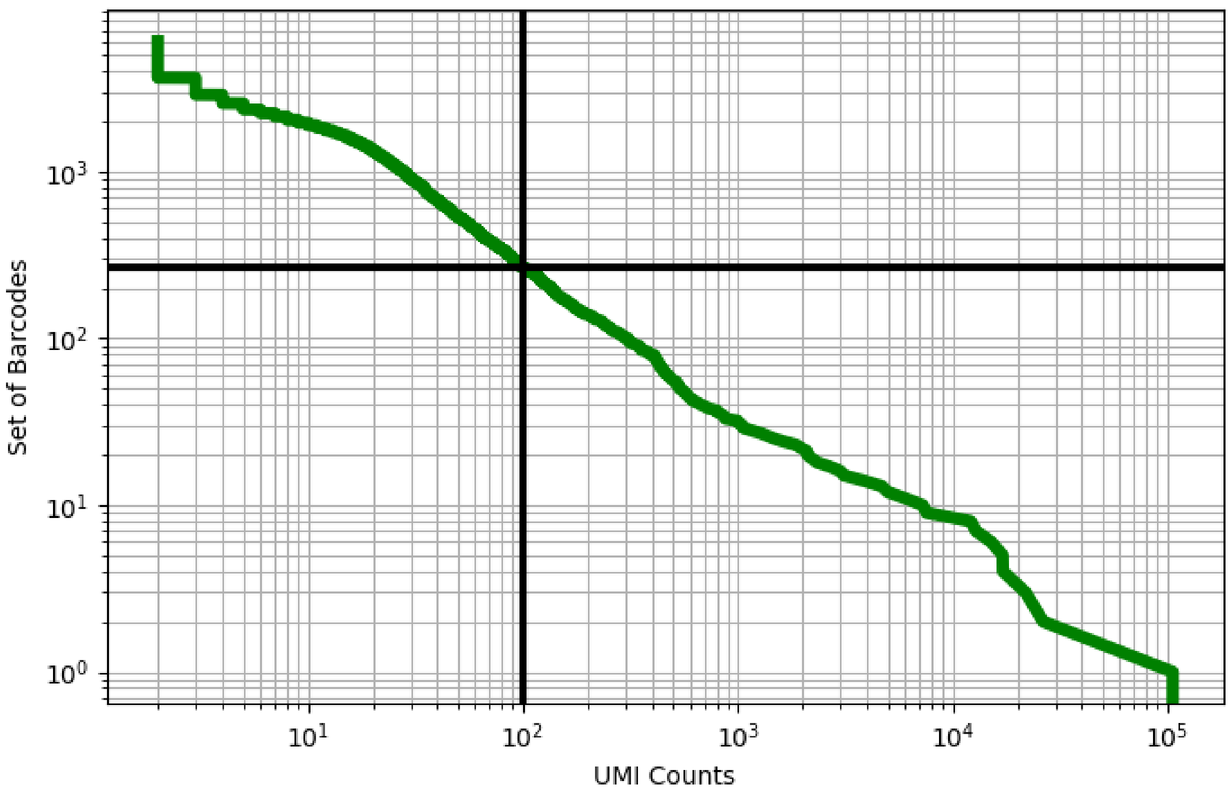


Figure 175: eye/GSM4037990

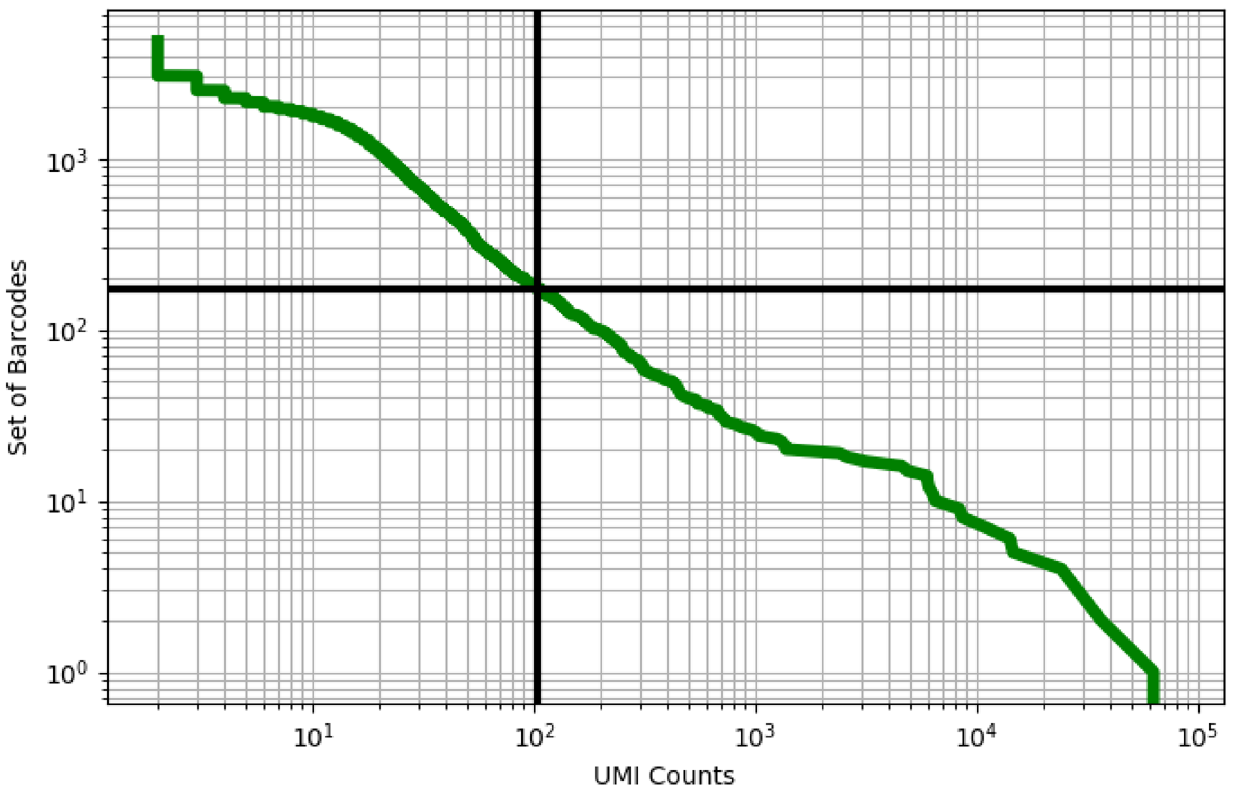


Figure 176: eye/GSM4037991

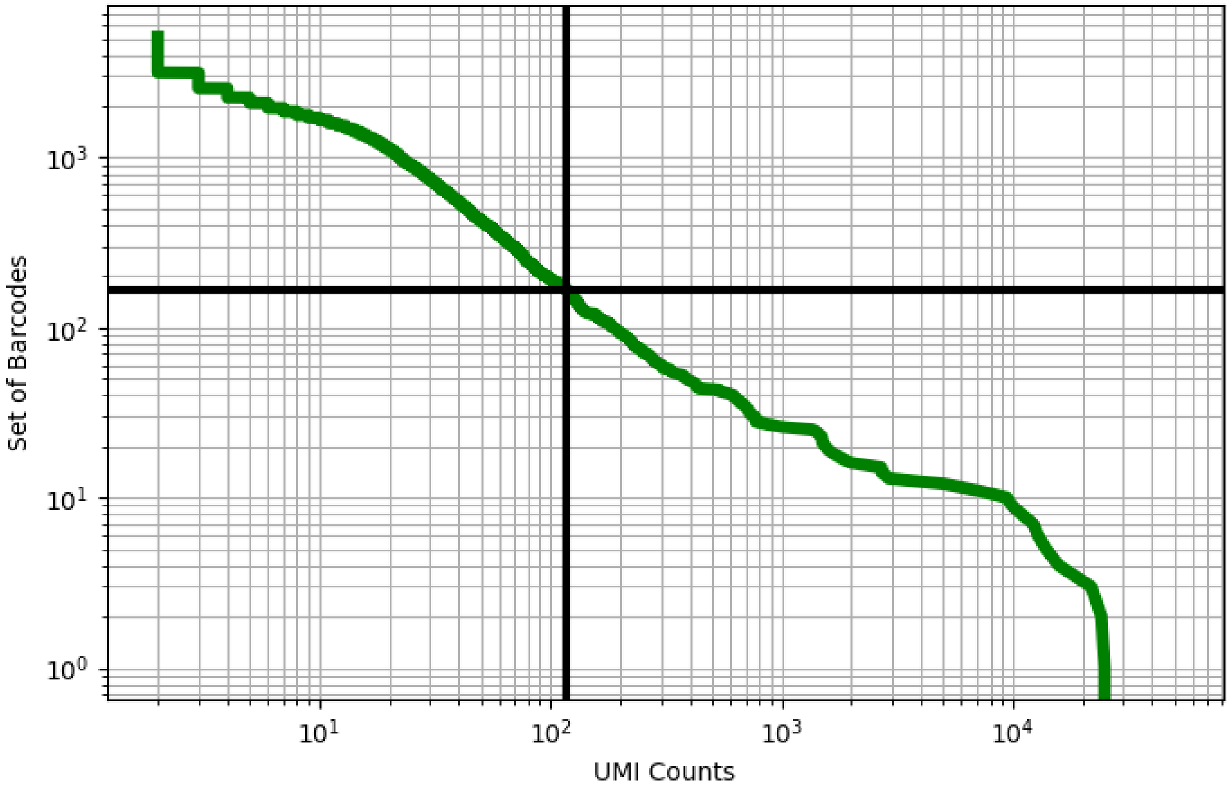


Figure 177: eye/GSM4037992

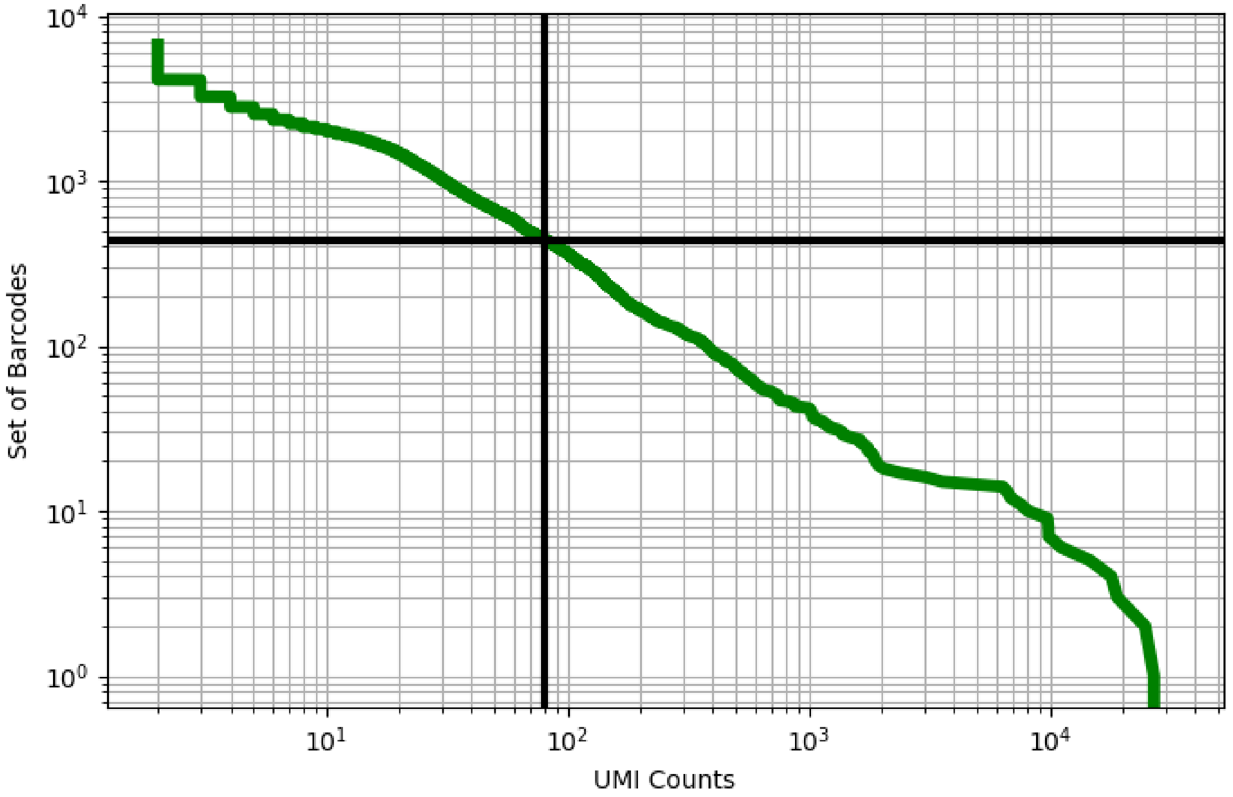


Figure 178: eye/GSM4037993

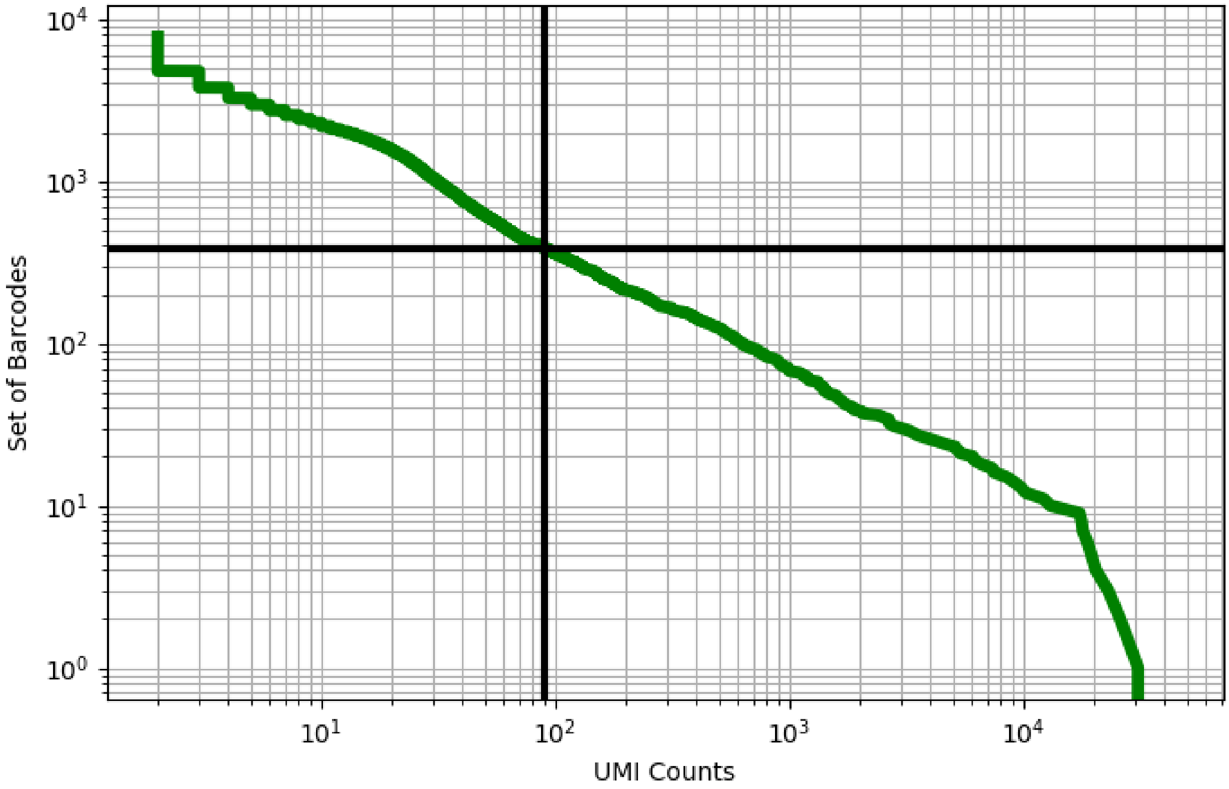


Figure 179: eye/GSM4037994

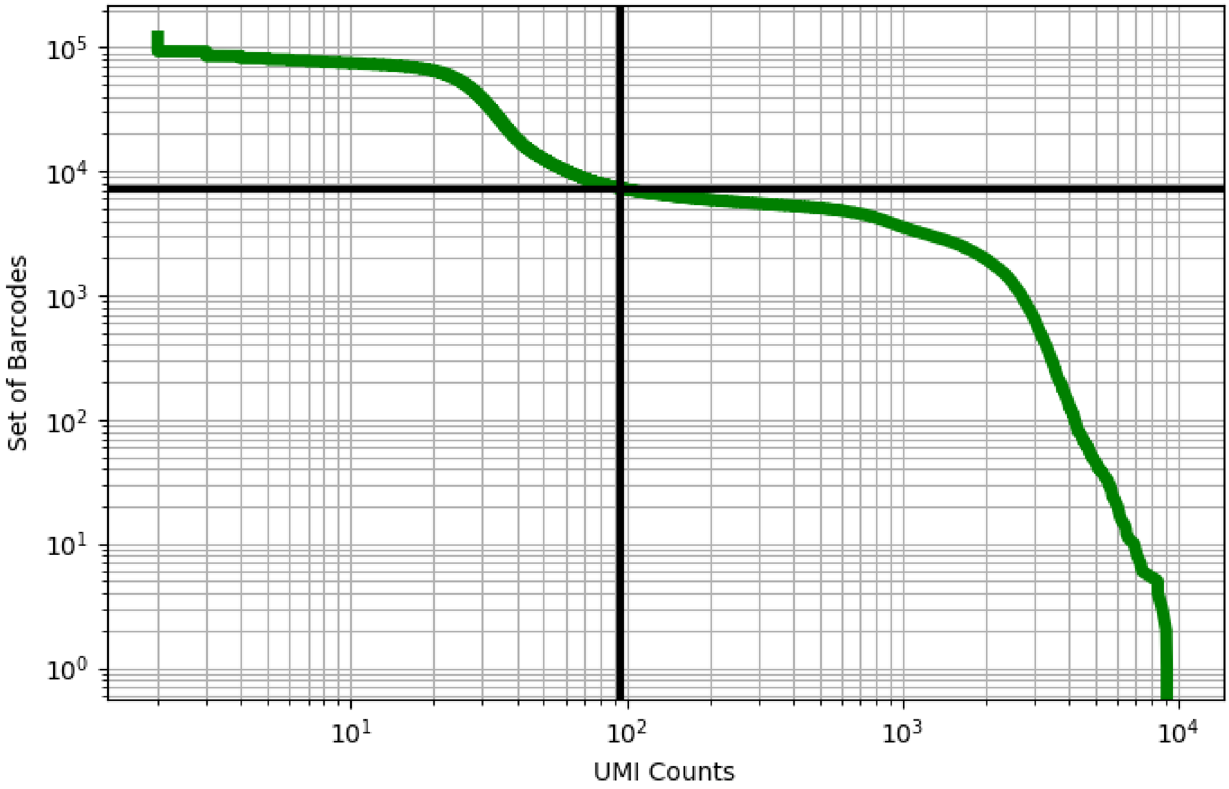


Figure 180: eye/GSM4463223

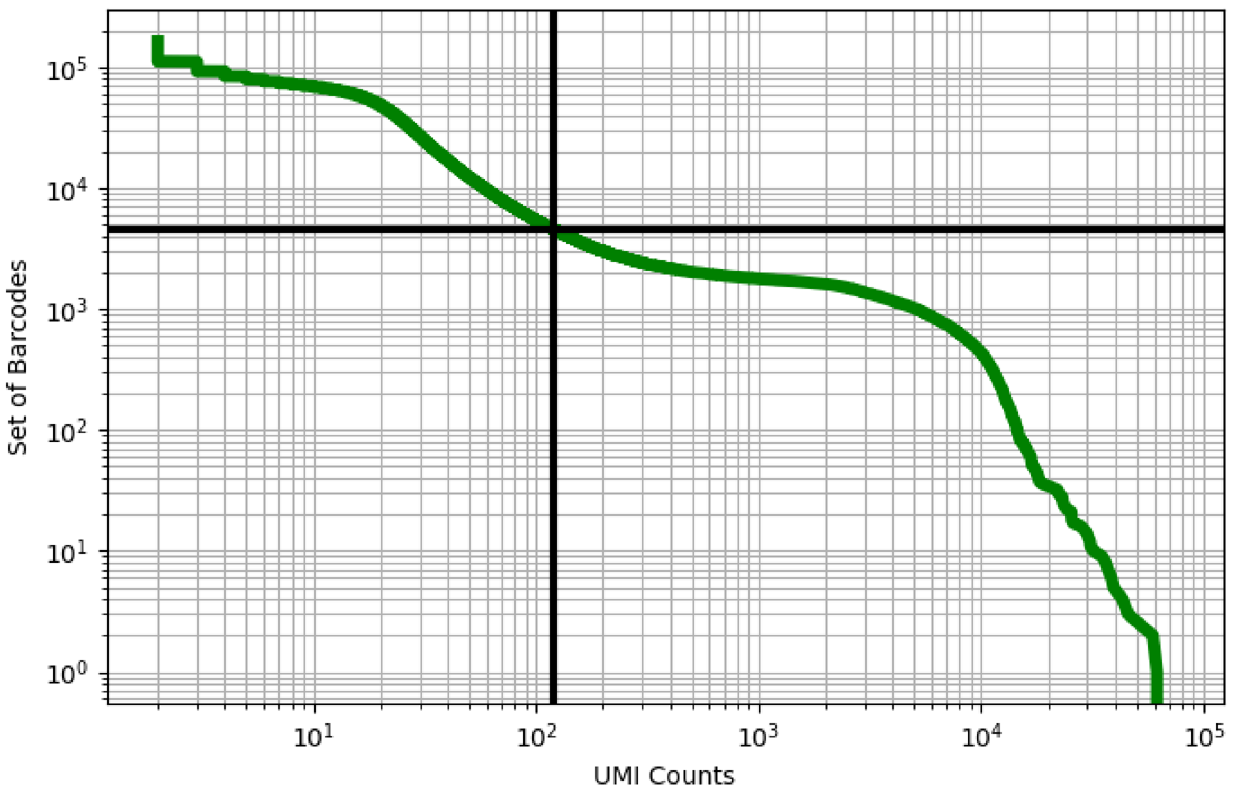


Figure 181: eye/GSM4463229

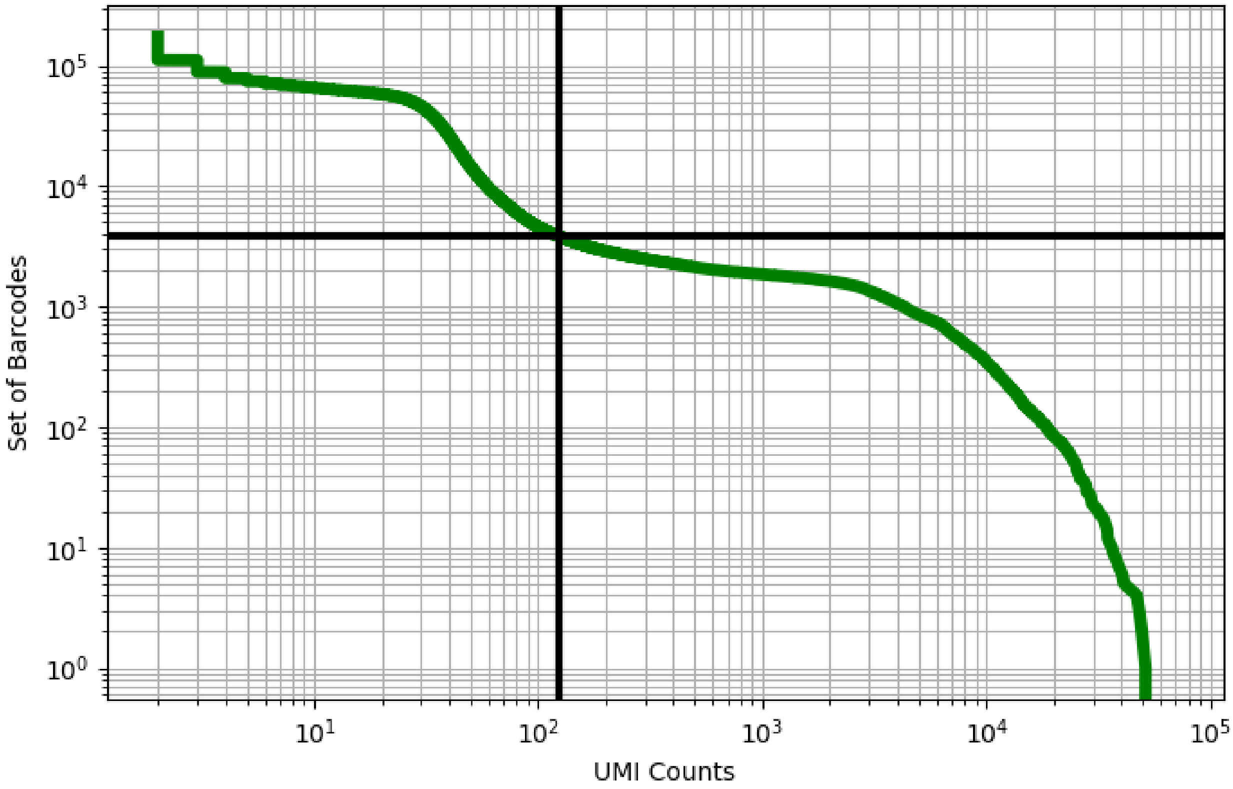


Figure 182: eye/GSM4463234

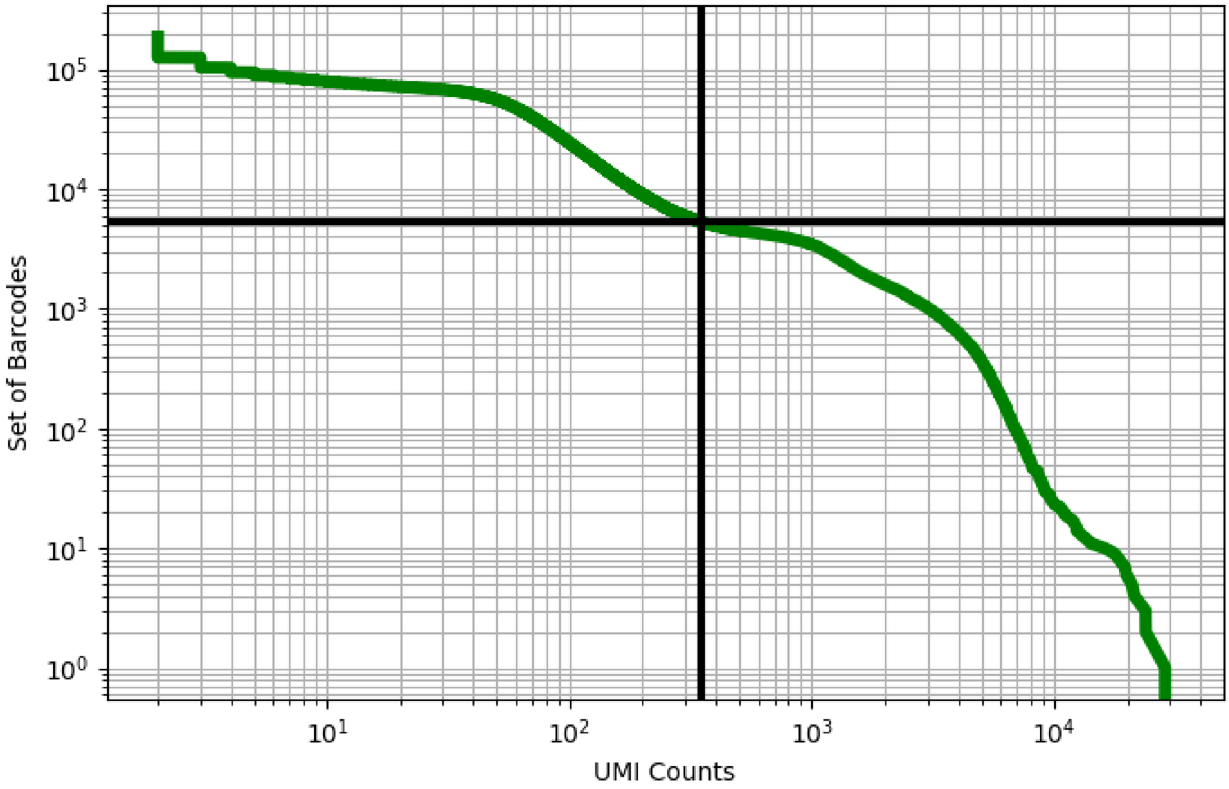


Figure 183: eye/GSM4463239

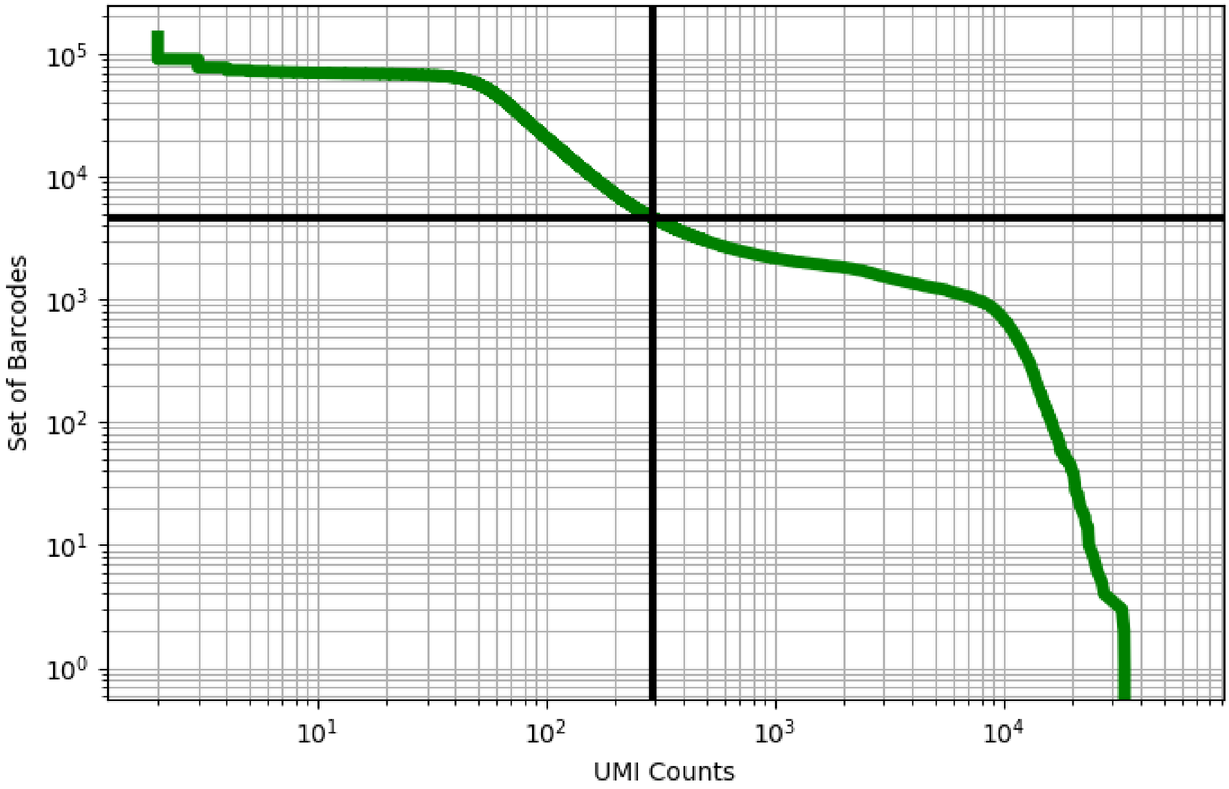


Figure 184: eye/GSM4463244

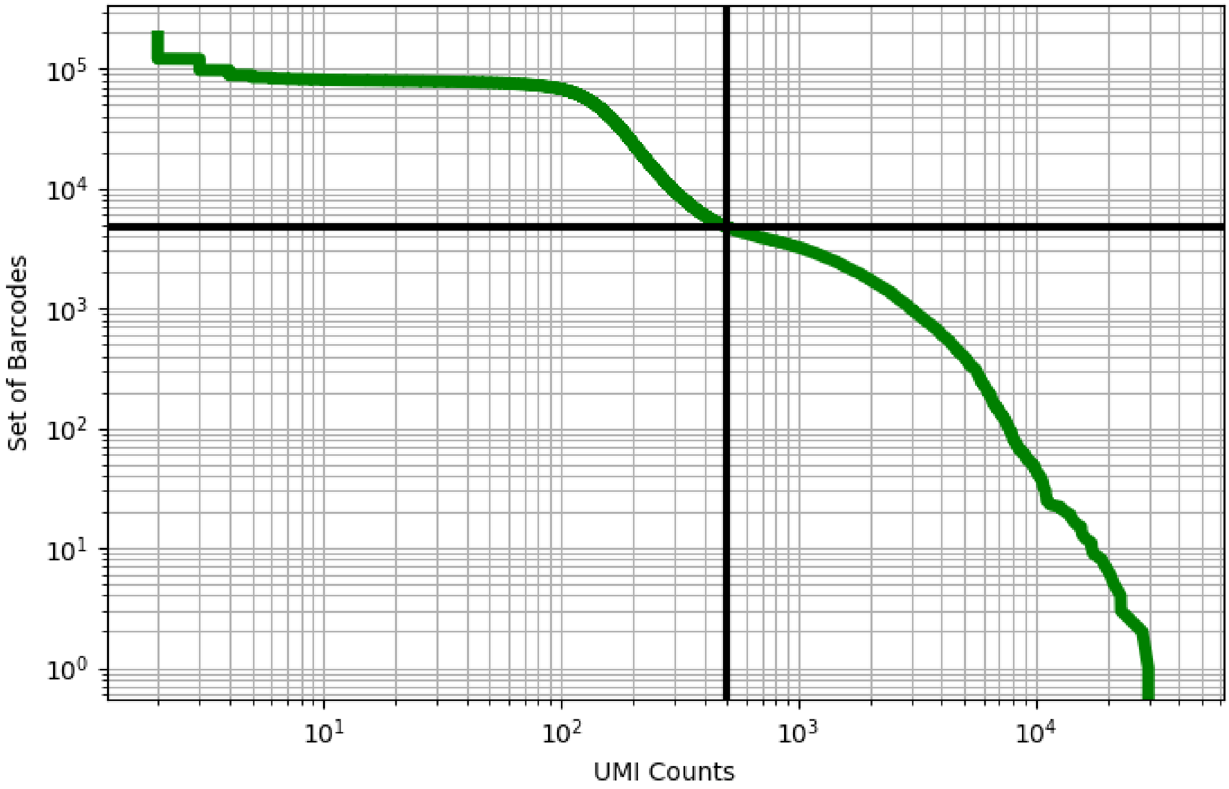


Figure 185: eye/GSM4463249

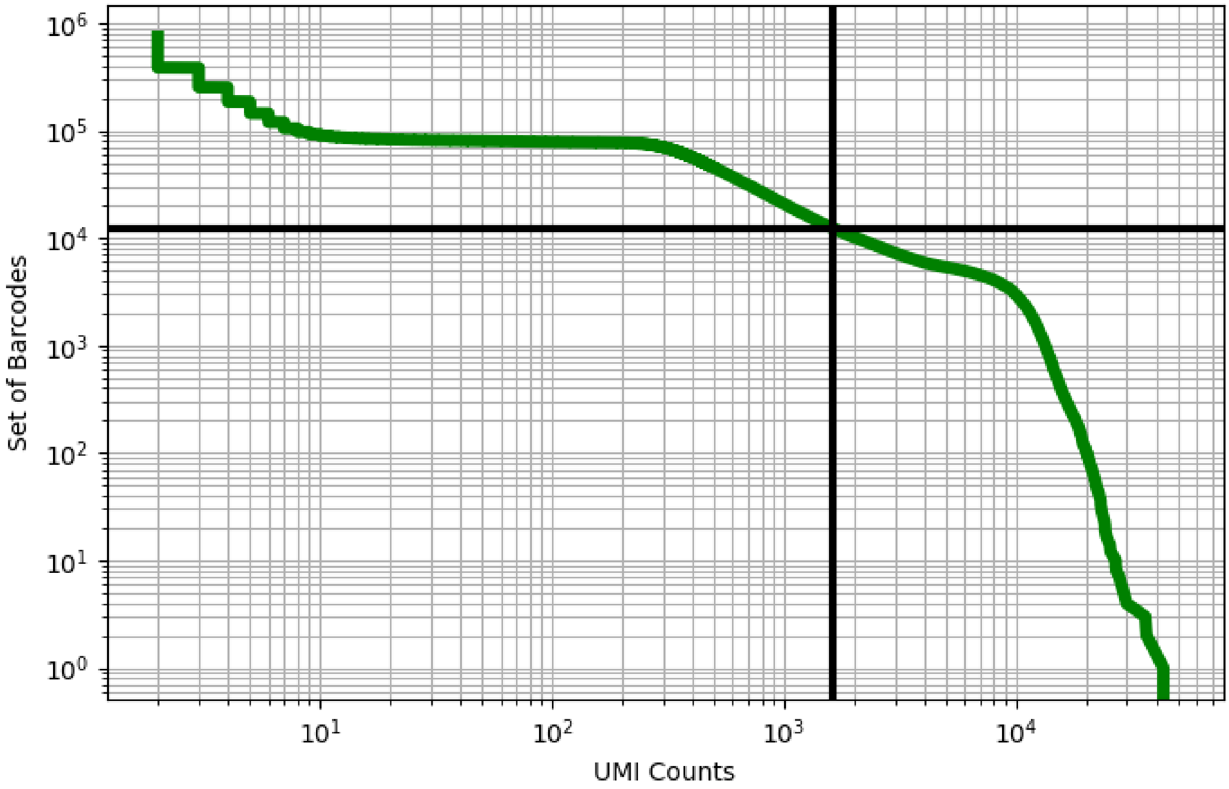


Figure 186: eye/GSM4463253

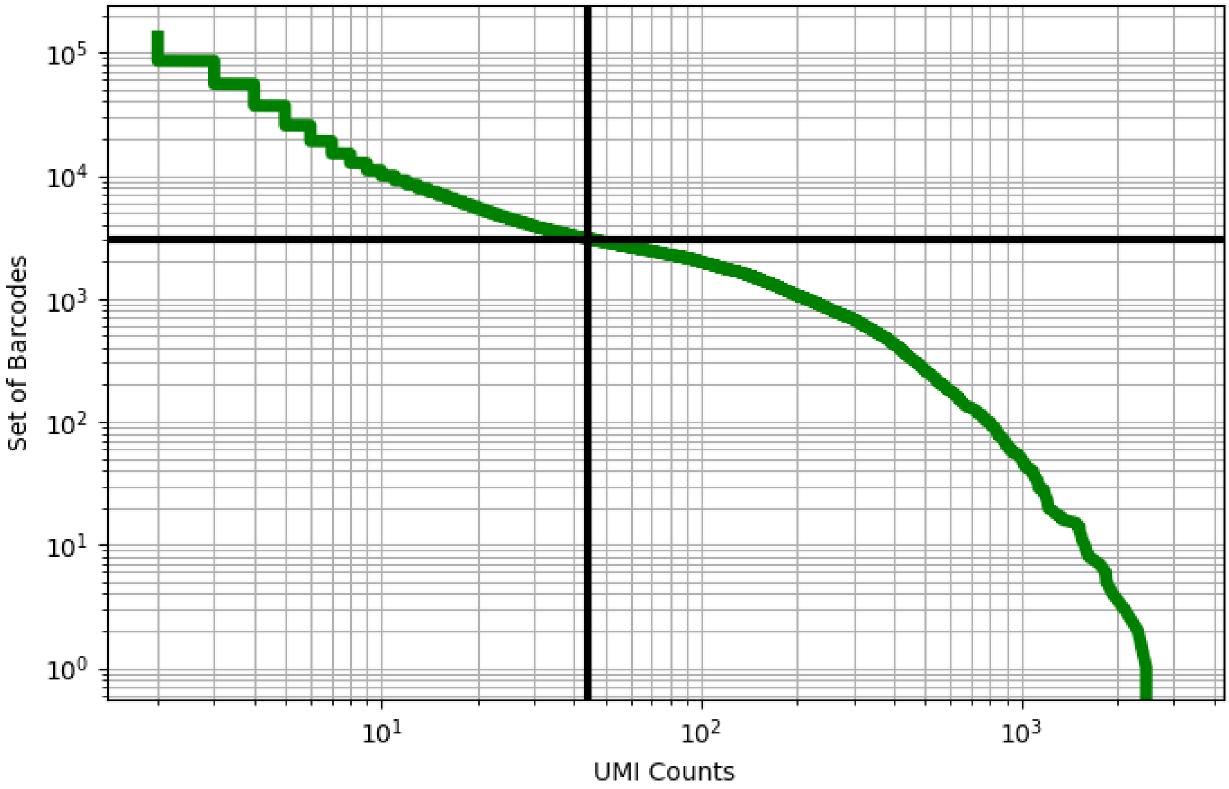


Figure 187: ileum/GSM3576426

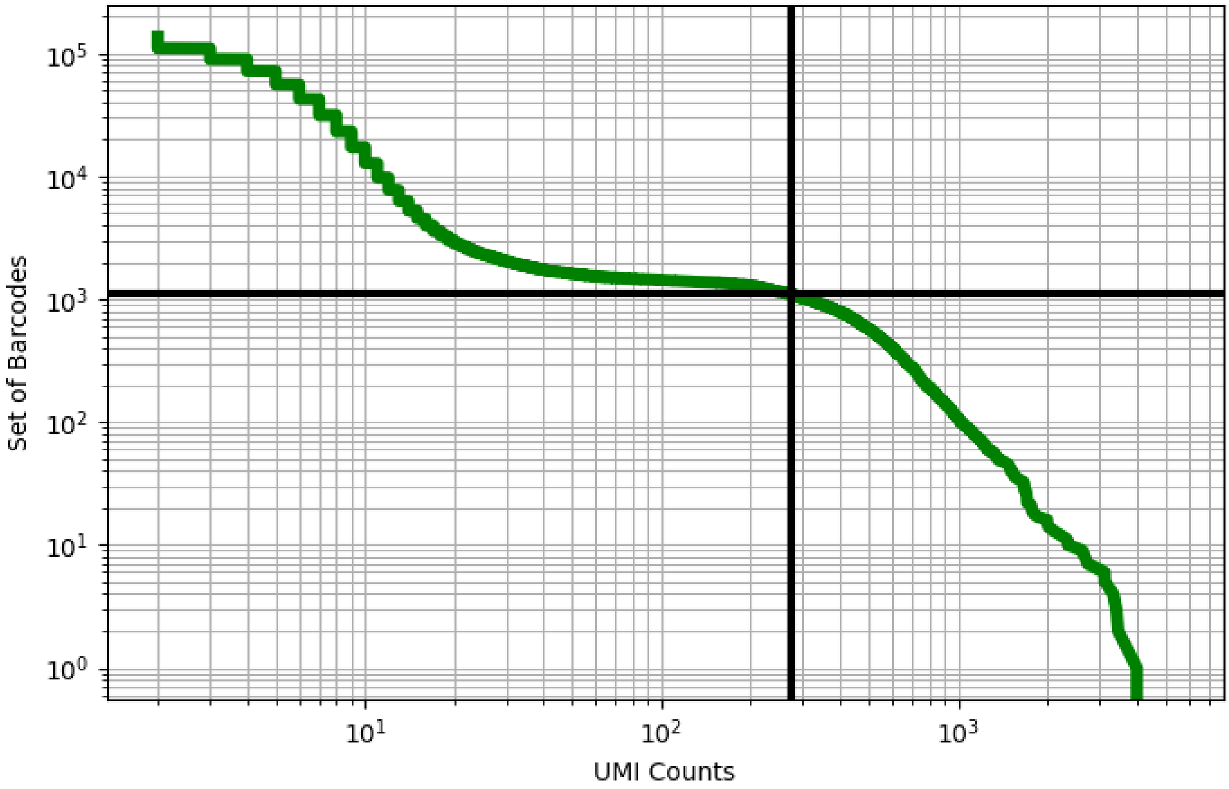


Figure 188: ileum/GSM3576427

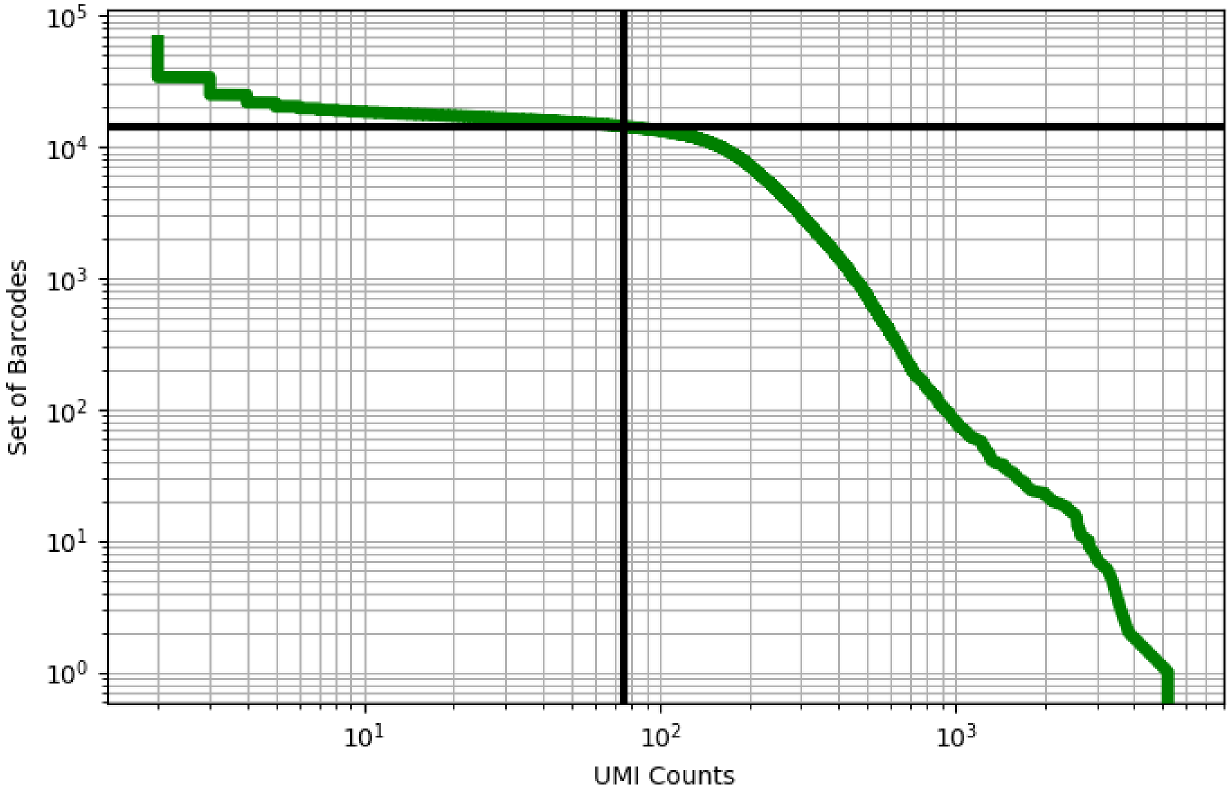


Figure 189: ileum/GSM3576428

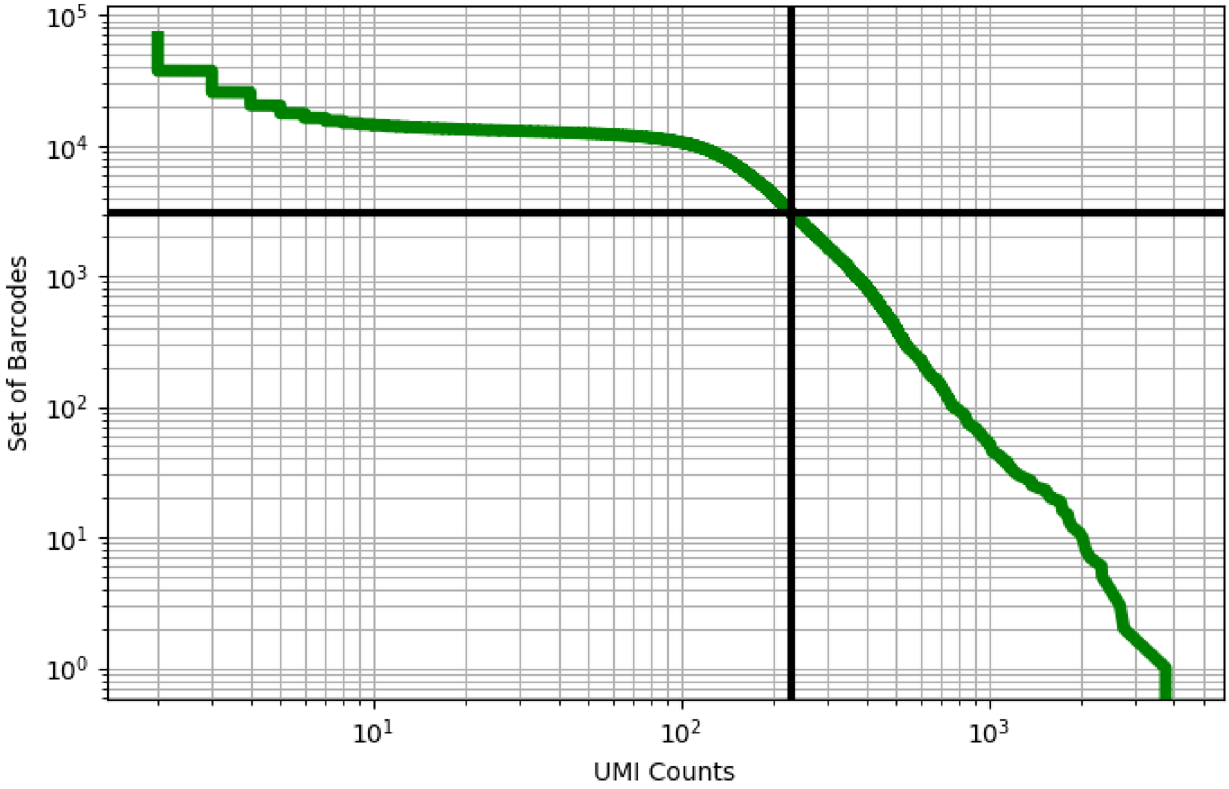


Figure 190: ileum/GSM3576429

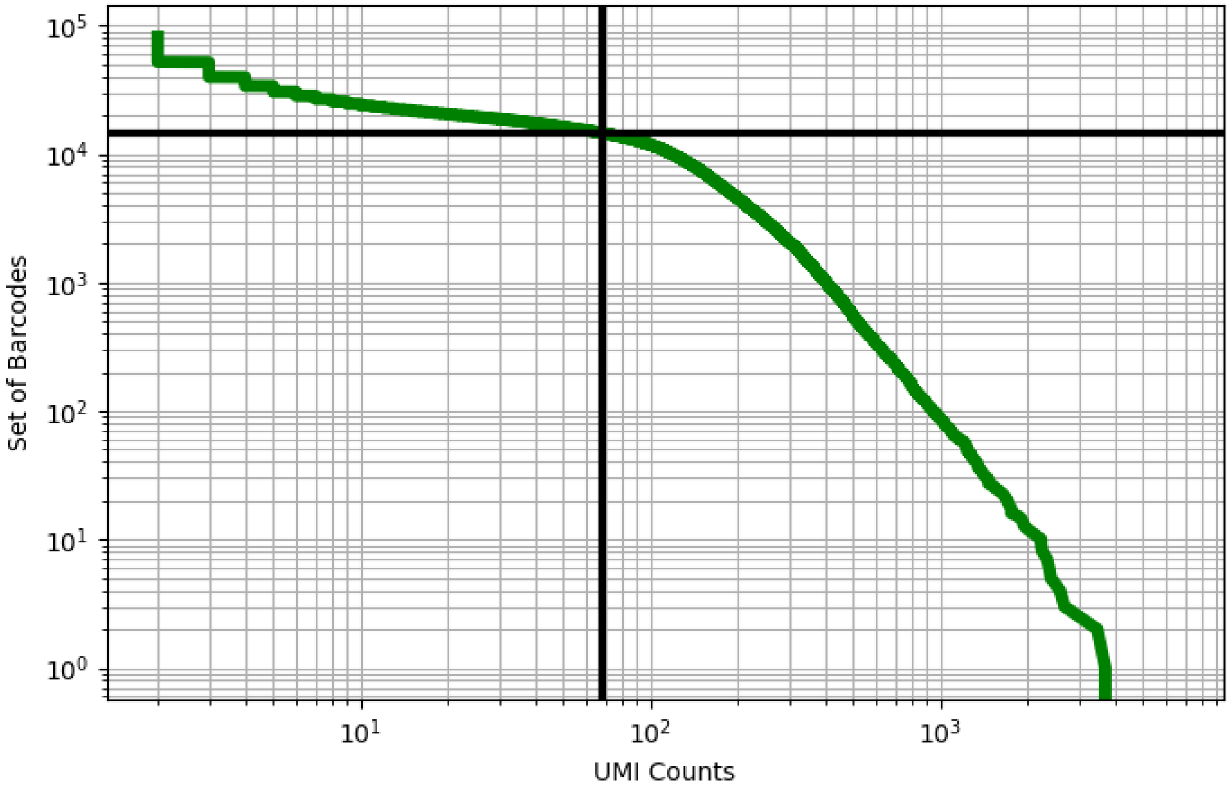


Figure 191: ileum/GSM3576430

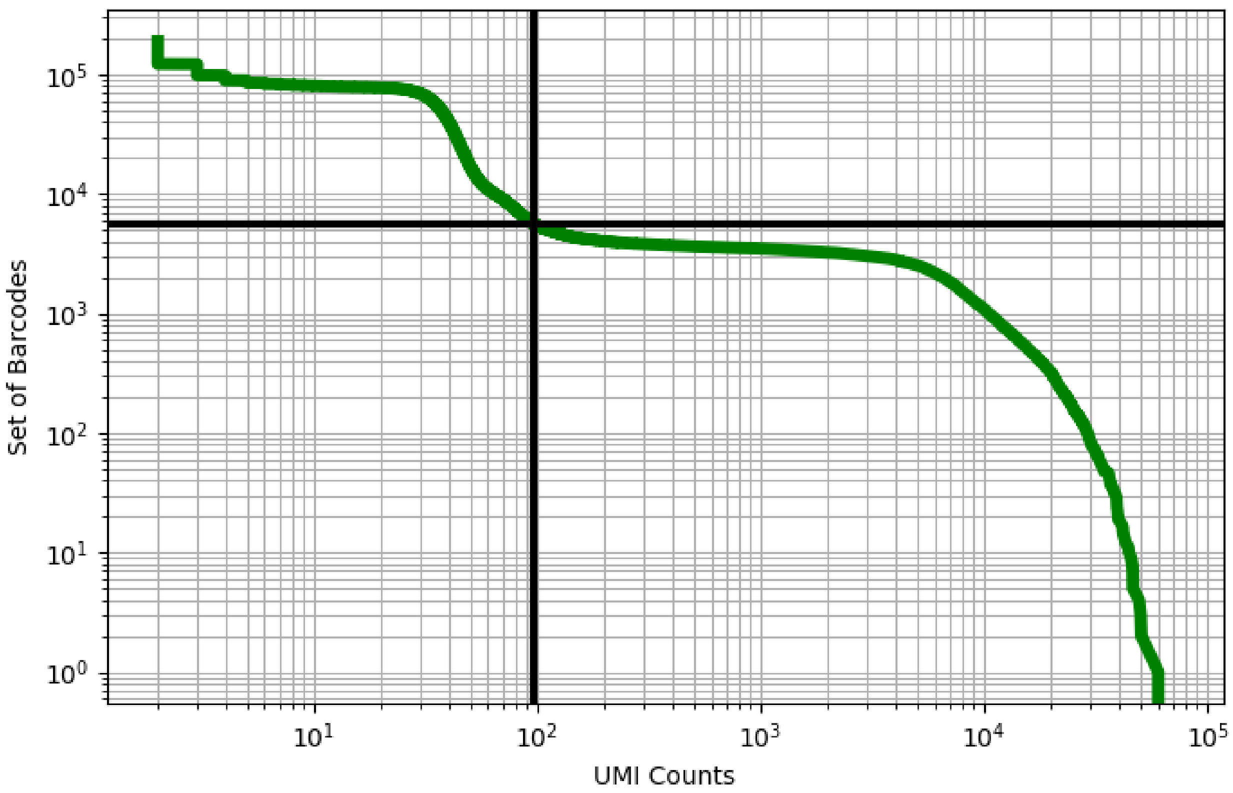


Figure 192: ileum/GSM3587008

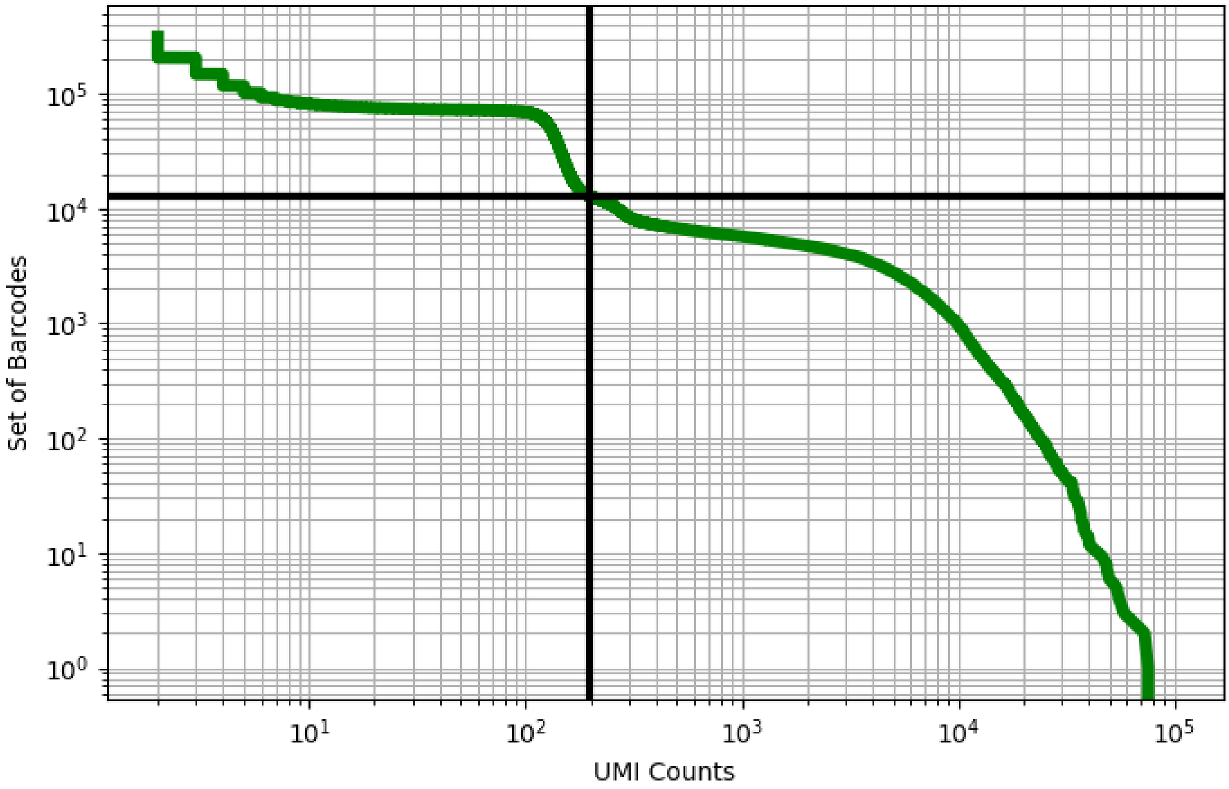


Figure 193: ileum/GSM3587009

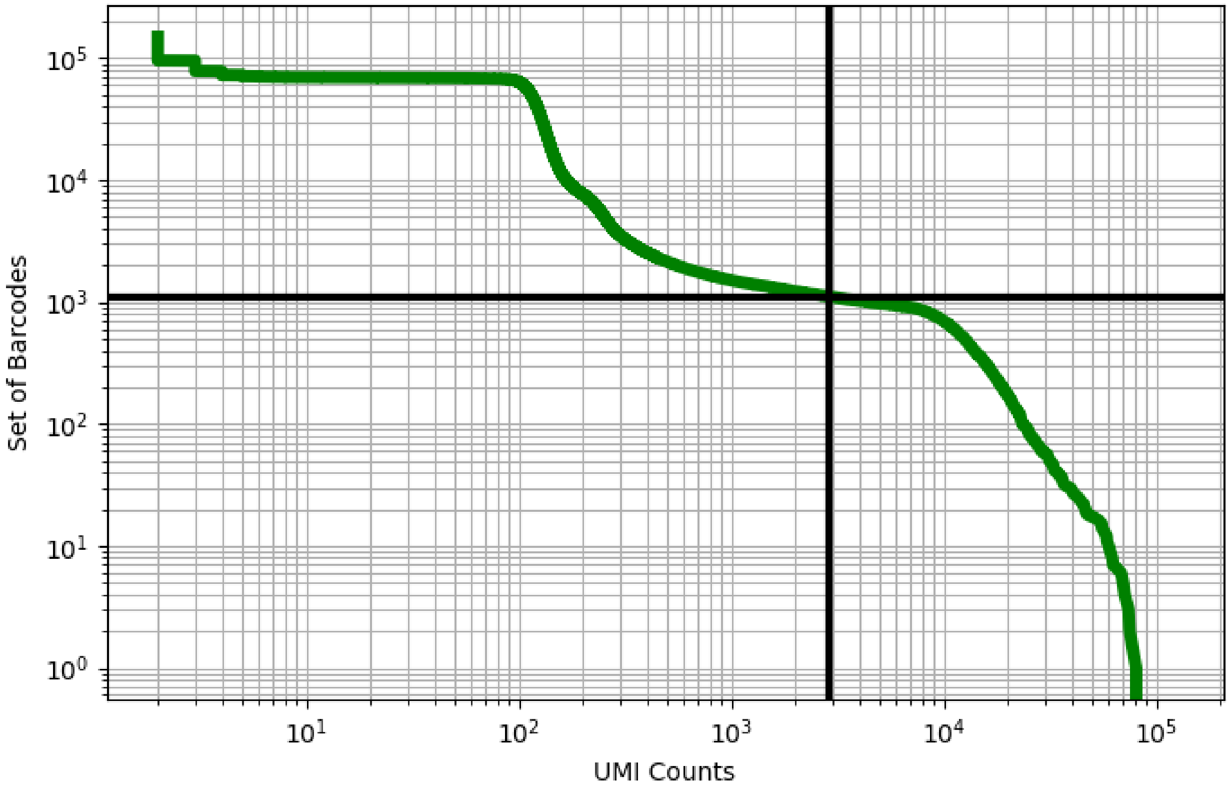


Figure 194: kidney/ERS3861771

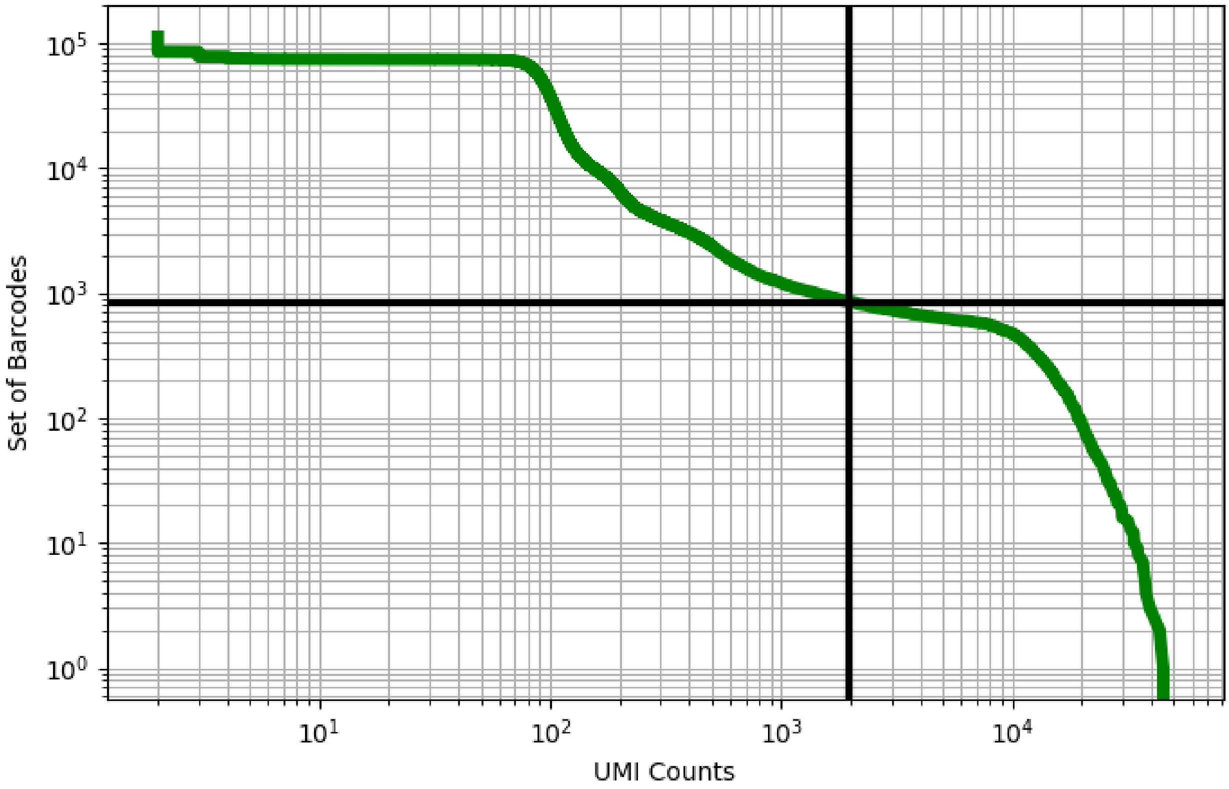


Figure 195: kidney/ERS3861772

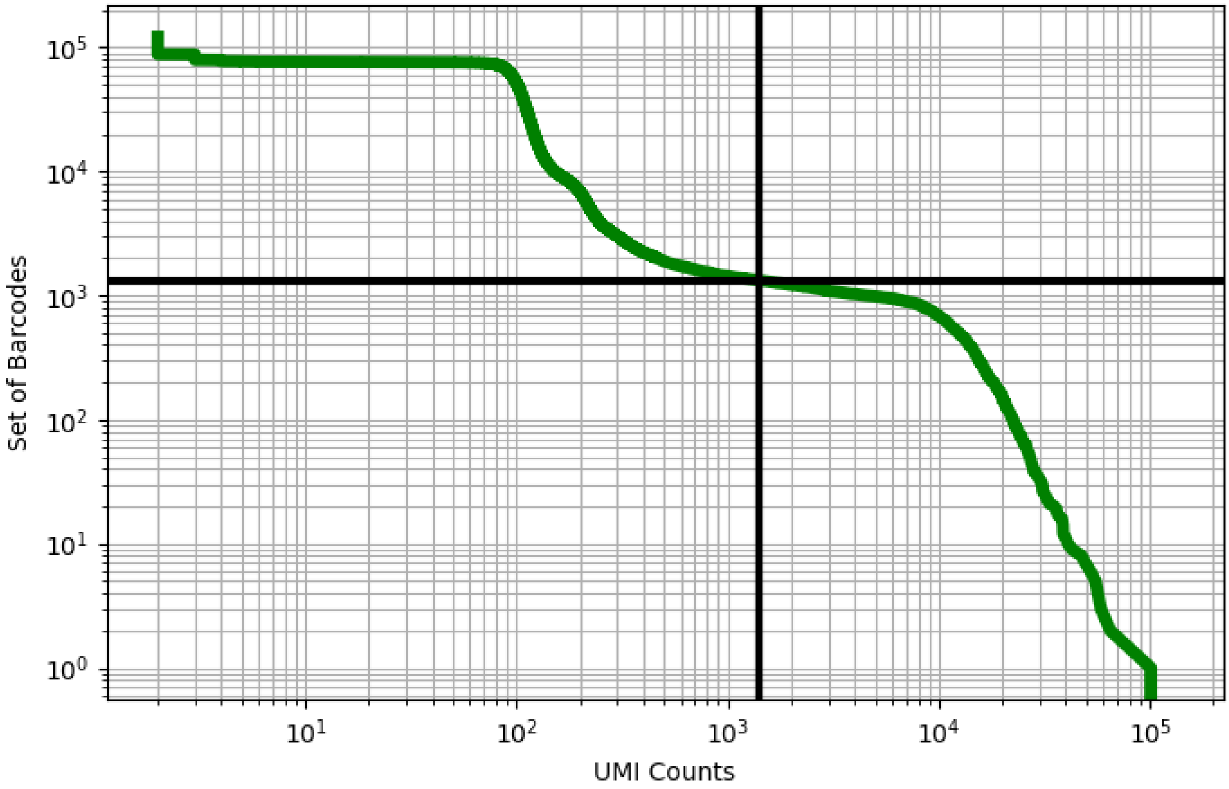


Figure 196: kidney/ERS3861777

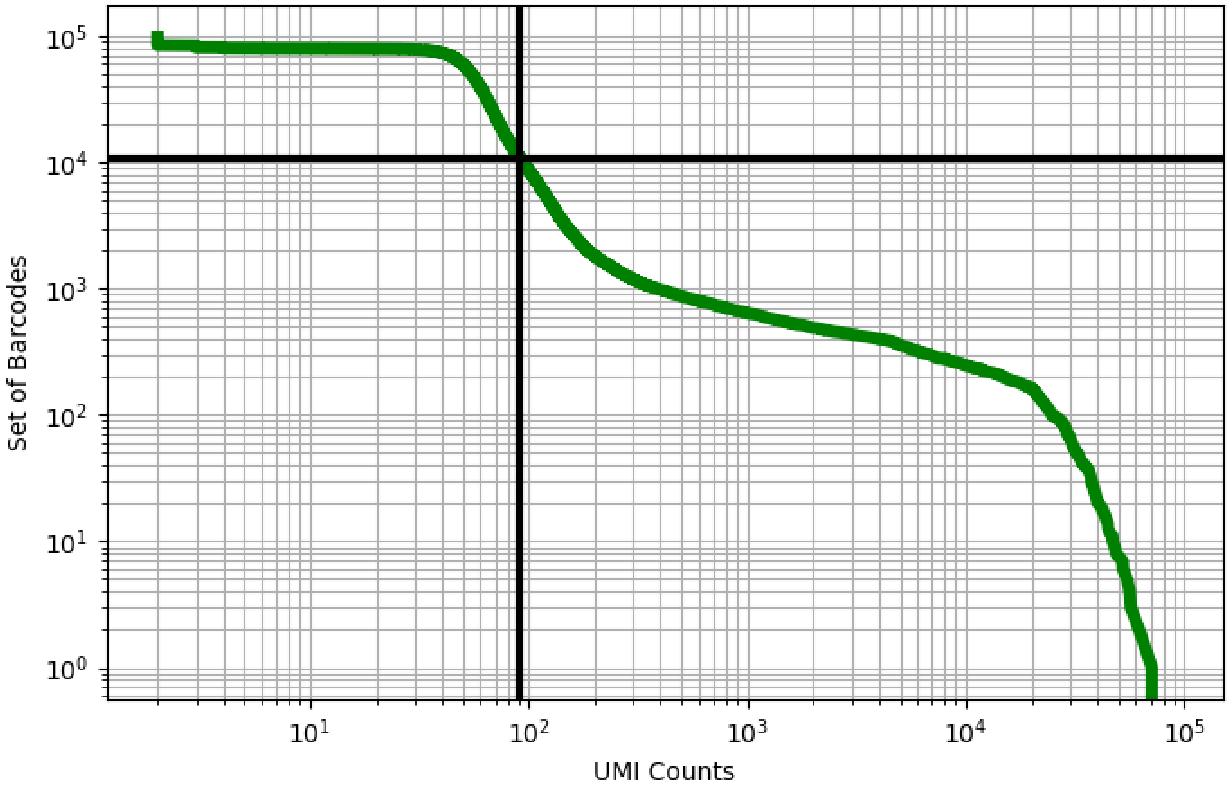


Figure 197: kidney/ERS3861780

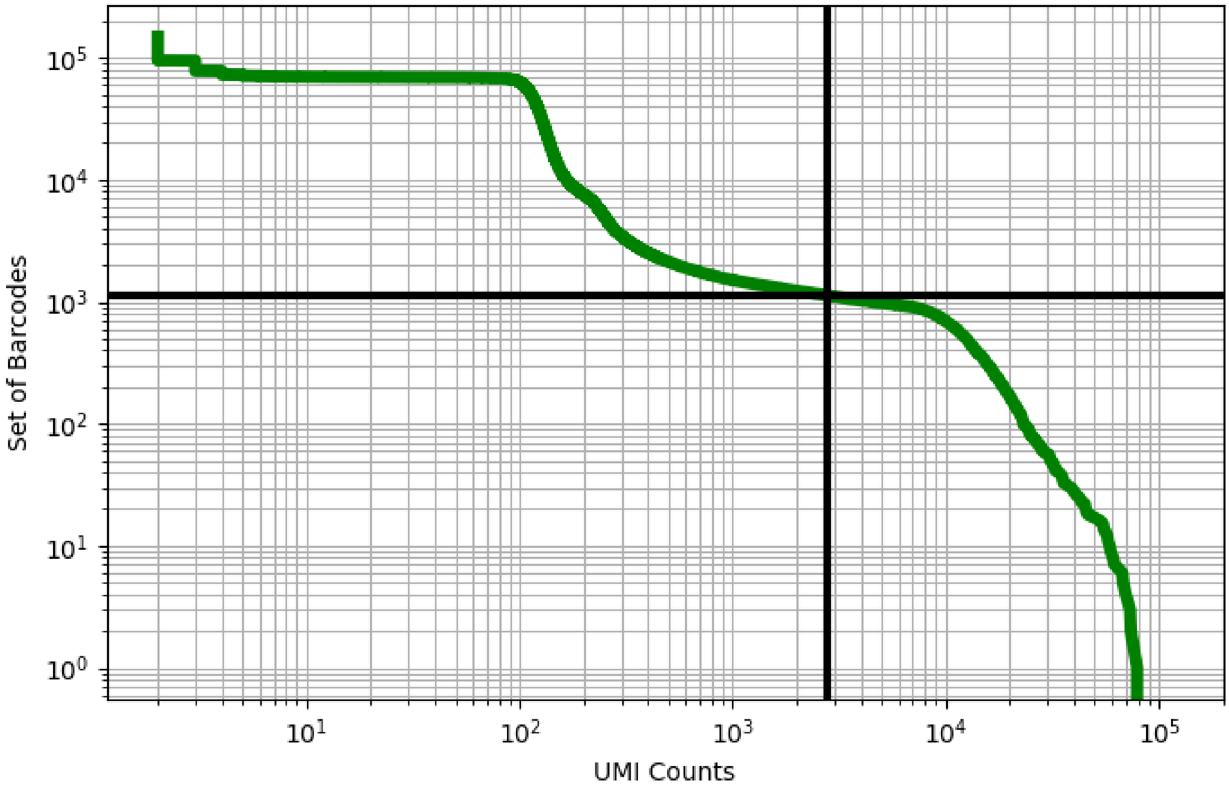


Figure 198: kidney/ERS3861781

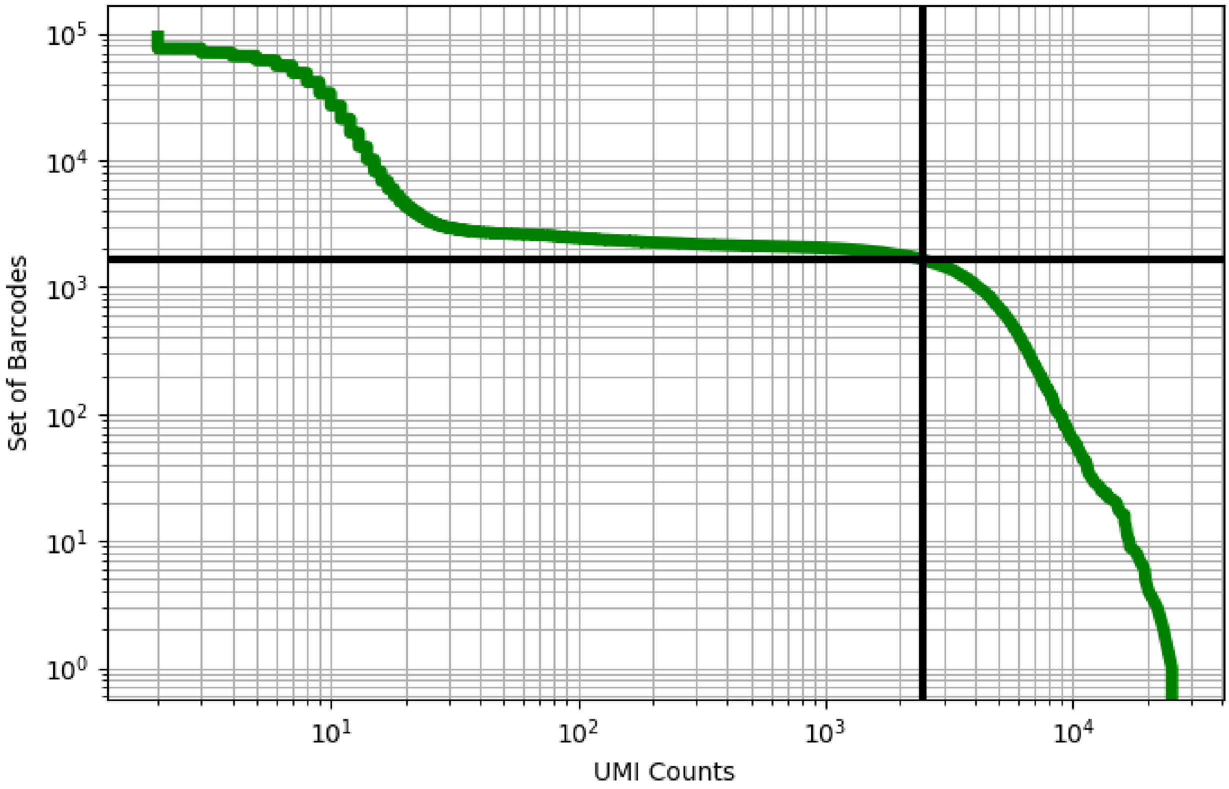


Figure 199: kidney/ERS3861828

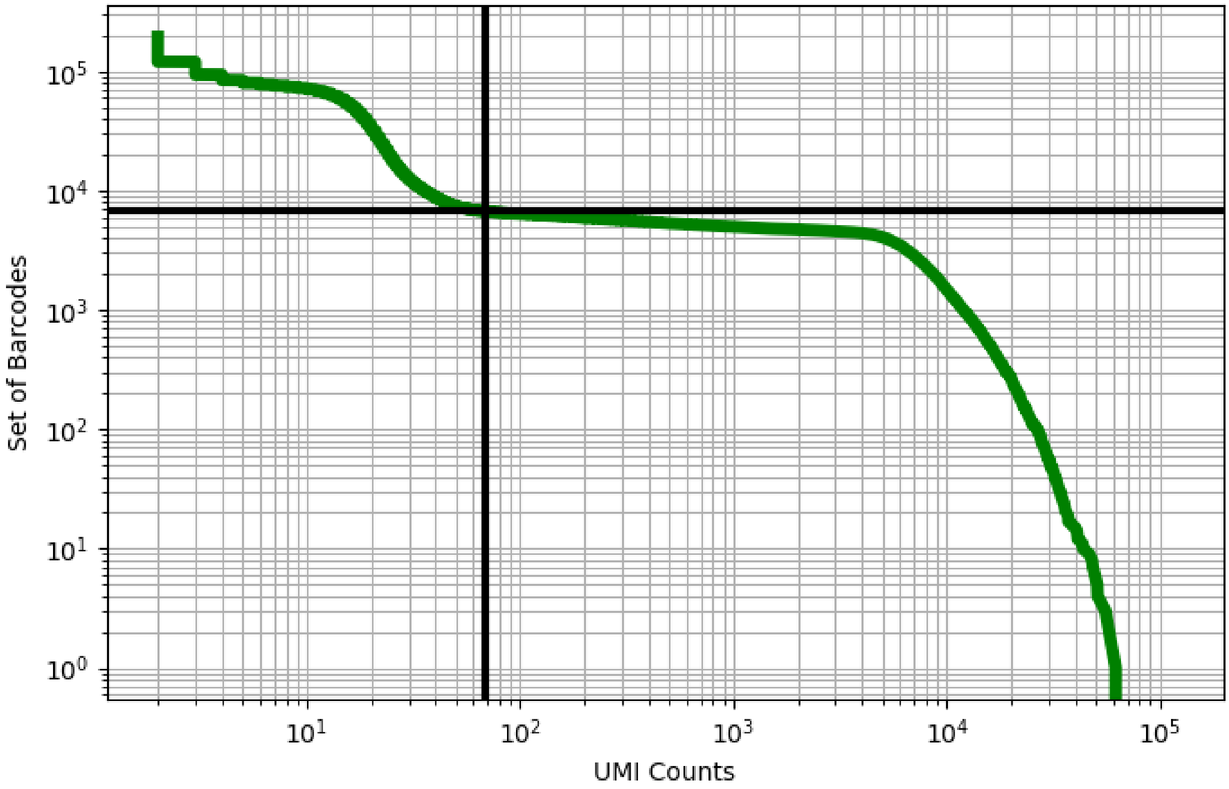


Figure 200: kidney/ERS3861829

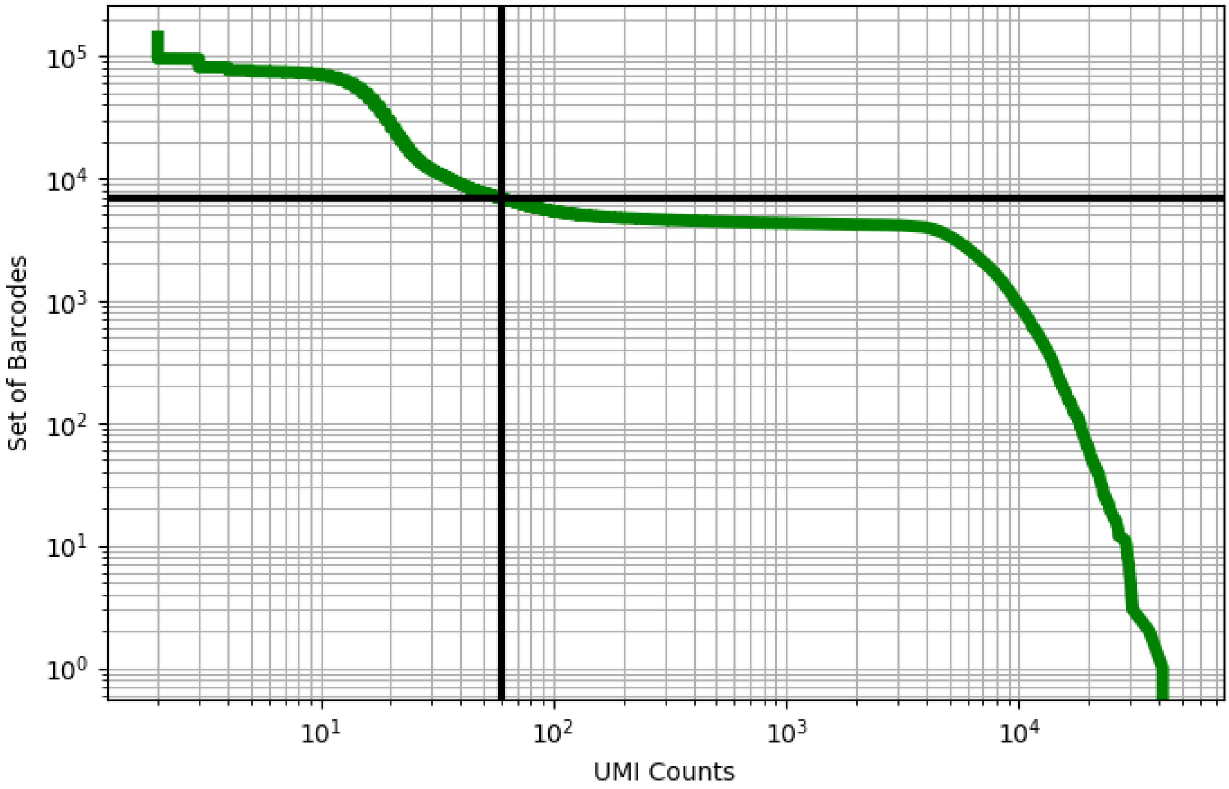


Figure 201: kidney/GSM3073088

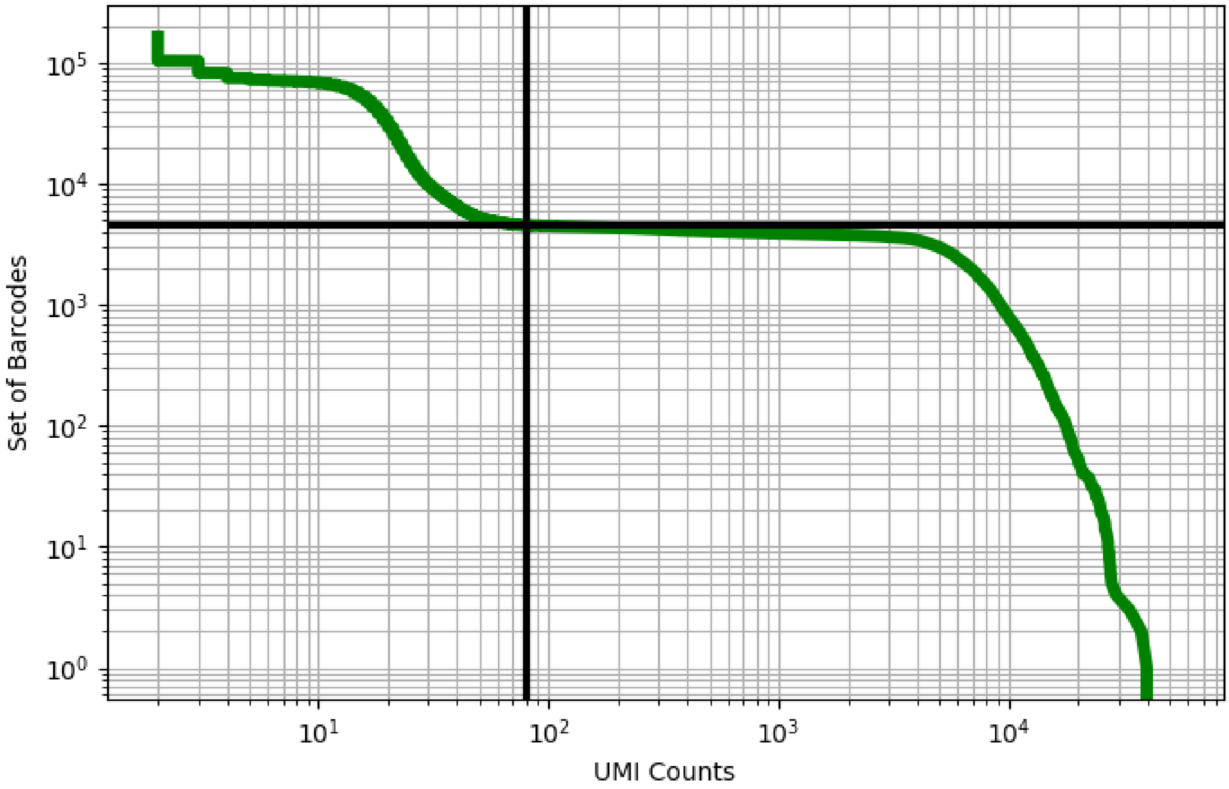


Figure 202: kidney/GSM3073089

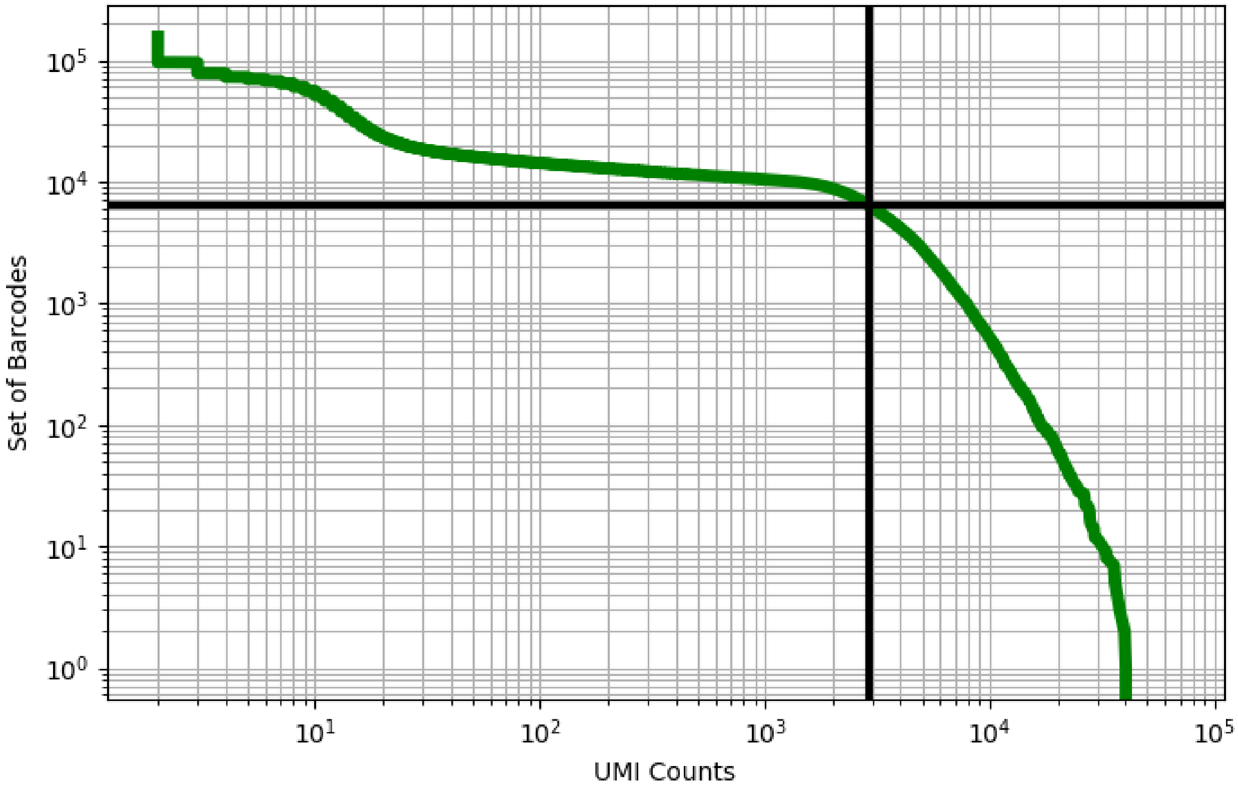


Figure 203: kidney/GSM3143601

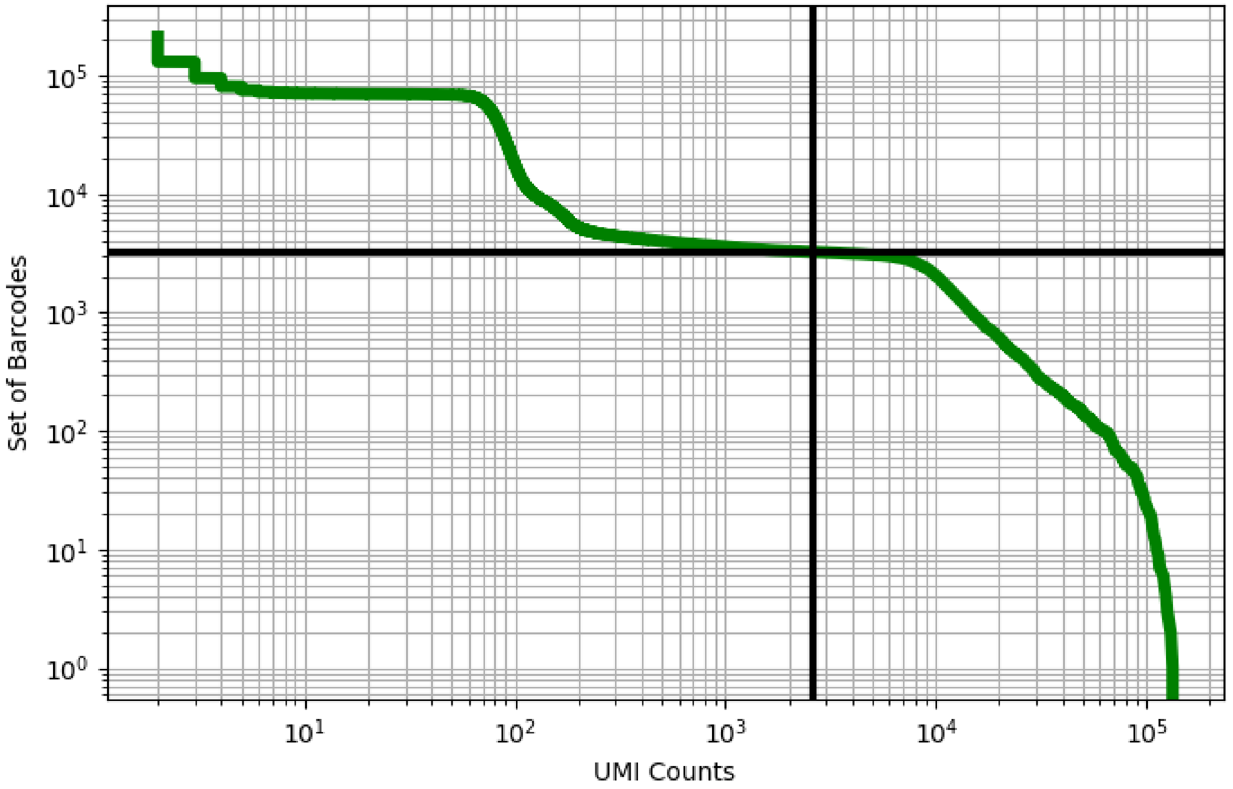


Figure 204: kidney/GSM3273570

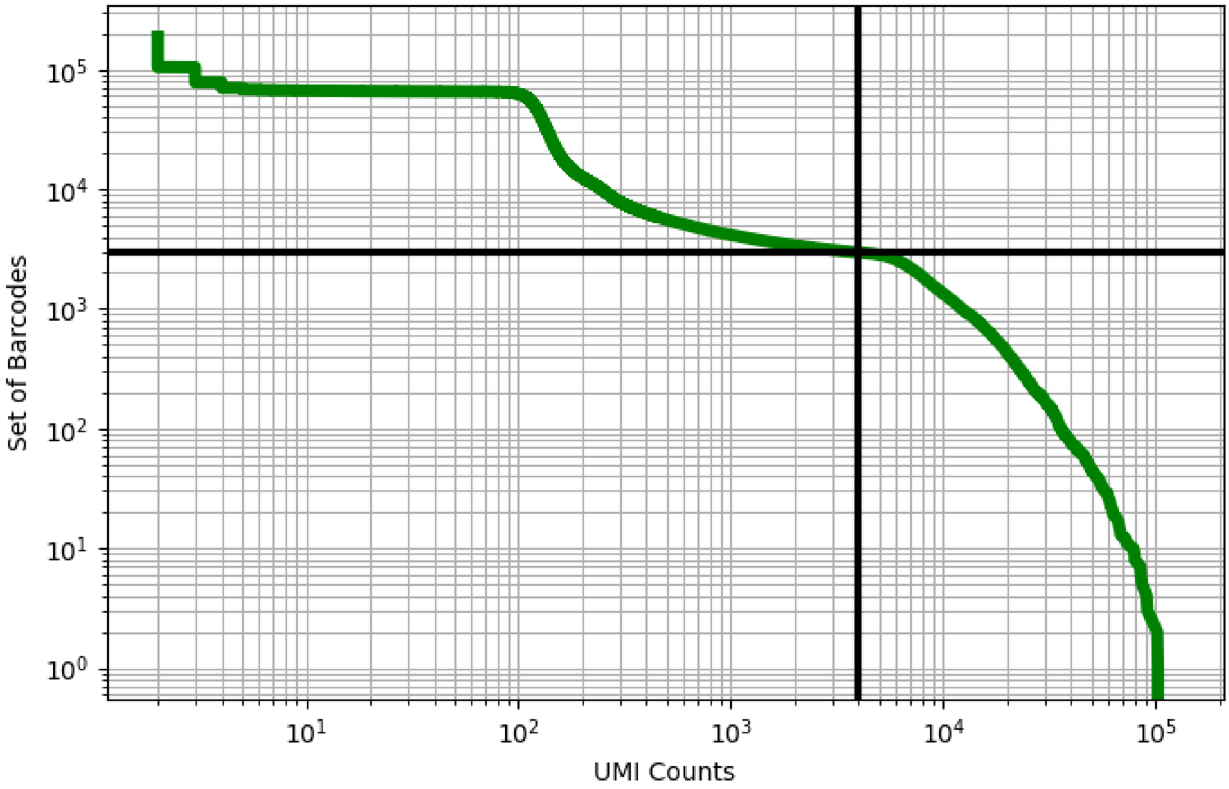


Figure 205: kidney/GSM3377671

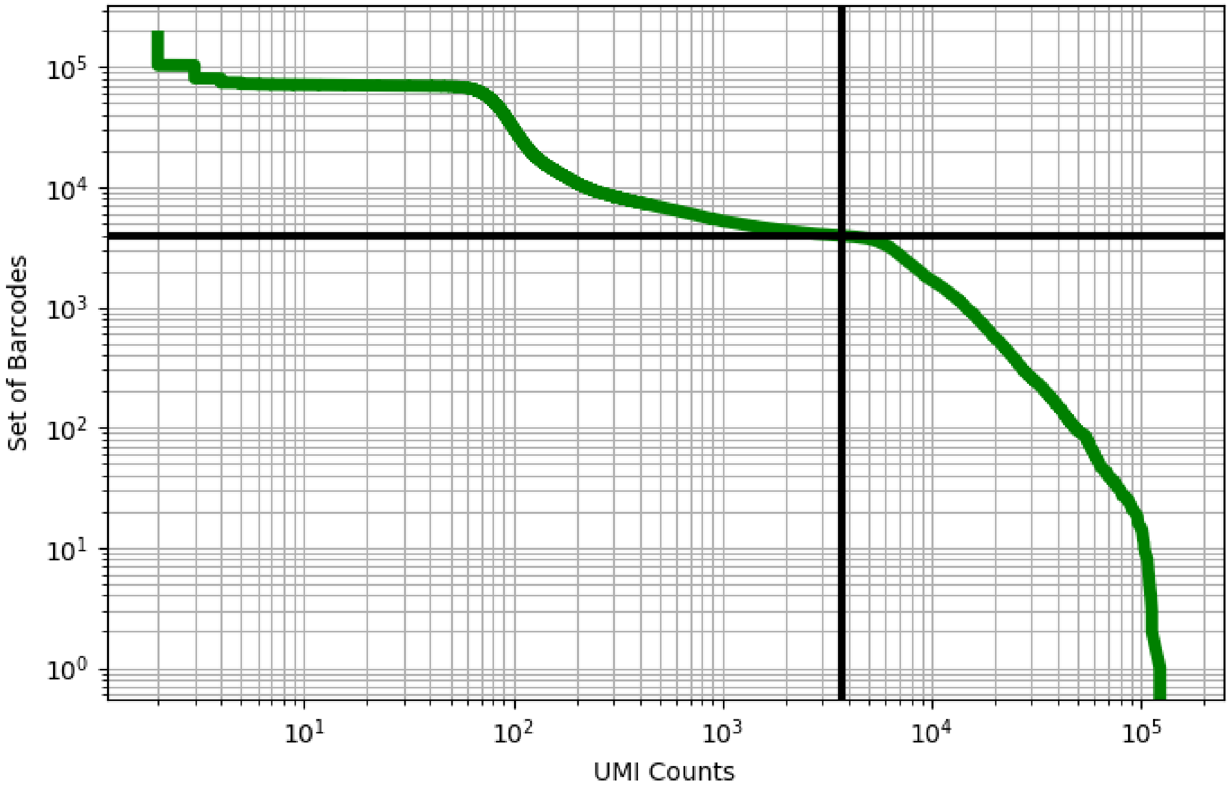


Figure 206: kidney/GSM3377672

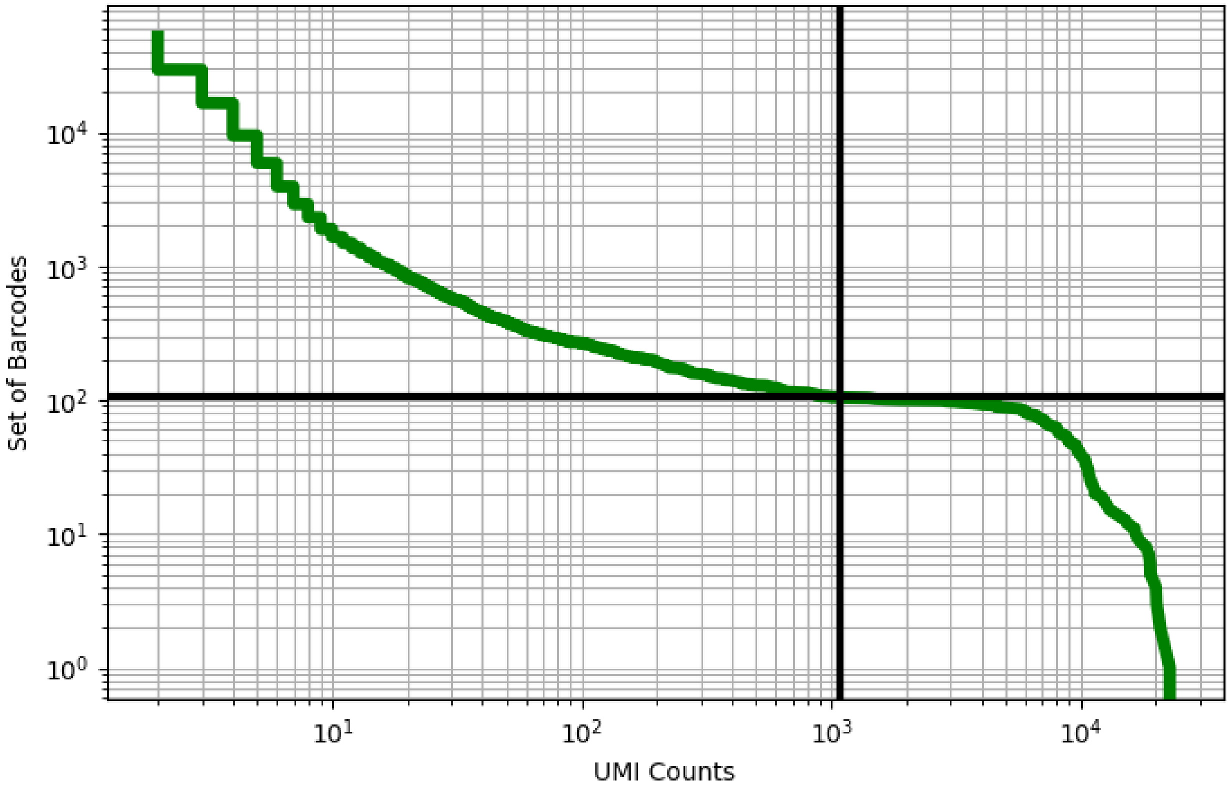


Figure 207: kidney/GSM3509837

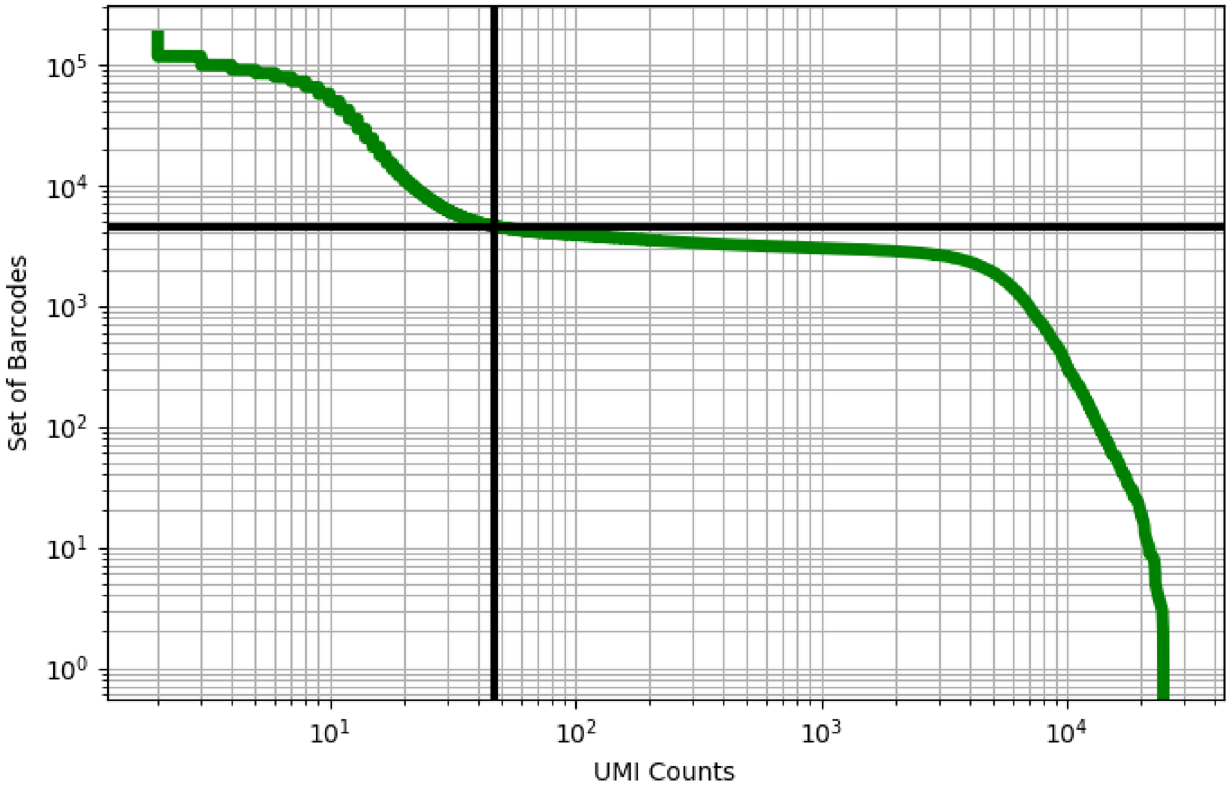


Figure 208: kidney/GSM3509838

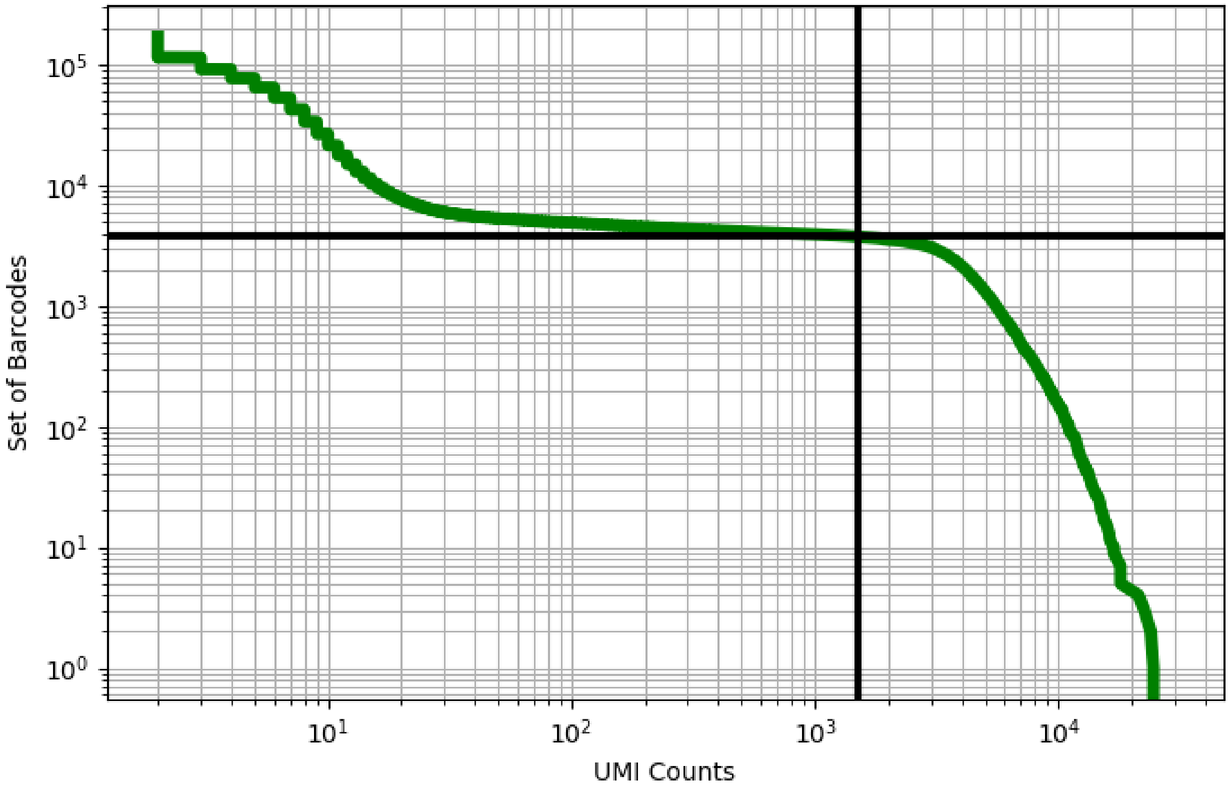


Figure 209: kidney/GSM3509839

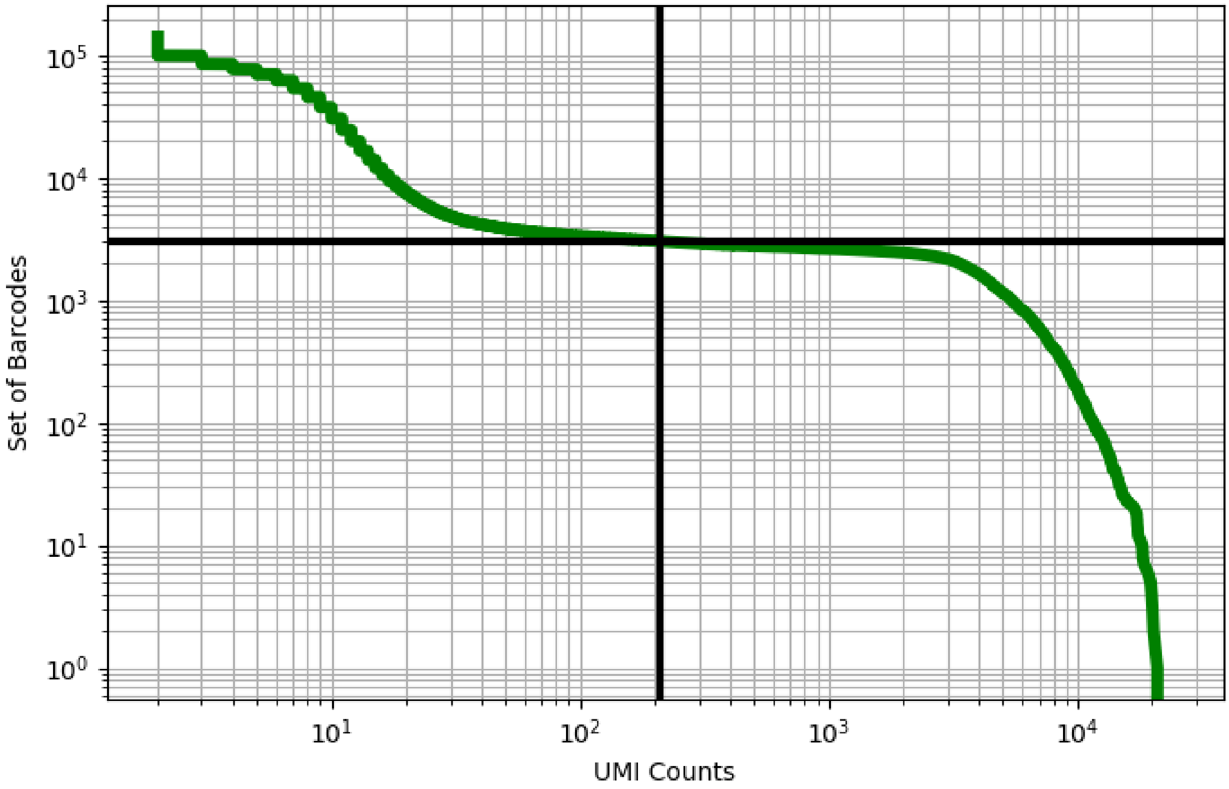


Figure 210: kidney/GSM3509840

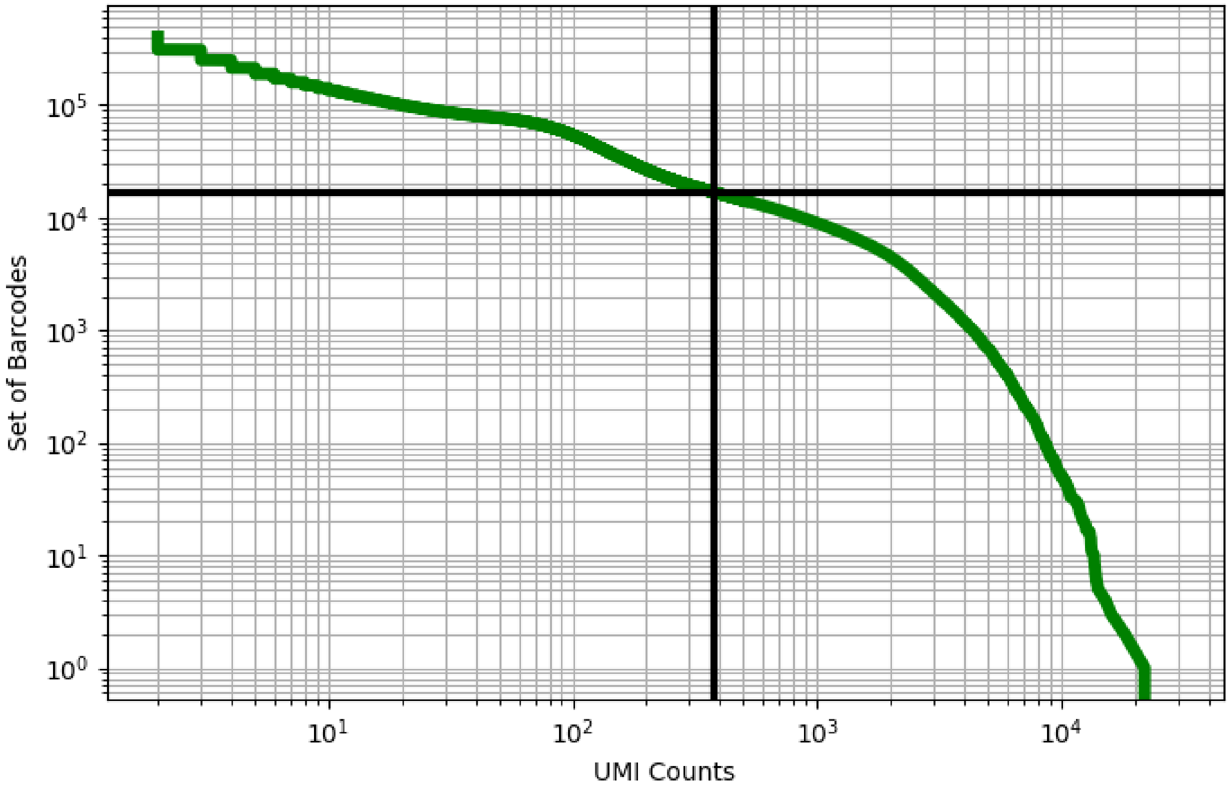


Figure 211: kidney/GSM4145204

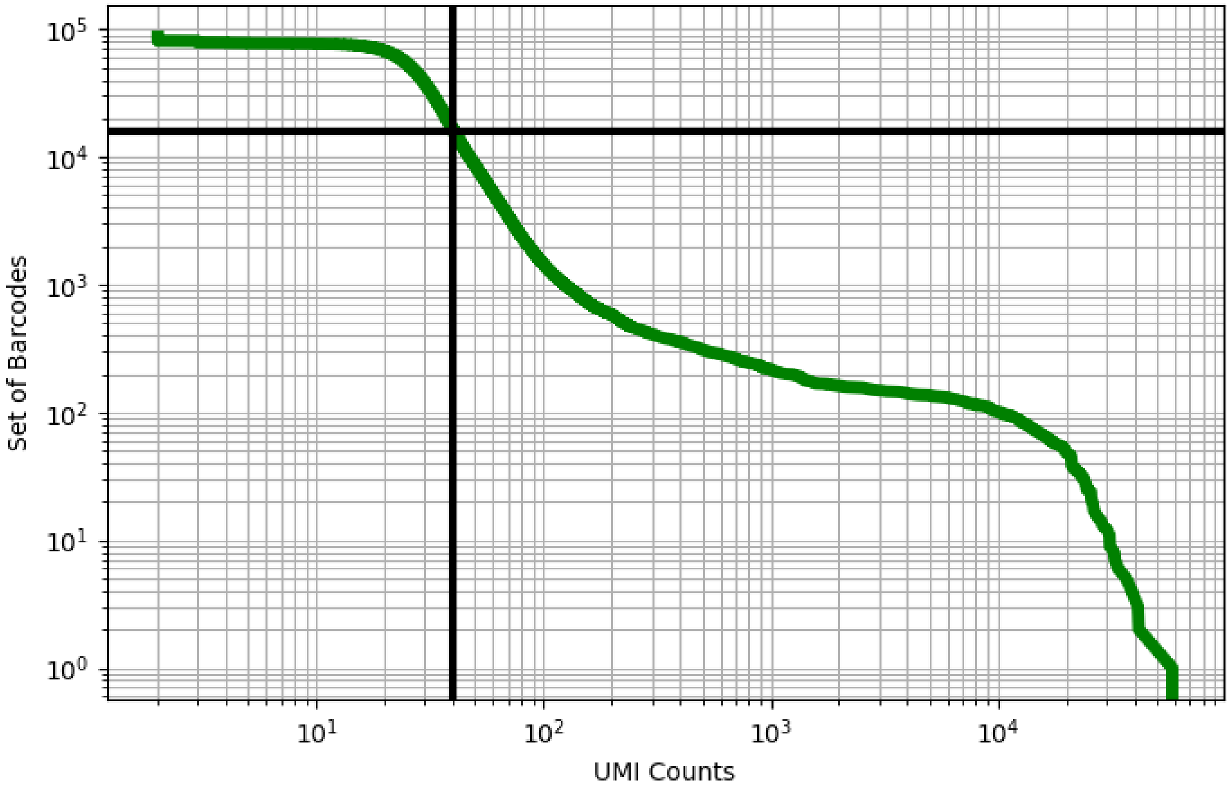


Figure 212: liver/ERS3861773

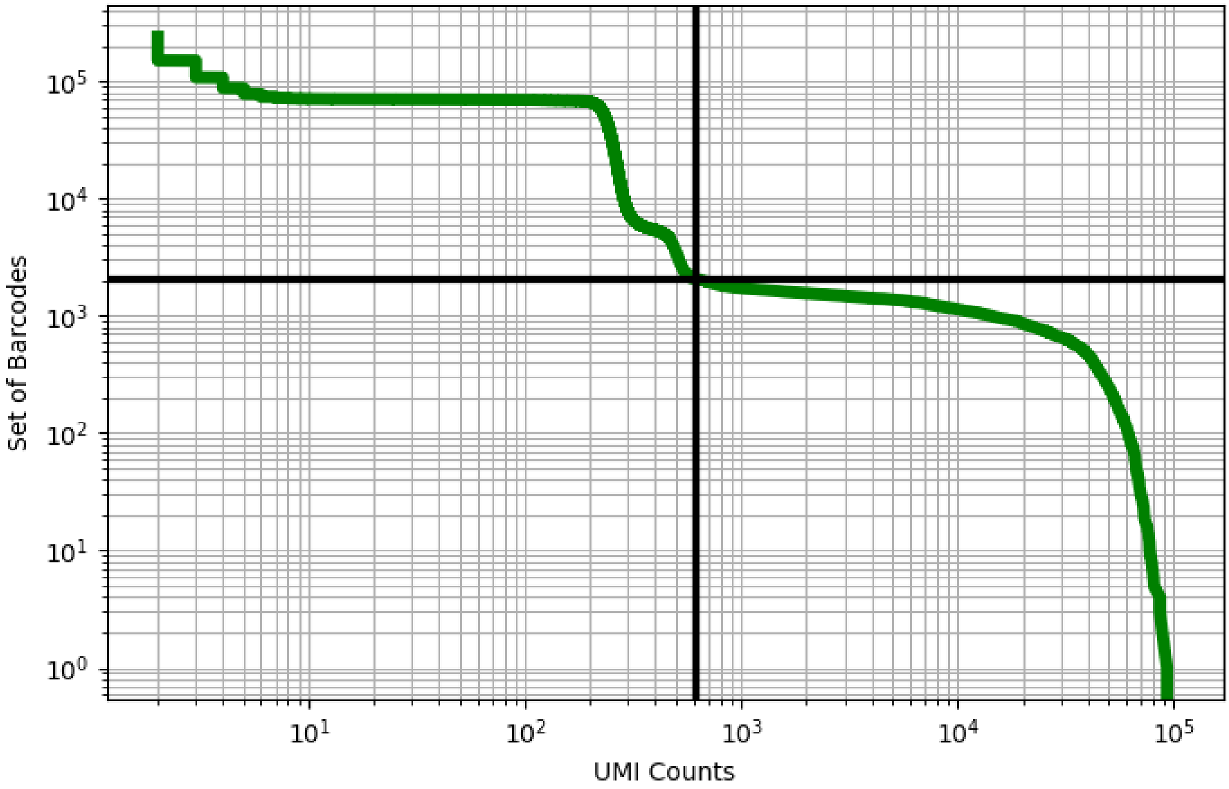


Figure 213: liver/ERS3861774

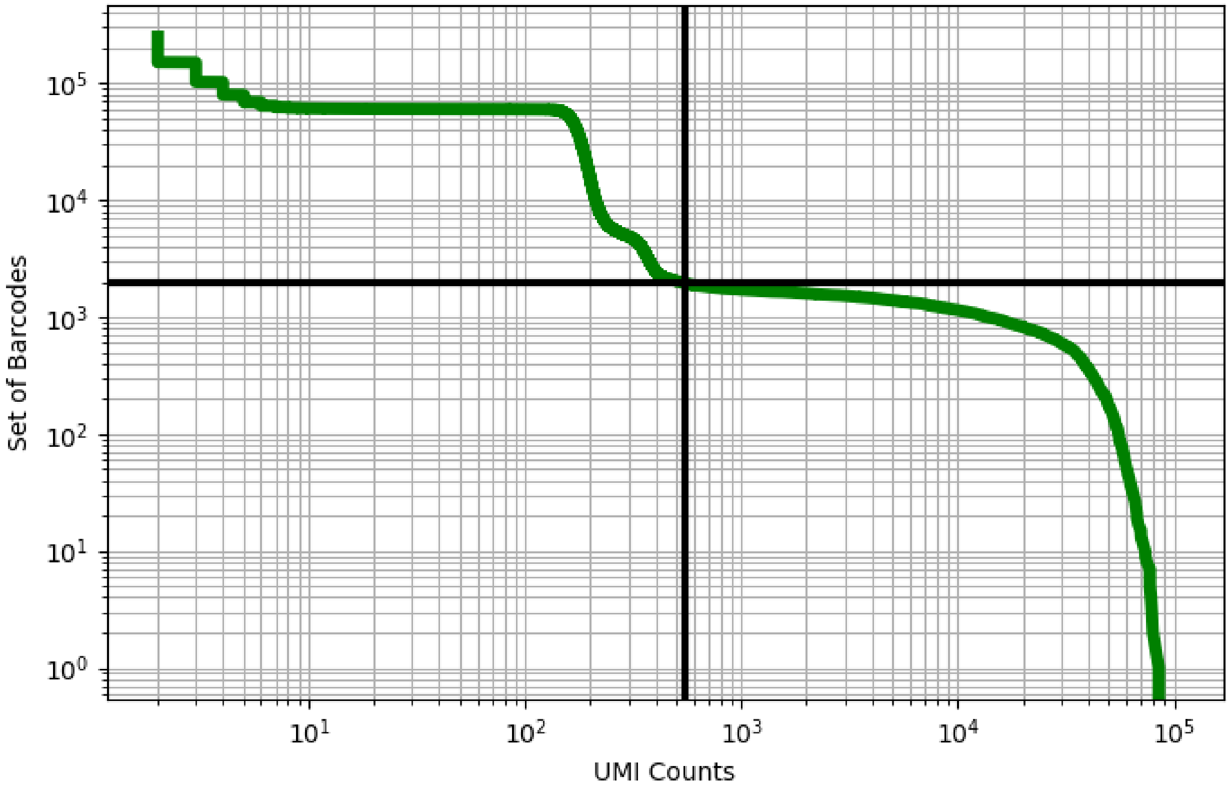


Figure 214: liver/ERS3861778

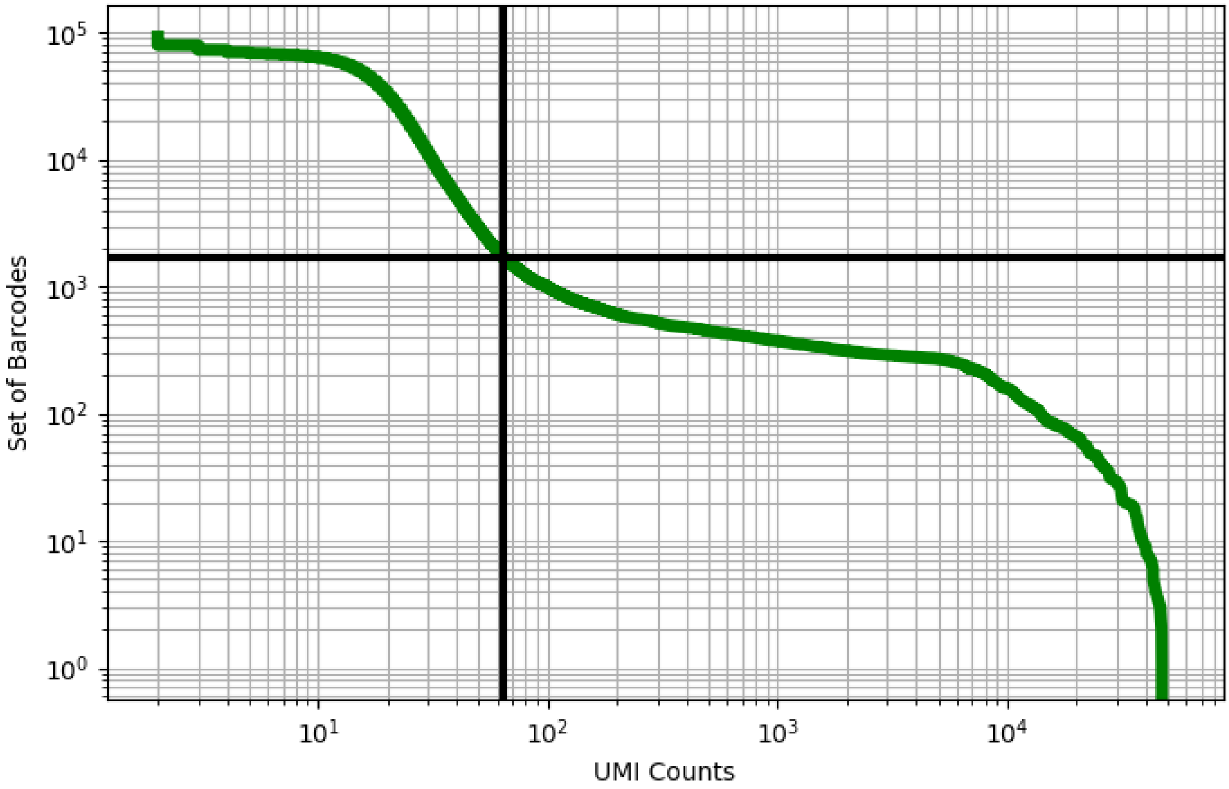


Figure 215: liver/ERS3861782

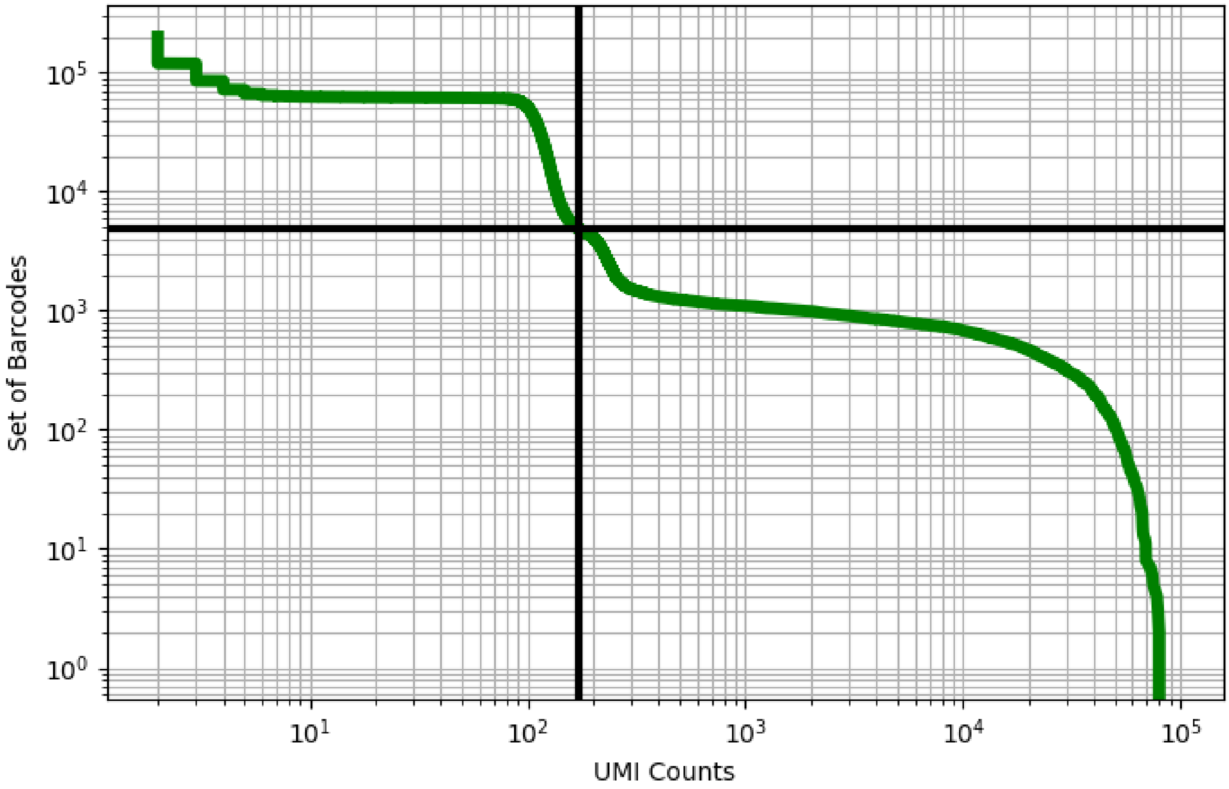


Figure 216: liver/ERS3861783

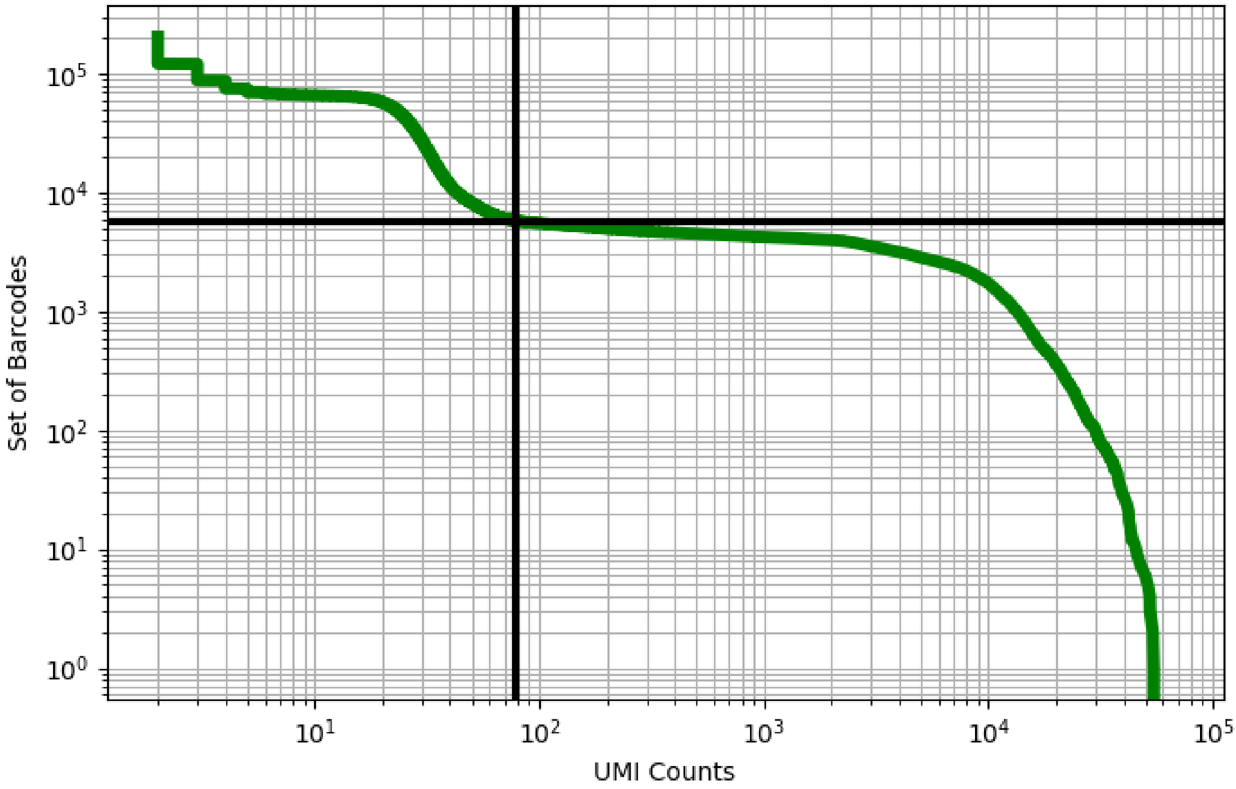


Figure 217: liver/ERS3861786

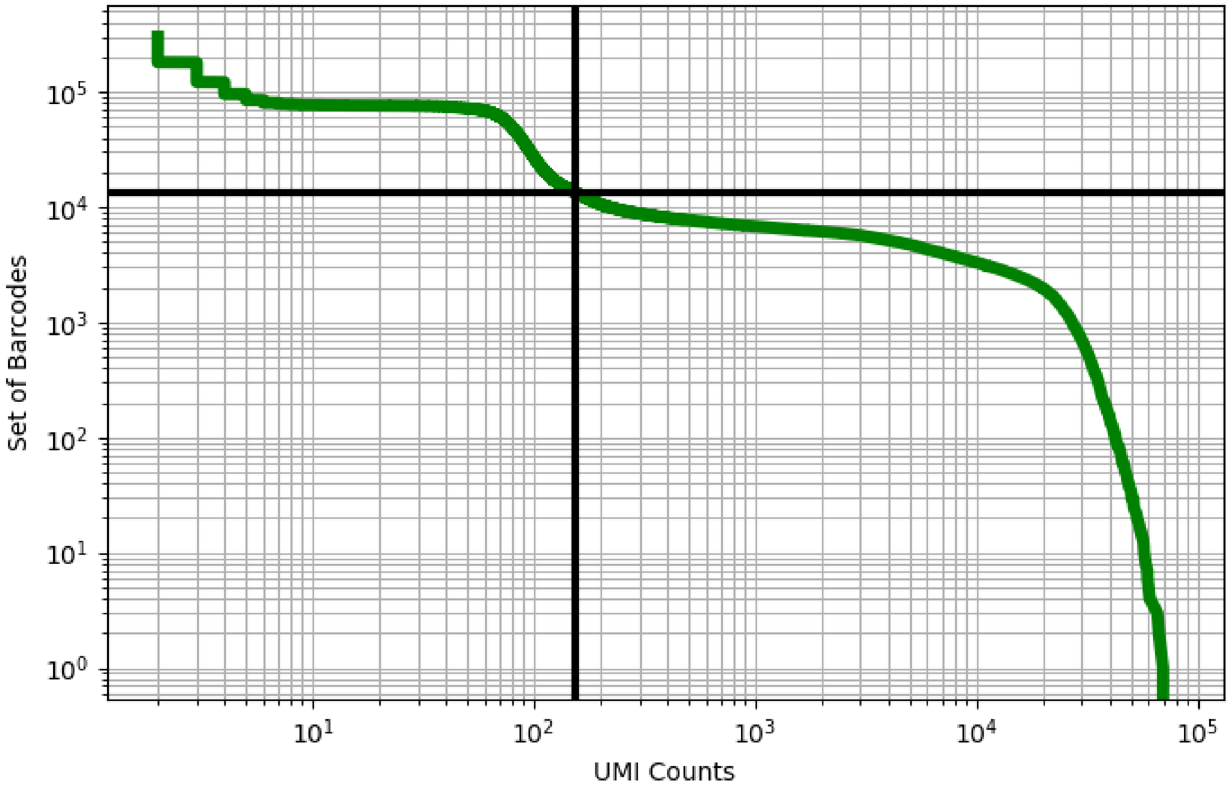


Figure 218: liver/ERS3861787

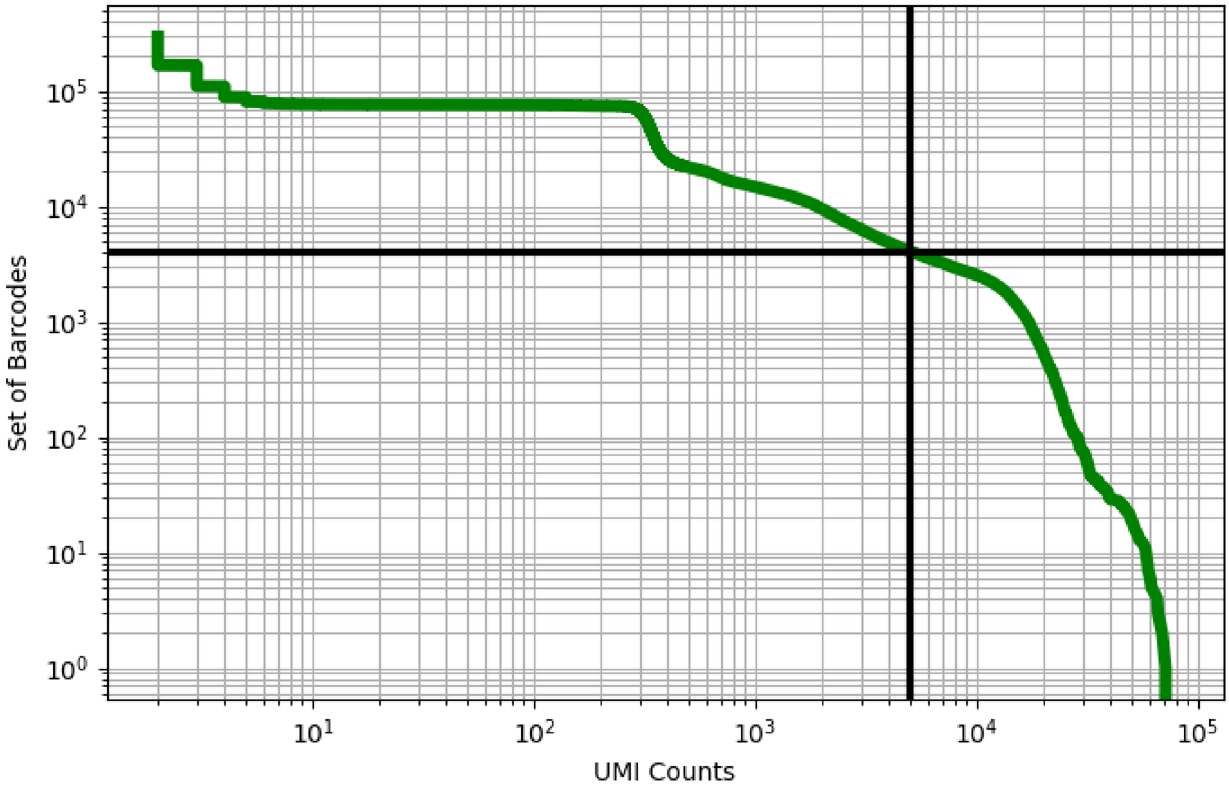


Figure 219: liver/ERS3861788

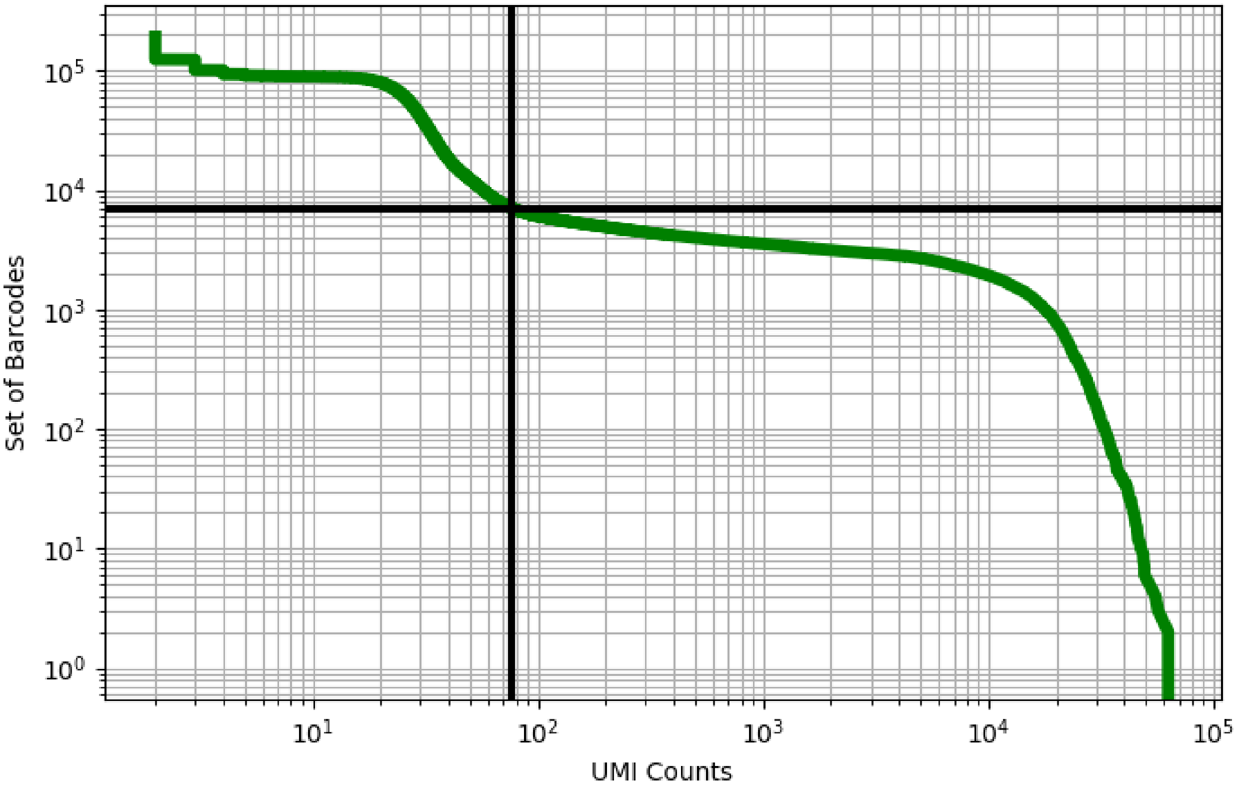


Figure 220: liver/ERS3861789

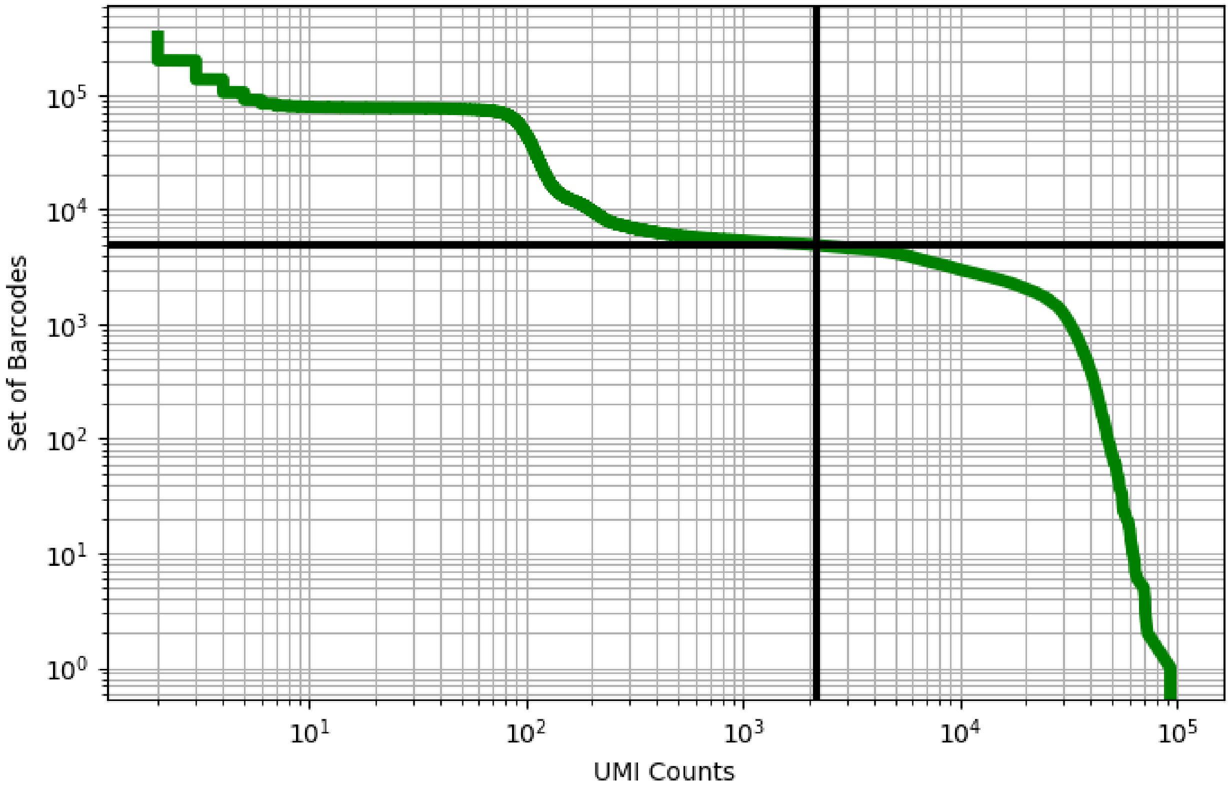


Figure 221: liver/ERS3861790

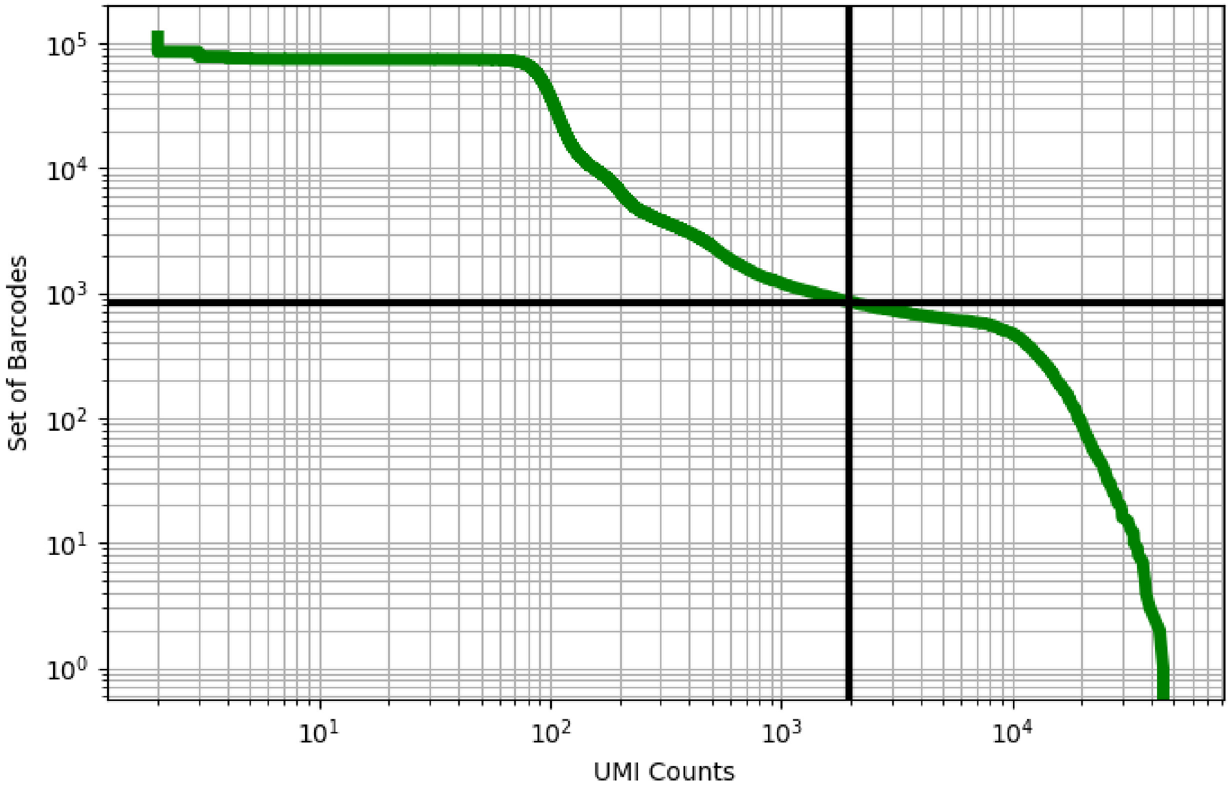


Figure 222: liver/ERS3861791

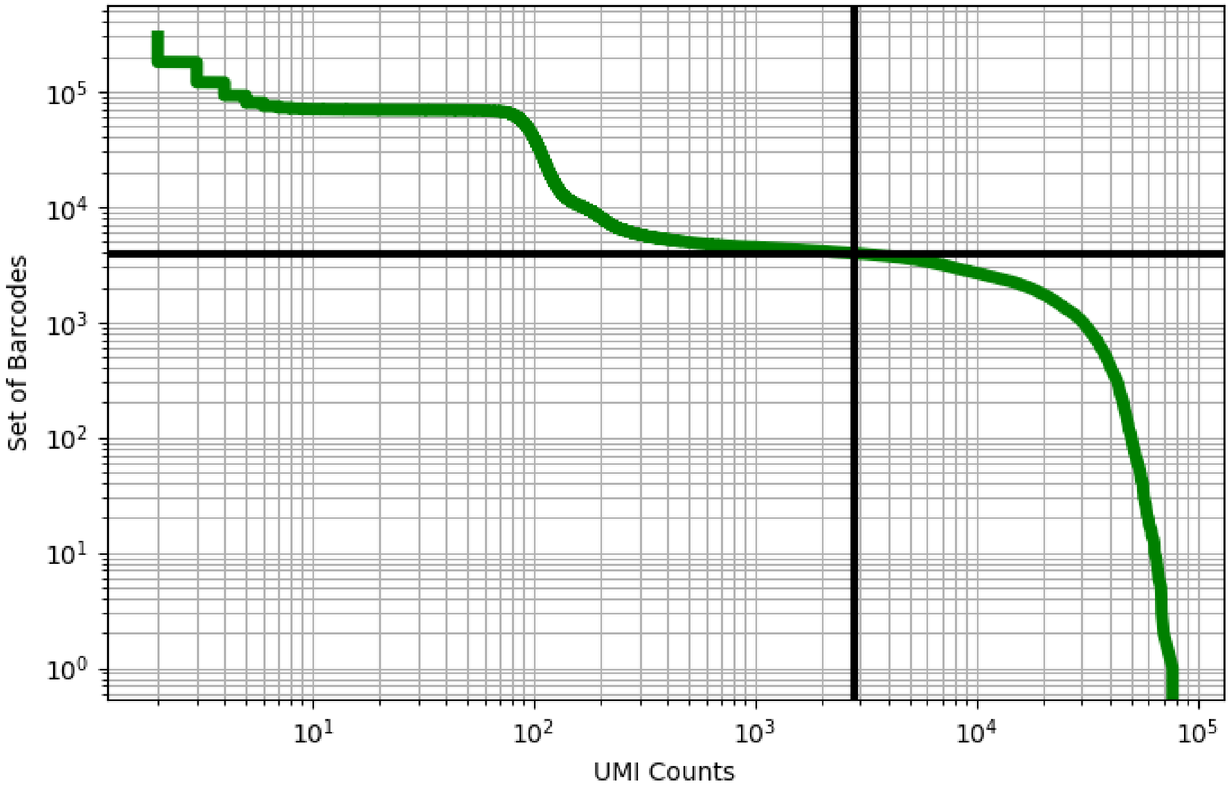


Figure 223: liver/ERS3861792

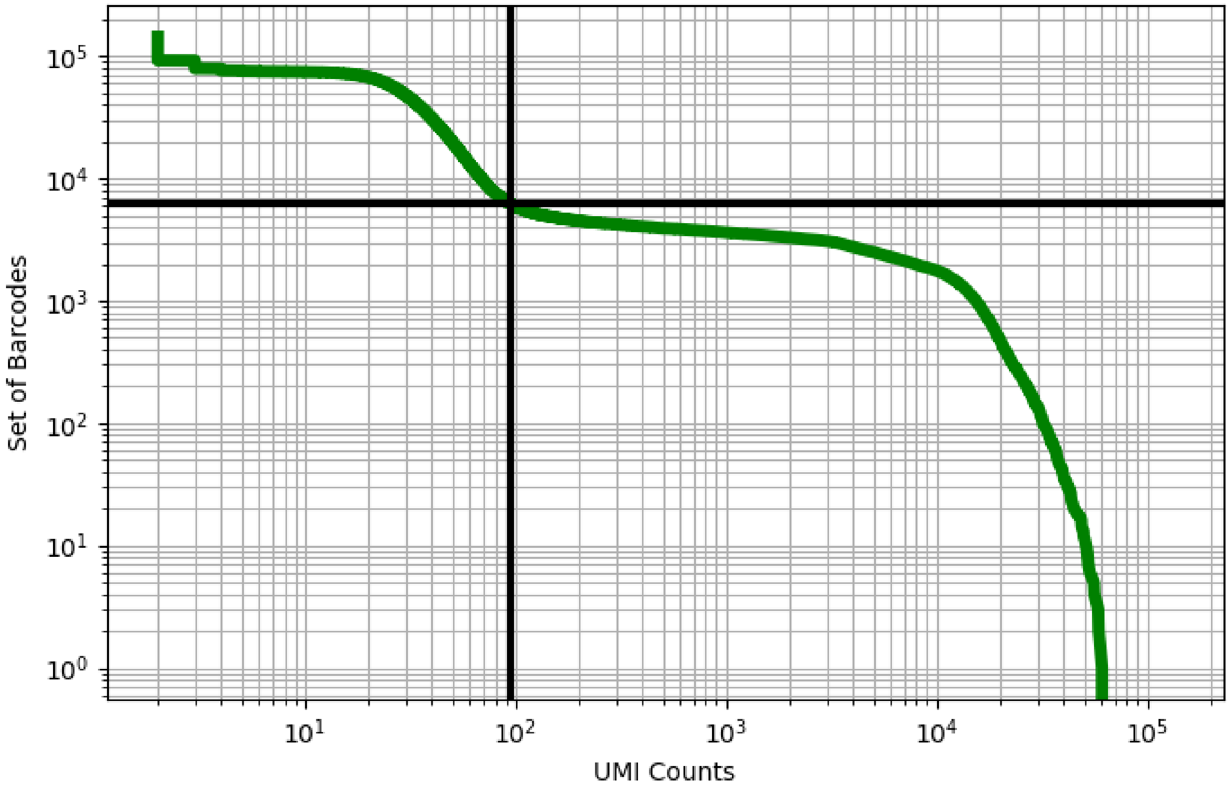


Figure 224: liver/ERS3861795

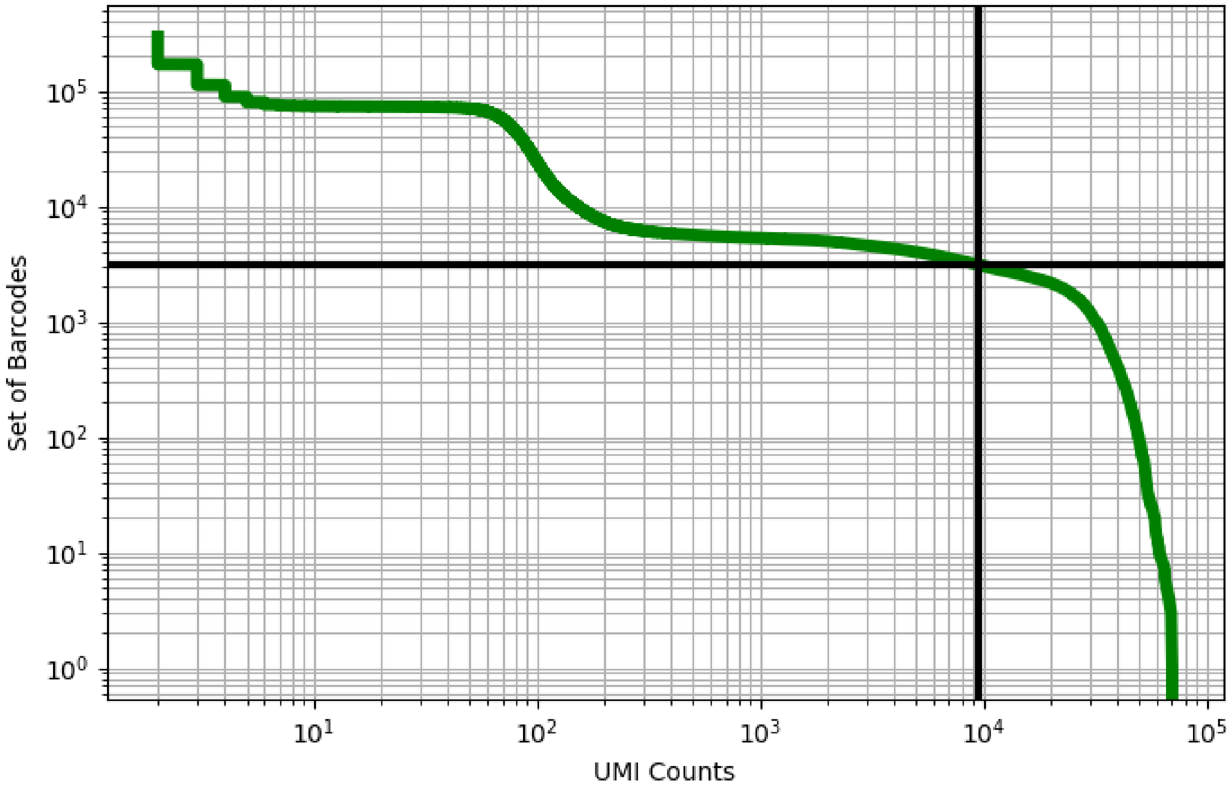


Figure 225: liver/ERS3861796

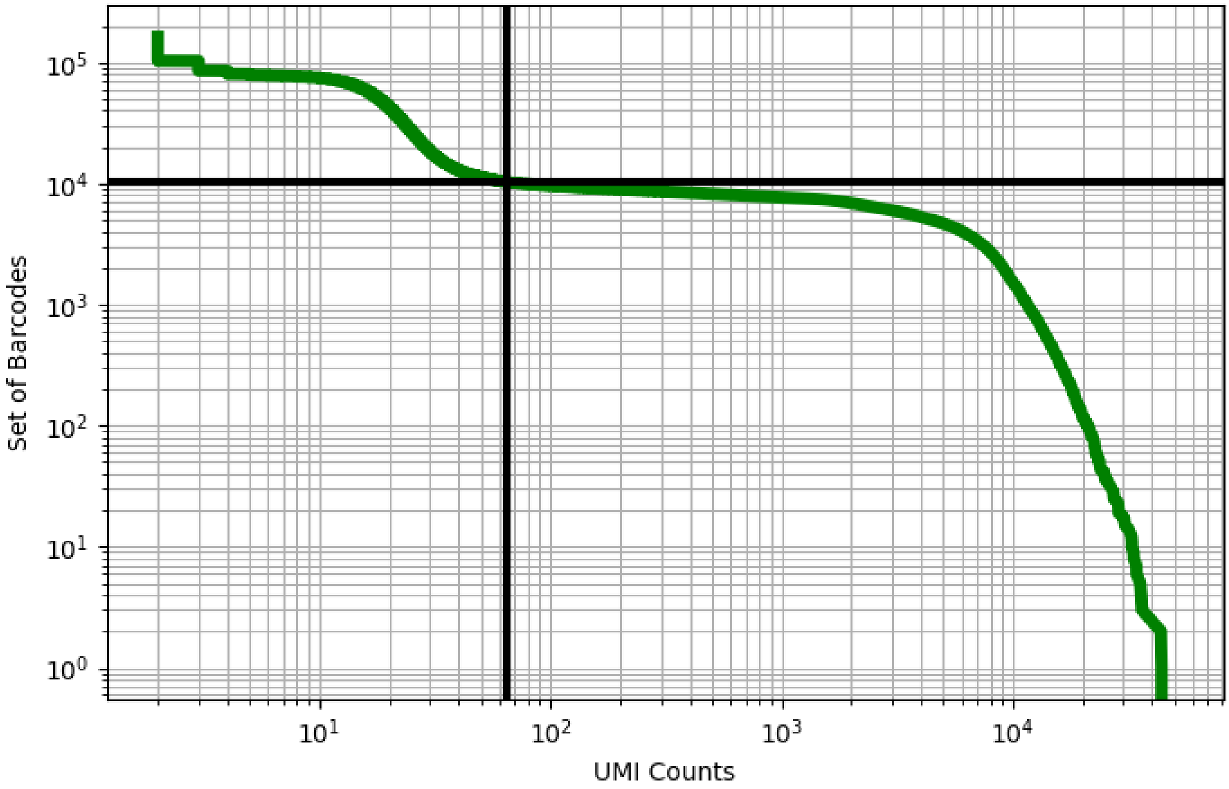


Figure 226: liver/ERS3861797

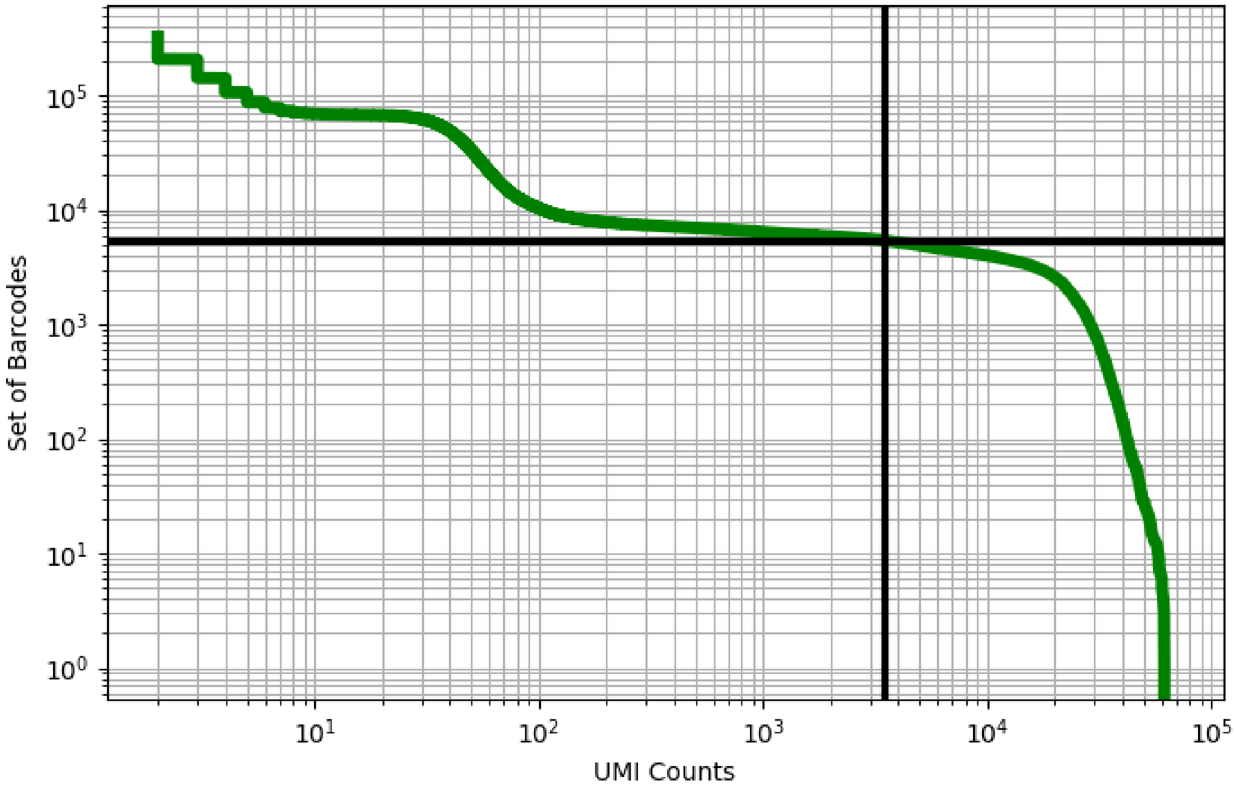


Figure 227: liver/ERS3861798

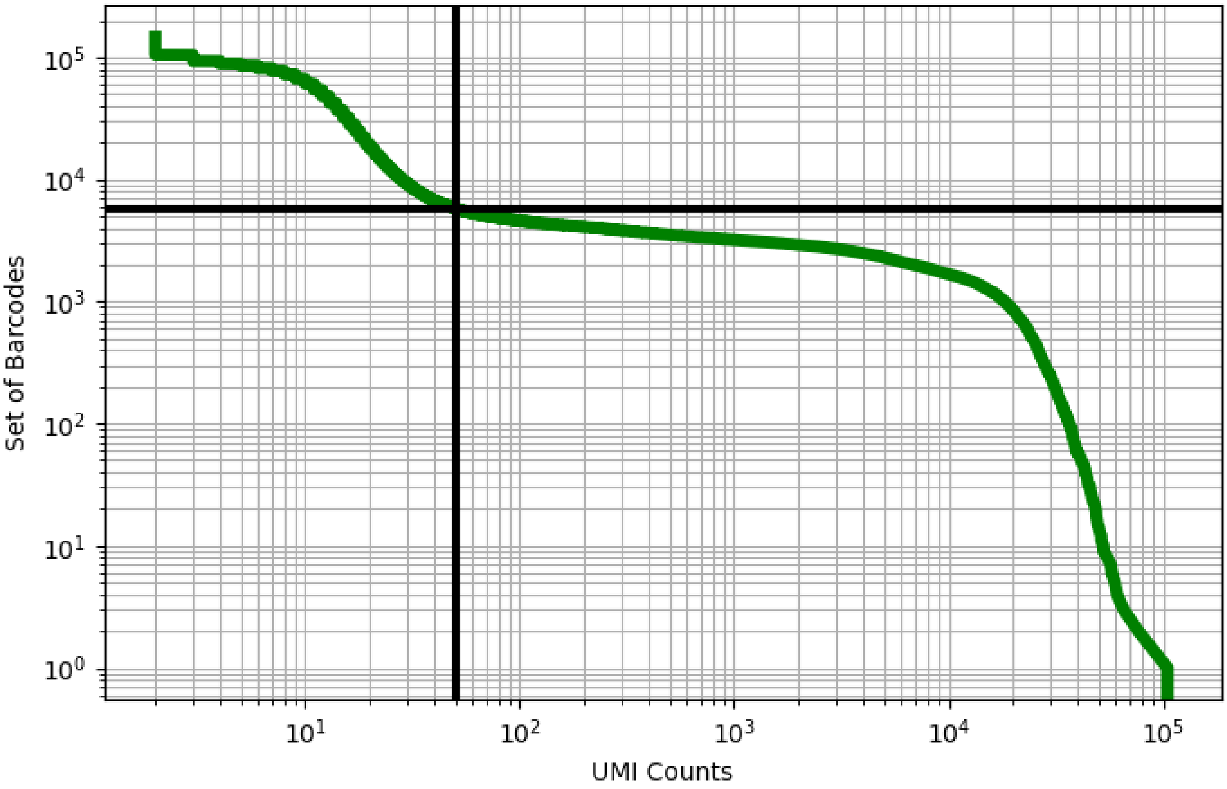


Figure 228: liver/ERS3861799

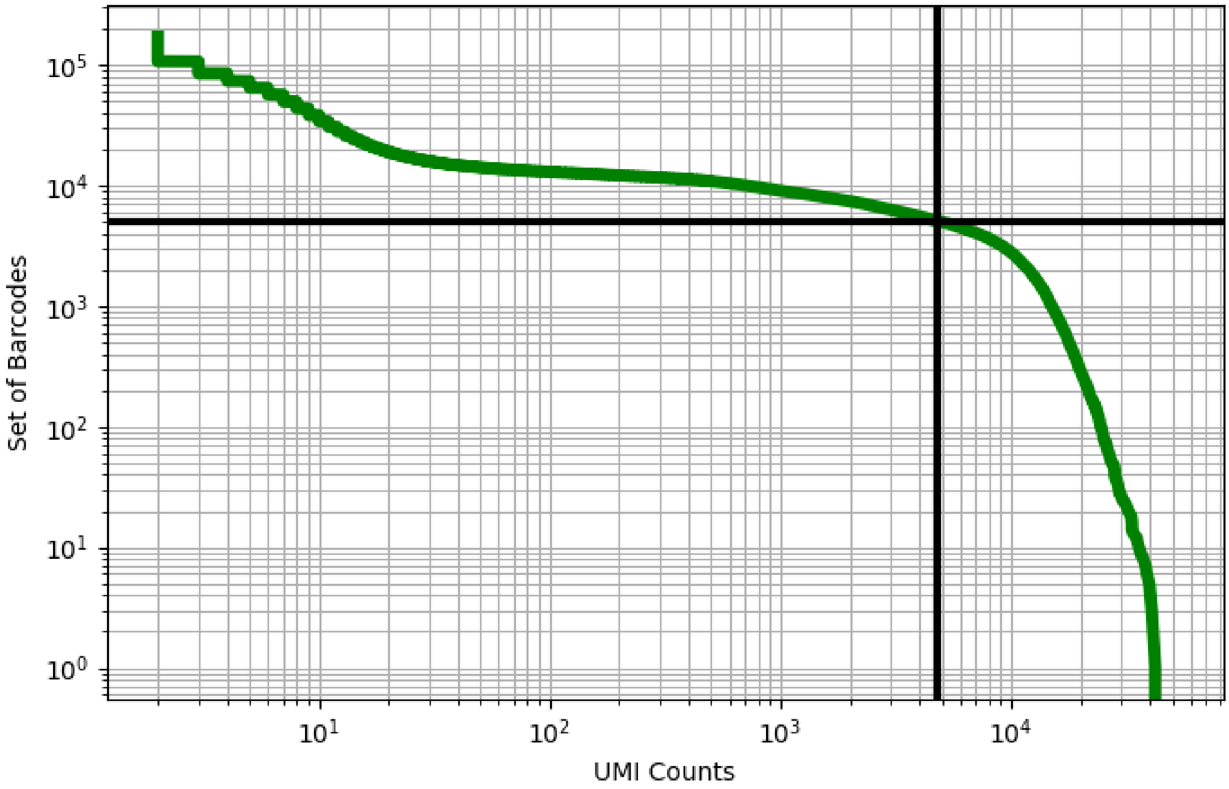


Figure 229: liver/ERS3861800

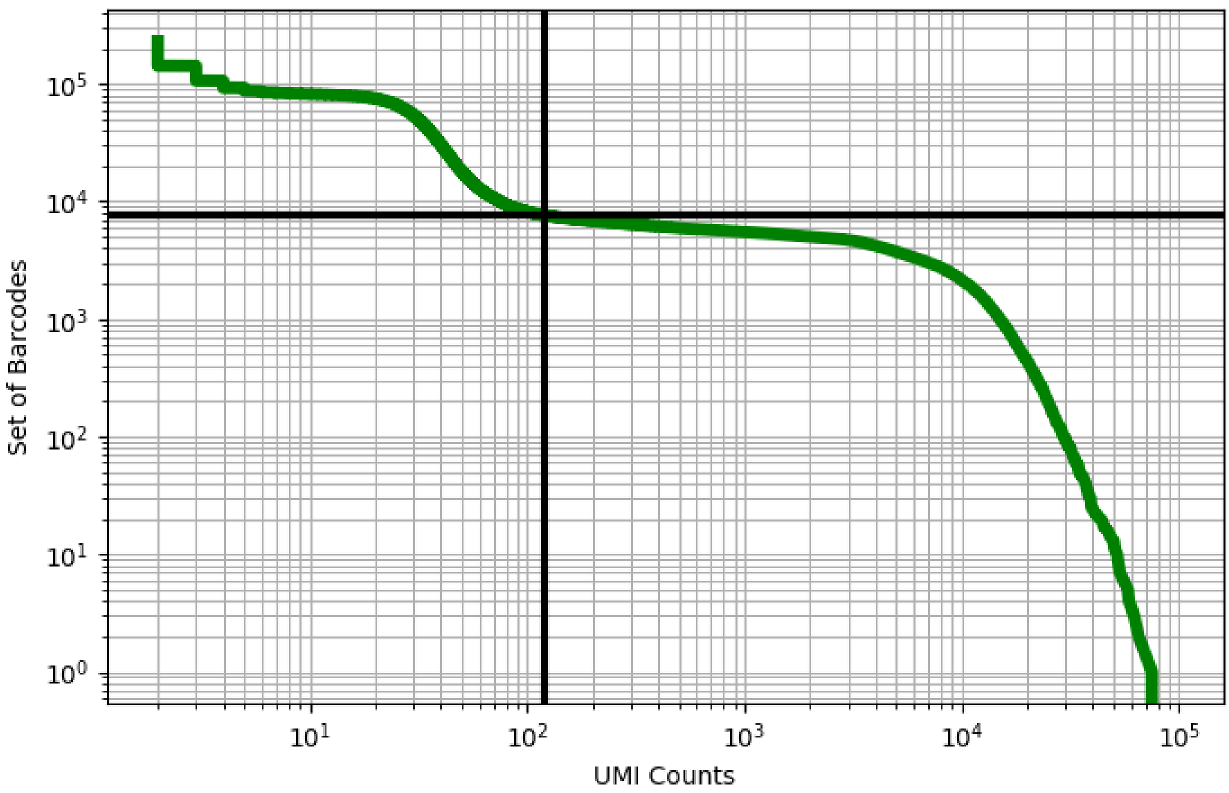


Figure 230: liver/ERS3861804

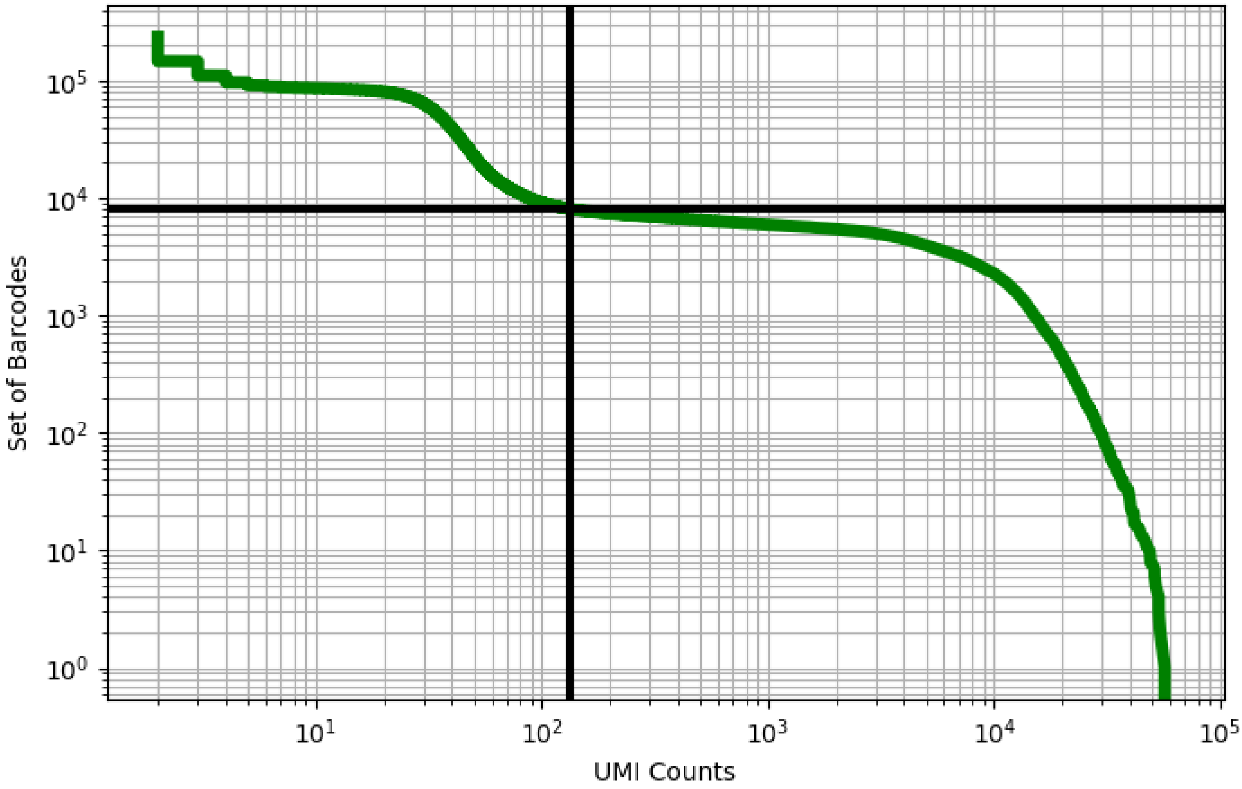


Figure 231: liver/ERS3861805

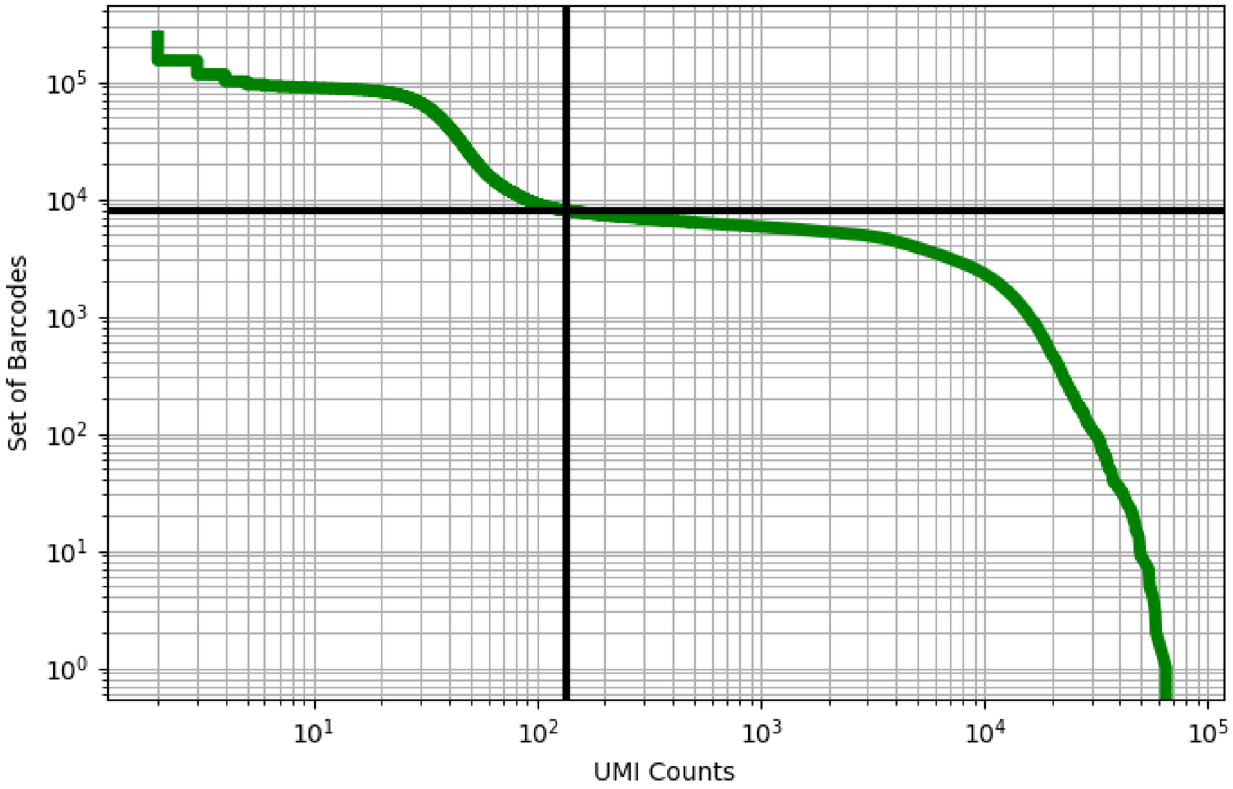


Figure 232: liver/ERS3861806

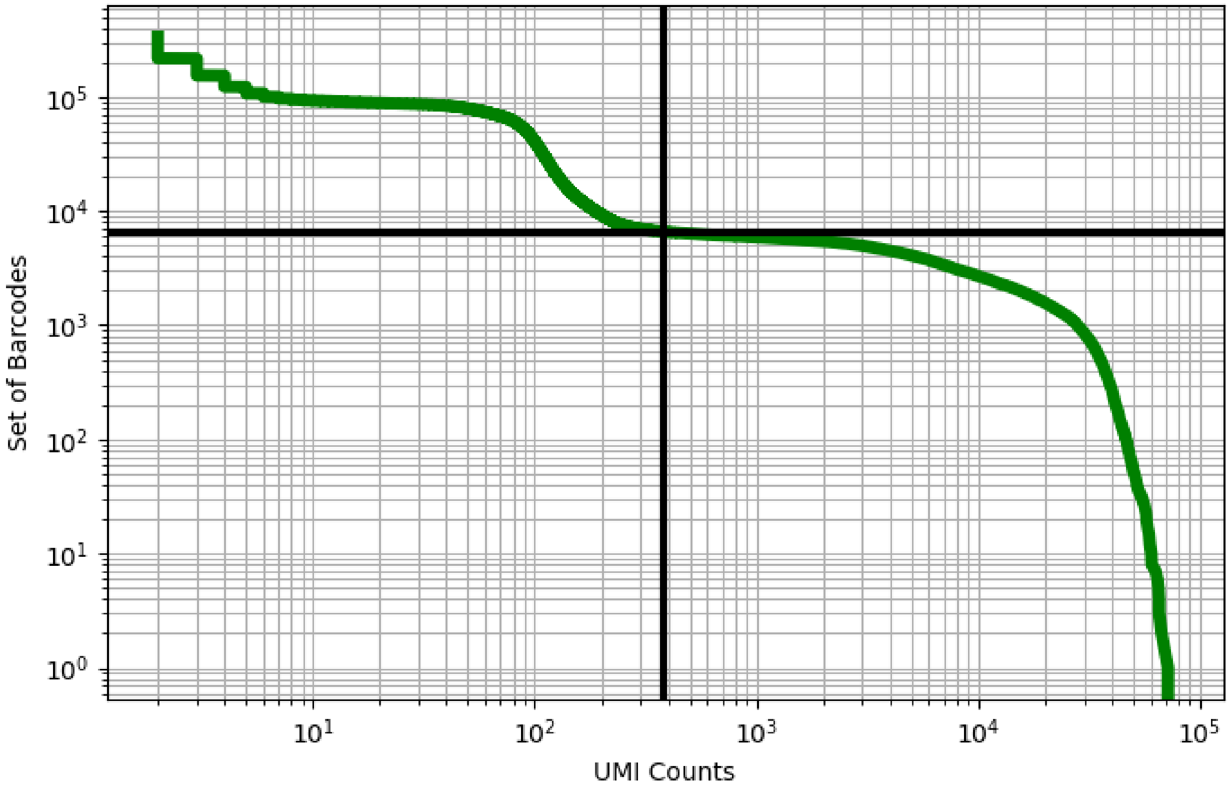


Figure 233: liver/ERS3861807

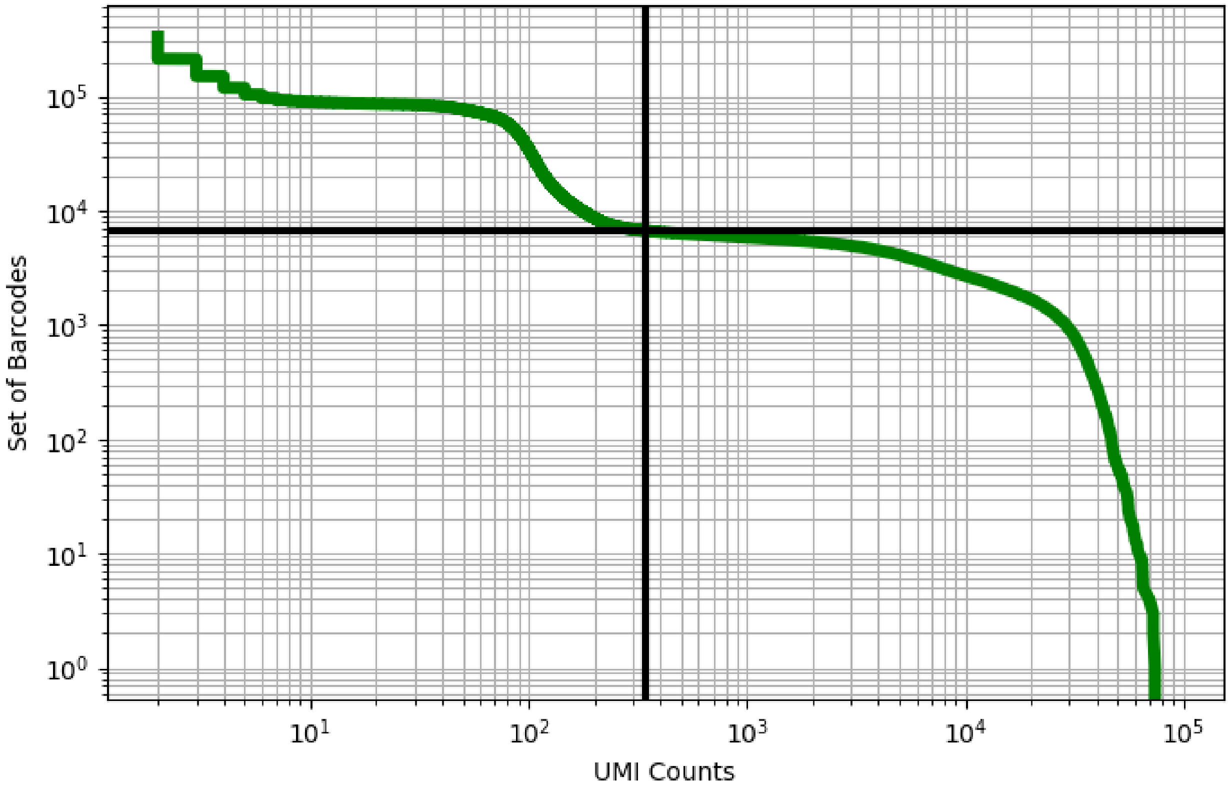


Figure 234: liver/ERS3861808

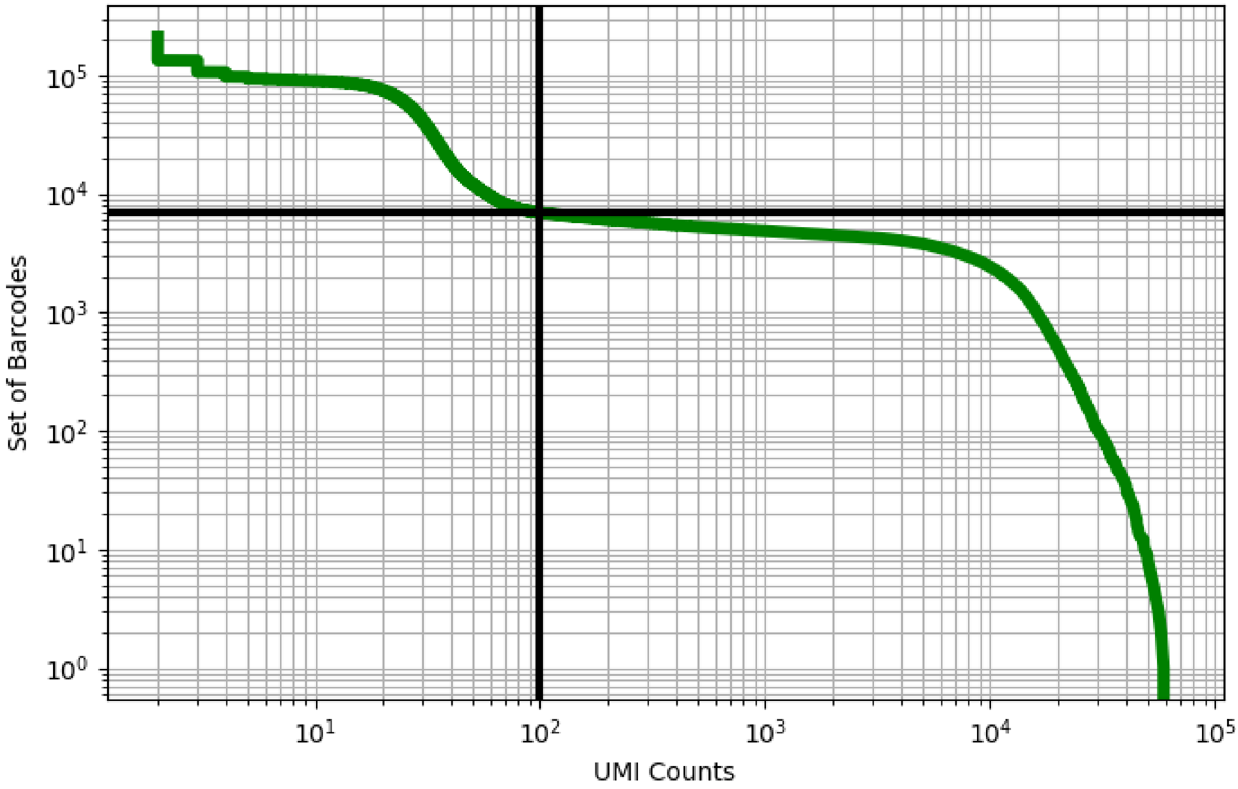


Figure 235: liver/ERS3861809

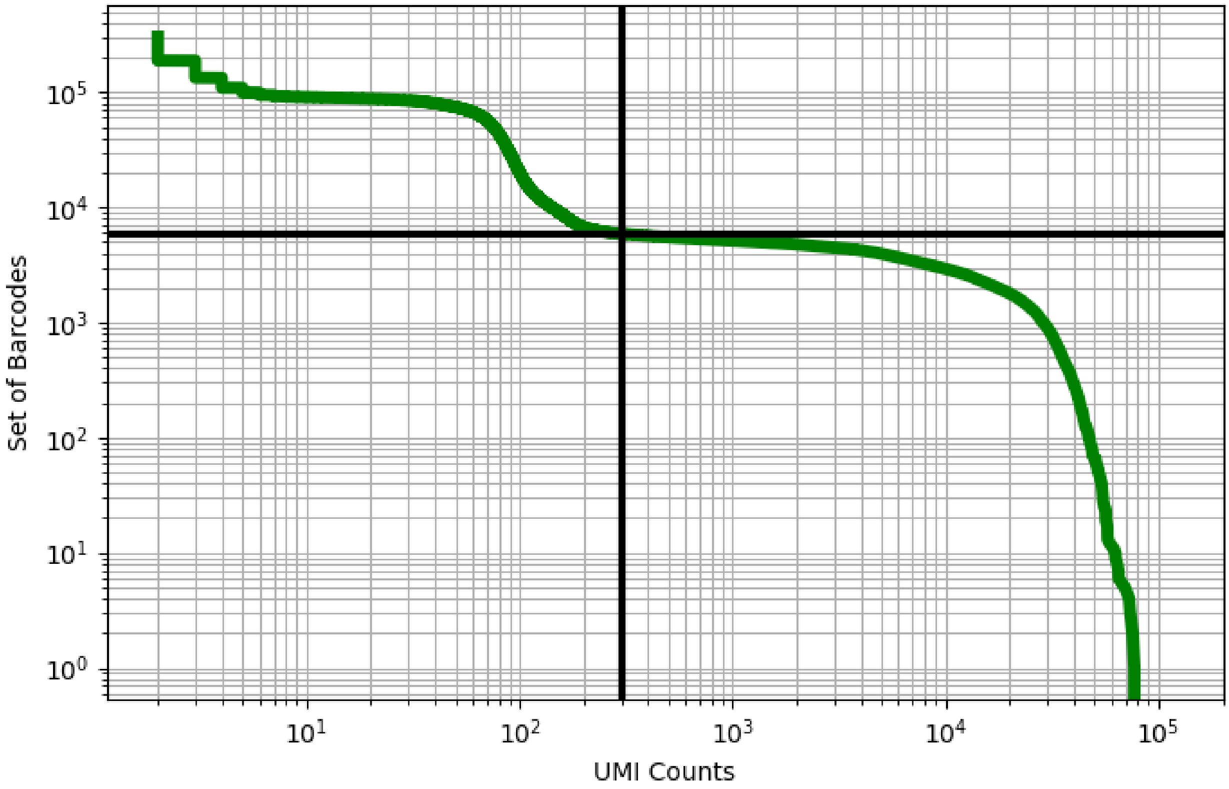


Figure 236: liver/ERS3861810

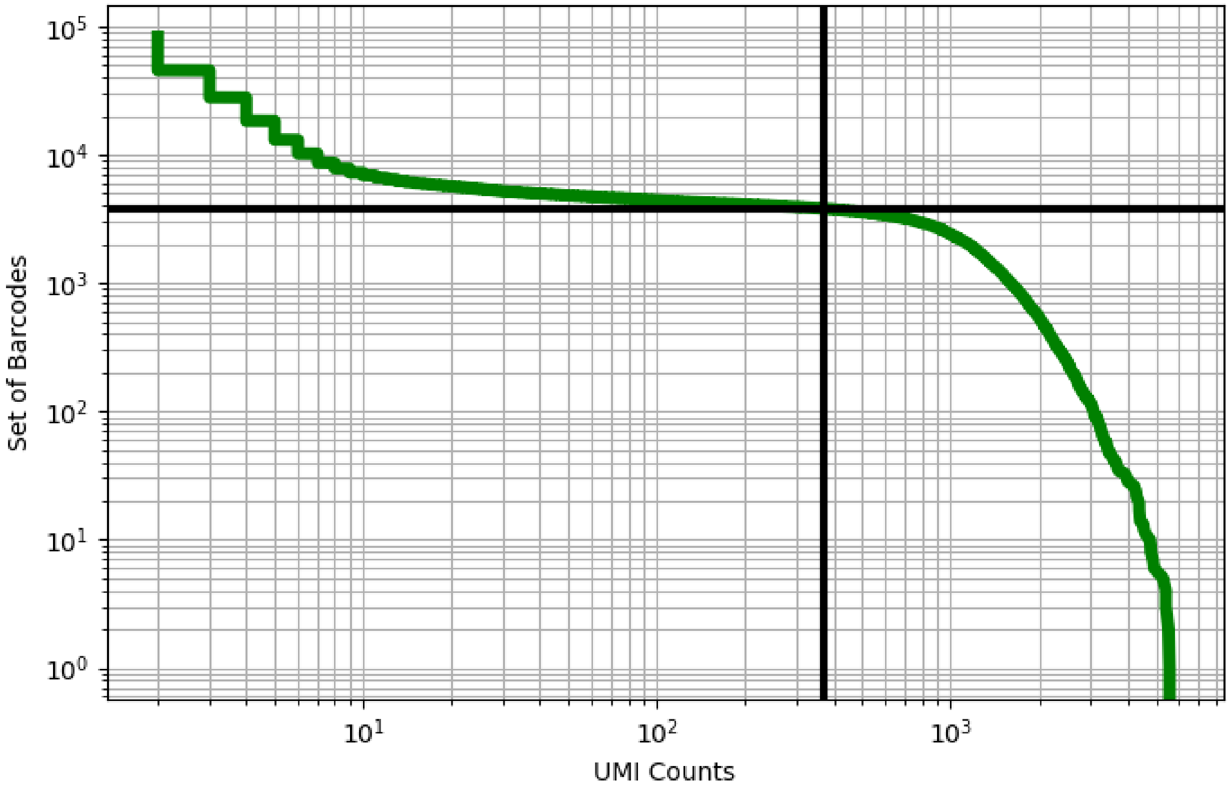


Figure 237: liver/ERS3861814

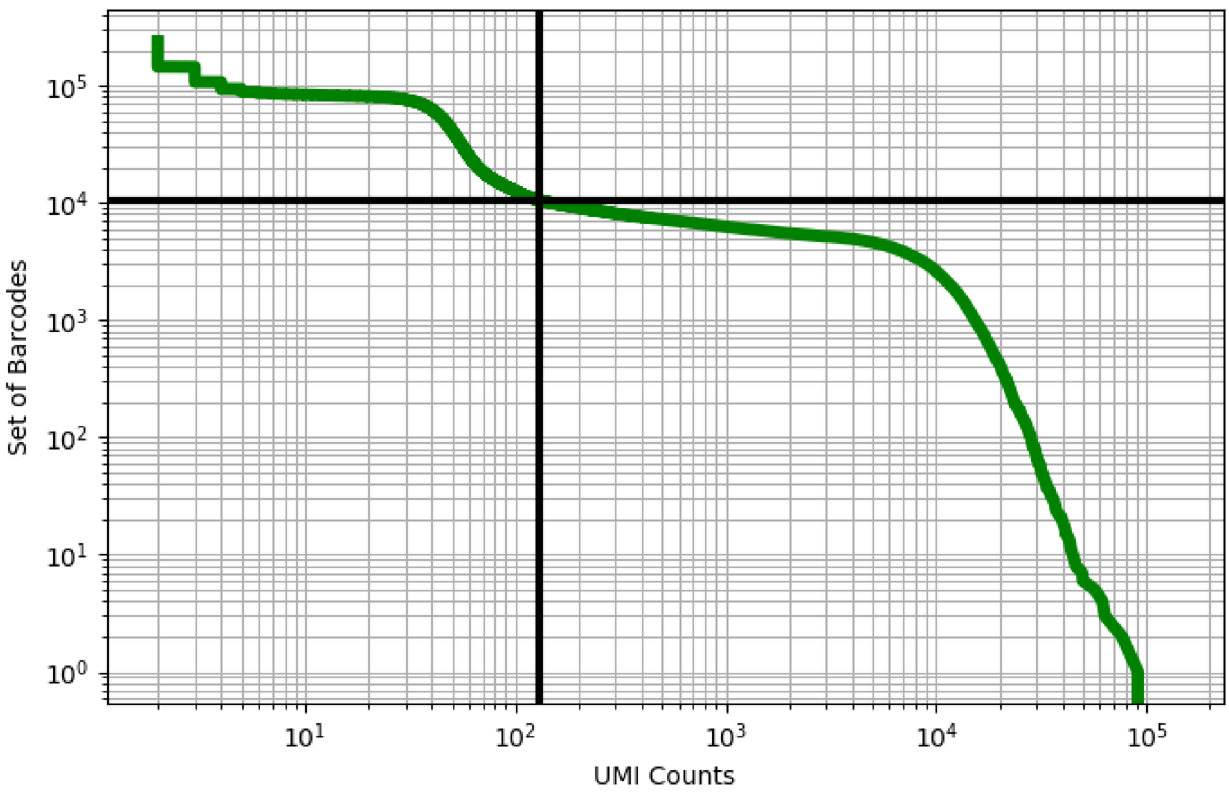


Figure 238: liver/ERS3861818

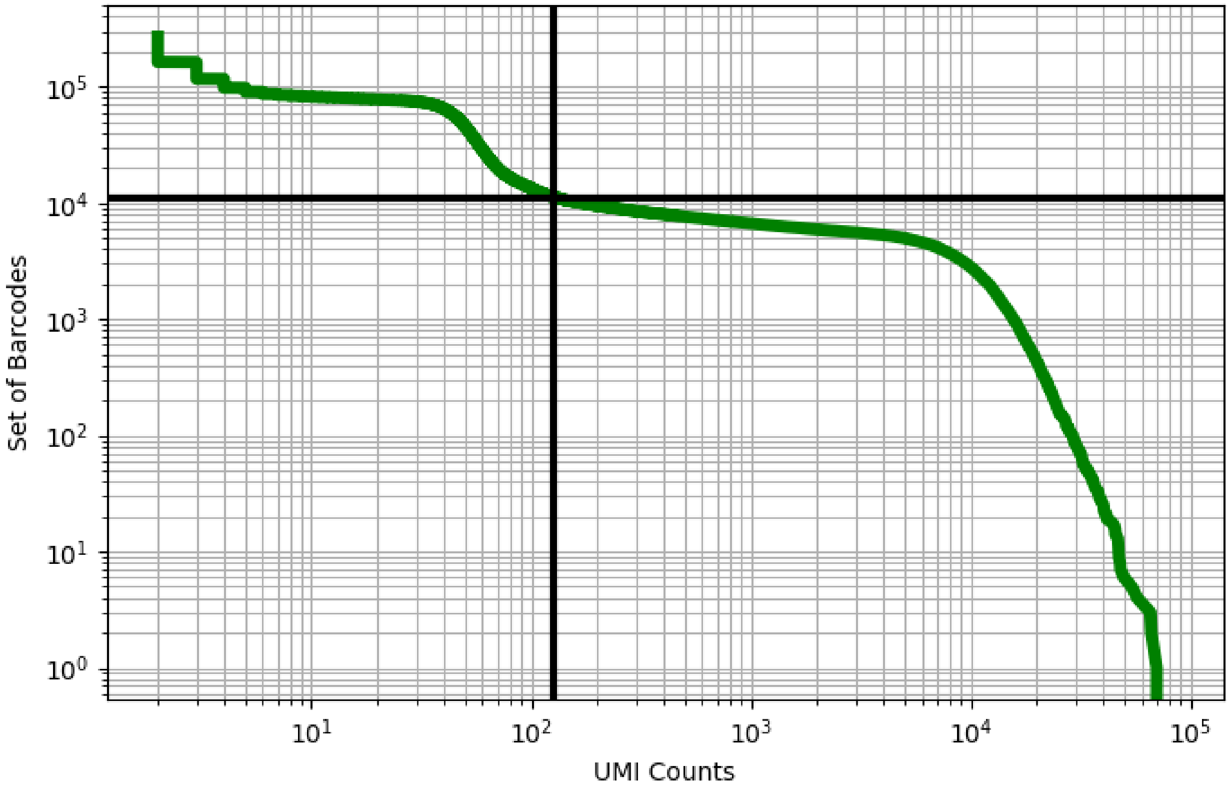


Figure 239: liver/ERS3861819

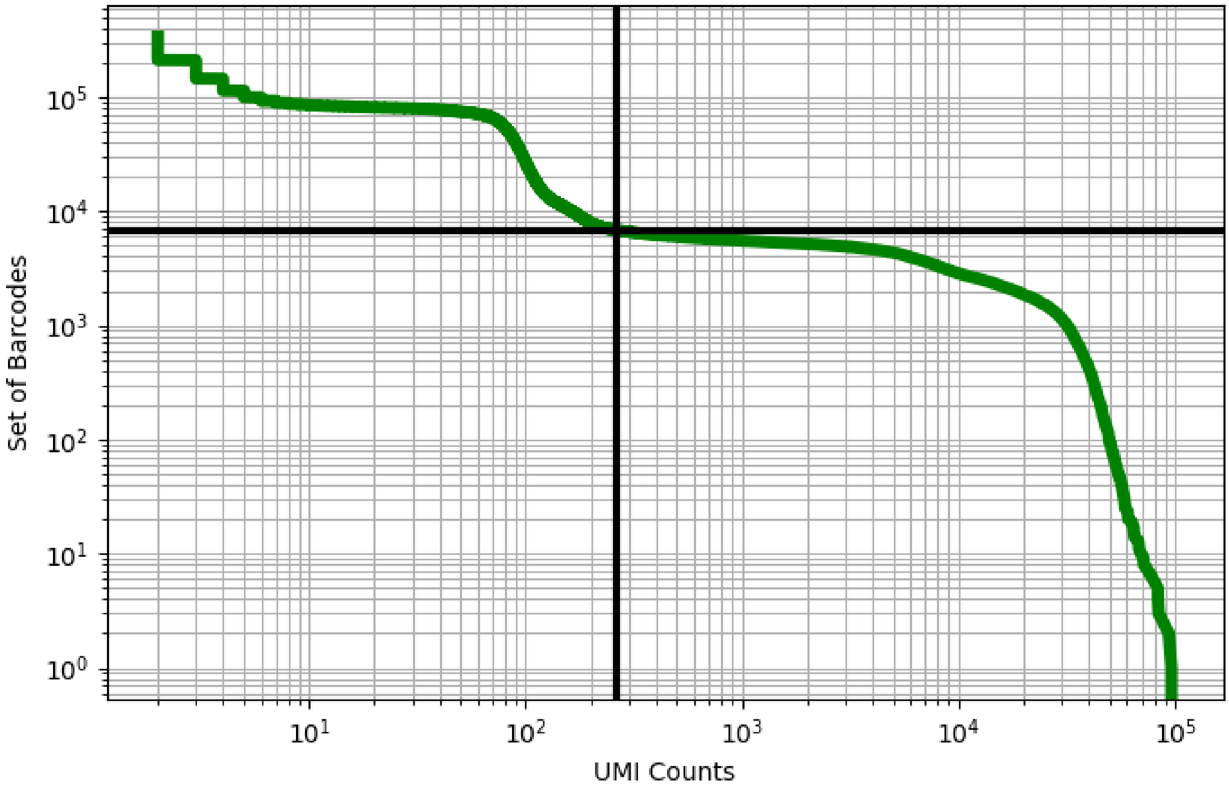


Figure 240: liver/ERS3861820

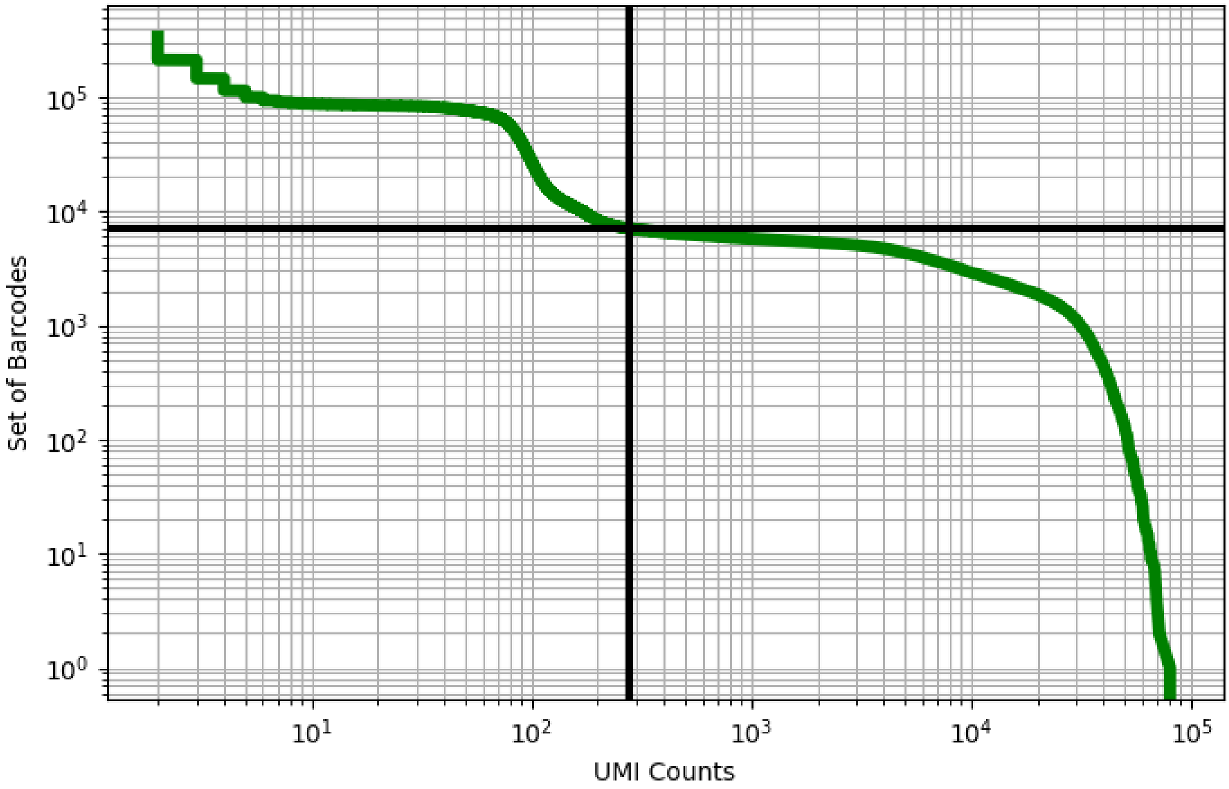


Figure 241: liver/ERS3861821

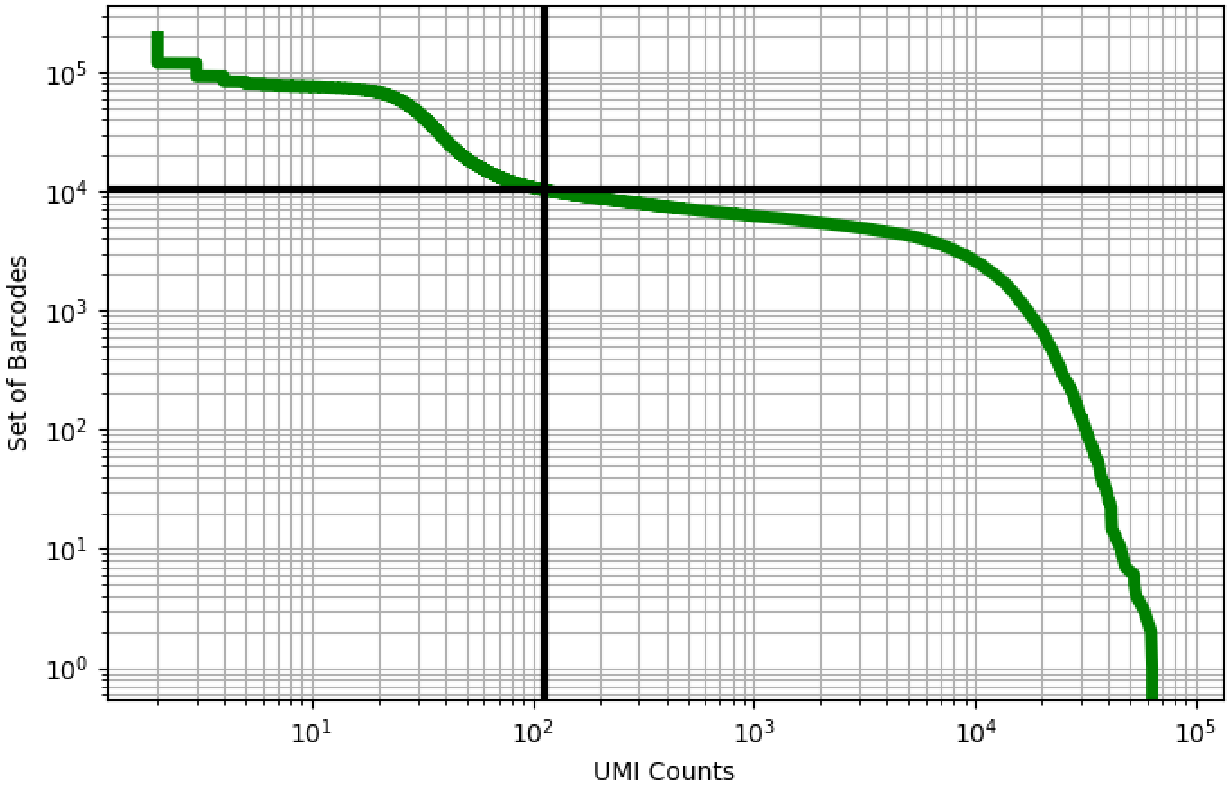


Figure 242: liver/ERS3861822

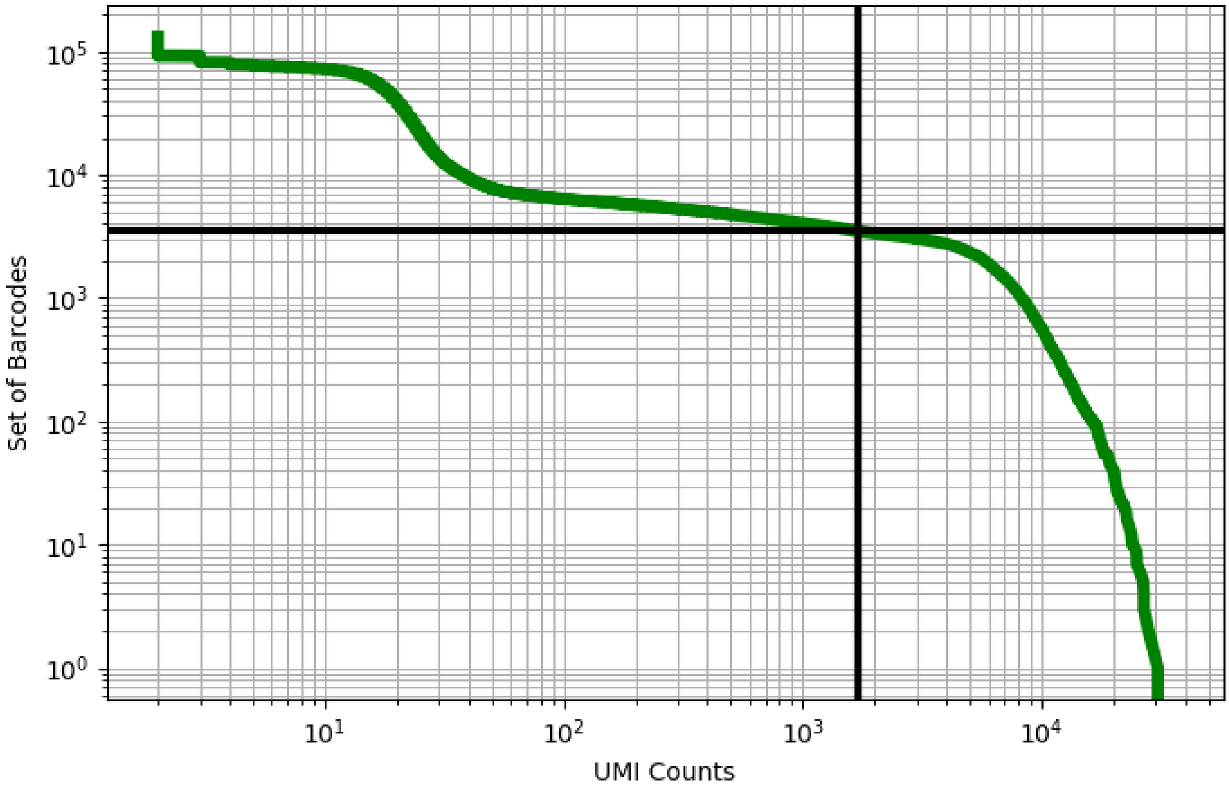

Figure 243: liver/ERS3861823

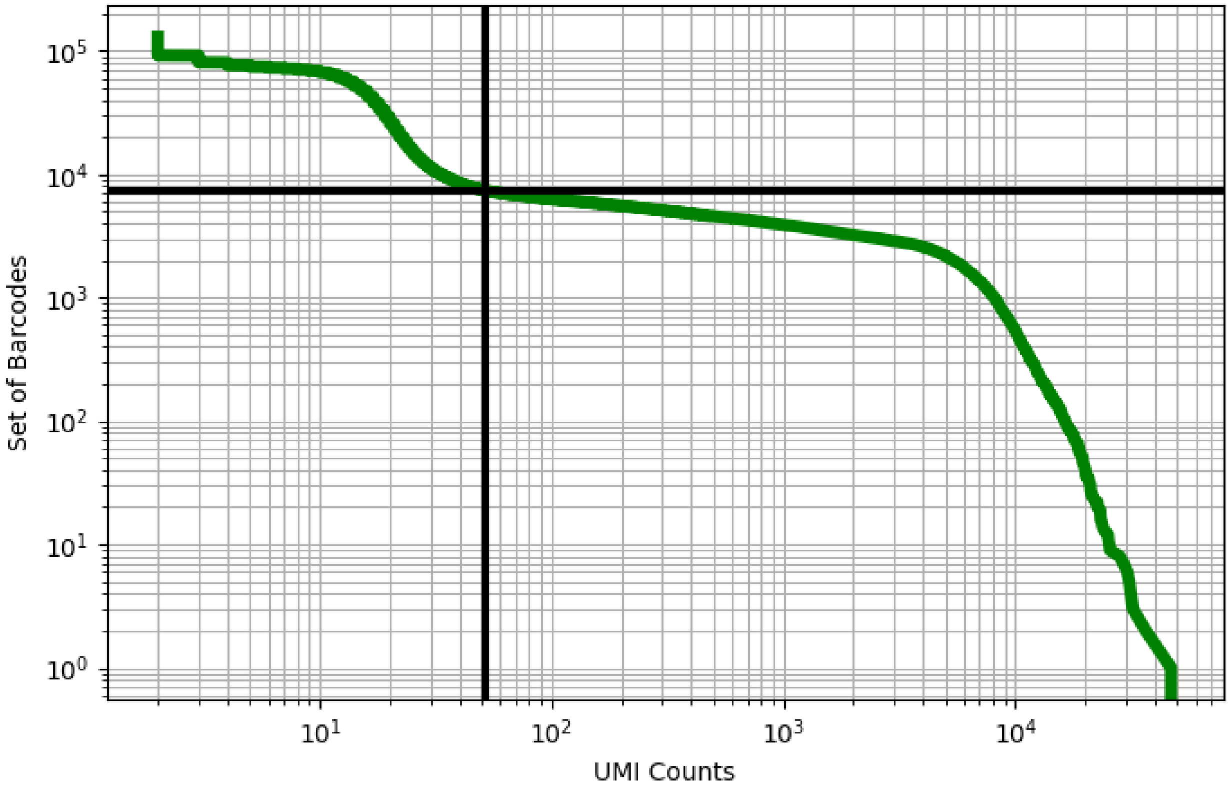


Figure 244: liver/ERS3861824

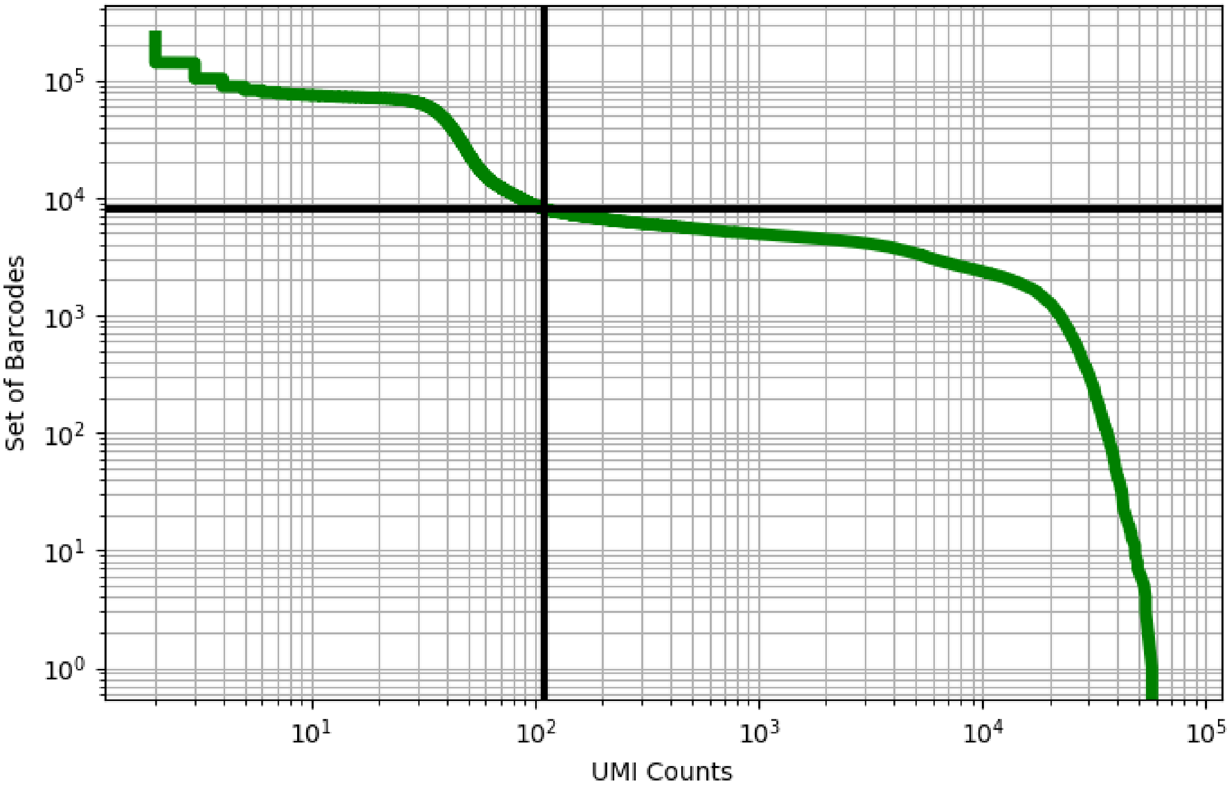


Figure 245: liver/ERS3861825

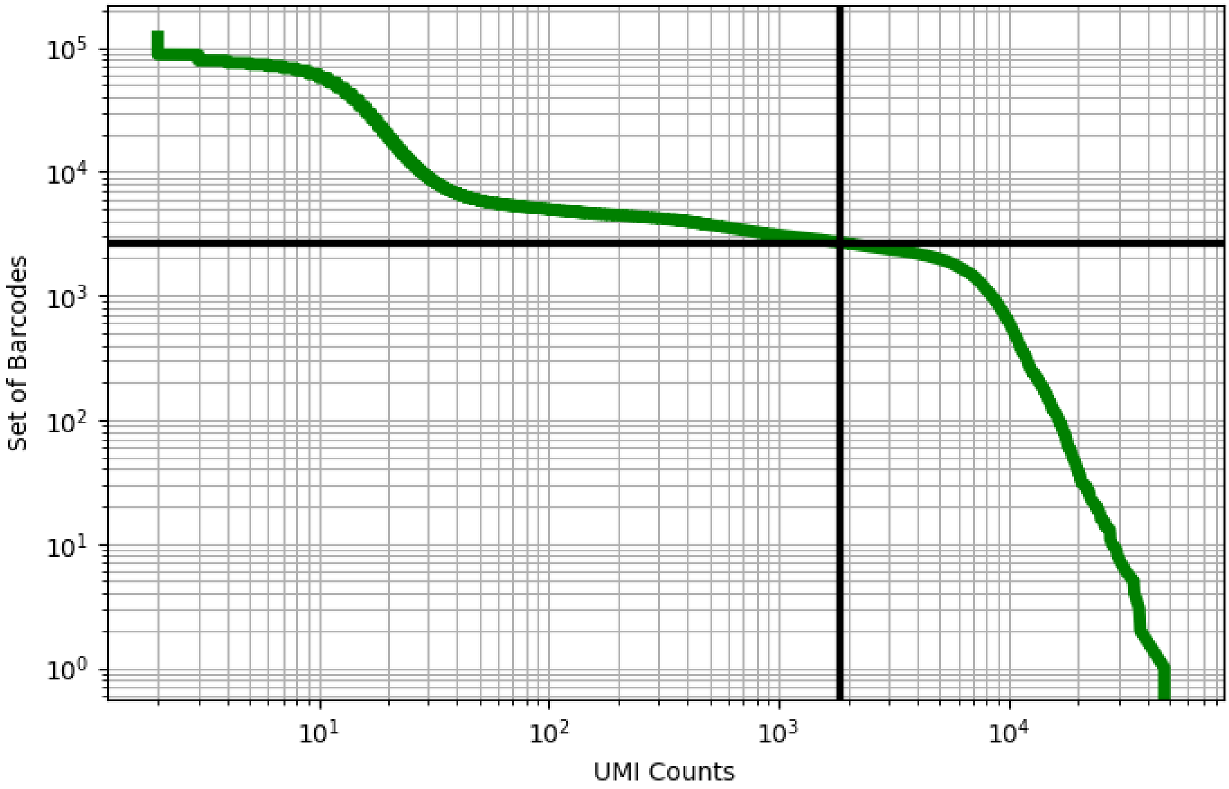


Figure 246: liver/ERS3861836

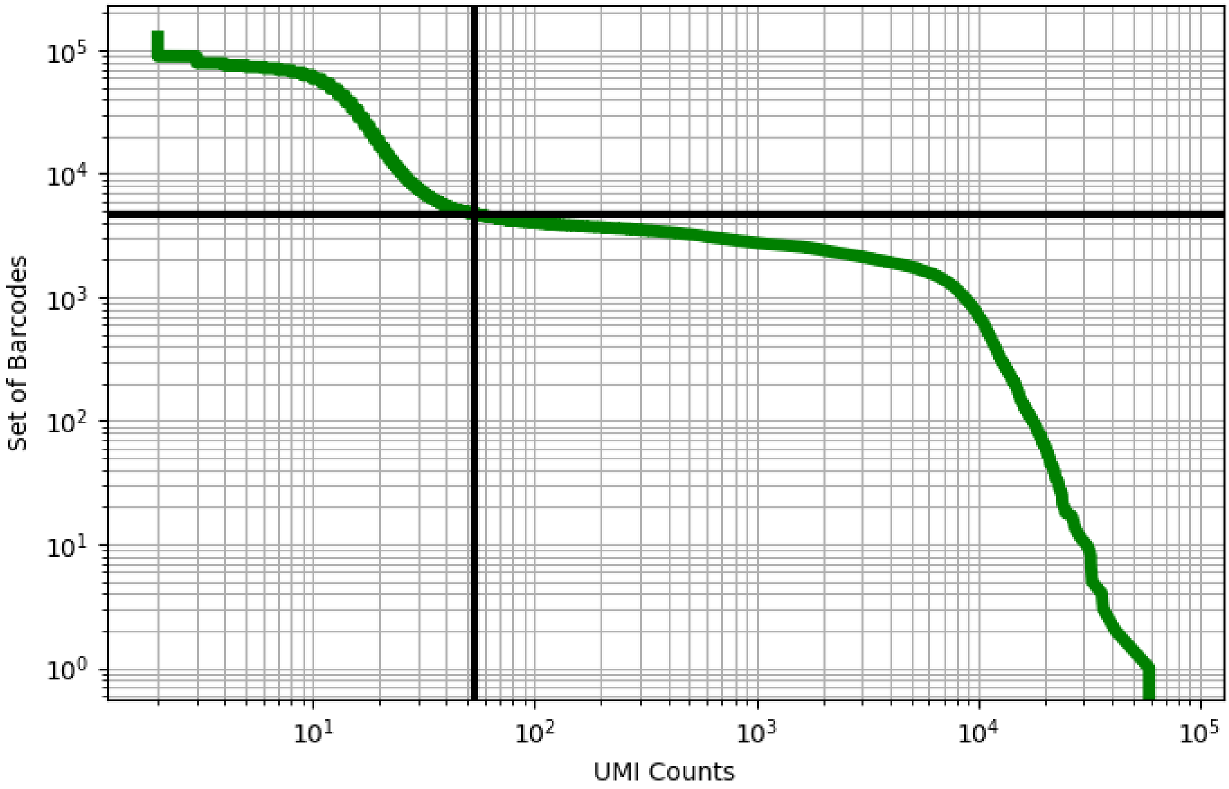


Figure 247: liver/ERS3861837

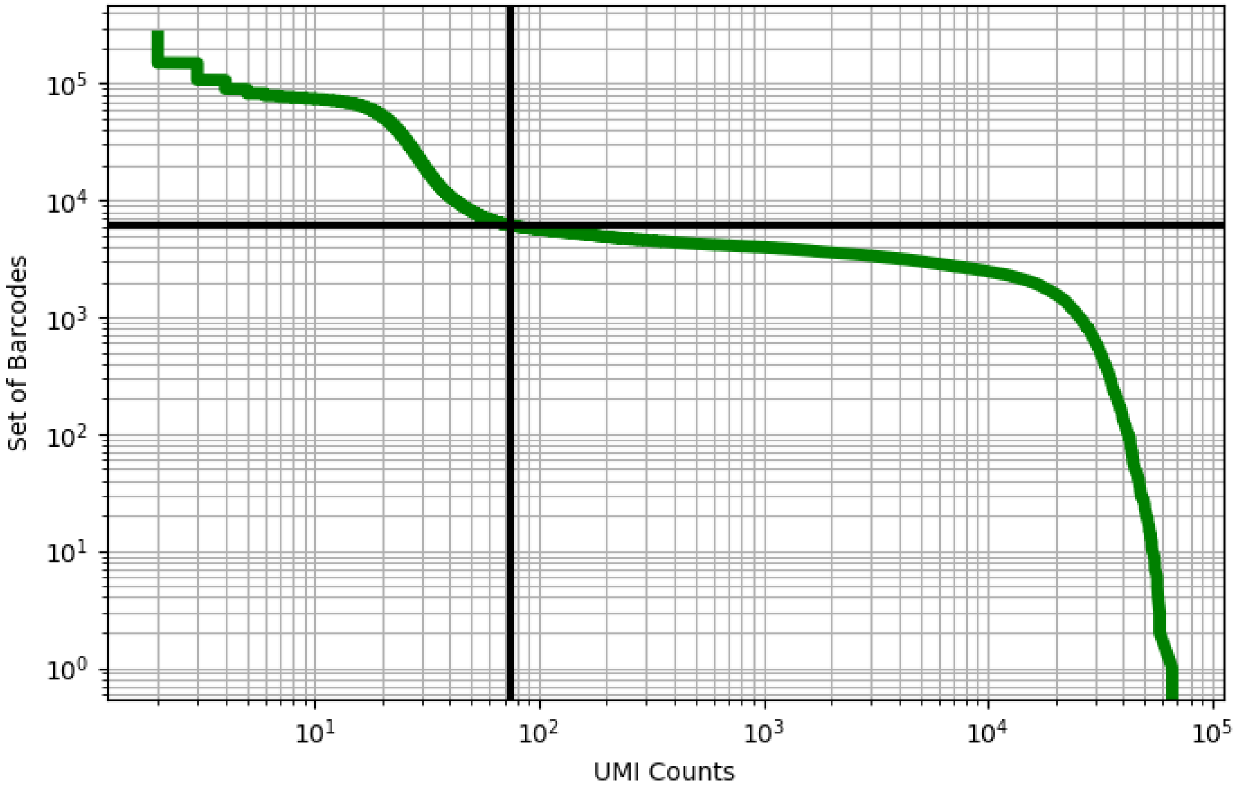


Figure 248: liver/ERS3861838

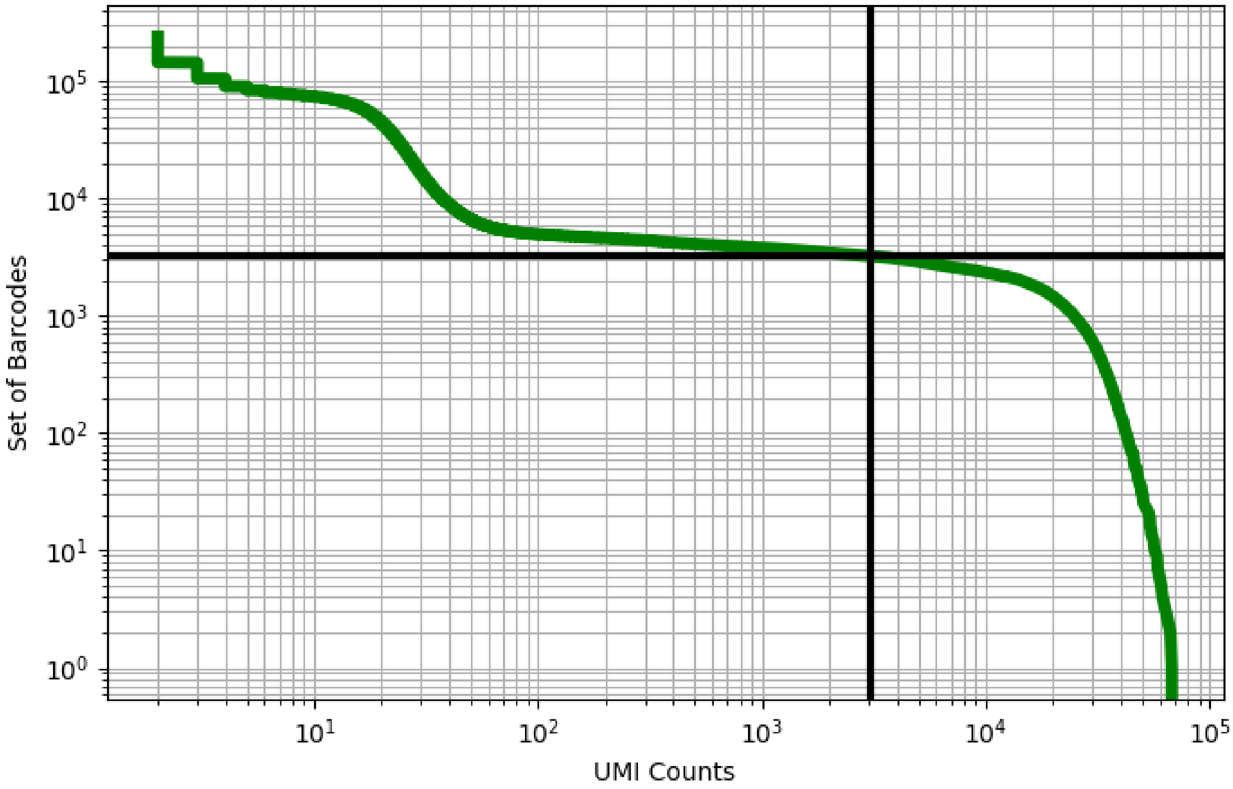


Figure 249: liver/ERS3861839

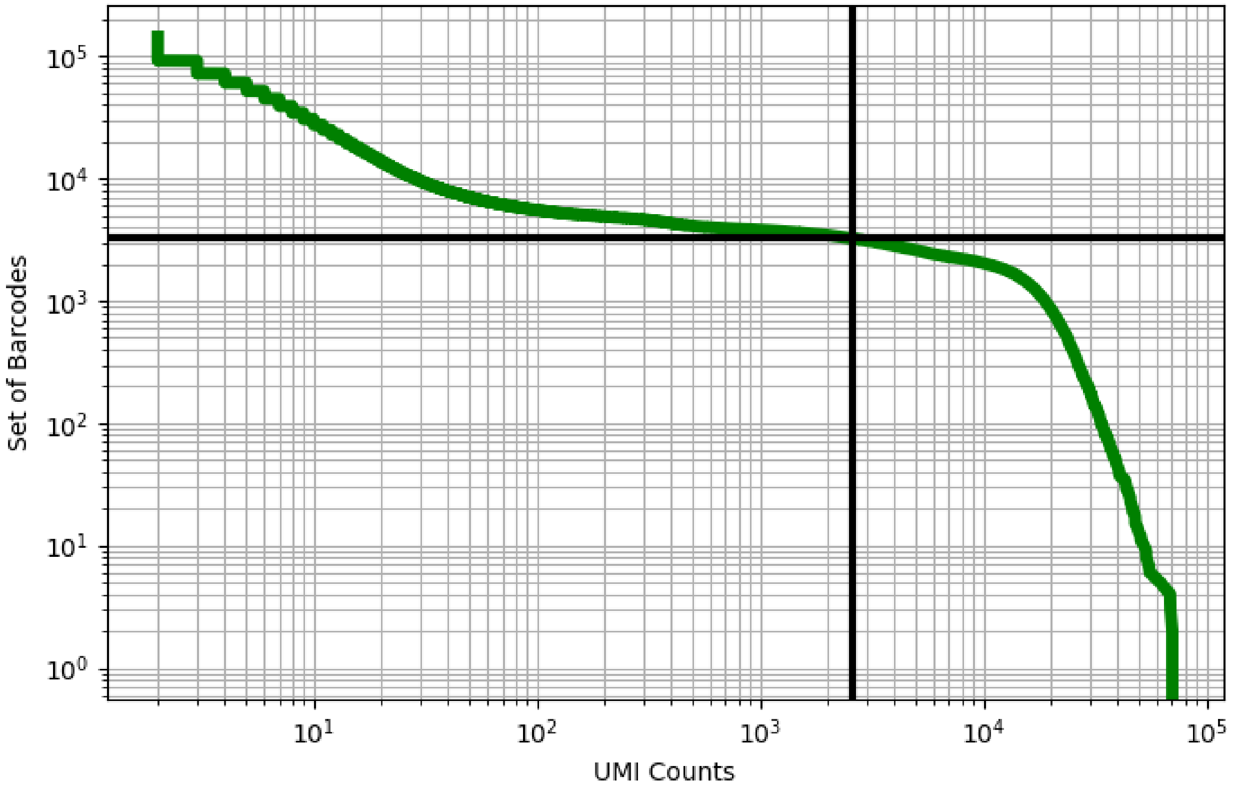


Figure 250: liver/ERS3861840

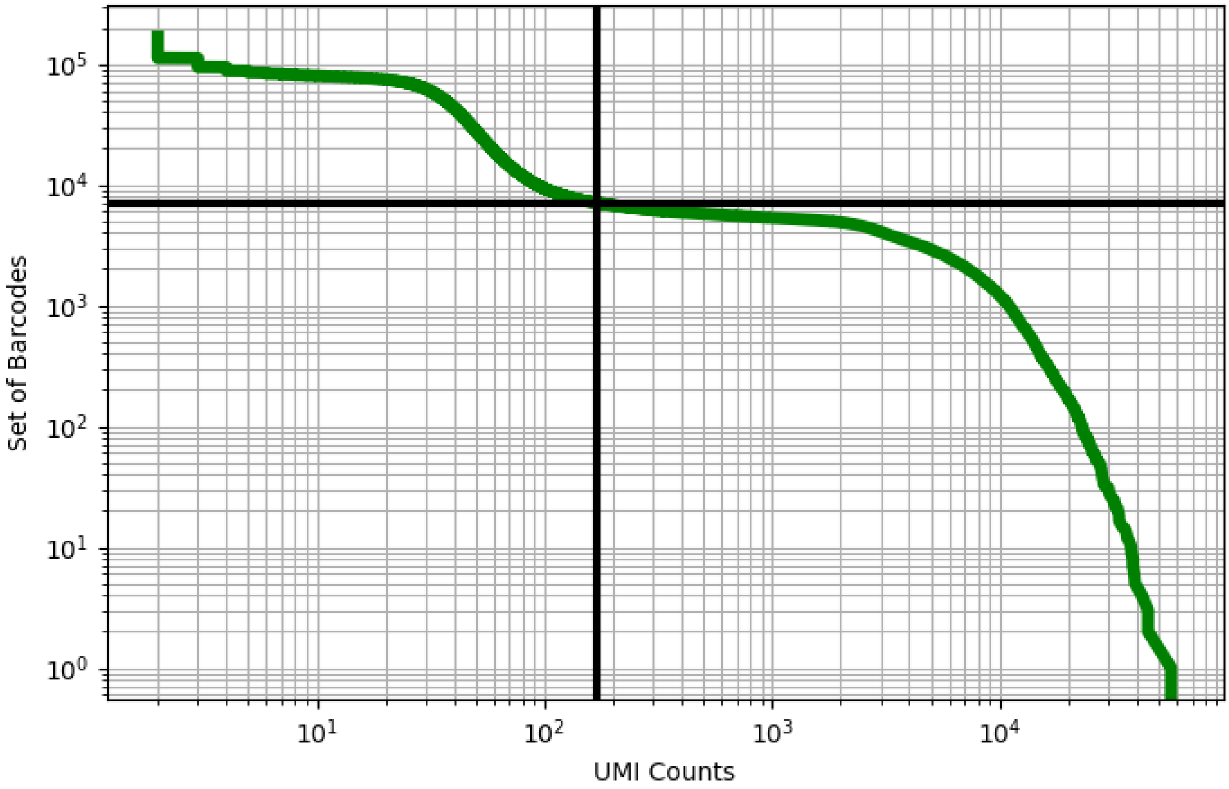


Figure 251: liver/ERS3861841

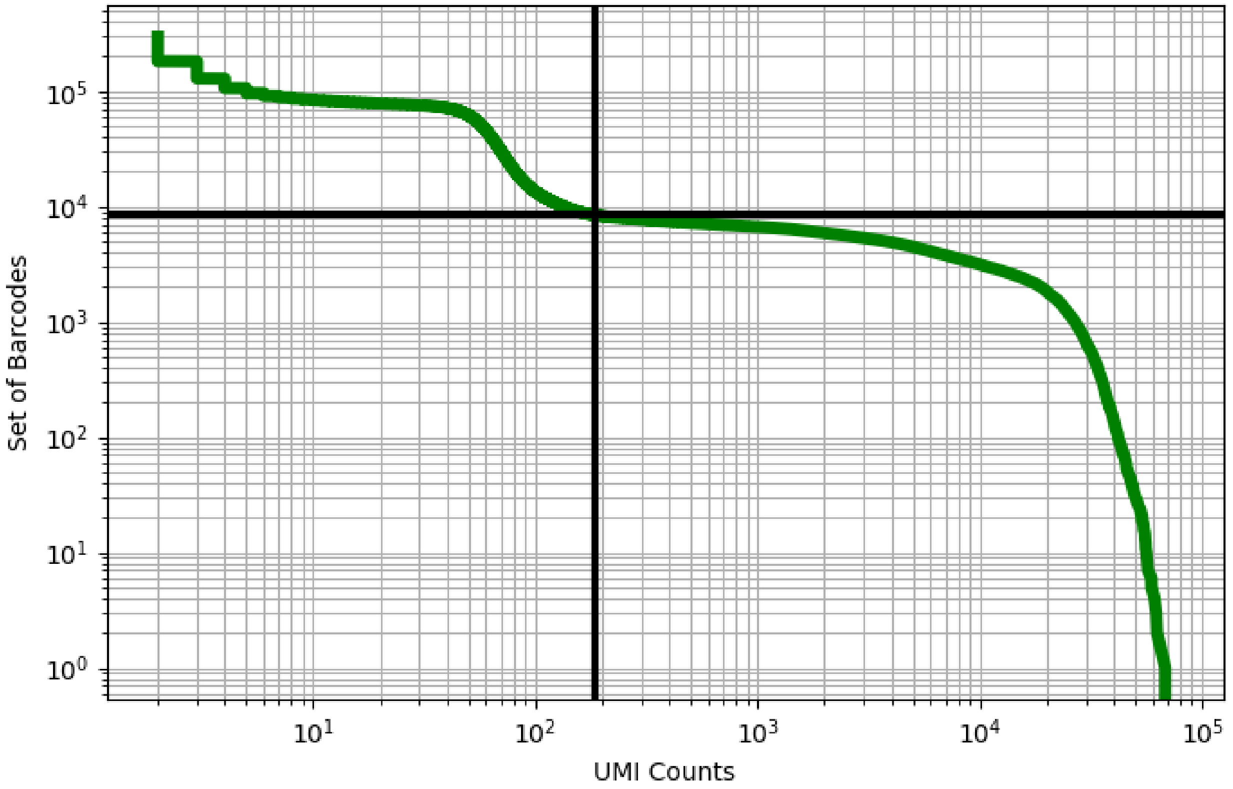


Figure 252: liver/ERS3861842

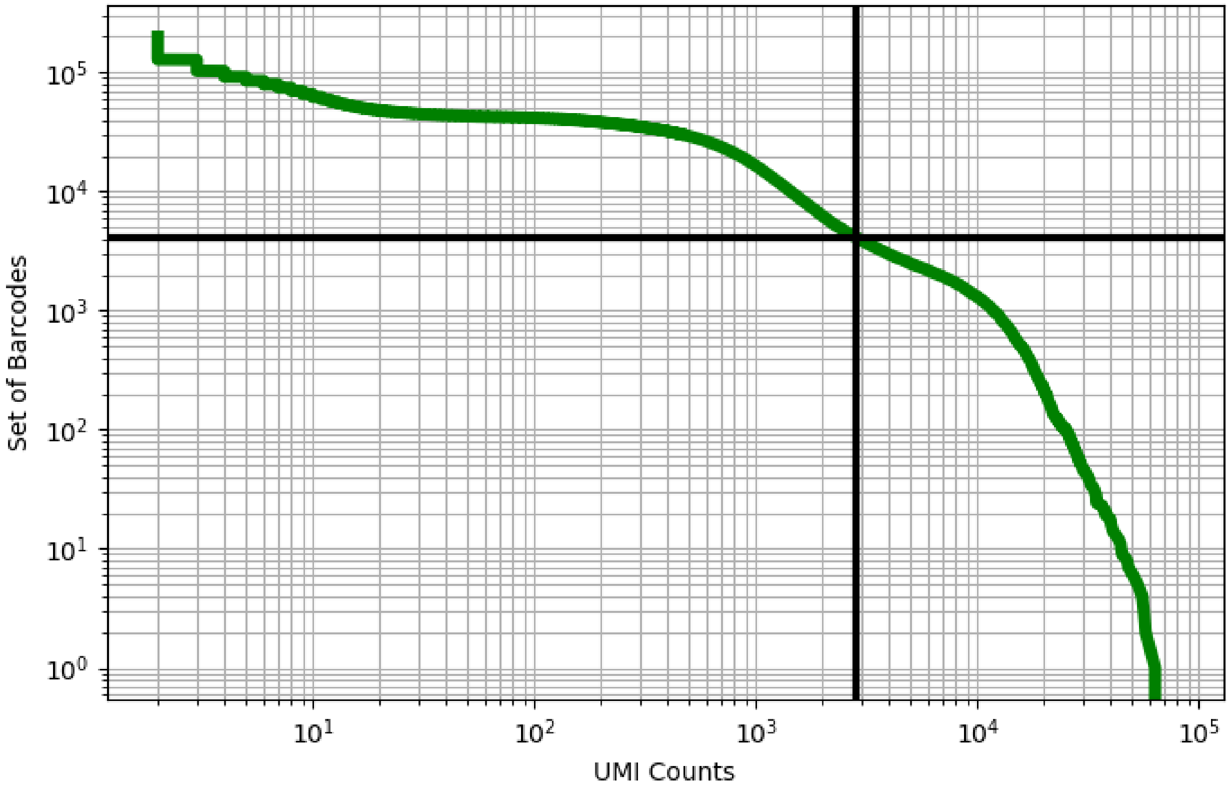


Figure 253: liver/ERS3861843

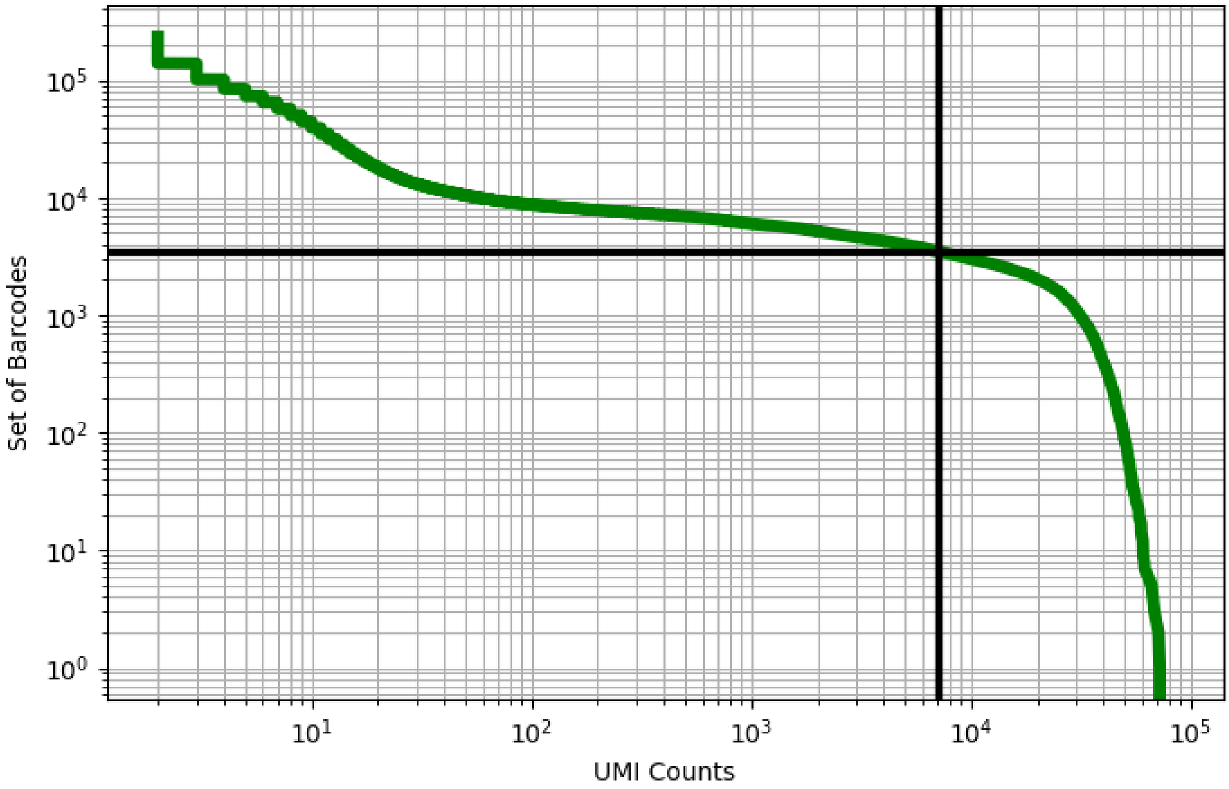


Figure 254: liver/ERS3861844

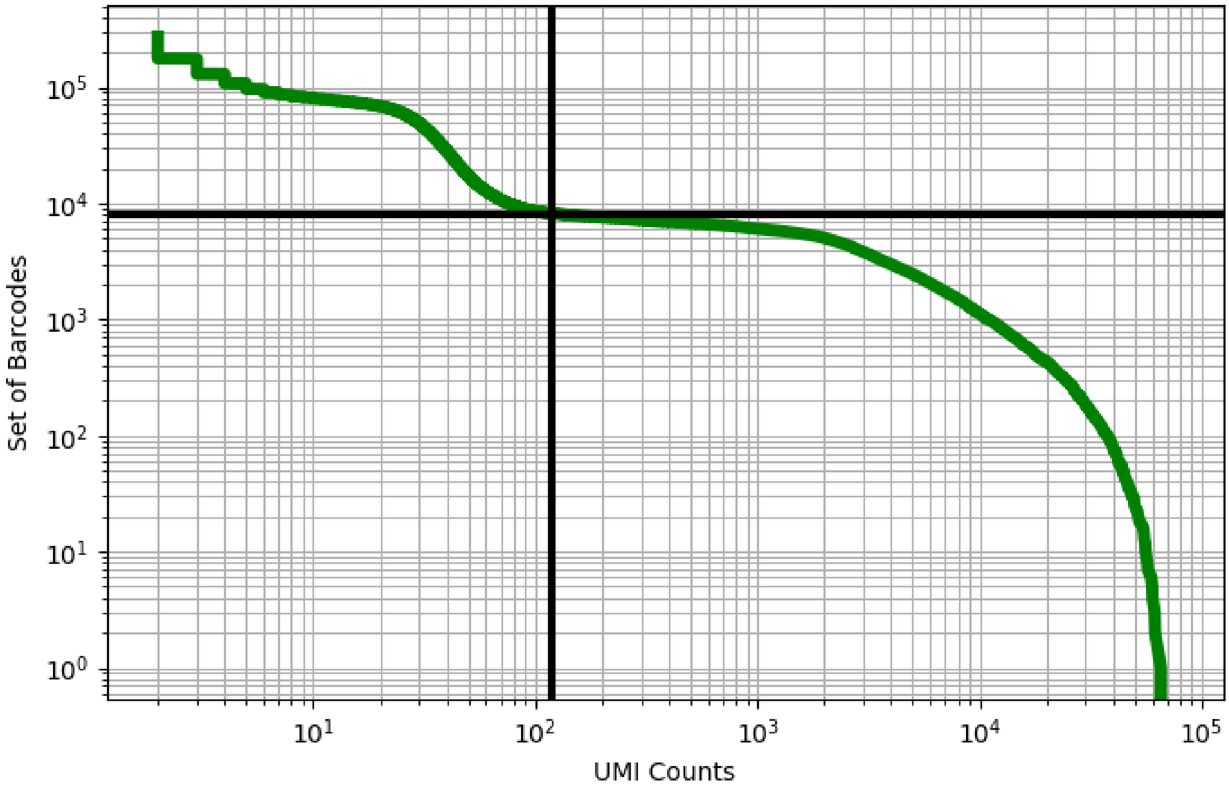


Figure 255: liver/ERS3861845

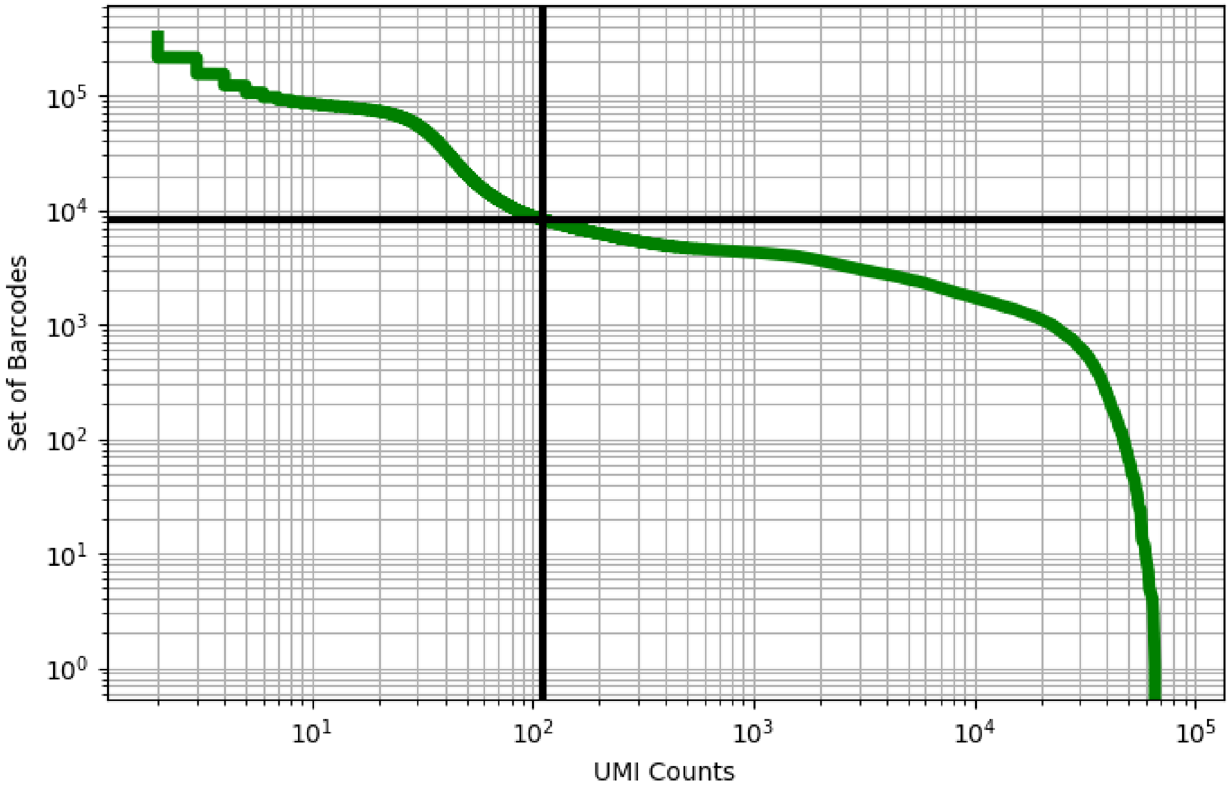


Figure 256: liver/ERS3861846

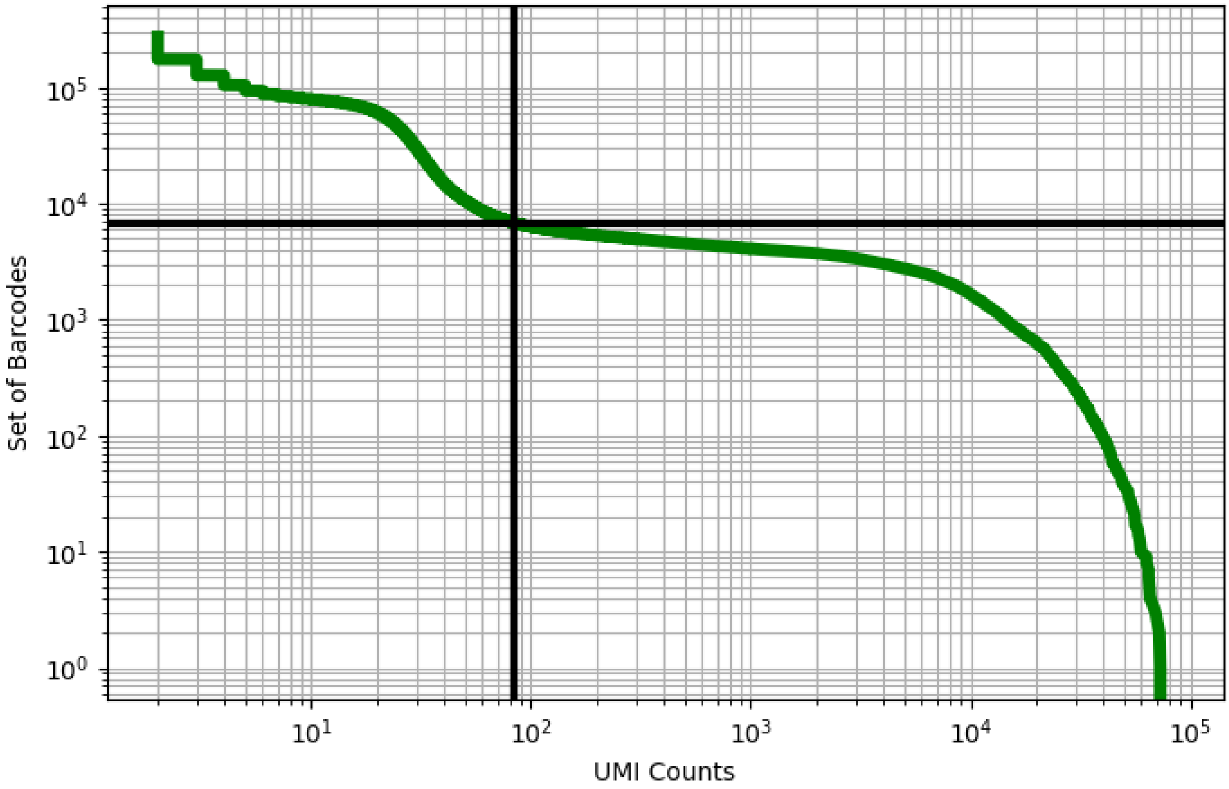


Figure 257: liver/ERS3861847

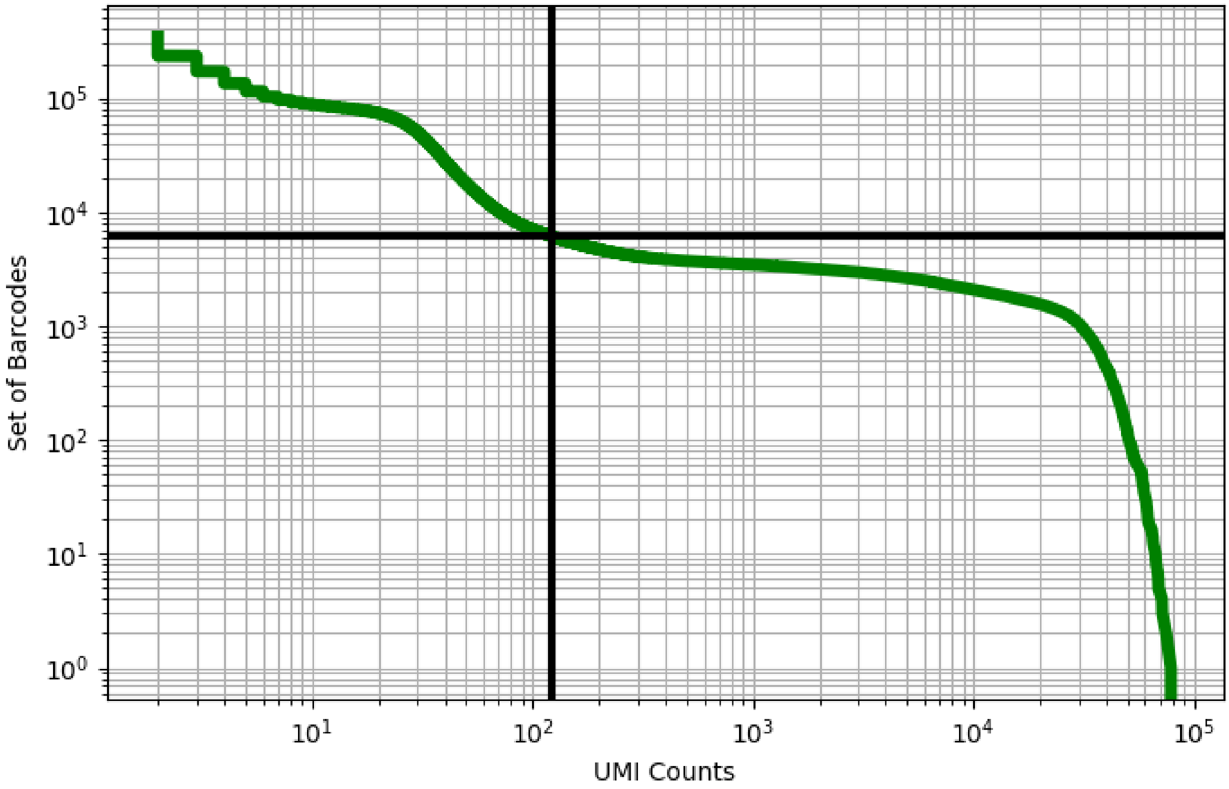


Figure 258: liver/ERS3861848

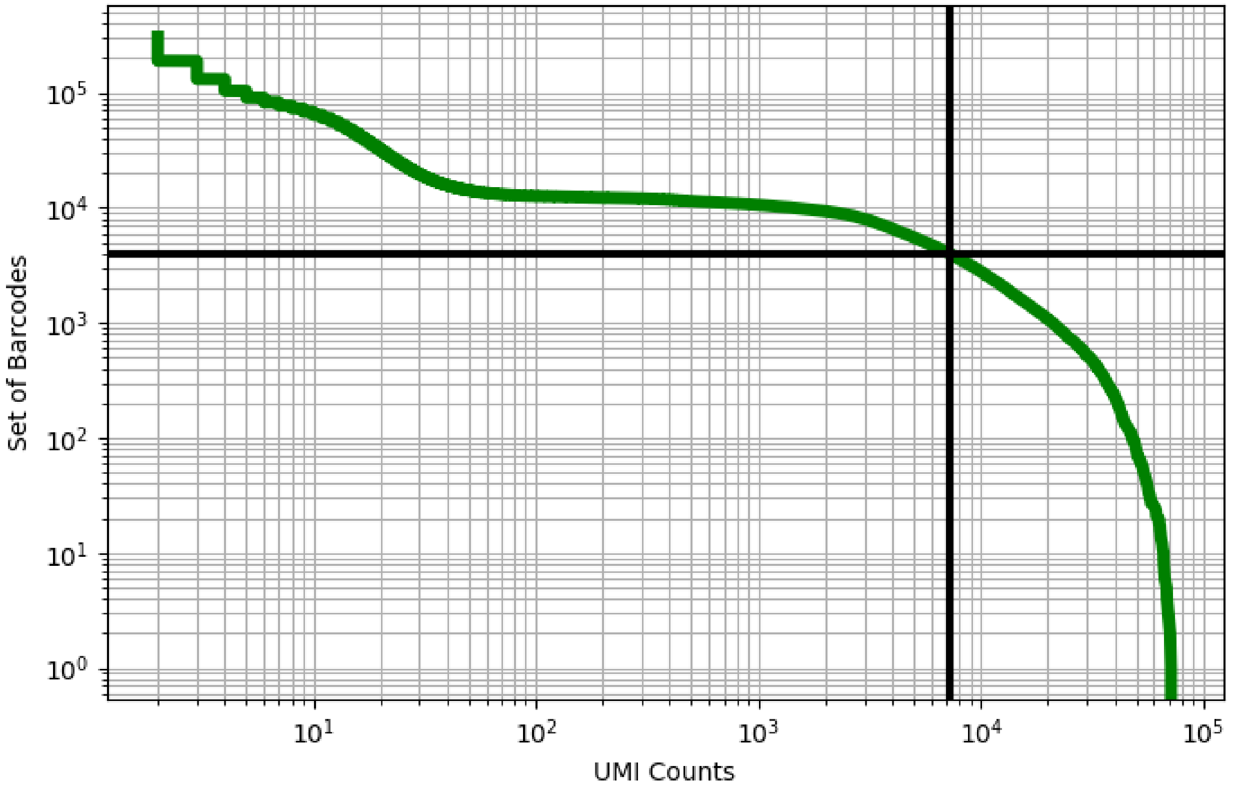


Figure 259: liver/ERS3861849

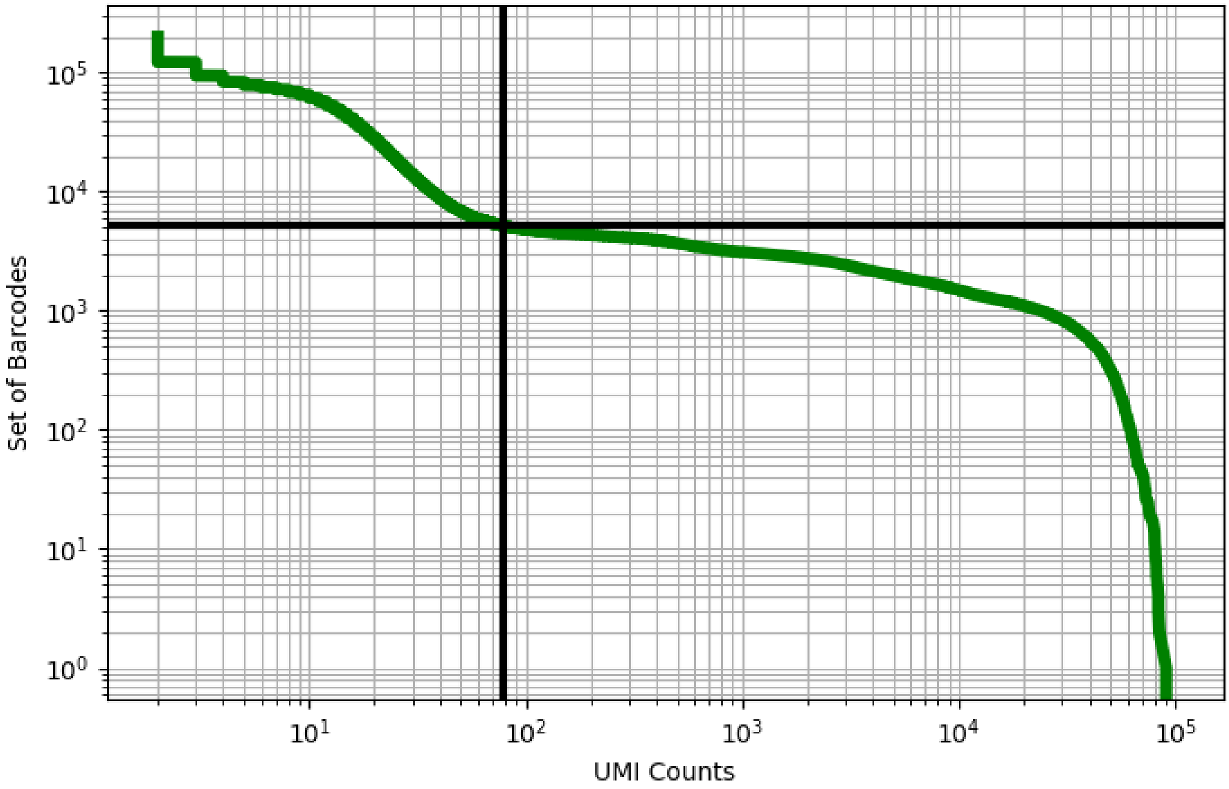


Figure 260: liver/ERS3861850

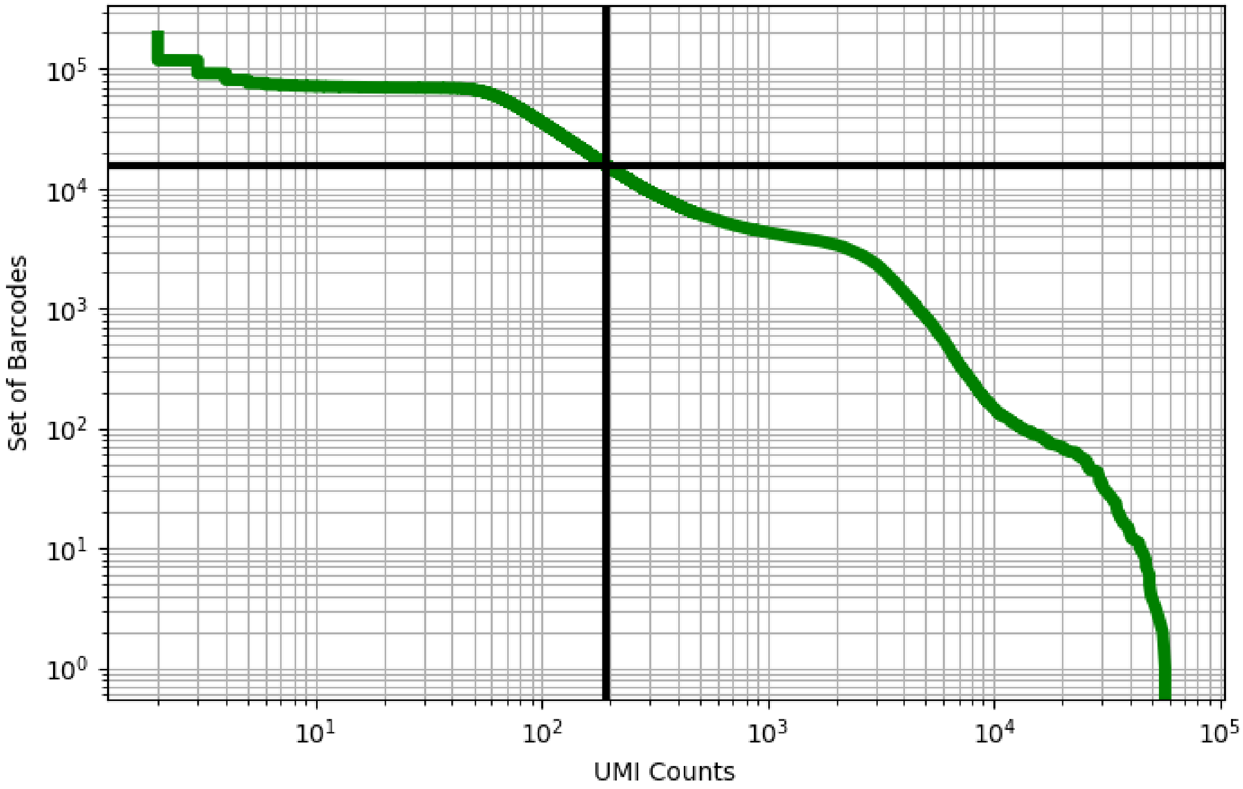


Figure 261: liver/GSM3178782

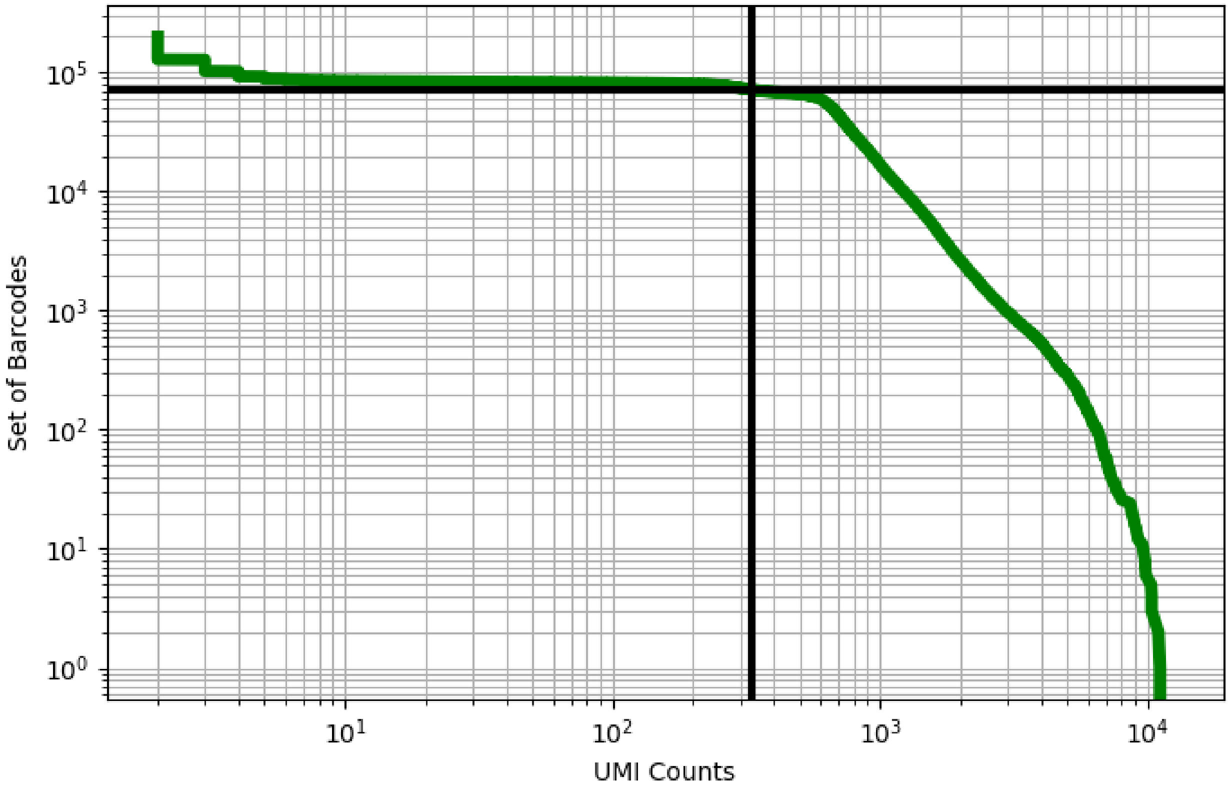


Figure 262: liver/GSM3178783

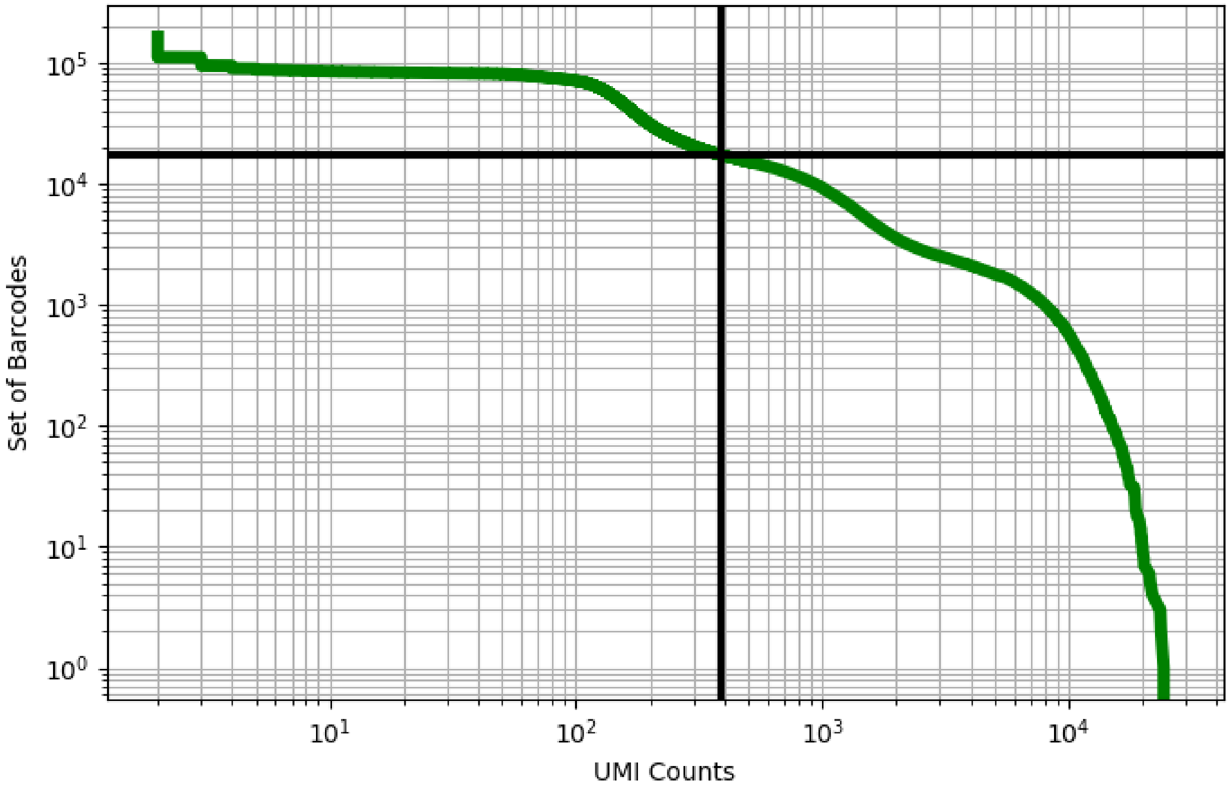


Figure 263: liver/GSM3178784

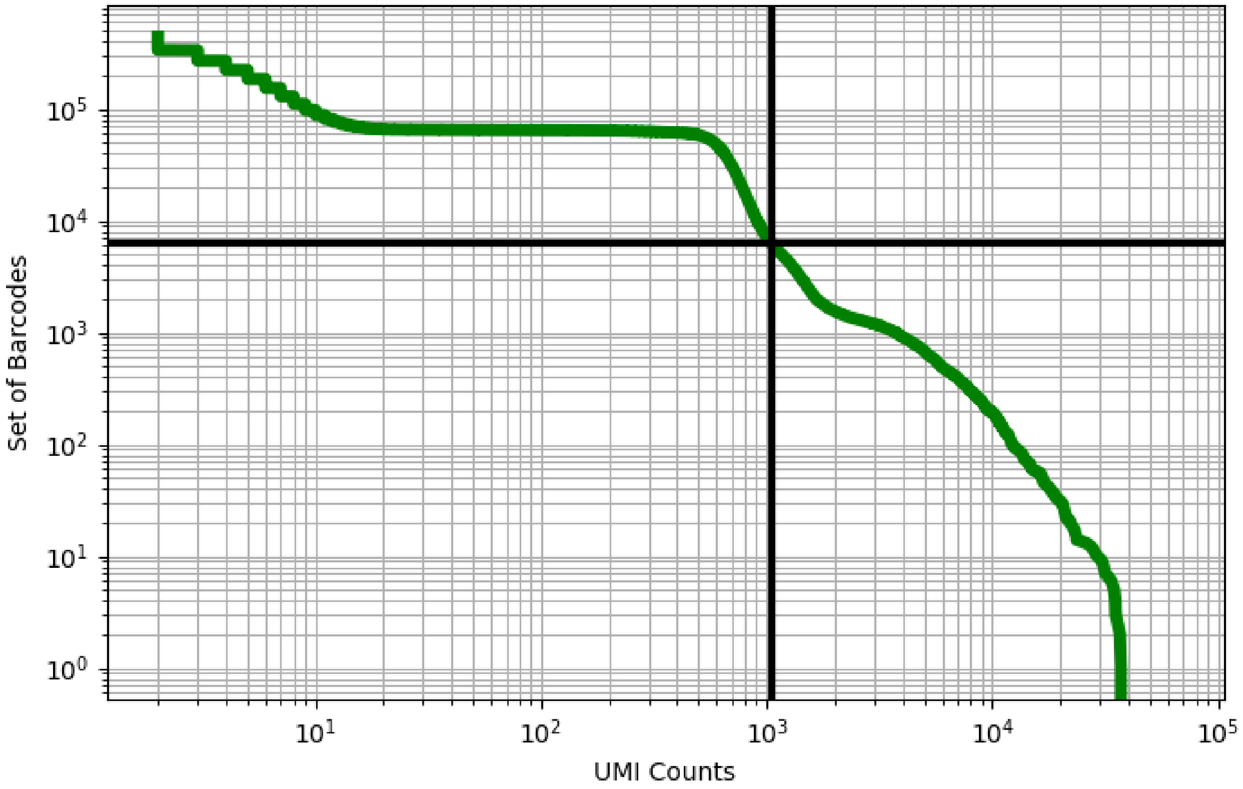


Figure 264: liver/GSM3178785

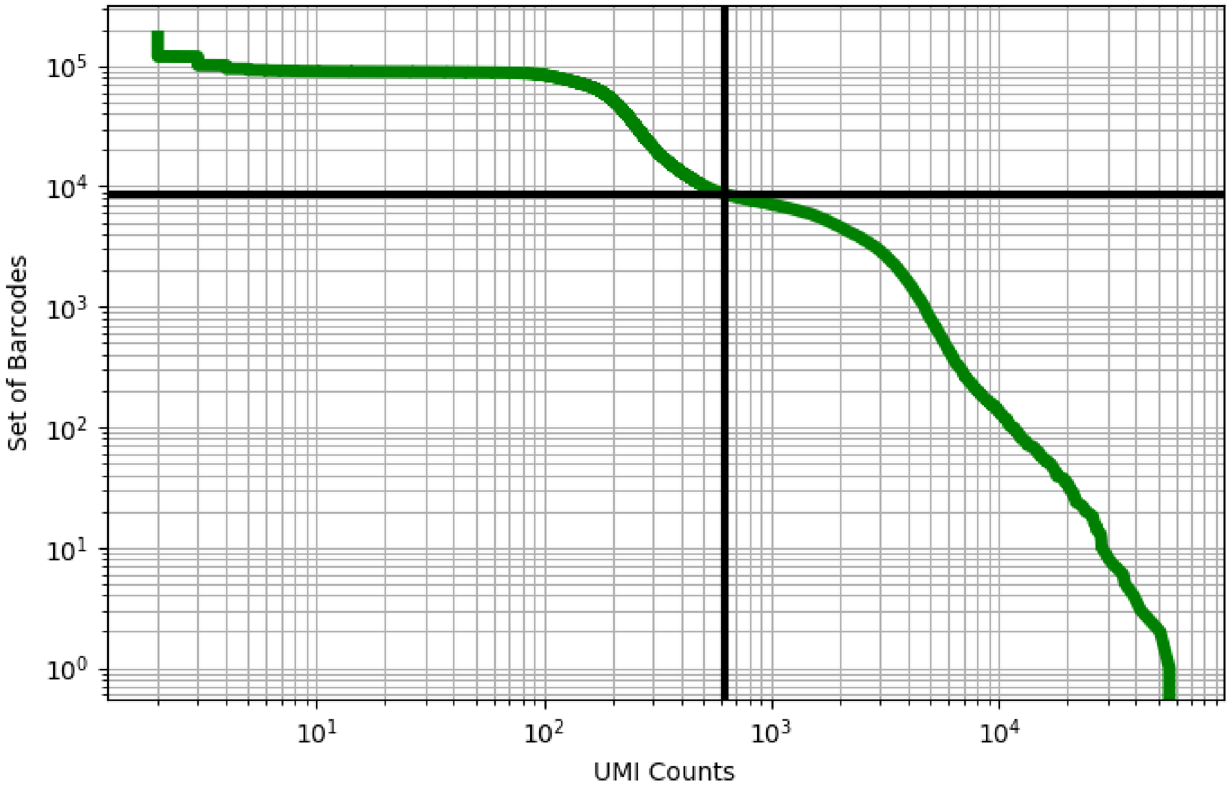


Figure 265: liver/GSM3178786

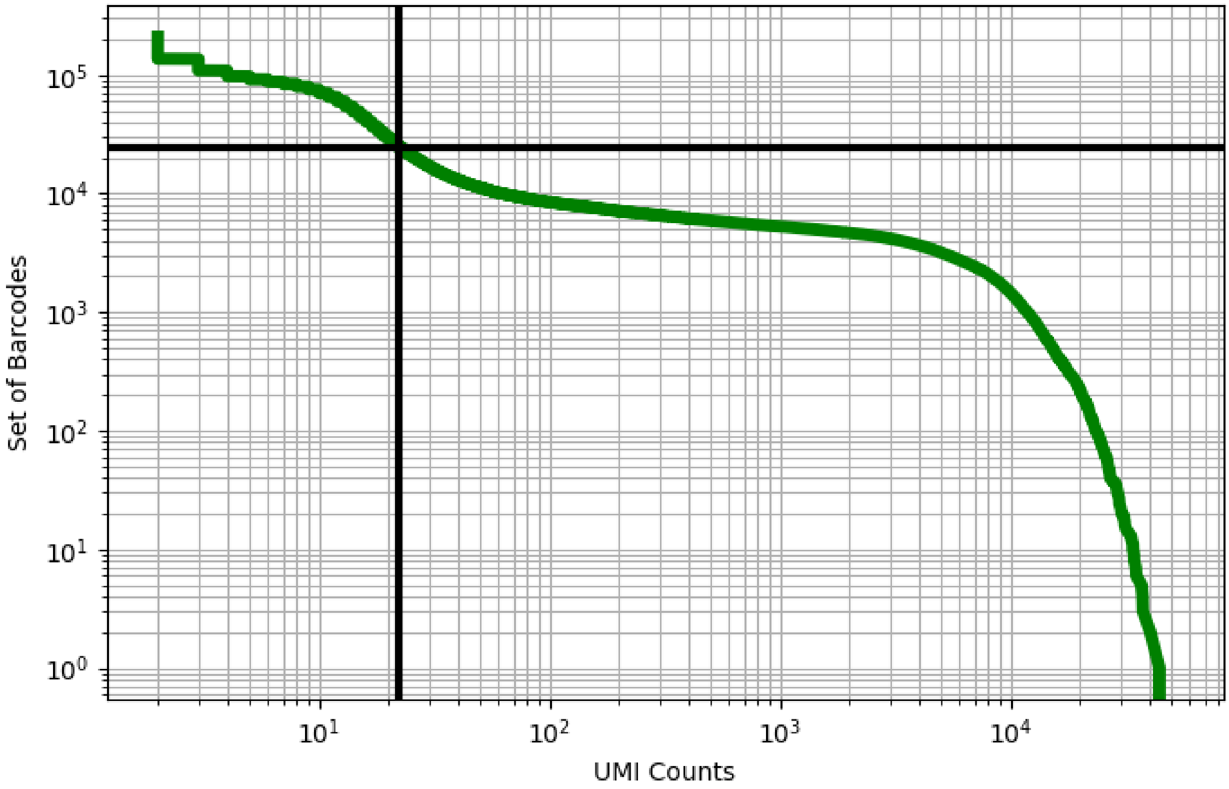


Figure 266: liver/GSM3731527

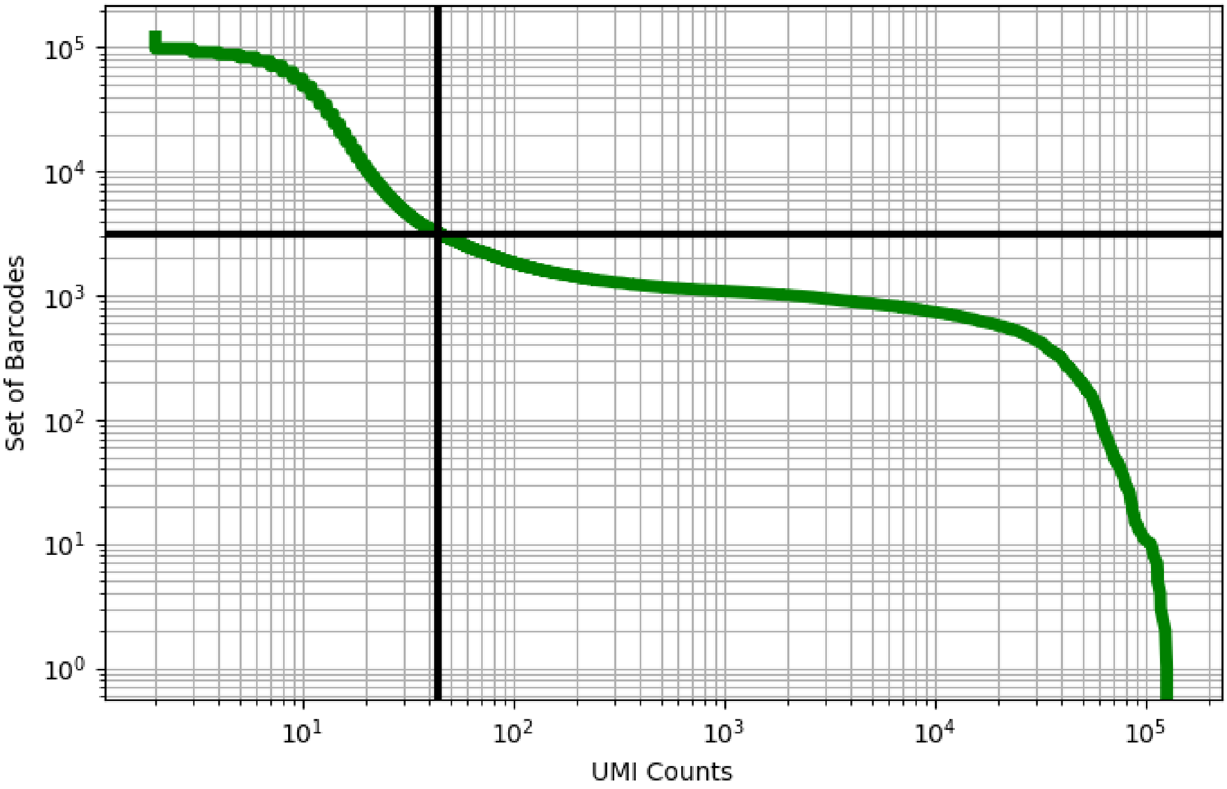


Figure 267: lung/GSM2786157

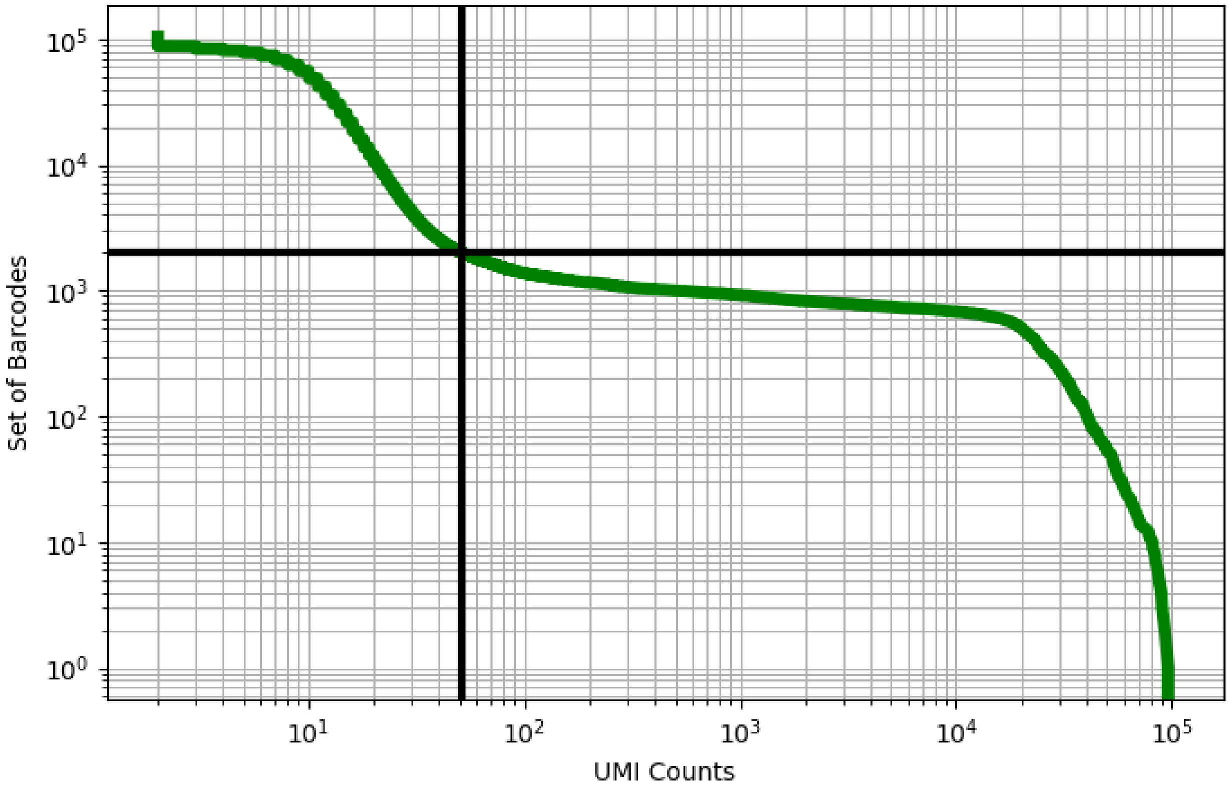


Figure 268: lung/GSM2786158

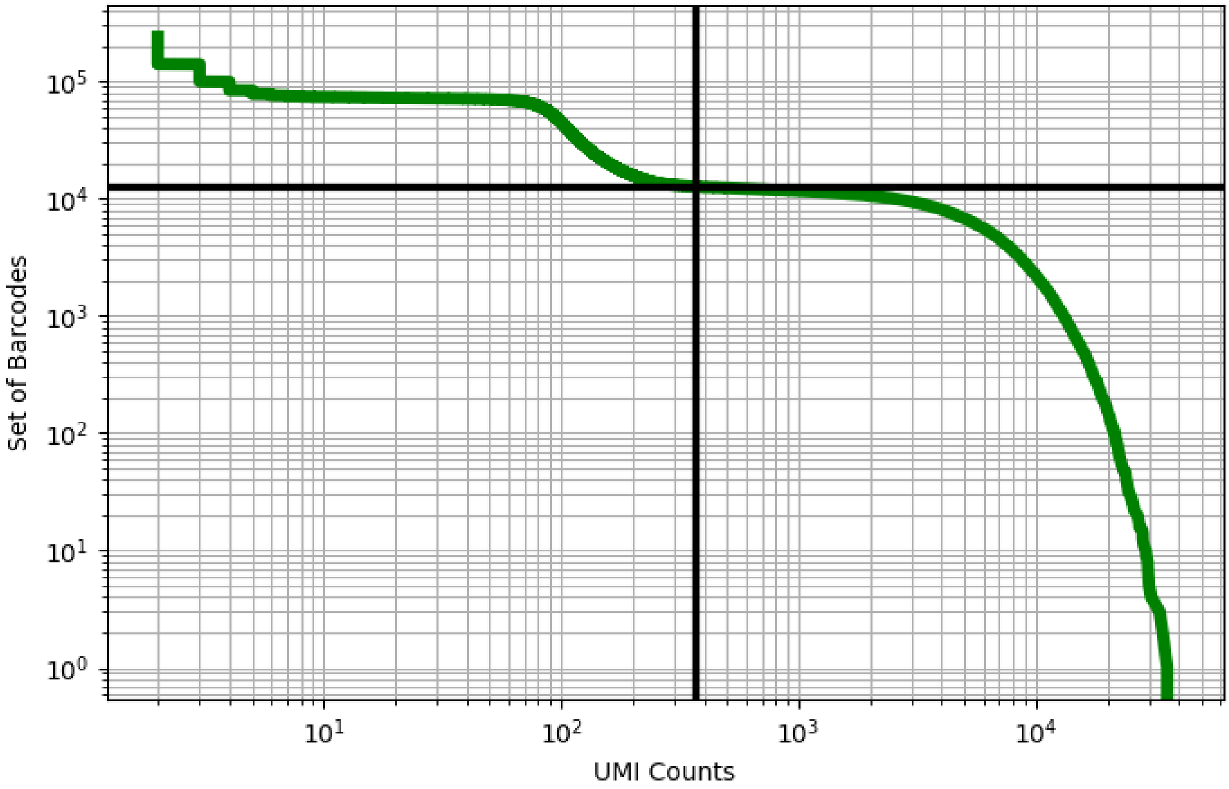


Figure 269: lung/GSM2894834

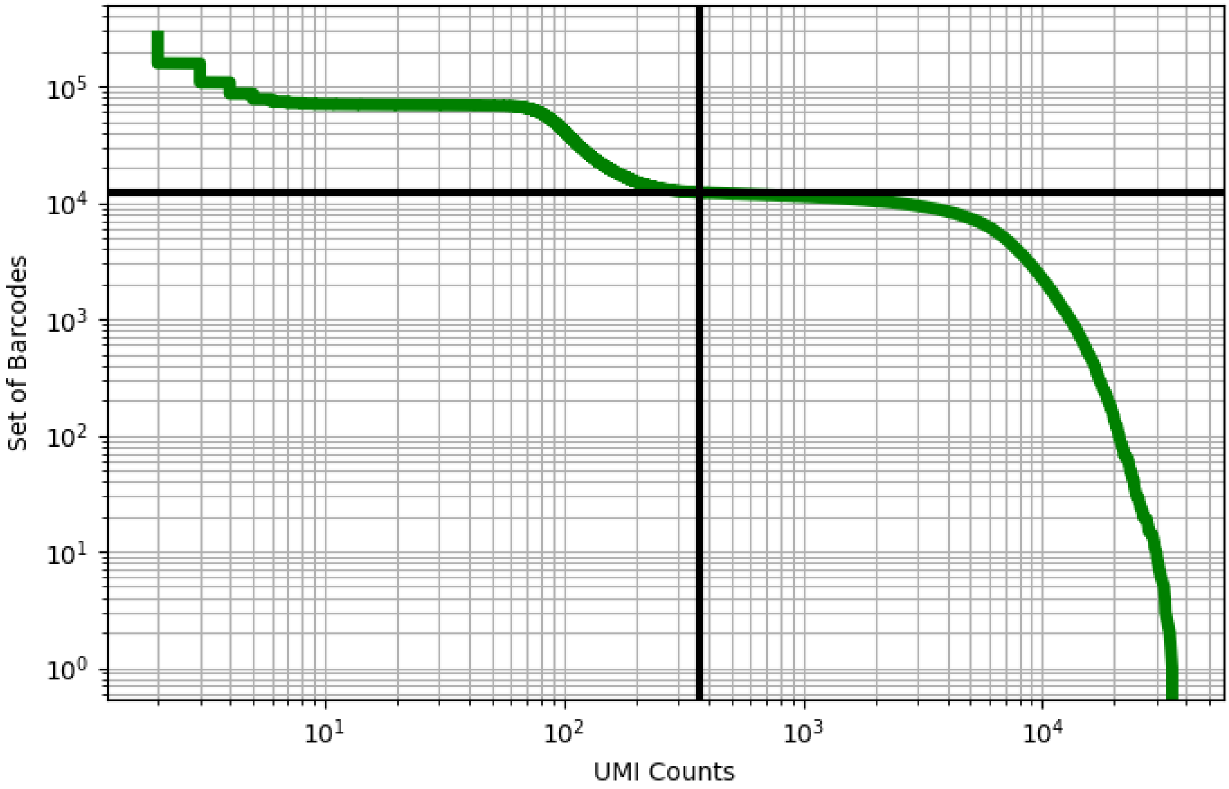


Figure 270: lung/GSM2894835

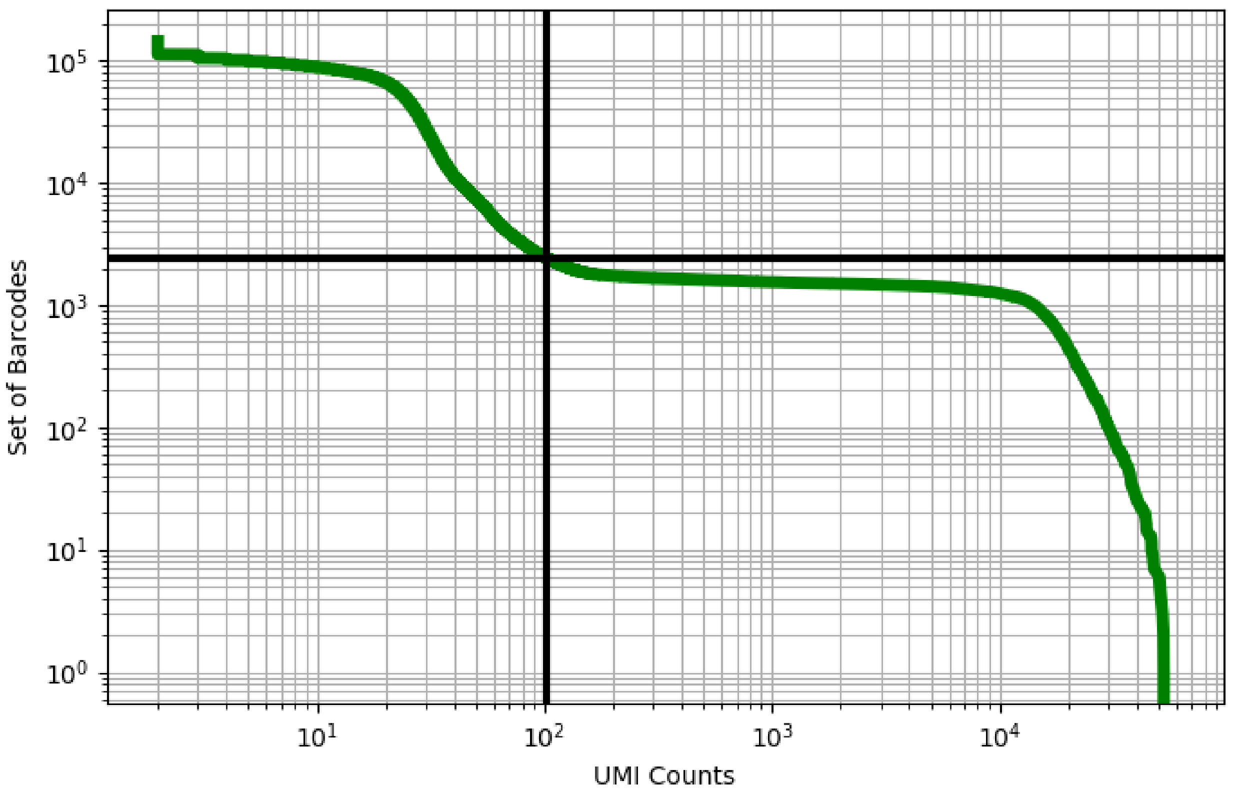


Figure 271: lung/GSM3439913

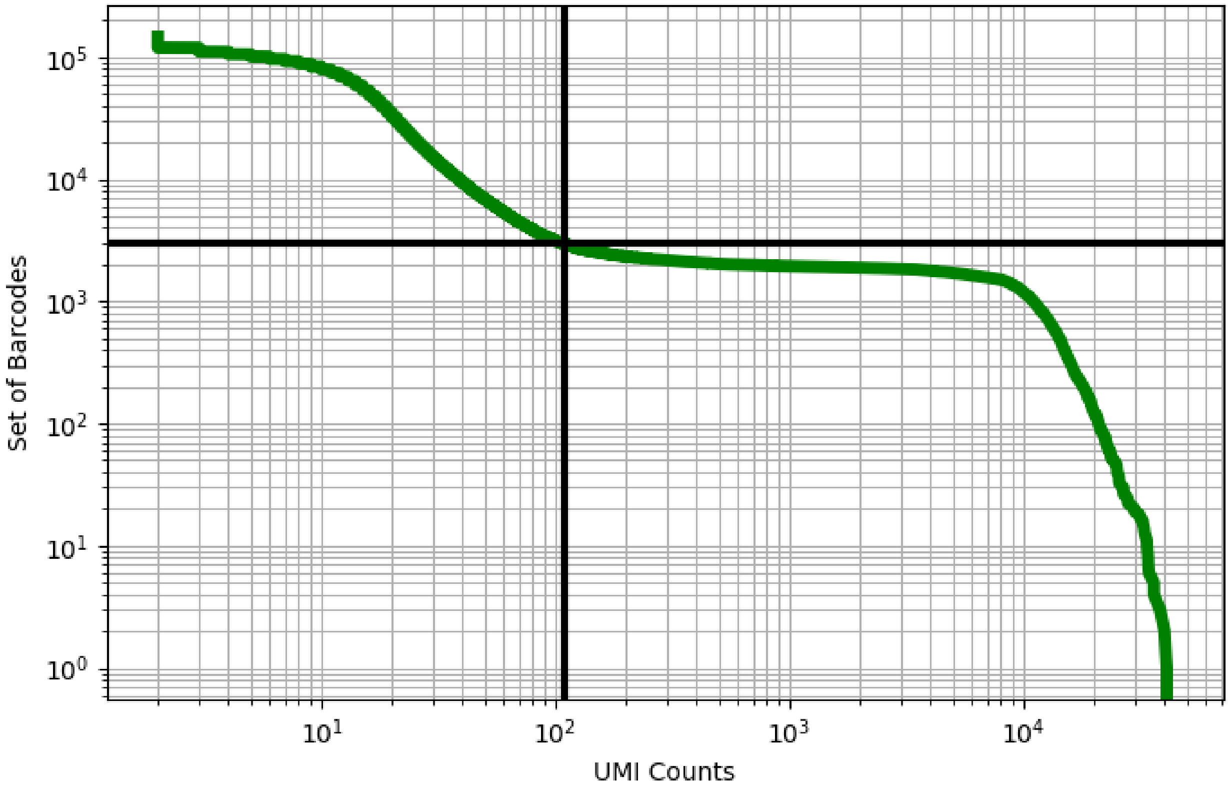


Figure 272: lung/GSM3439914

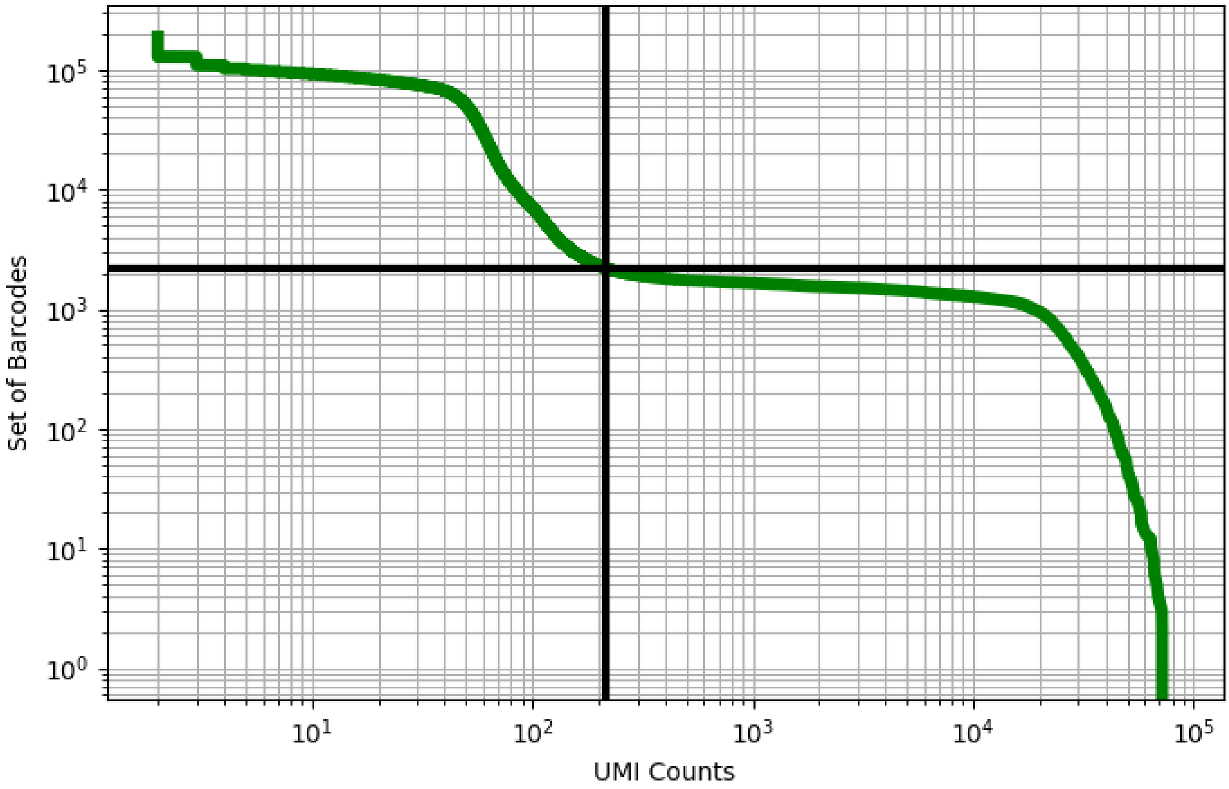


Figure 273: lung/GSM3439915

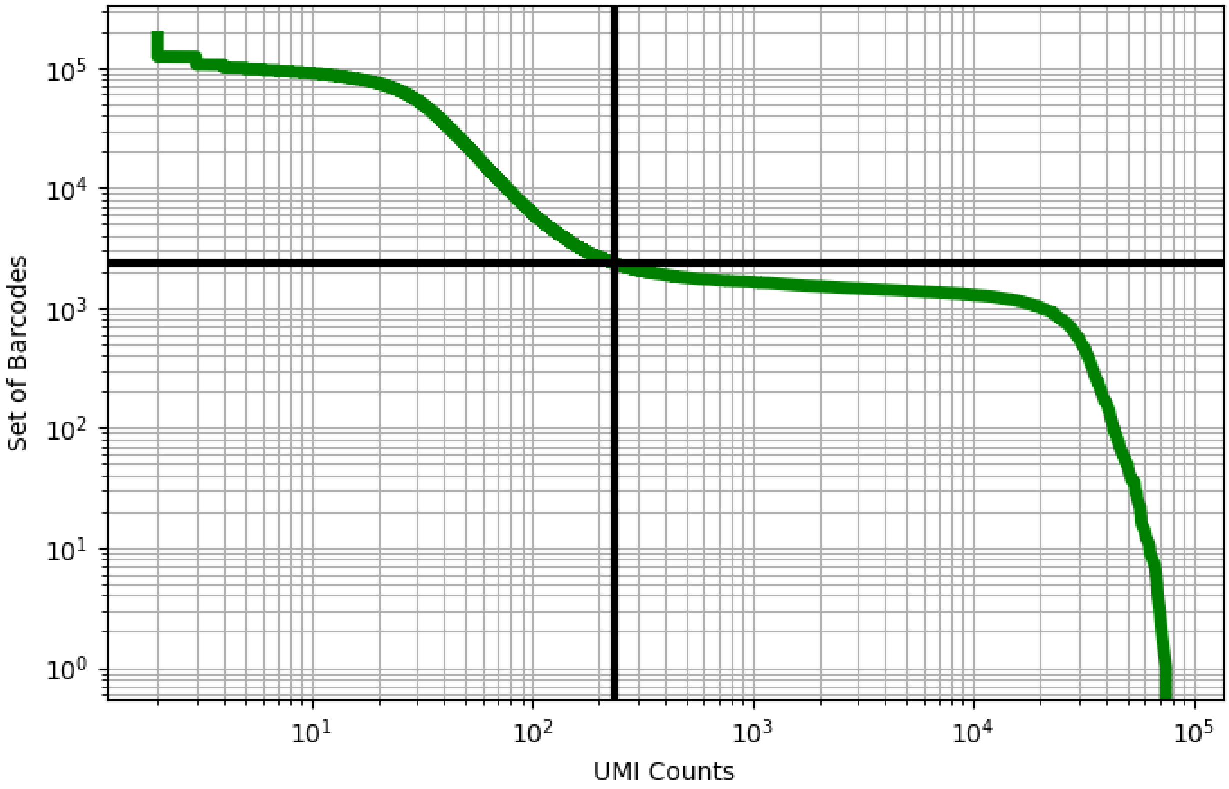


Figure 274: lung/GSM3439916

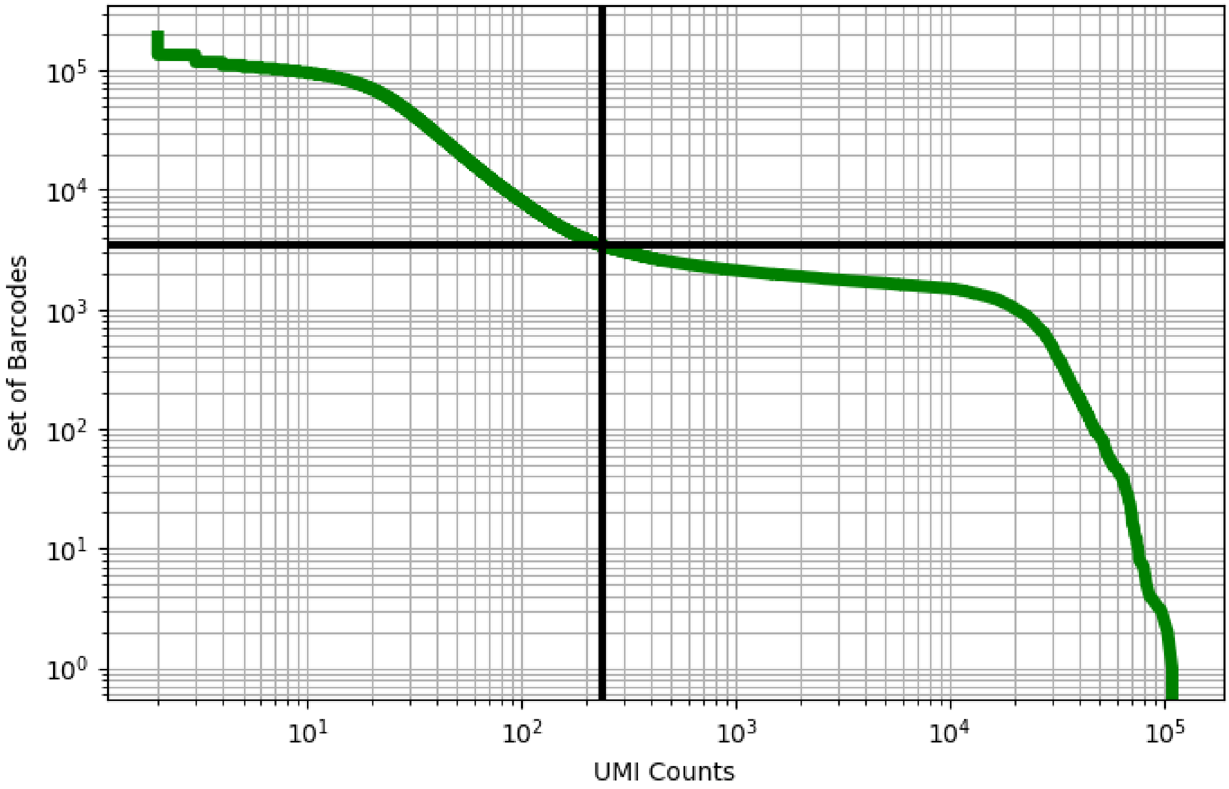


Figure 275: lung/GSM3439917

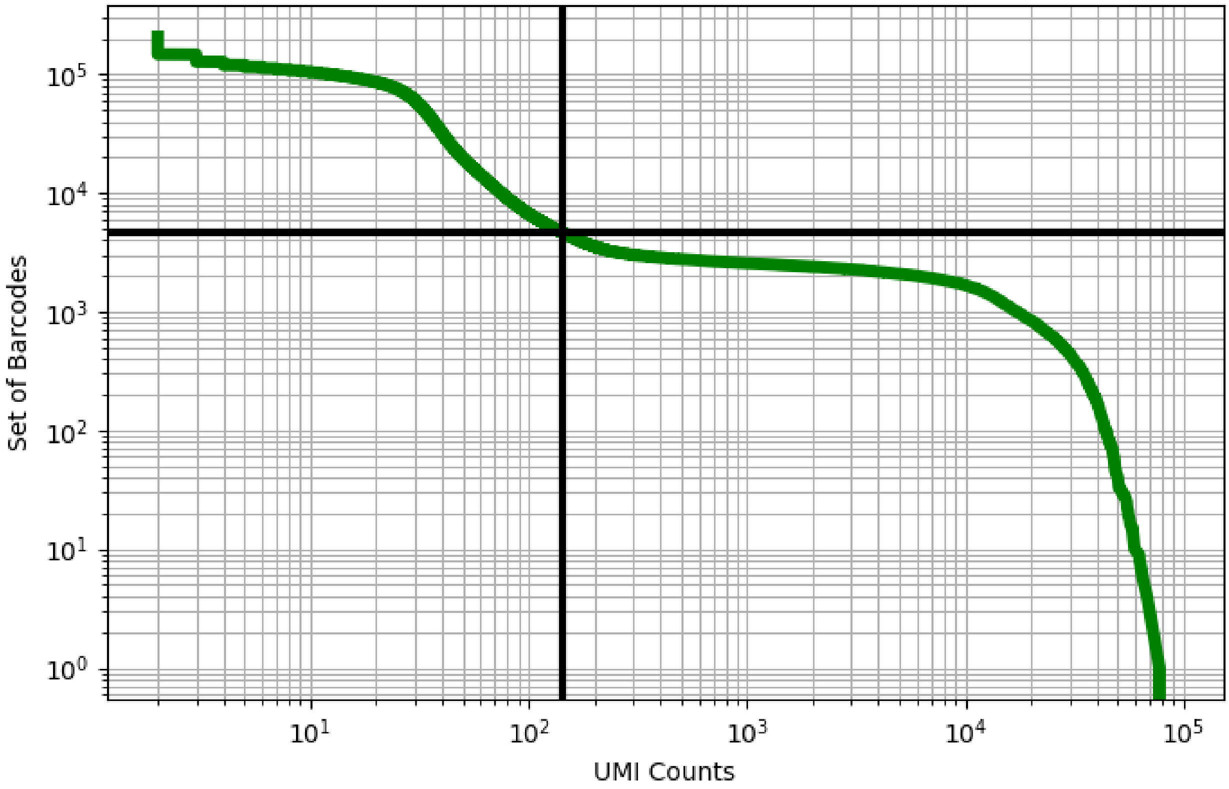


Figure 276: lung/GSM3439918

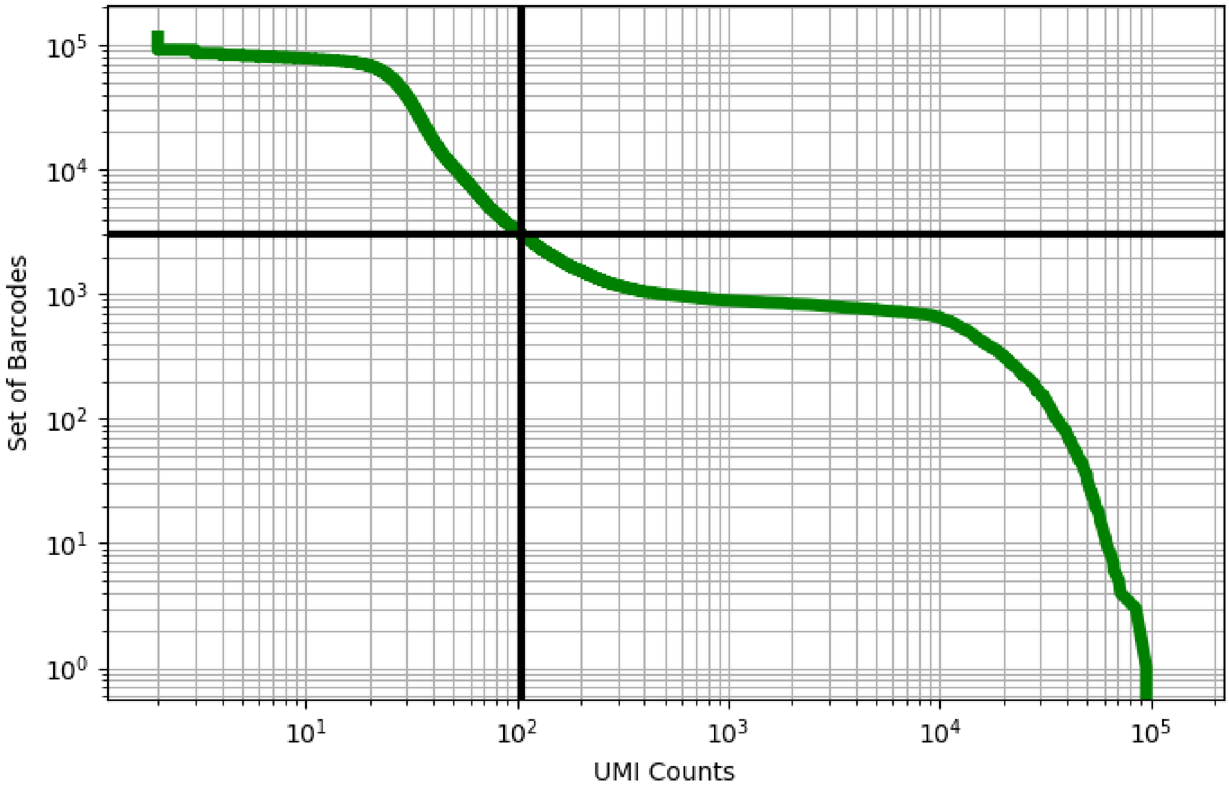


Figure 277: lung/GSM3439919

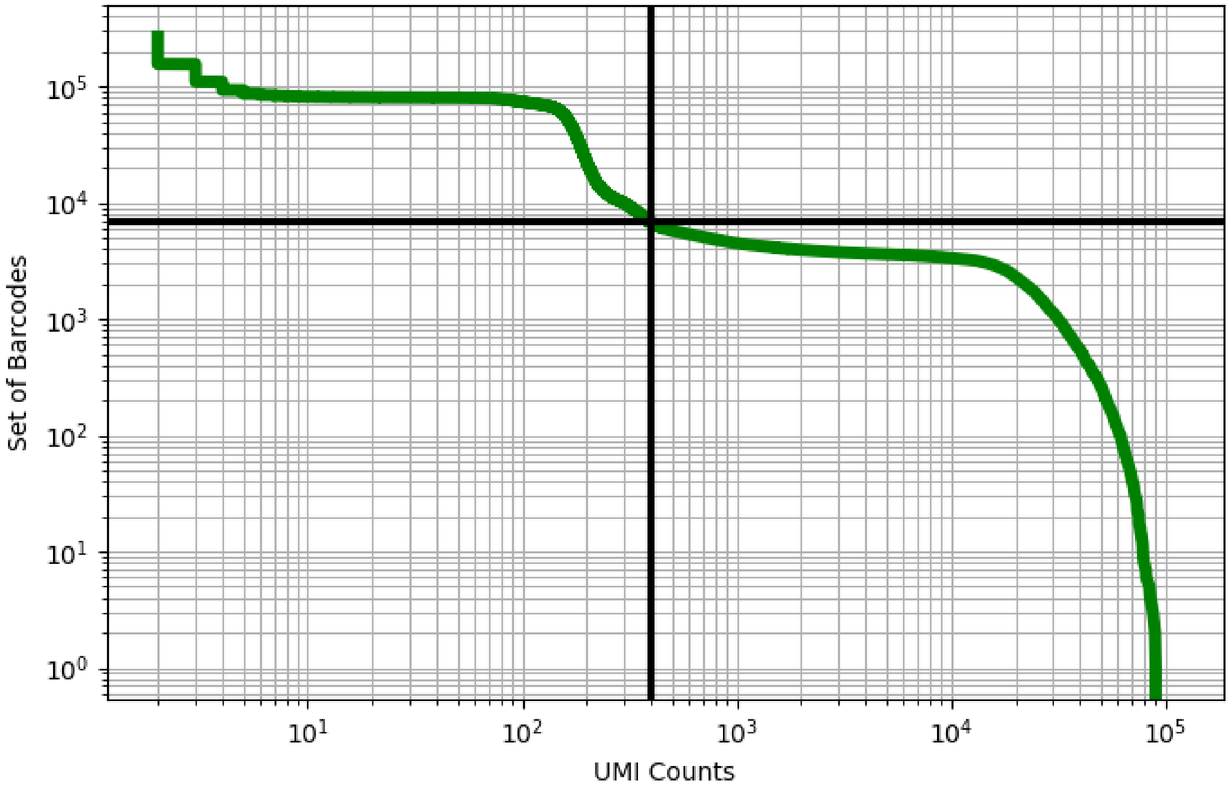


Figure 278: lung/GSM3439920

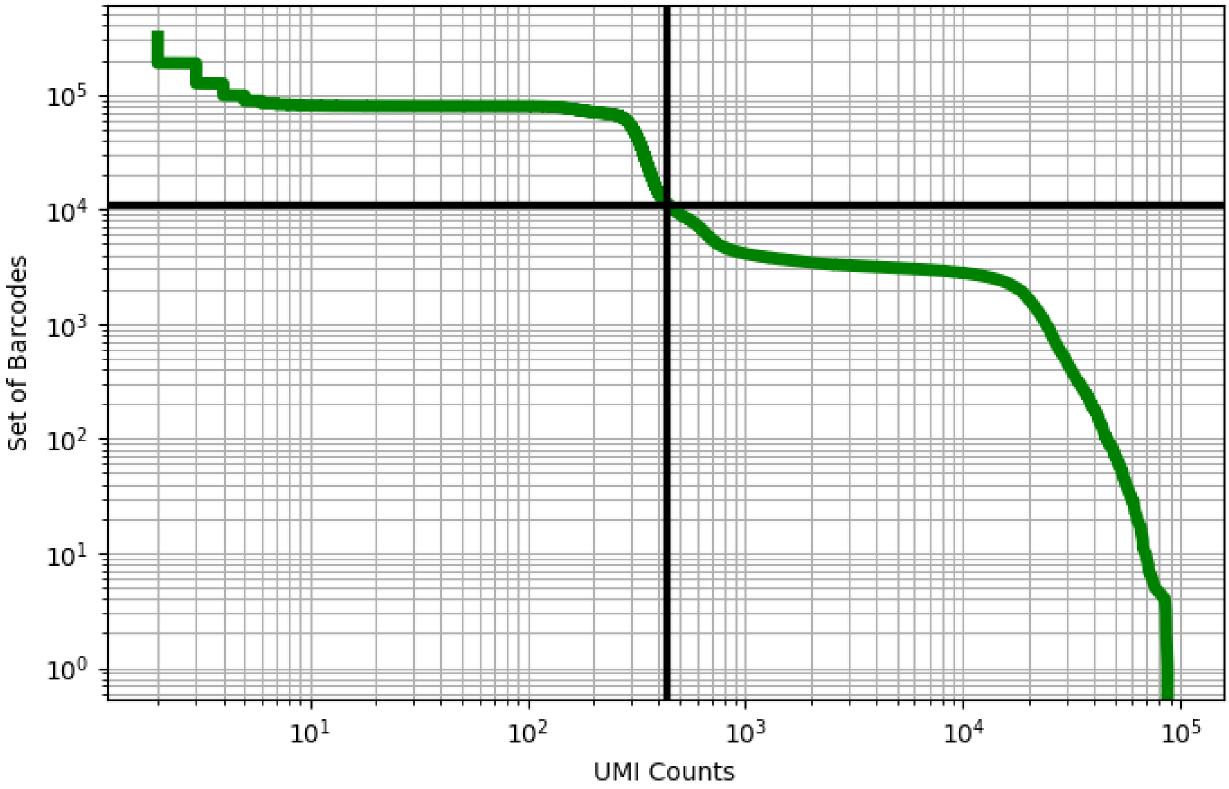


Figure 279: lung/GSM3439921

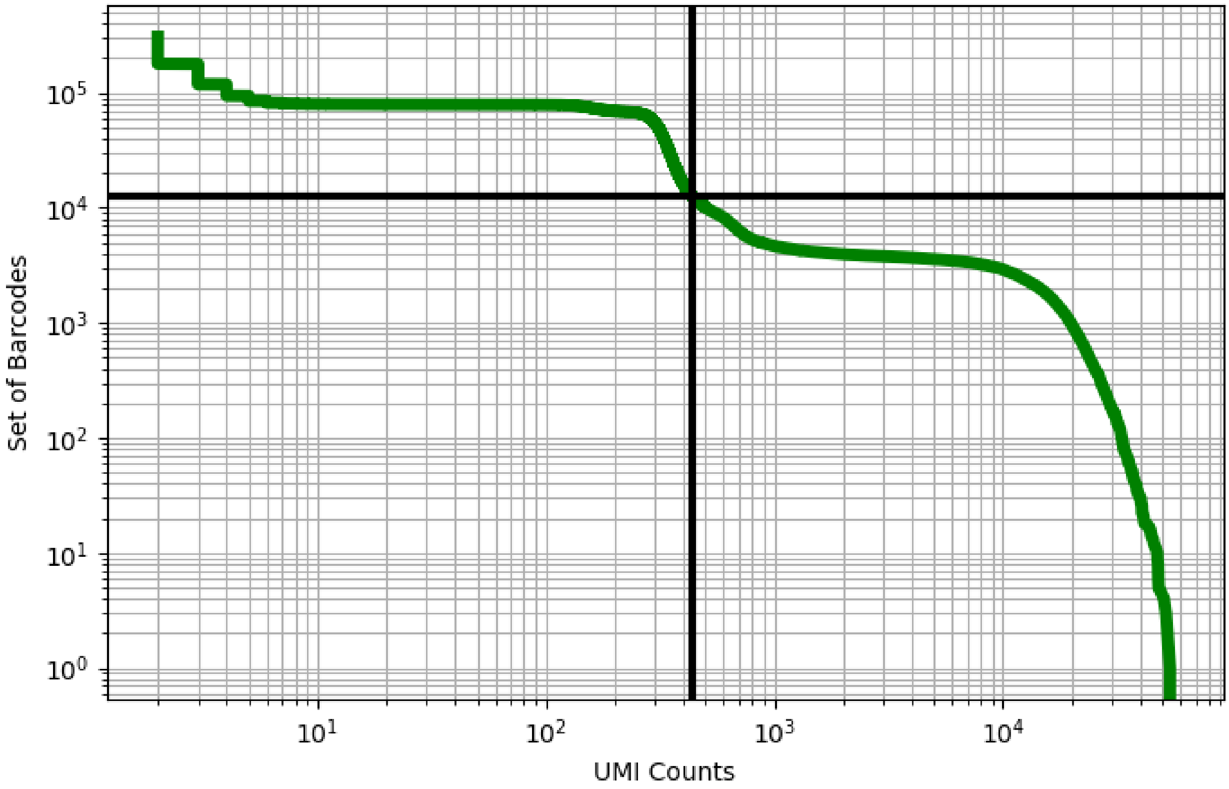


Figure 280: lung/GSM3439922

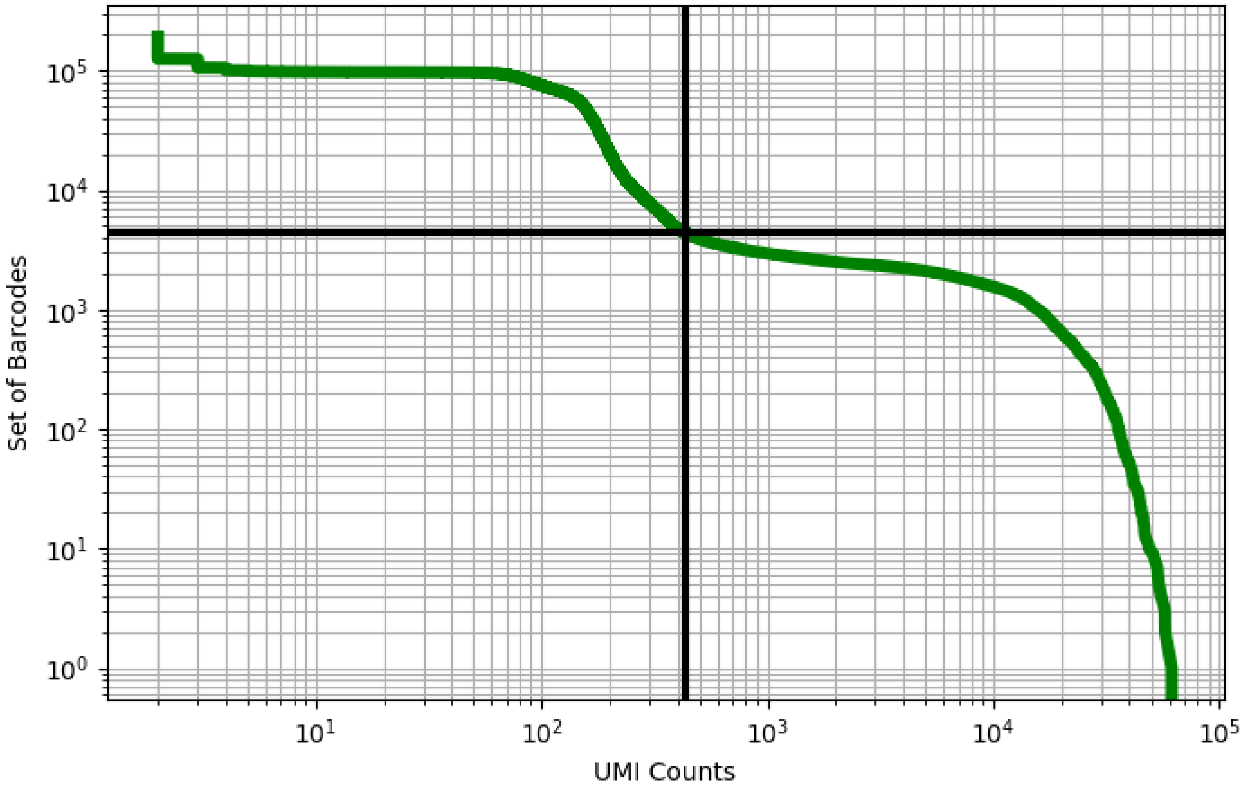


Figure 281: lung/GSM3439923

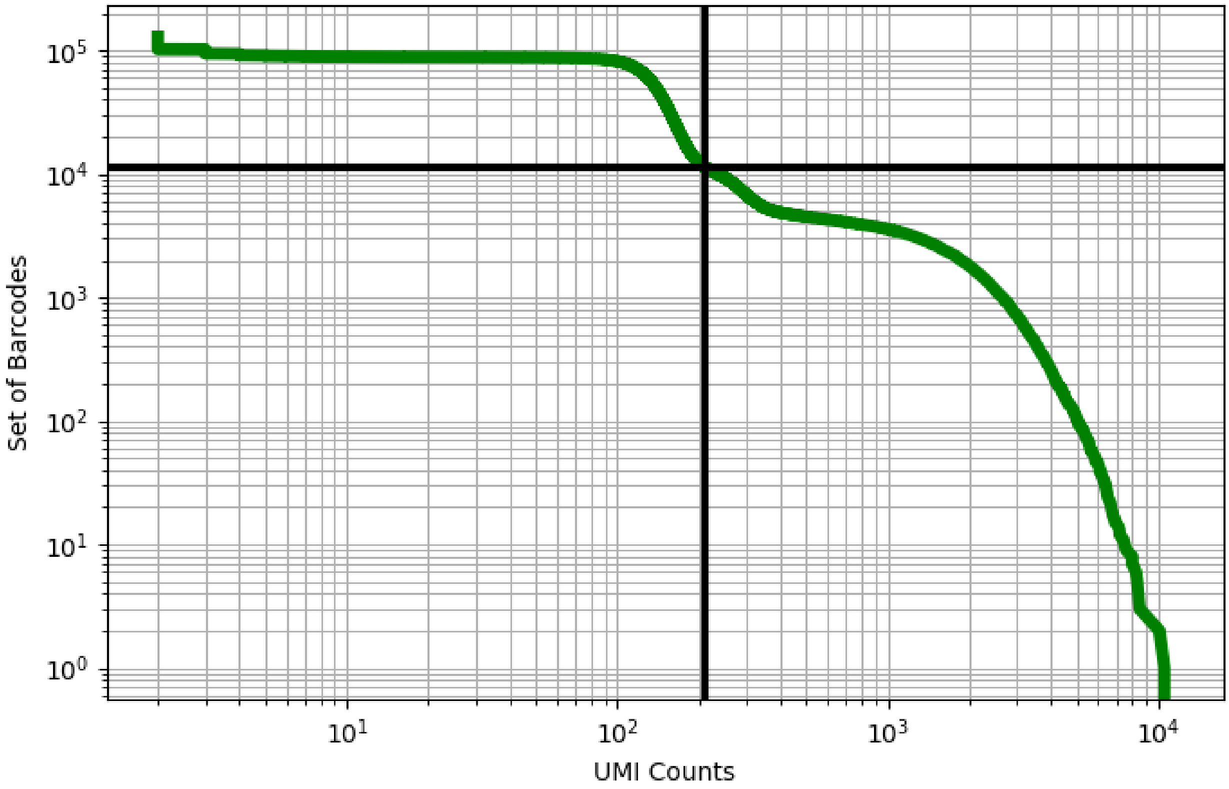


Figure 282: lung/GSM3439925

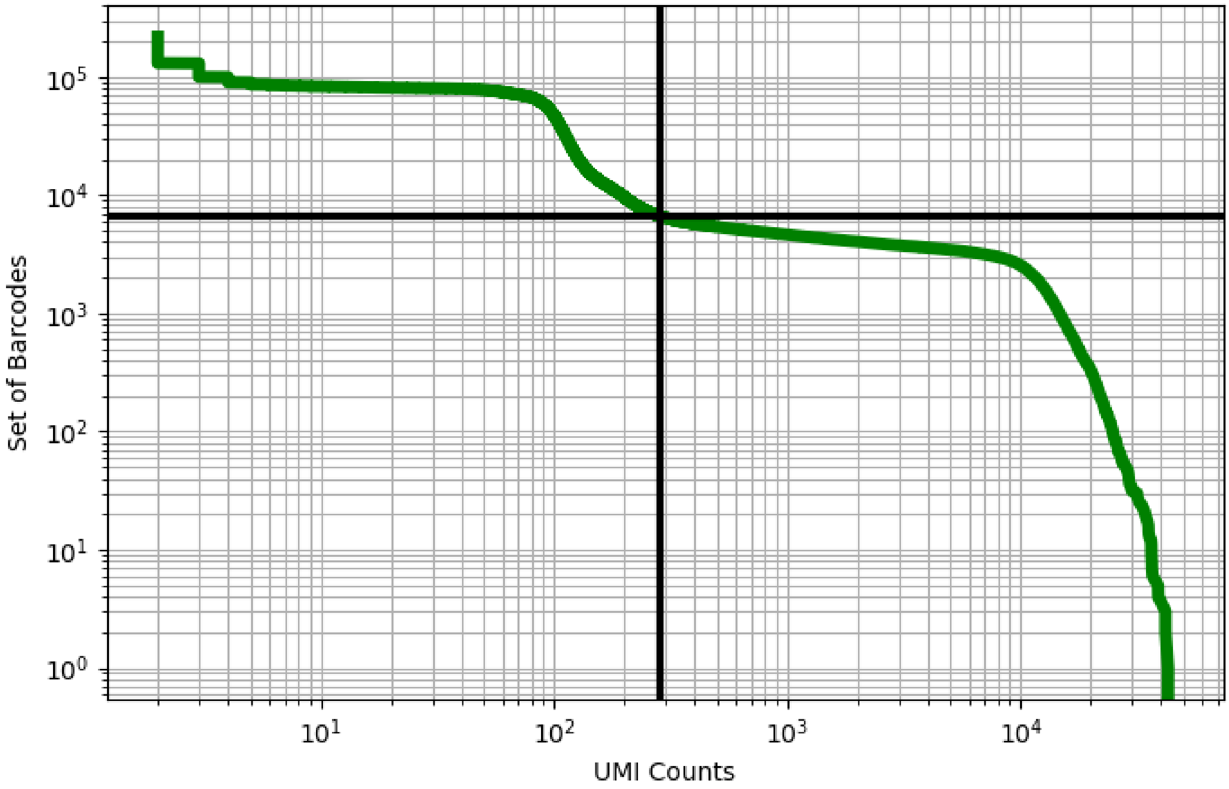


Figure 283: lung/GSM3439926

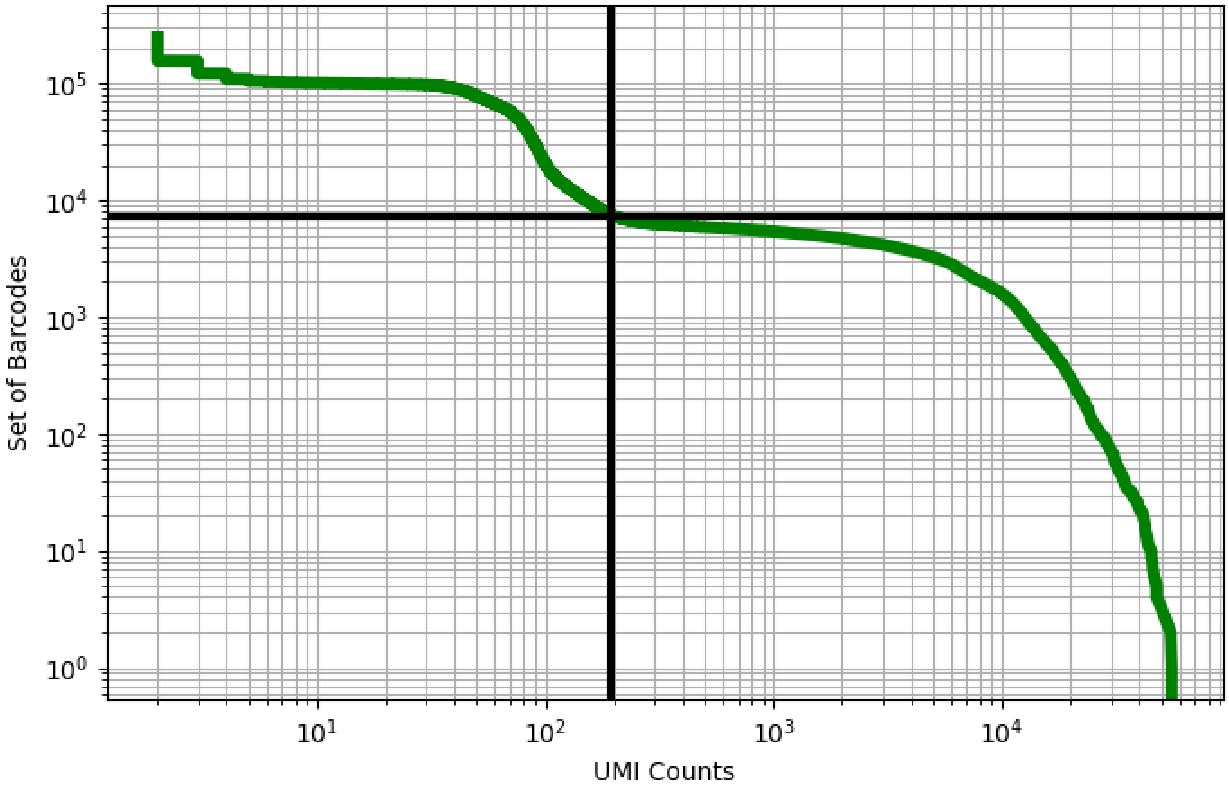


Figure 284: lung/GSM3439927

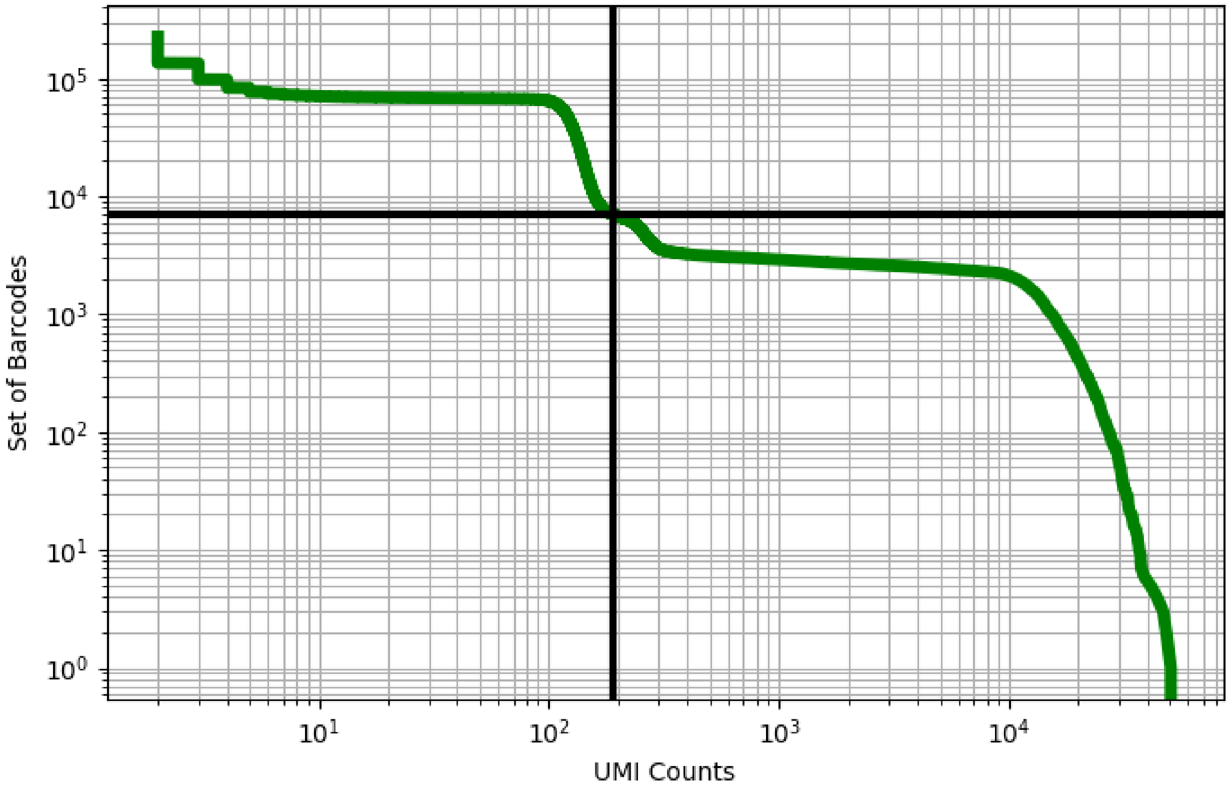


Figure 285: lung/GSM3439987

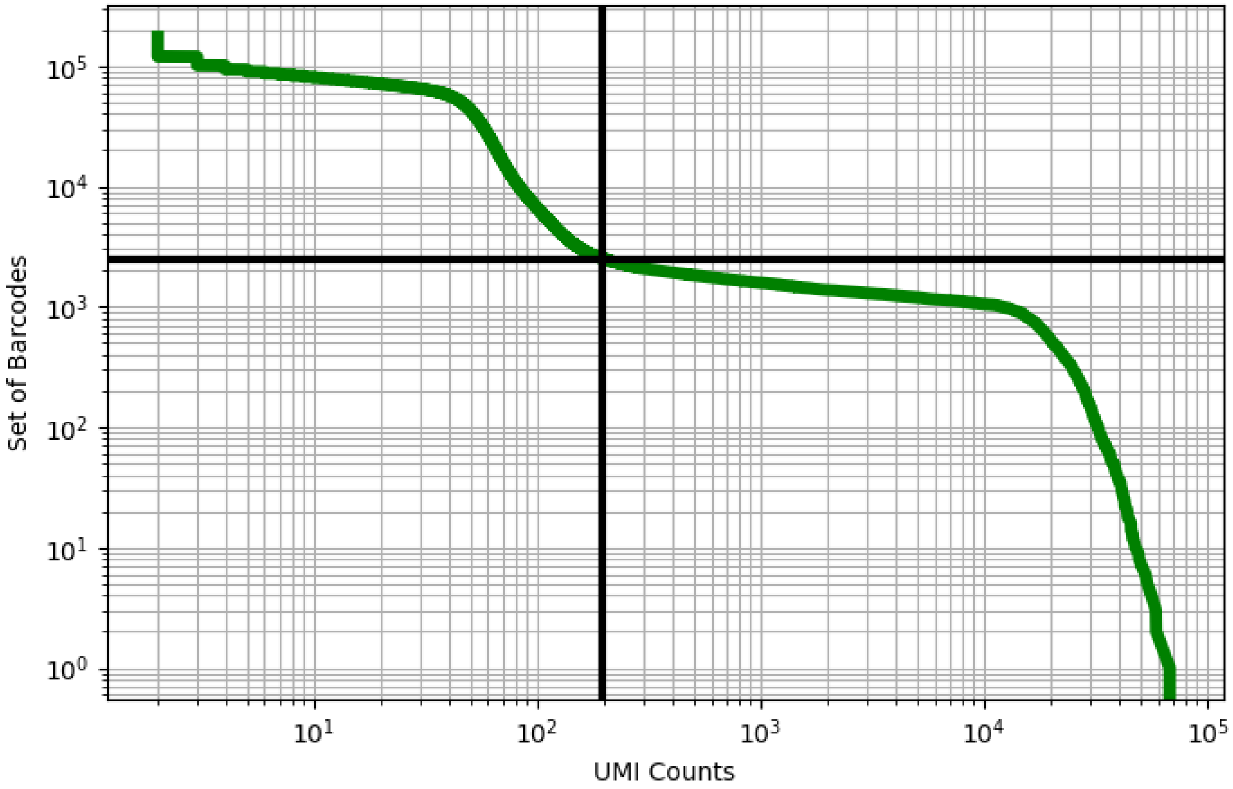


Figure 286: lung/GSM3439988

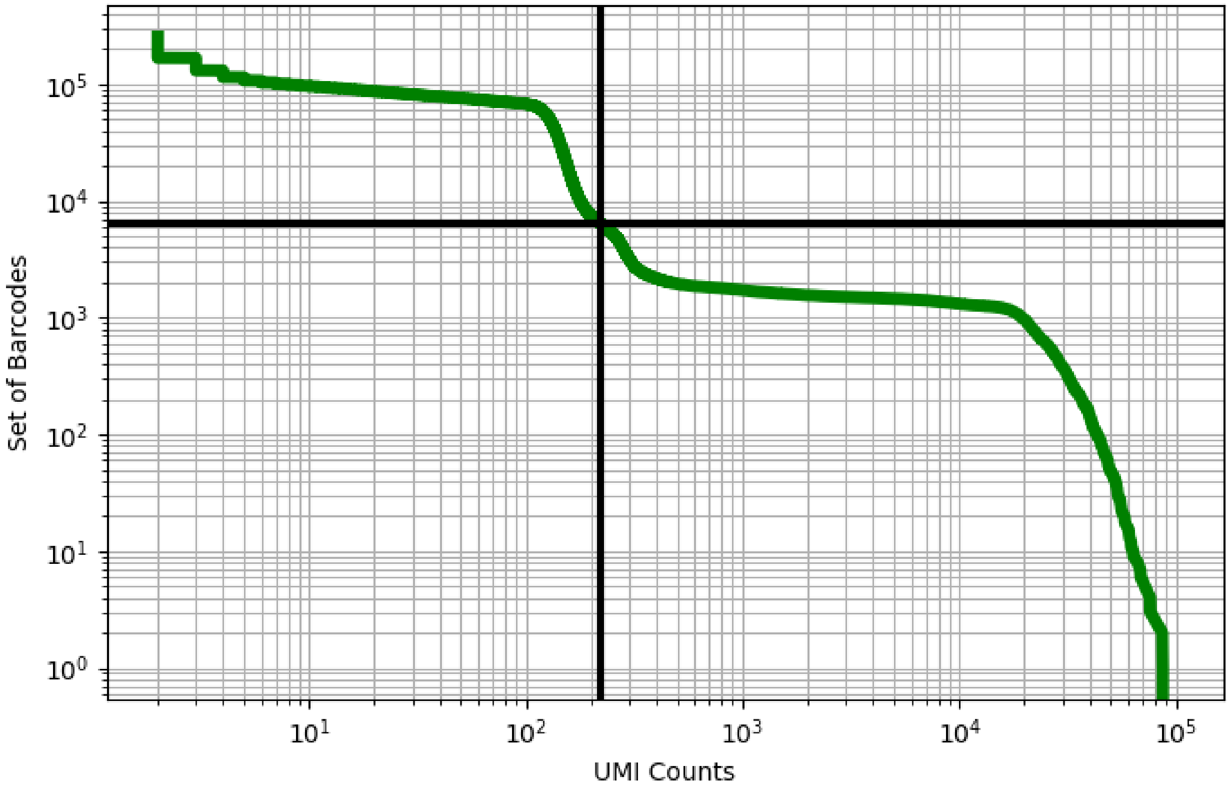


Figure 287: lung/GSM3439989

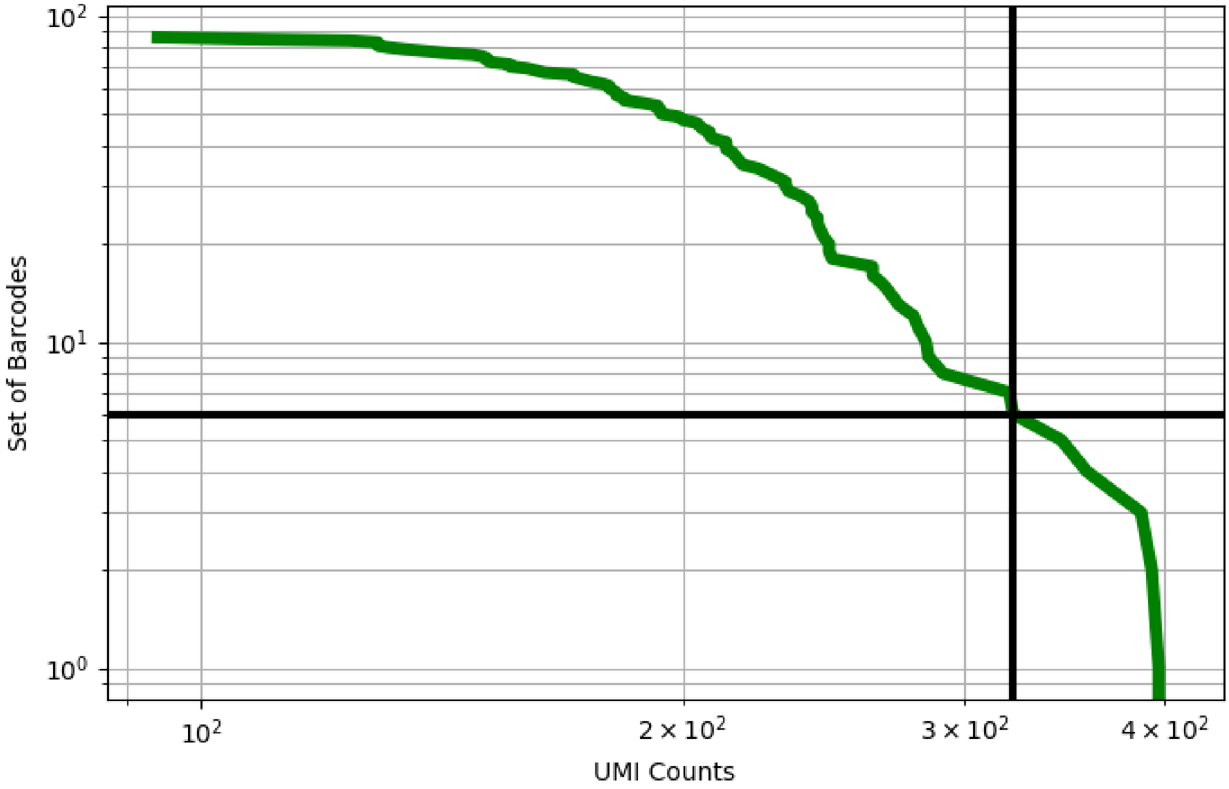


Figure 288: lung/GSM3773108

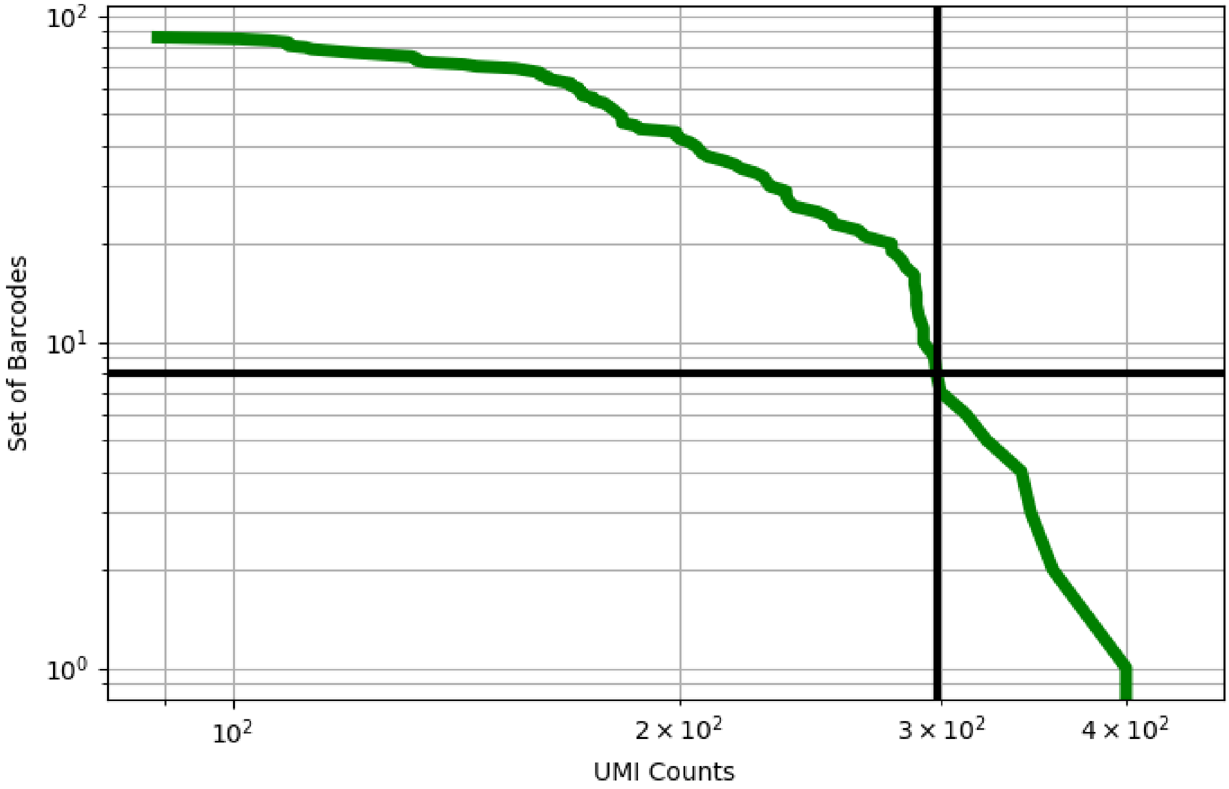


Figure 289: lung/GSM3773111

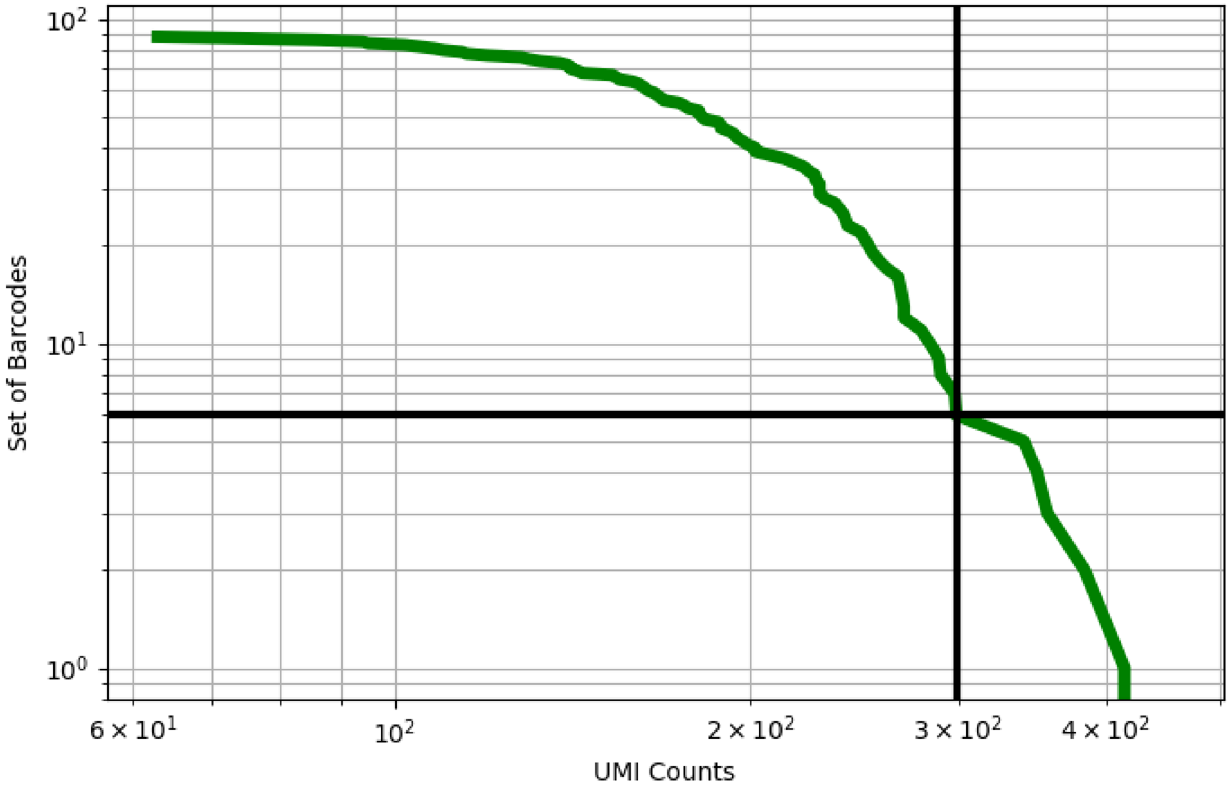


Figure 290: lung/GSM3773112

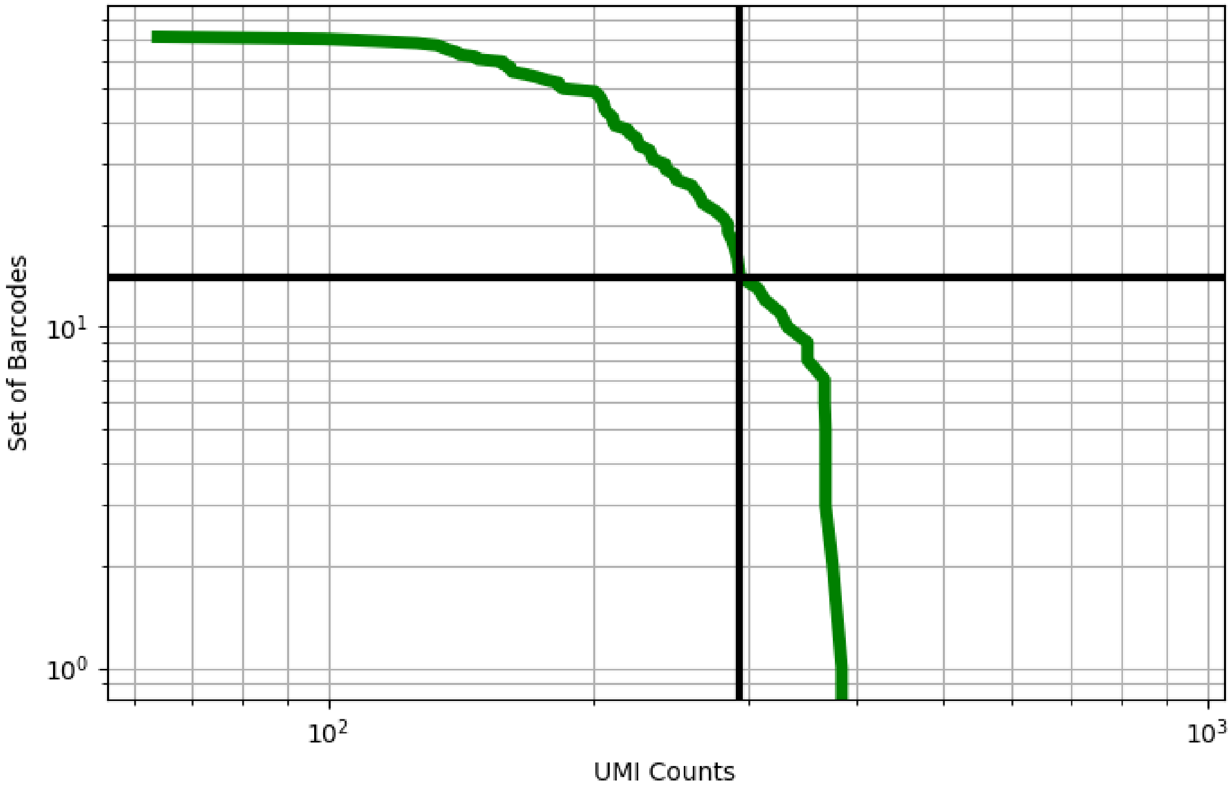


Figure 291: lung/GSM3773114

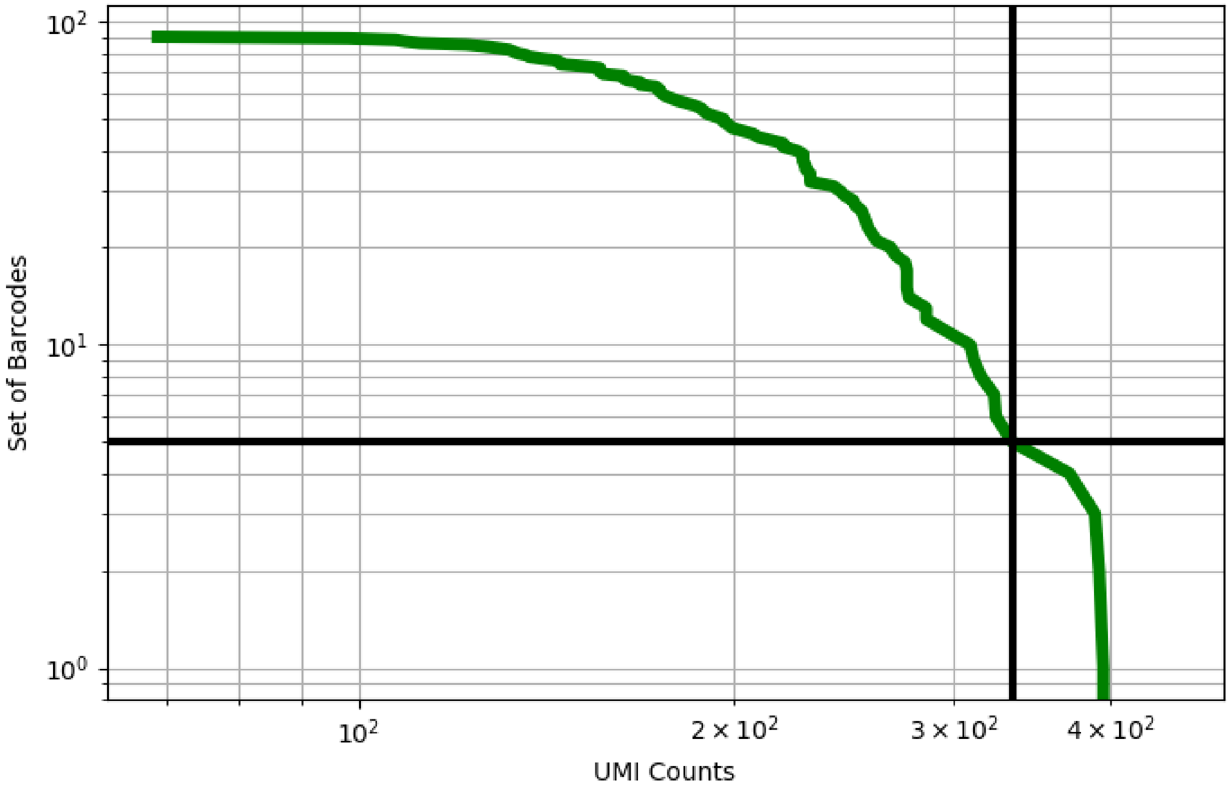


Figure 292: lung/GSM3773116

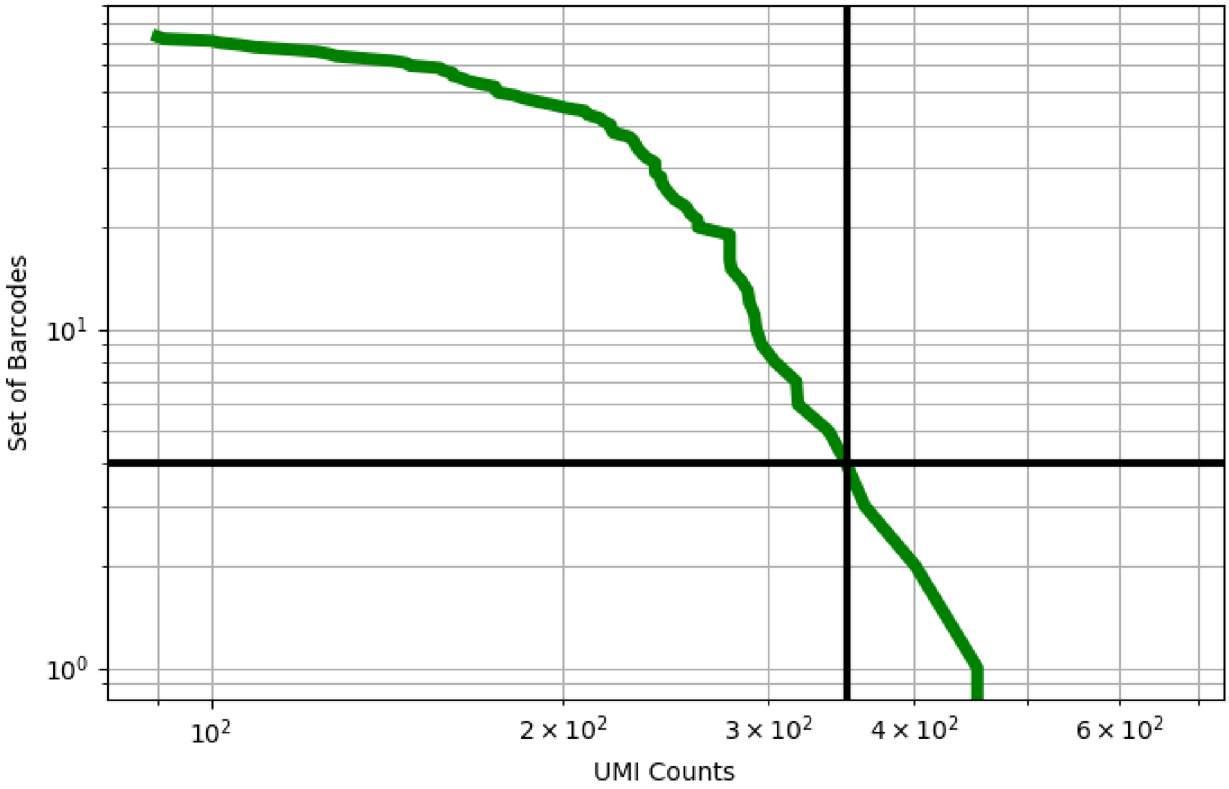


Figure 293: lung/GSM3773117

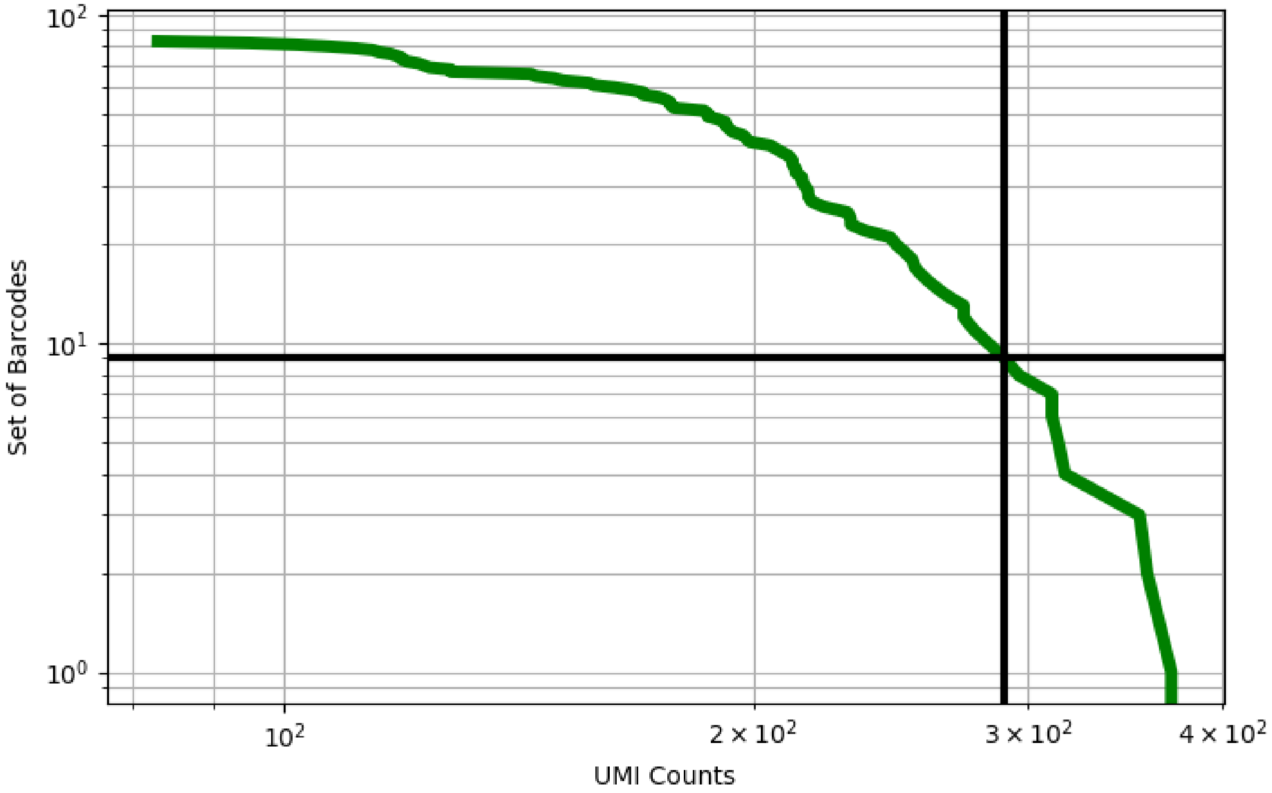


Figure 294: lung/GSM3773119

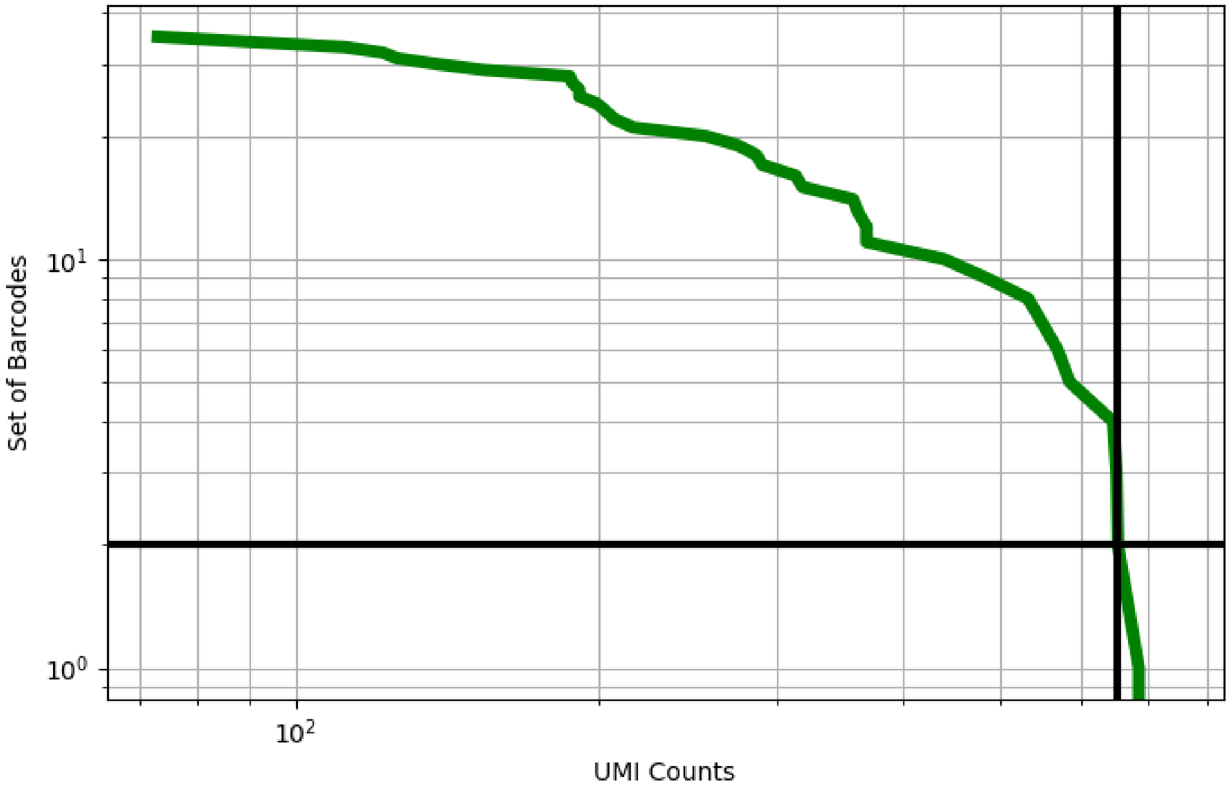


Figure 295: lung/GSM3773121

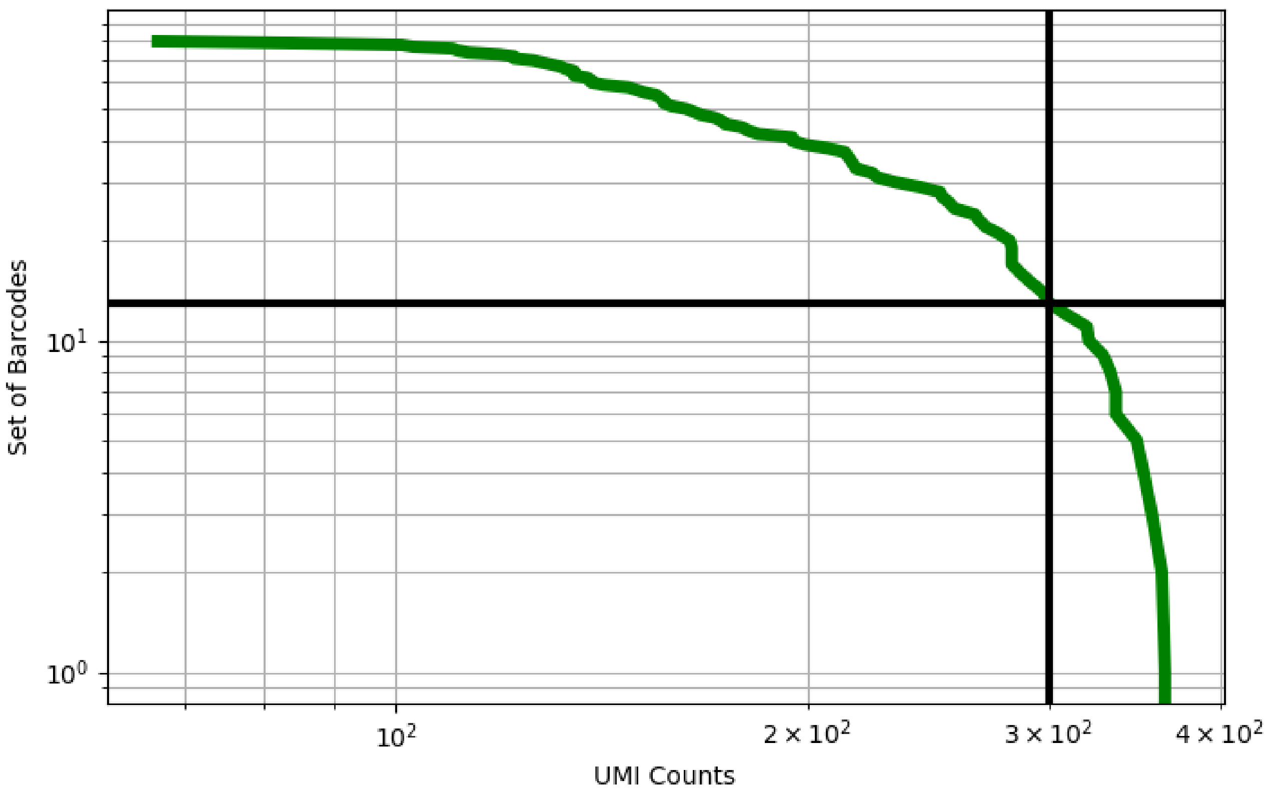


Figure 296: lung/GSM3773124

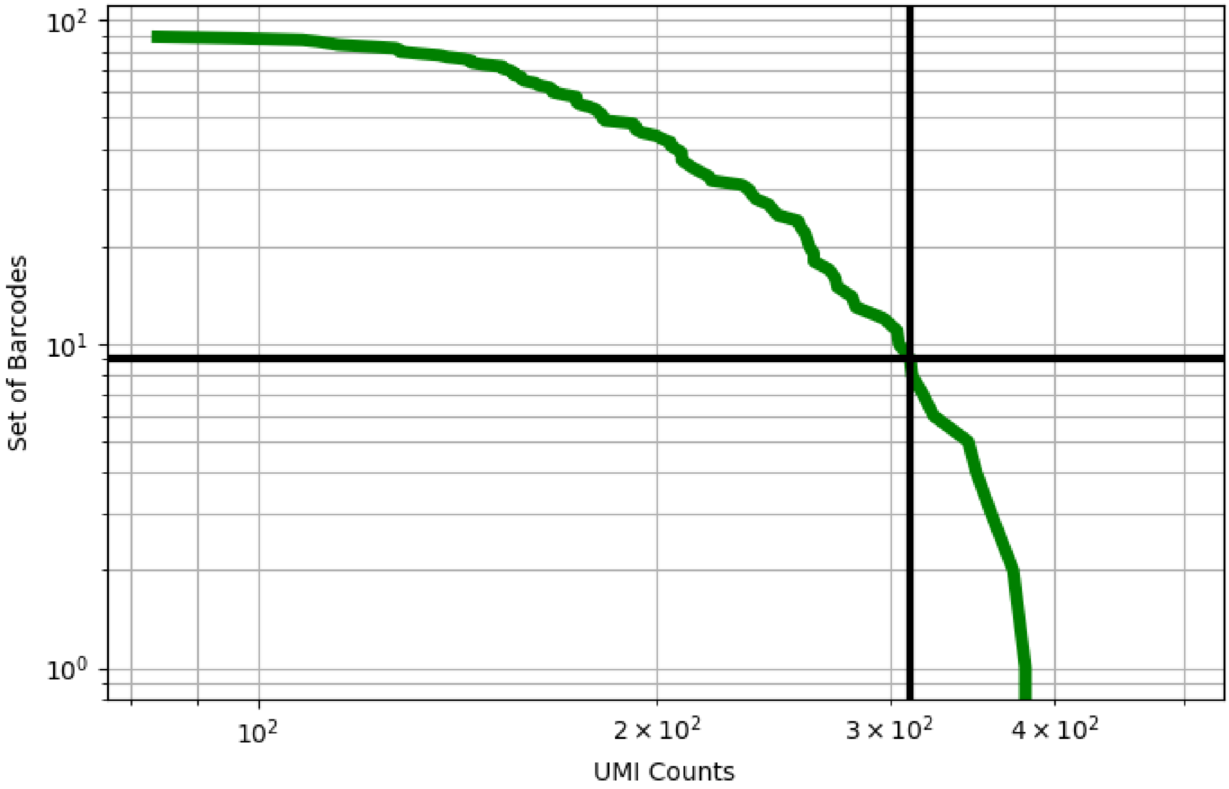


Figure 297: lung/GSM3773125

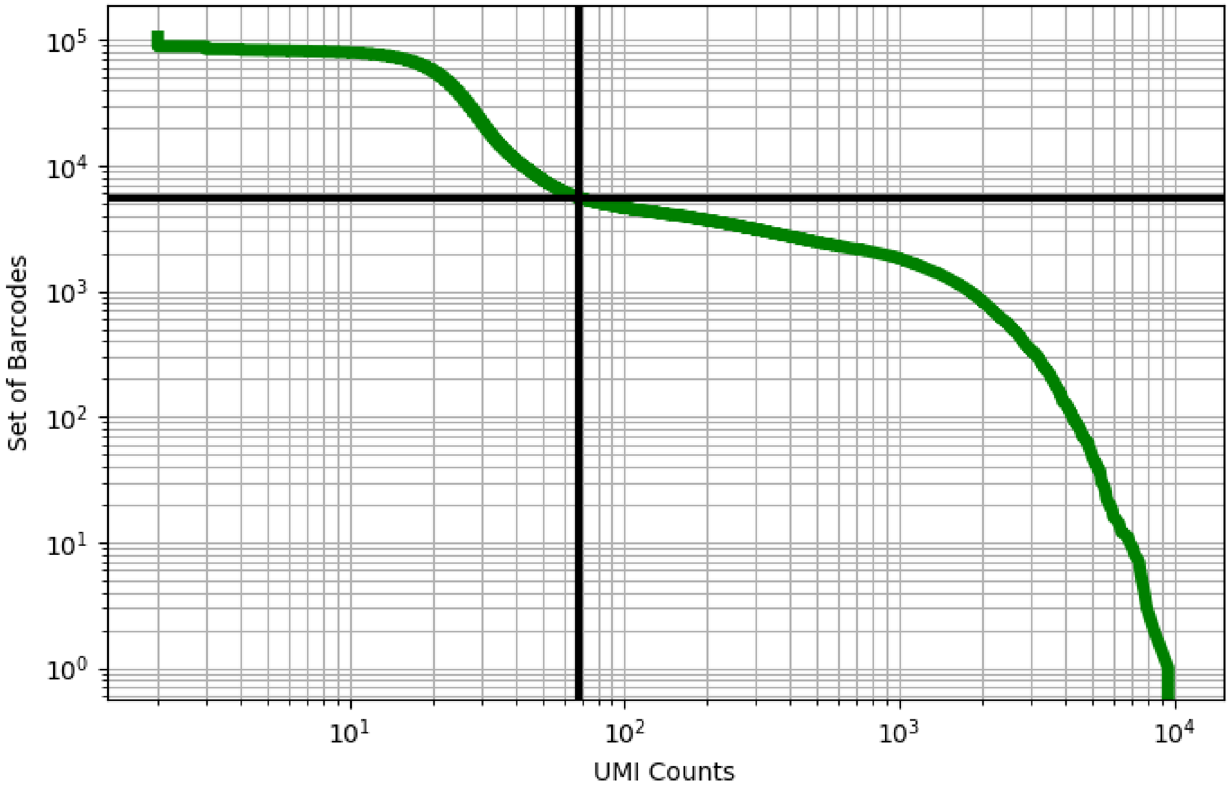


Figure 298: lung/GSM4037301

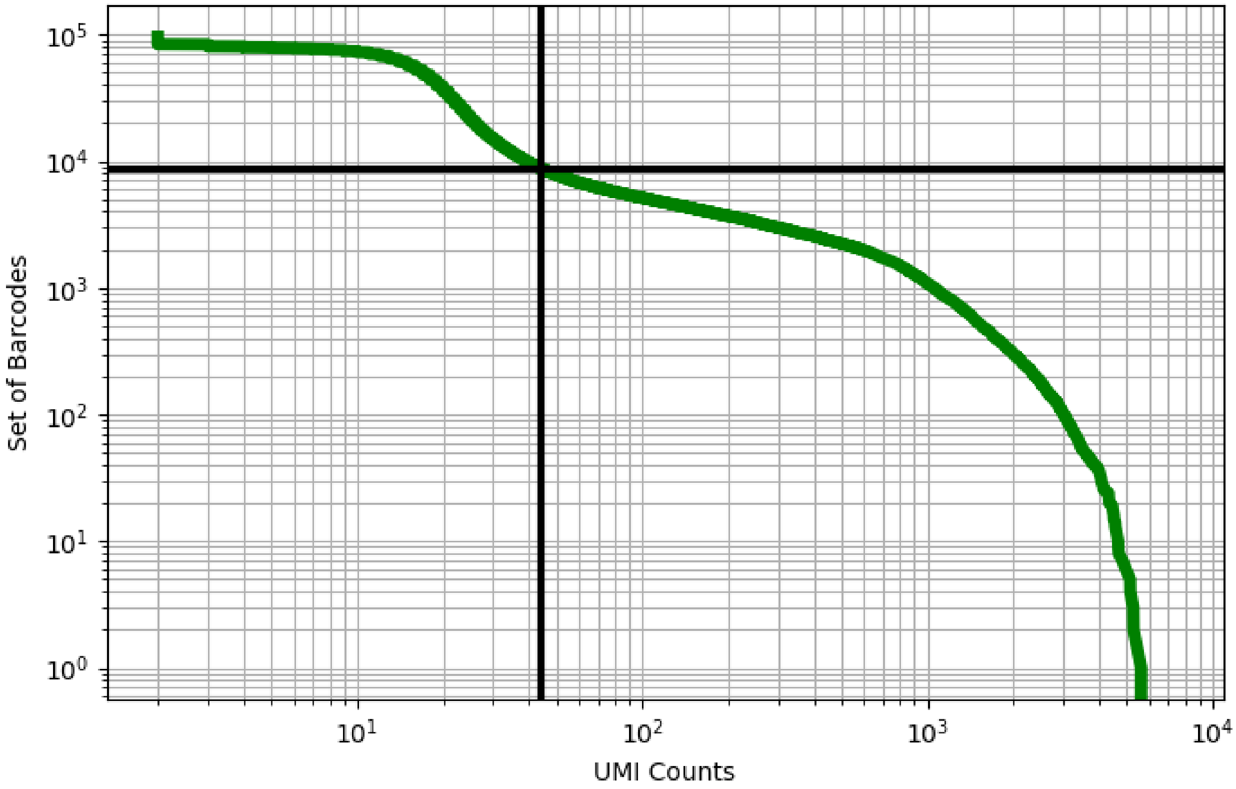


Figure 299: lung/GSM4037302

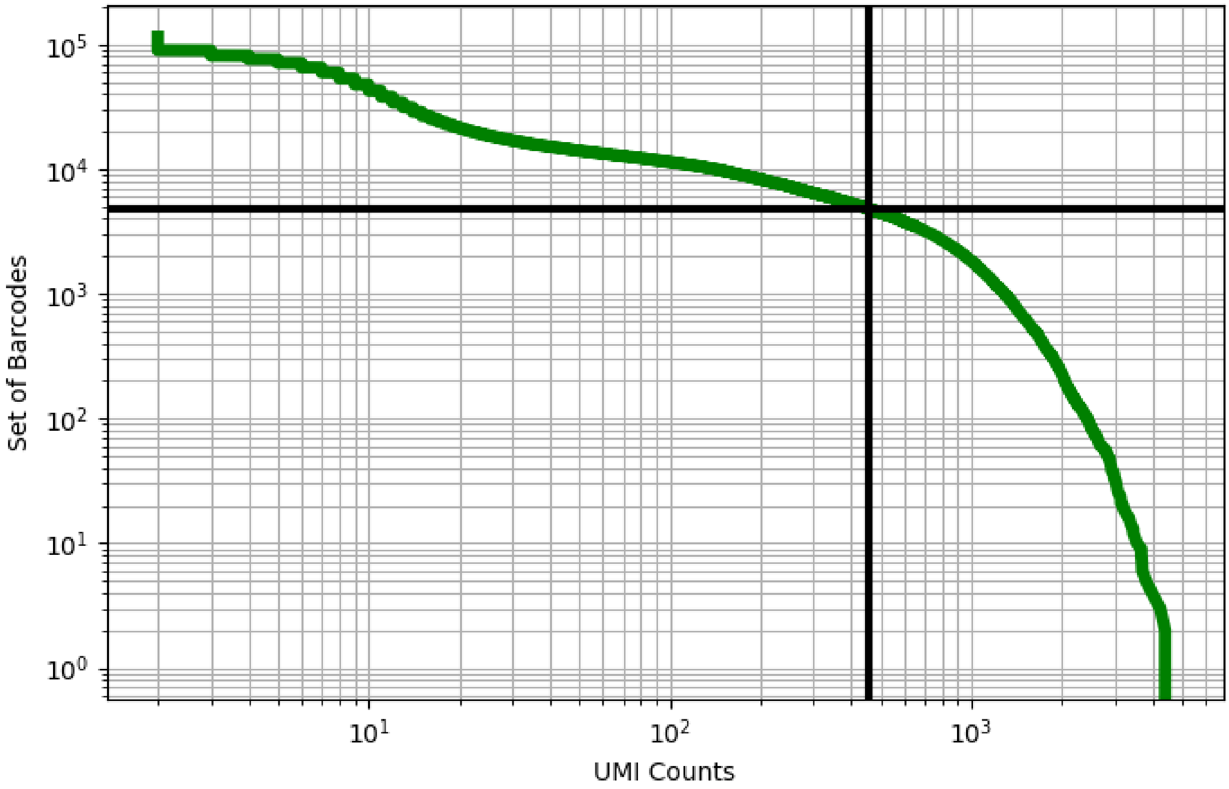


Figure 300: lung/GSM4037305

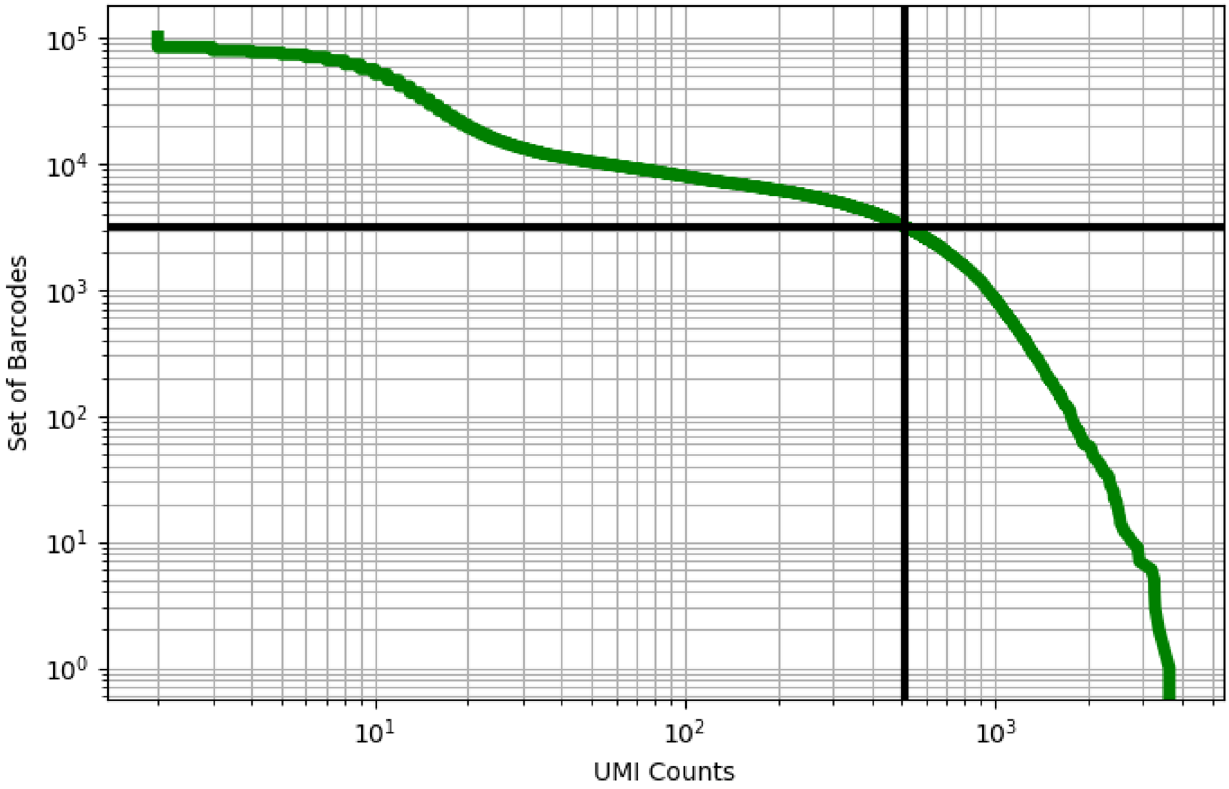


Figure 301: lung/GSM4037306

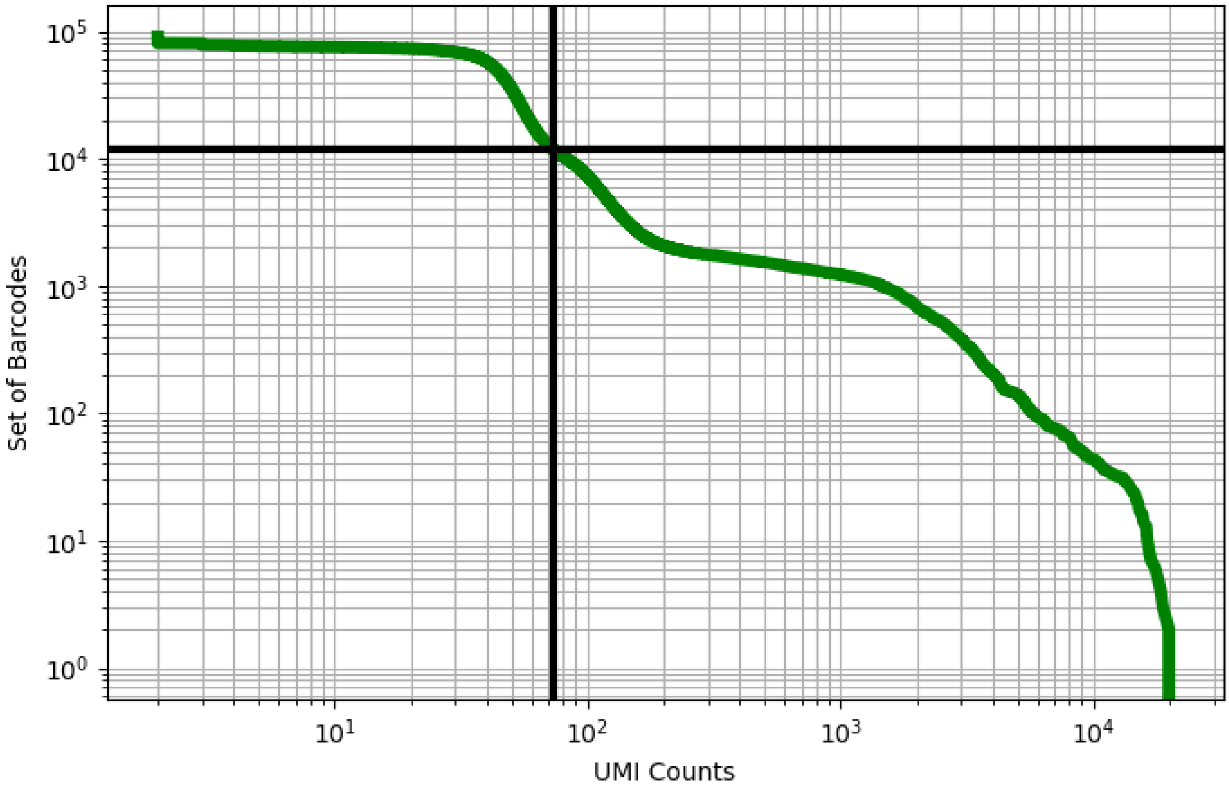


Figure 302: lung/GSM4037309

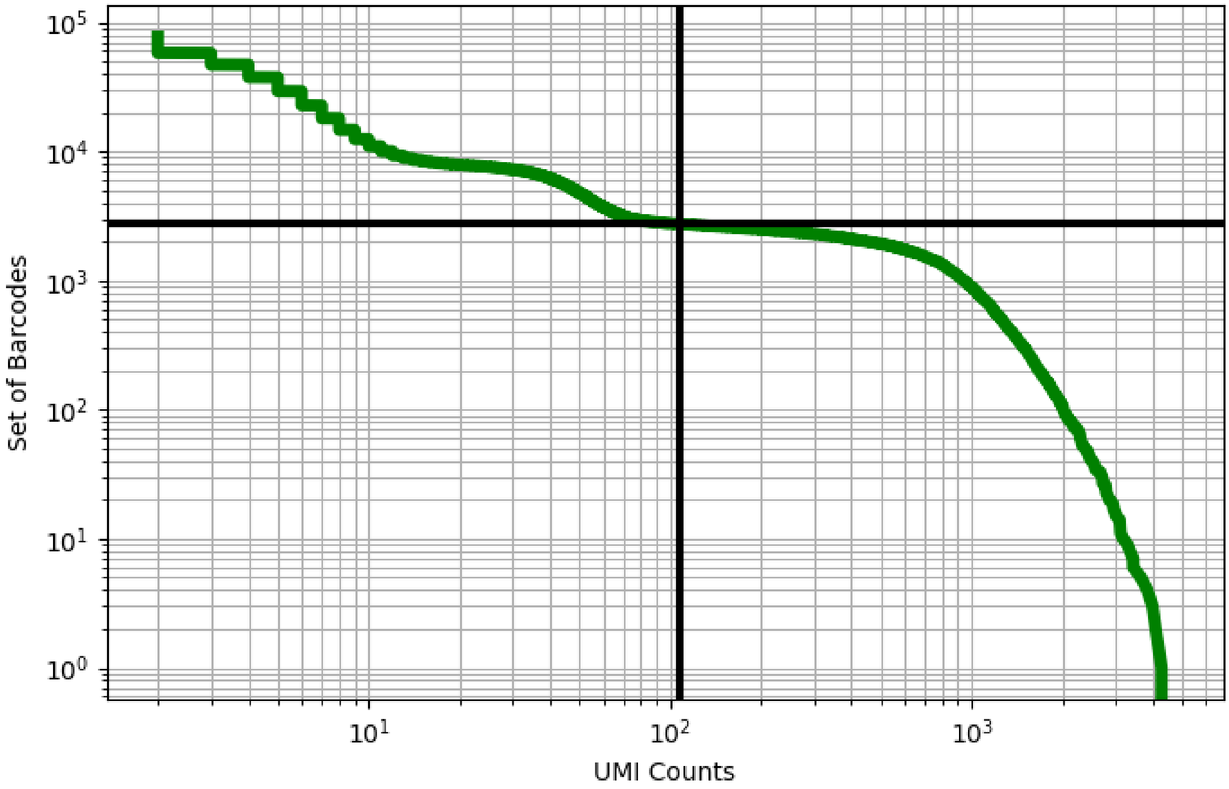


Figure 303: lung/GSM4037313

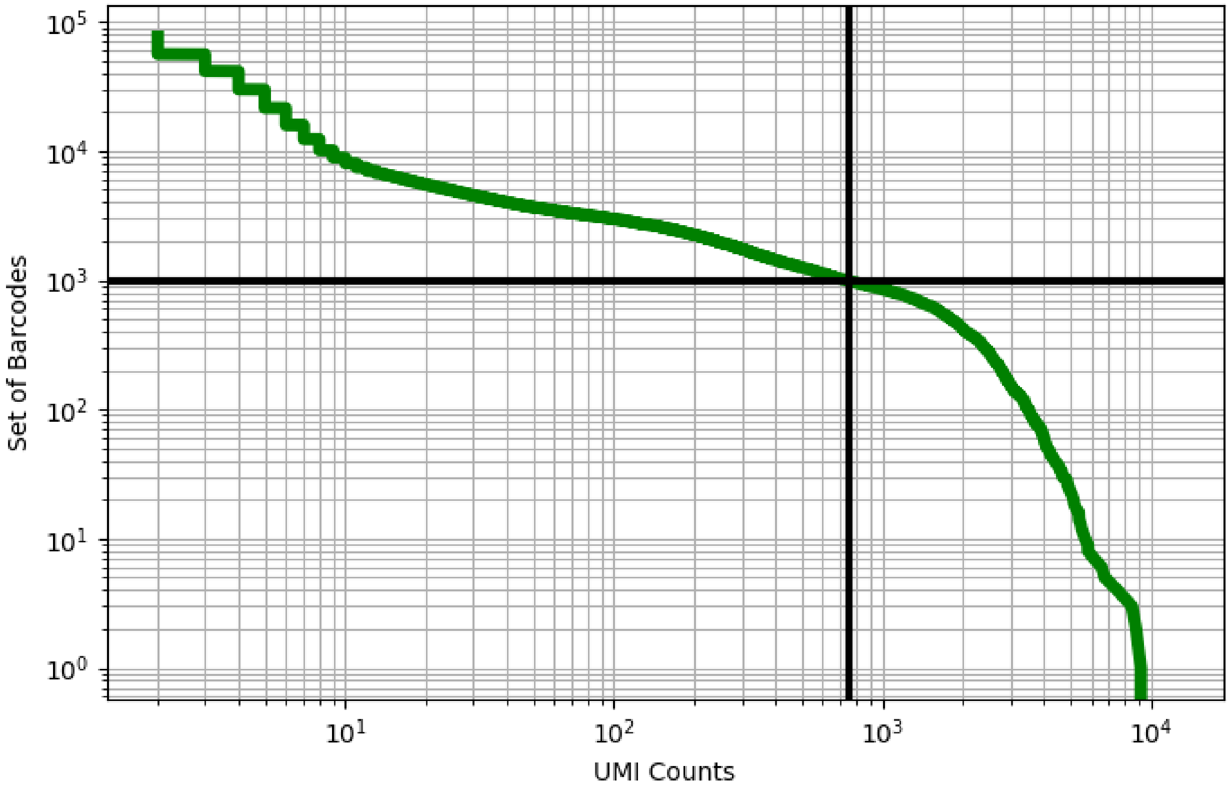


Figure 304: lung/GSM4037316

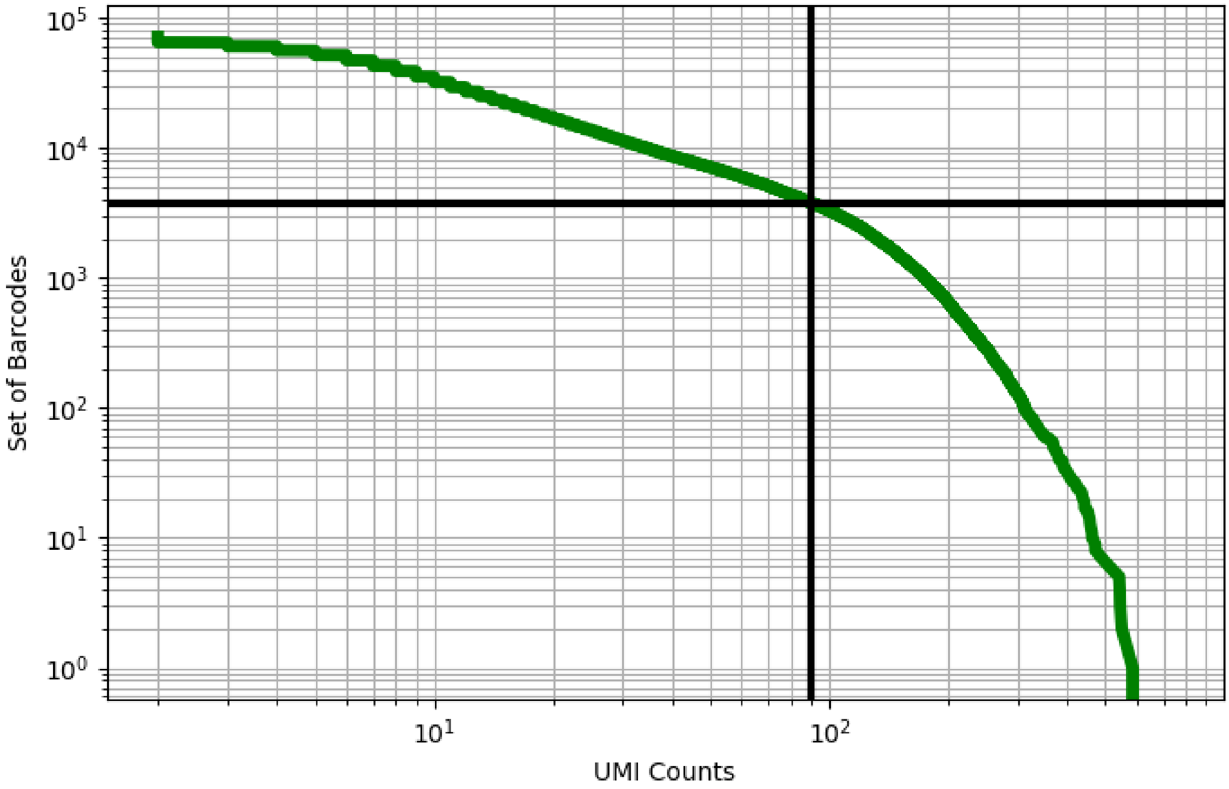


Figure 305: lung/GSM4037323

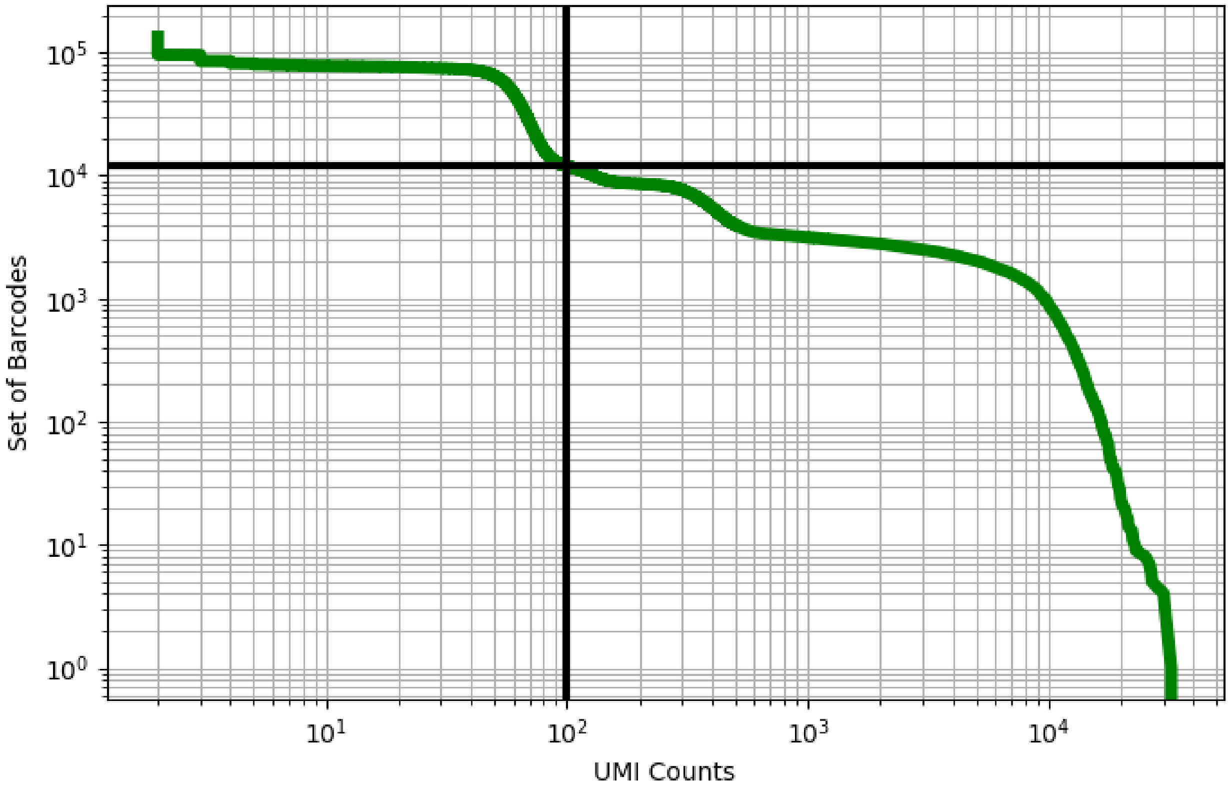


Figure 306: lung/GSM4037324

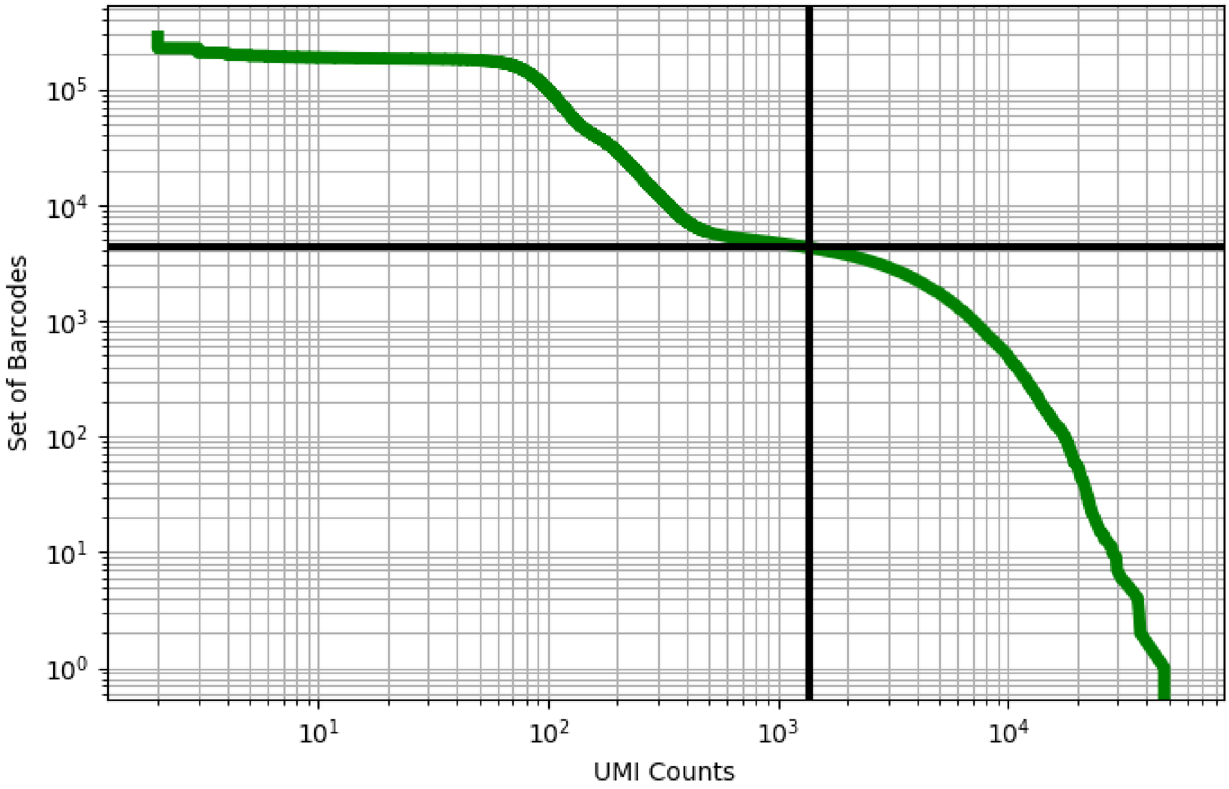


Figure 307: lung/GSM4037325

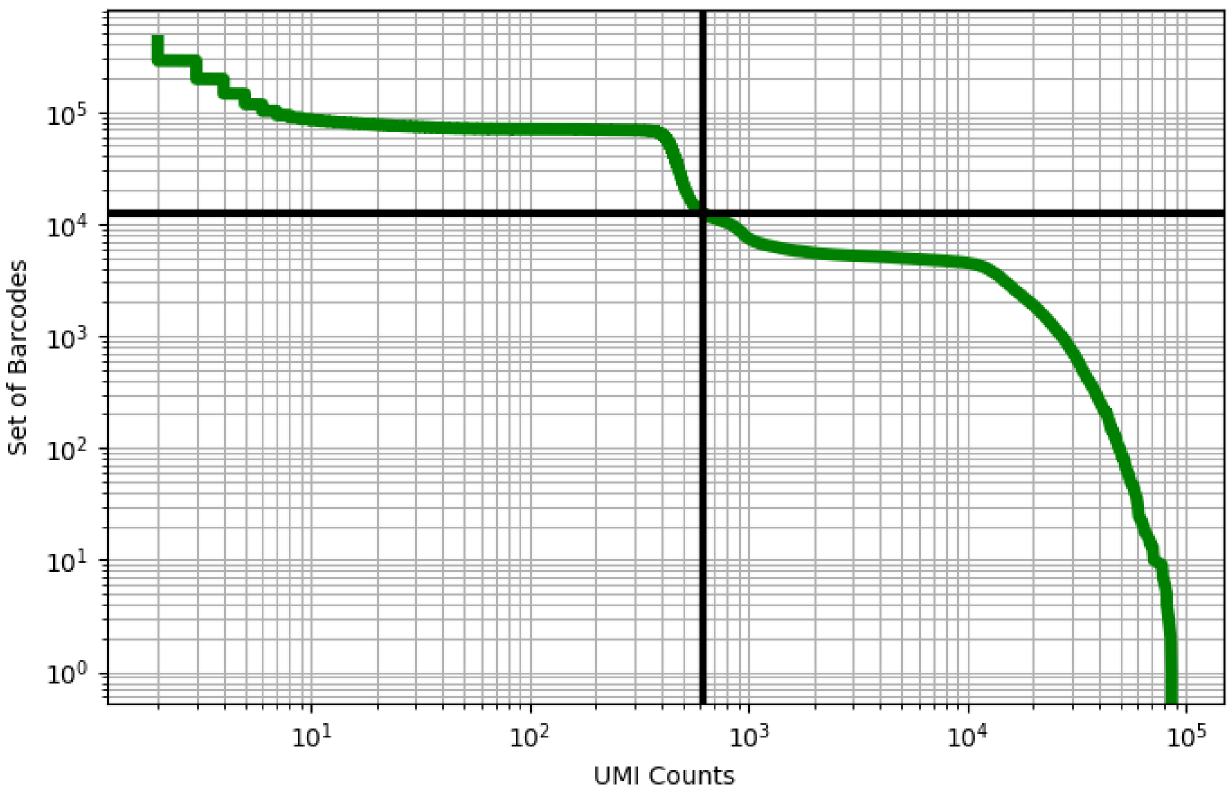


Figure 308: lung/GSM4080085

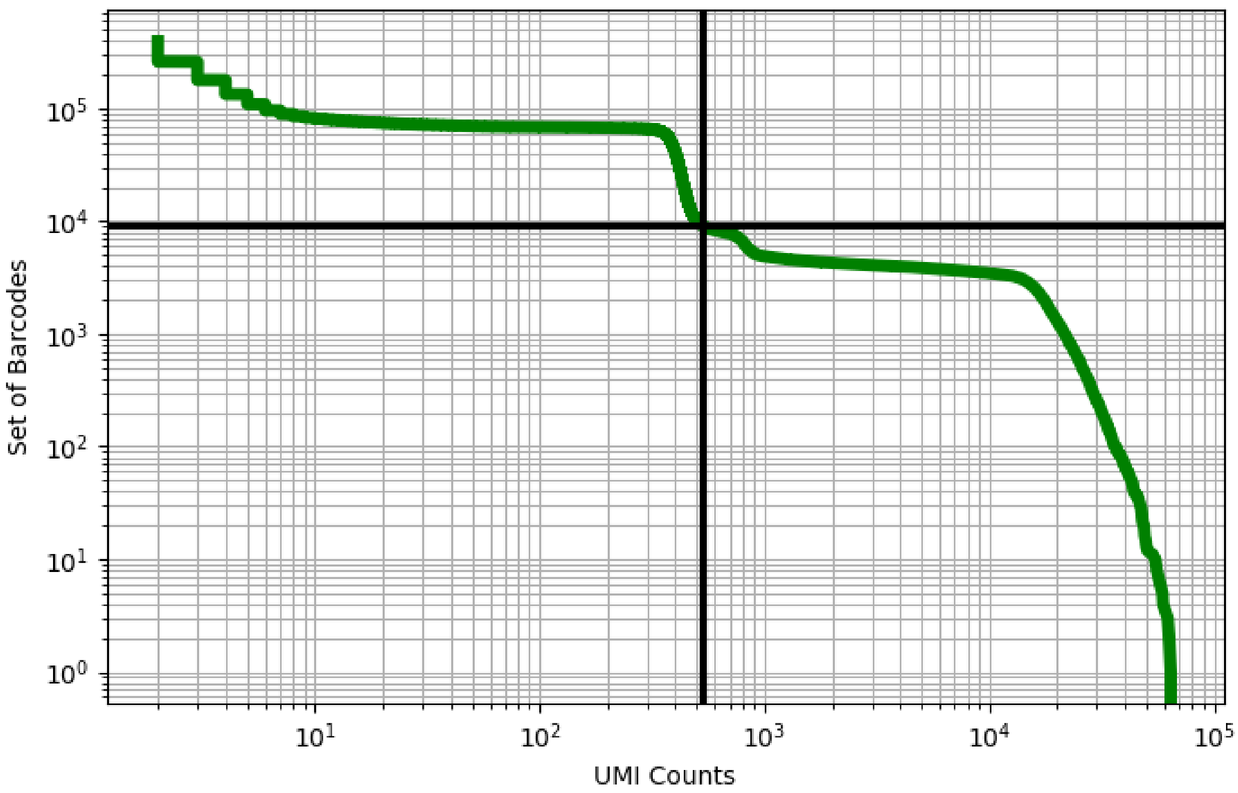


Figure 309: lung/GSM4080086

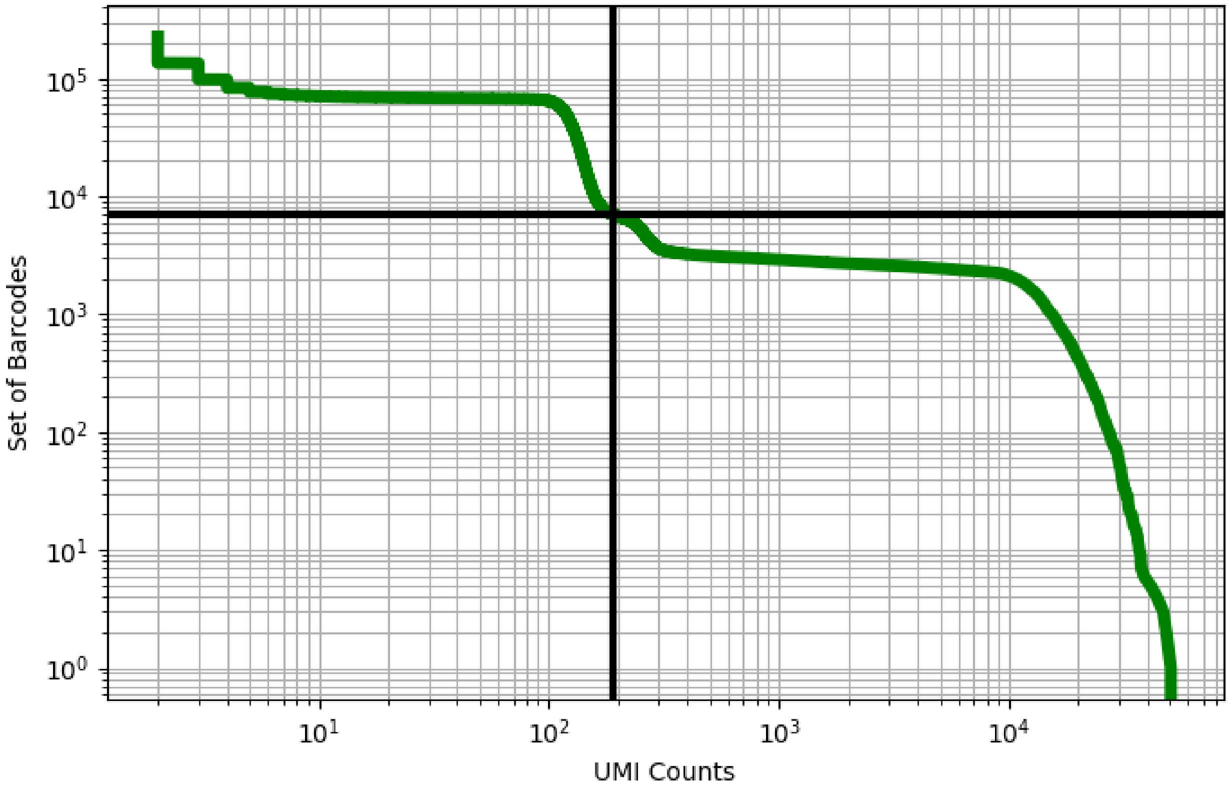


Figure 310: lung/GSM4080087

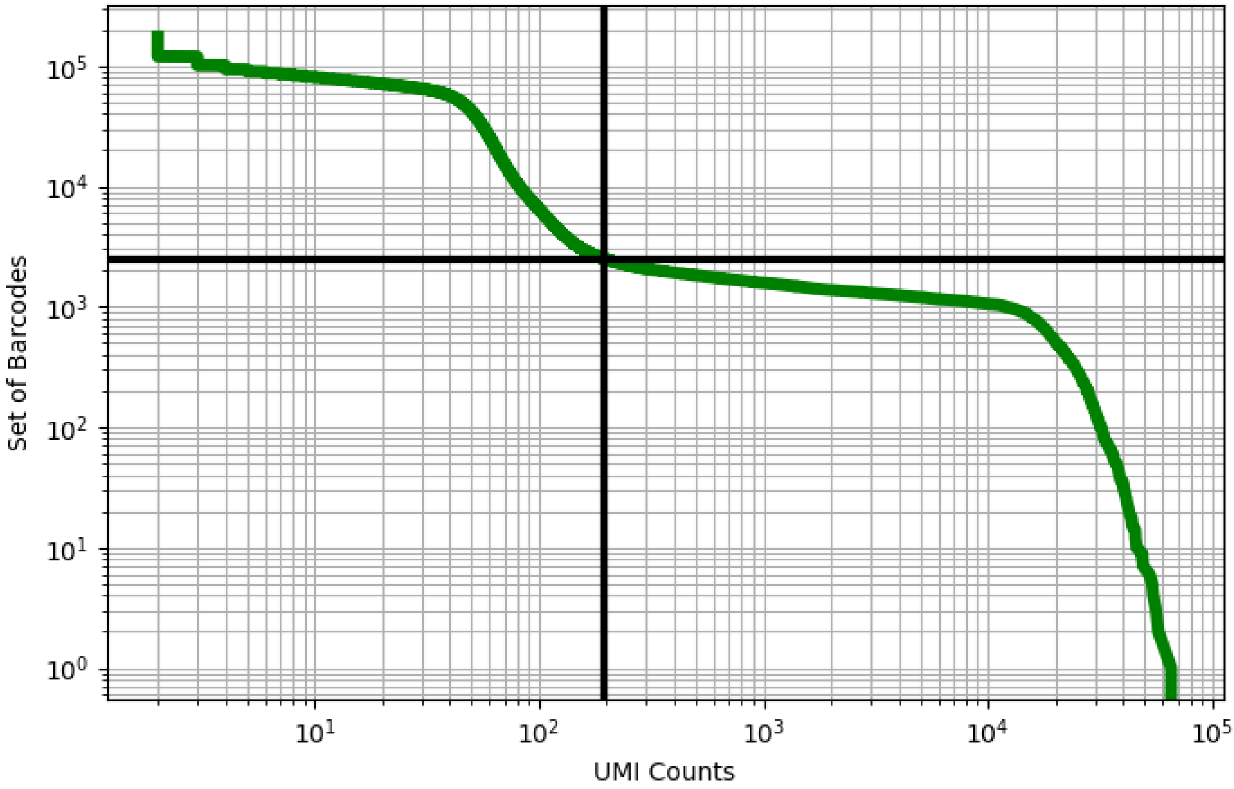


Figure 311: lung/GSM4080088

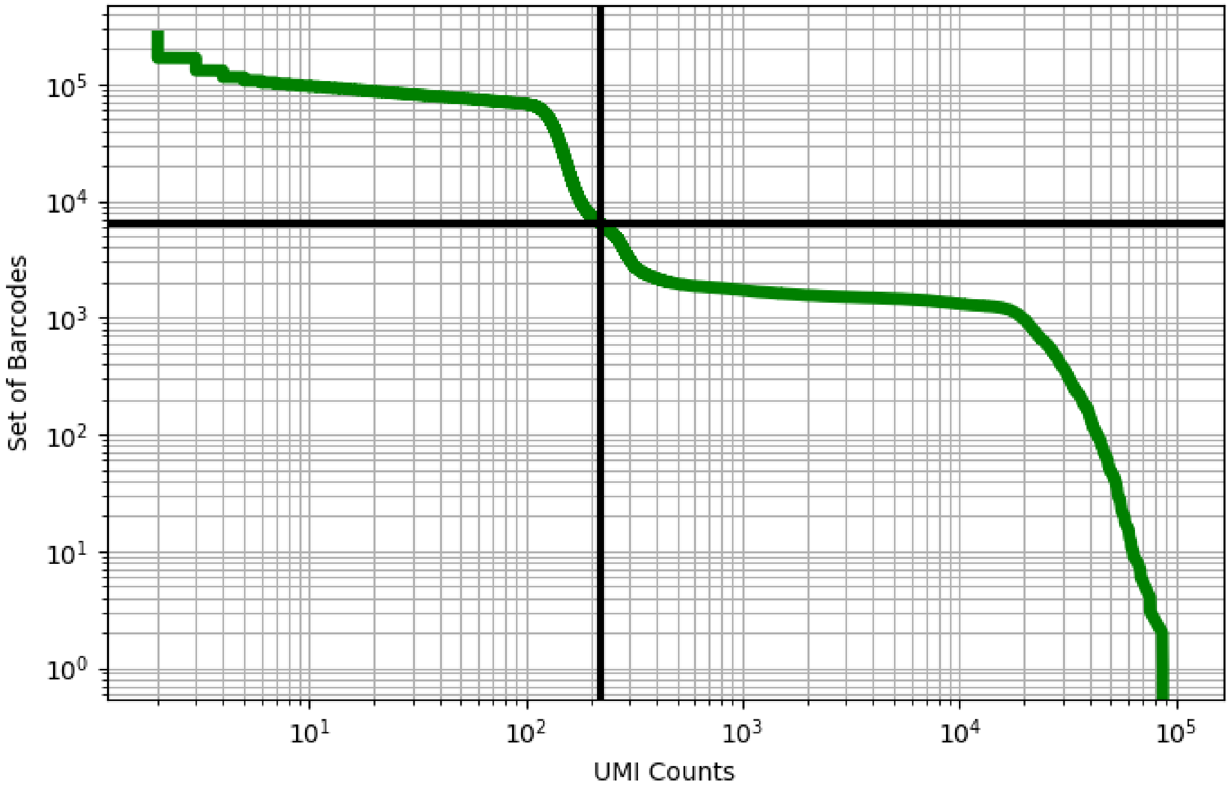


Figure 312: lung/GSM4080089

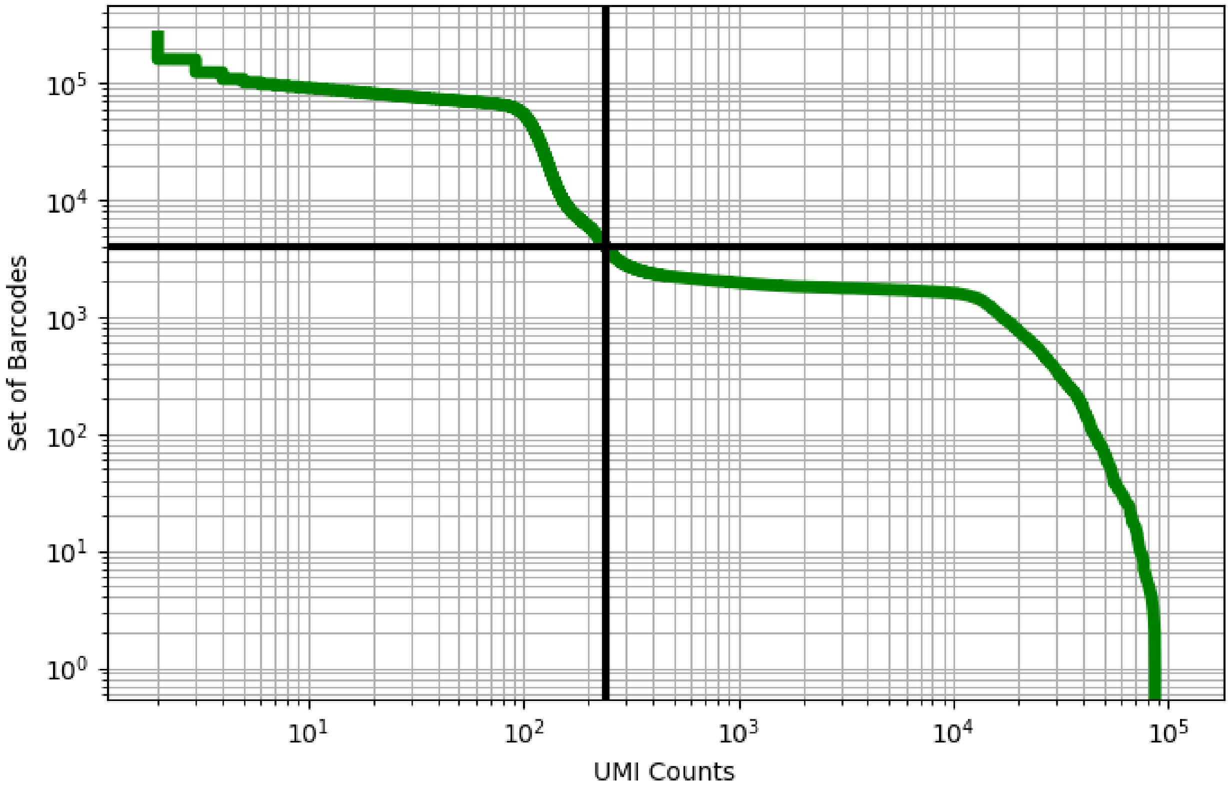


Figure 313: lung/GSM4080090

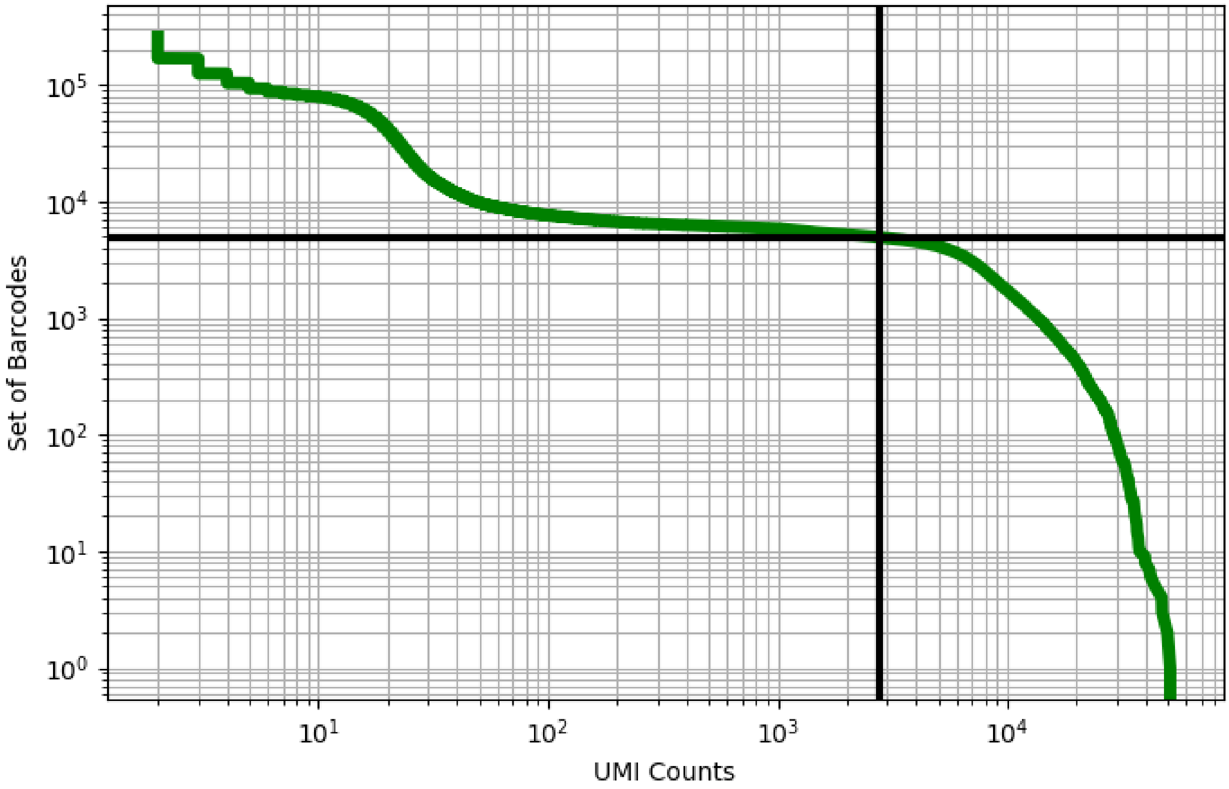


Figure 314: lung/GSM4142870

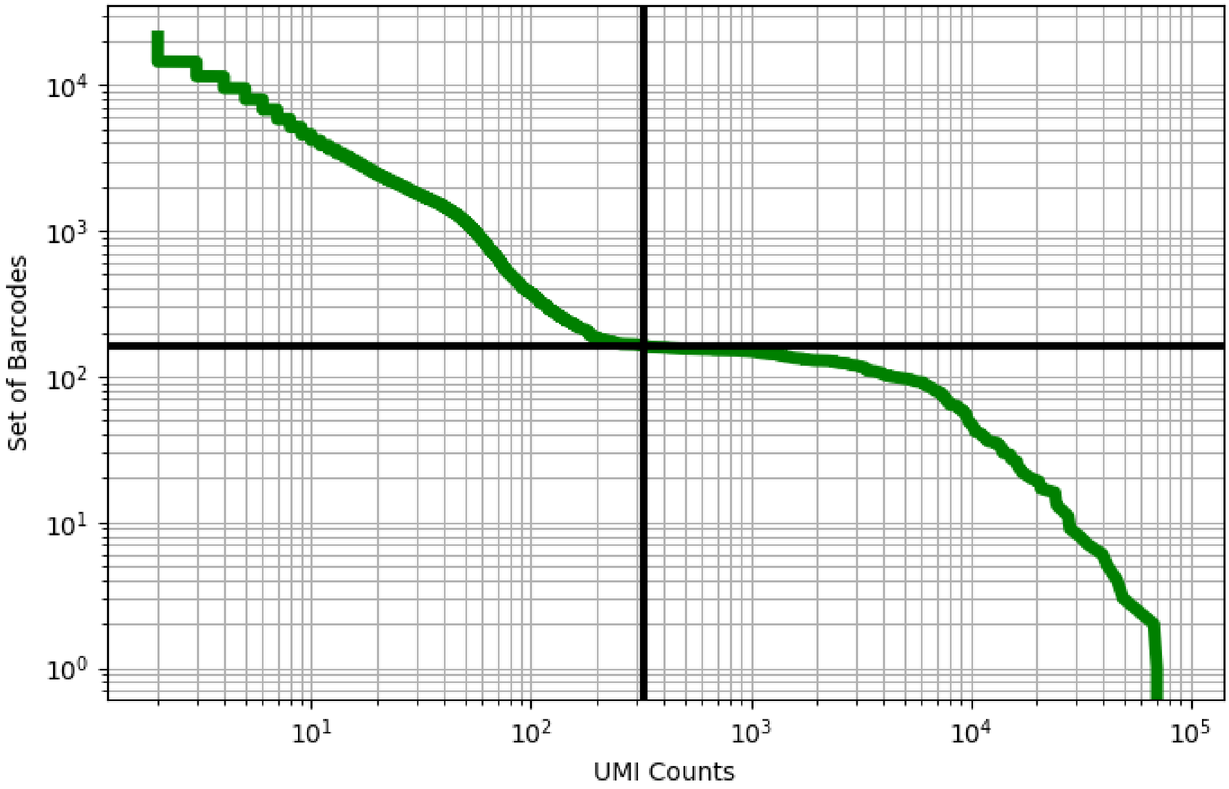


Figure 315: lung/GSM4142871

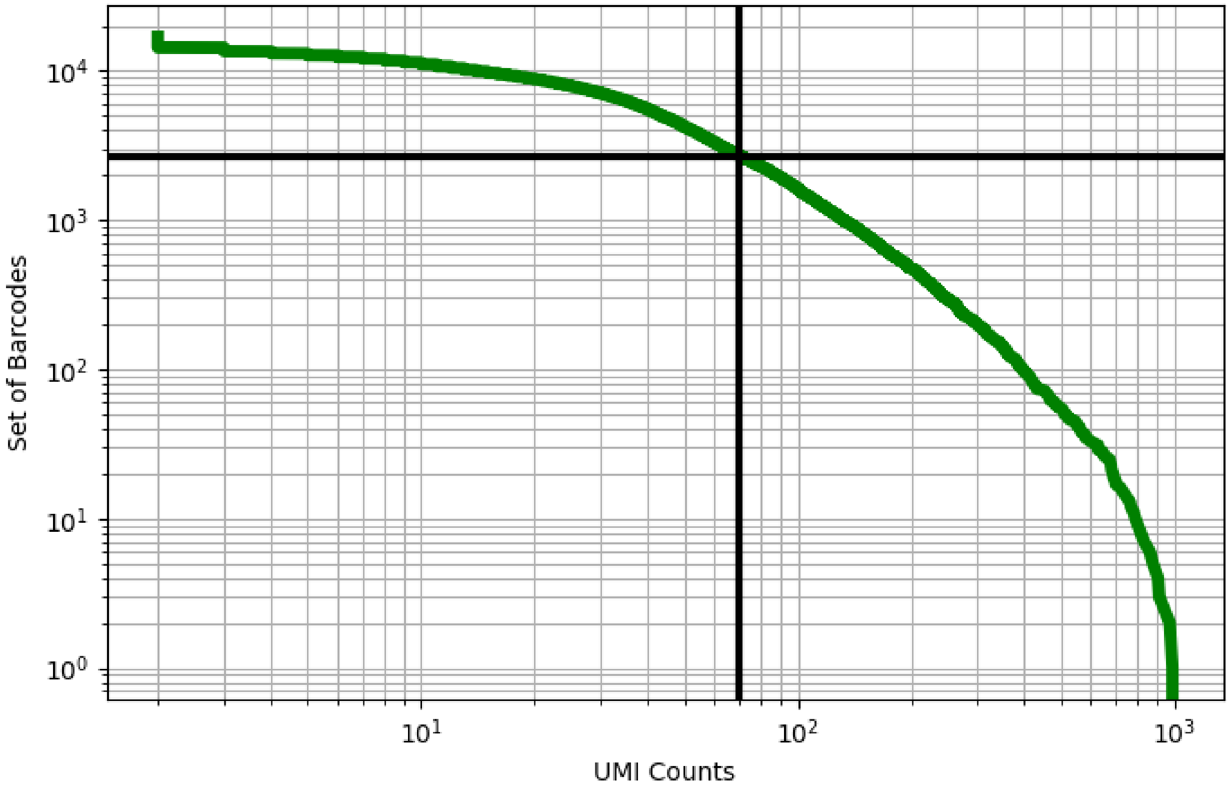


Figure 316: lung/GSM4142872

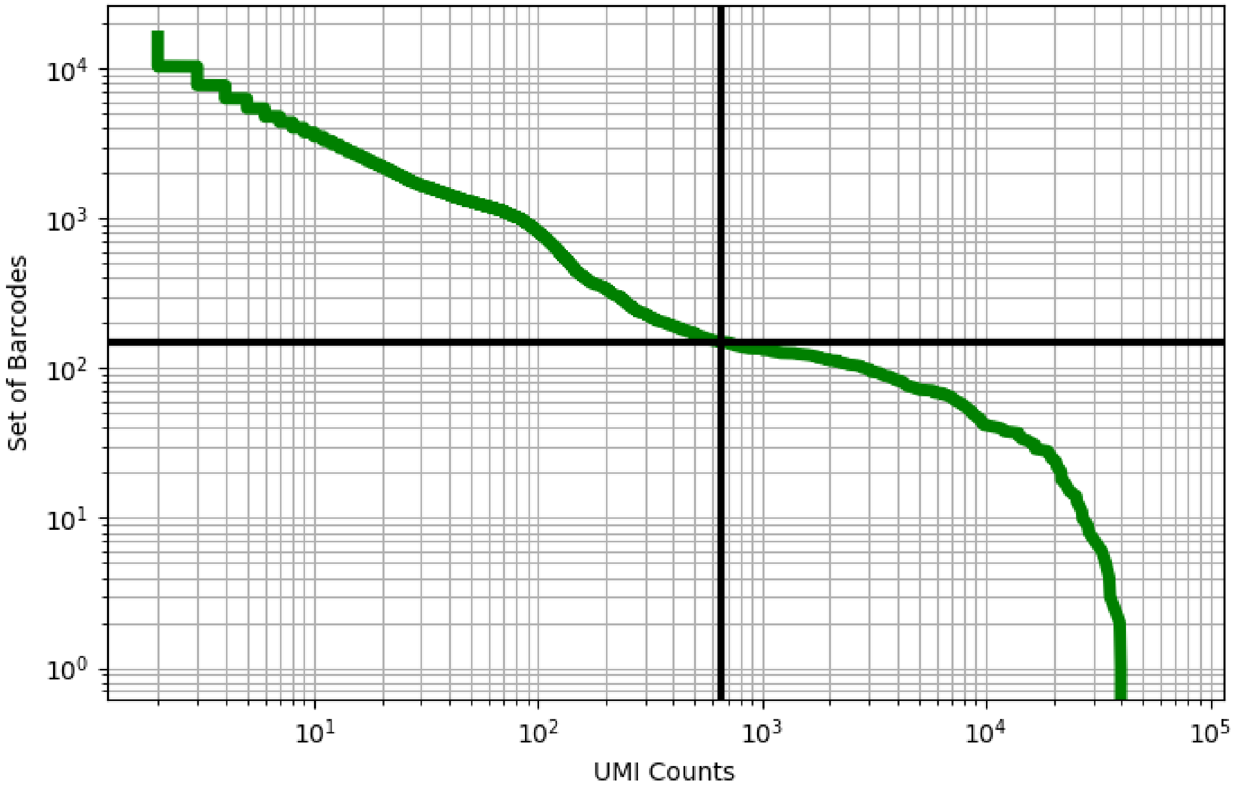


Figure 317: lung/GSM4142873

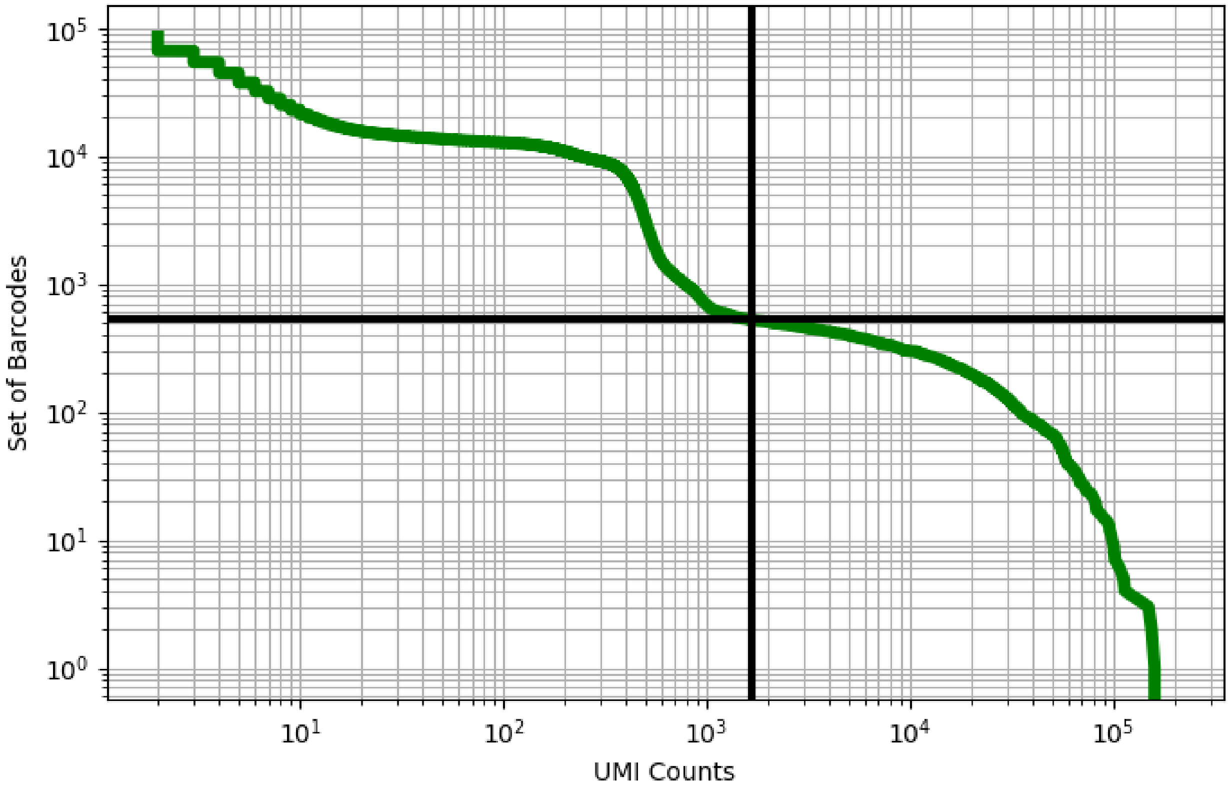


Figure 318: lung/GSM4213824

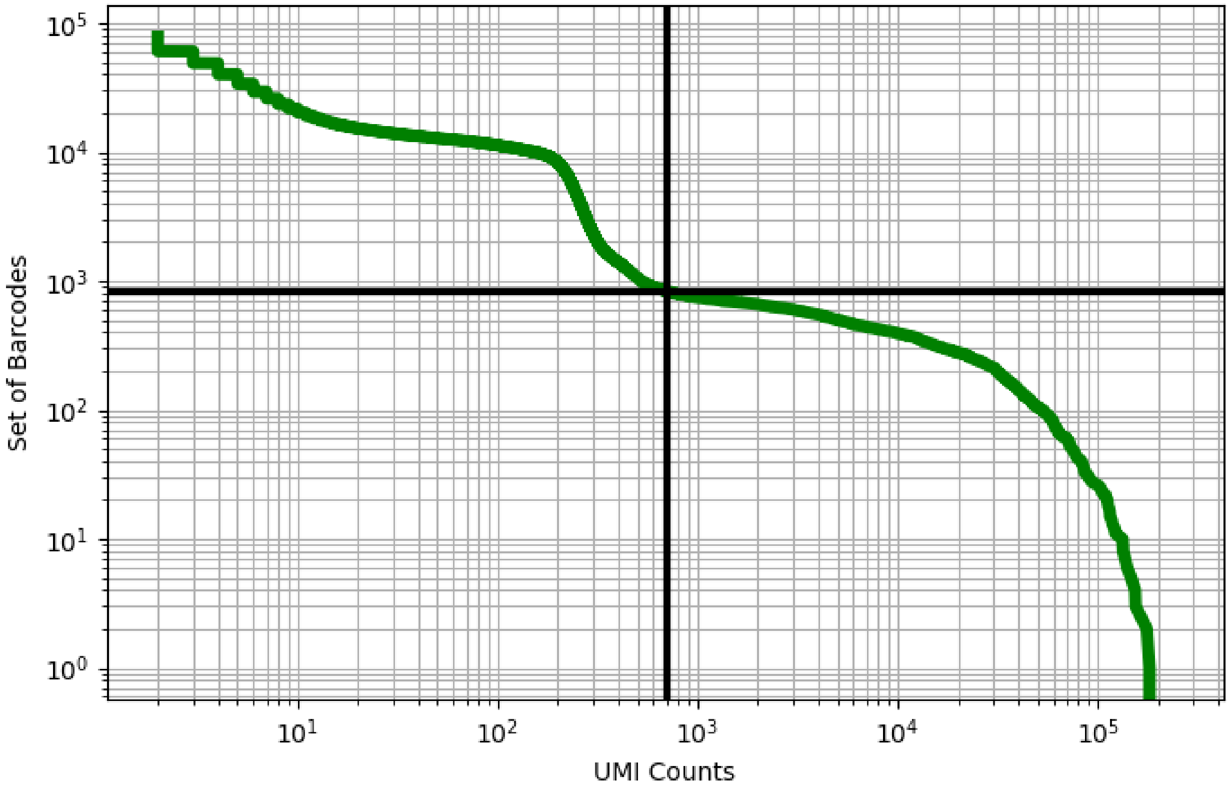


Figure 319: lung/GSM4213826

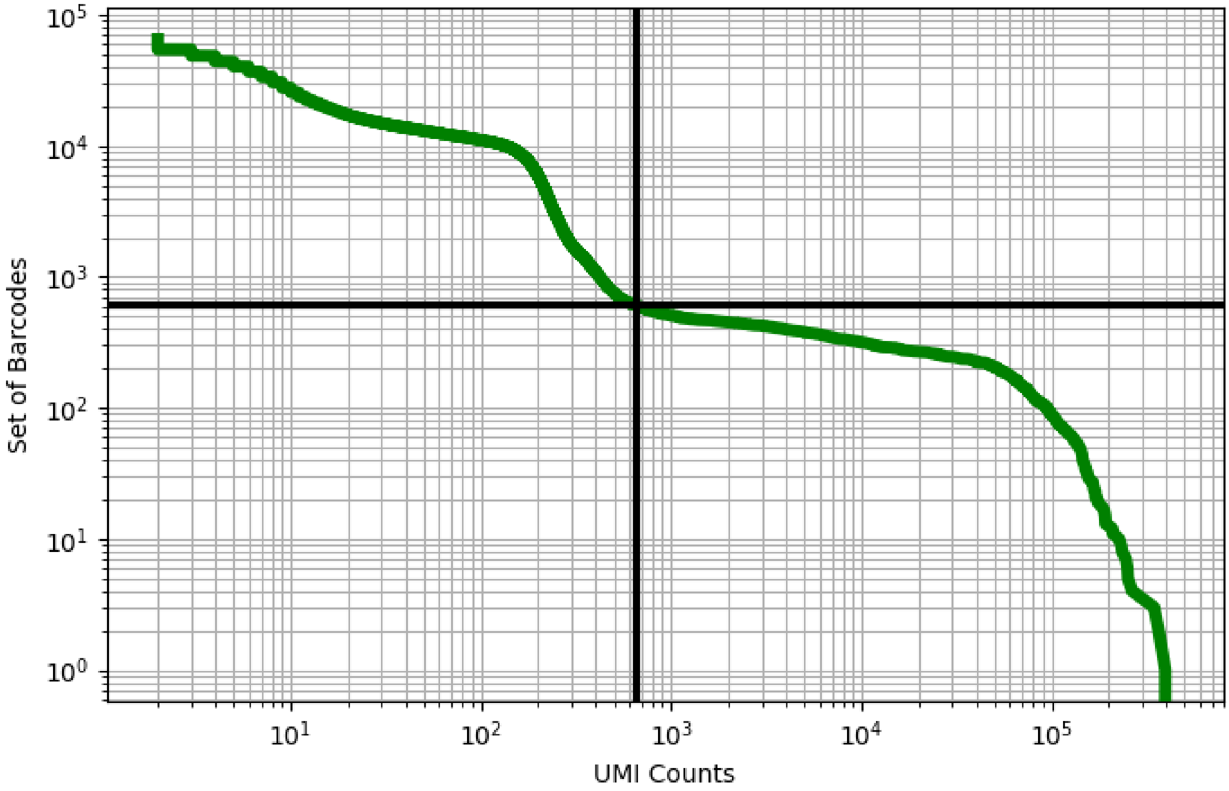


Figure 320: lung/GSM4213827

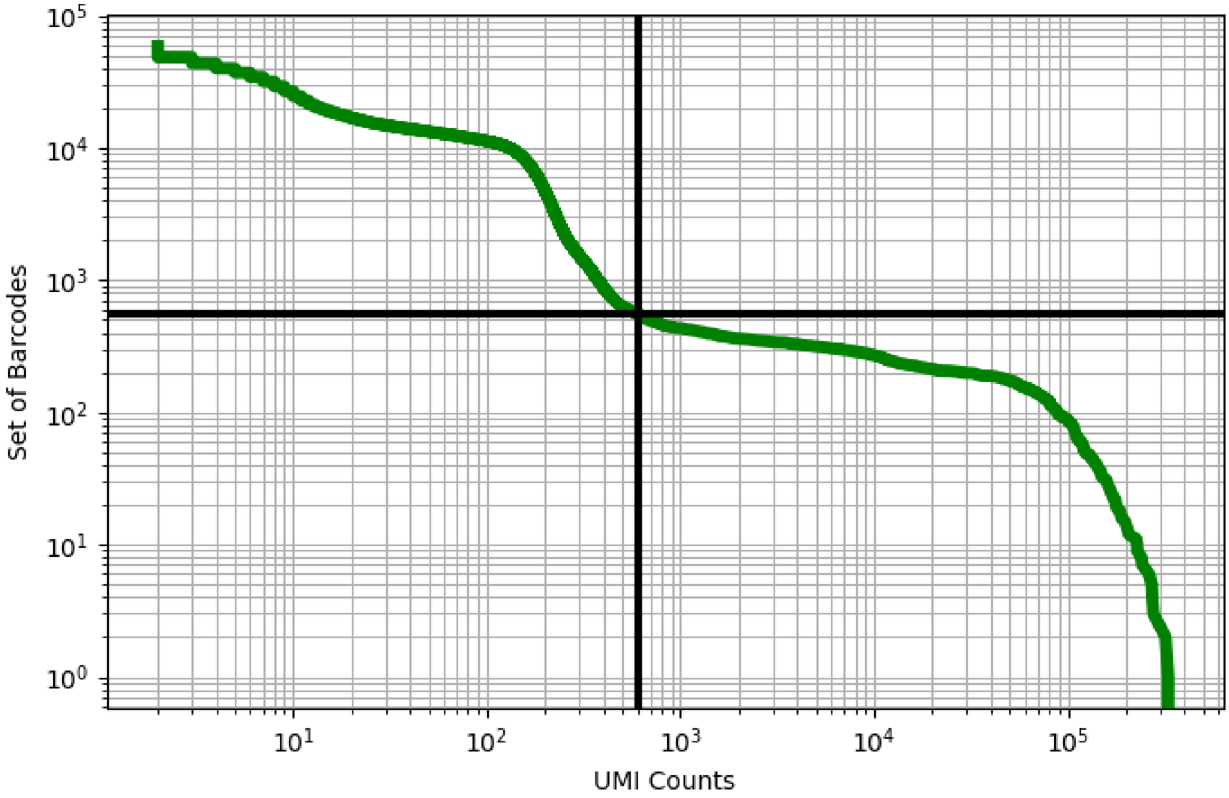


Figure 321: lung/GSM4213828

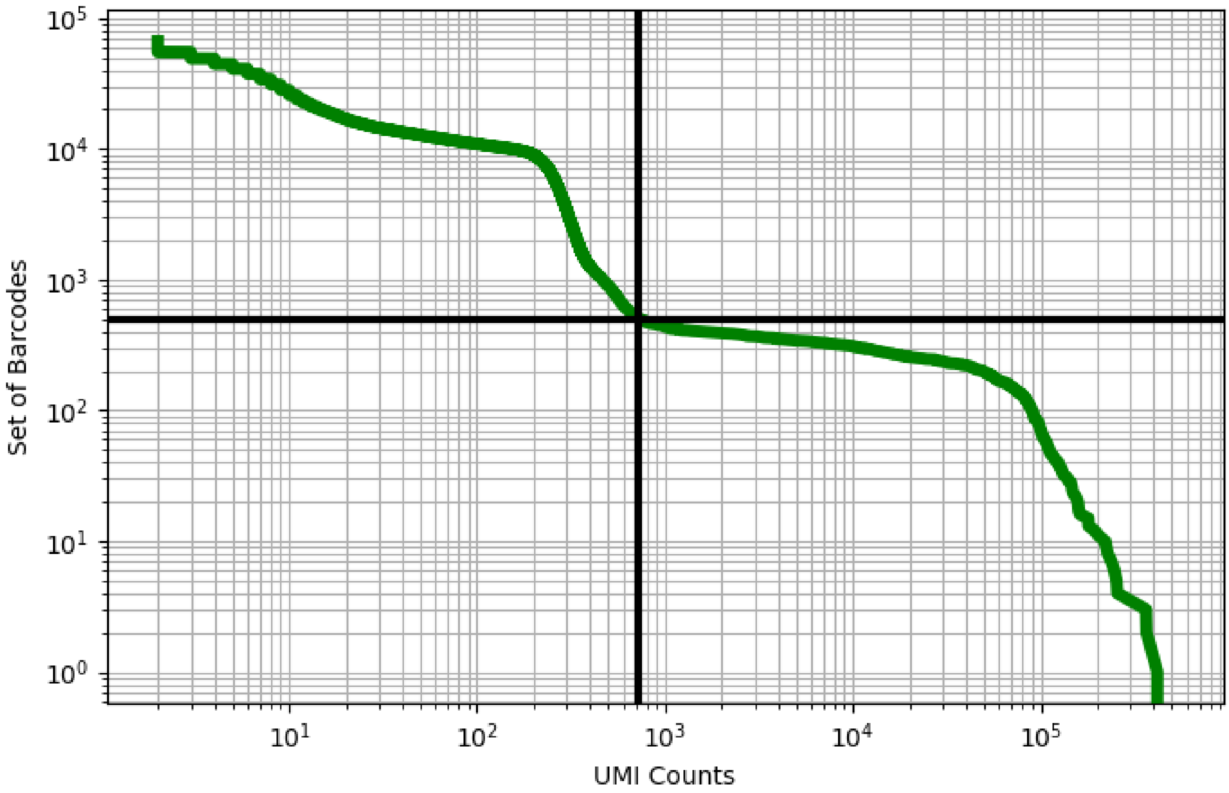


Figure 322: lung/GSM4213829

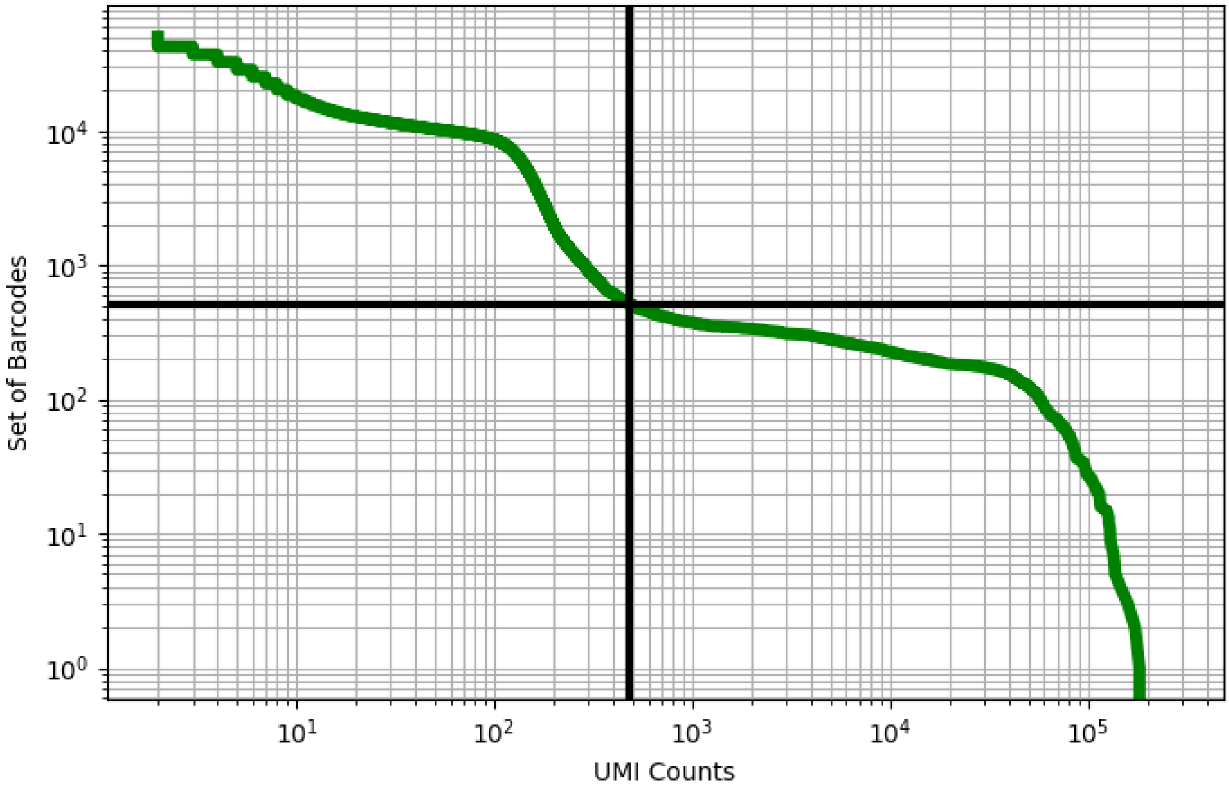


Figure 323: lung/GSM4213830

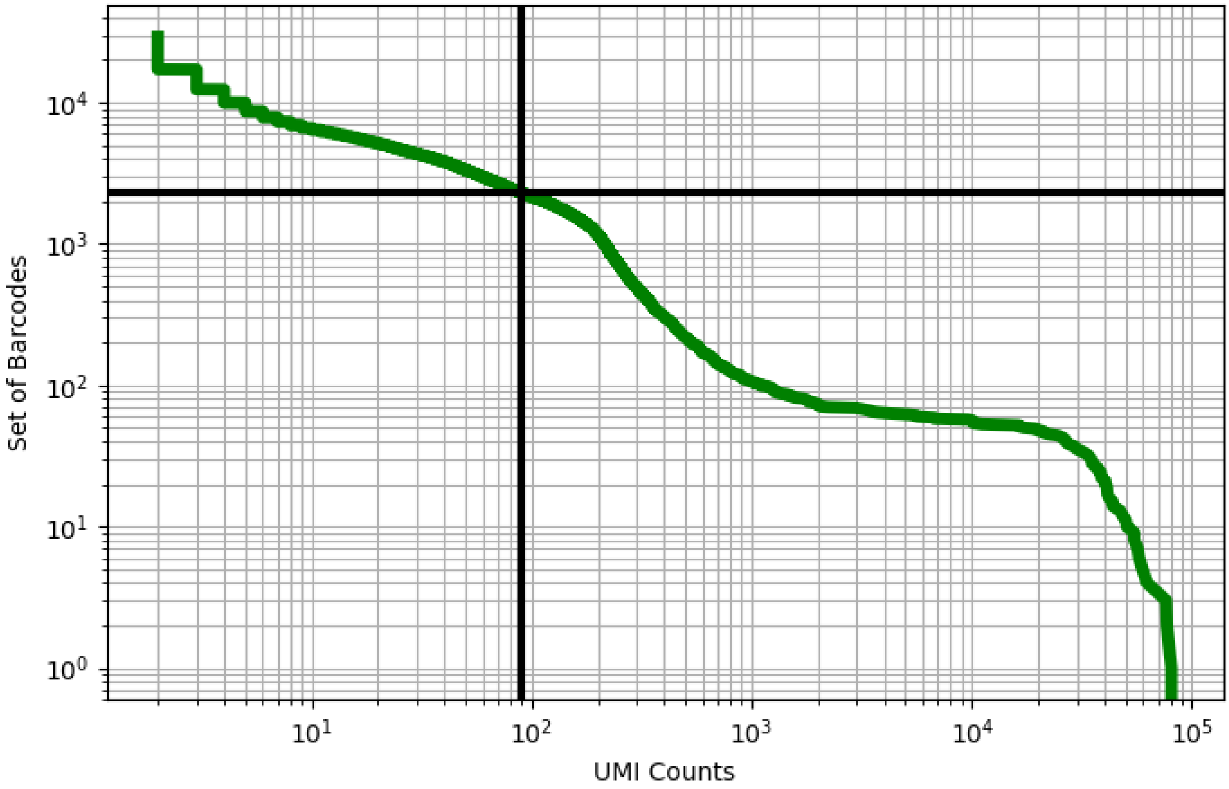


Figure 324: lung/GSM4213832

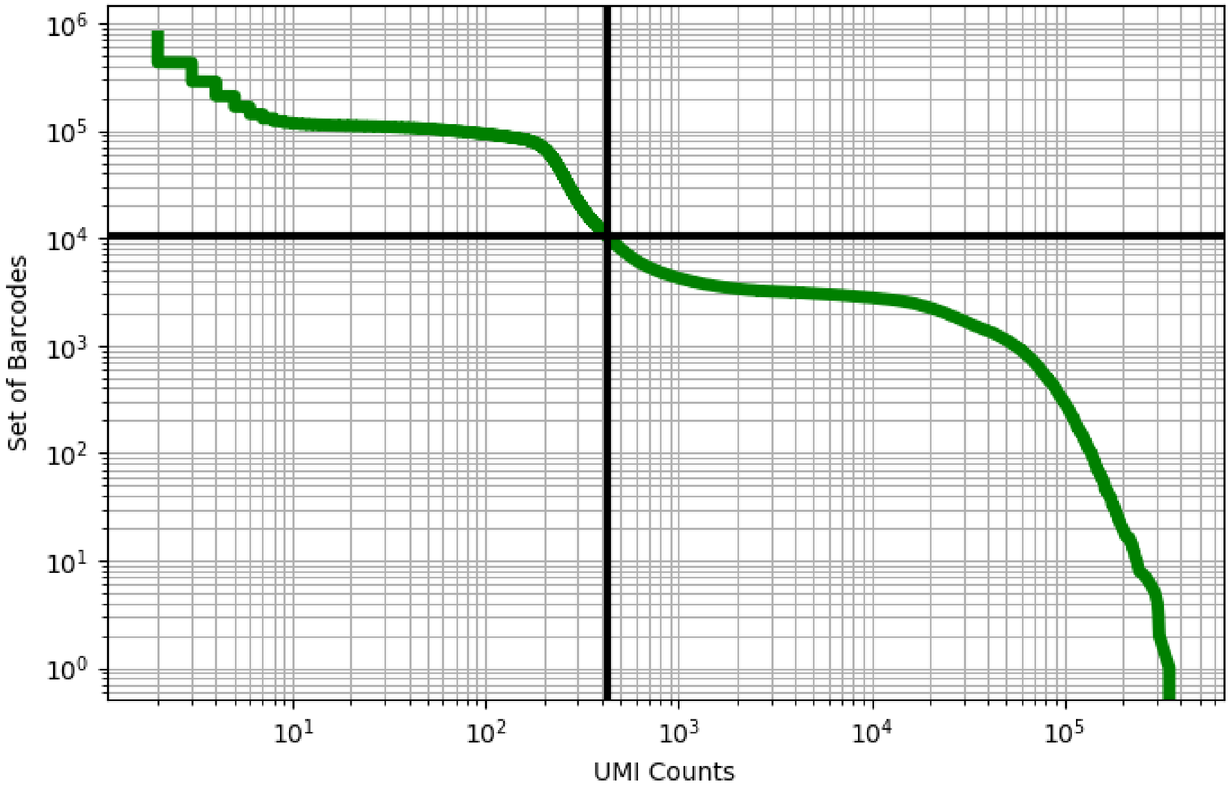


Figure 325: lung/GSM4213834

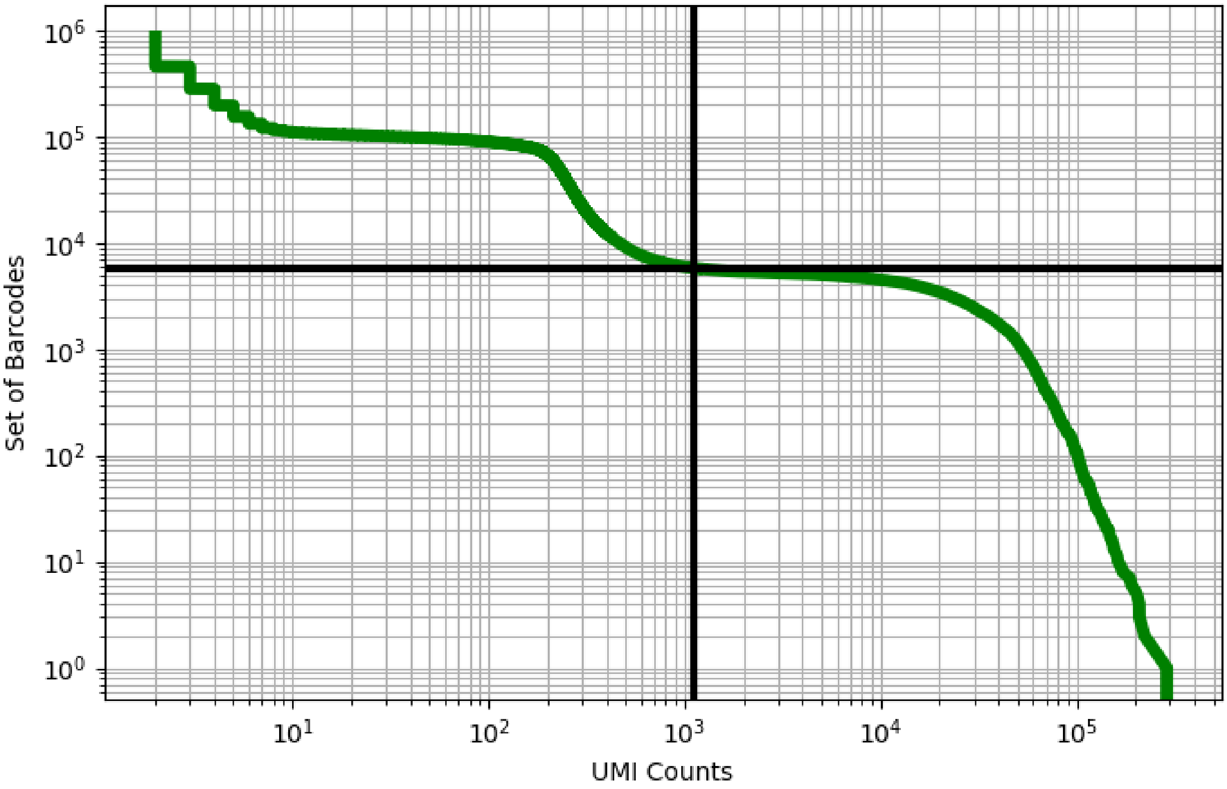


Figure 326: lung/GSM4213835

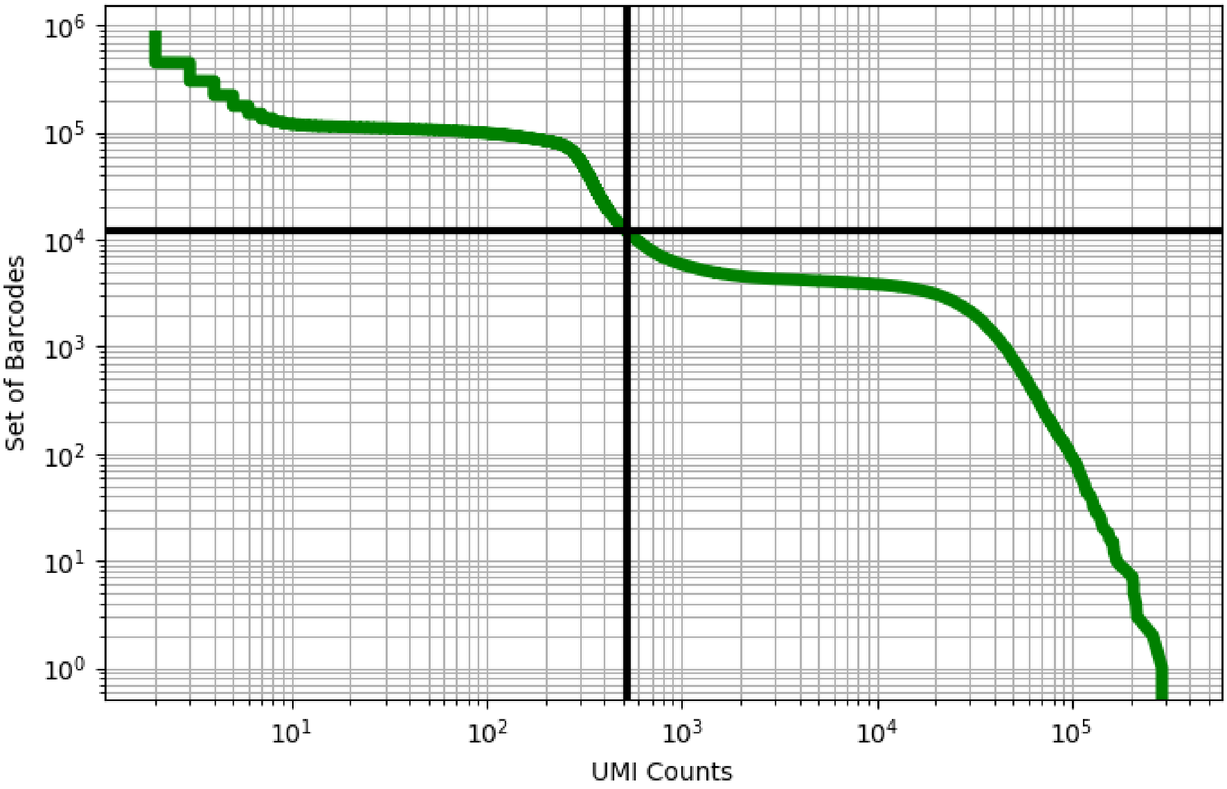


Figure 327: lung/GSM4213836

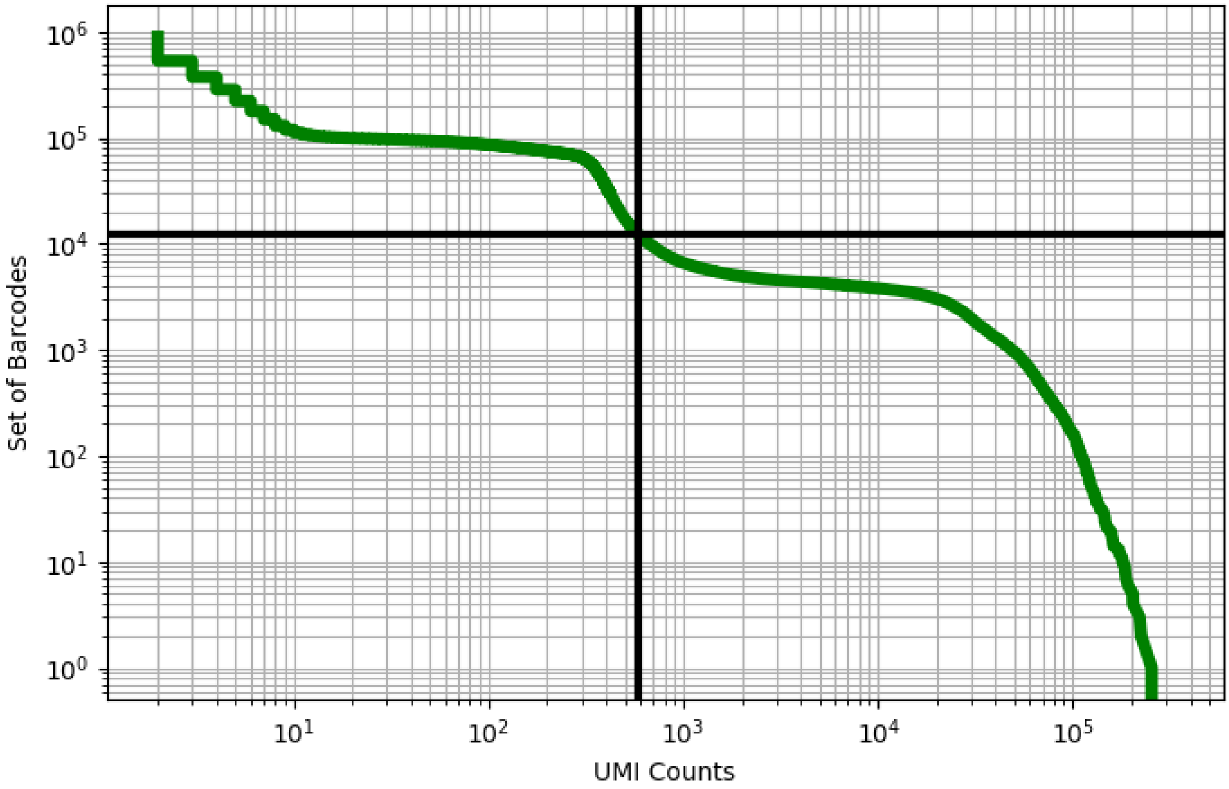


Figure 328: lung/GSM4213837

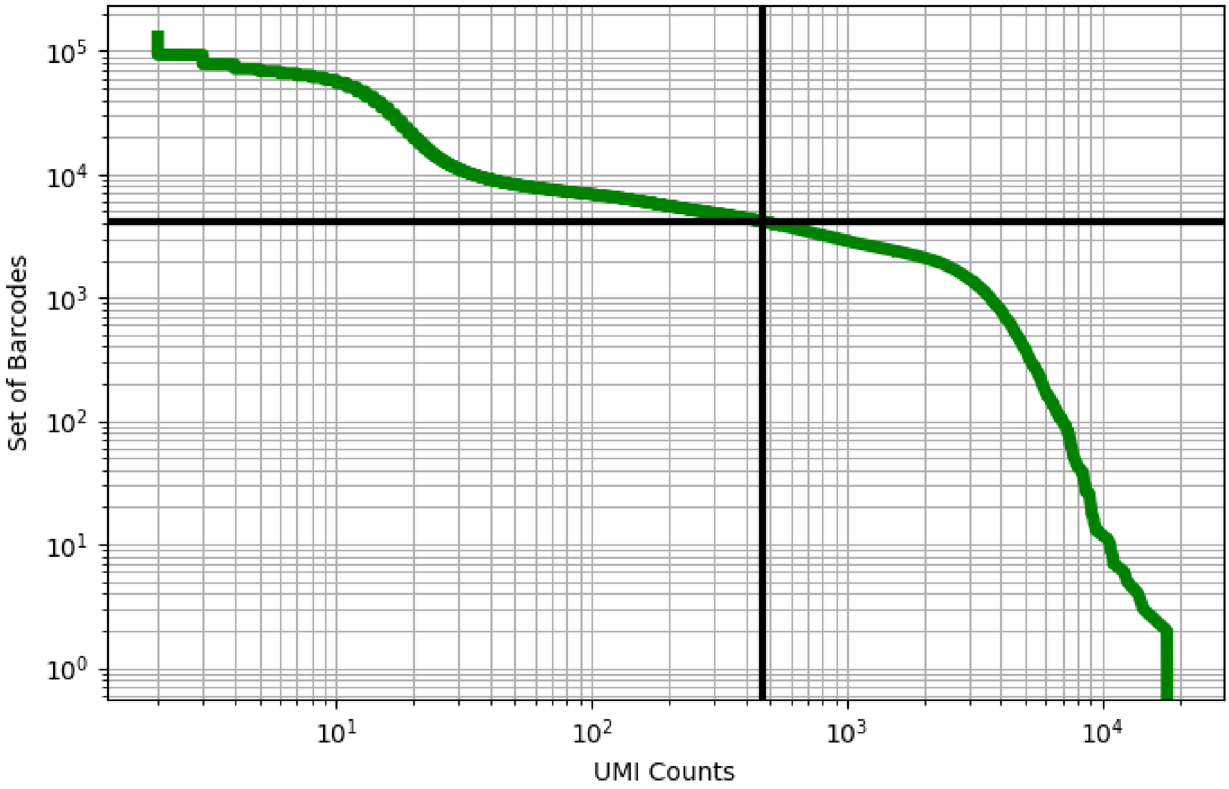


Figure 329: lung/GSM4339769

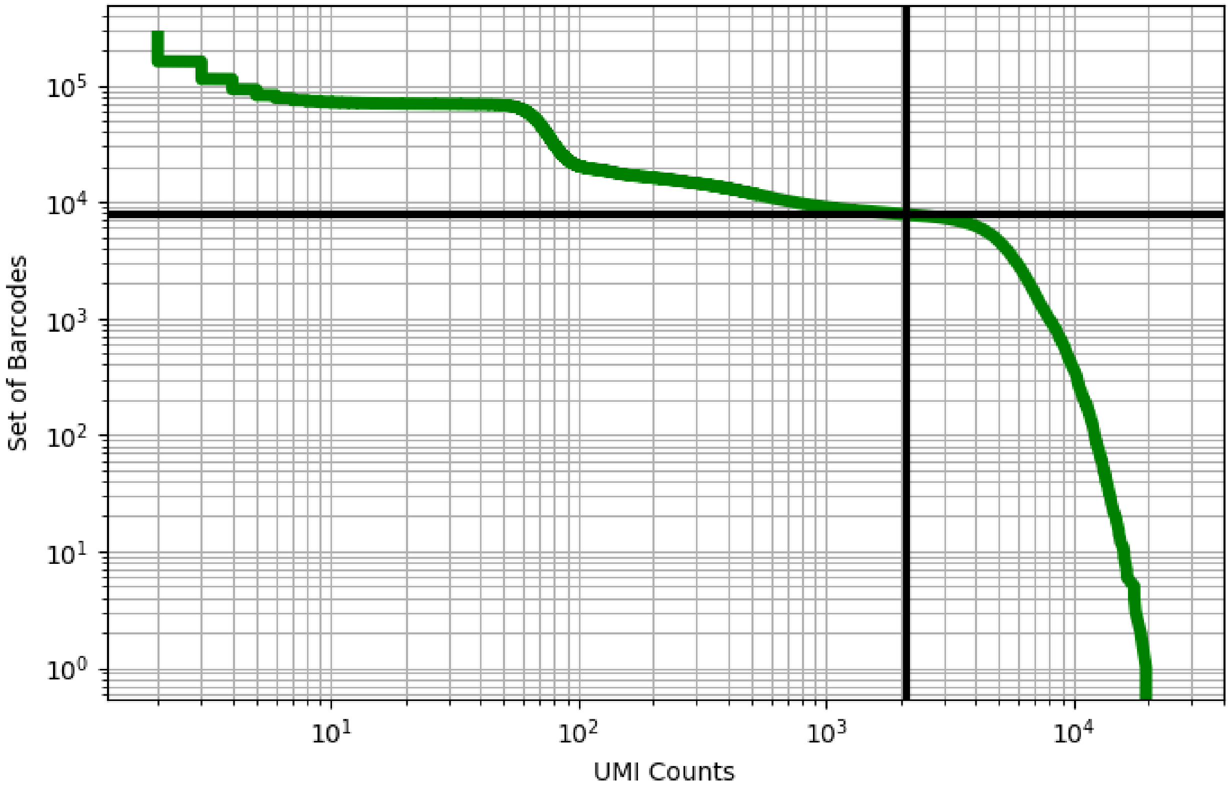


Figure 330: lung/GSM4475048

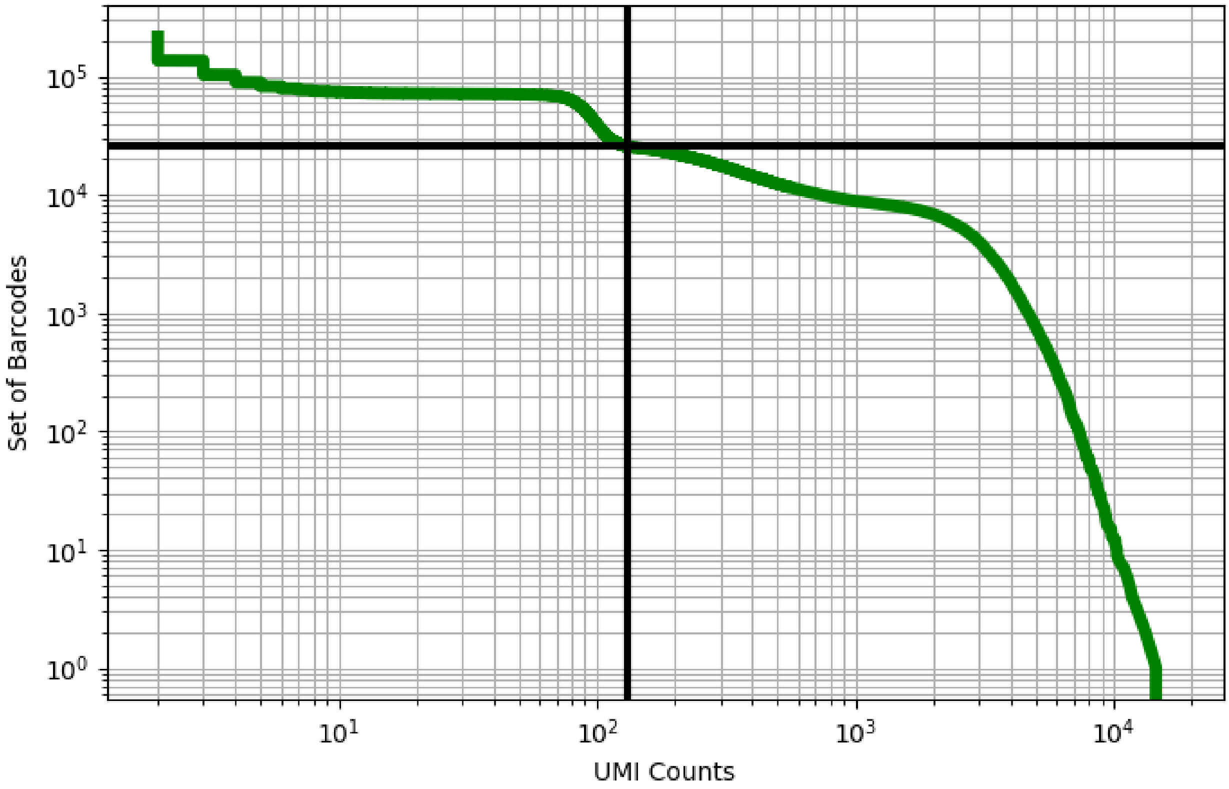


Figure 331: lung/GSM4475049

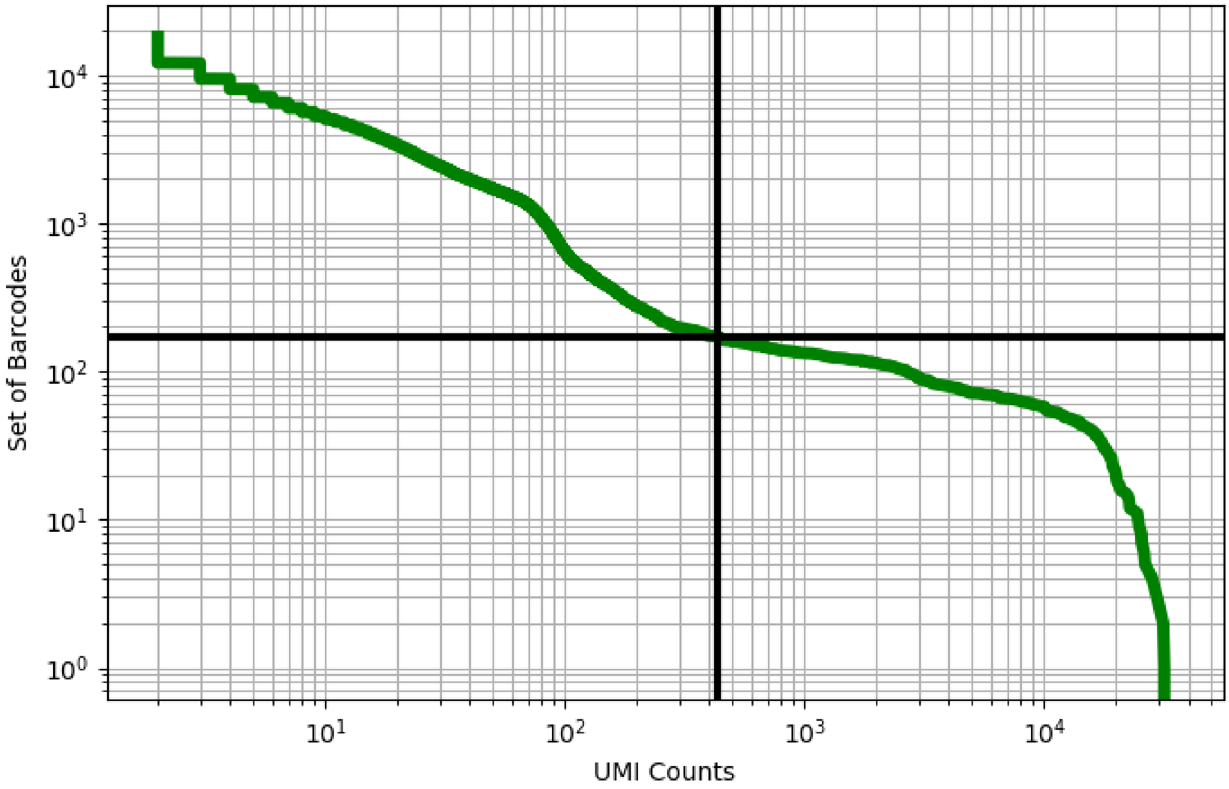


Figure 332: lung/GSM4475050

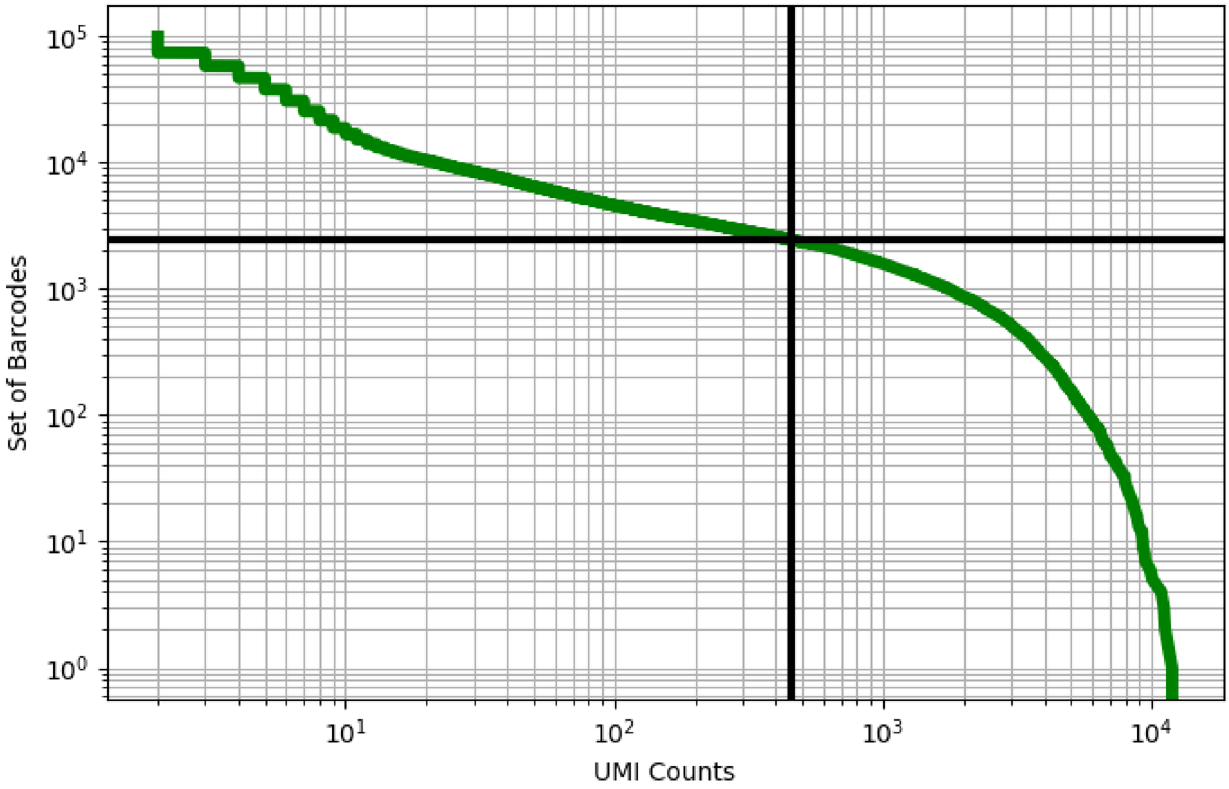


Figure 333: lung/GSM4475051

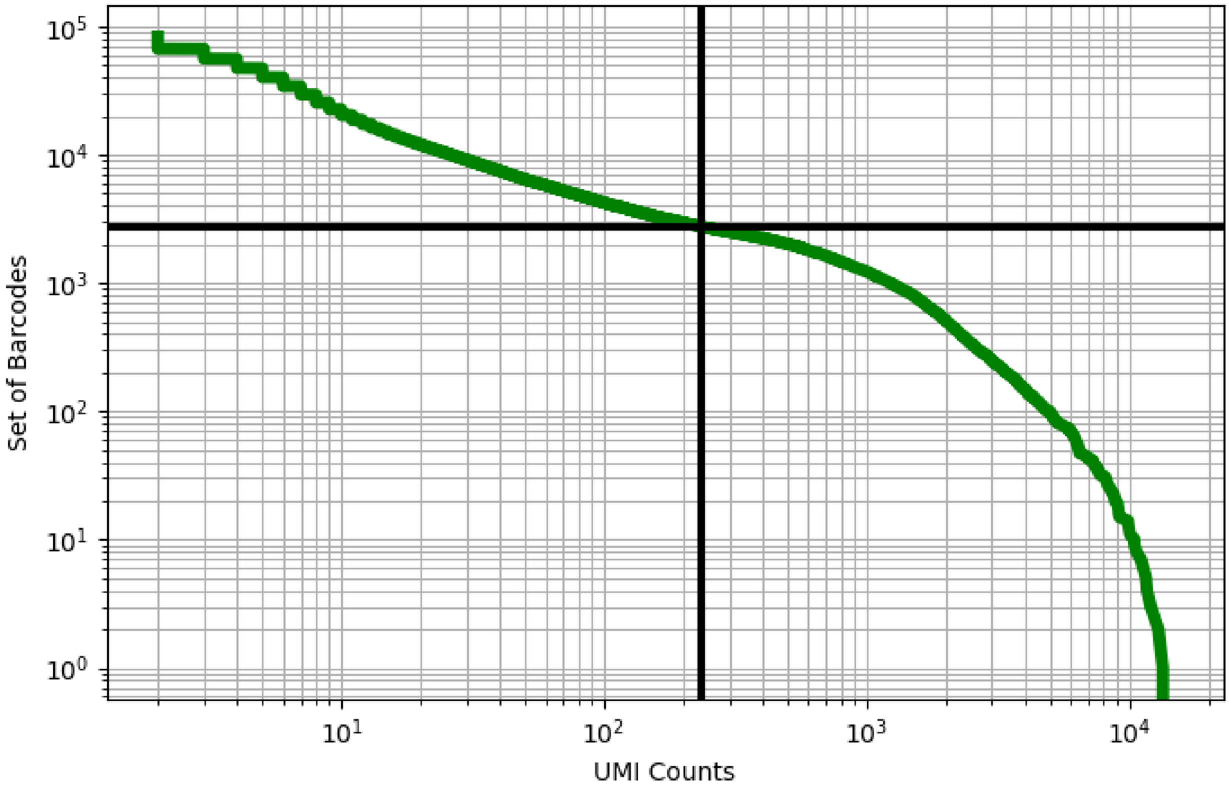


Figure 334: lung/GSM4475052

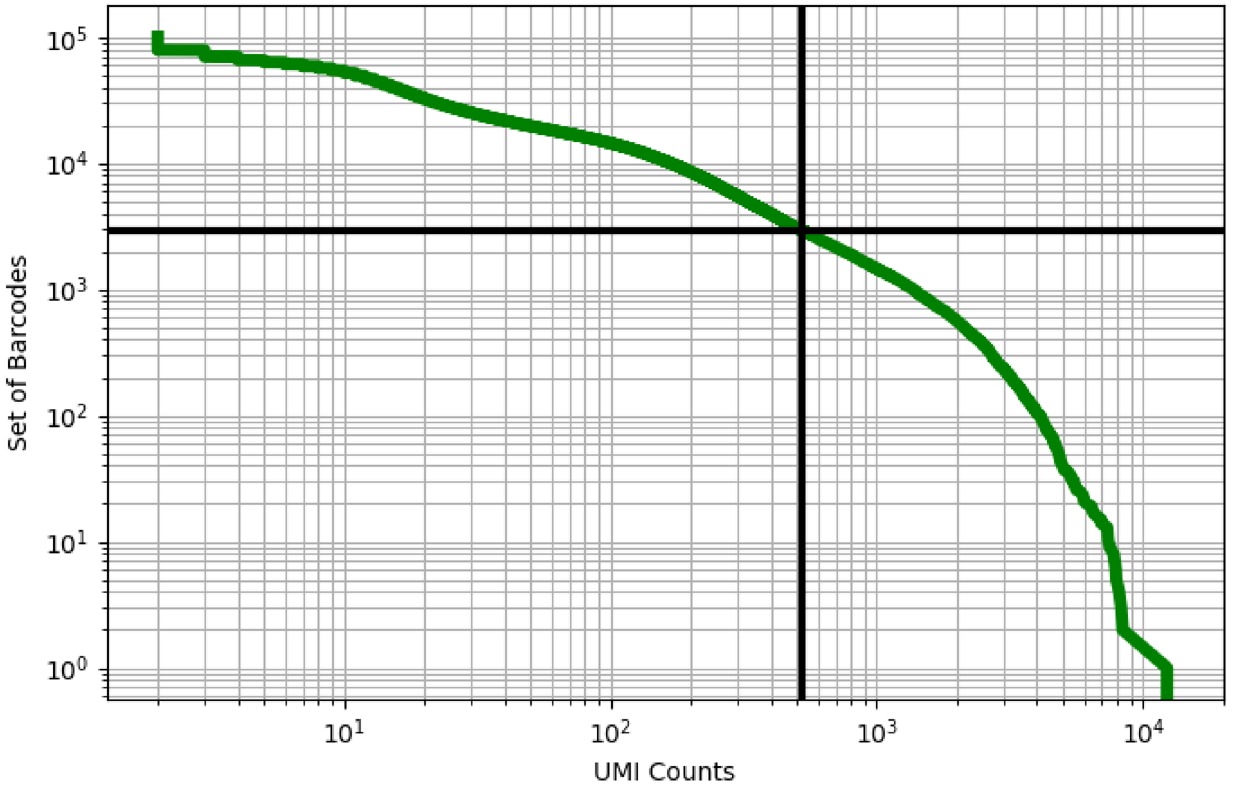


Figure 335: lung/GSM4475053

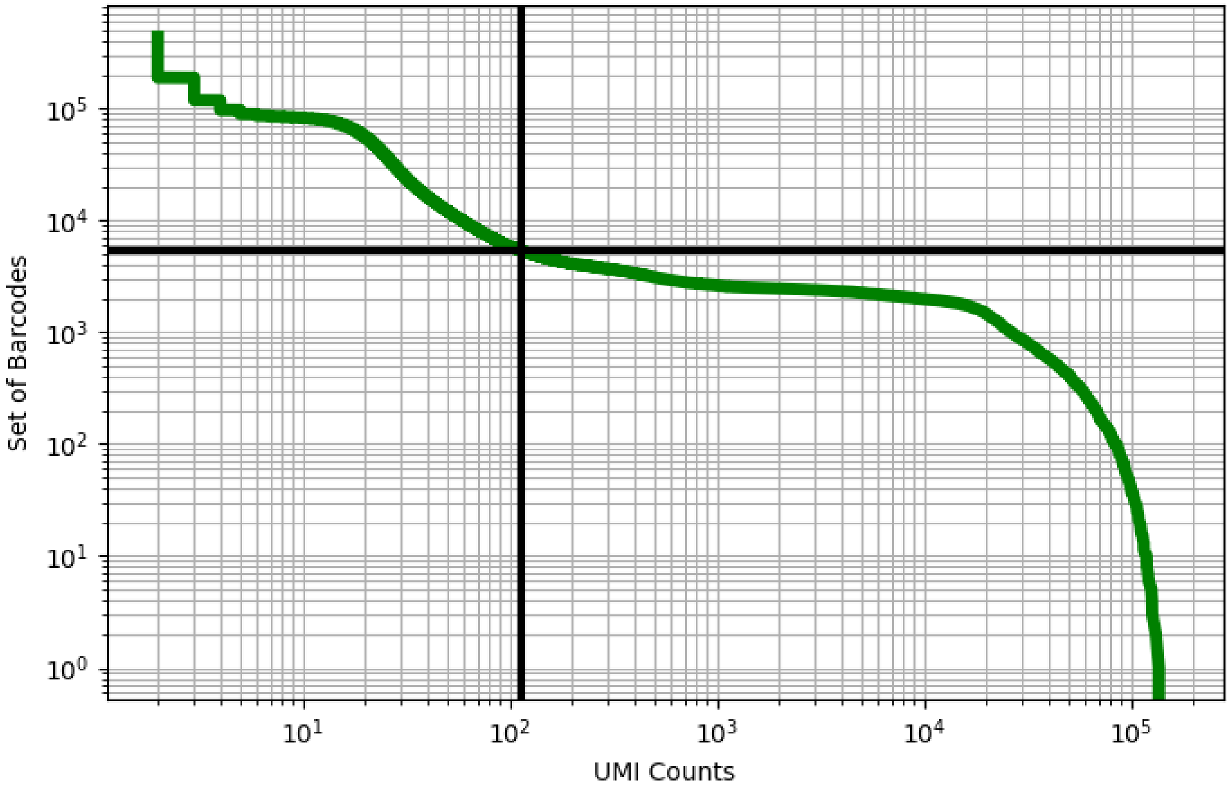


Figure 336: lung/GSM4769387

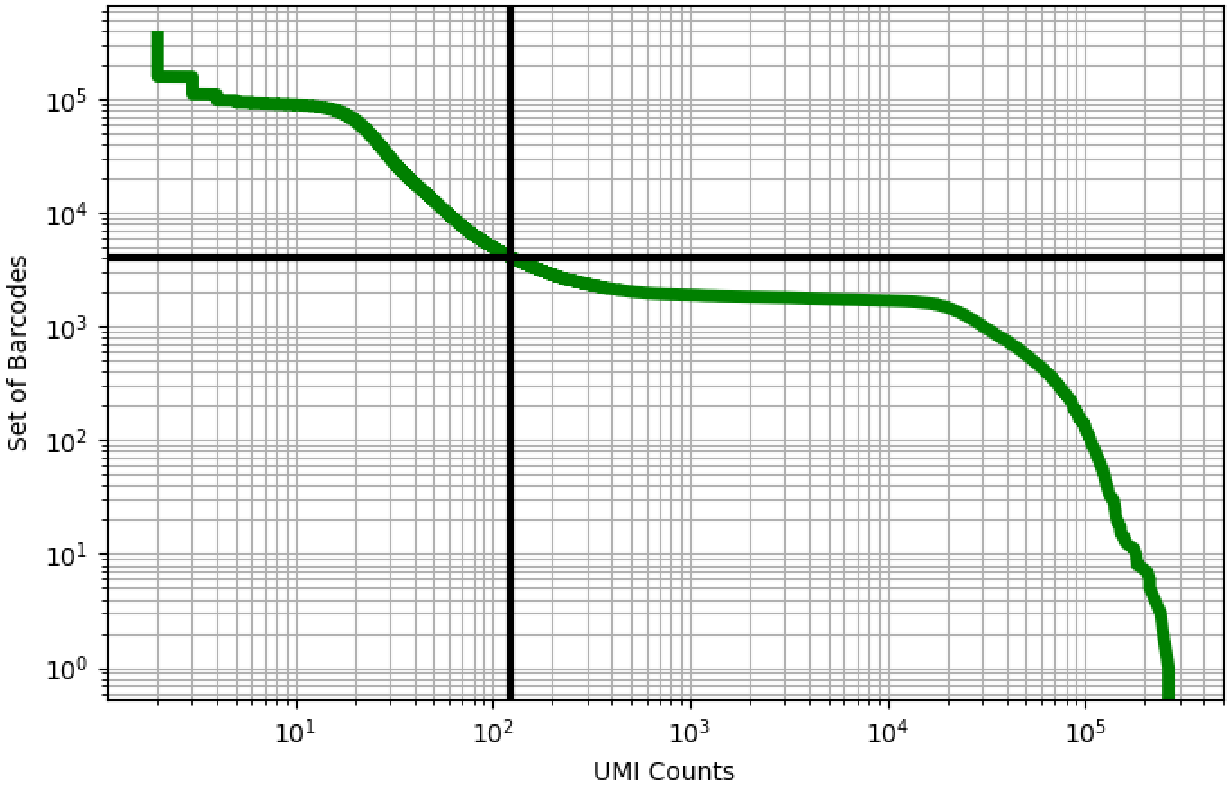


Figure 337: lung/GSM4769388

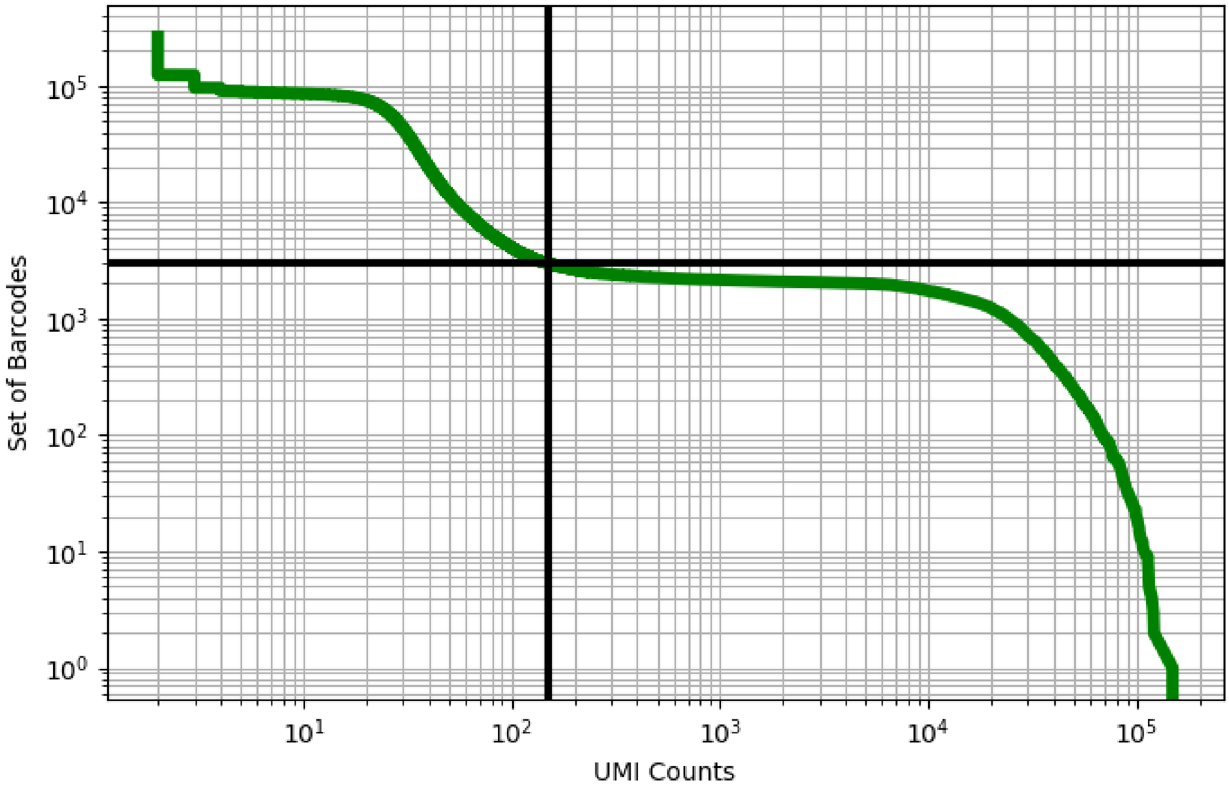


Figure 338: lung/GSM4769389

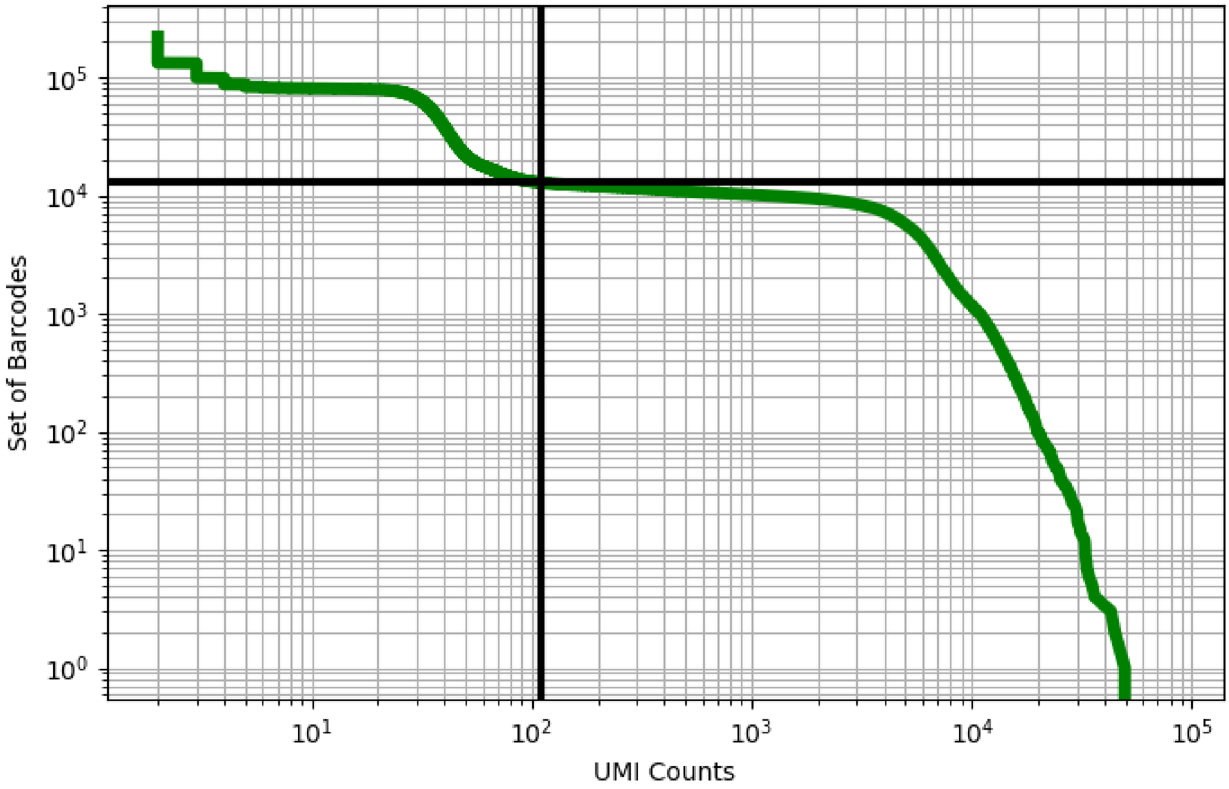


Figure 339: lymph_node/GSM3535276

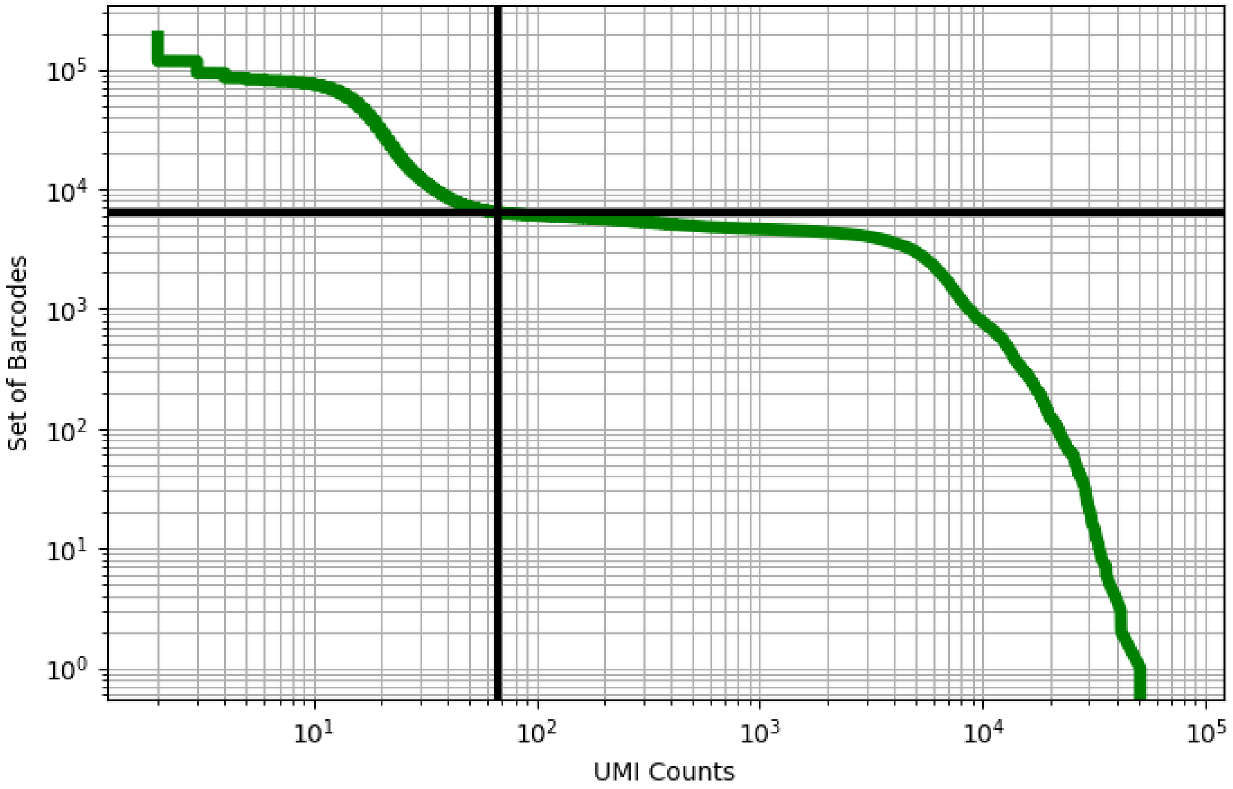


Figure 340: lymph_node/GSM3535277

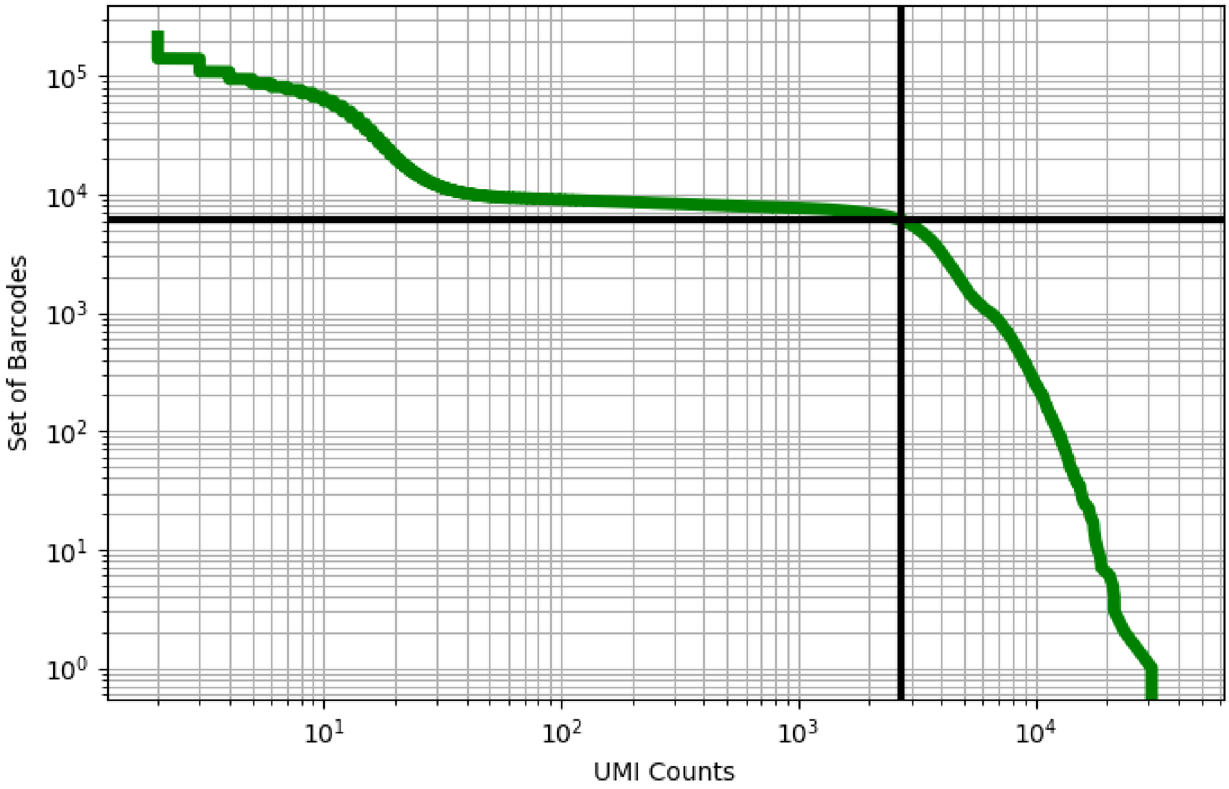


Figure 341: lymph_node/GSM3535278

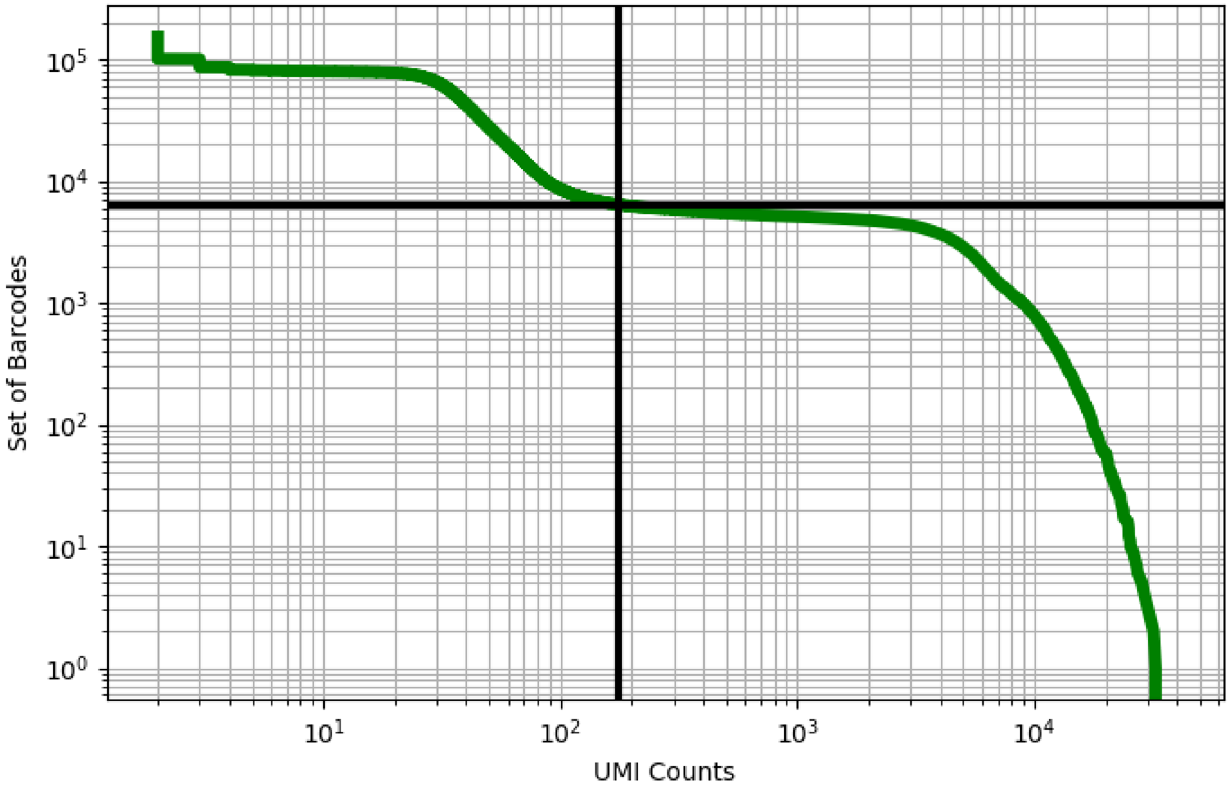


Figure 342: lymph_node/GSM3535279

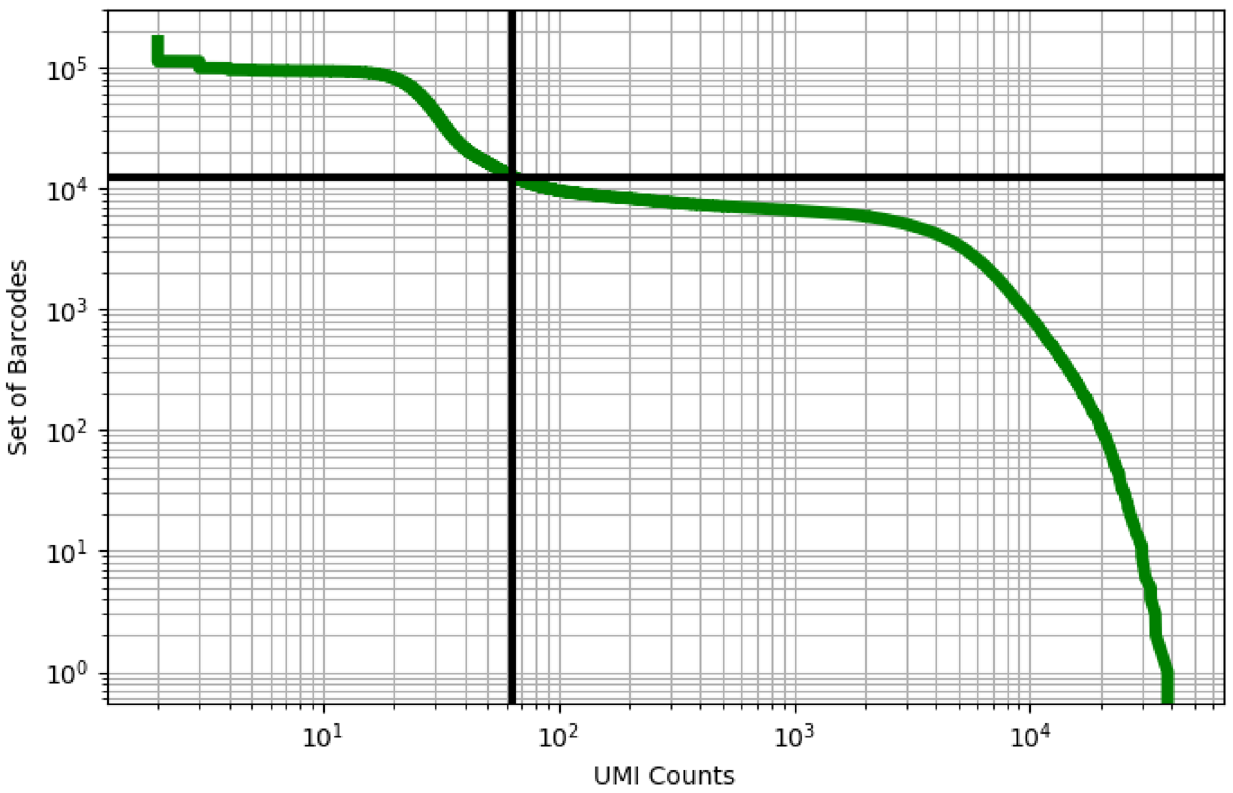


Figure 343: lymph_node/GSM3535280

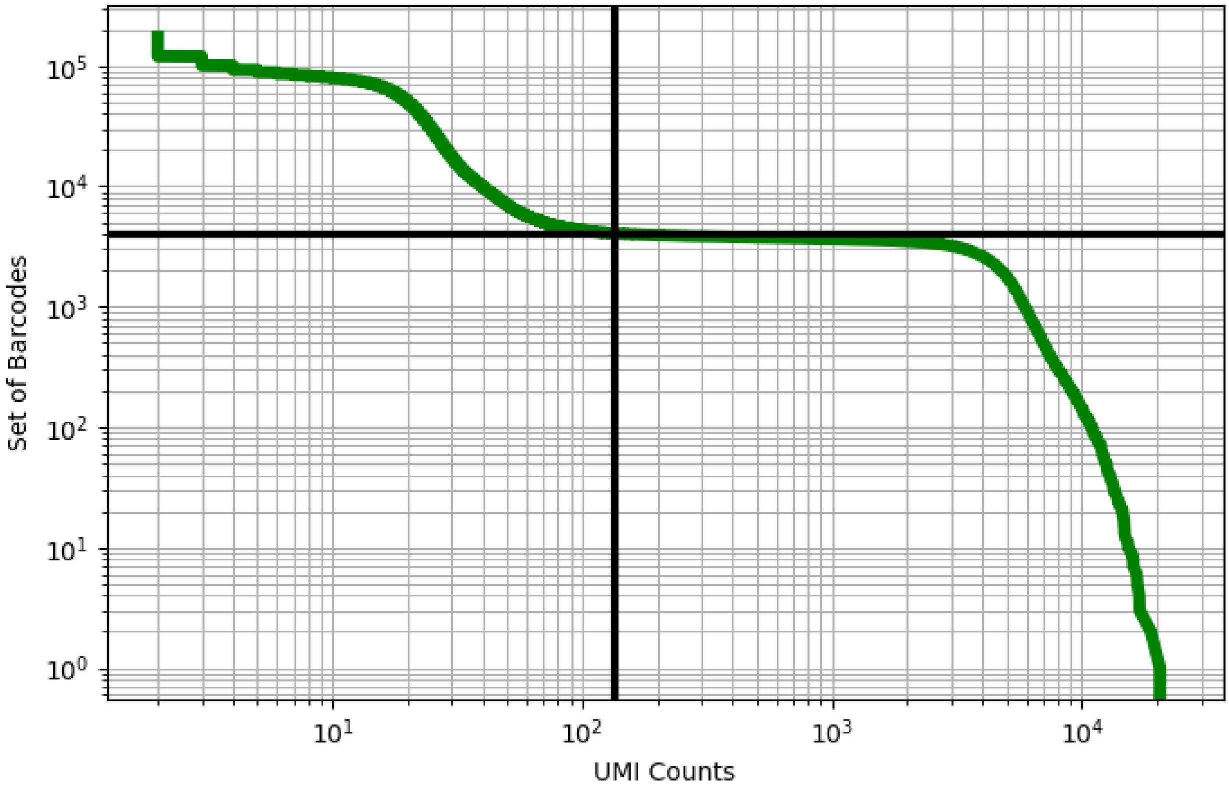


Figure 344: lymph_node/GSM3535281

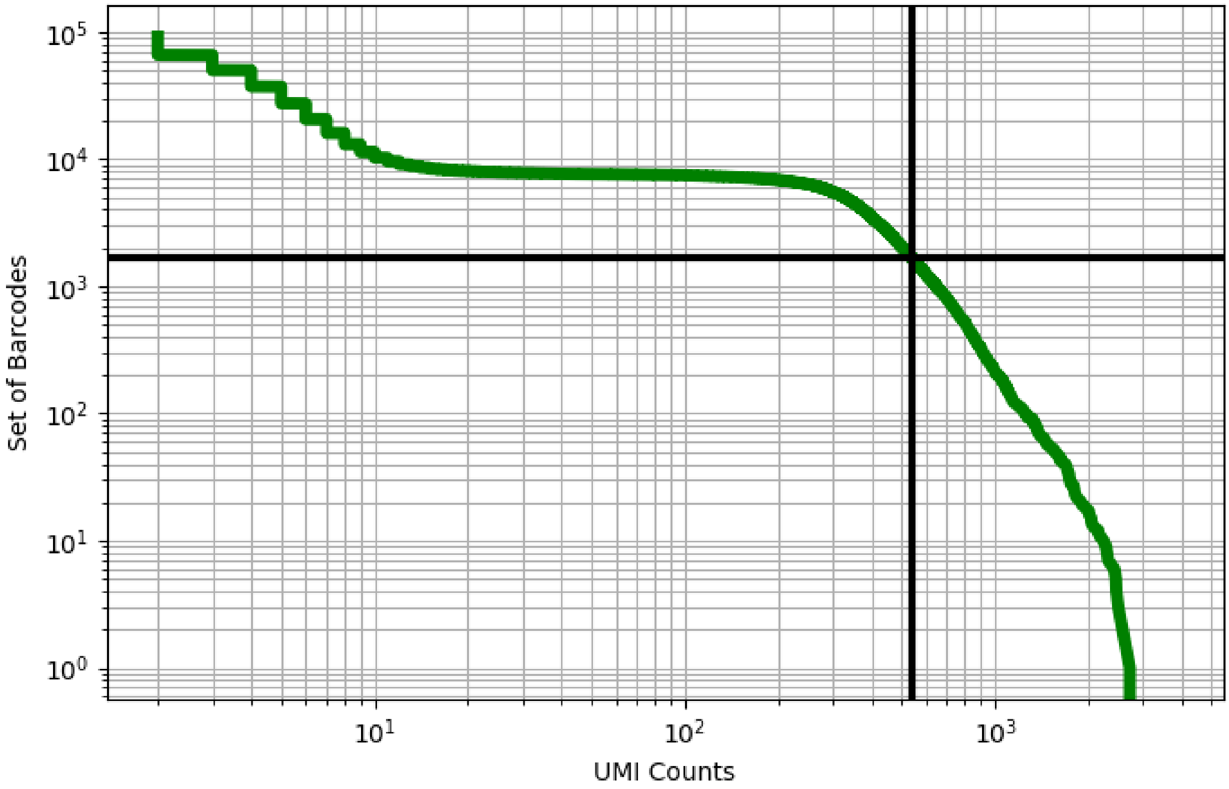


Figure 345: mammary/GSM3148575

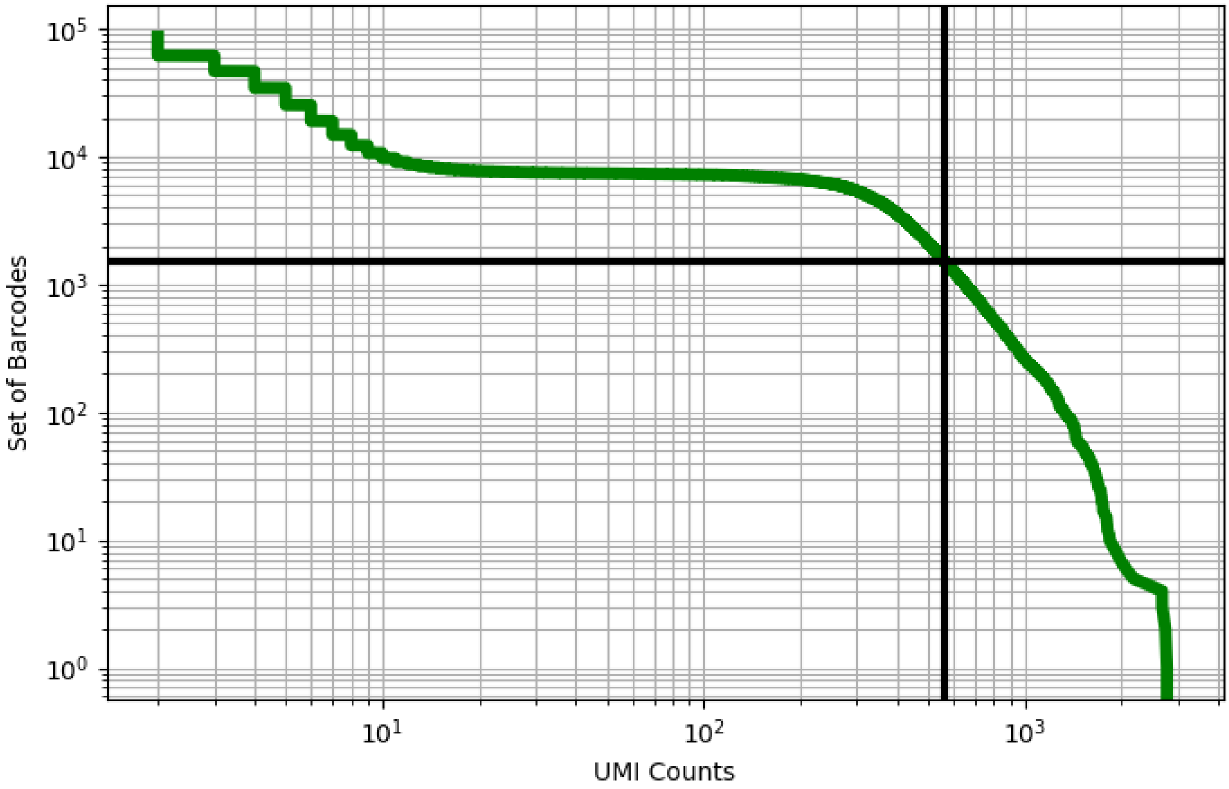


Figure 346: mammary/GSM3148576

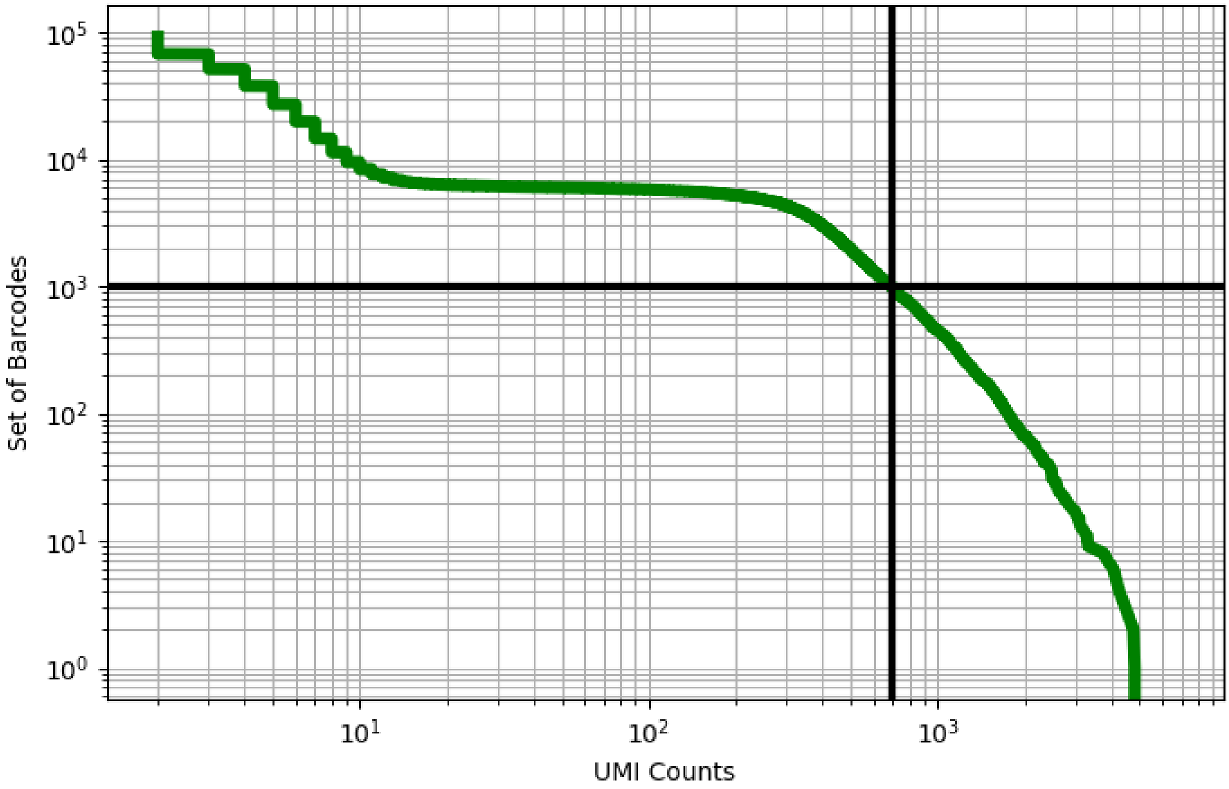


Figure 347: mammary/GSM3148577

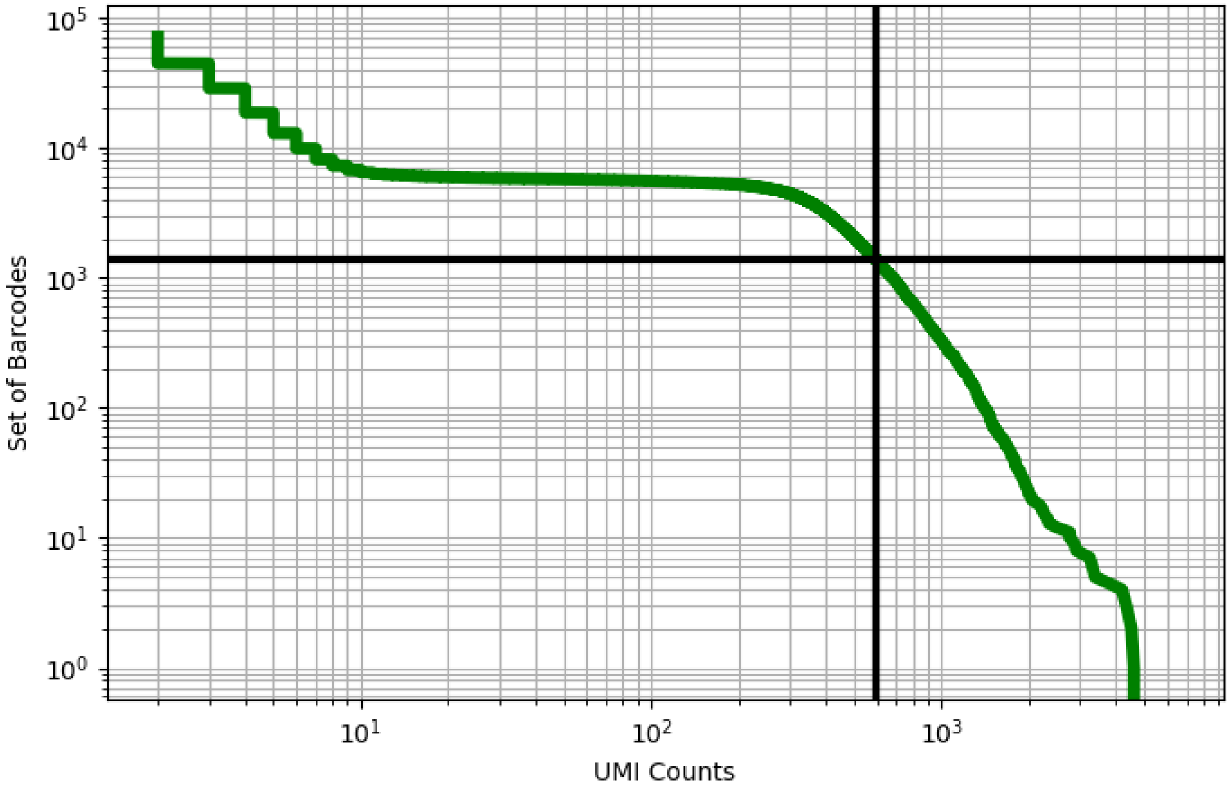


Figure 348: mammary/GSM3148578

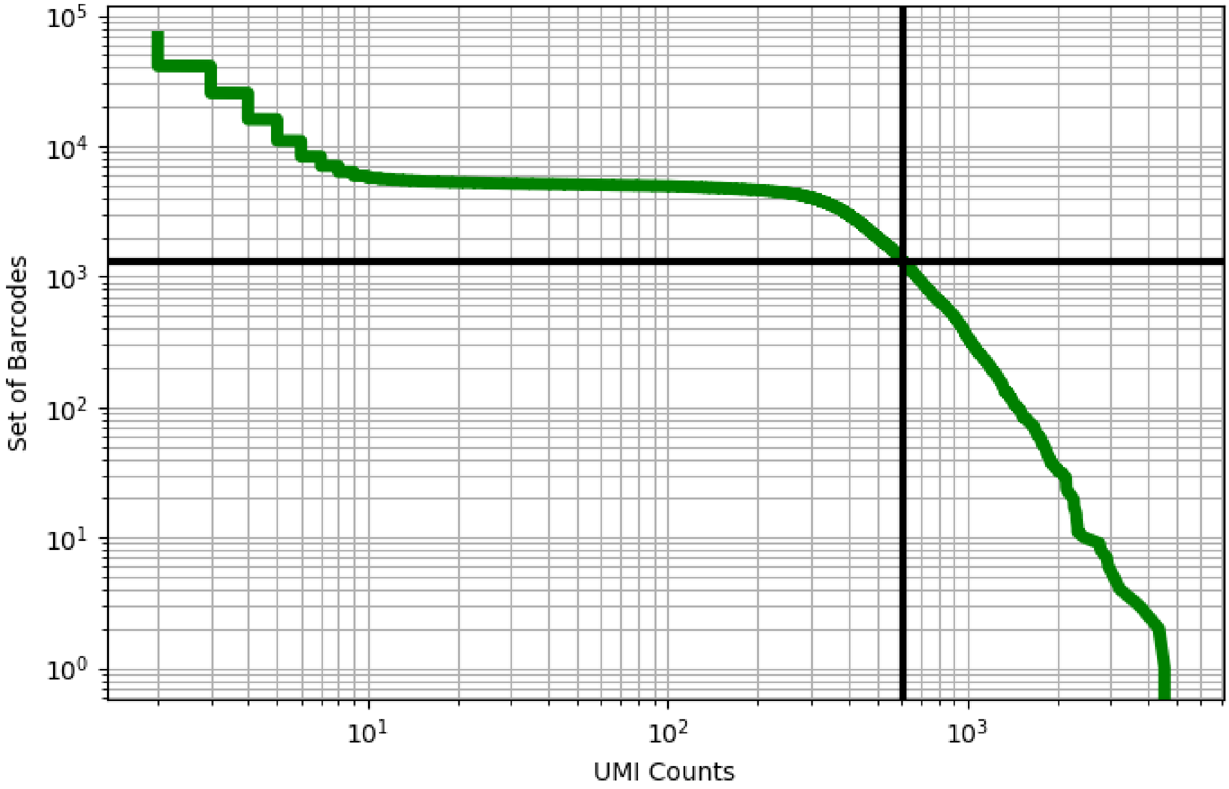


Figure 349: mammary/GSM3148579

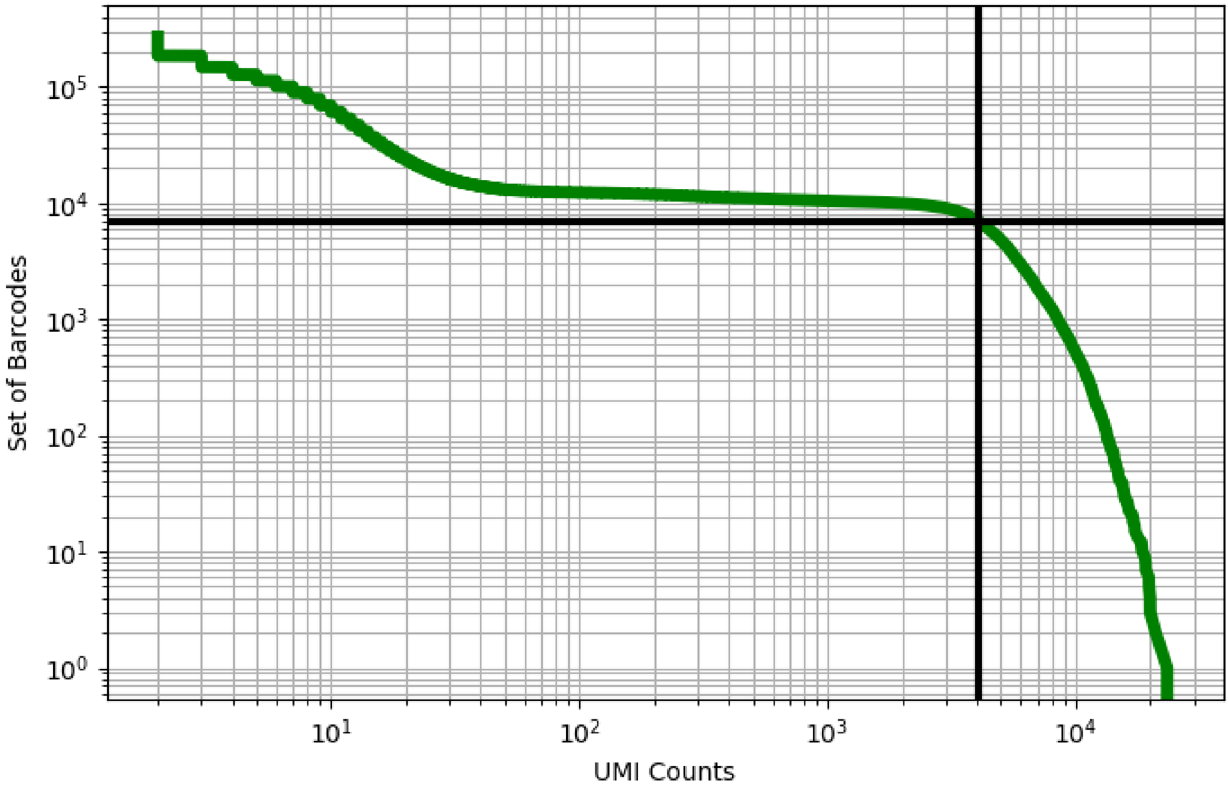


Figure 350: mammary/GSM3516947

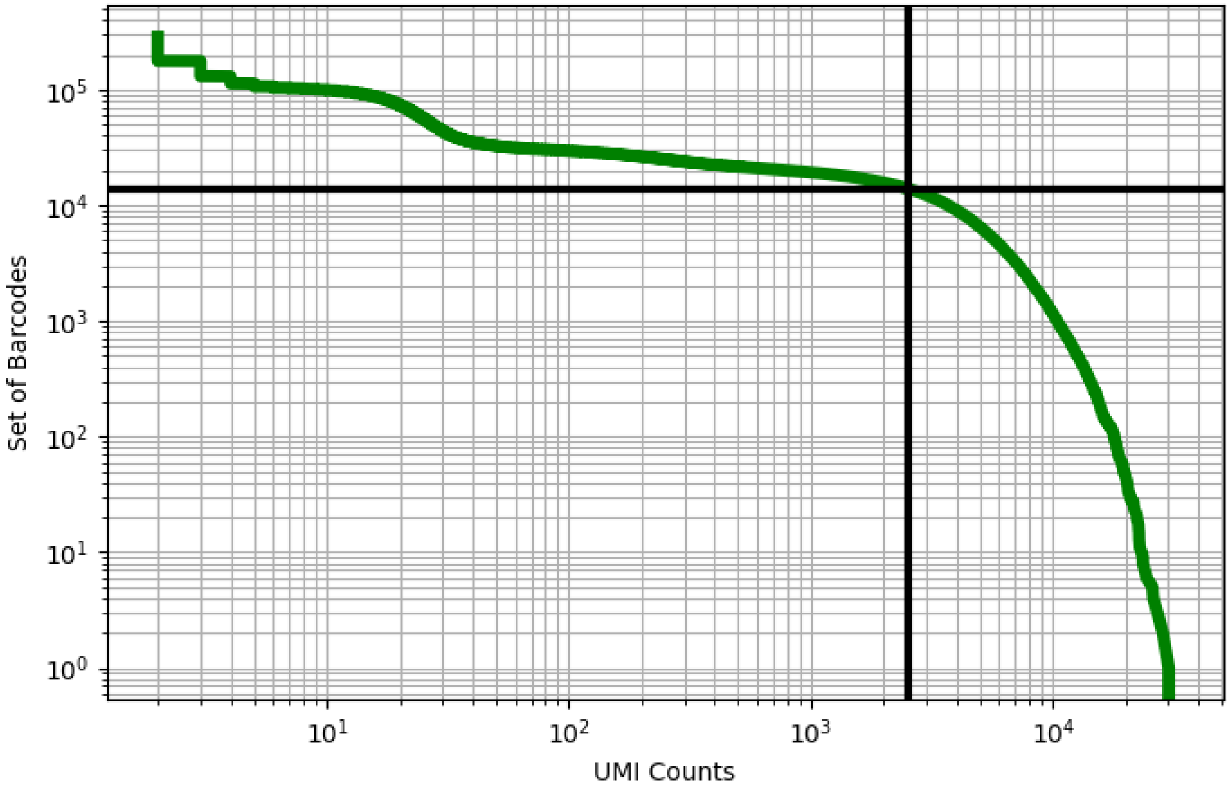


Figure 351: mammary/GSM3516948

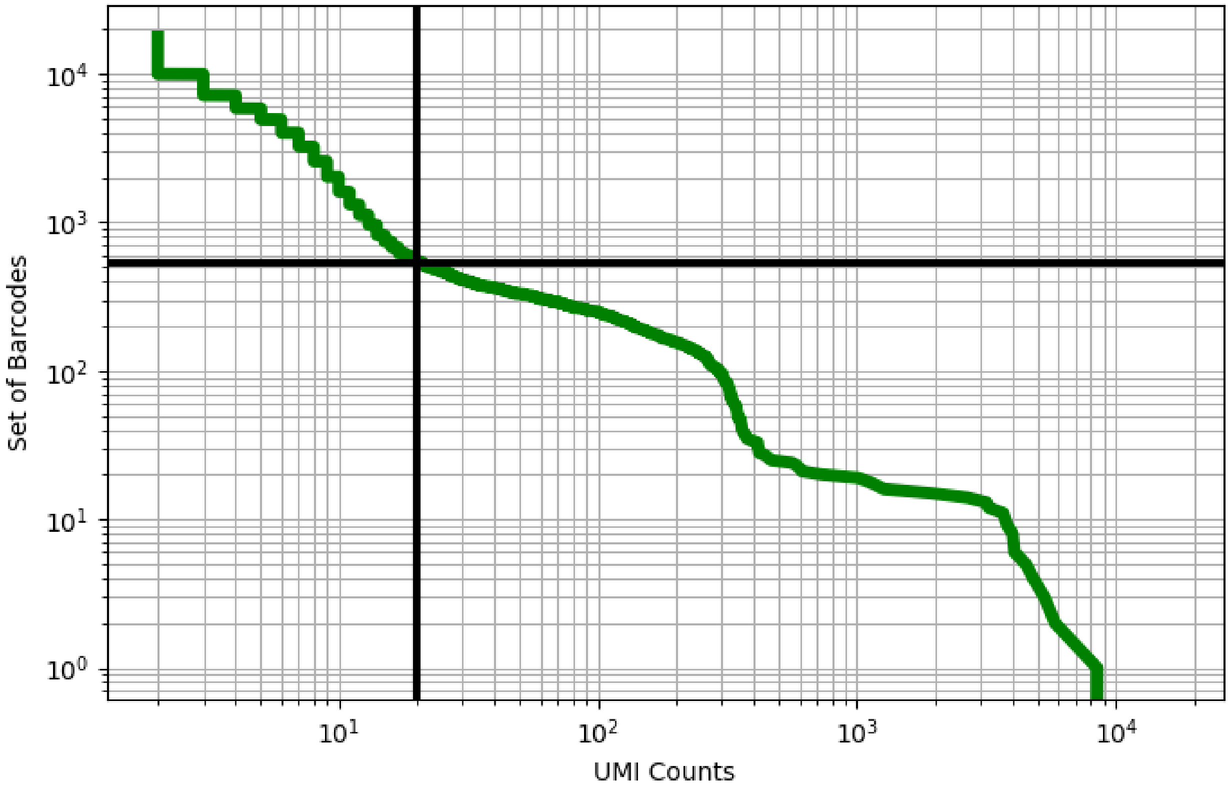


Figure 352: muscle/GSM4272895

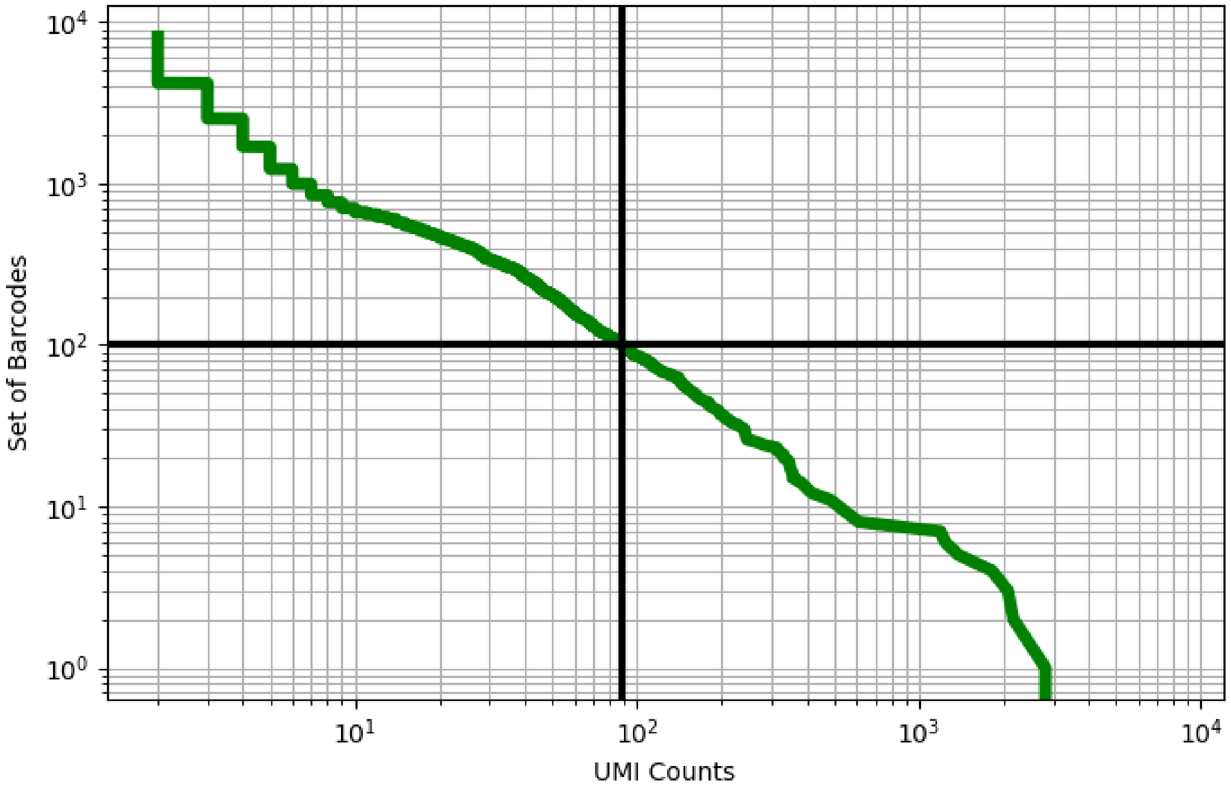


Figure 353: muscle/GSM4272896

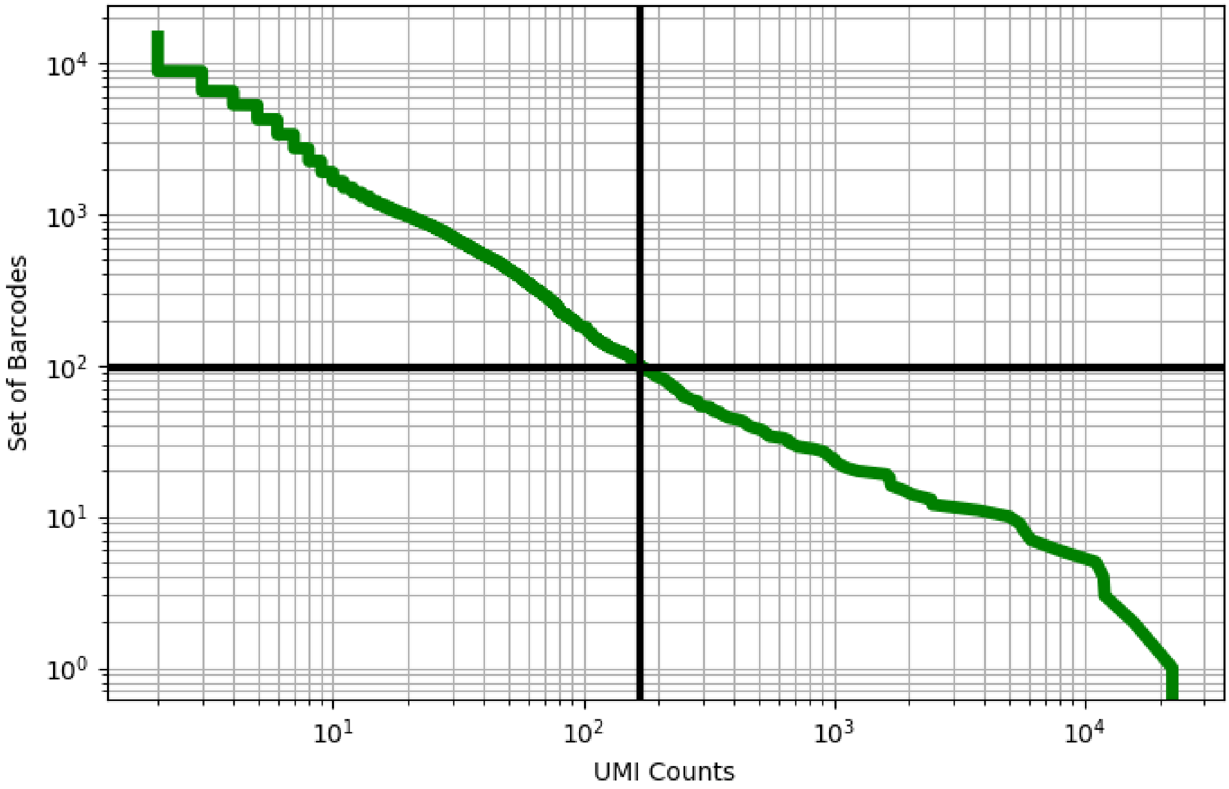


Figure 354: muscle/GSM4272897

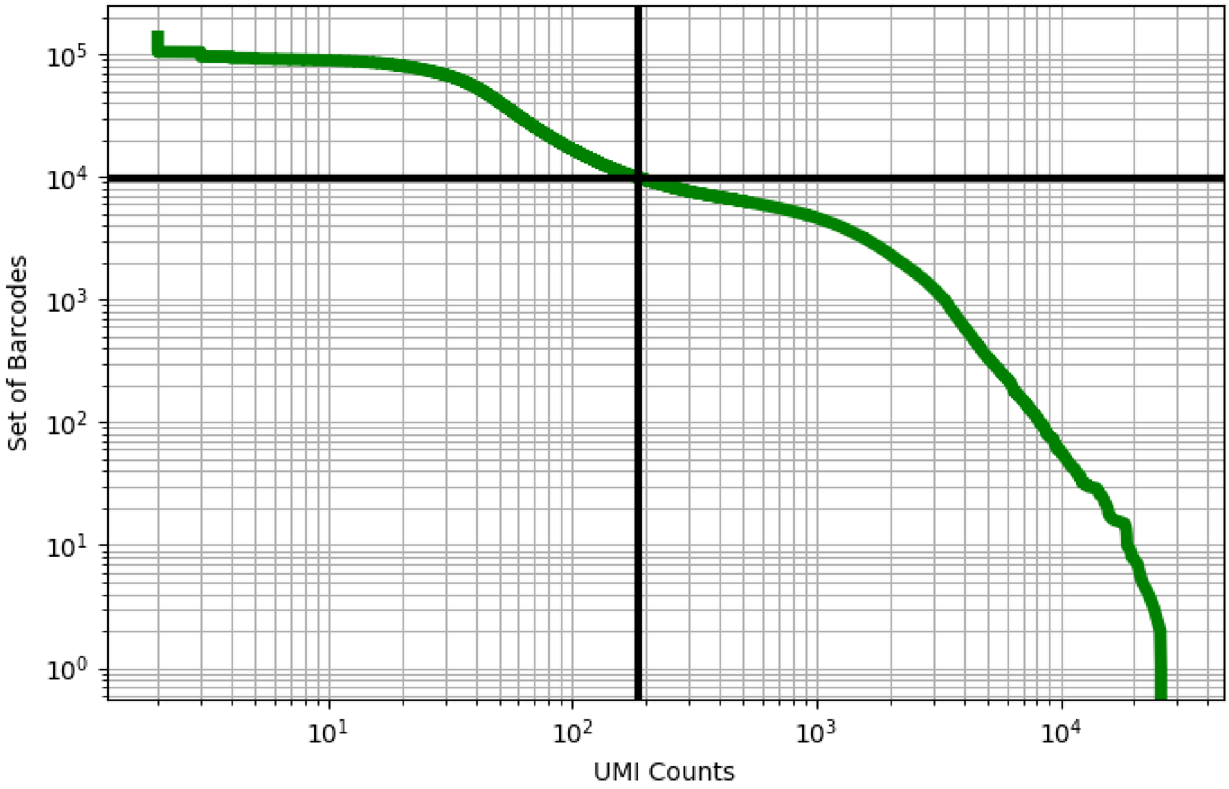


Figure 355: muscle/GSM4272898

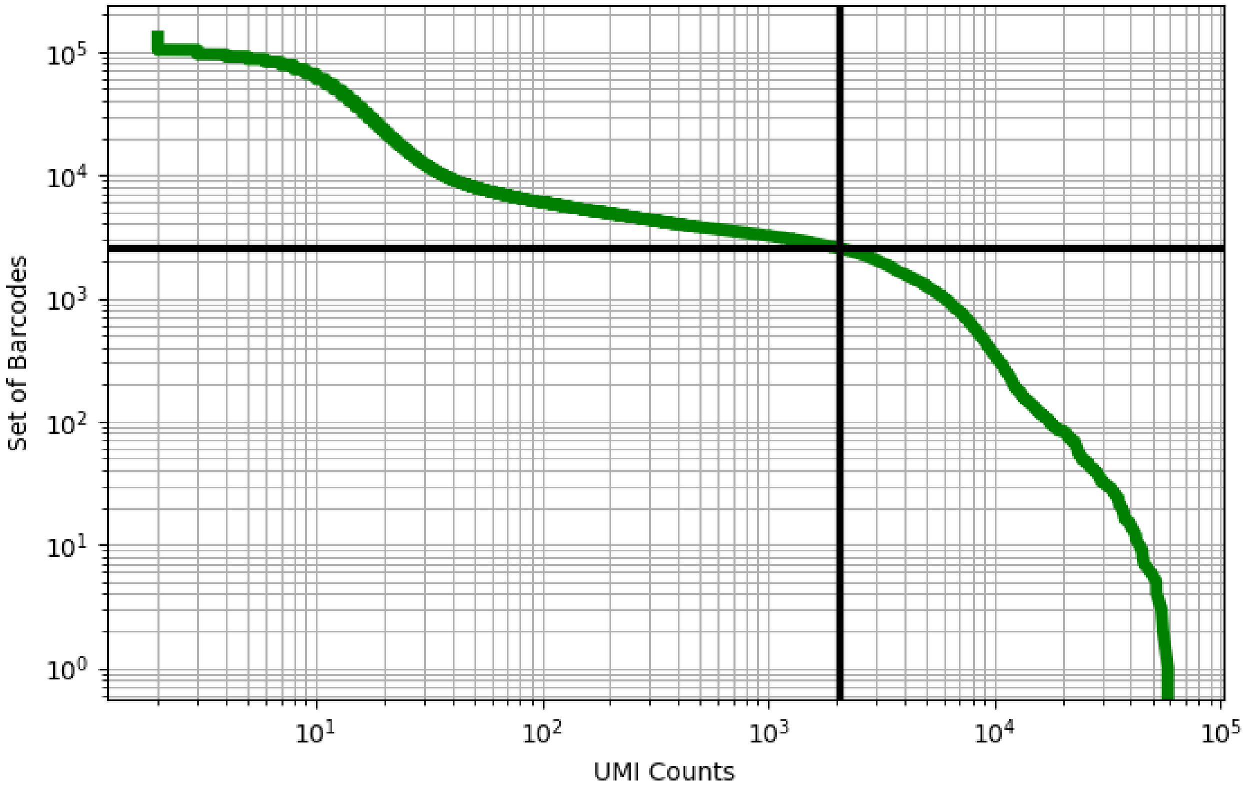


Figure 356: muscle/GSM4272899

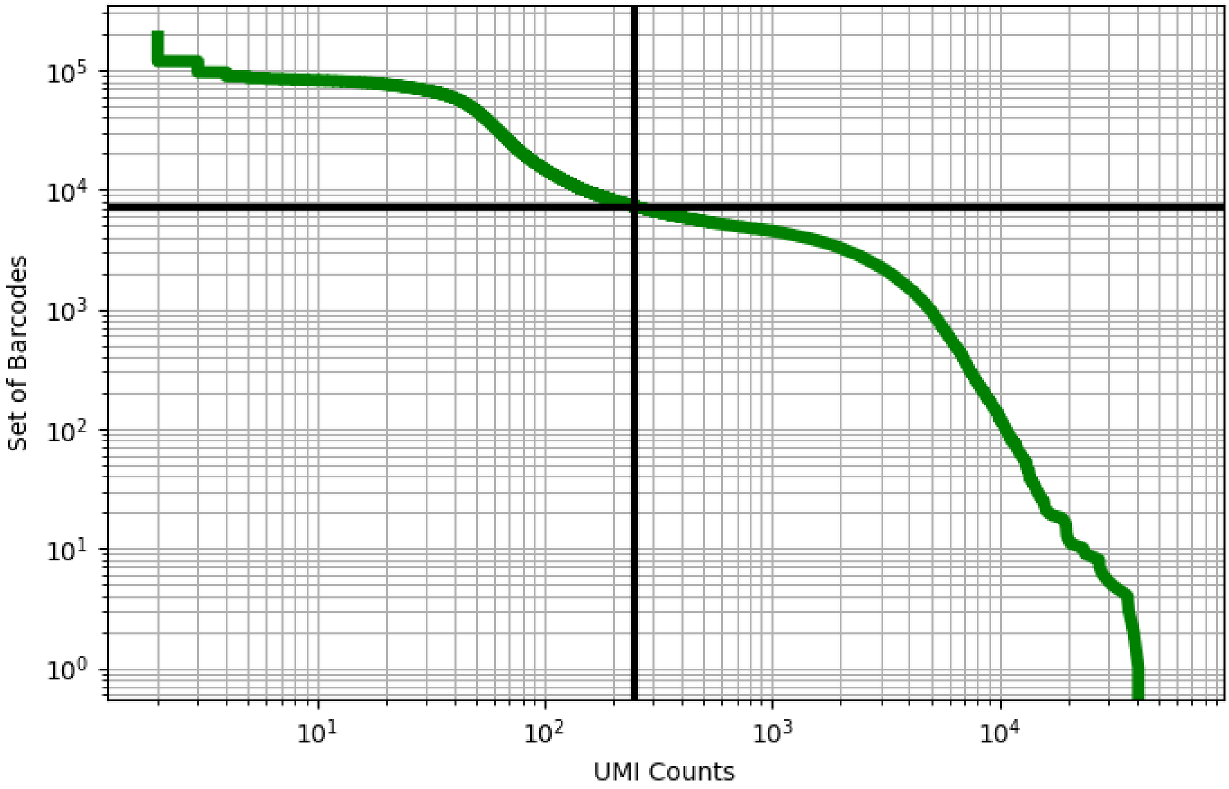


Figure 357: muscle/GSM4272900

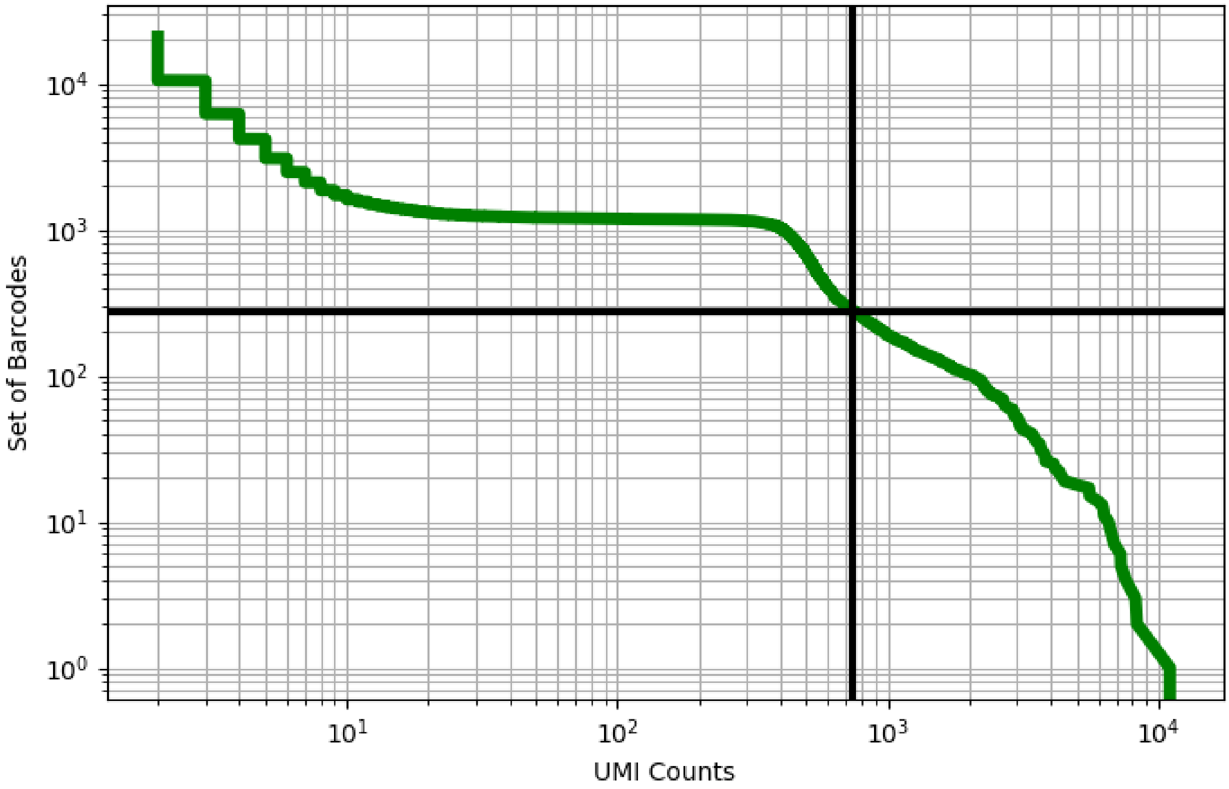


Figure 358: muscle/GSM4272901

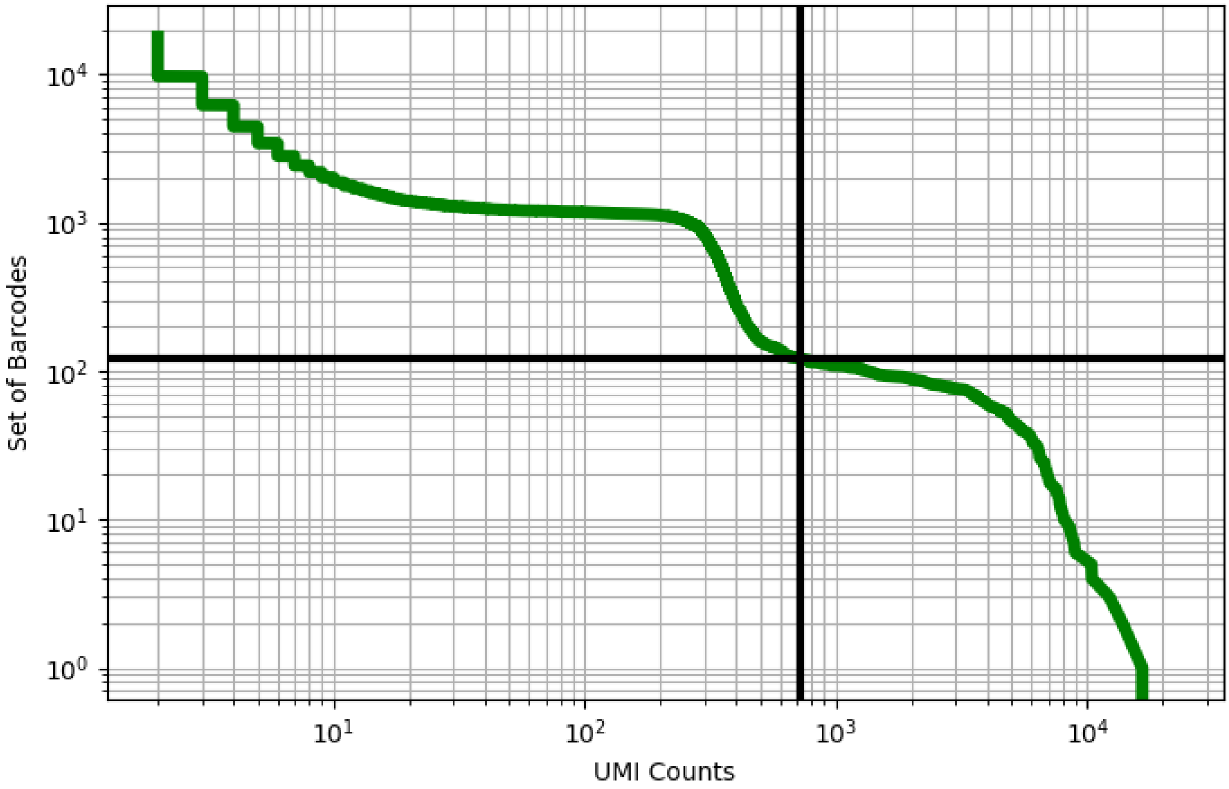


Figure 359: muscle/GSM4272902

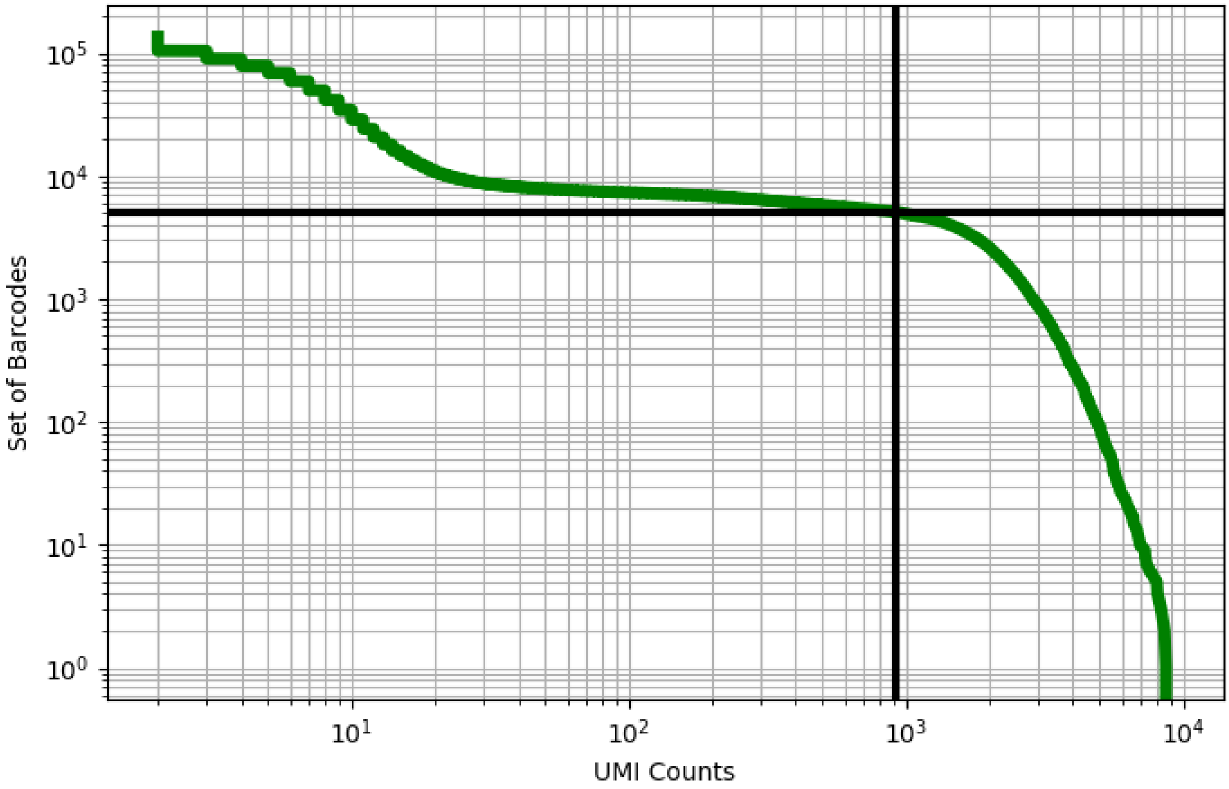


Figure 360: ovary/GSM3319032

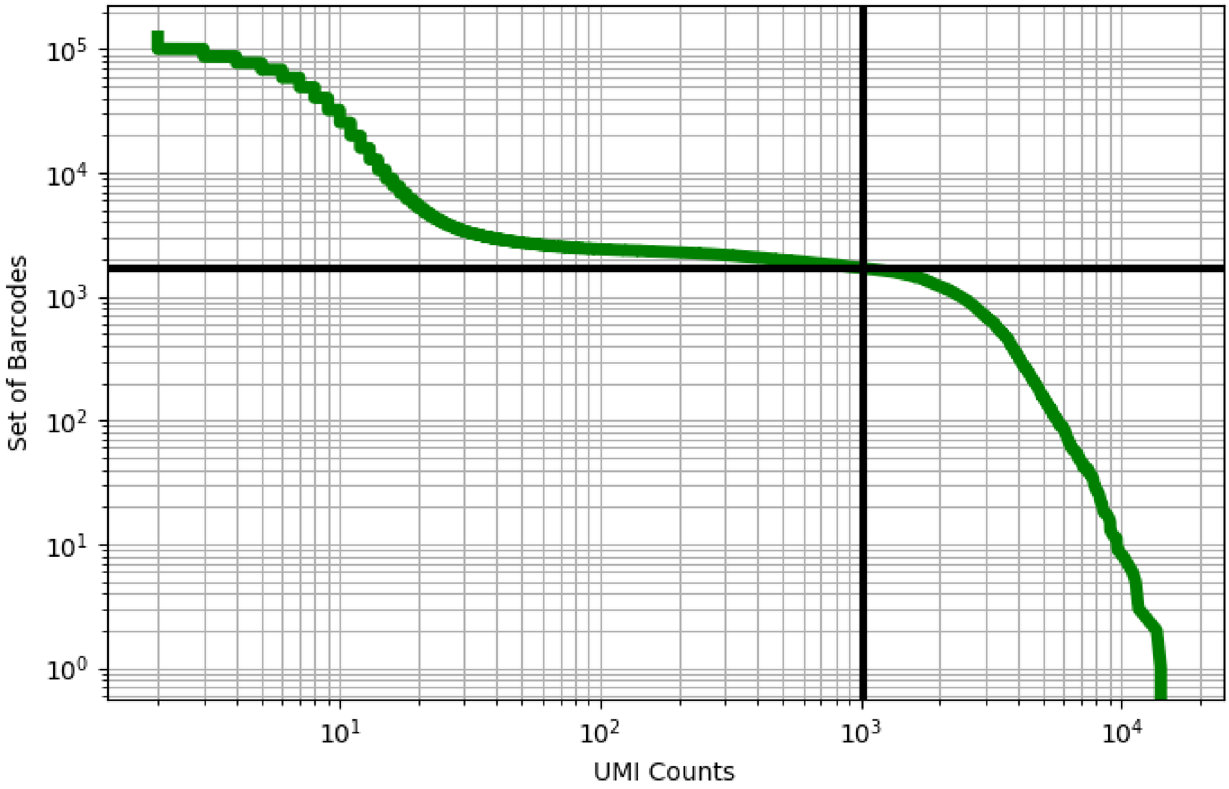


Figure 361: ovary/GSM3319033

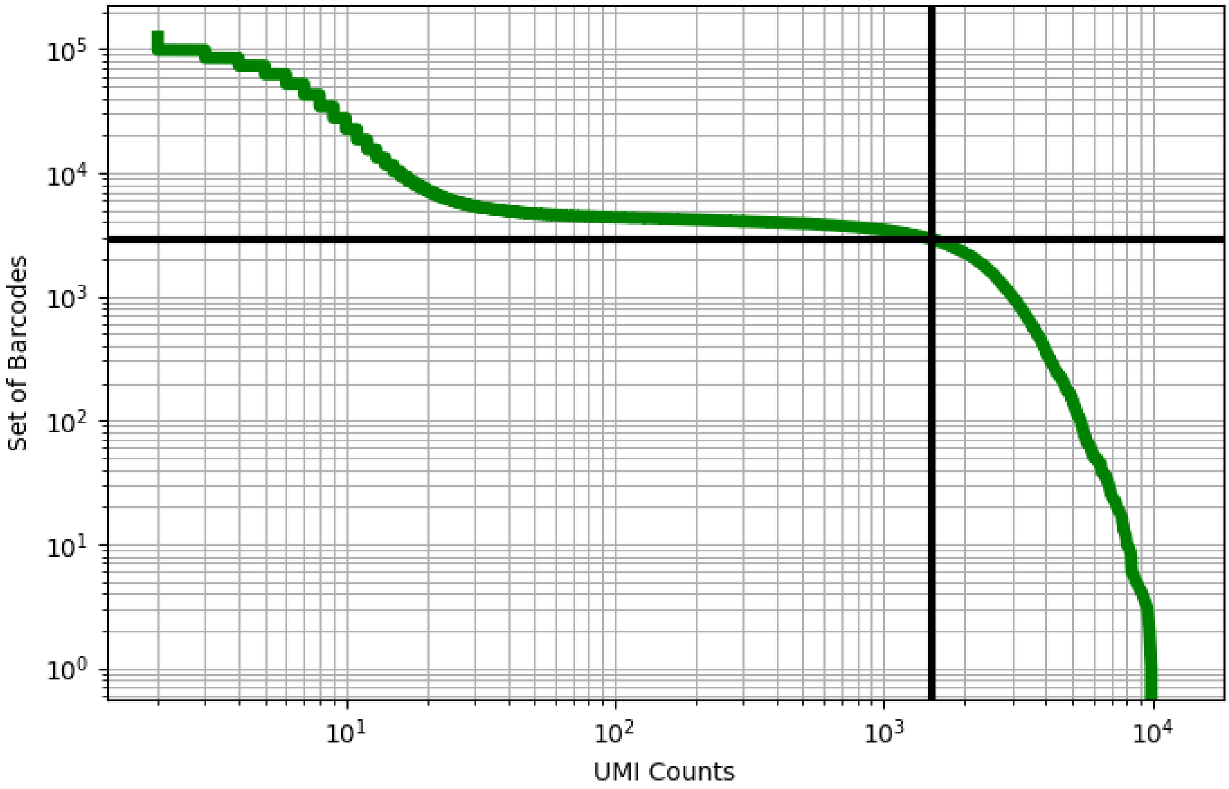


Figure 362: ovary/GSM3319034

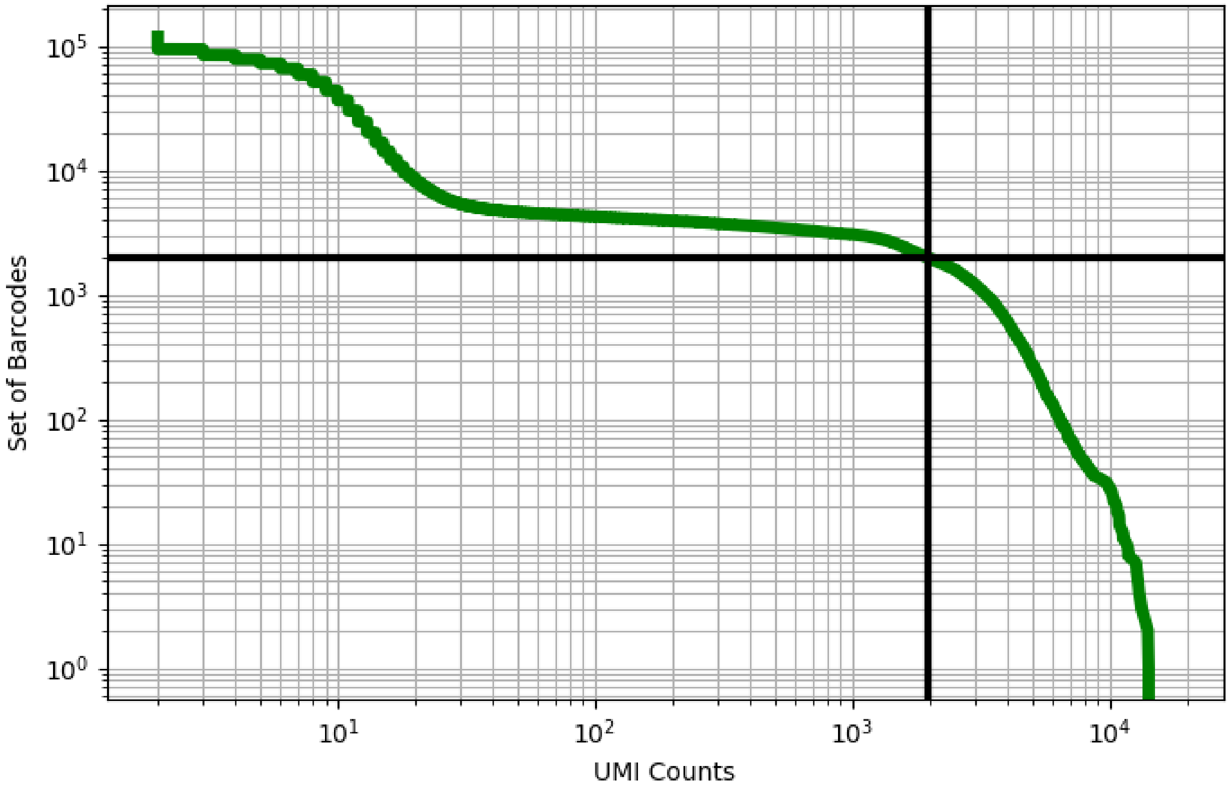


Figure 363: ovary/GSM3319035

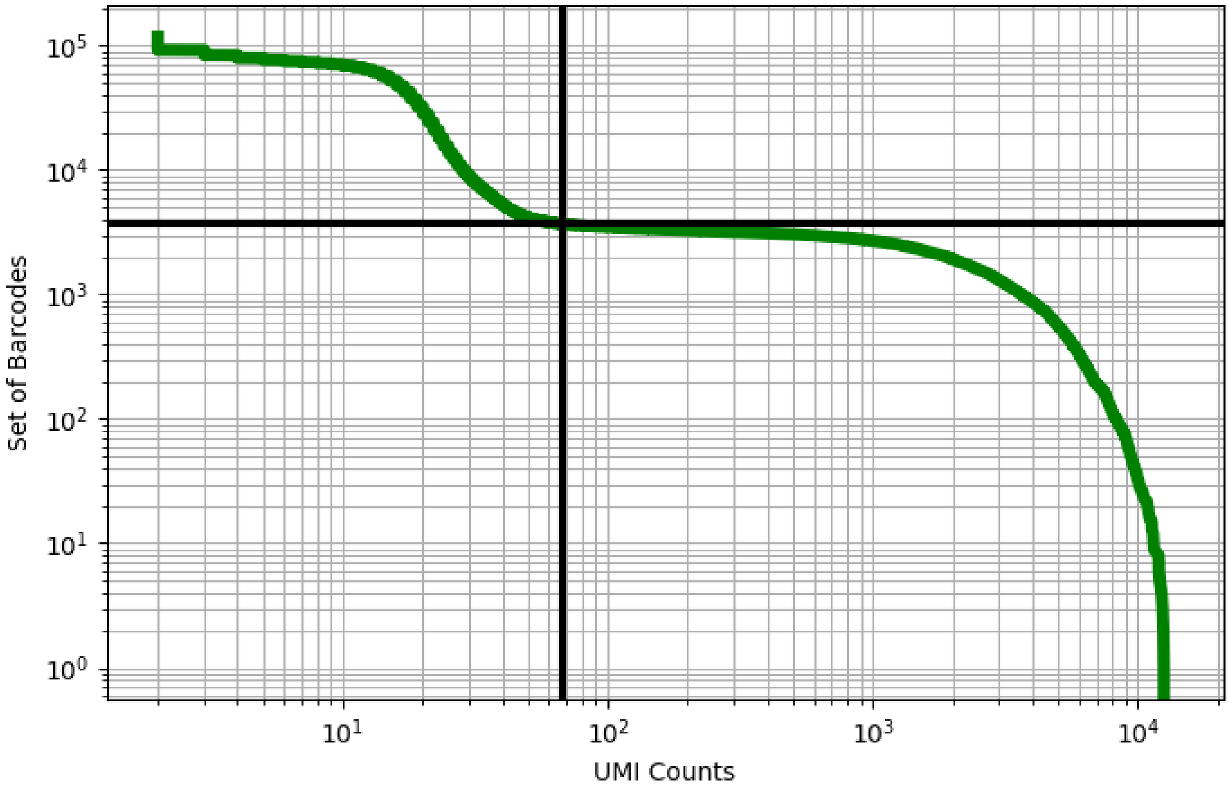


Figure 364: ovary/GSM3319036

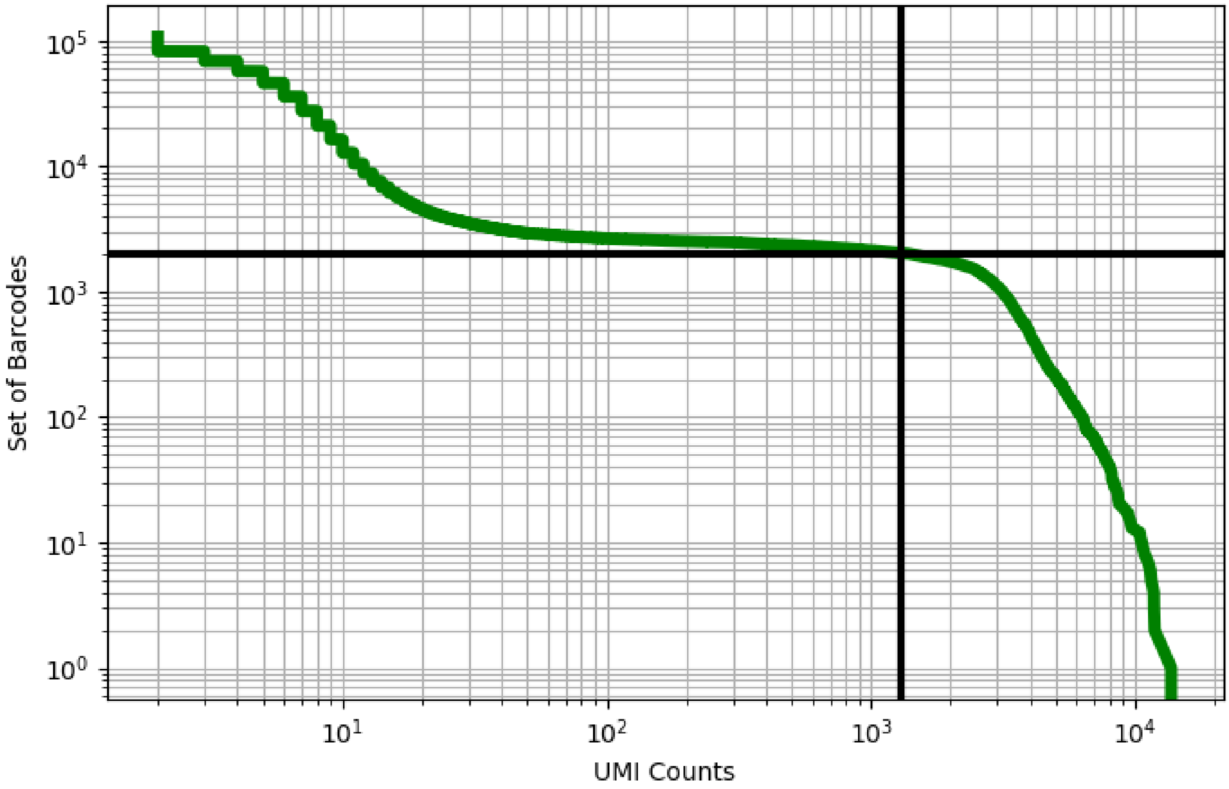


Figure 365: ovary/GSM3319037

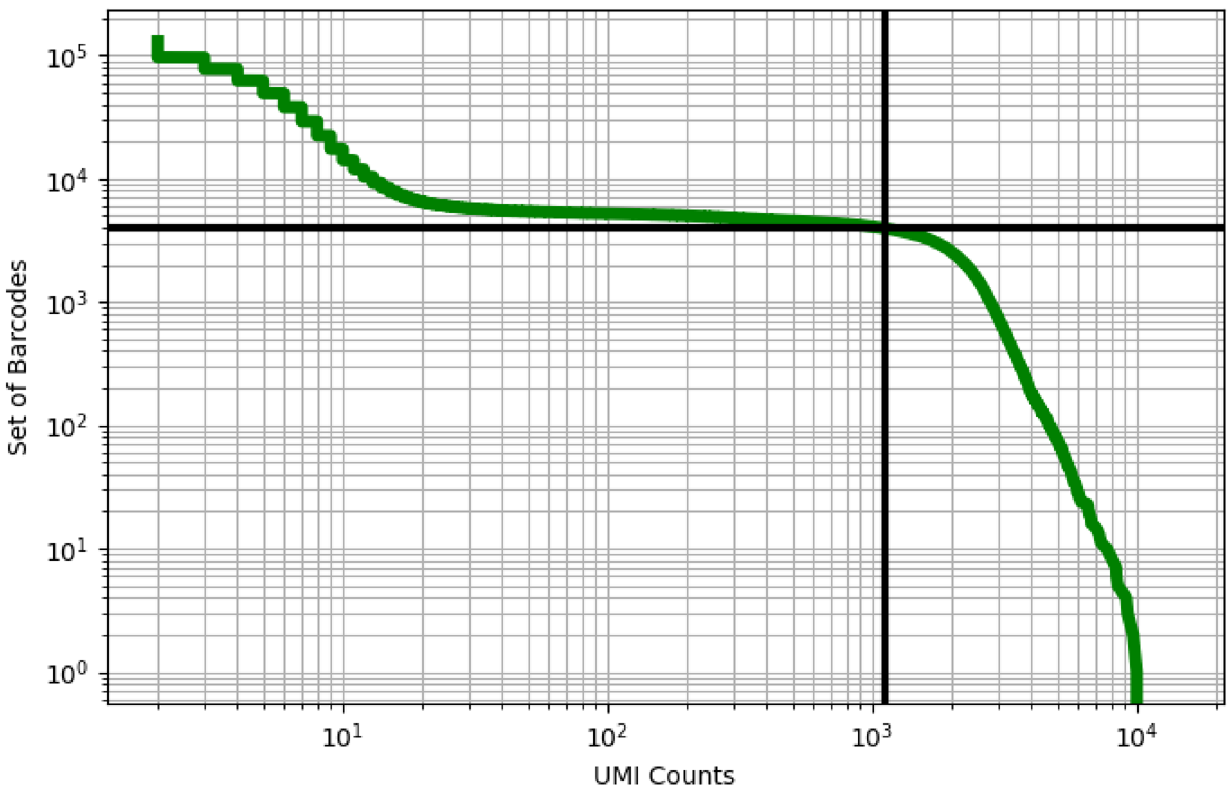


Figure 366: ovary/GSM3319038

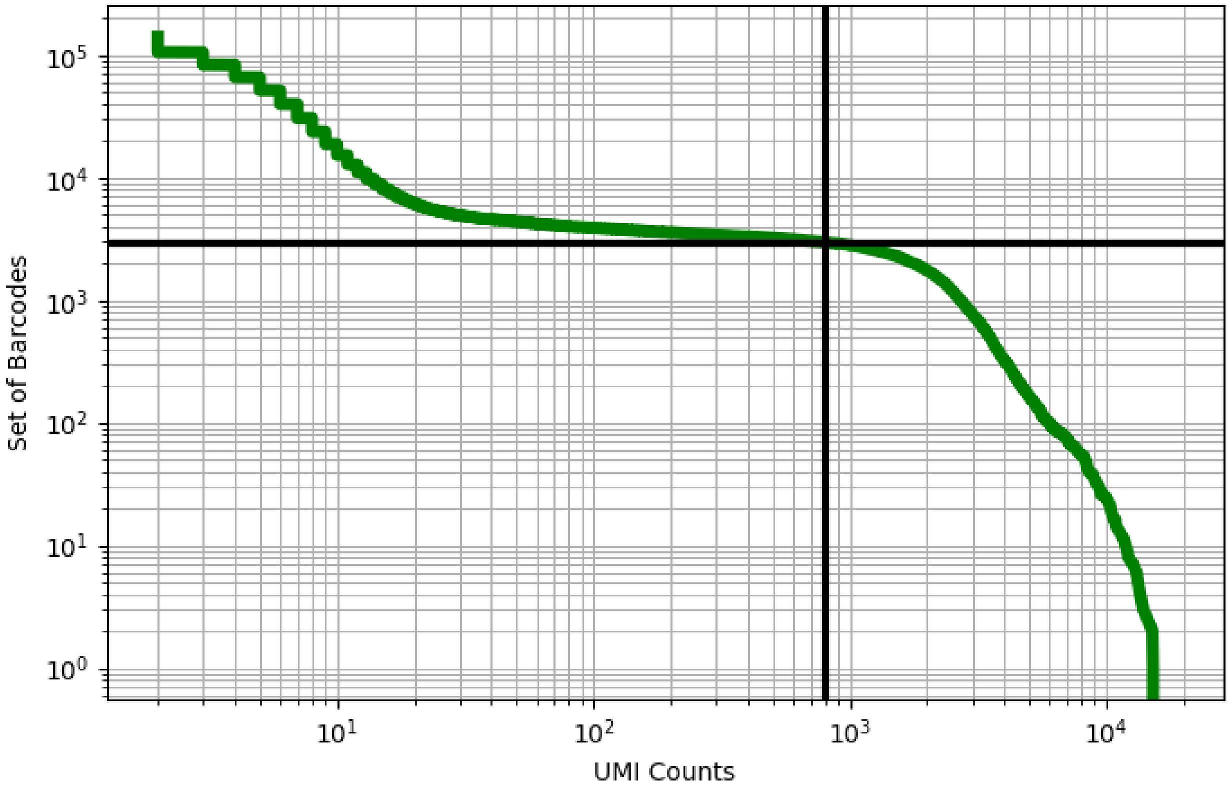


Figure 367: ovary/GSM3319039

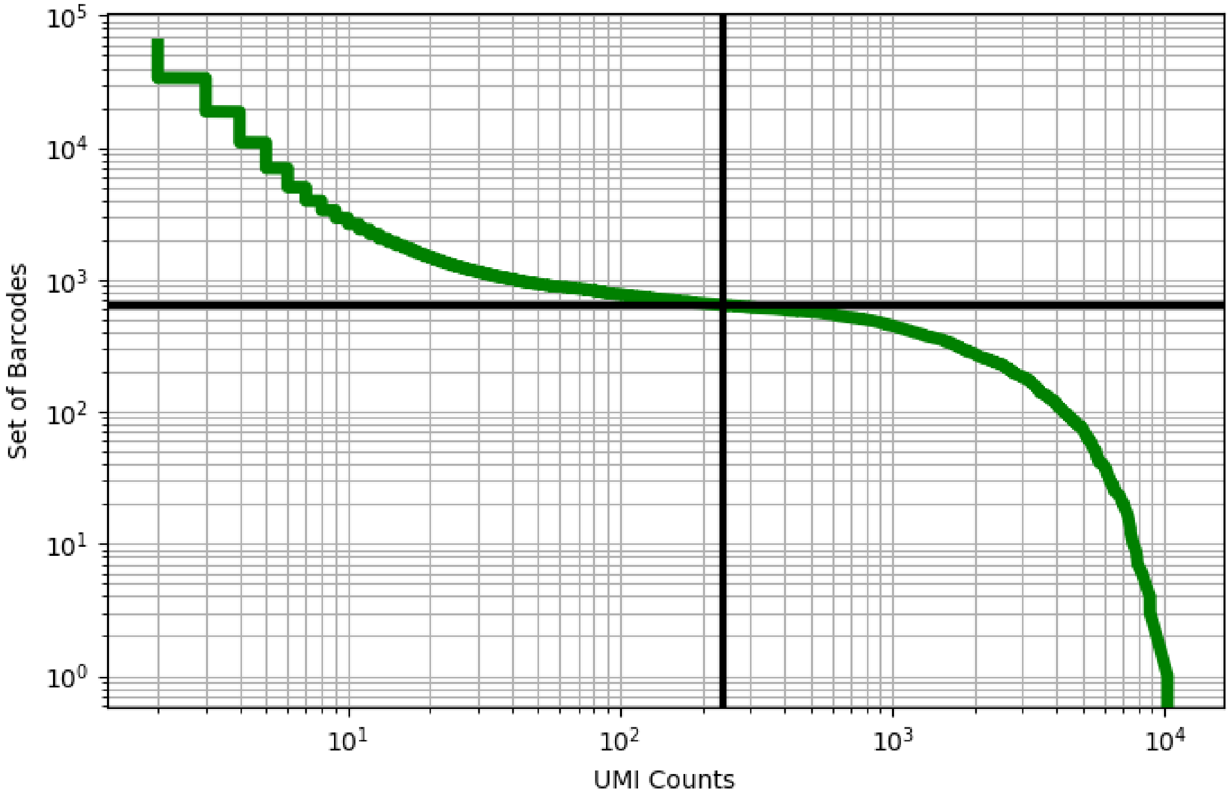


Figure 368: ovary/GSM3319040

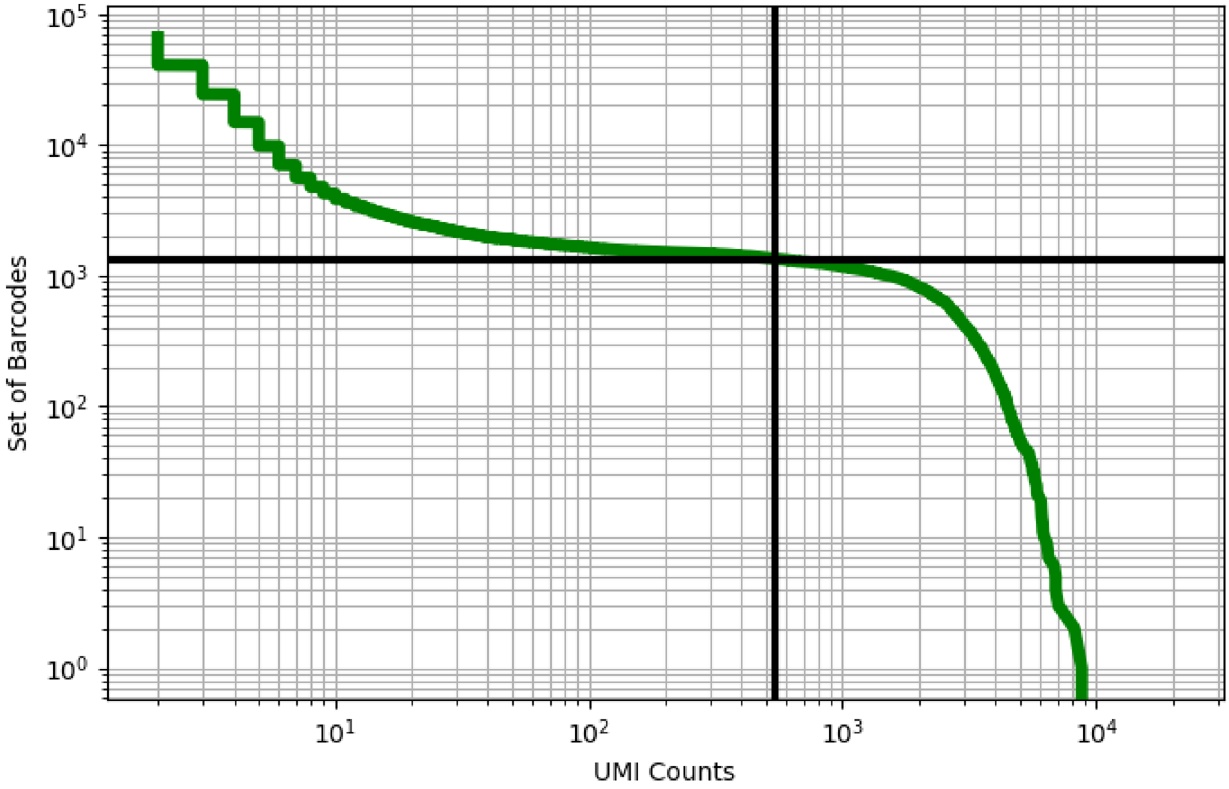


Figure 369: ovary/GSM3319041

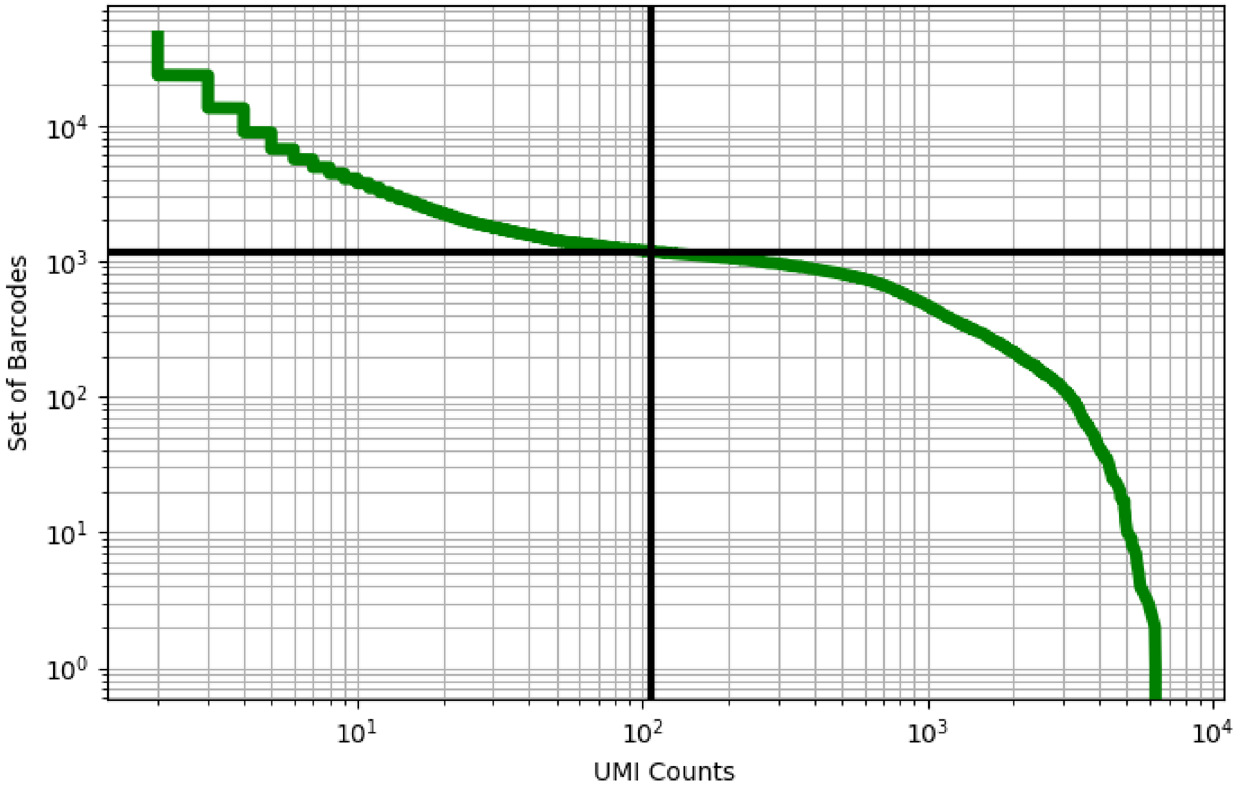


Figure 370: ovary/GSM3319042

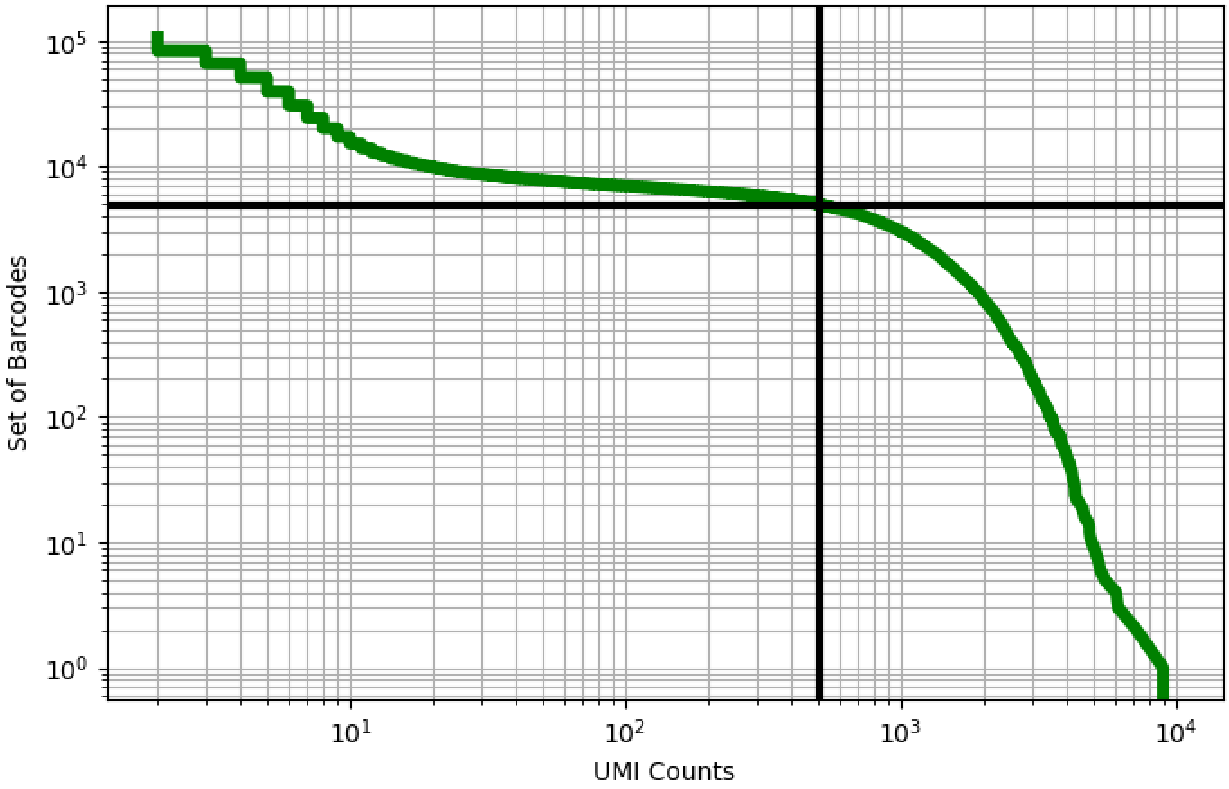


Figure 371: ovary/GSM3319043

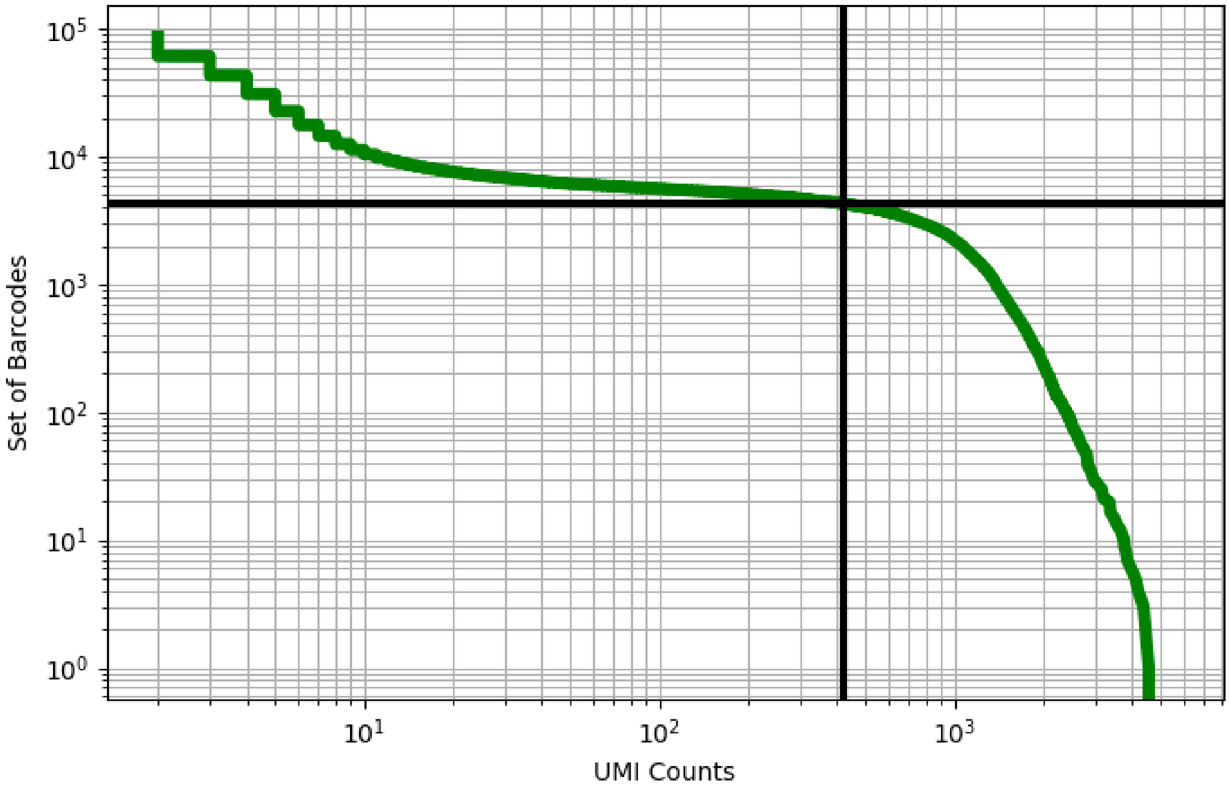


Figure 372: ovary/GSM3319044

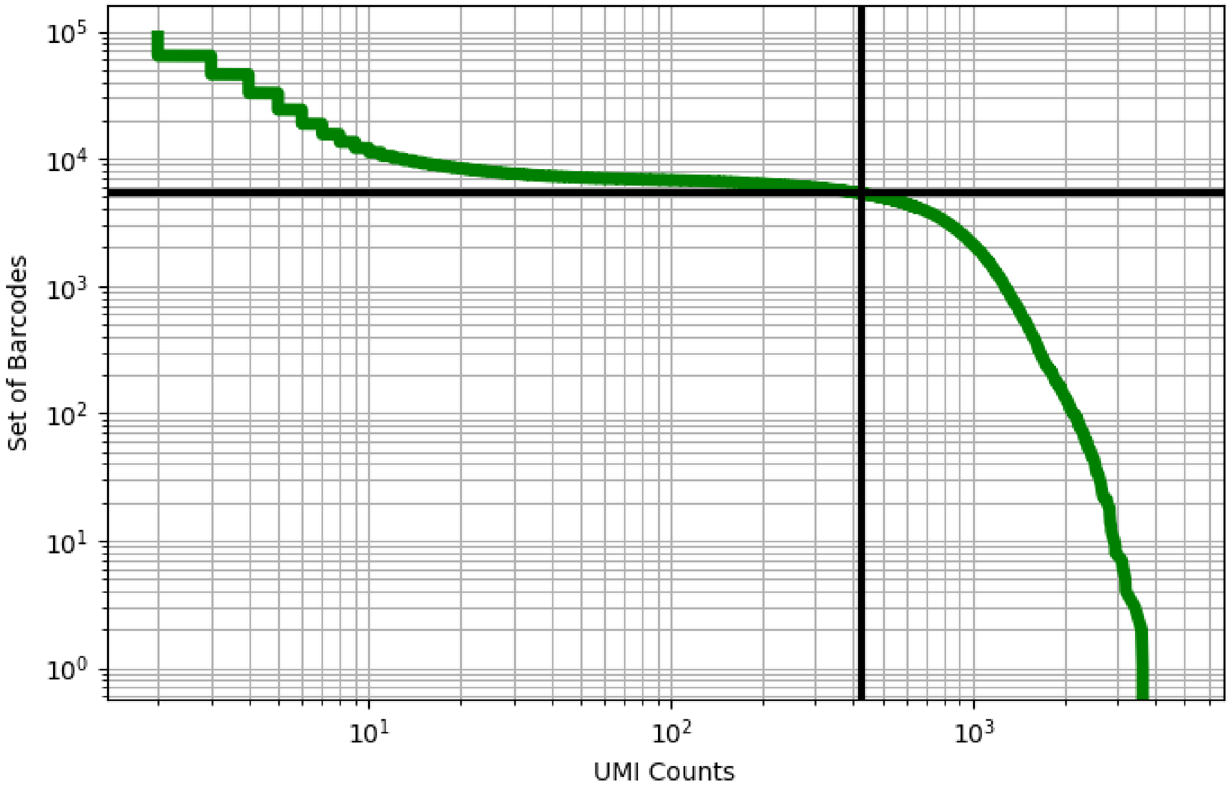


Figure 373: ovary/GSM3319045

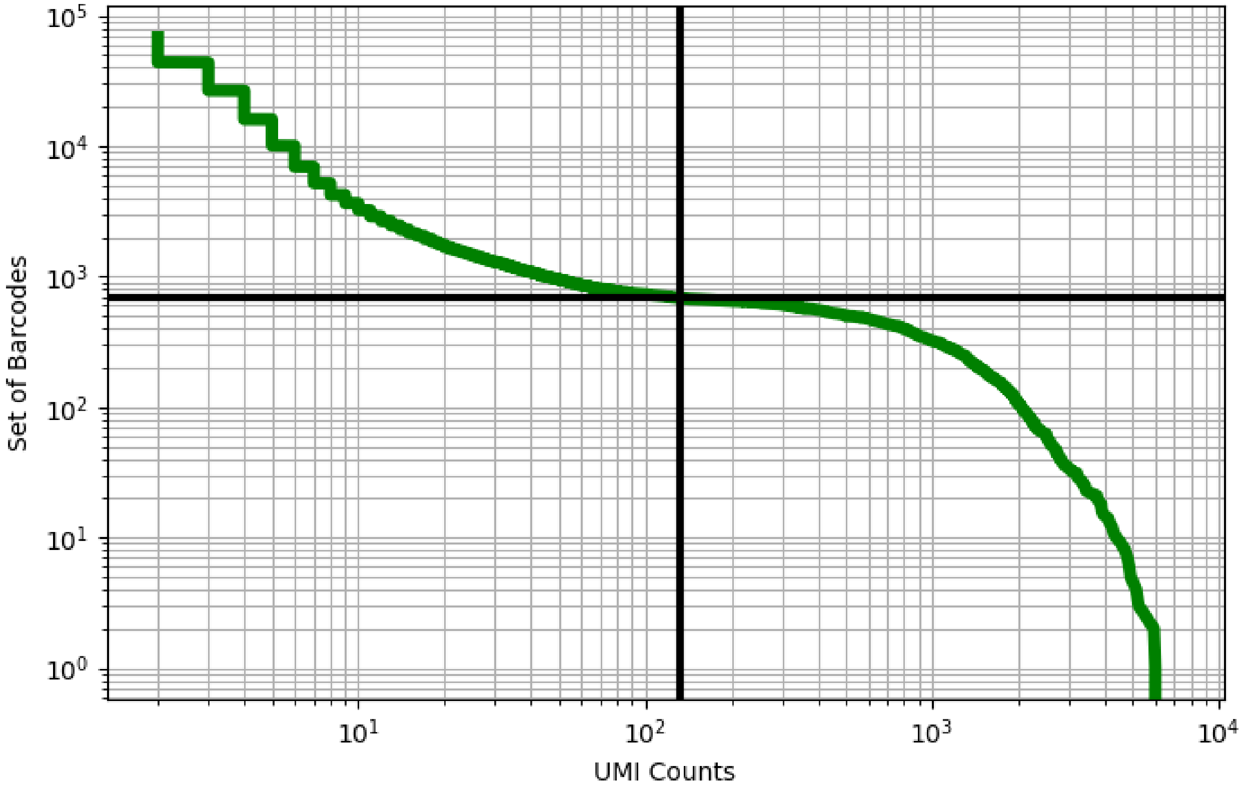


Figure 374: ovary/GSM3319046

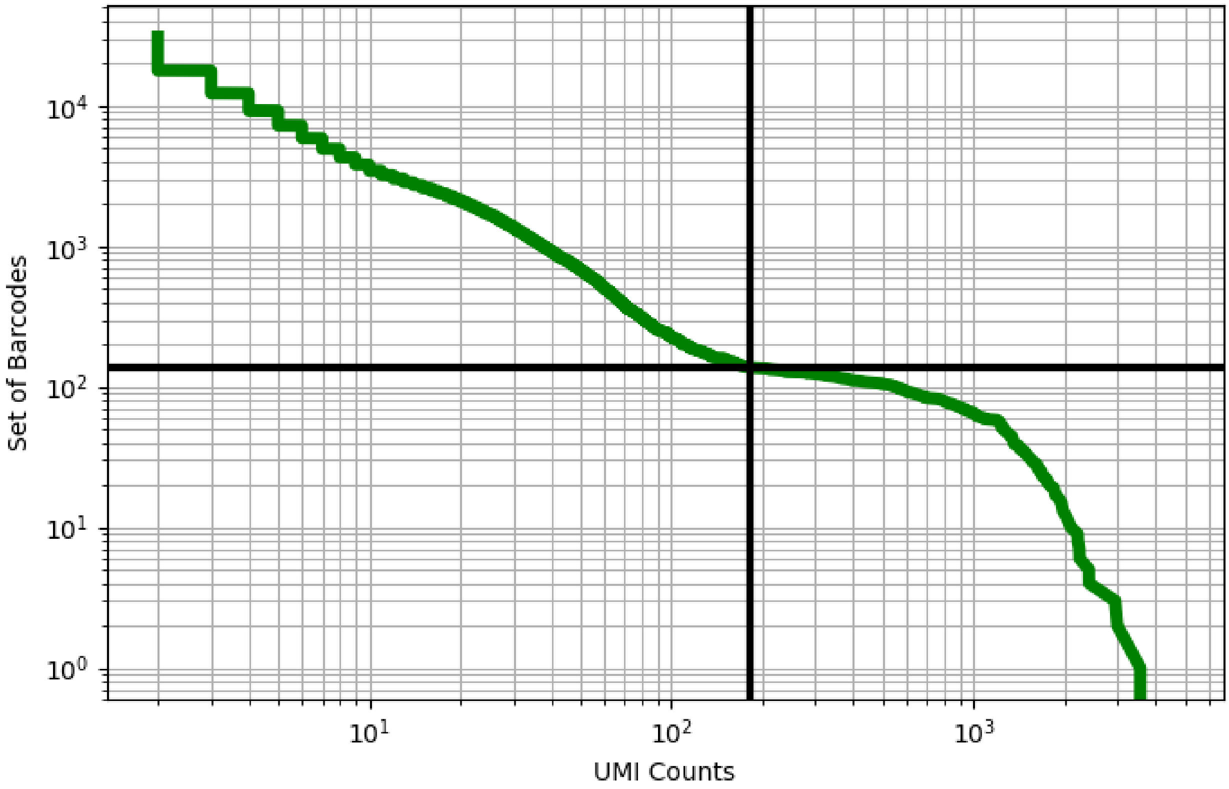


Figure 375: ovary/GSM3319047

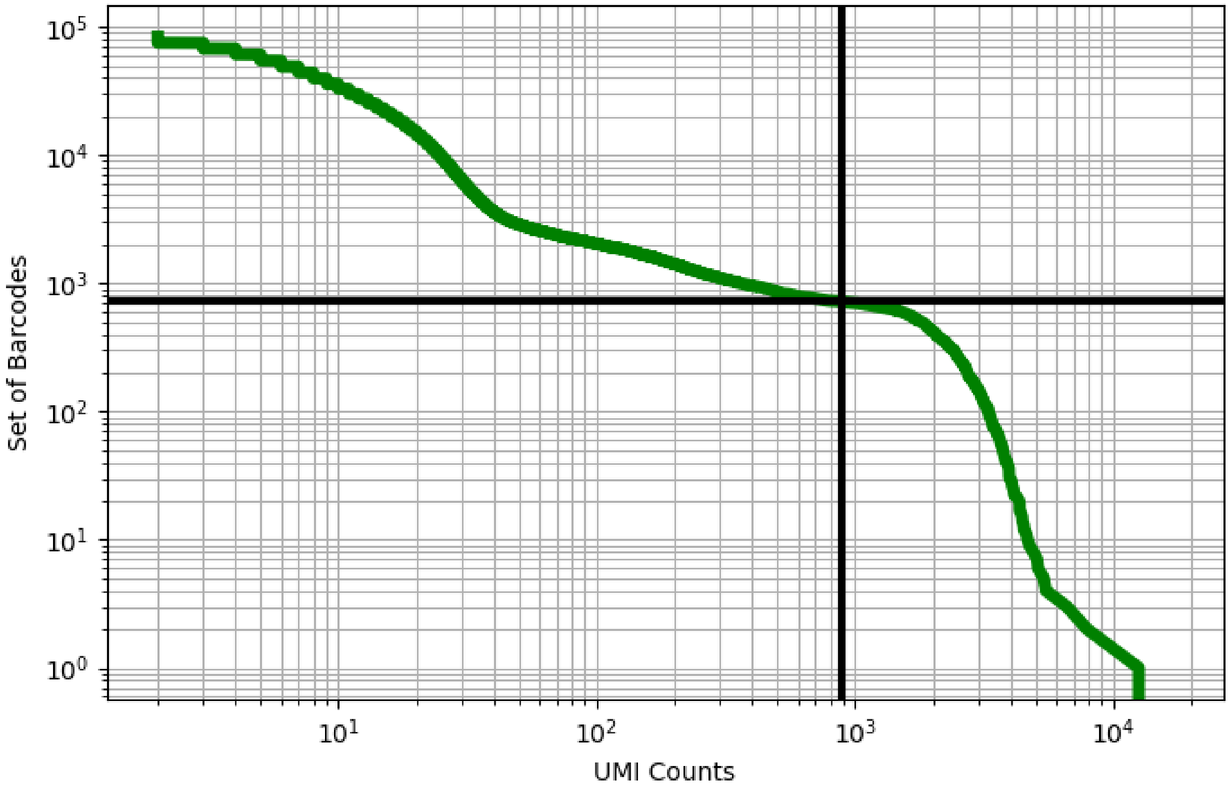


Figure 376: ovary/GSM3557959

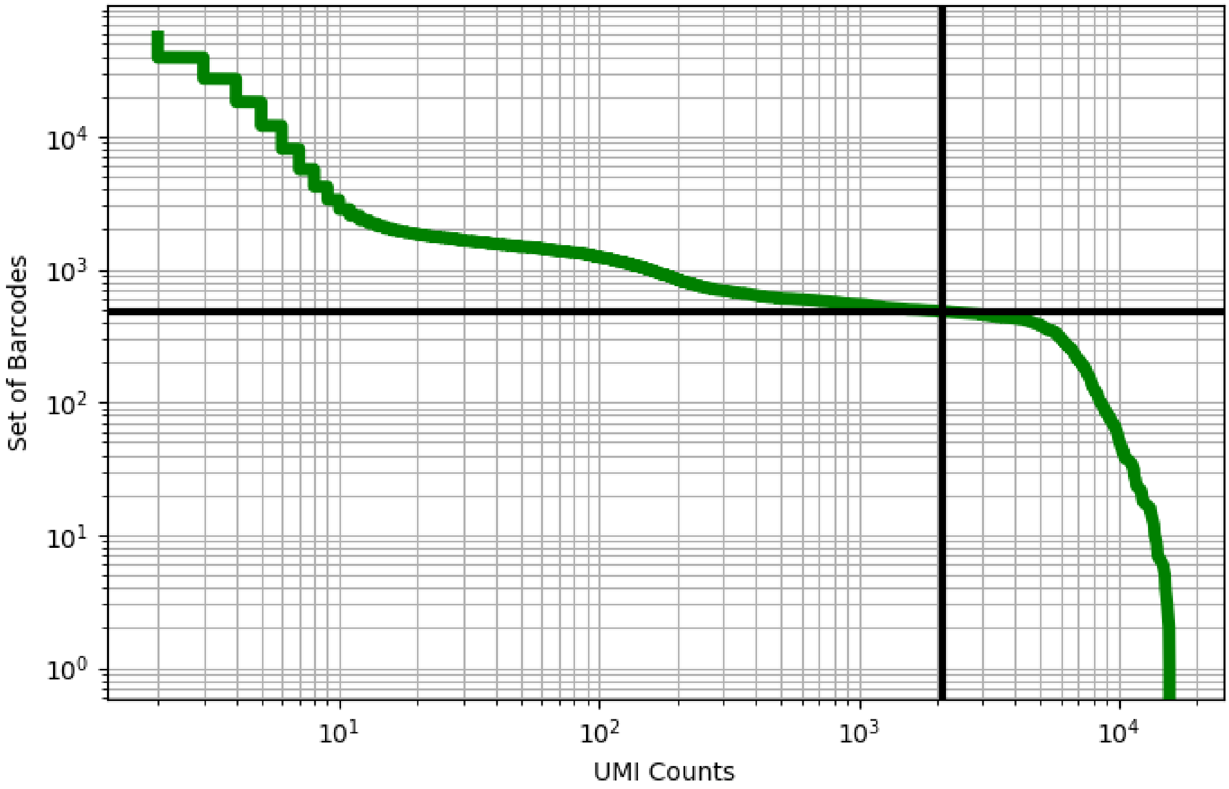


Figure 377: ovary/GSM3557960

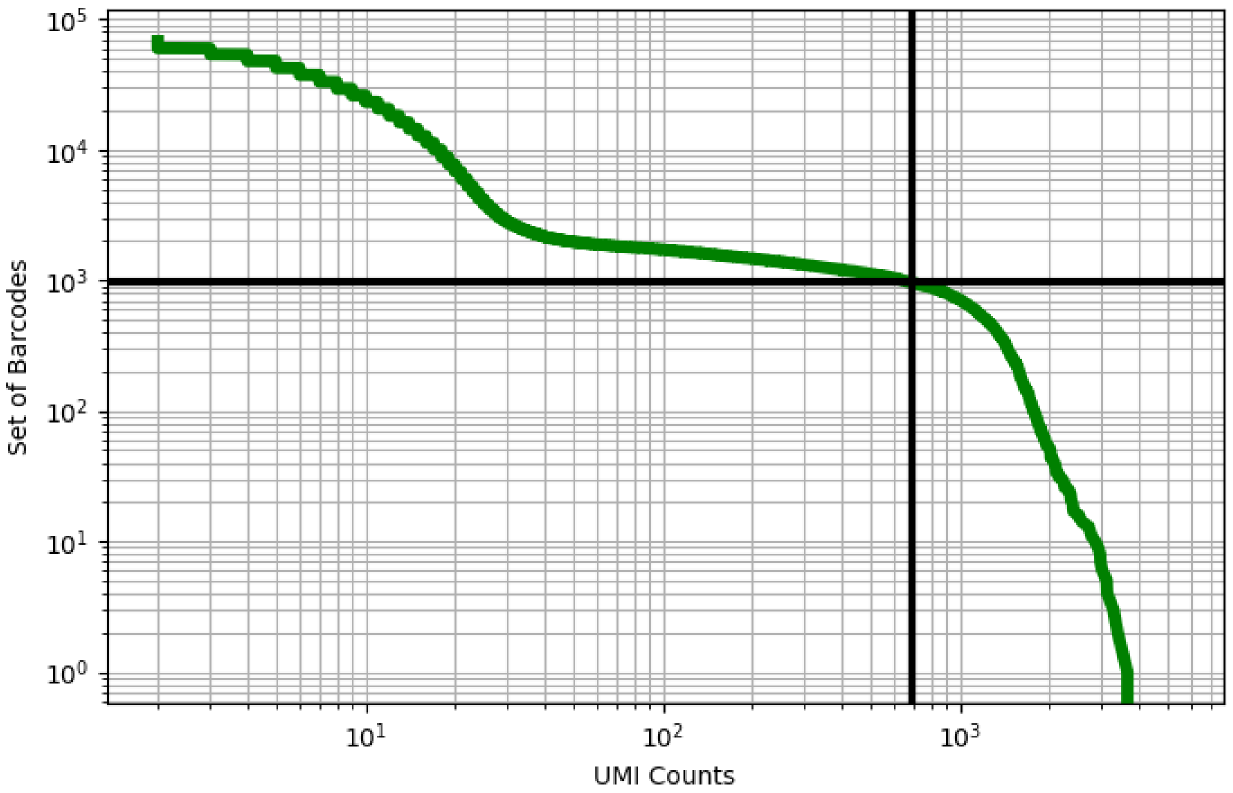


Figure 378: ovary/GSM3557961

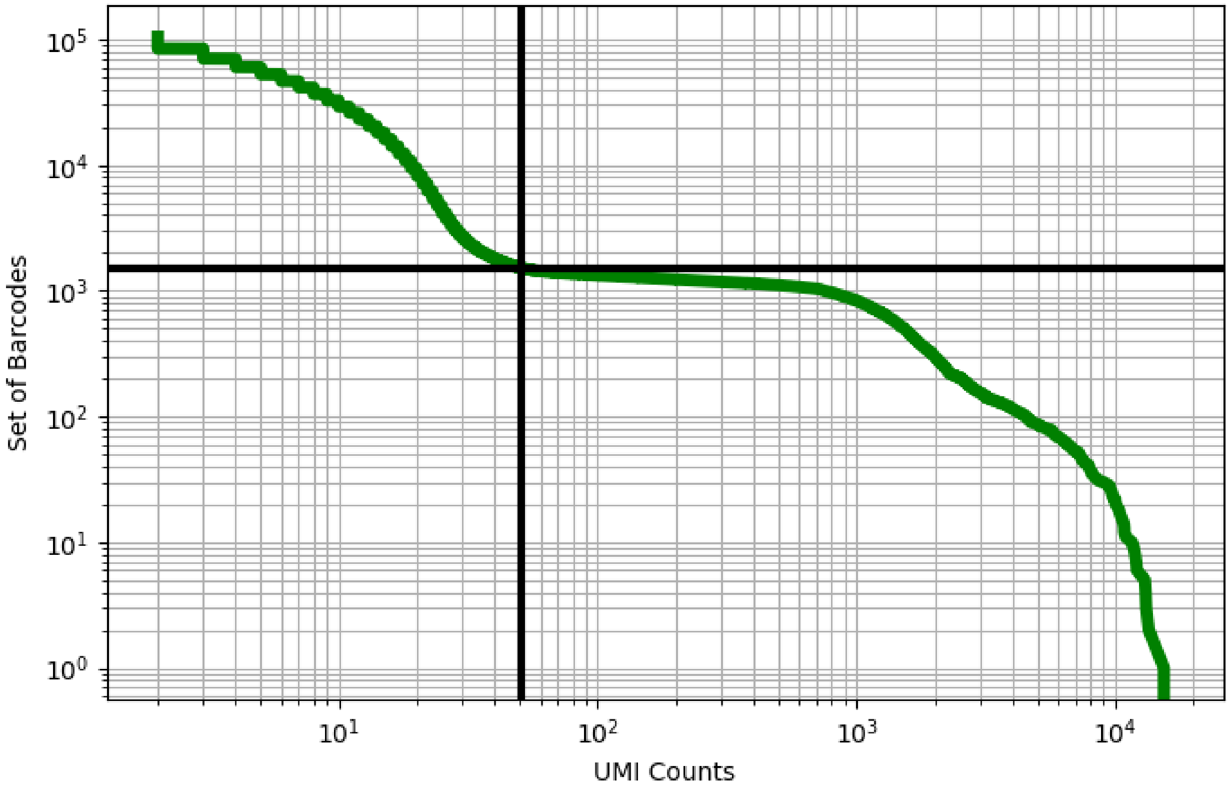


Figure 379: ovary/GSM3557962

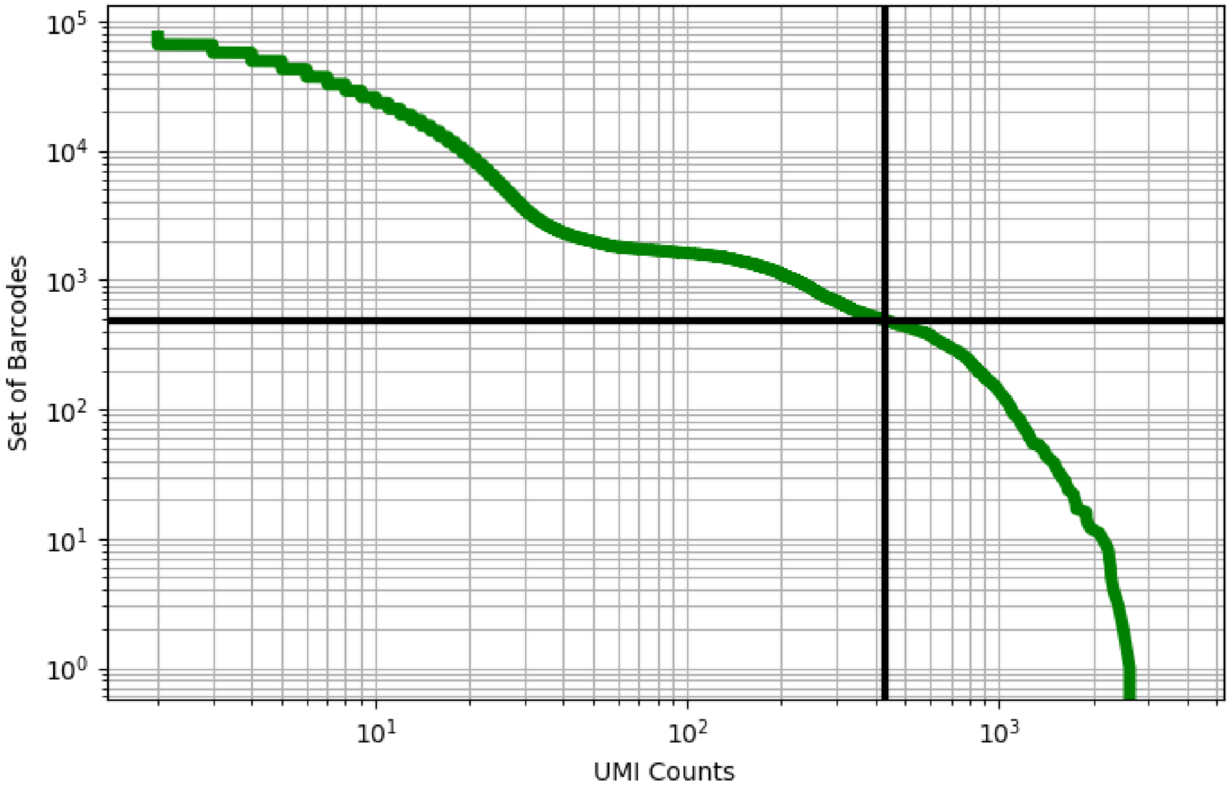


Figure 380: ovary/GSM3557963

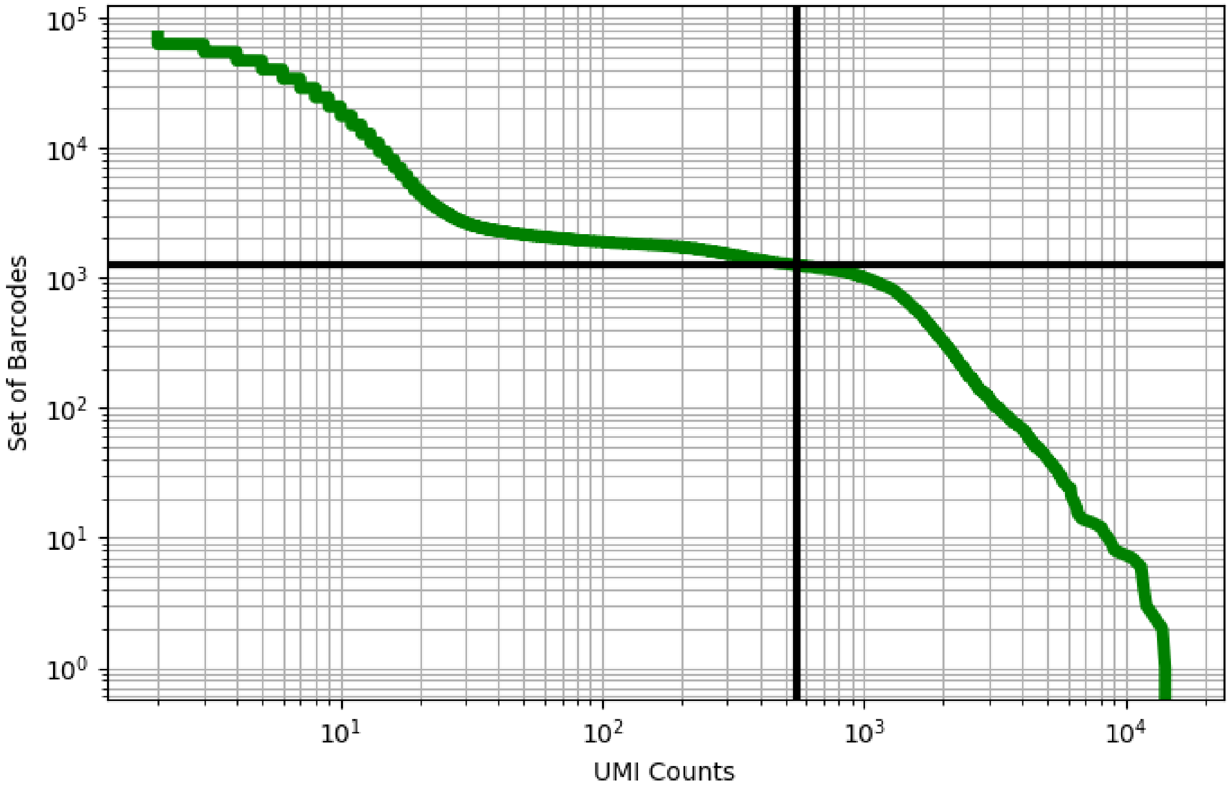


Figure 381: ovary/GSM3557964

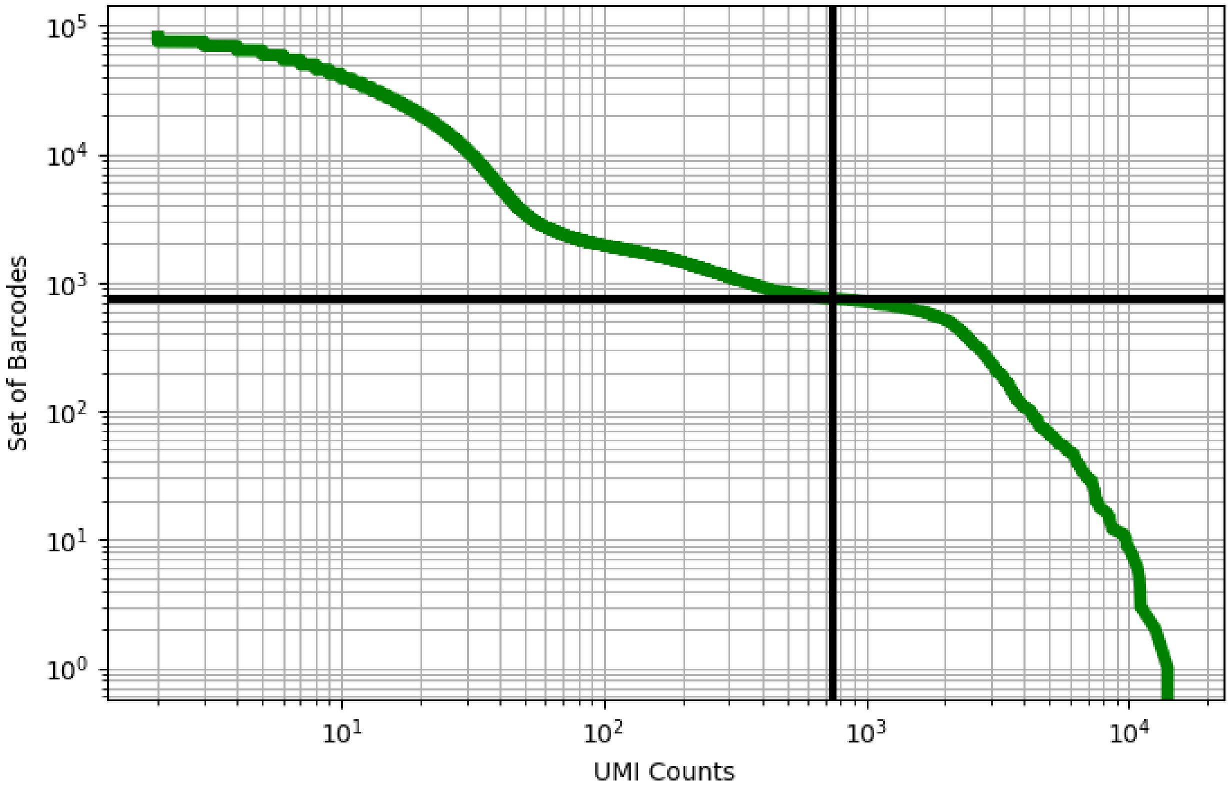


Figure 382: ovary/GSM3557965

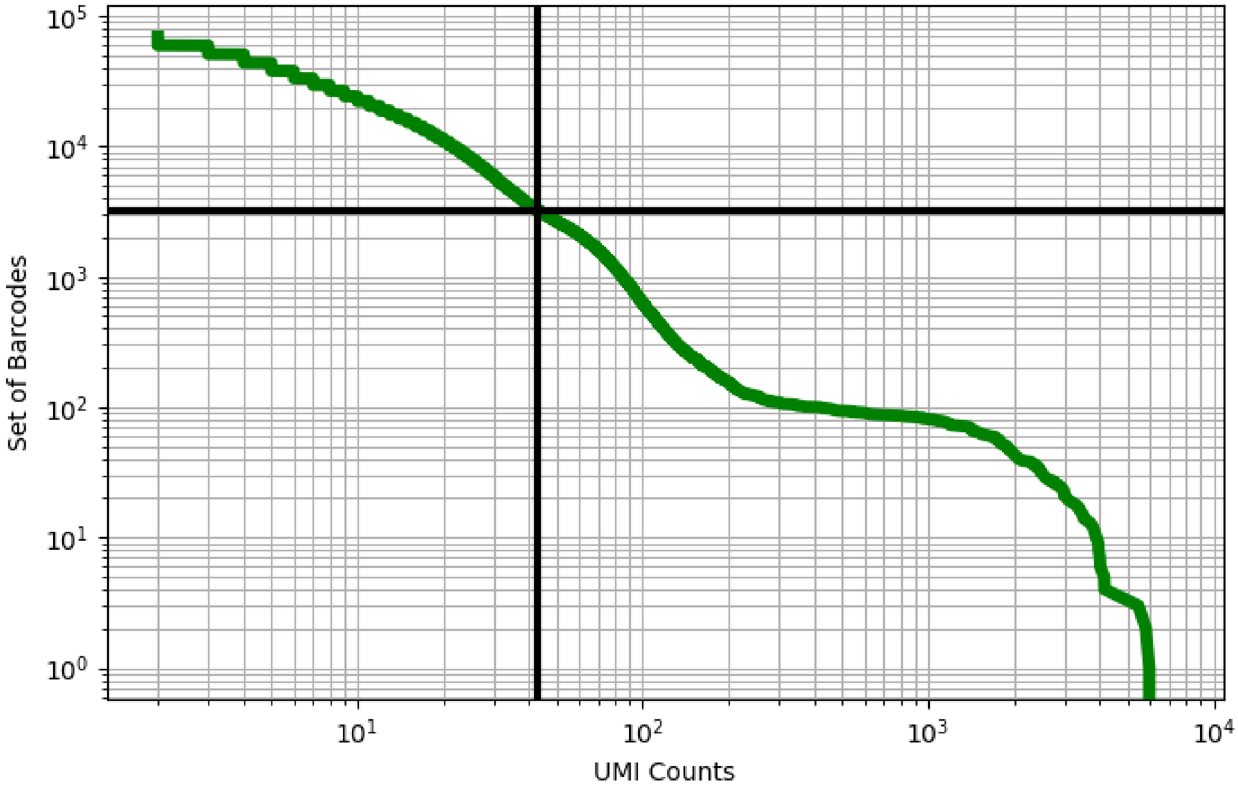


Figure 383: ovary/GSM3557966

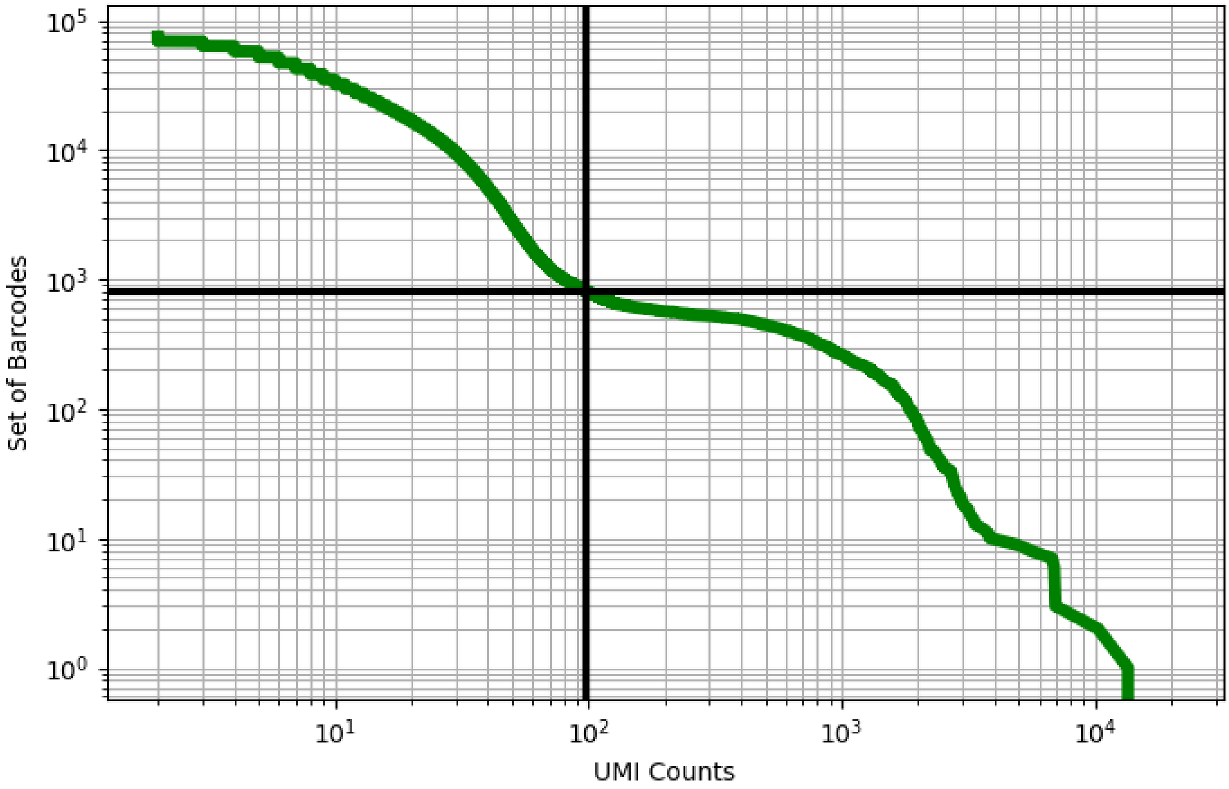


Figure 384: ovary/GSM3557967

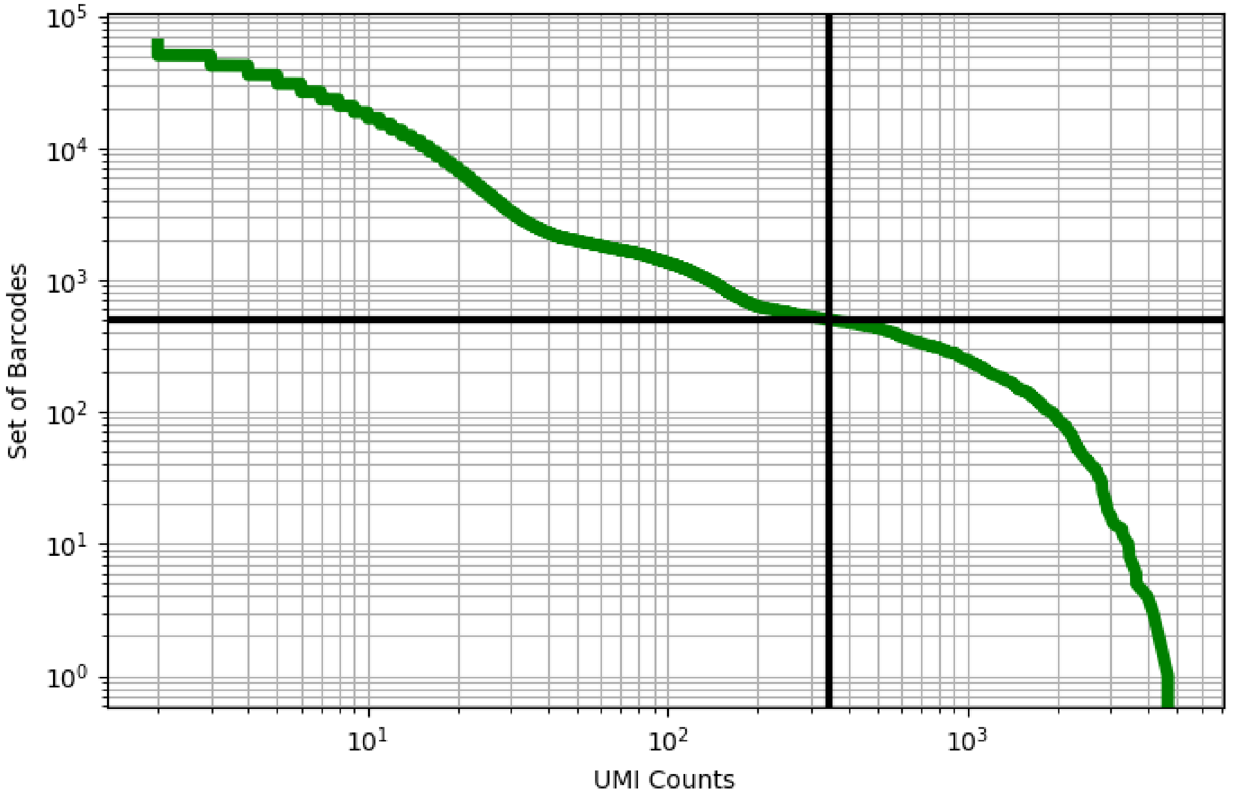


Figure 385: ovary/GSM3557968

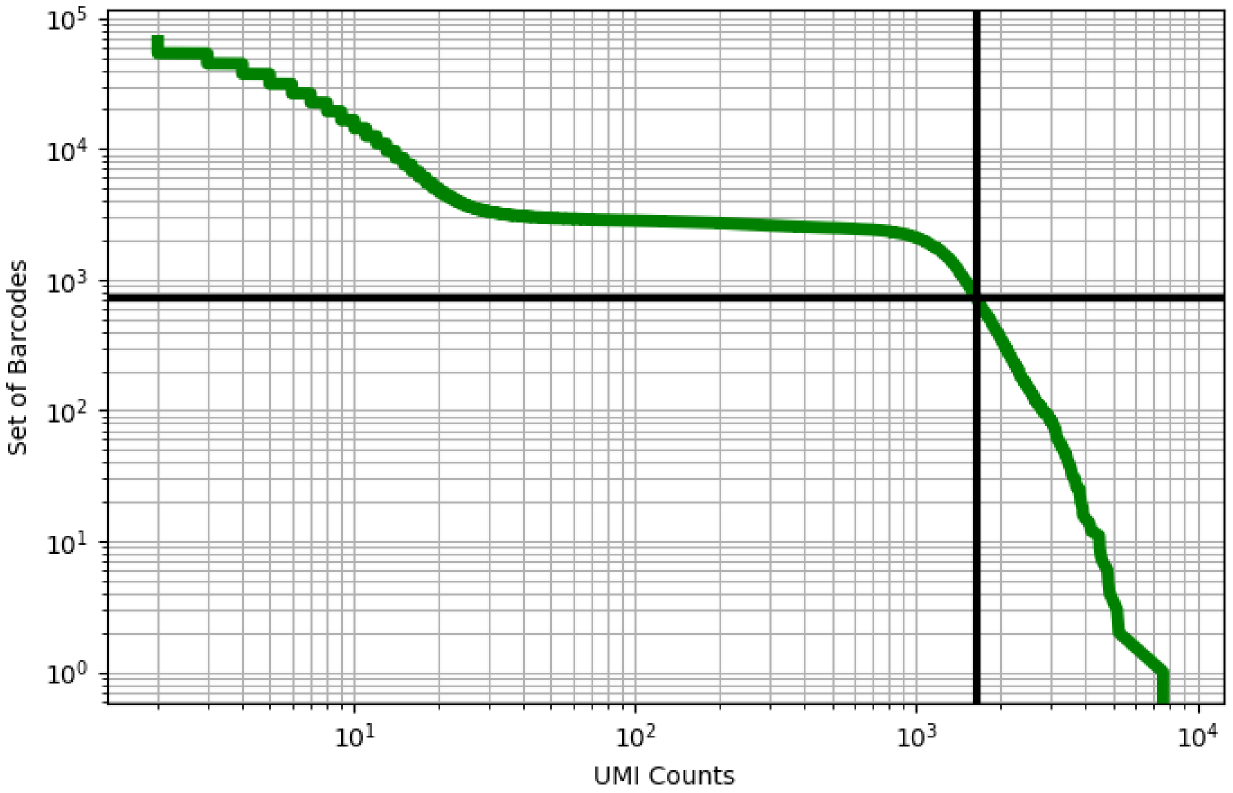


Figure 386: ovary/GSM3557969

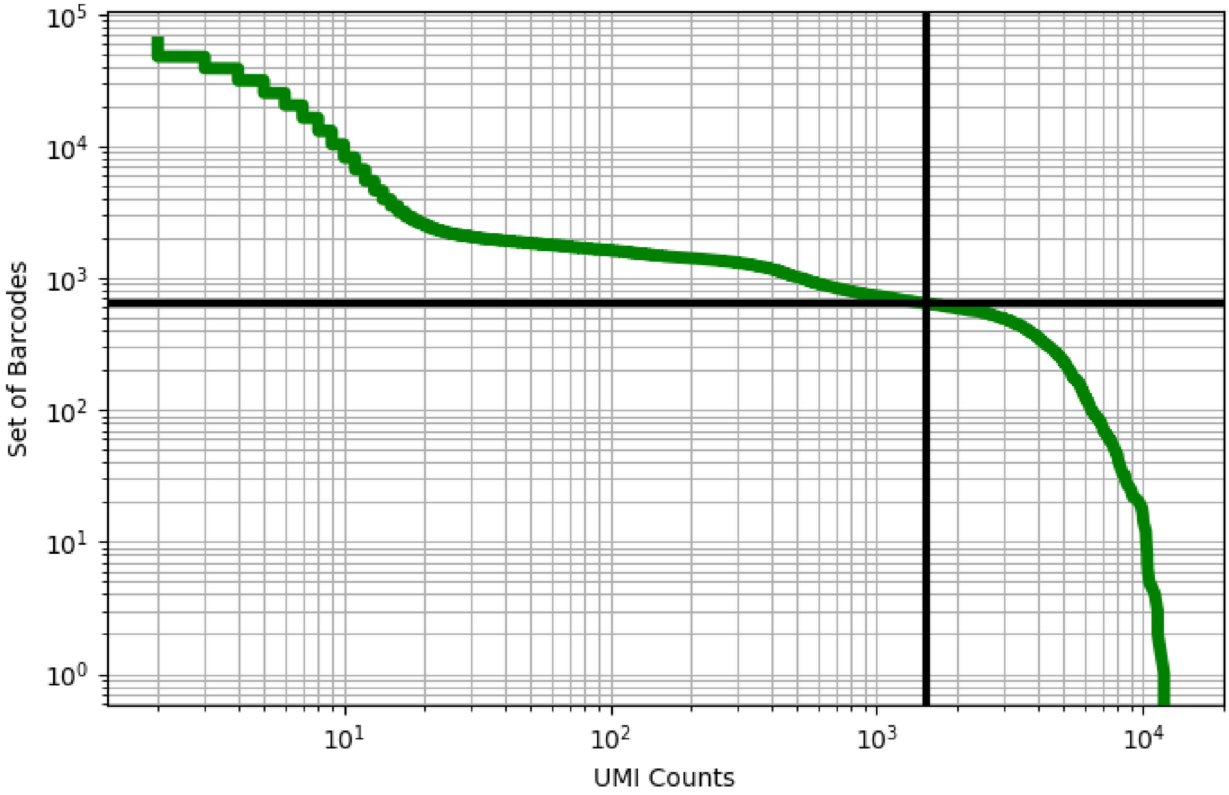


Figure 387: ovary/GSM3557970

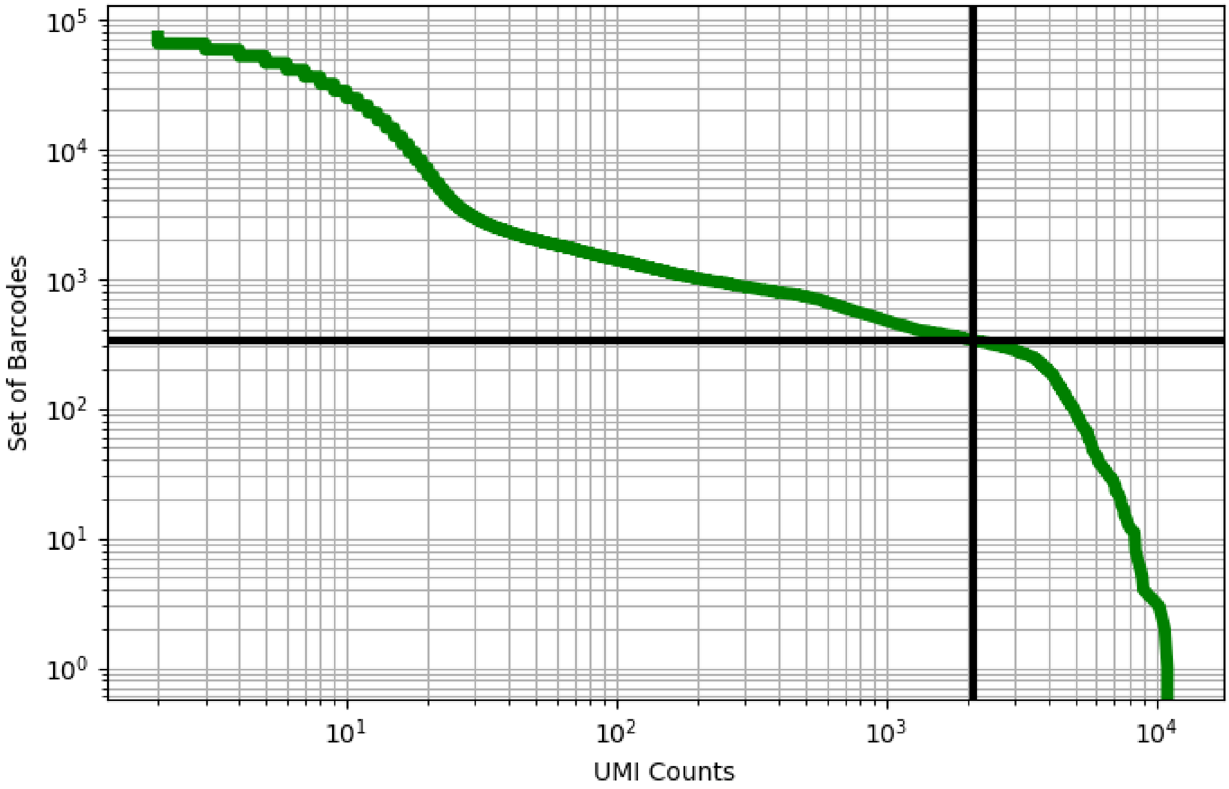


Figure 388: ovary/GSM3557971

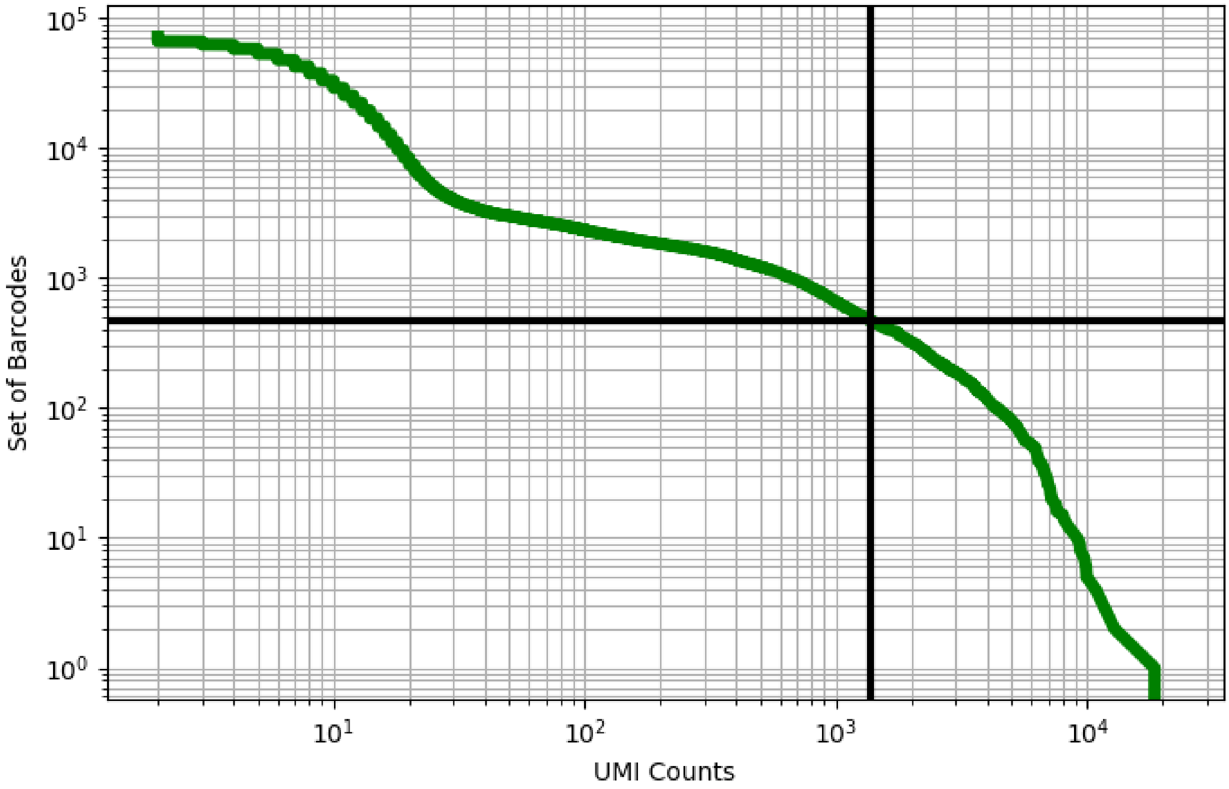


Figure 389: ovary/GSM3557972

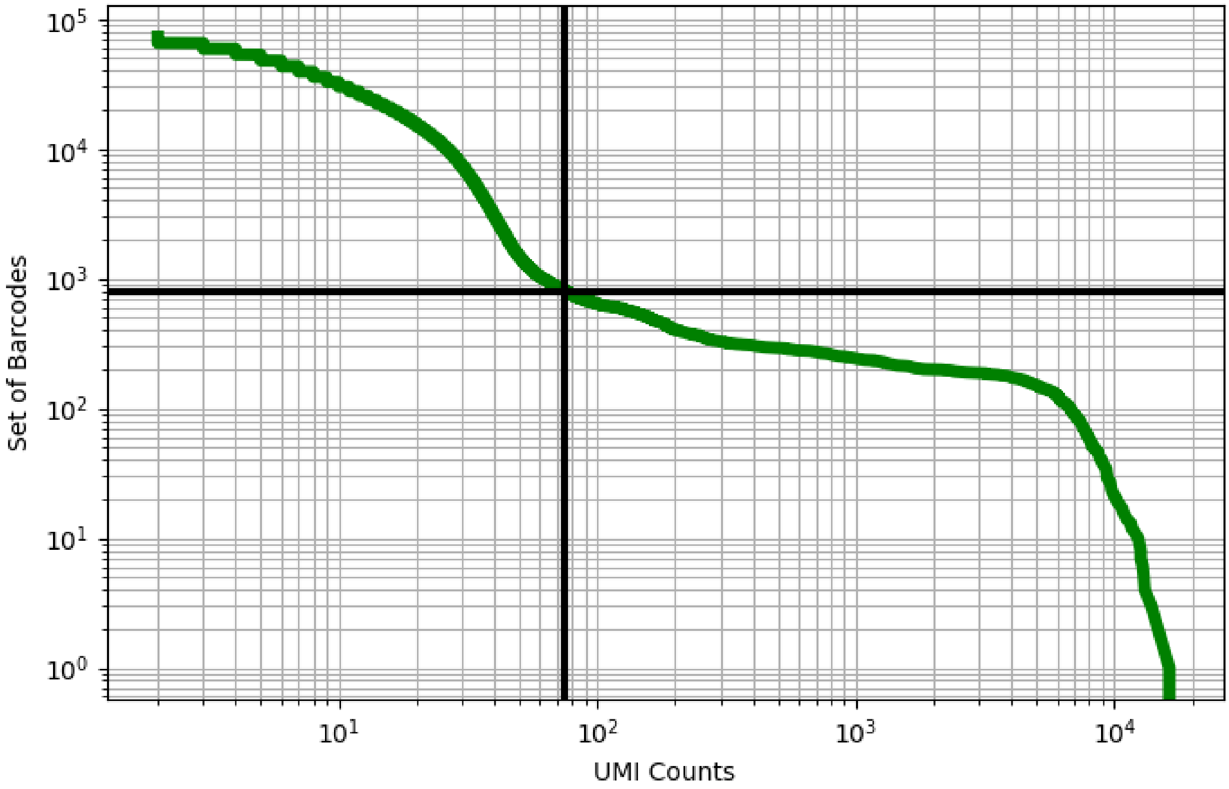


Figure 390: ovary/GSM3557973

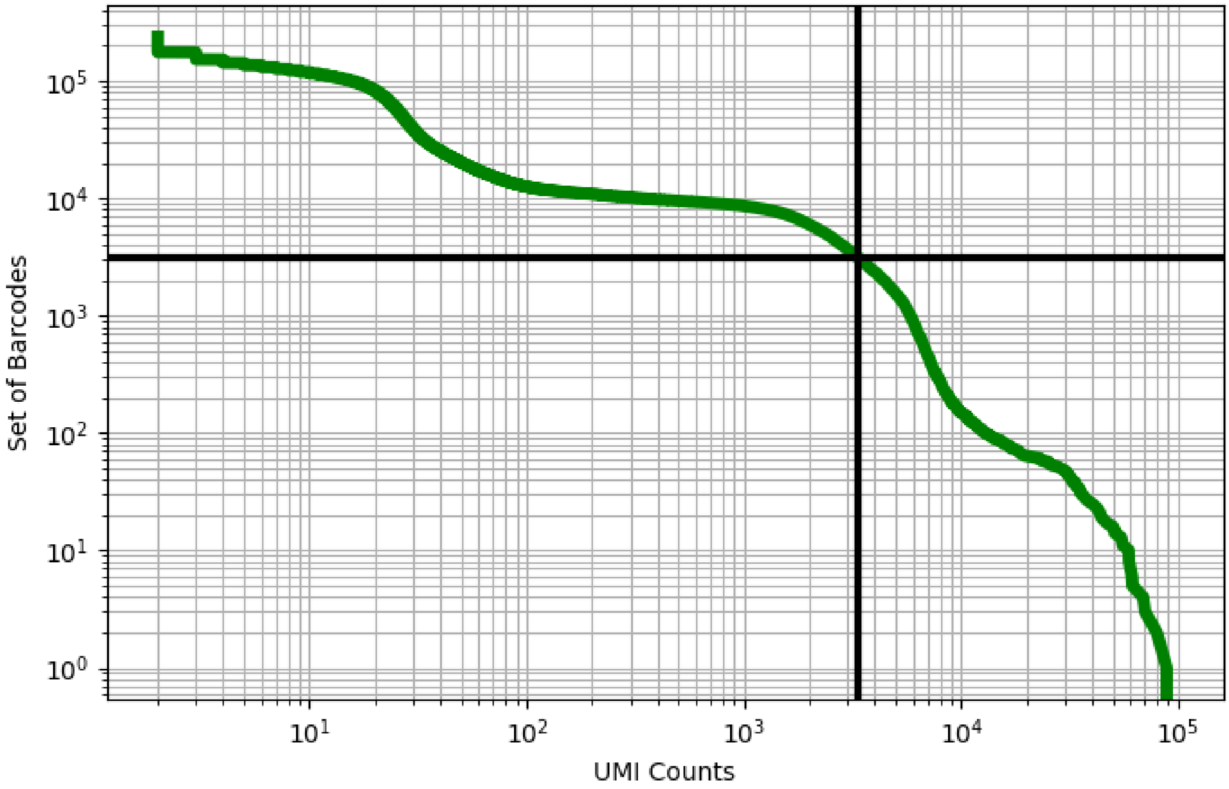


Figure 391: peritoneal/GSM3755687

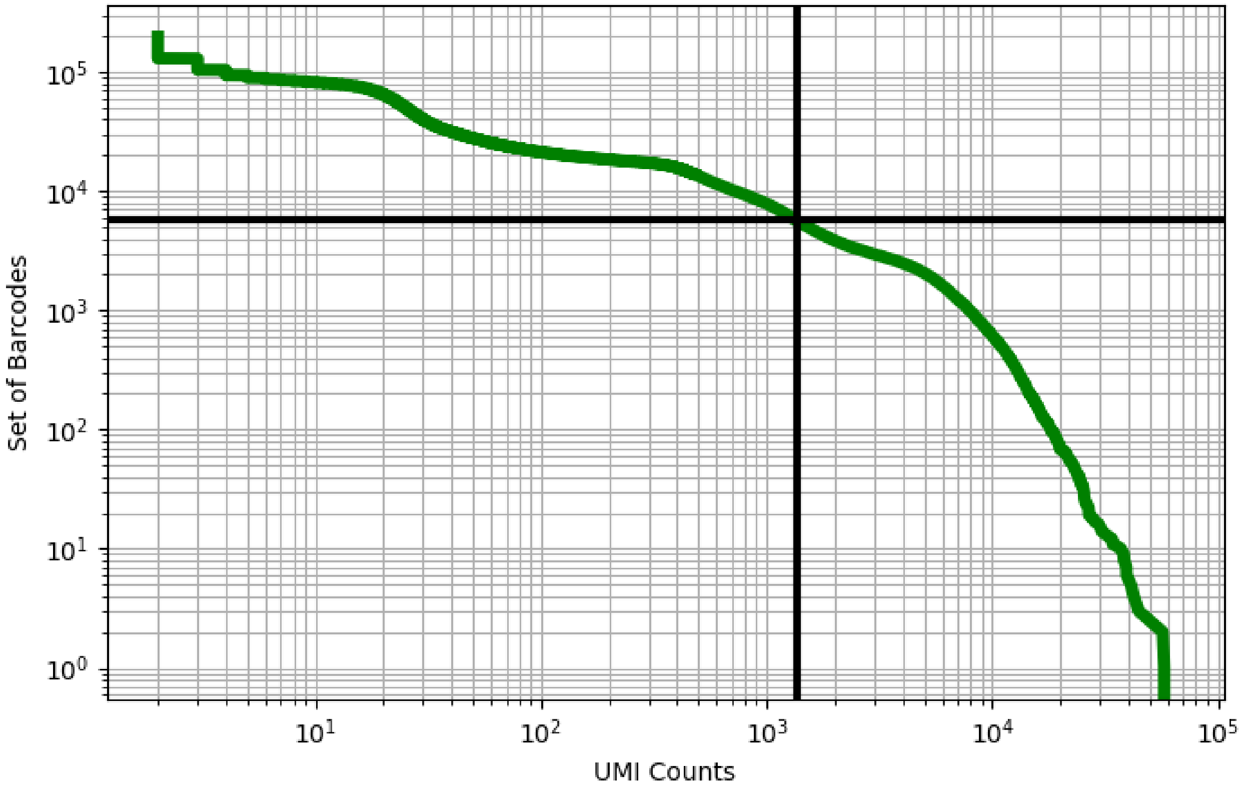


Figure 392: peritoneal/GSM3755690

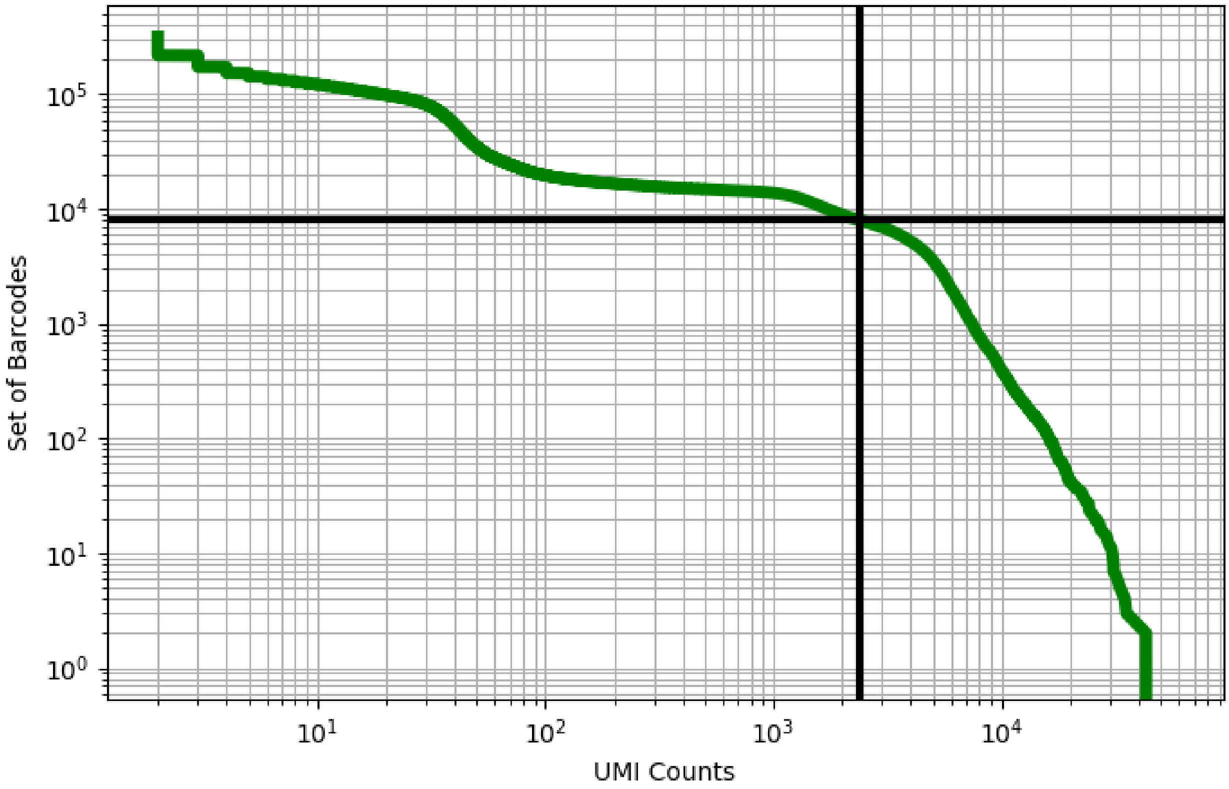


Figure 393: peritoneal/GSM3755691

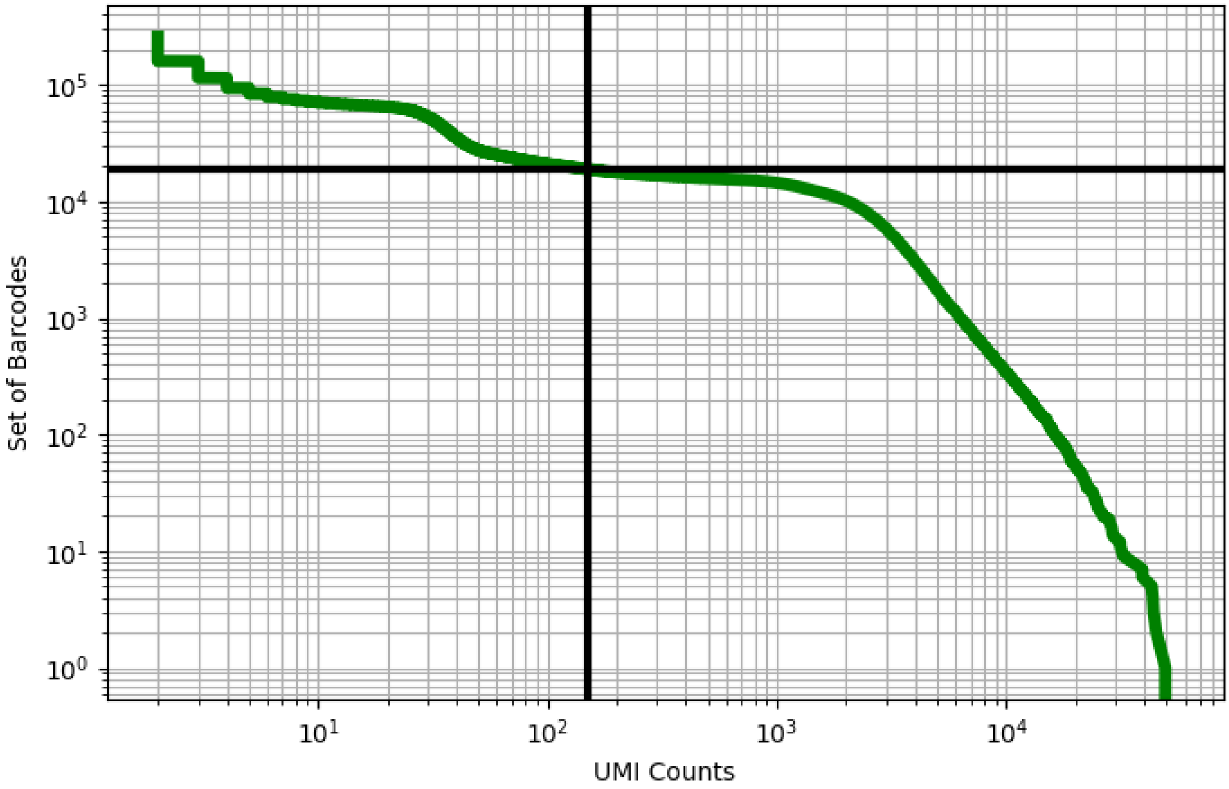


Figure 394: peritoneal/GSM3755692

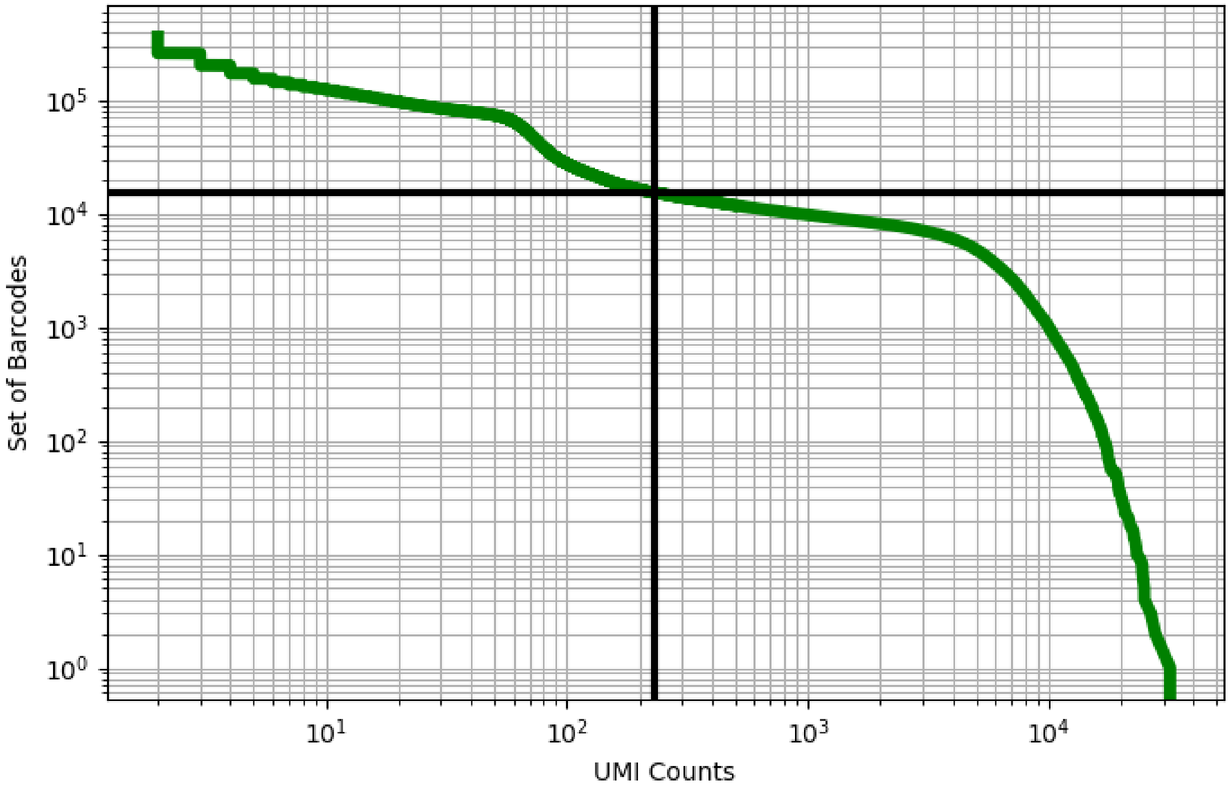


Figure 395: peritoneal/GSM3755693

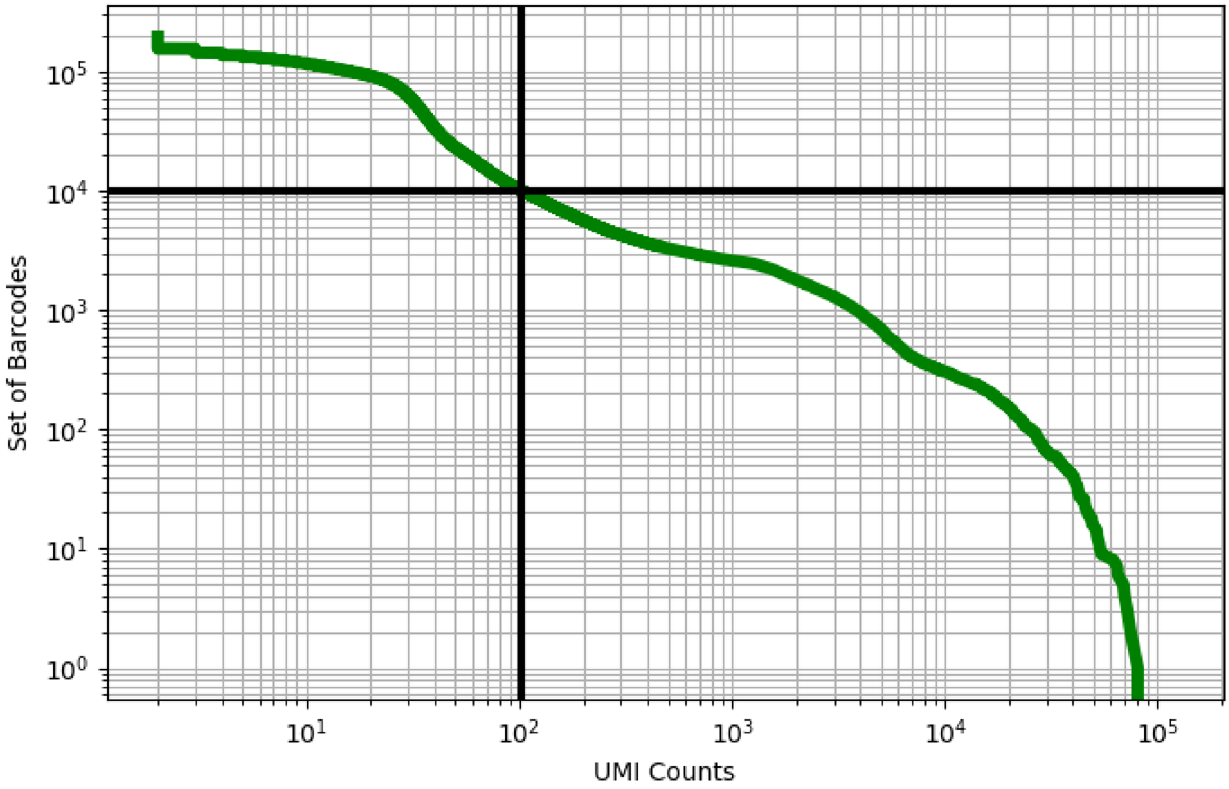


Figure 396: peritoneal/GSM3755696

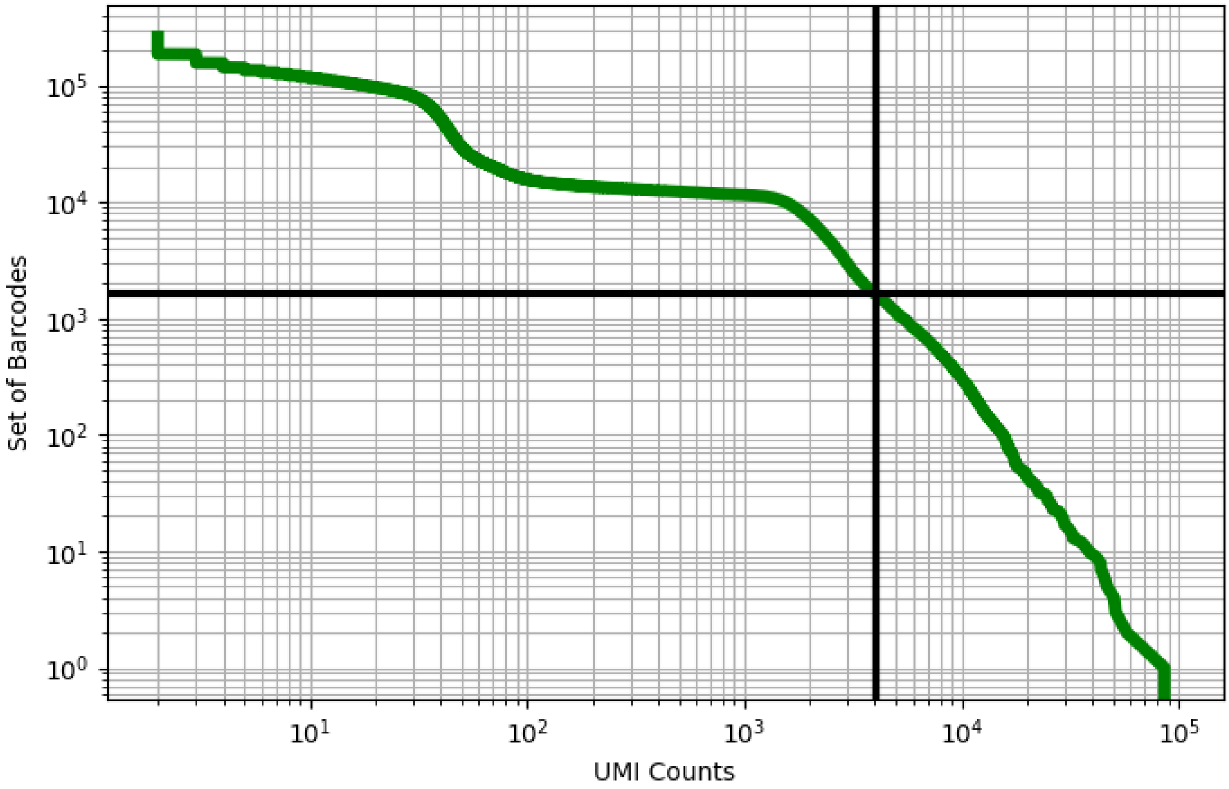


Figure 397: peritoneal/GSM3755697

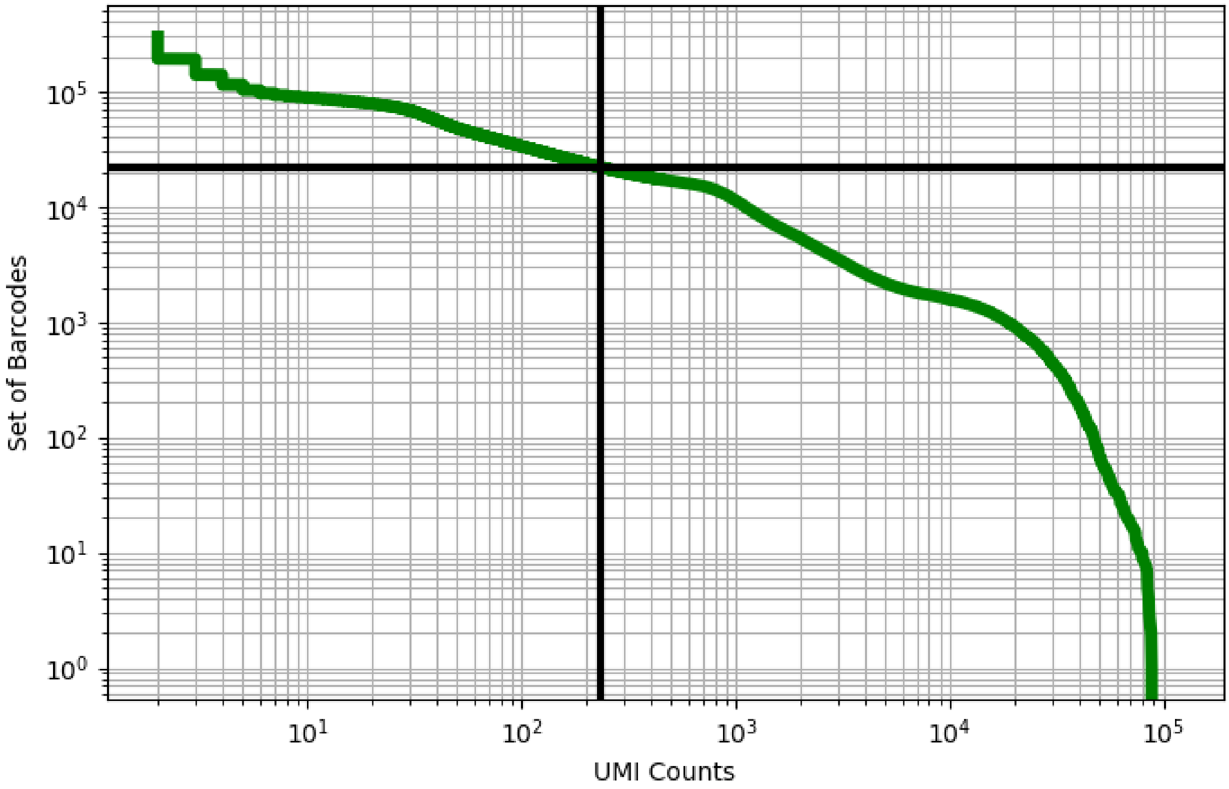


Figure 398: peritoneal/GSM3755698

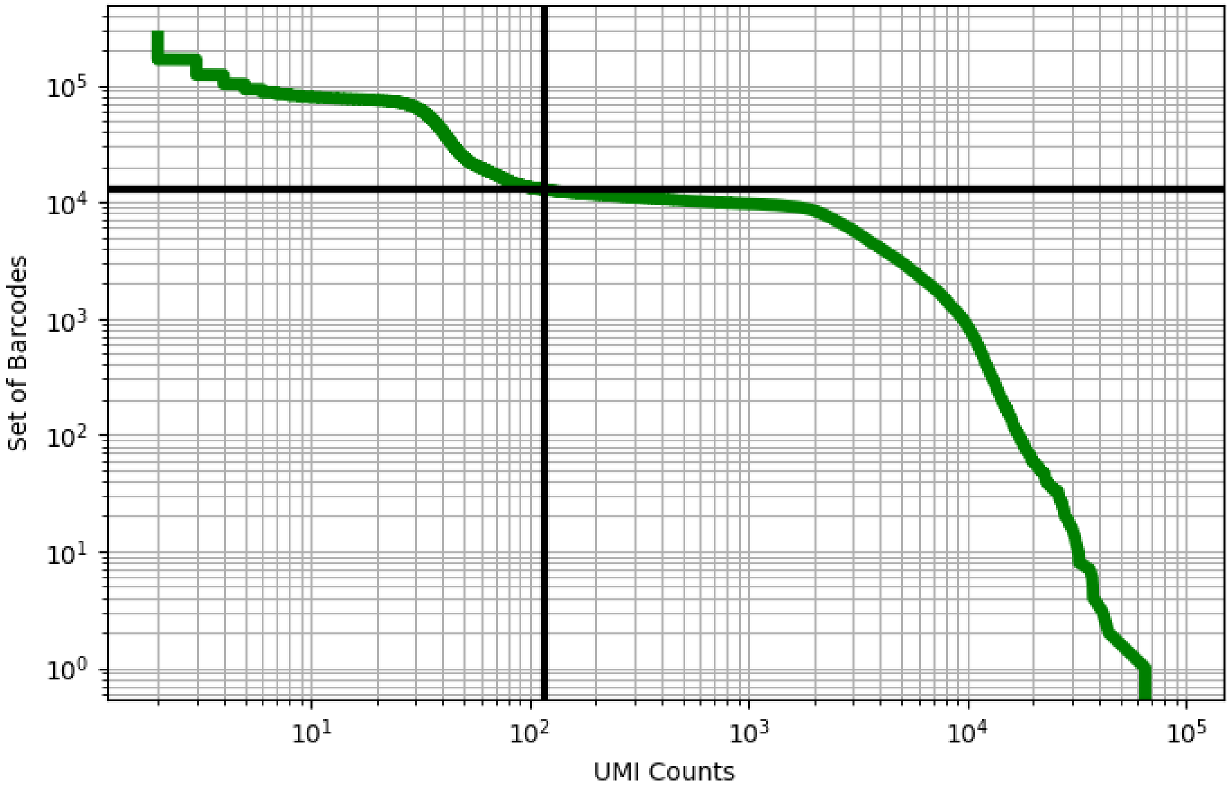


Figure 399: peritoneal/GSM3755699

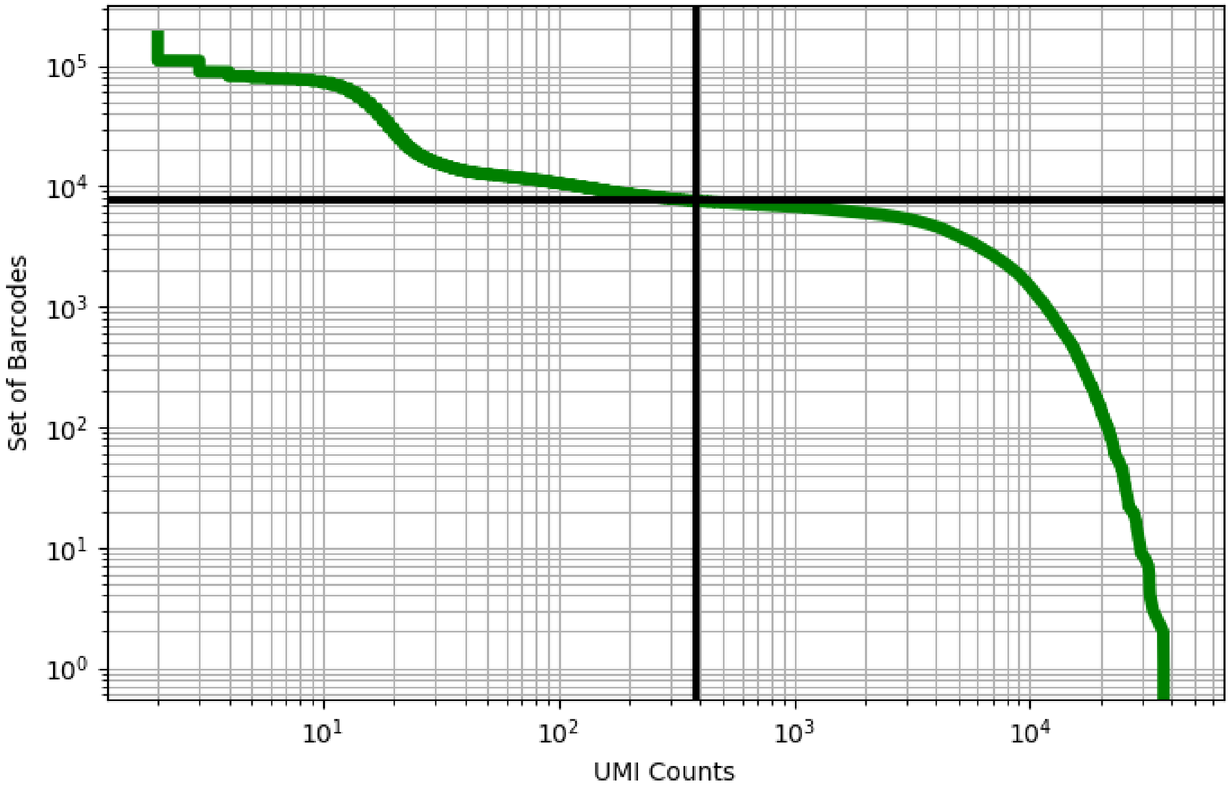


Figure 400: placenta/ERX2756723

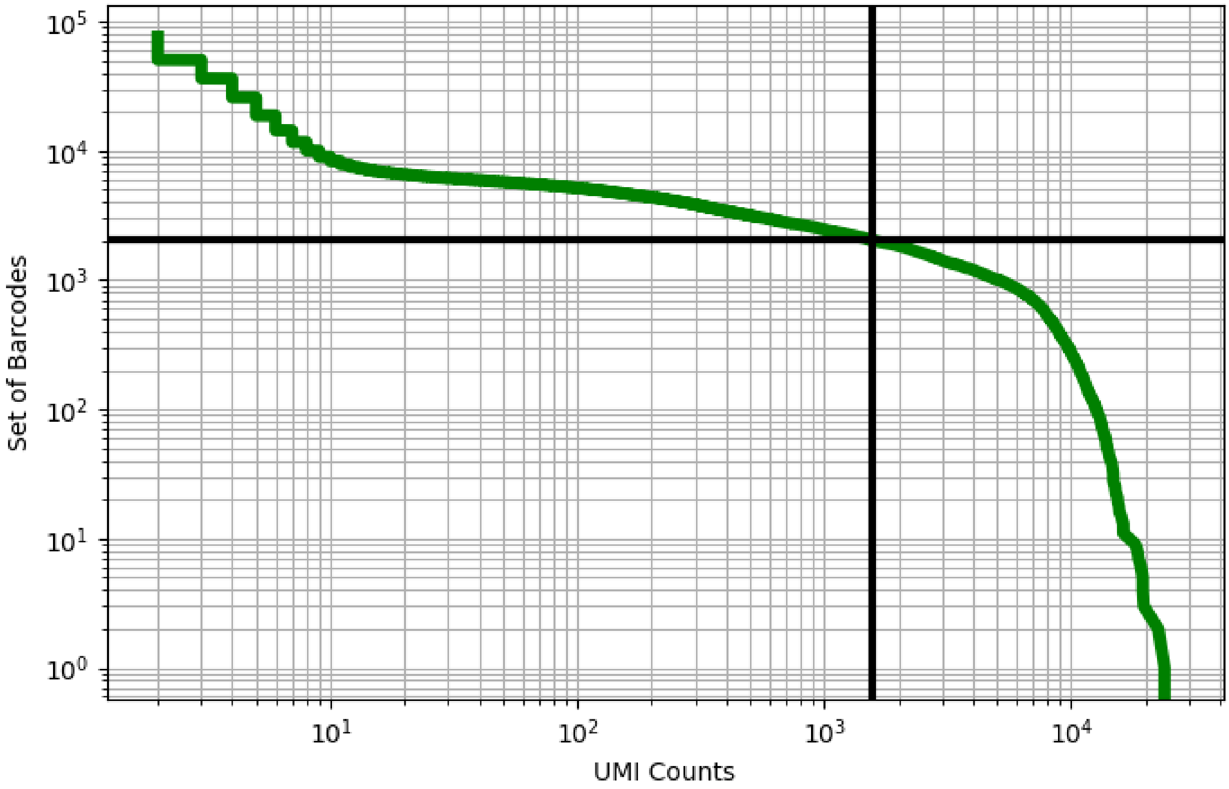


Figure 401: placenta/ERX2756726

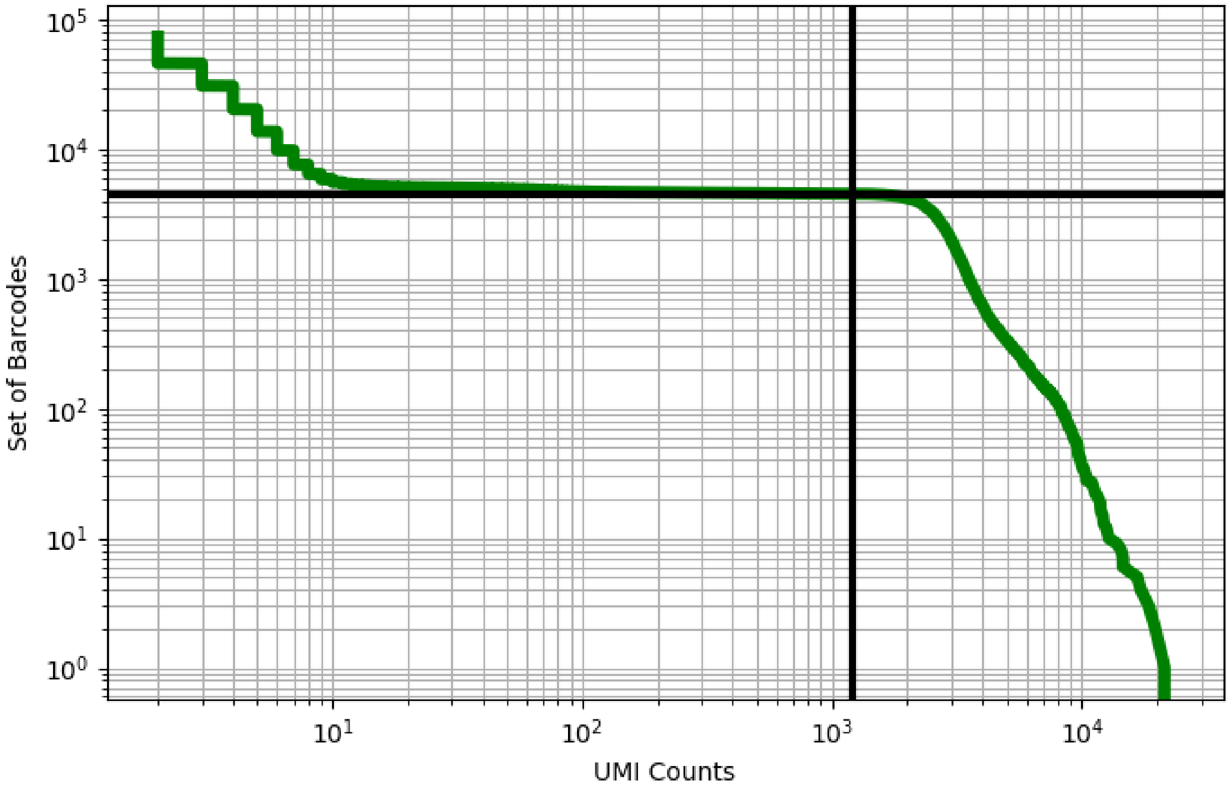


Figure 402: placenta/ERX2756728

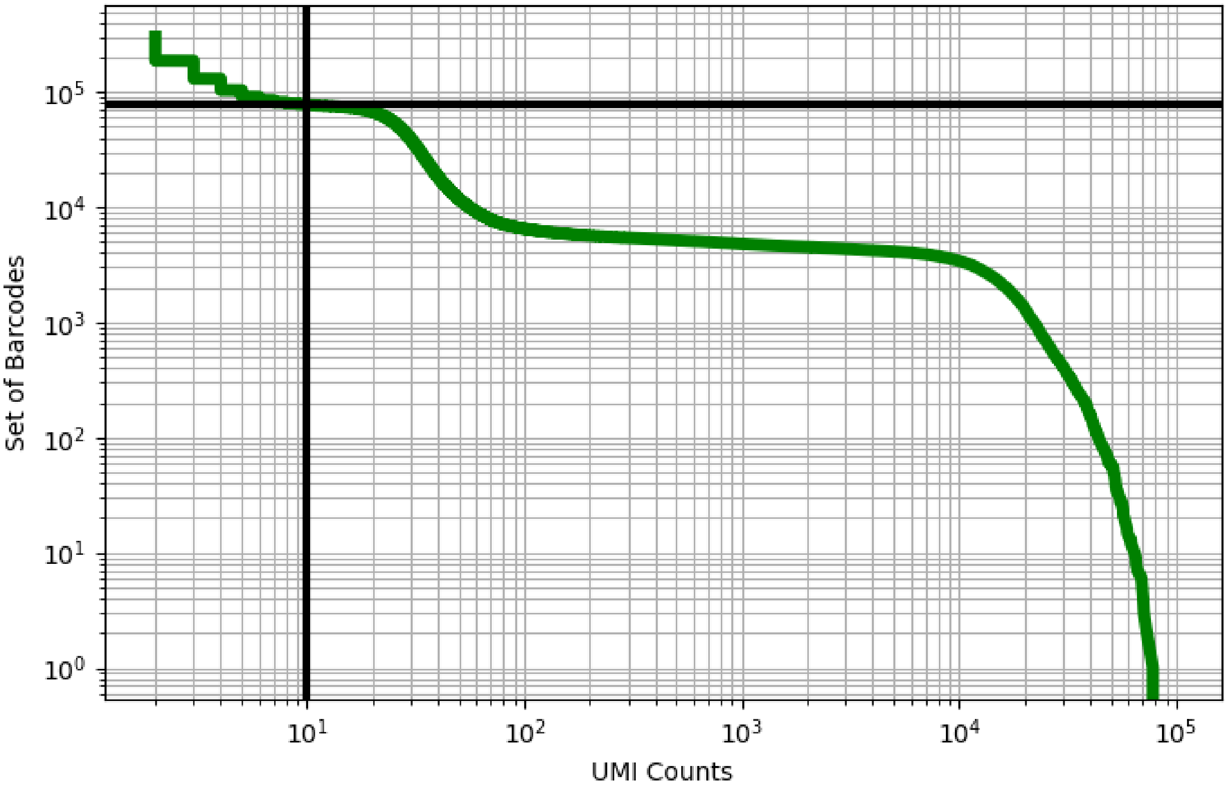


Figure 403: placenta/ERX2756731

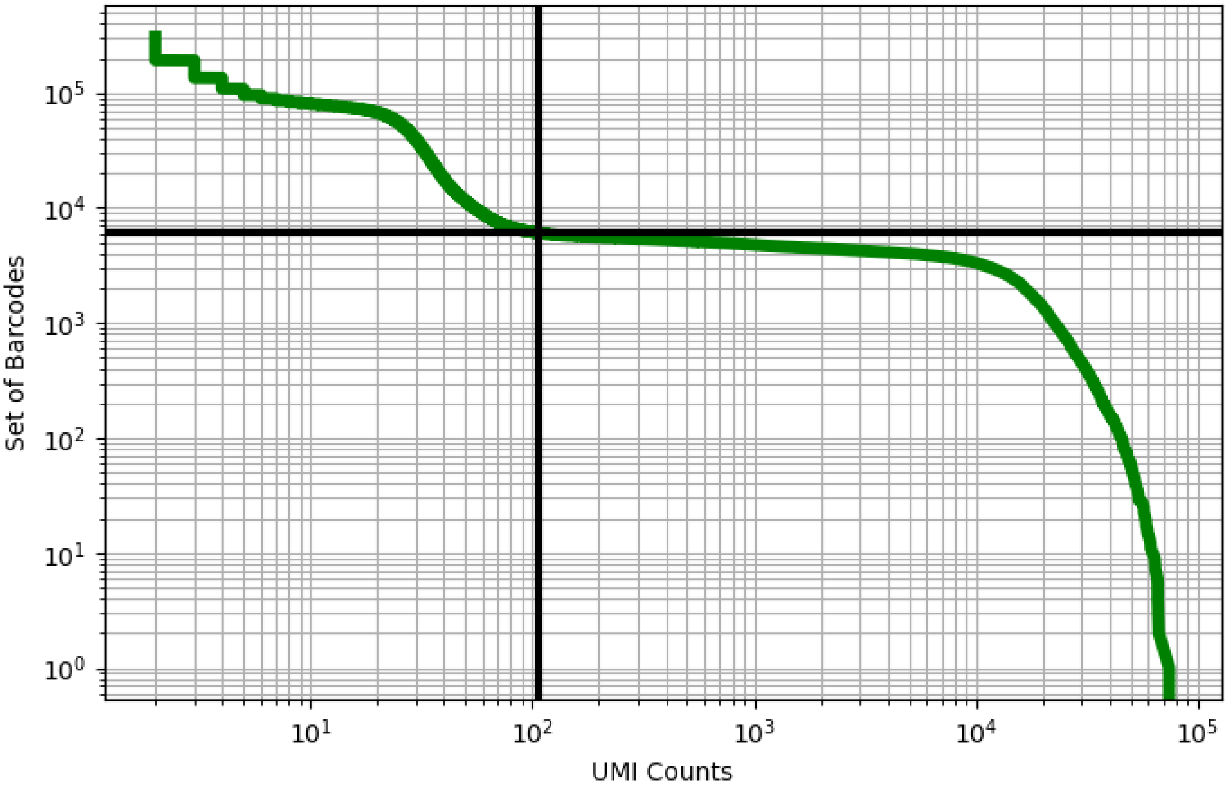


Figure 404: placenta/ERX2756732

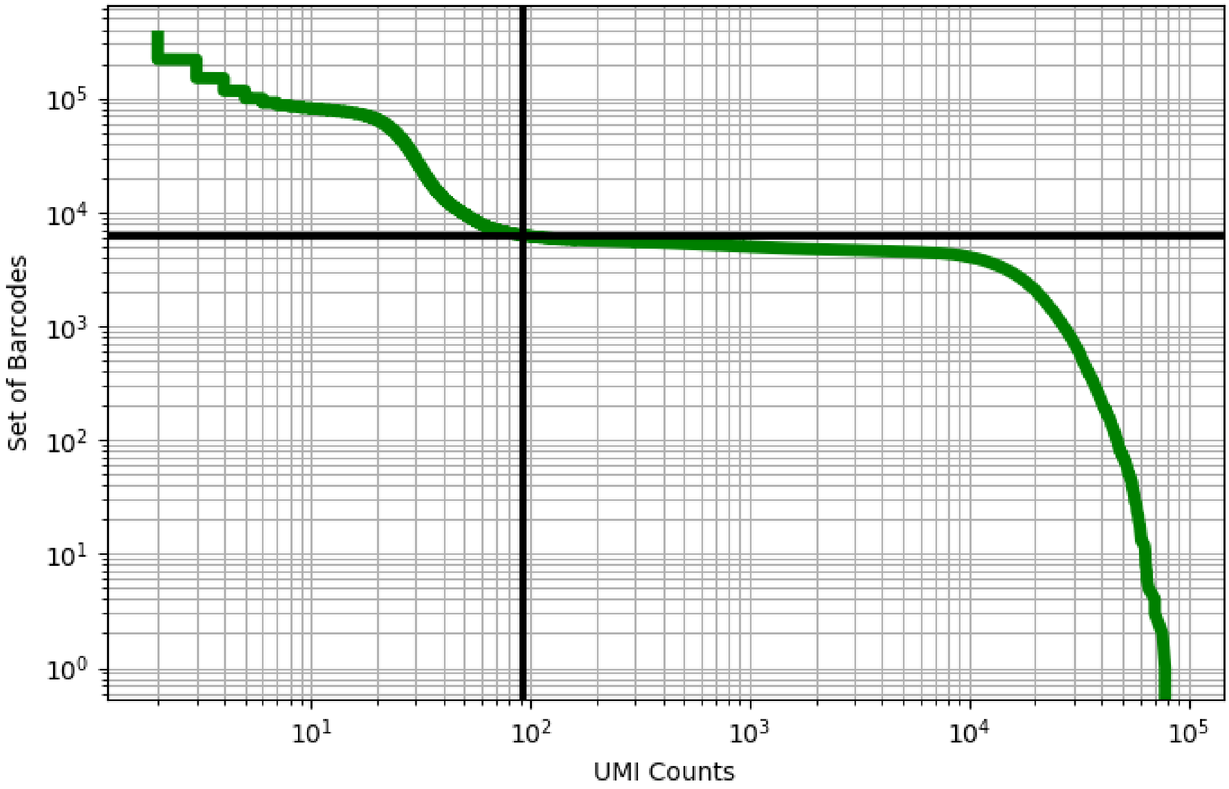


Figure 405: placenta/ERX2756733

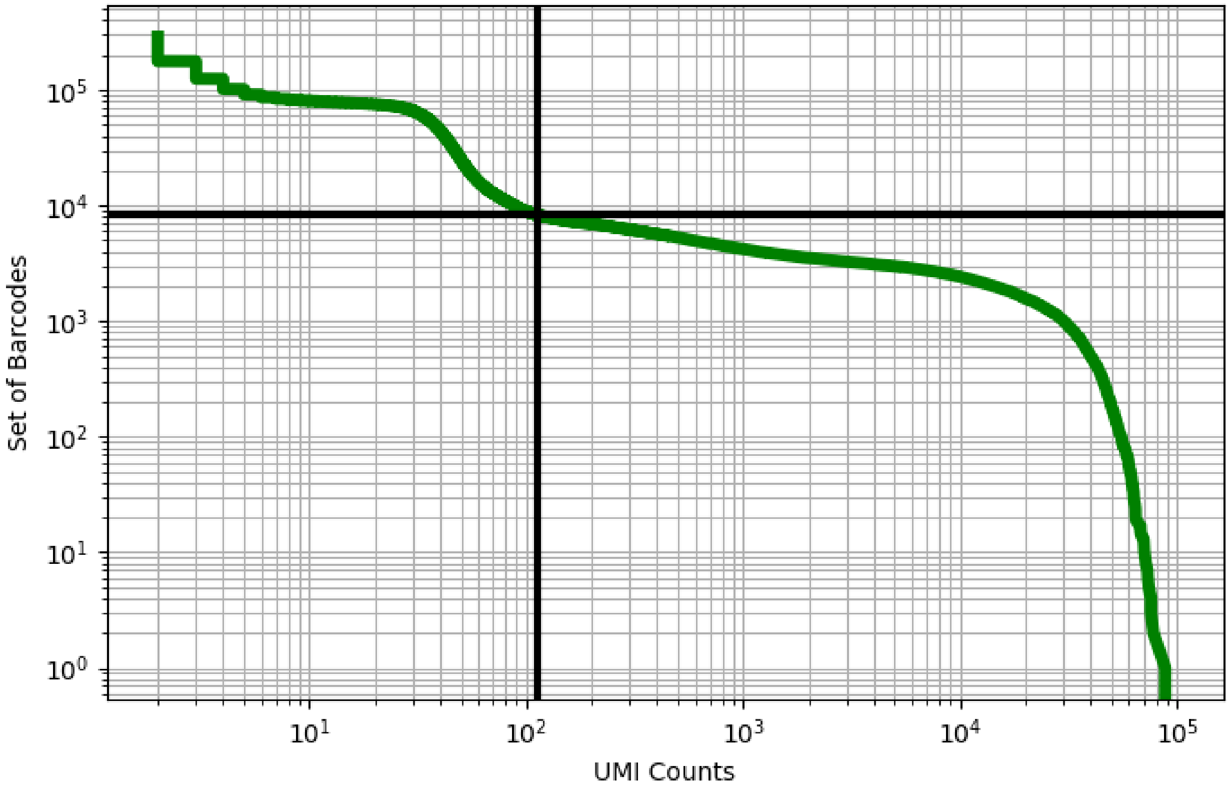


Figure 406: placenta/ERX2756735

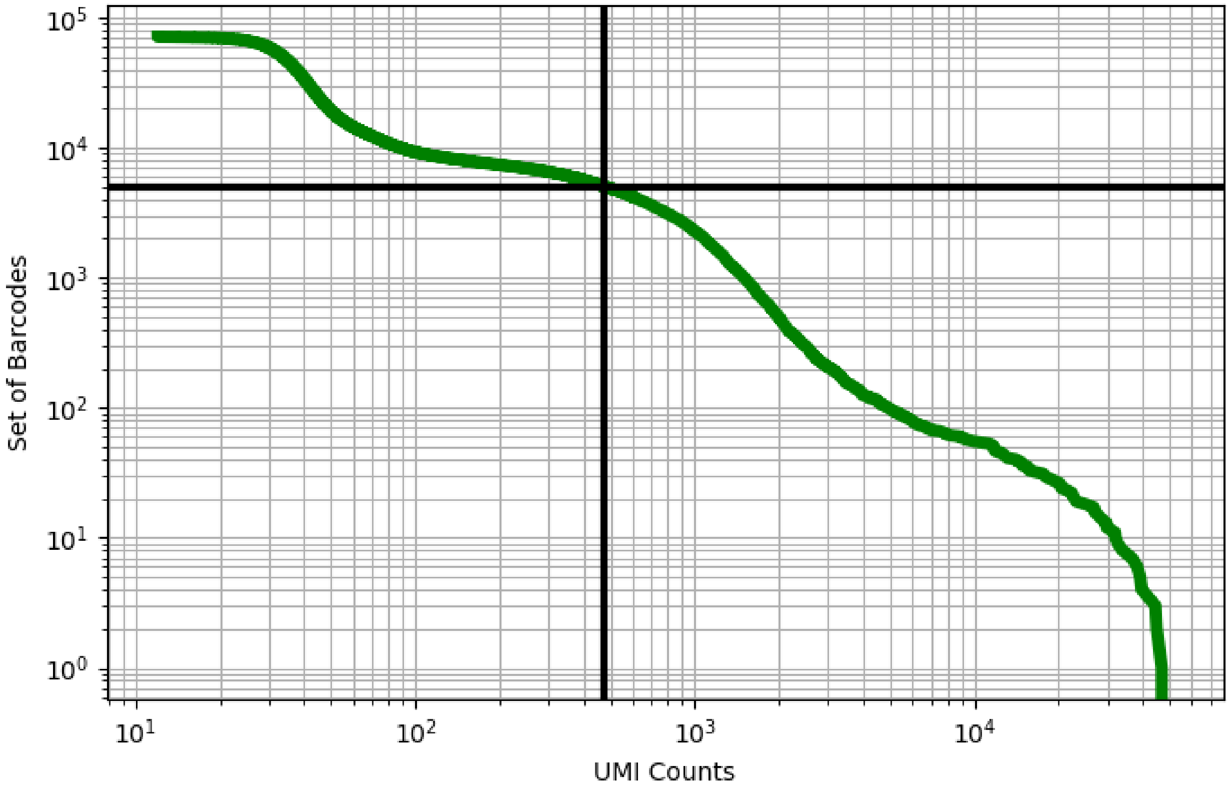


Figure 407: placenta/ERX2756736

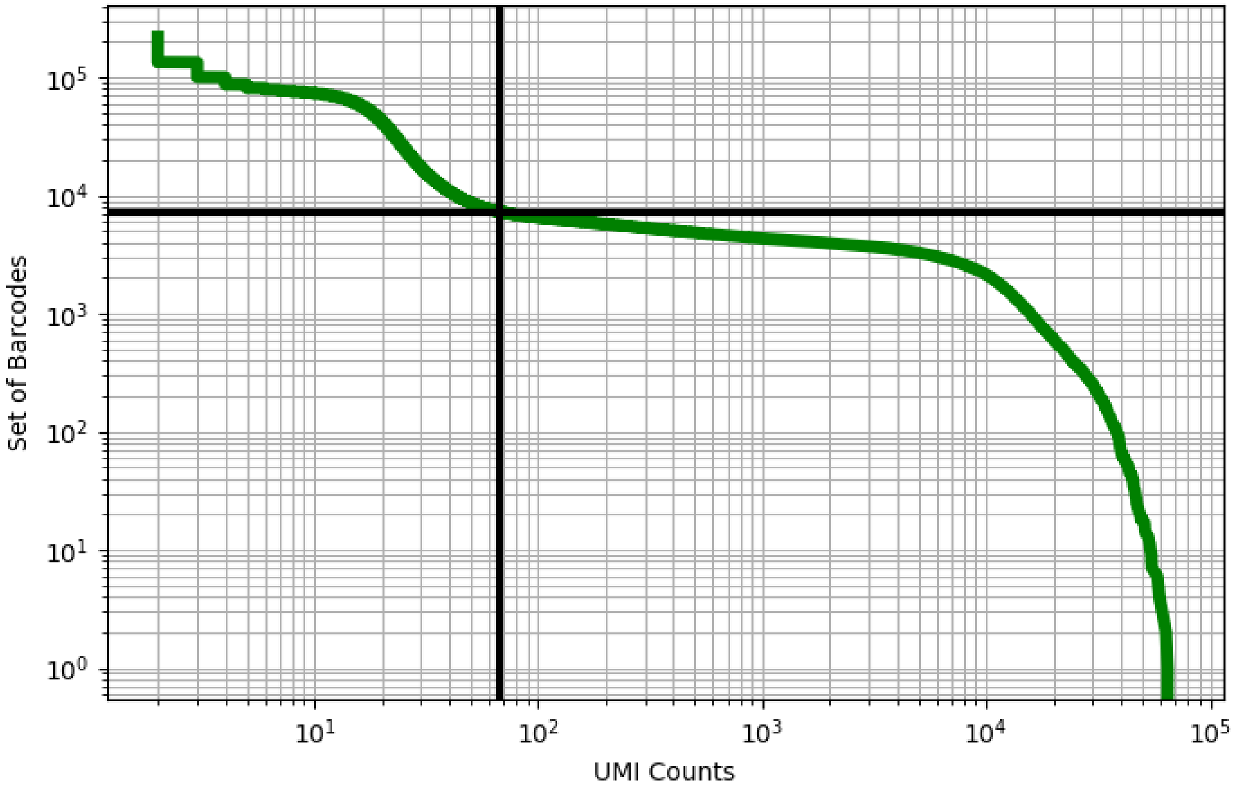


Figure 408: placenta/ERX2756740

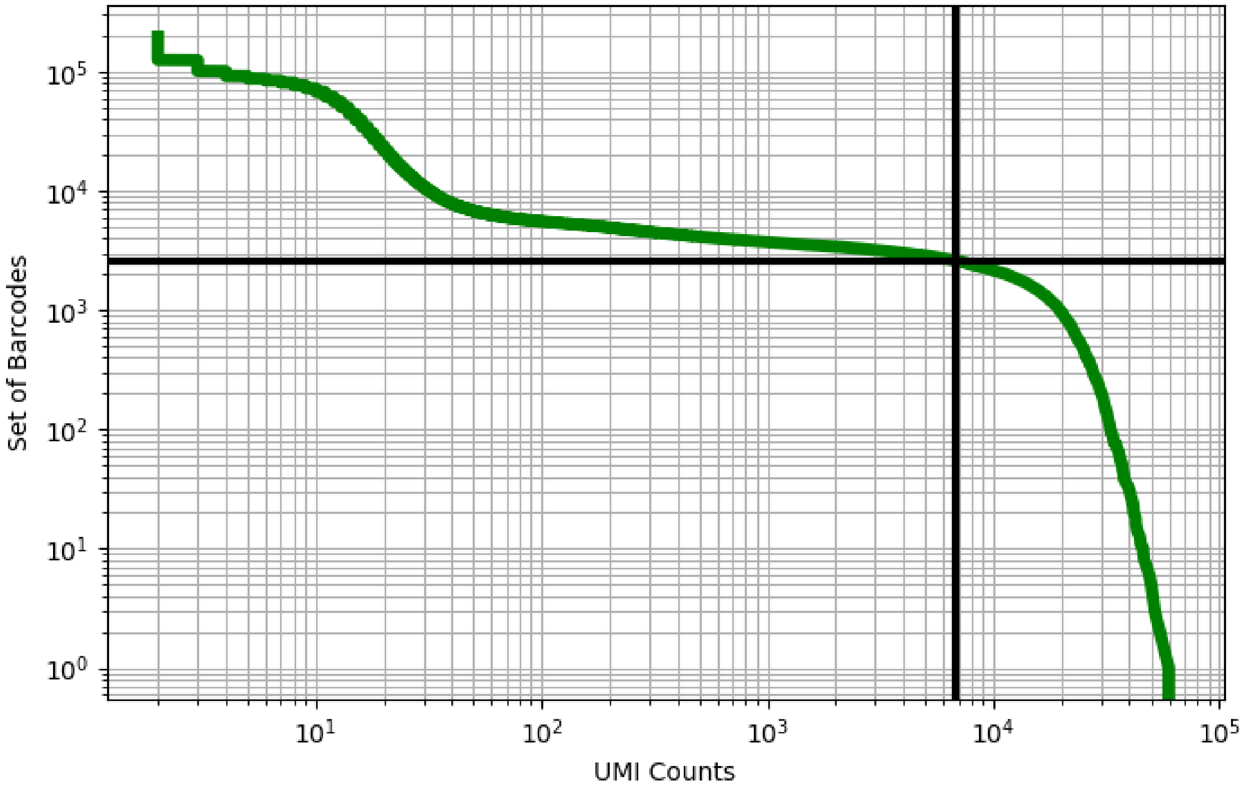


Figure 409: placenta/ERX2756741

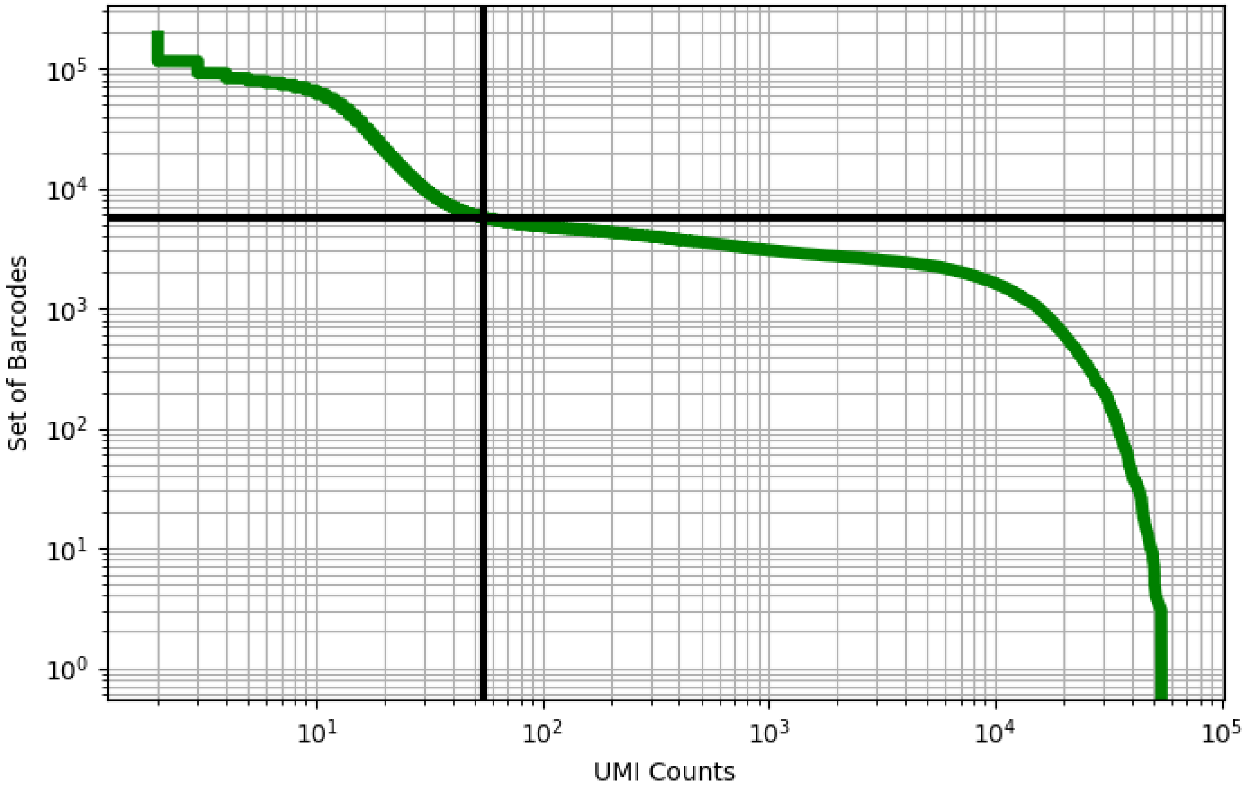


Figure 410: placenta/ERX2756742

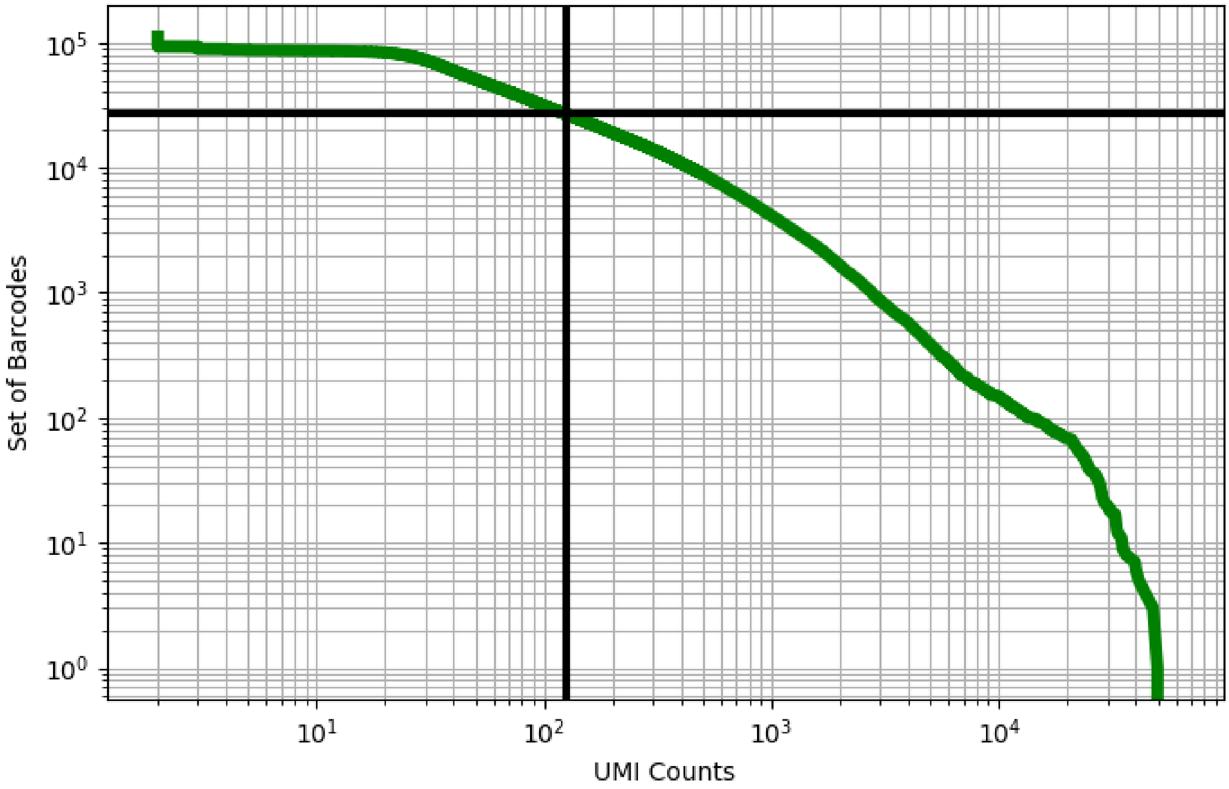


Figure 411: placenta/SRX4733402

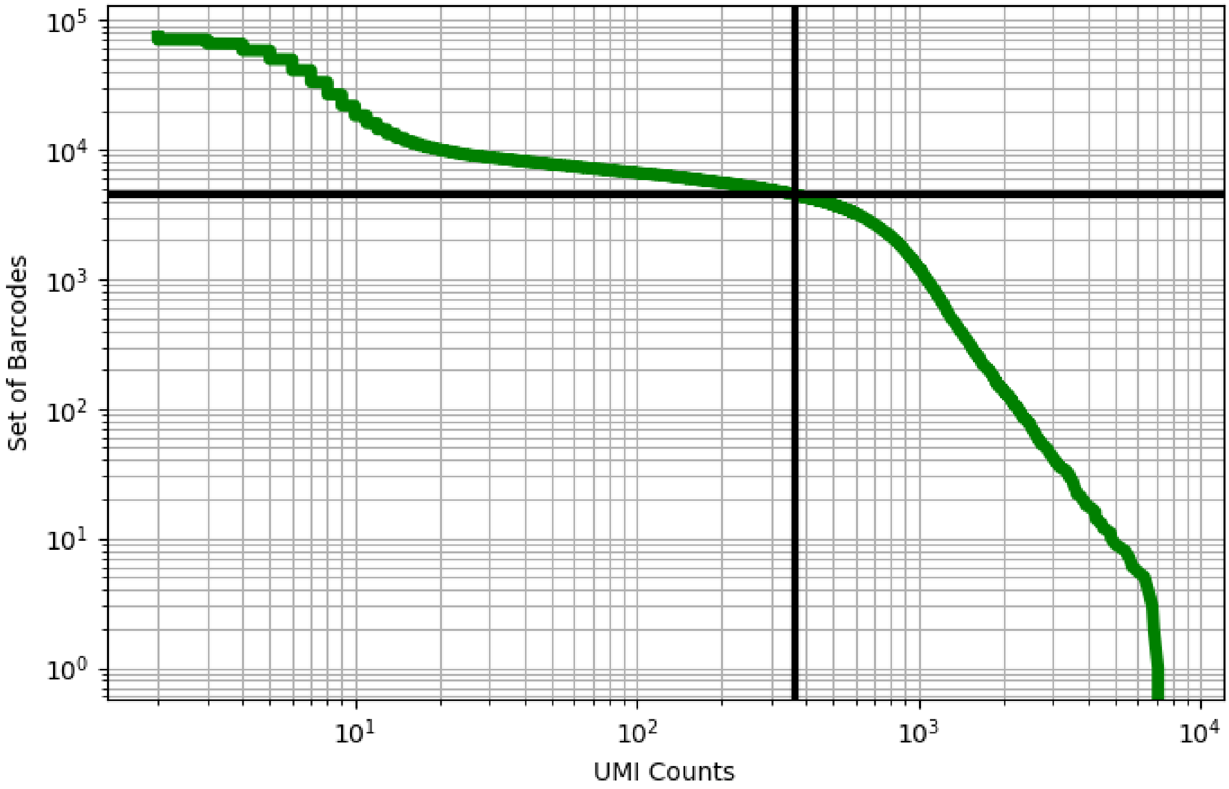


Figure 412: placenta/SRX4733403

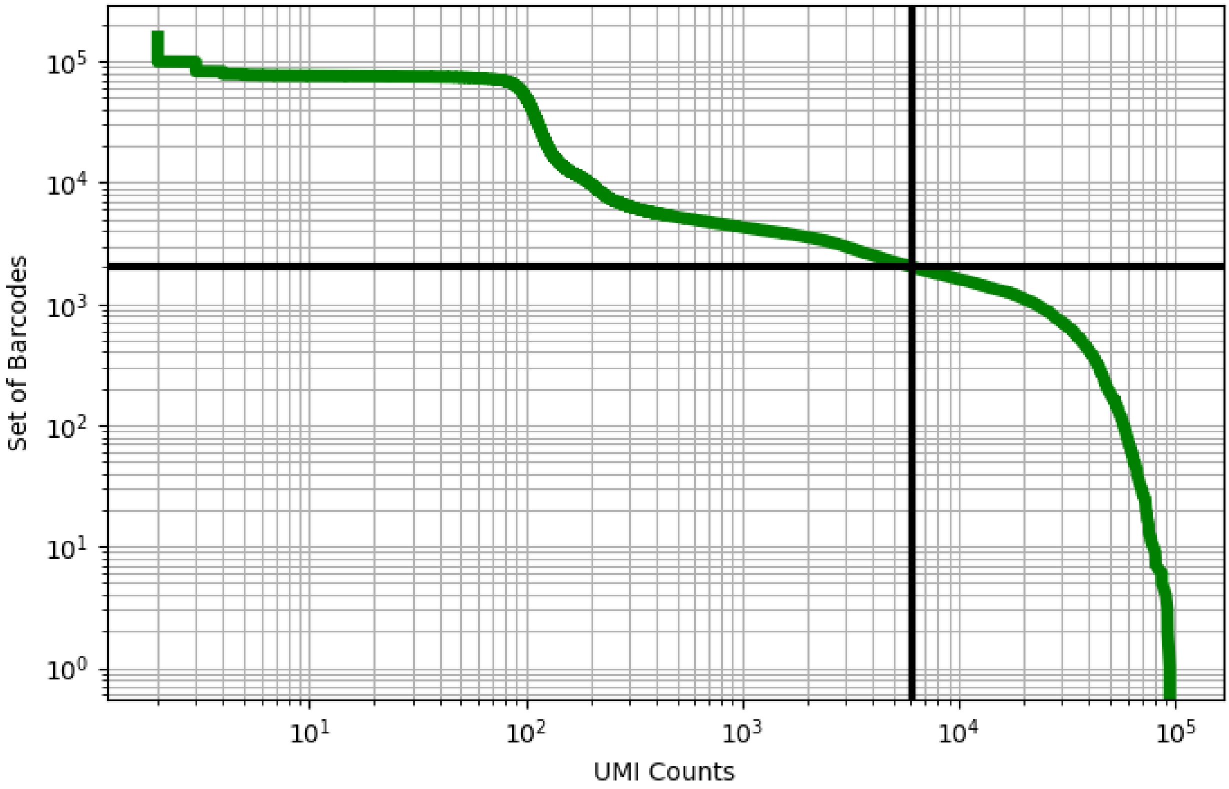


Figure 413: placenta/SRX4733411

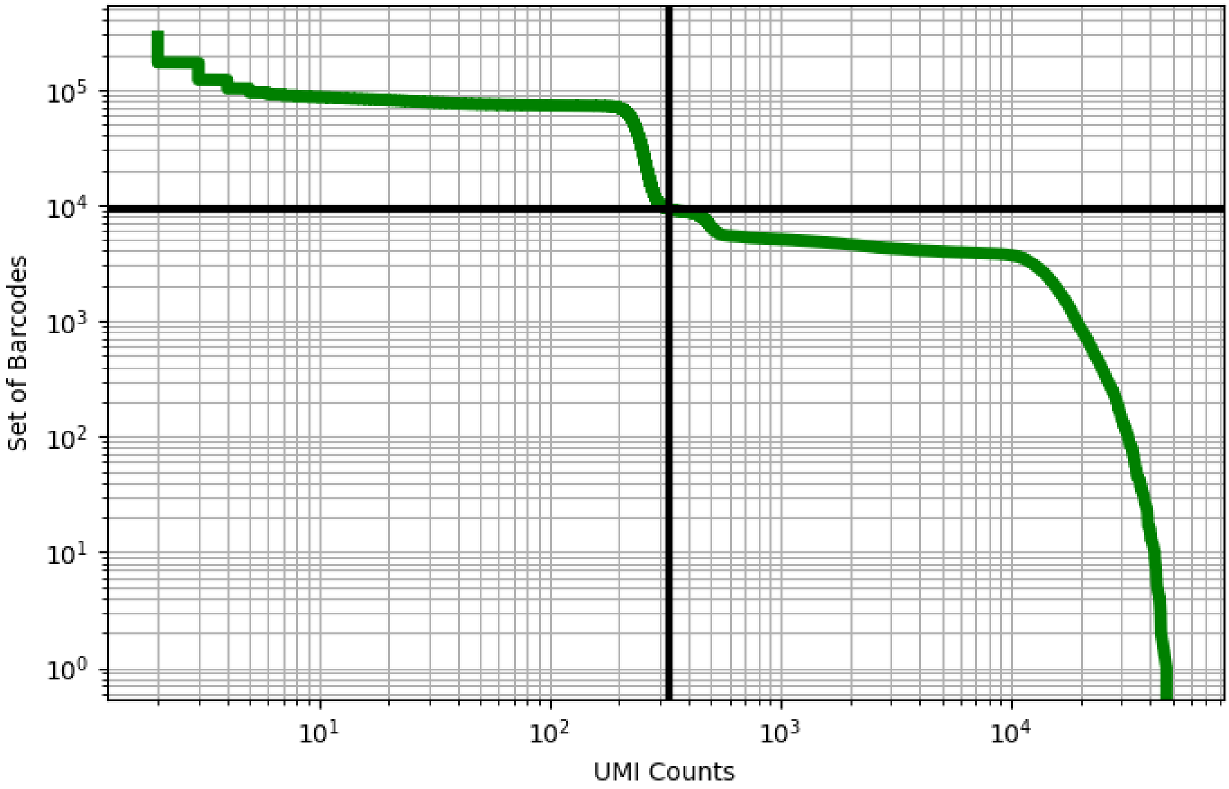


Figure 414: prostate/GSM3735993

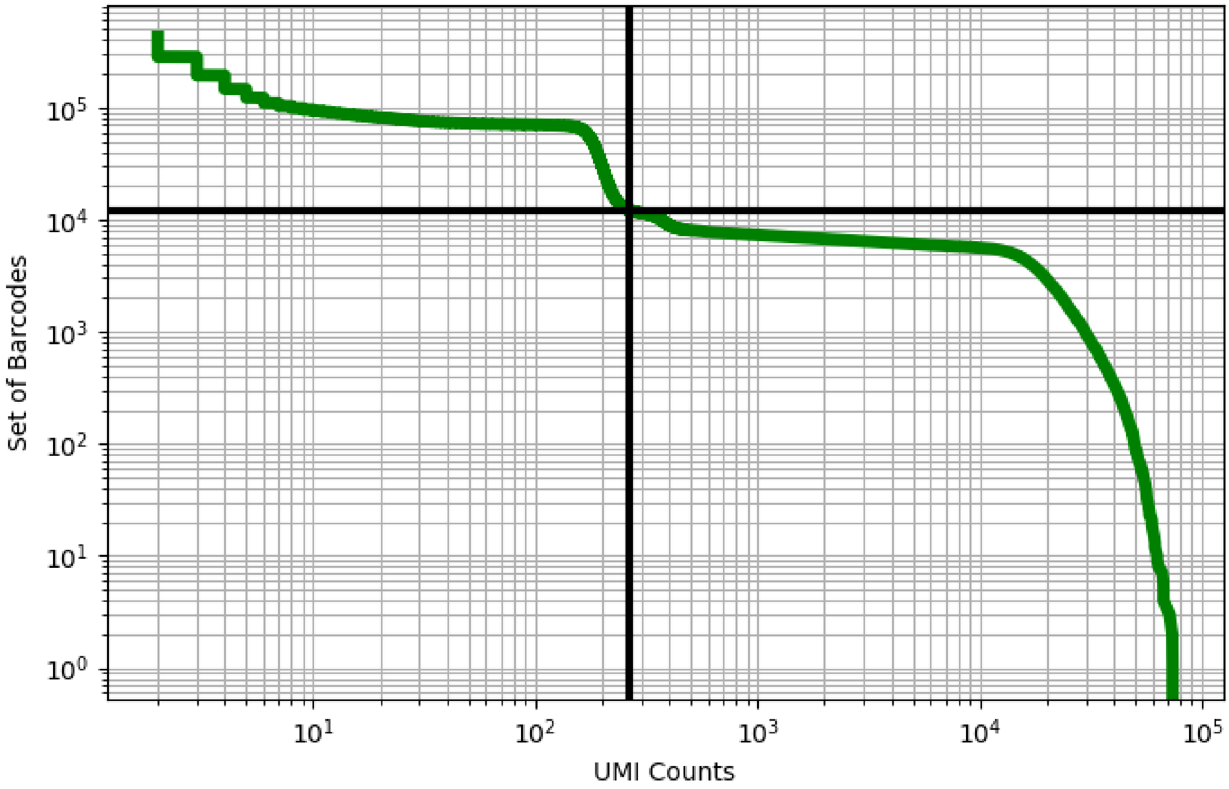


Figure 415: prostate/GSM3735994

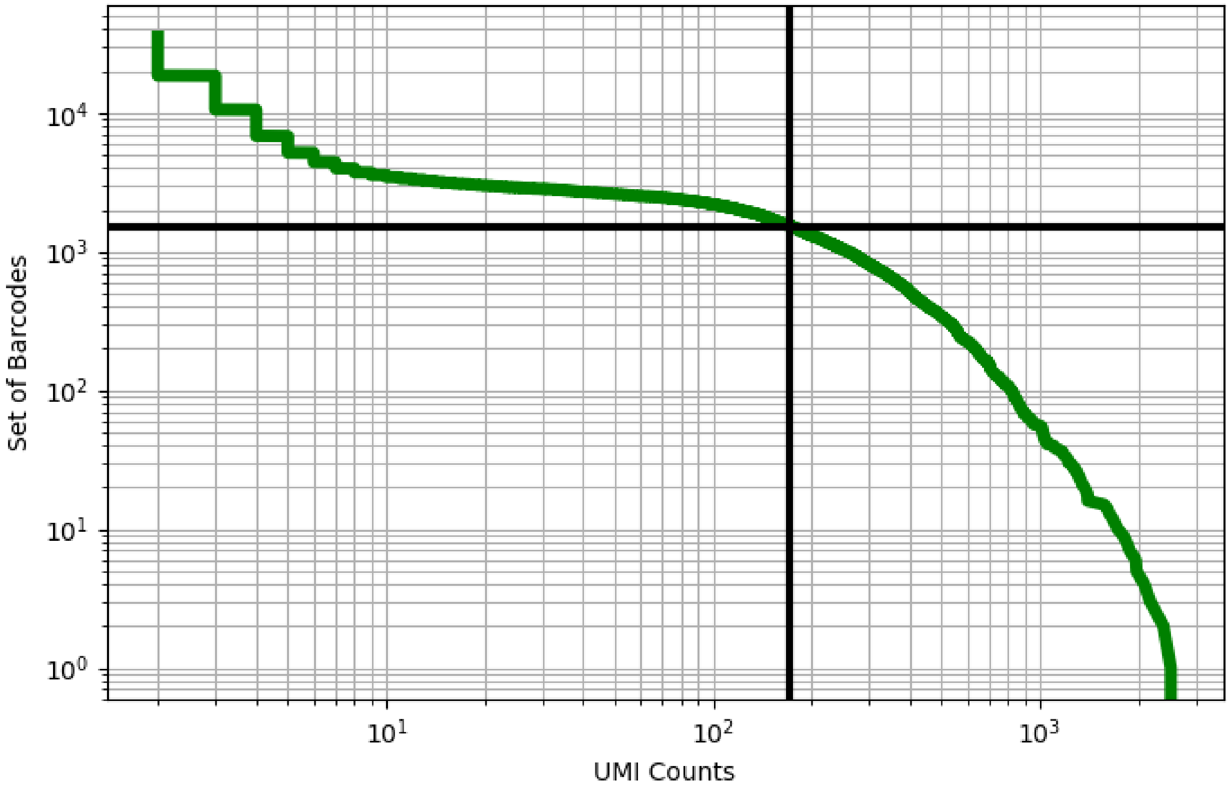


Figure 416: rectum/GSM3576397

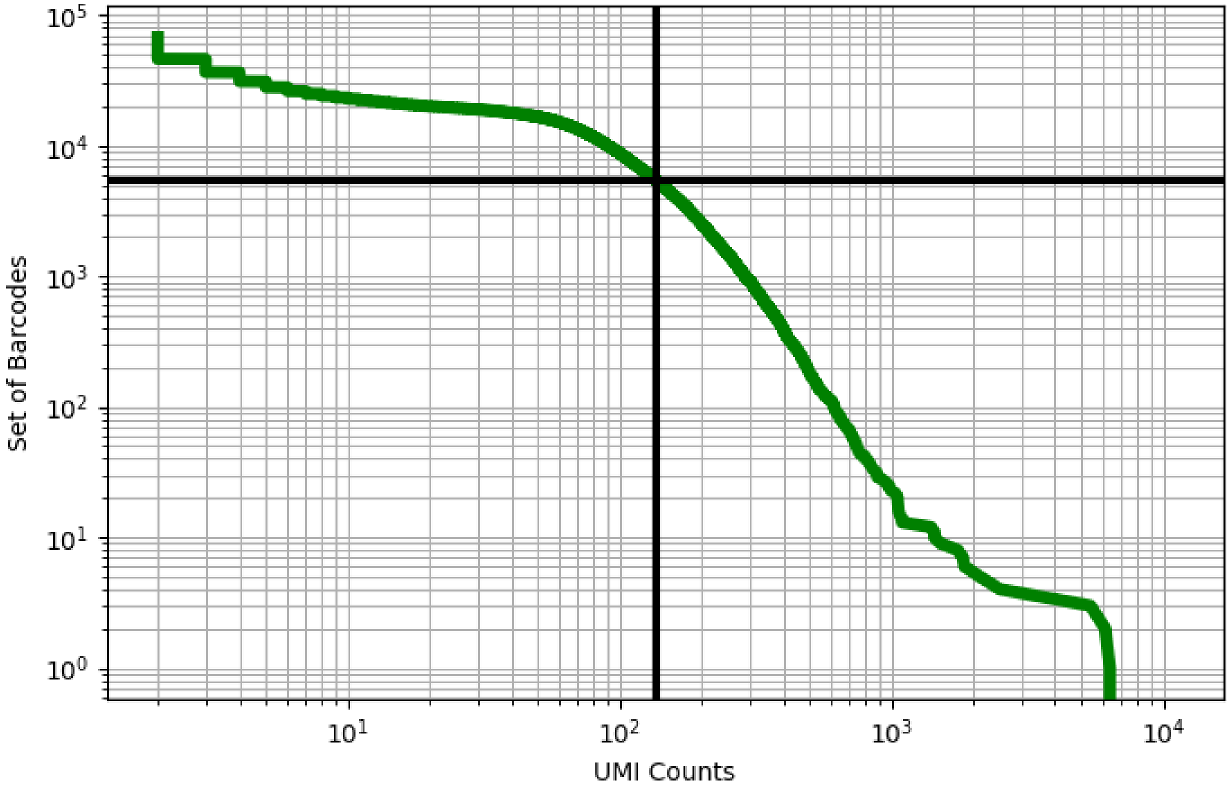


Figure 417: rectum/GSM3576398

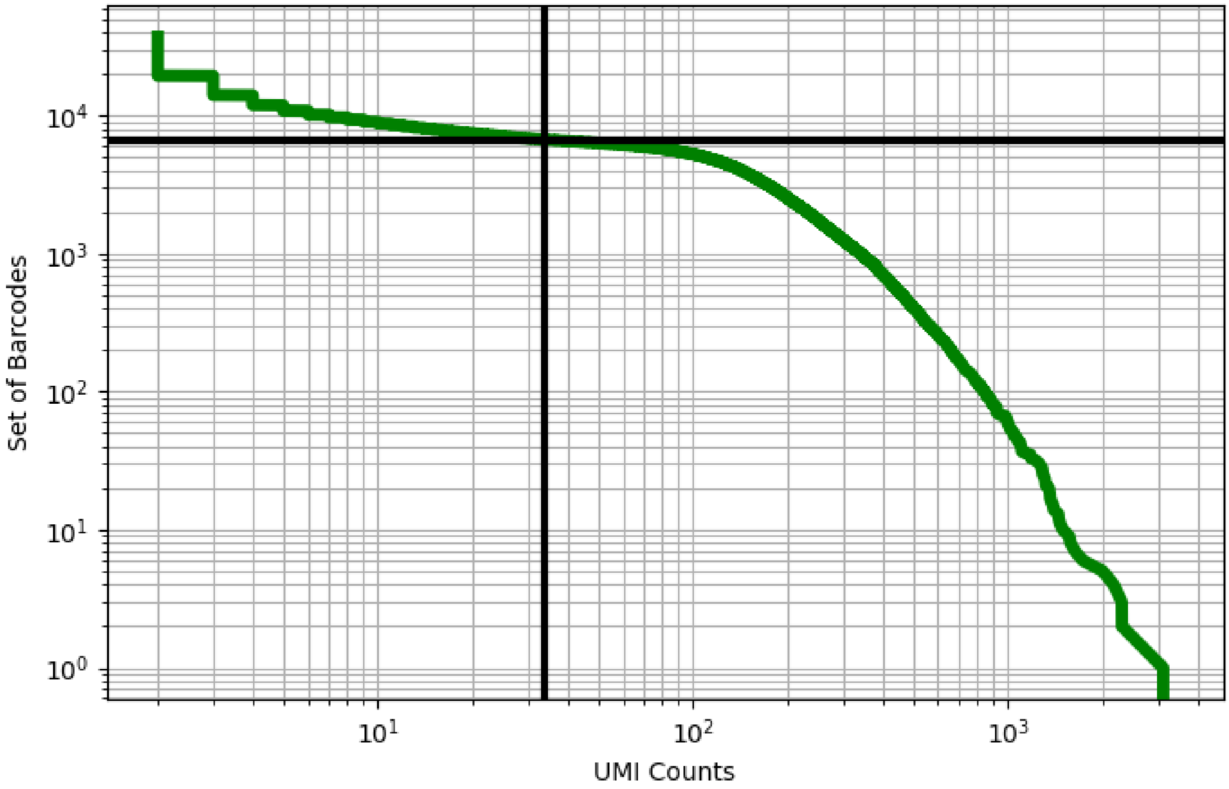


Figure 418: rectum/GSM3576399

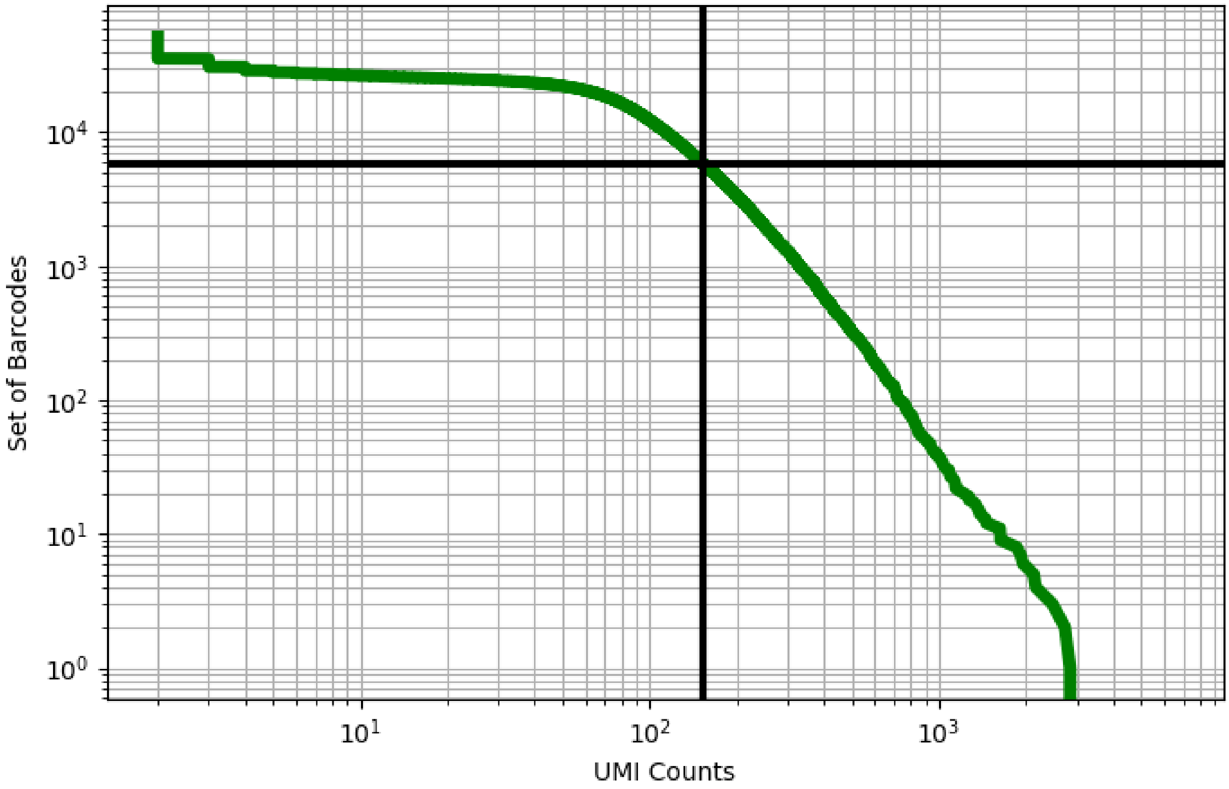


Figure 419: rectum/GSM3576400

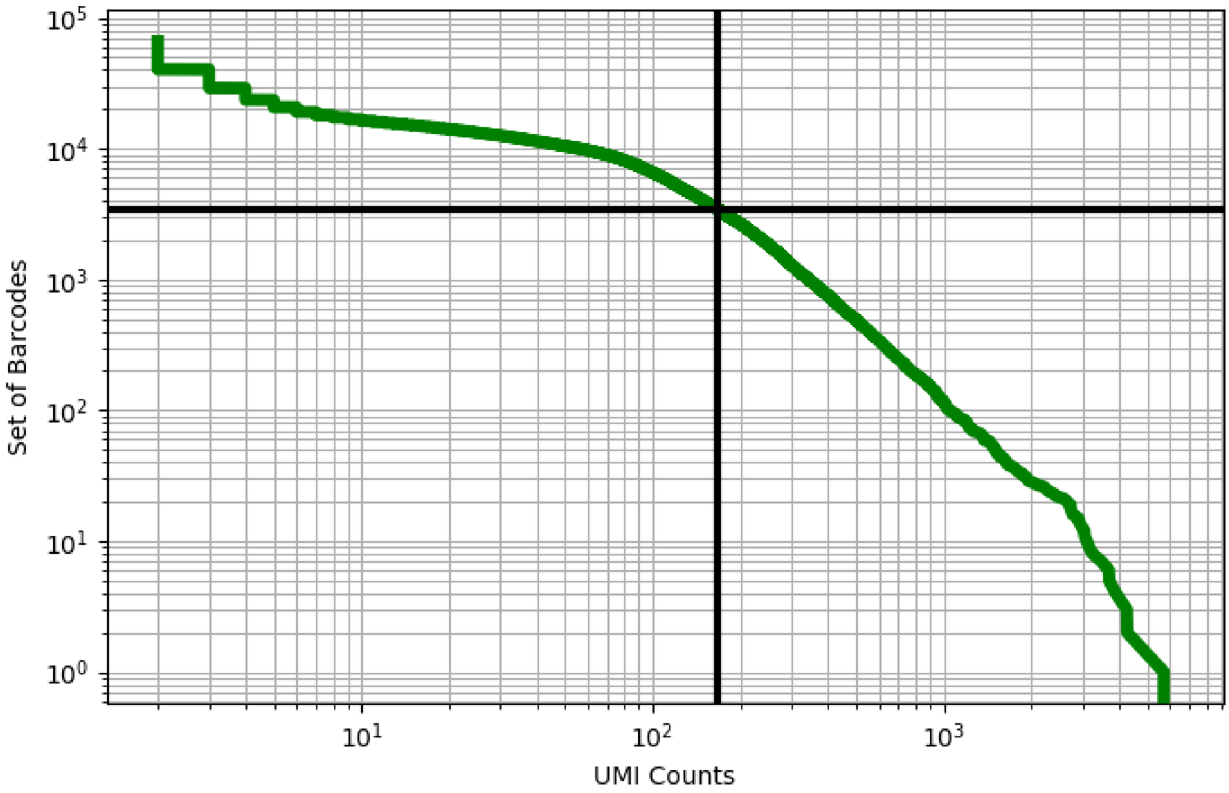


Figure 420: rectum/GSM3576401

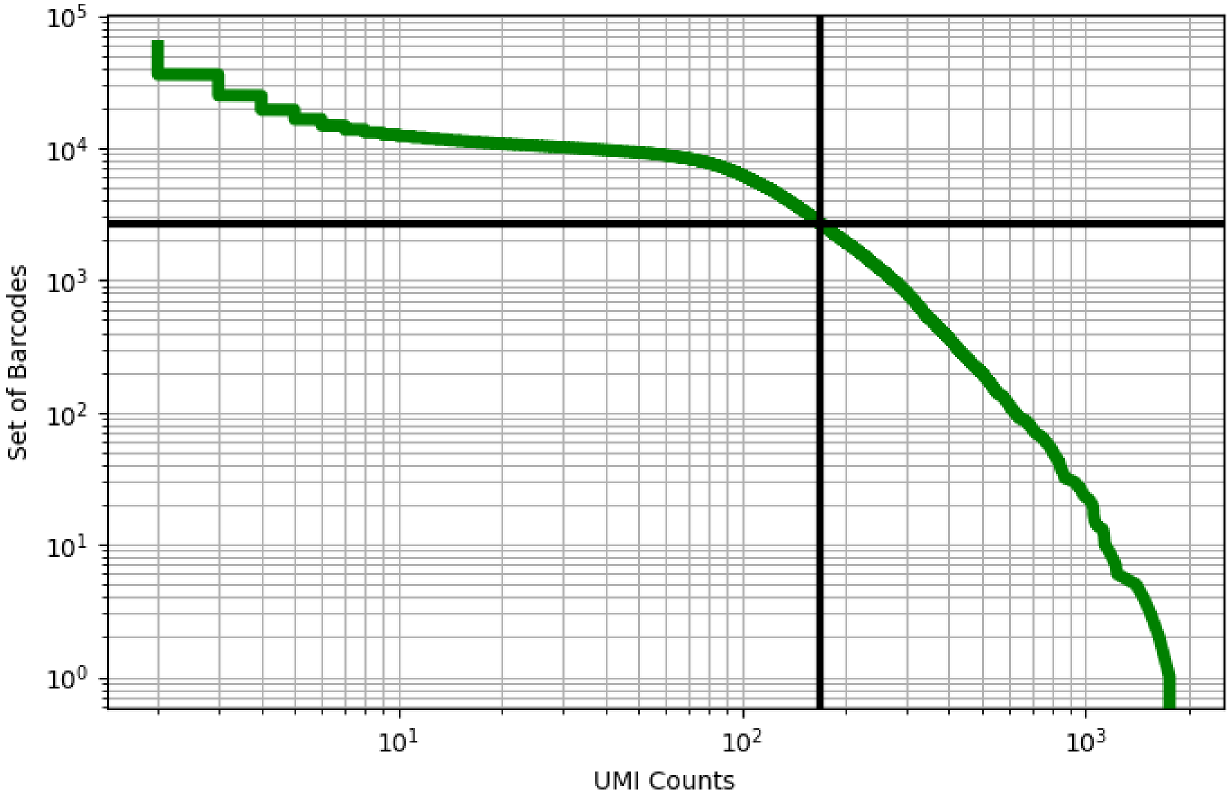


Figure 421: rectum/GSM3576402

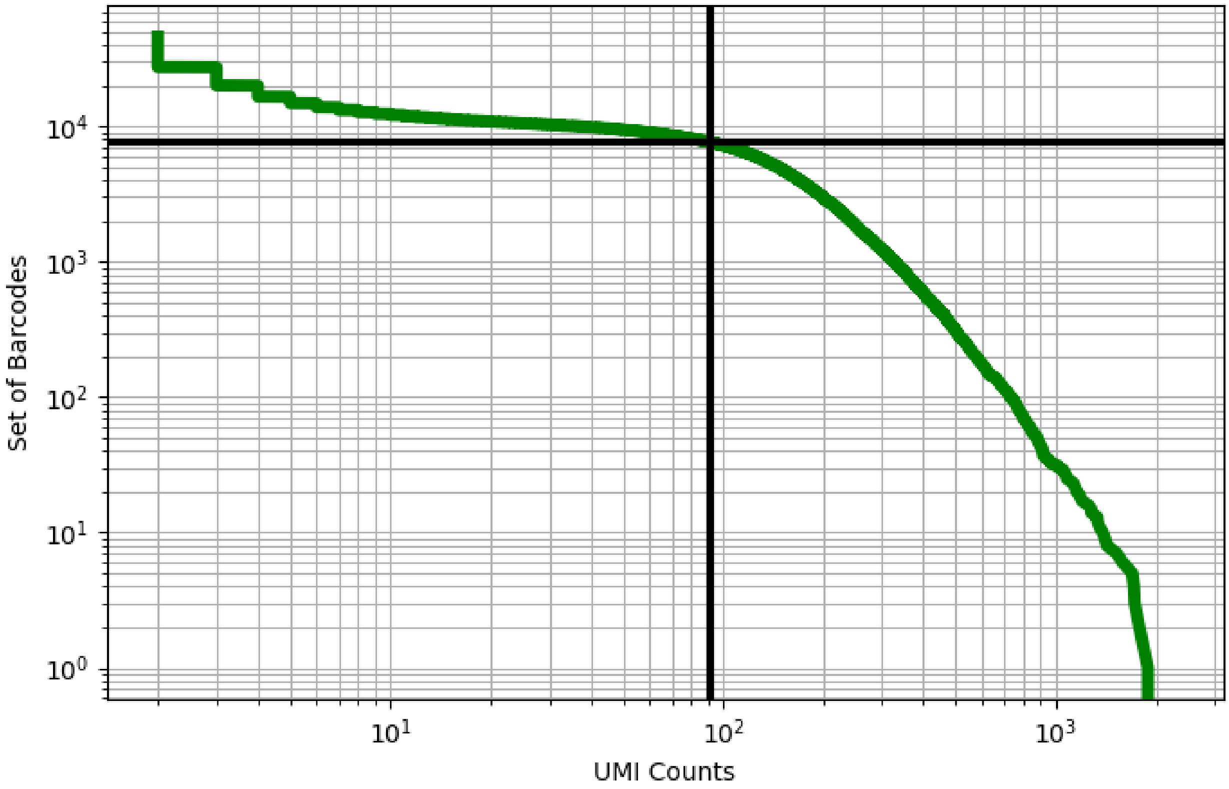


Figure 422: rectum/GSM3576403

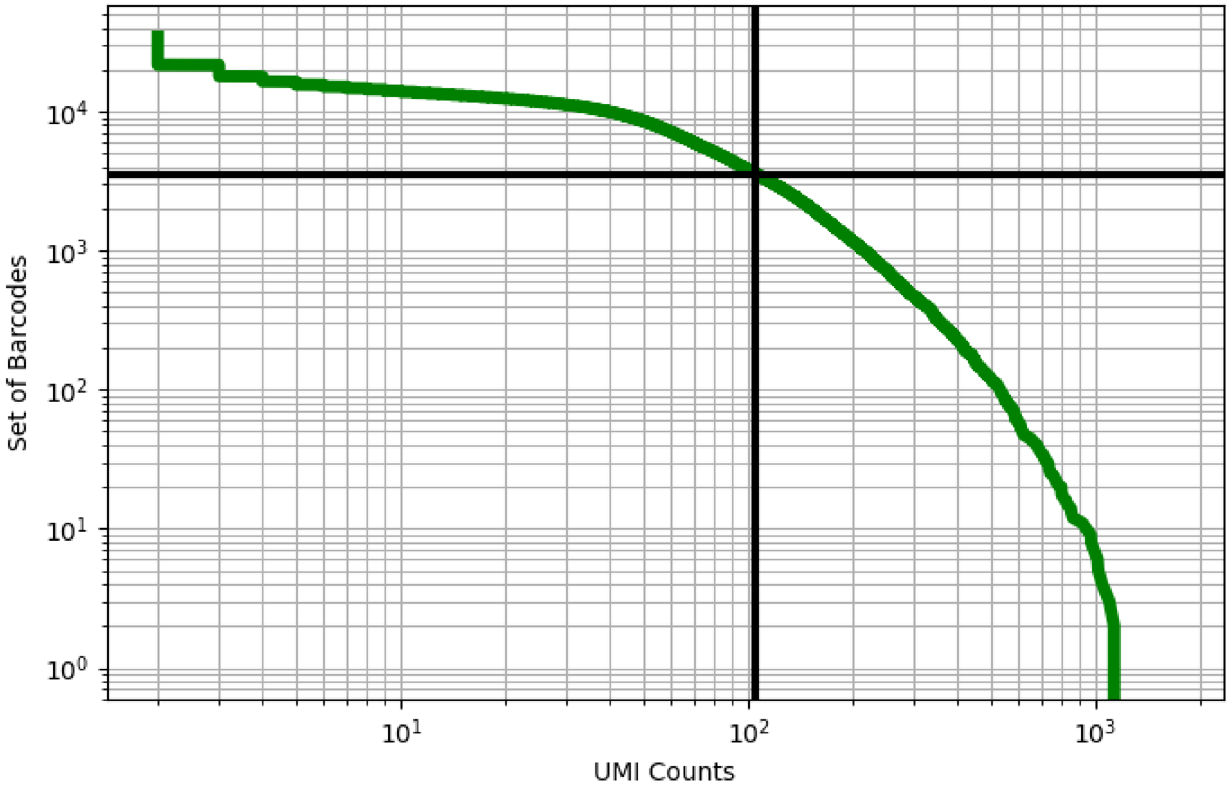


Figure 423: rectum/GSM3576404

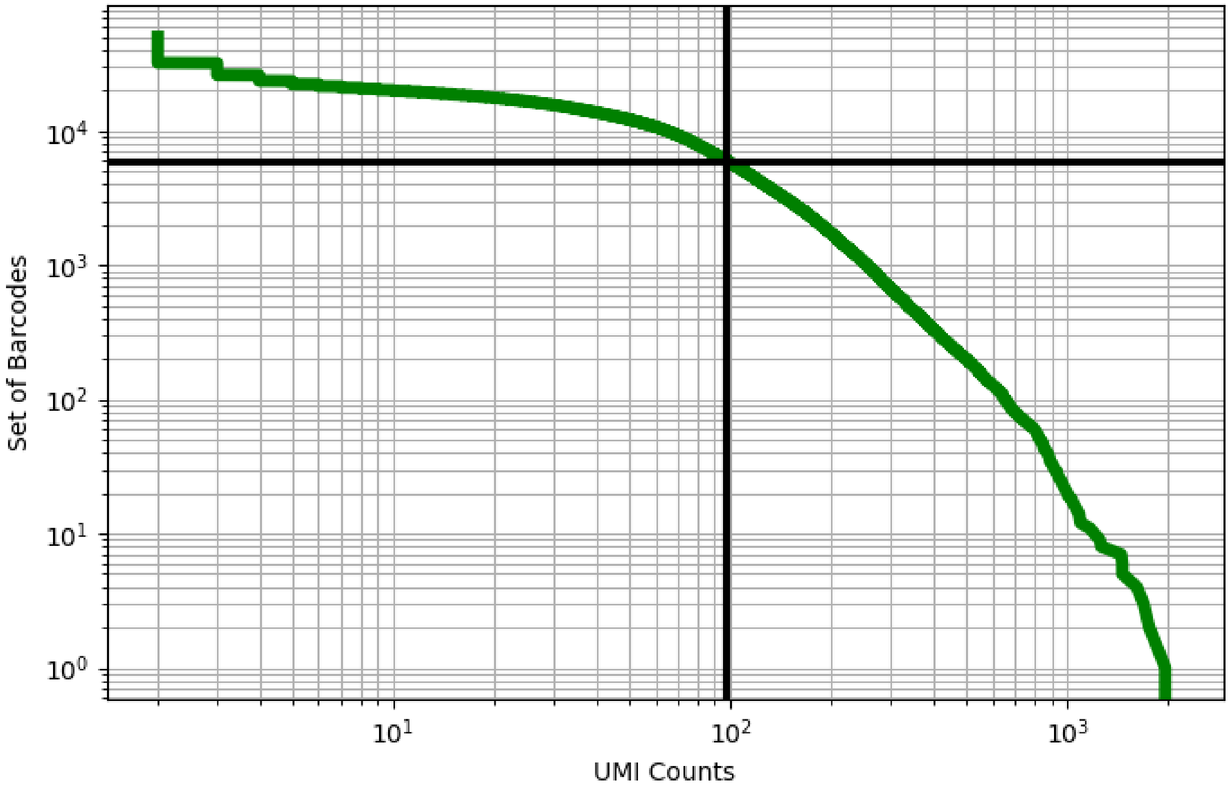


Figure 424: rectum/GSM3576405

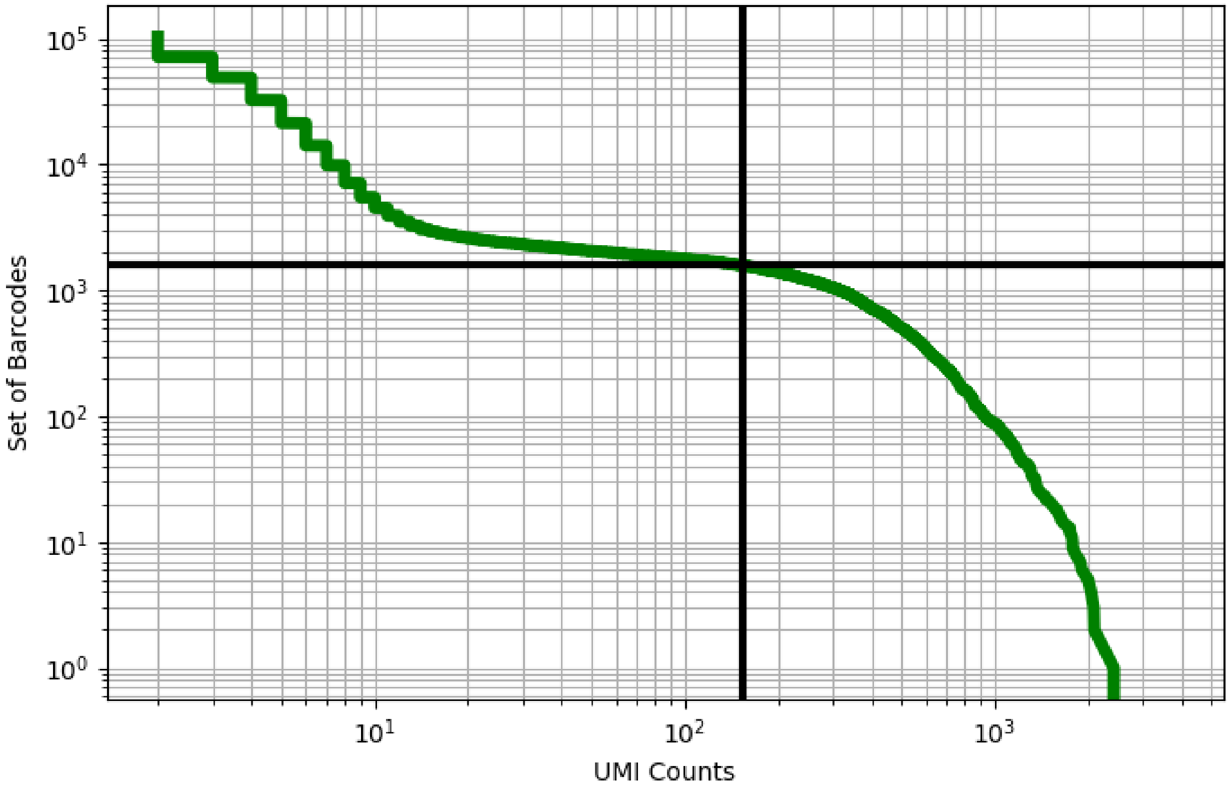


Figure 425: rectum/GSM3576406

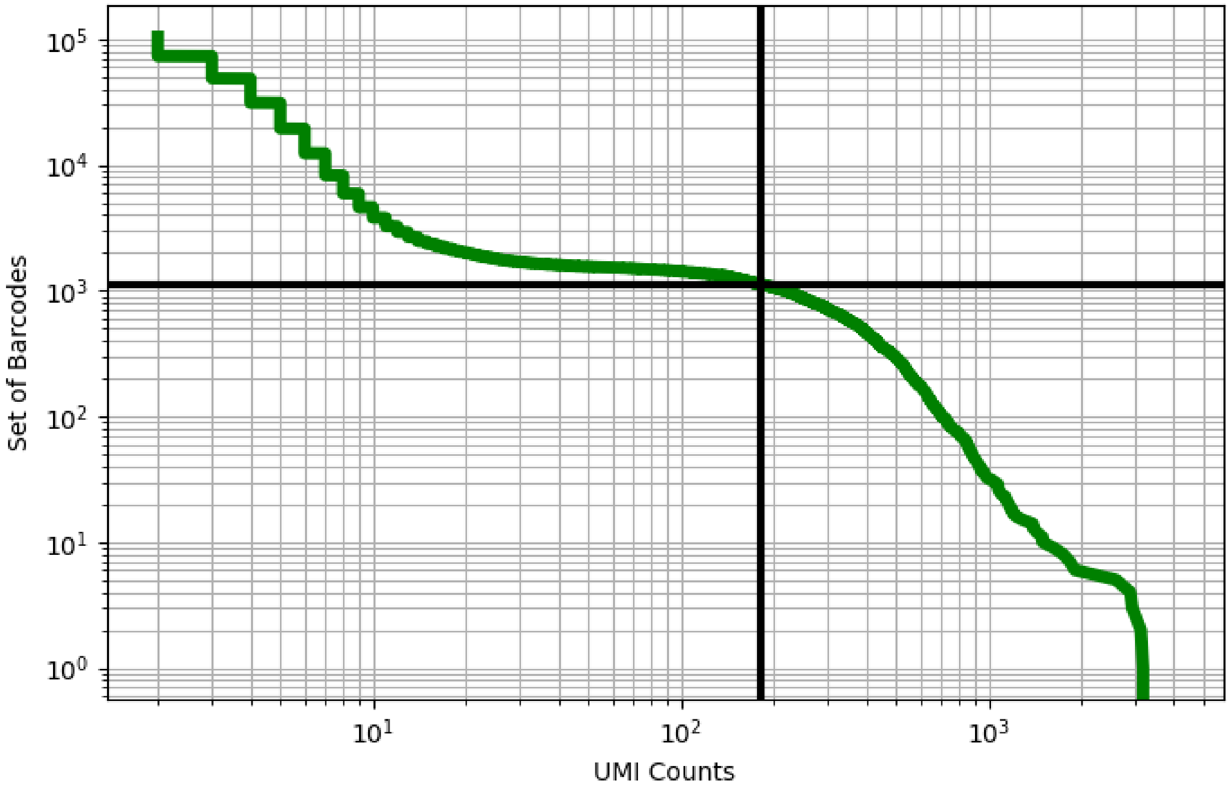


Figure 426: rectum/GSM3576407

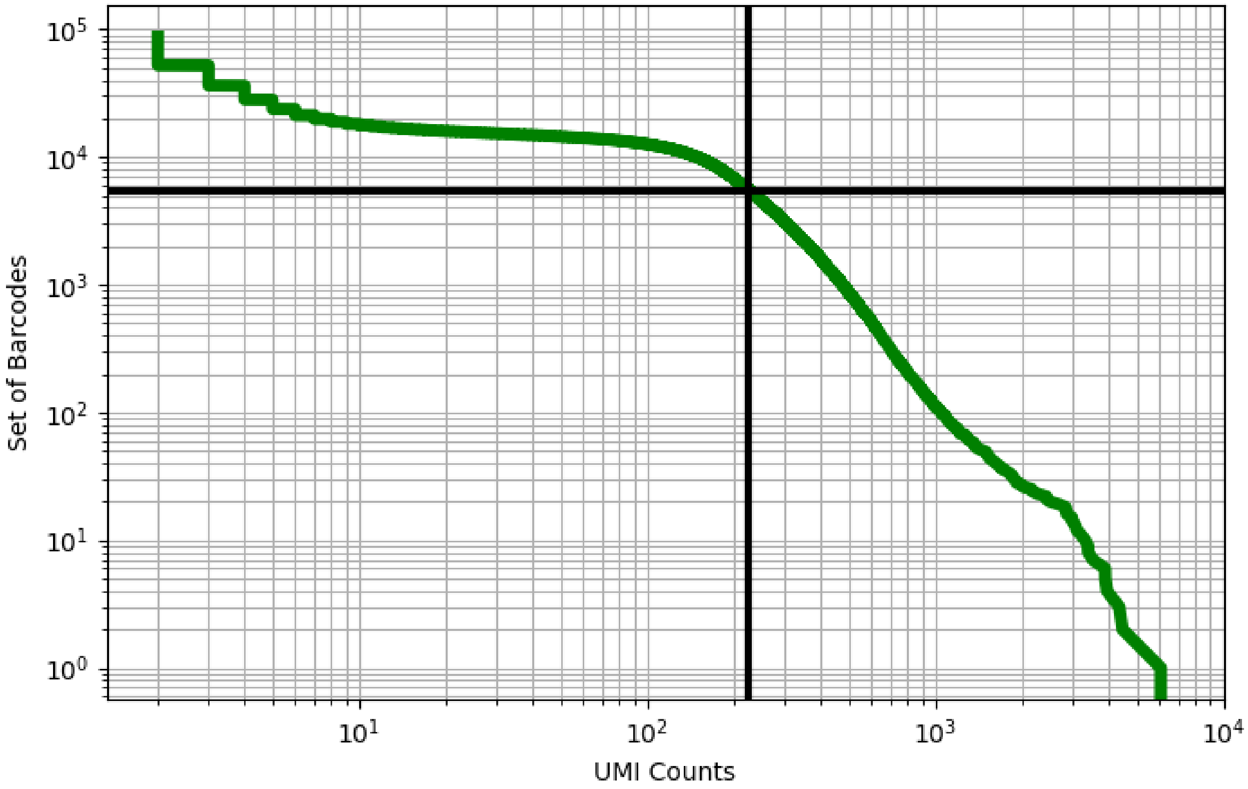


Figure 427: rectum/GSM3576408

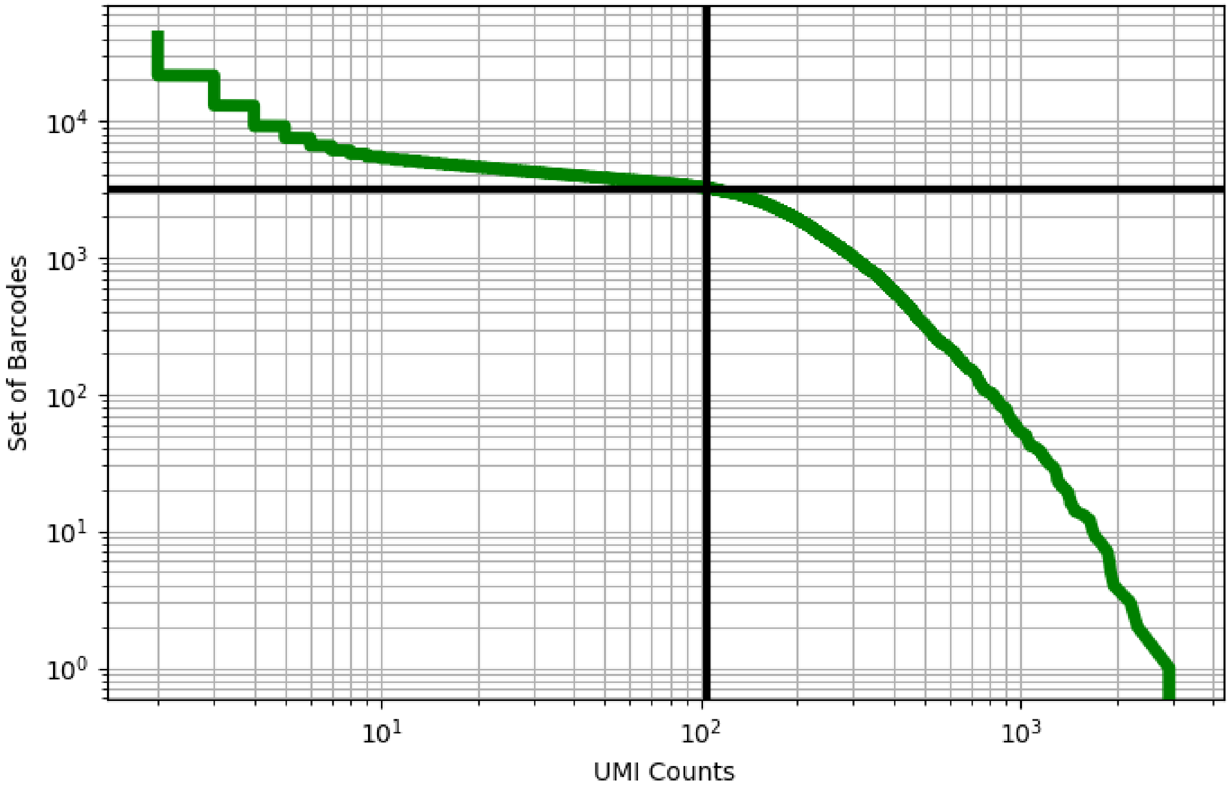


Figure 428: rectum/GSM3576409

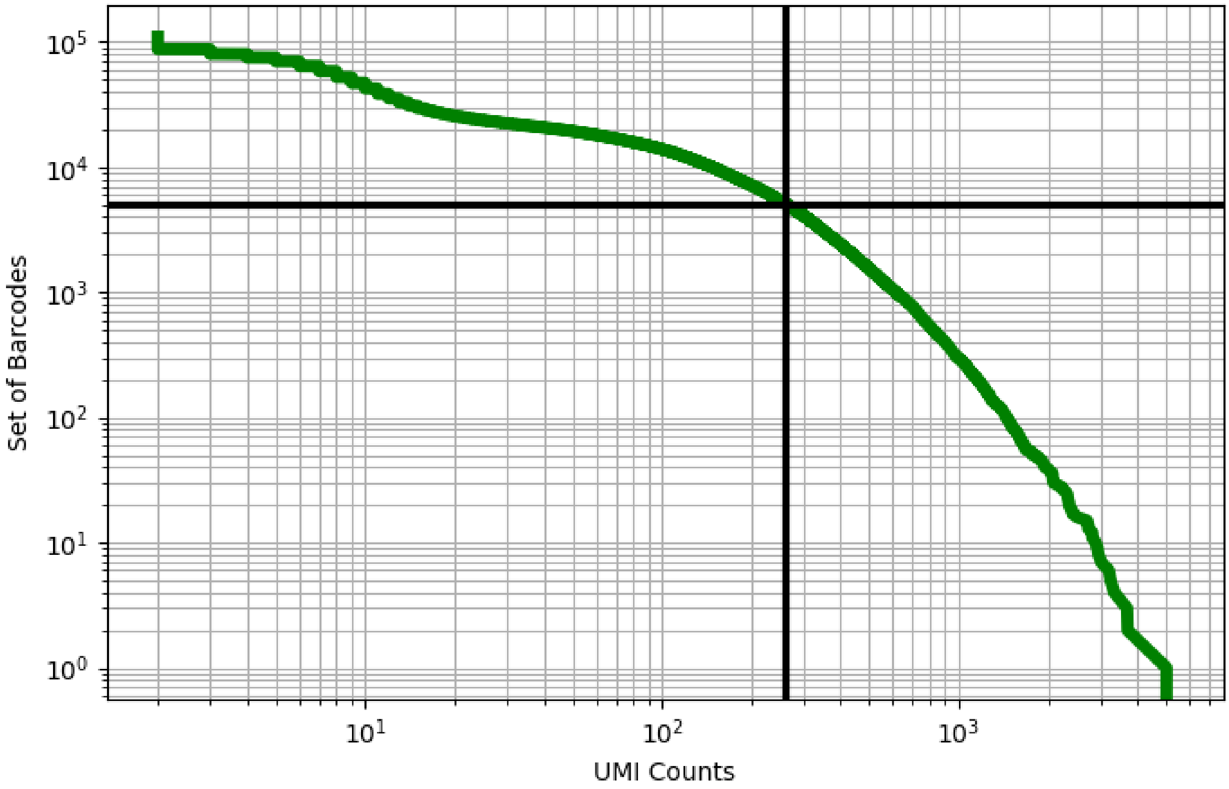


Figure 429: rectum/GSM3576410

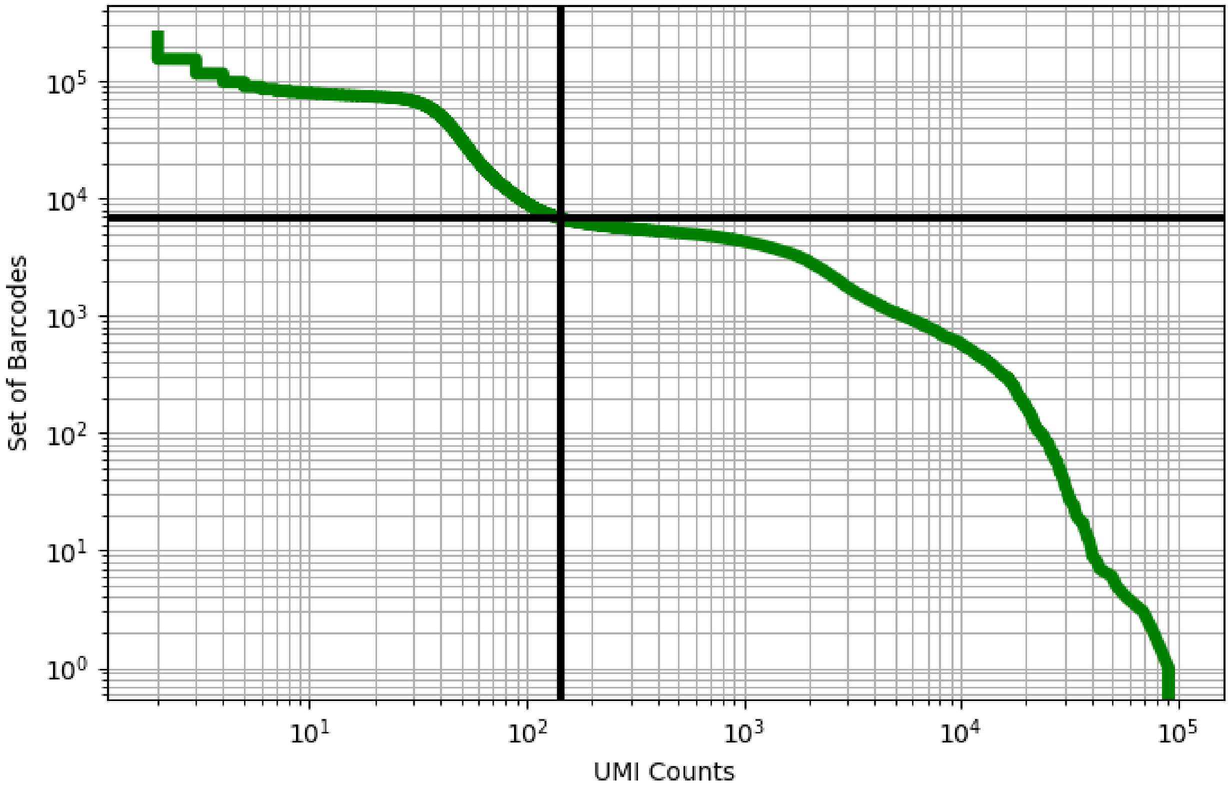


Figure 430: rectum/GSM3587013

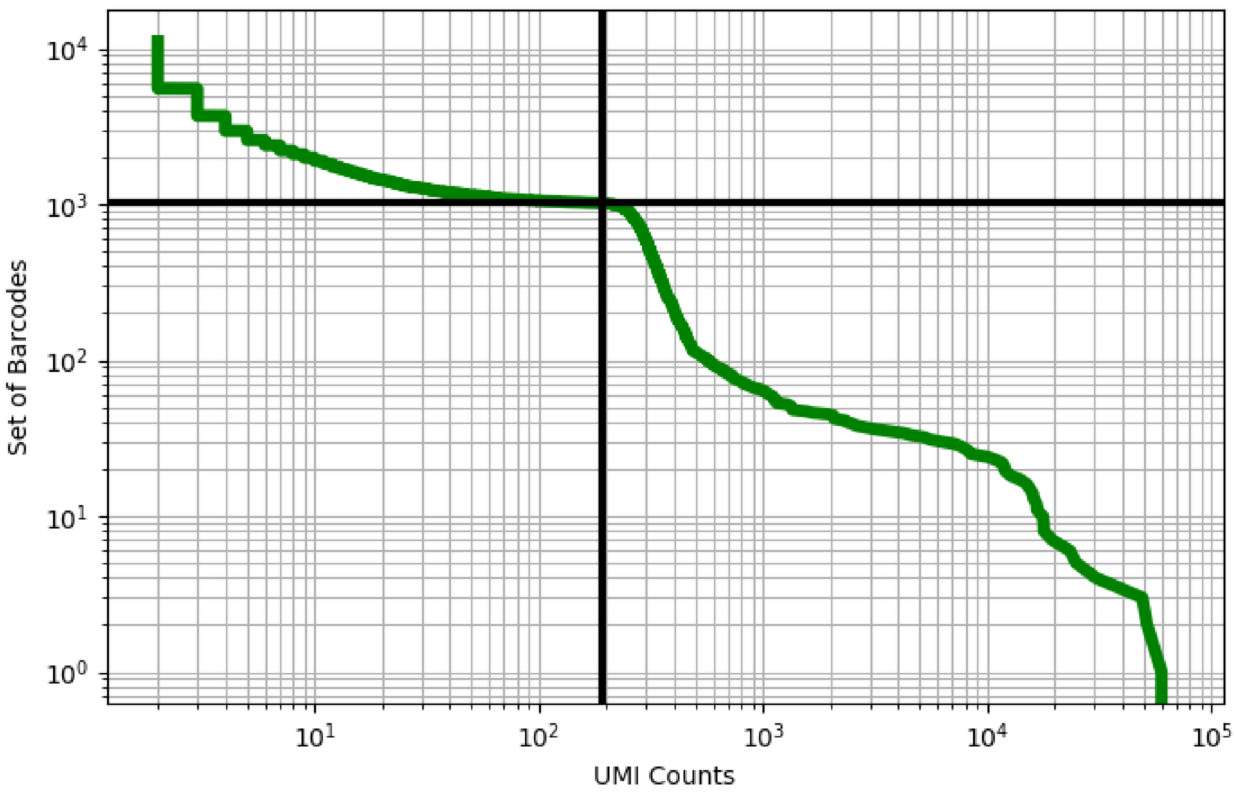


Figure 431: retina/GSM3745992

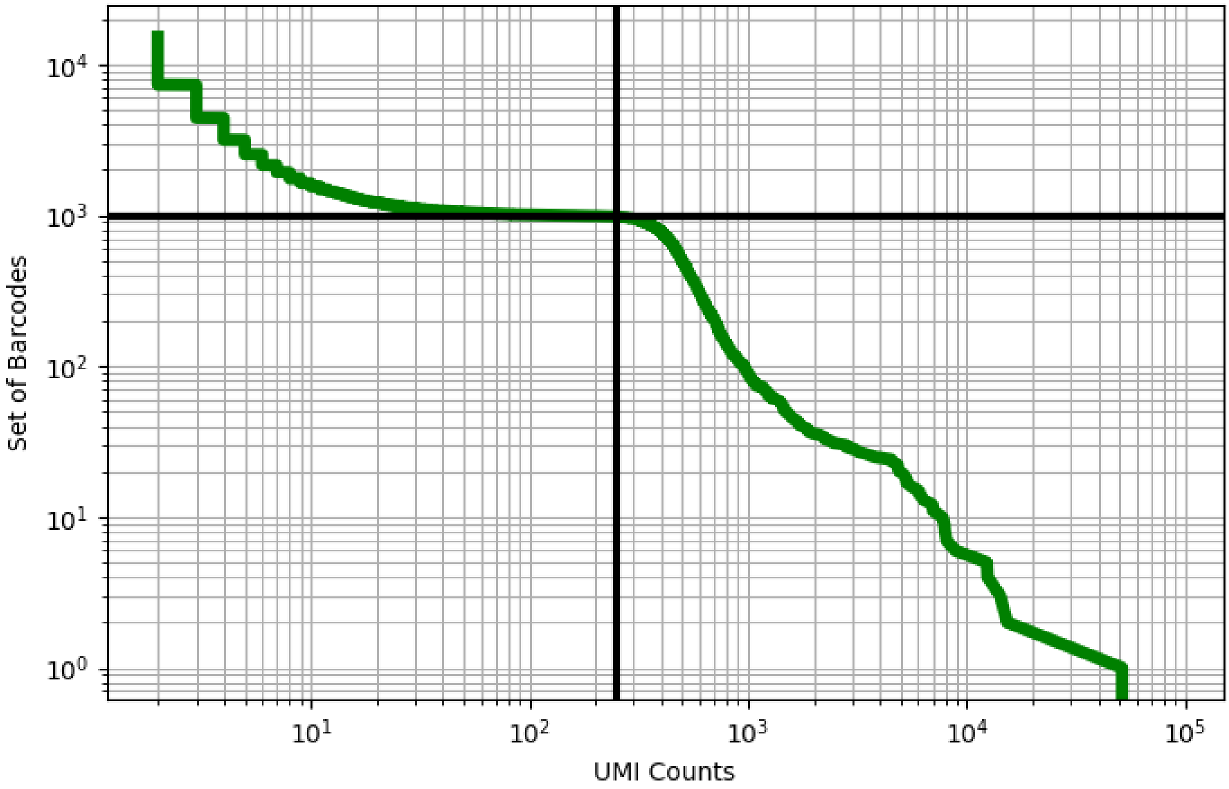


Figure 432: retina/GSM3745993

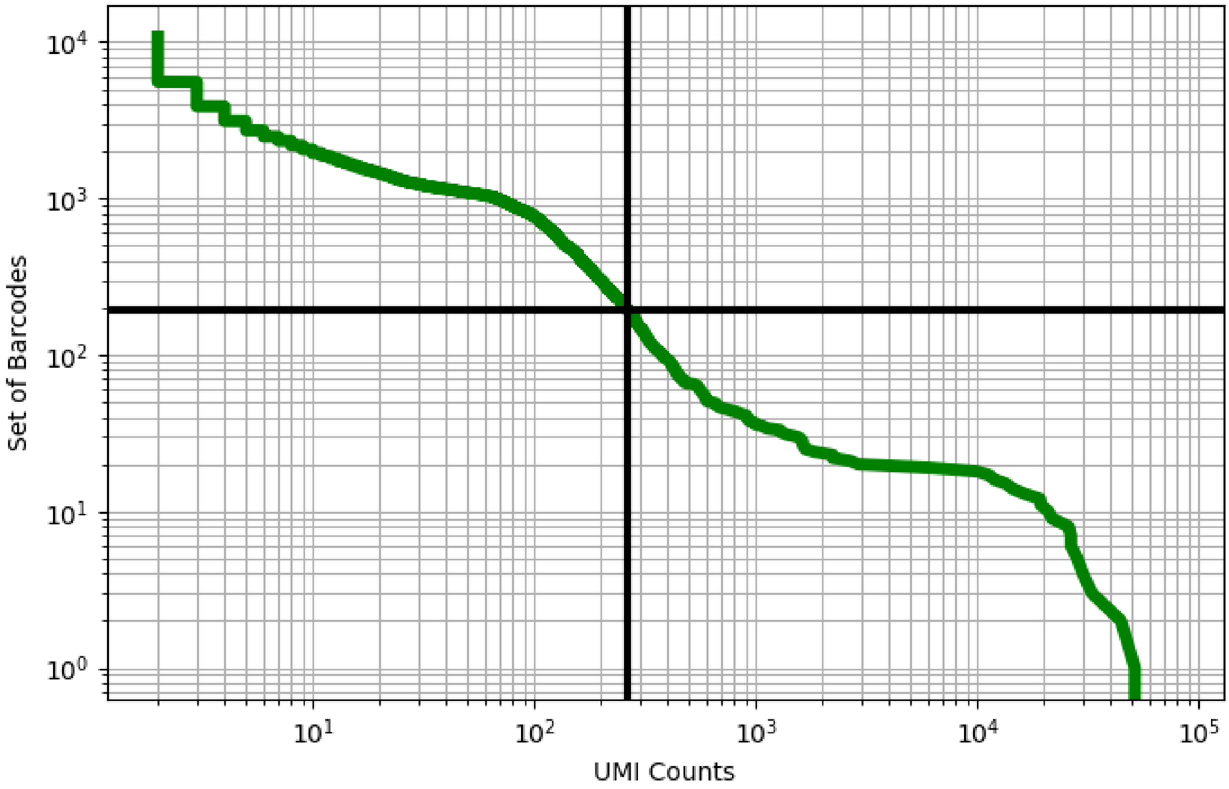


Figure 433: retina/GSM3745994

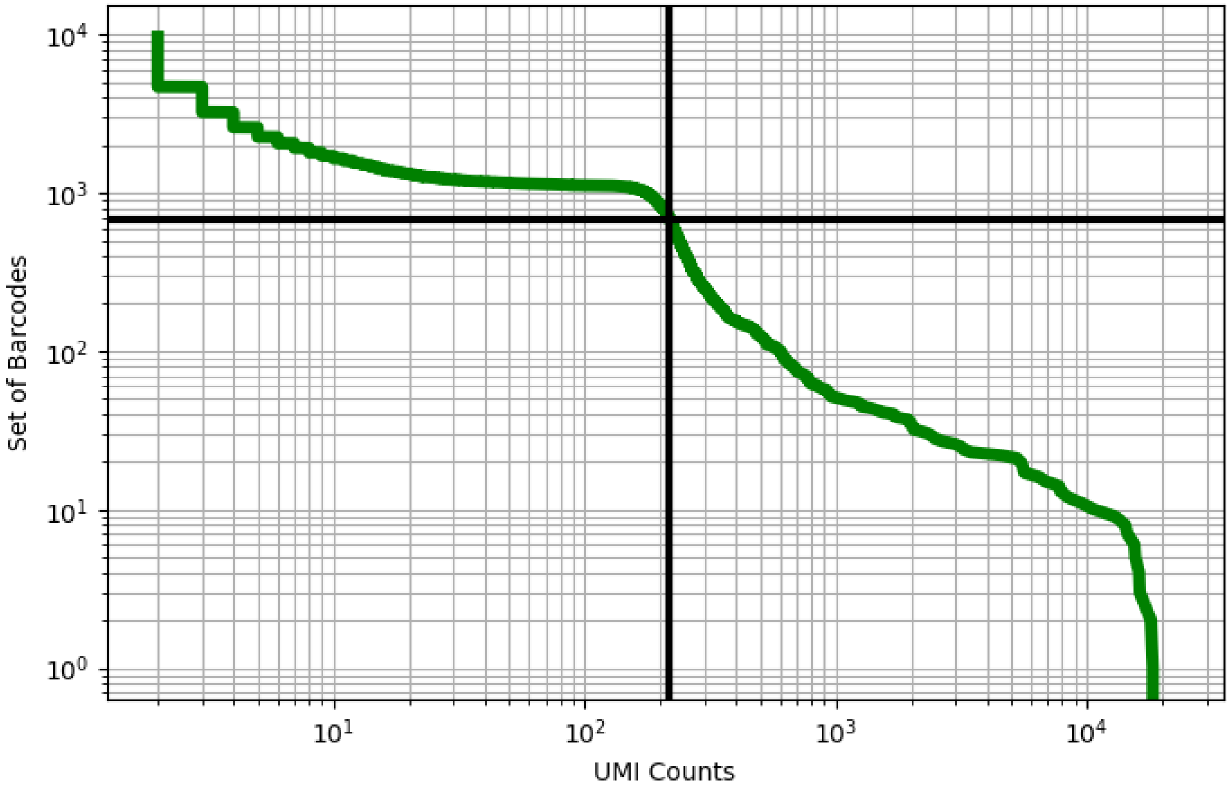


Figure 434: retina/GSM3745995

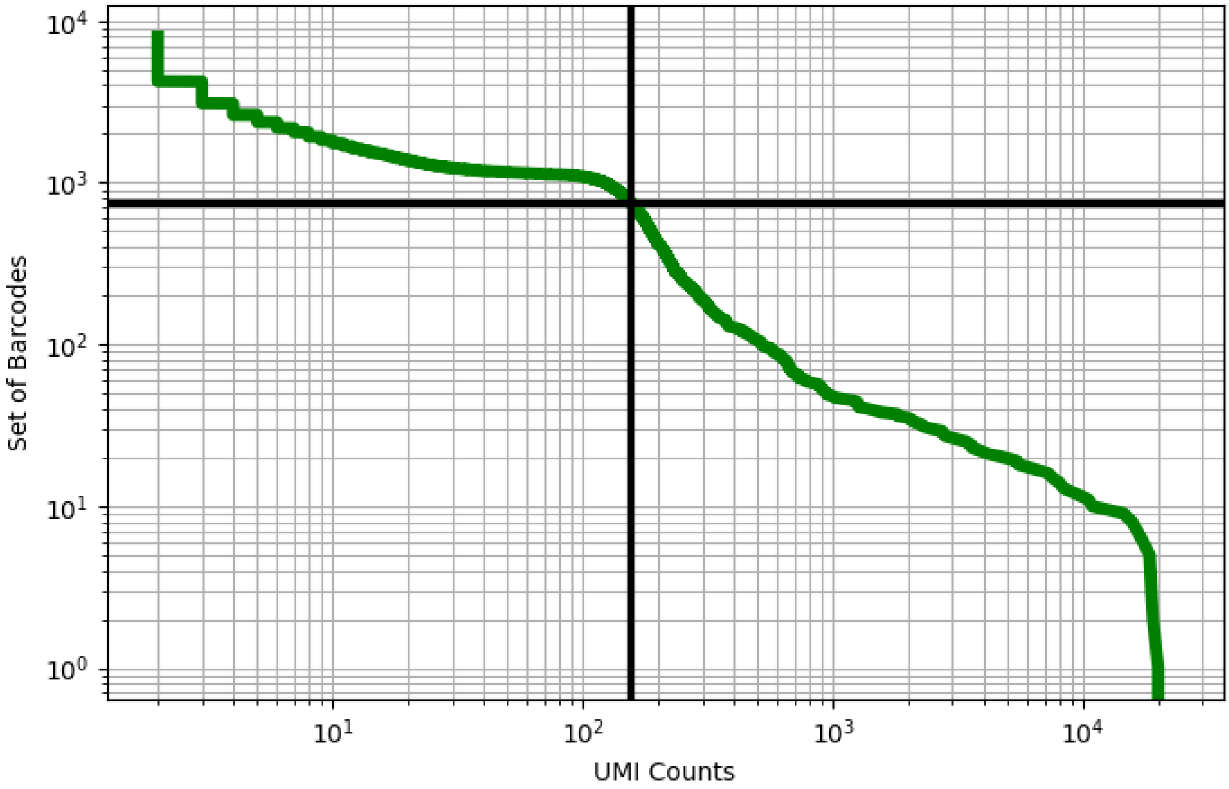


Figure 435: retina/GSM3745996

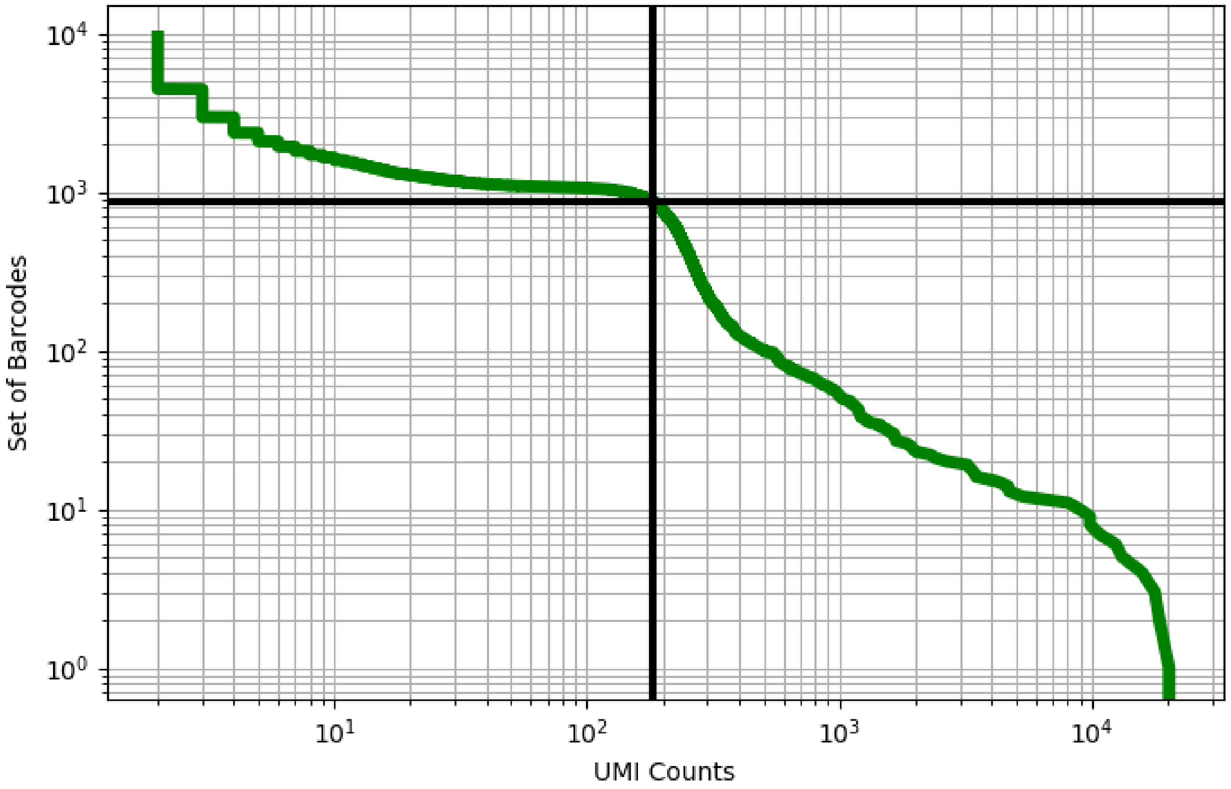


Figure 436: retina/GSM3745997

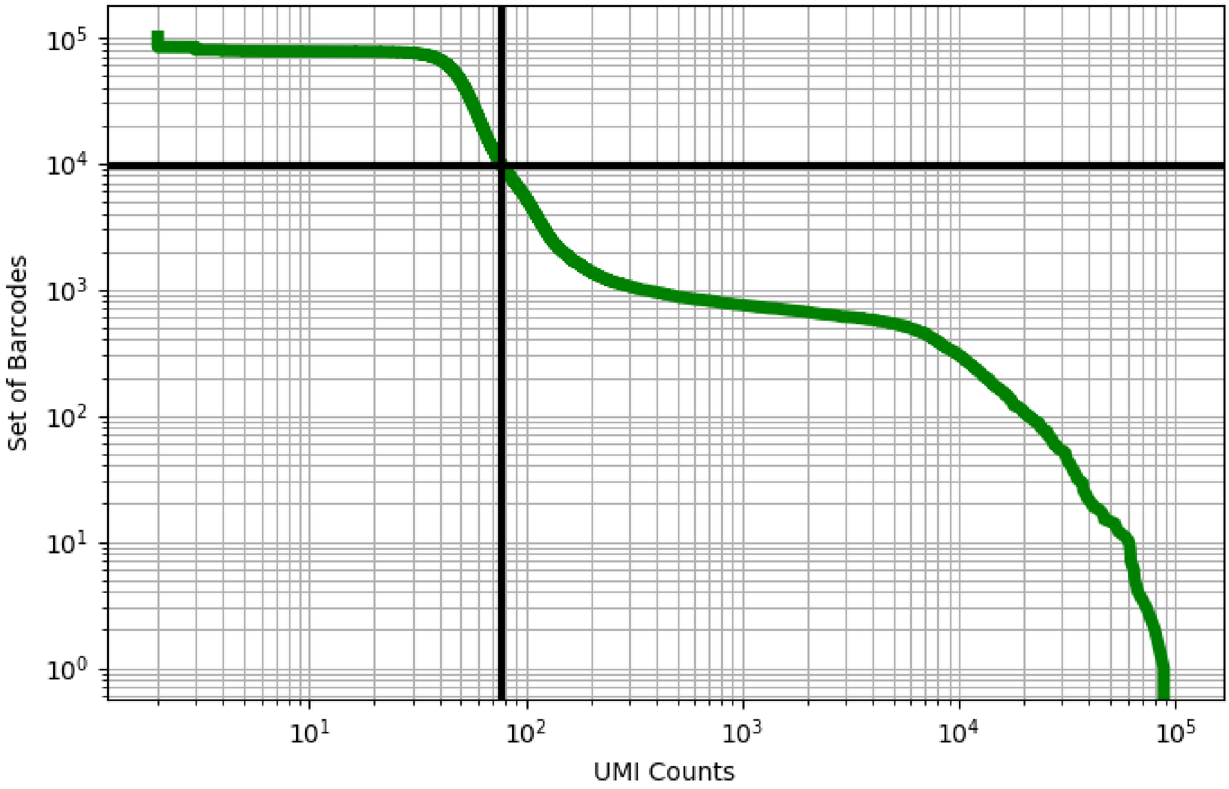


Figure 437: skin/ERS3861775

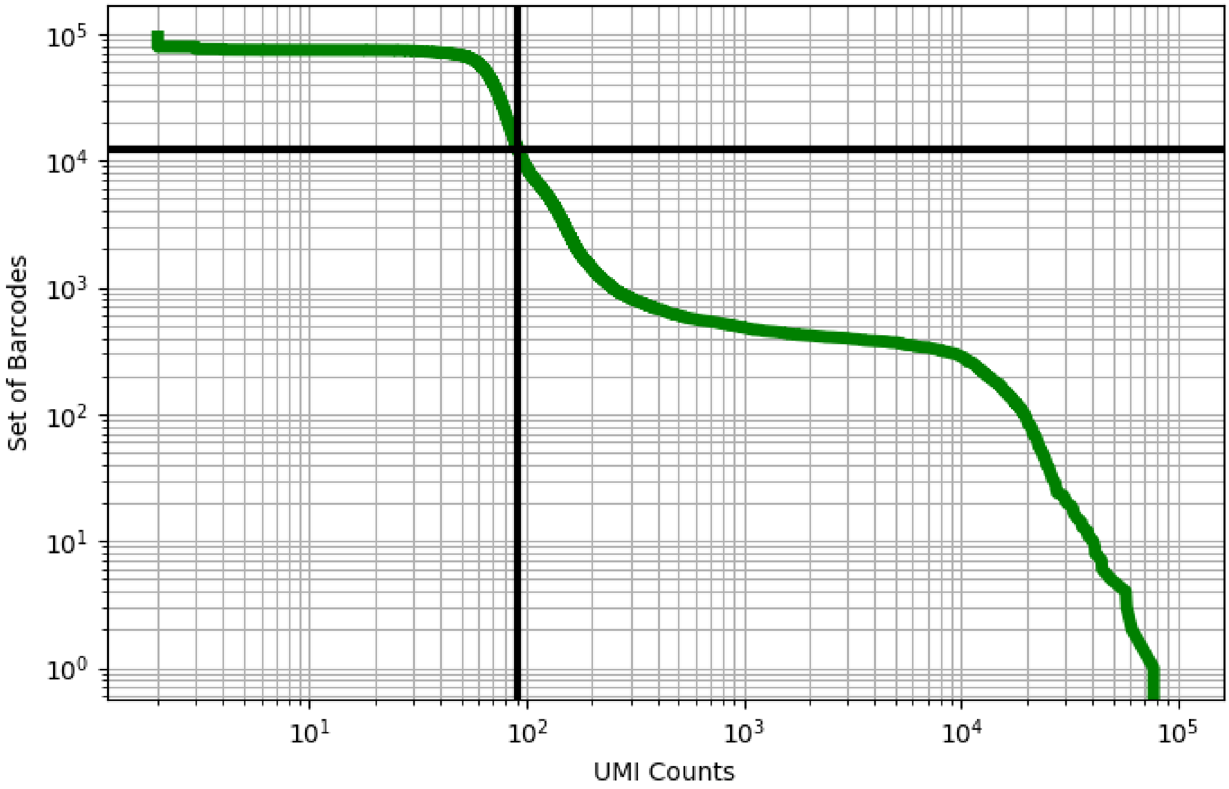


Figure 438: skin/ERS3861776

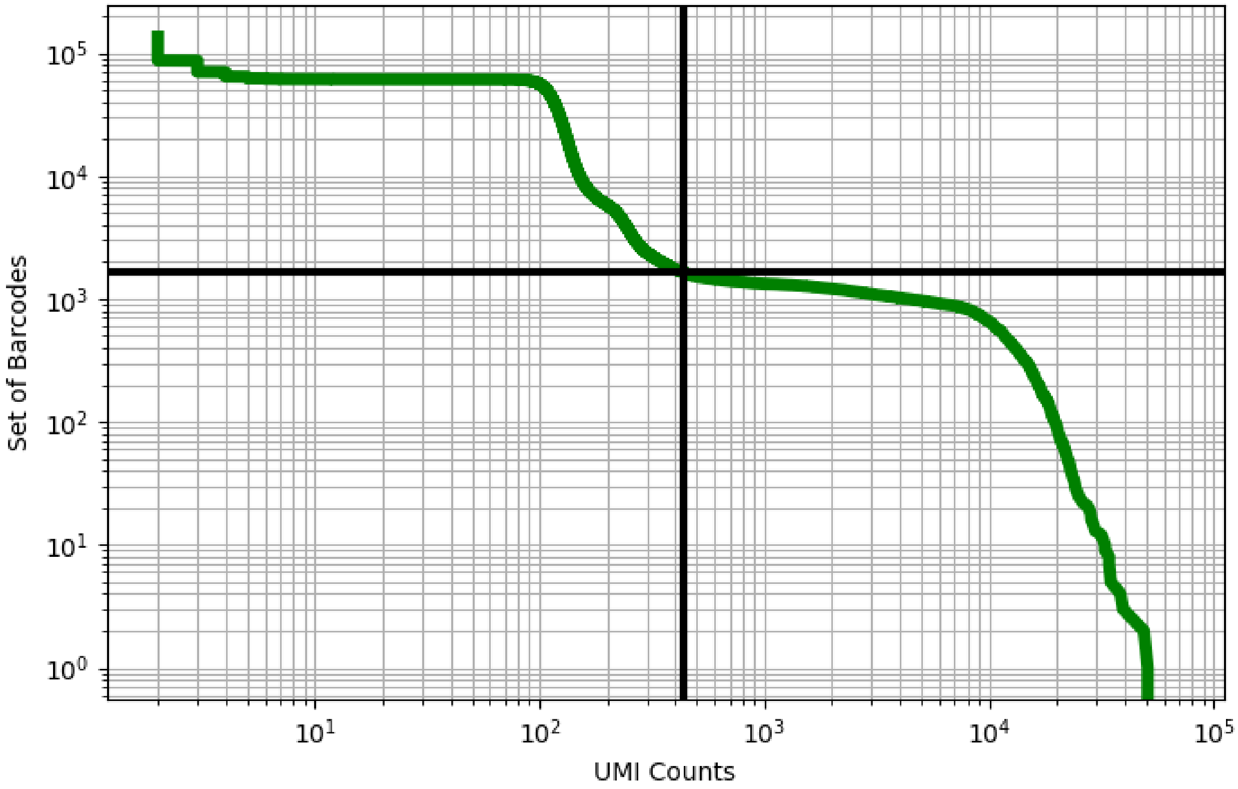


Figure 439: skin/ERS3861779

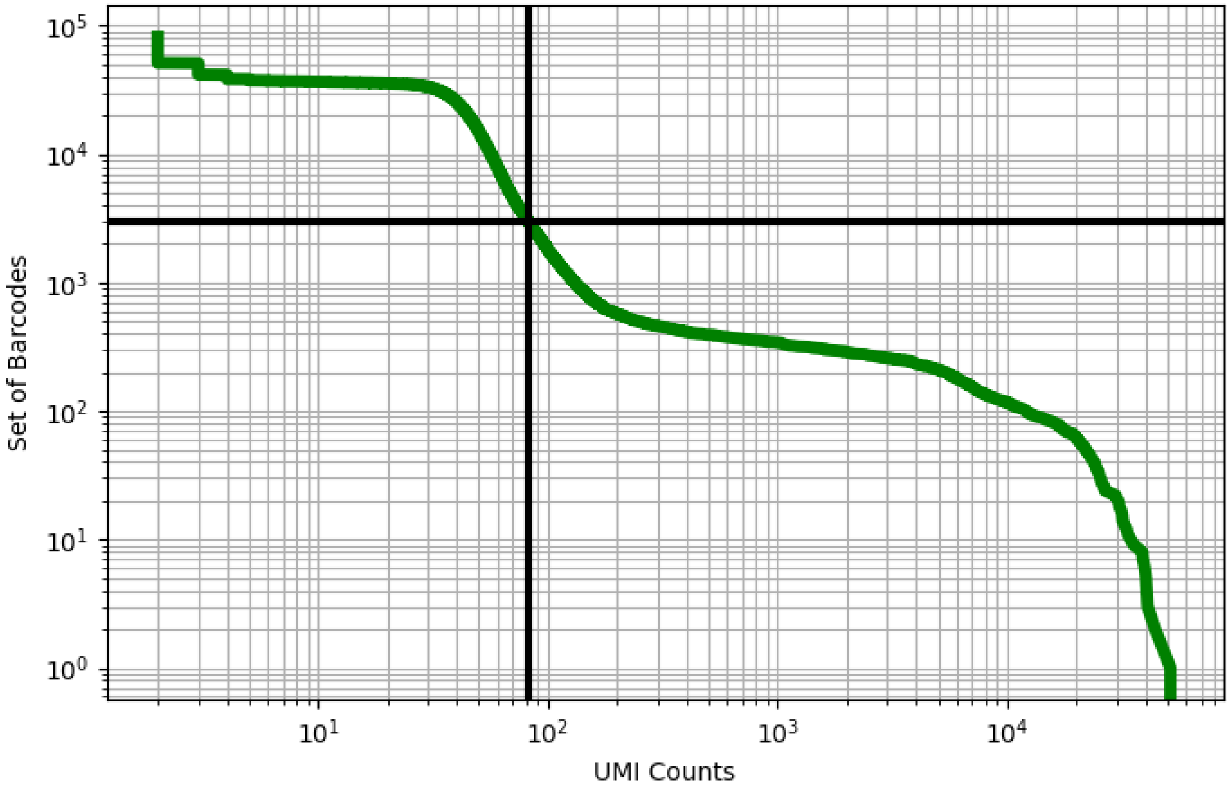


Figure 440: skin/ERS3861784

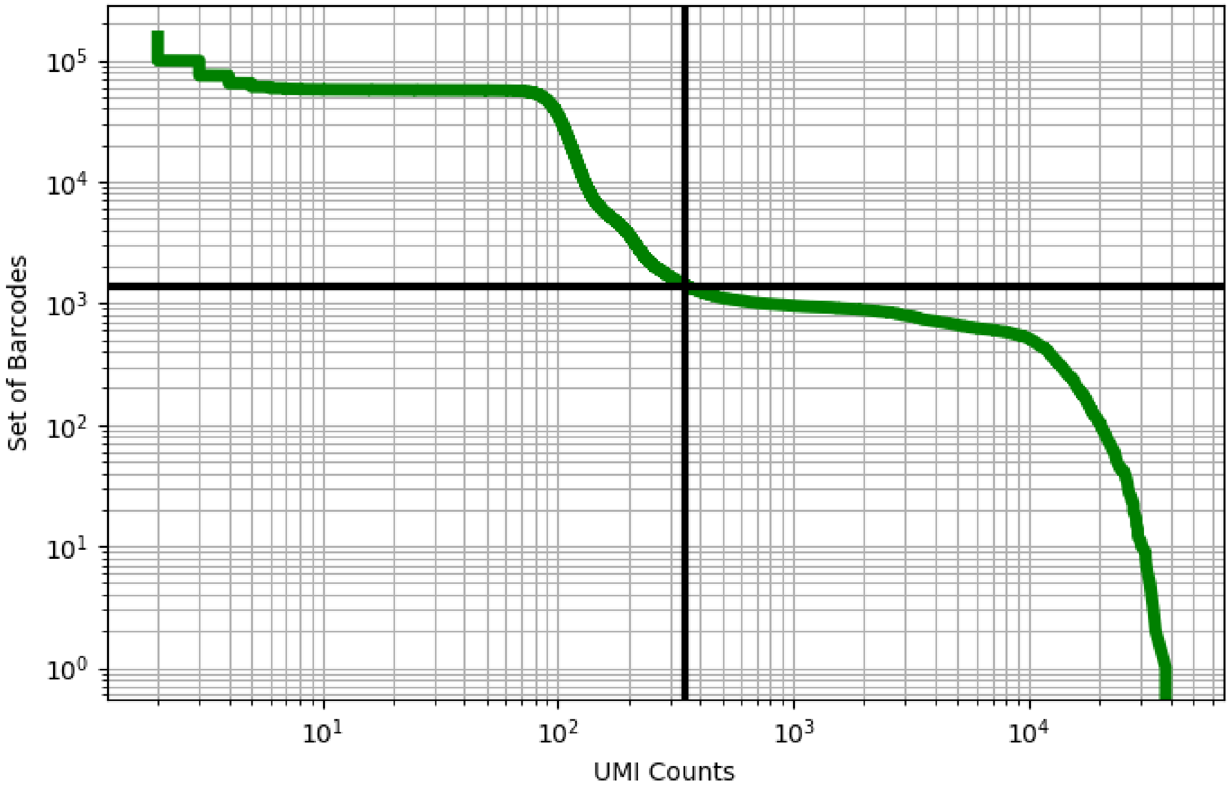


Figure 441: skin/ERS3861785

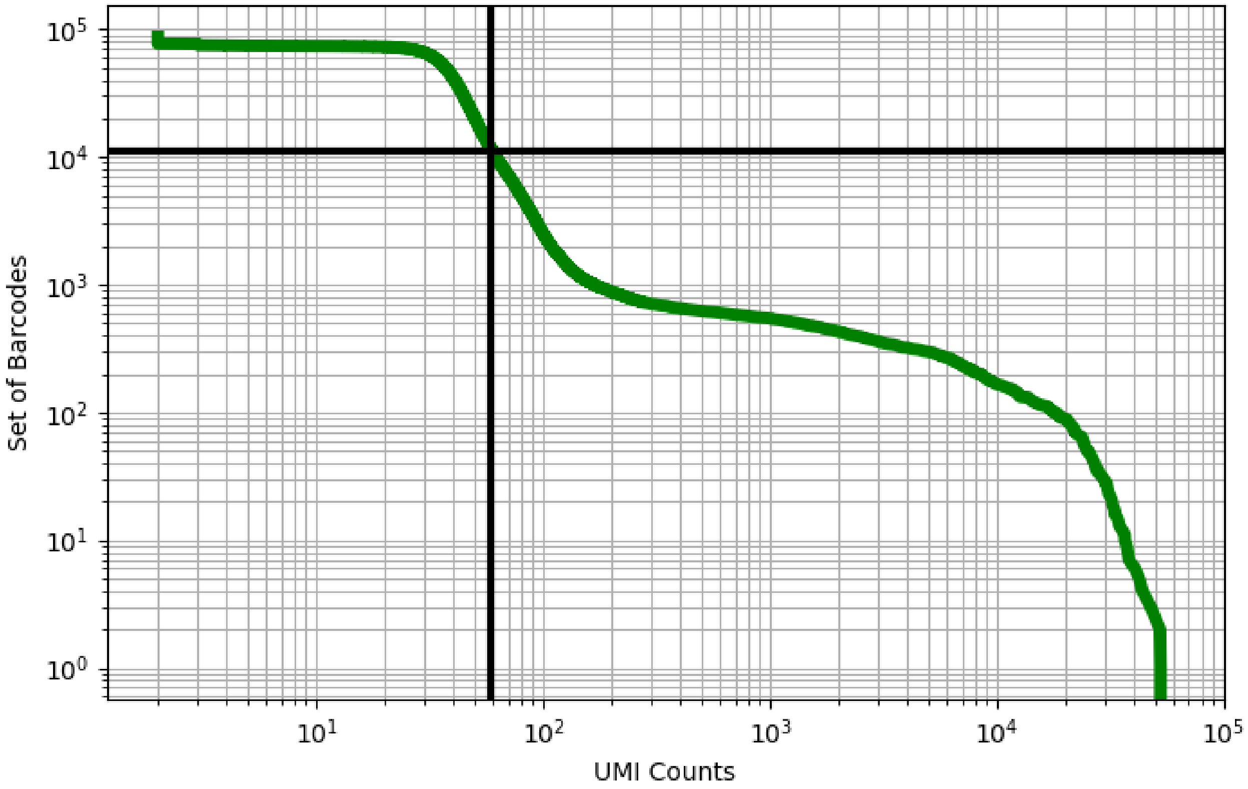


Figure 442: skin/ERS3861793

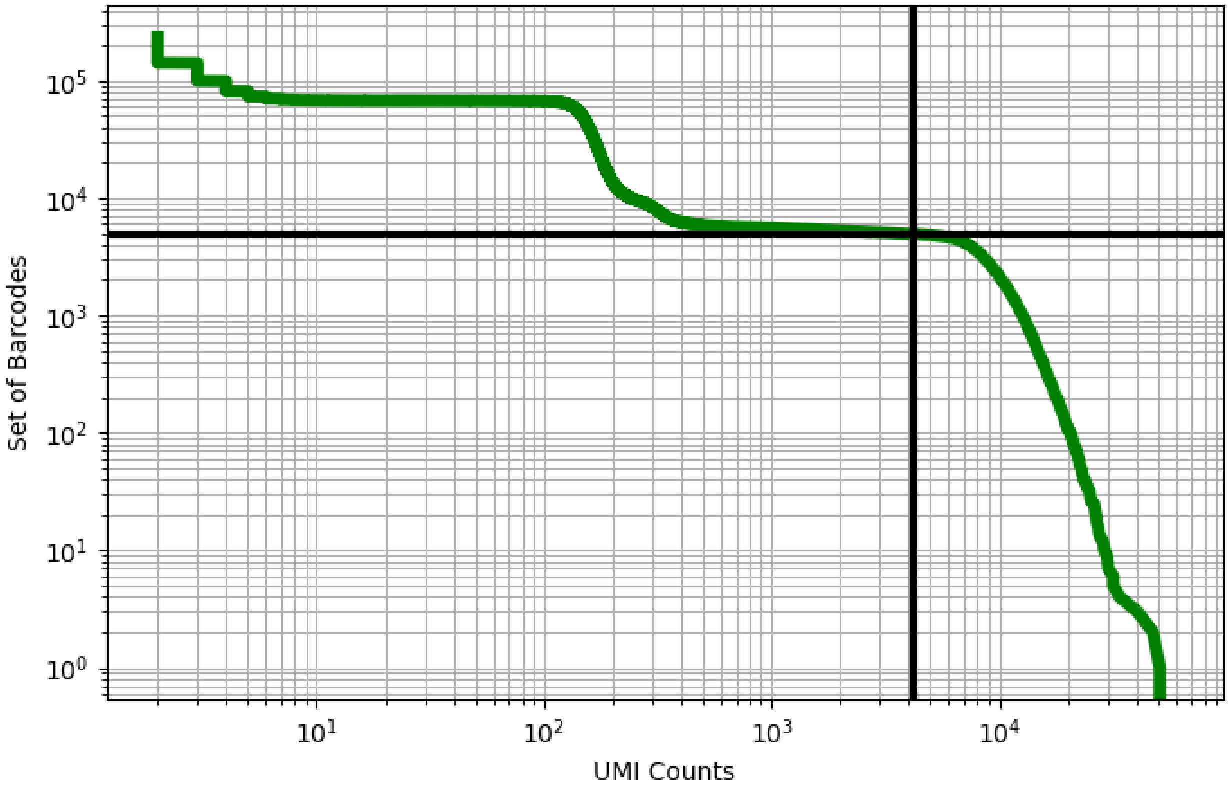


Figure 443: skin/ERS3861794

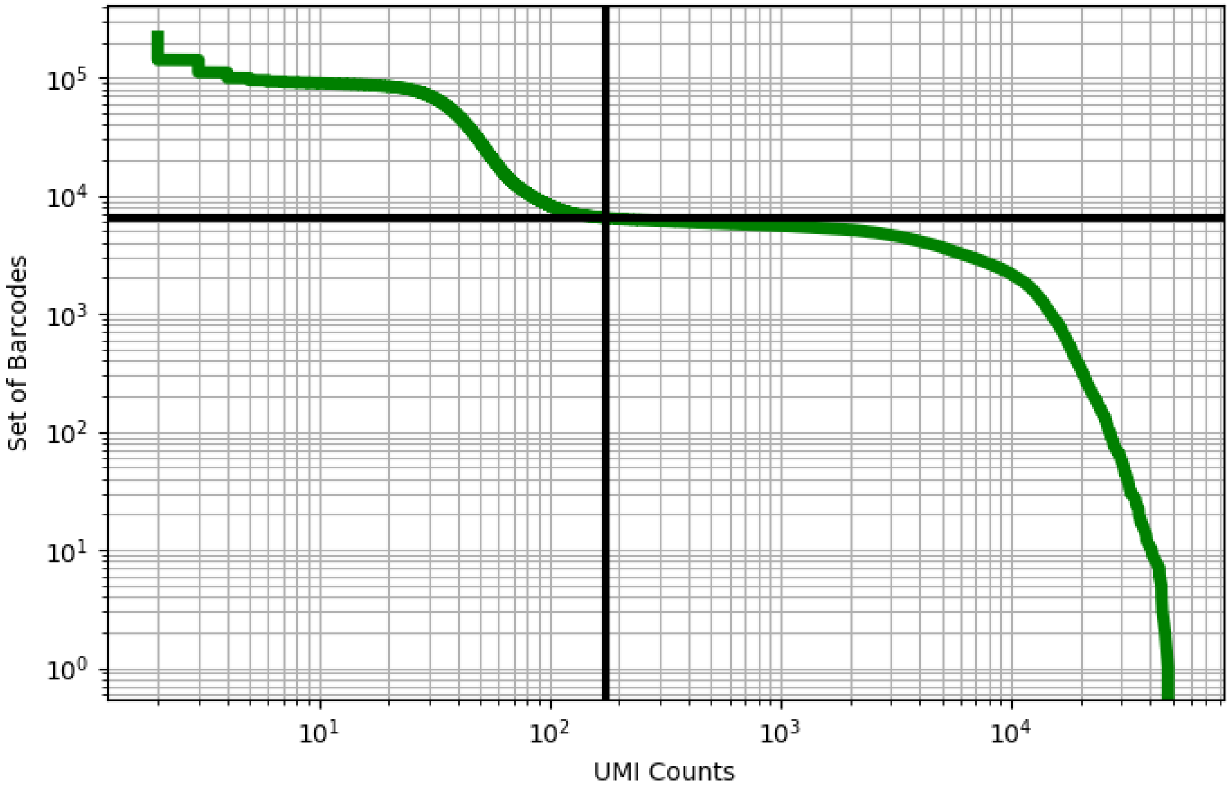


Figure 444: skin/ERS3861801

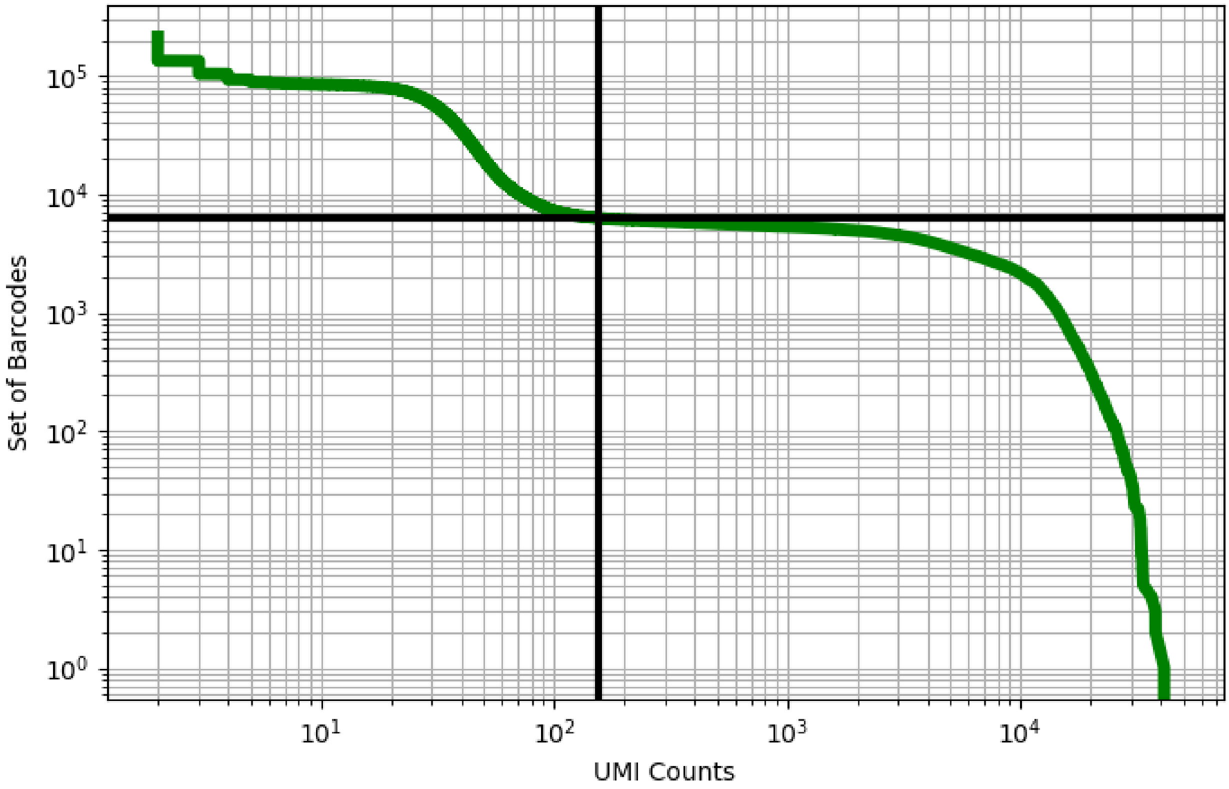


Figure 445: skin/ERS3861802

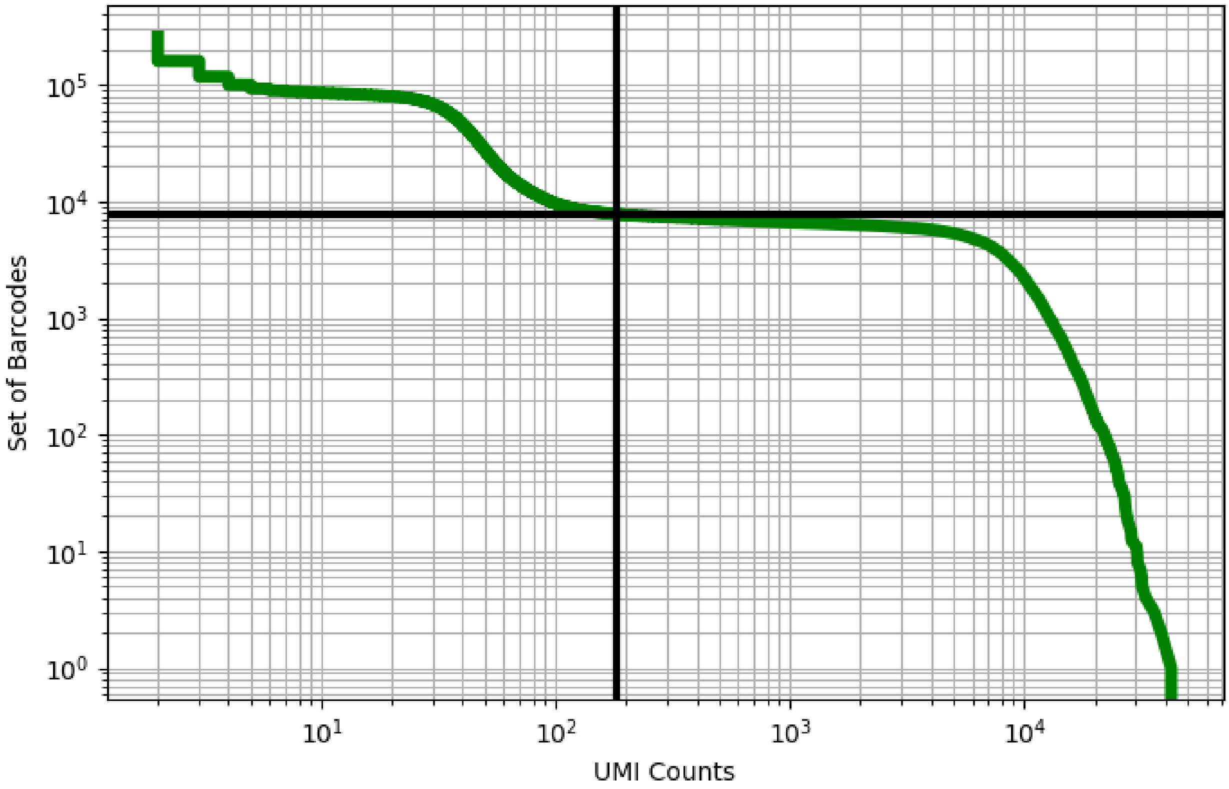


Figure 446: skin/ERS3861803

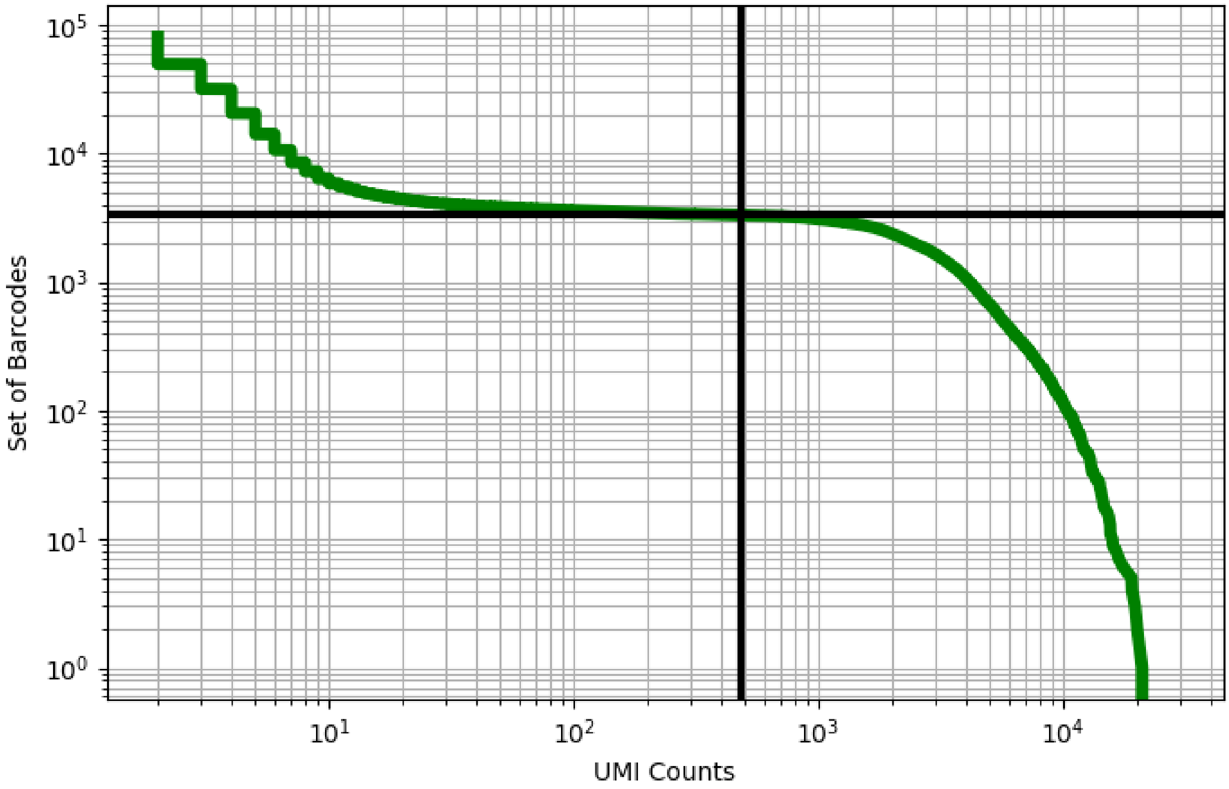


Figure 447: skin/ERS3861811

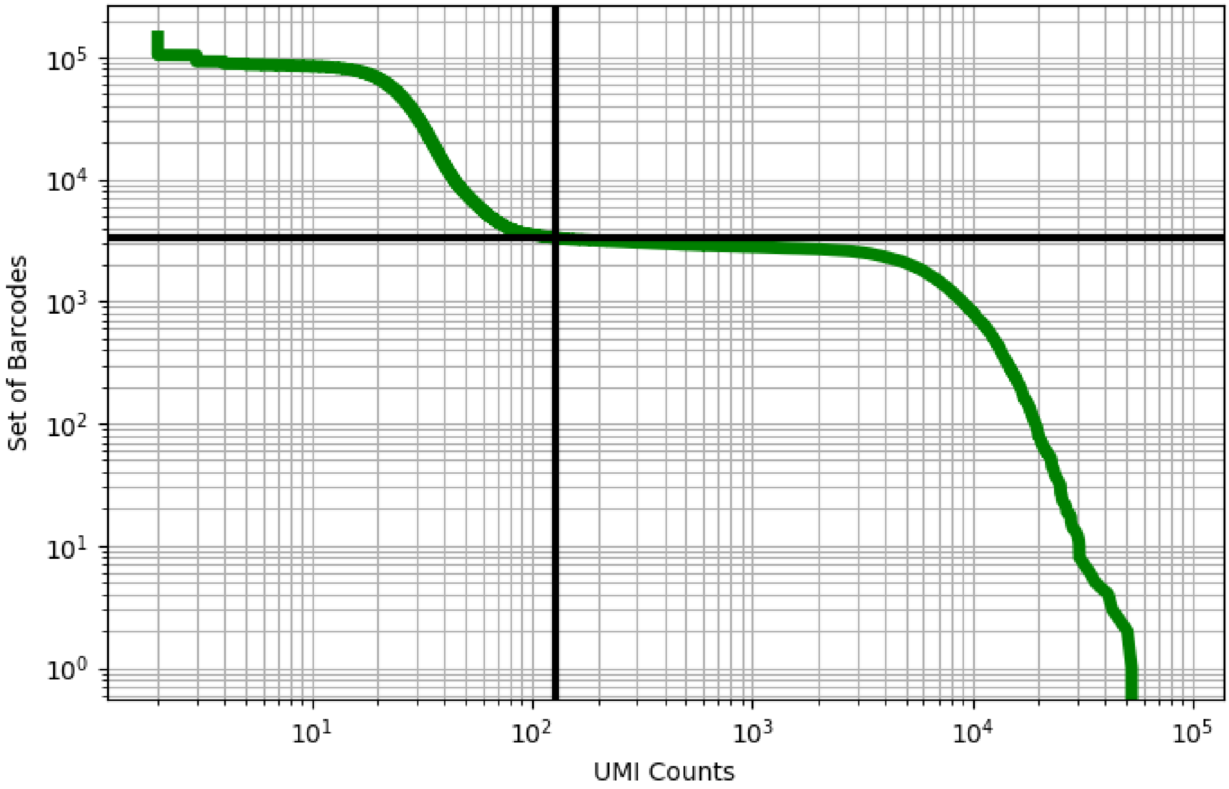


Figure 448: skin/ERS3861815

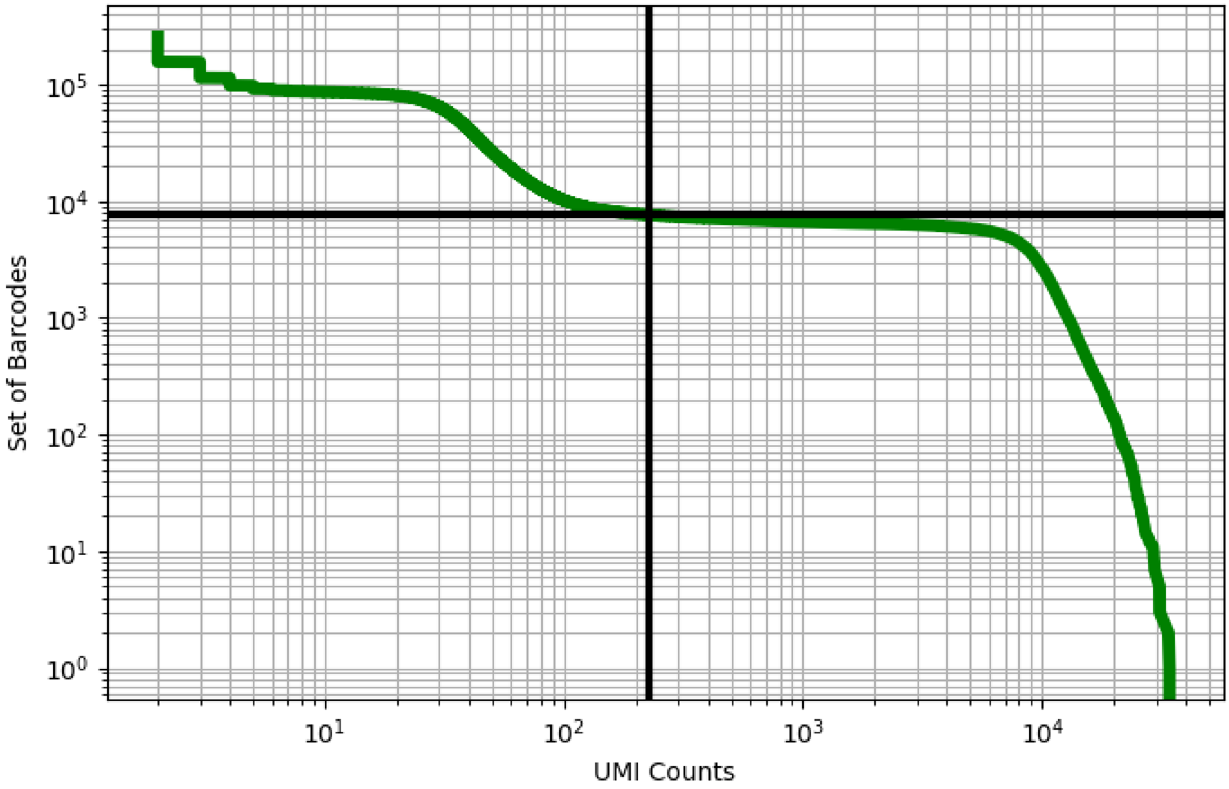


Figure 449: skin/ERS3861816

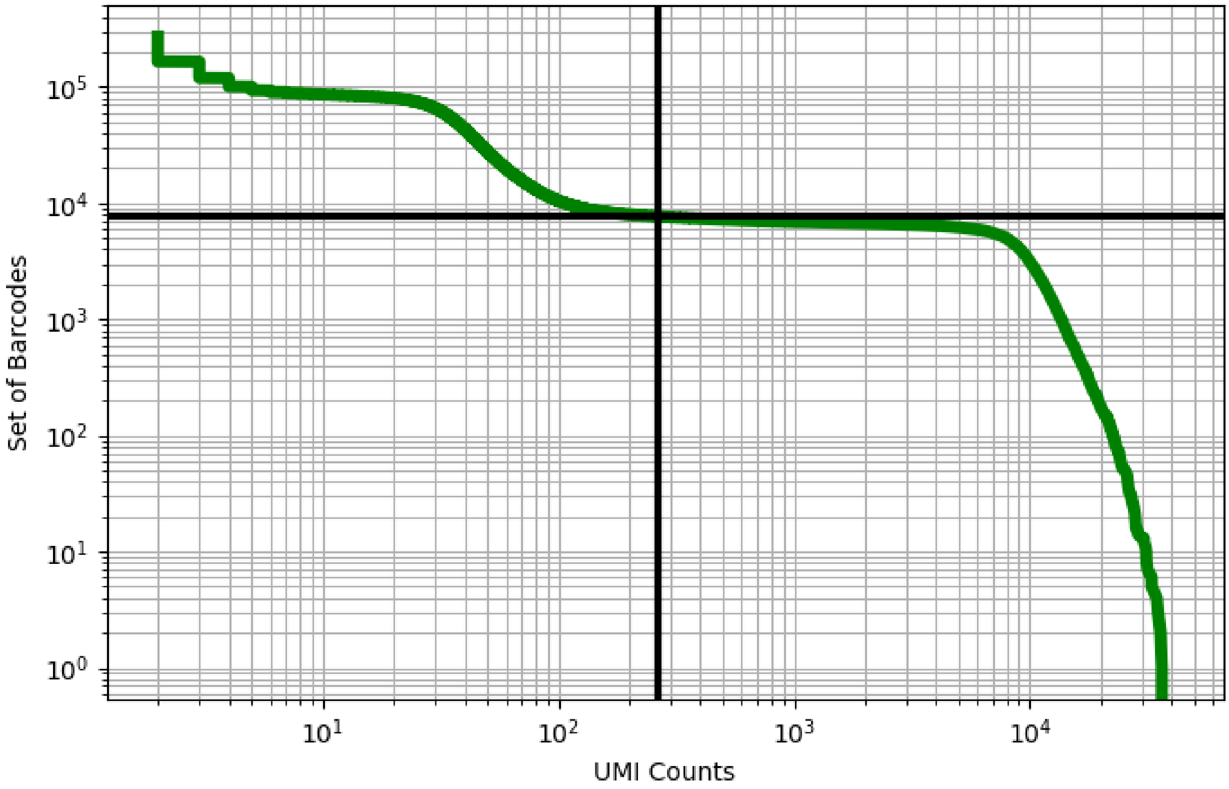


Figure 450: skin/ERS3861817

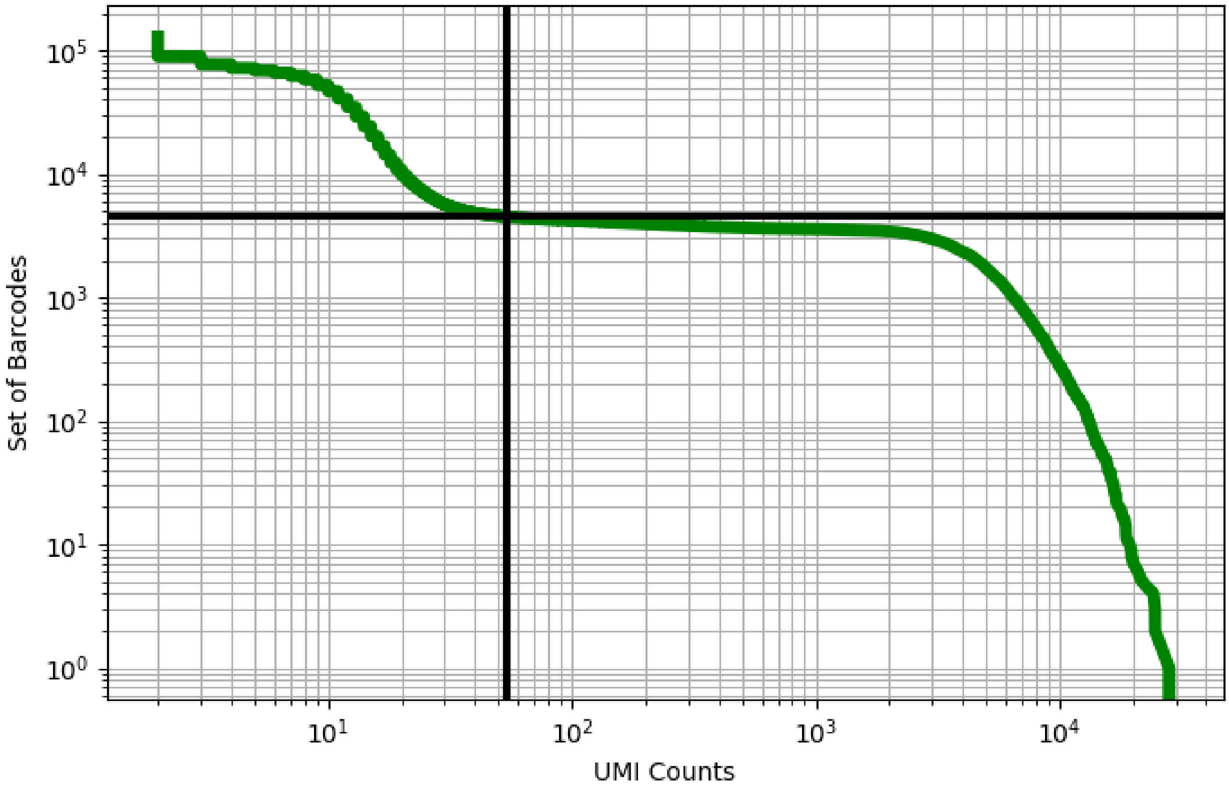


Figure 451: skin/ERS3861826

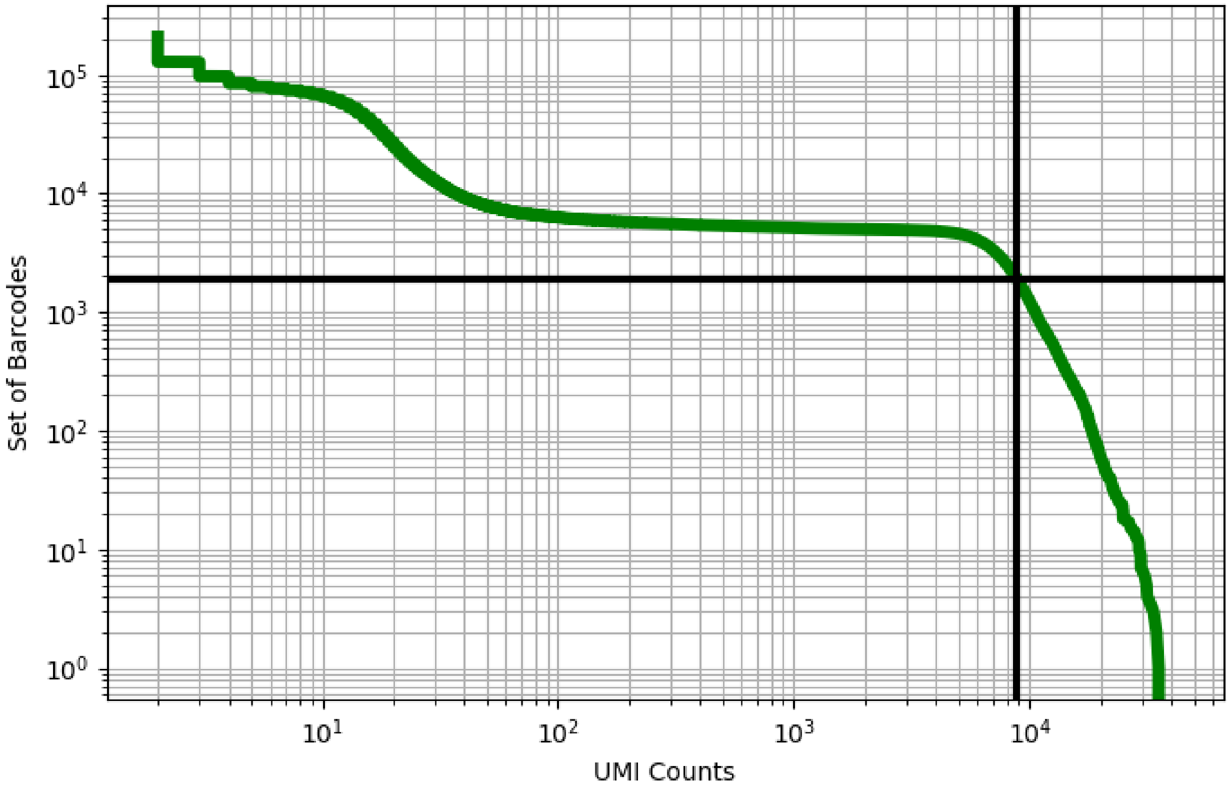


Figure 452: skin/ERS3861827

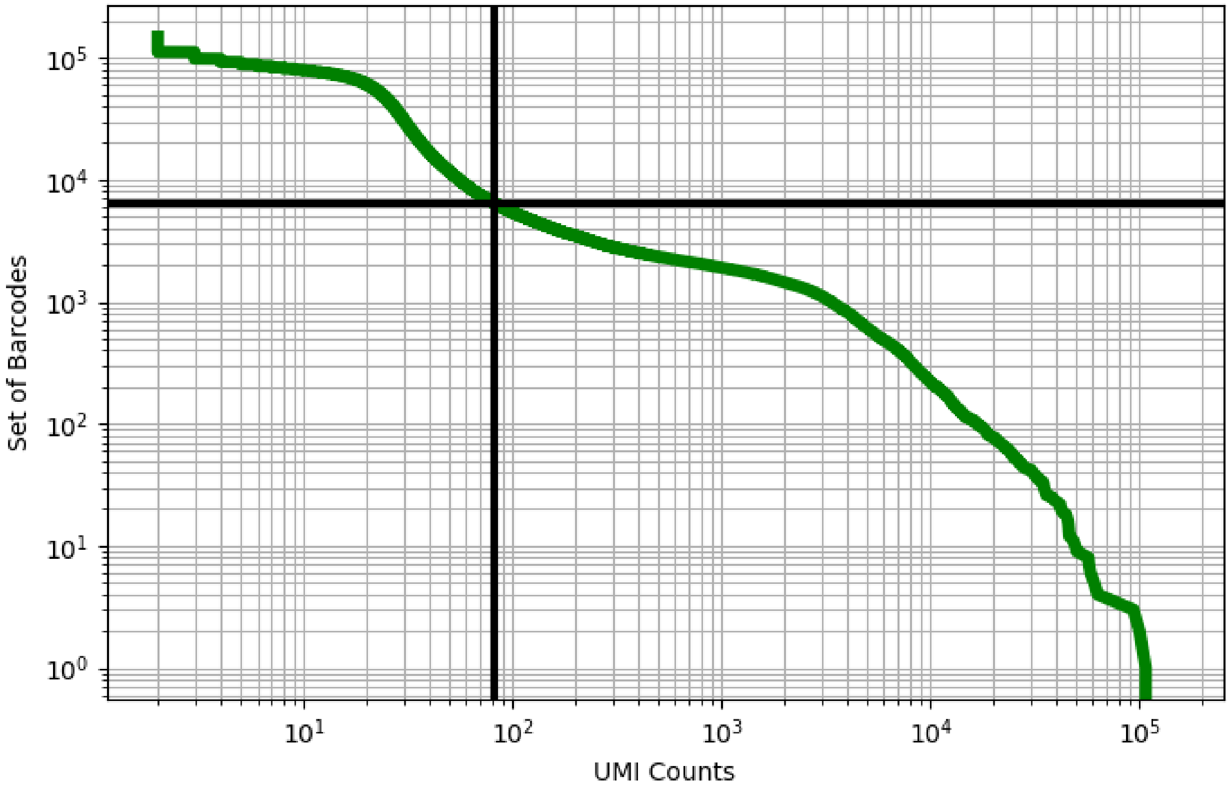


Figure 453: skin/GSM3667327

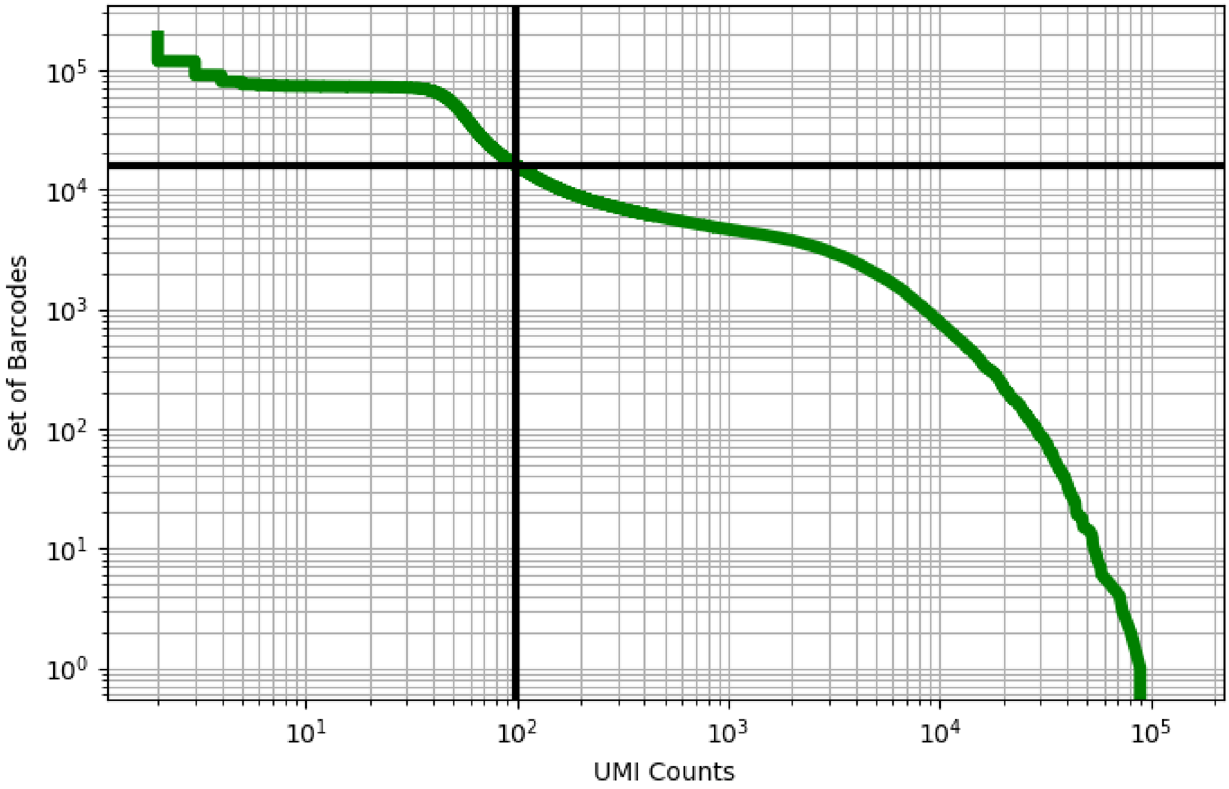


Figure 454: skin/GSM3667328

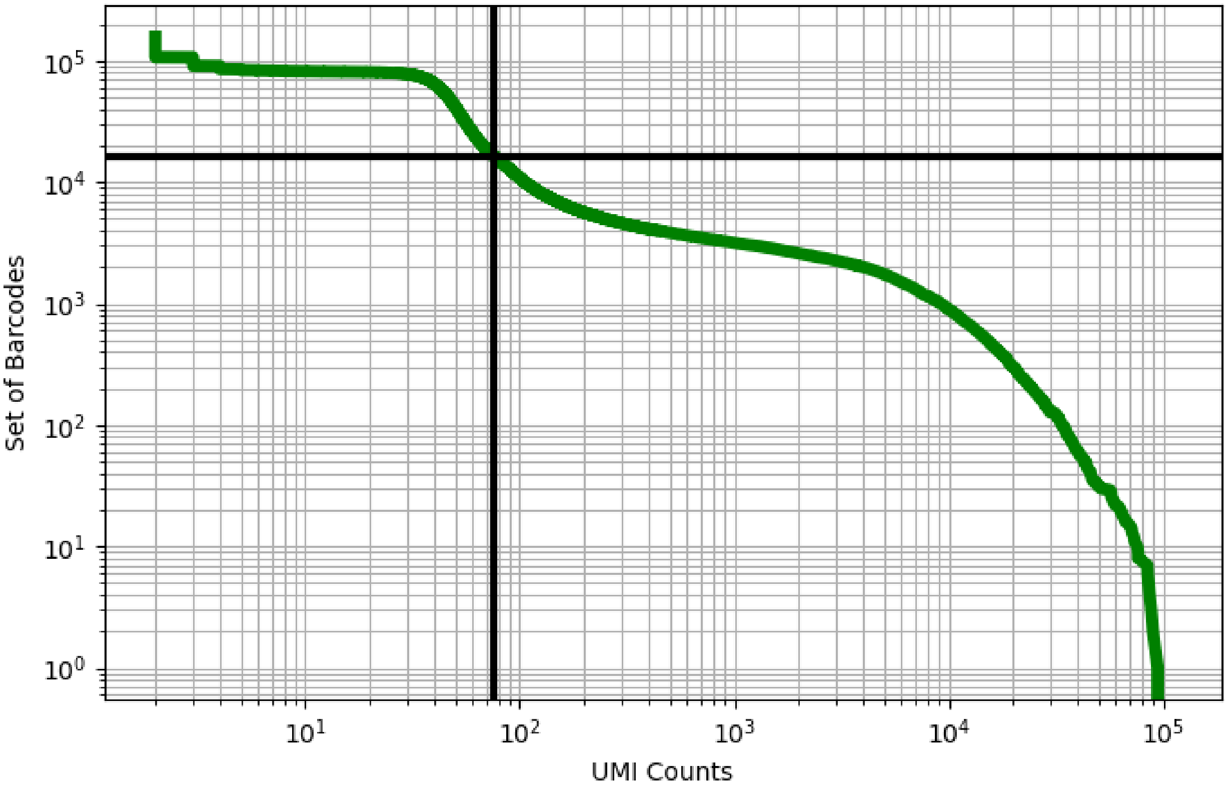


Figure 455: skin/GSM3667329

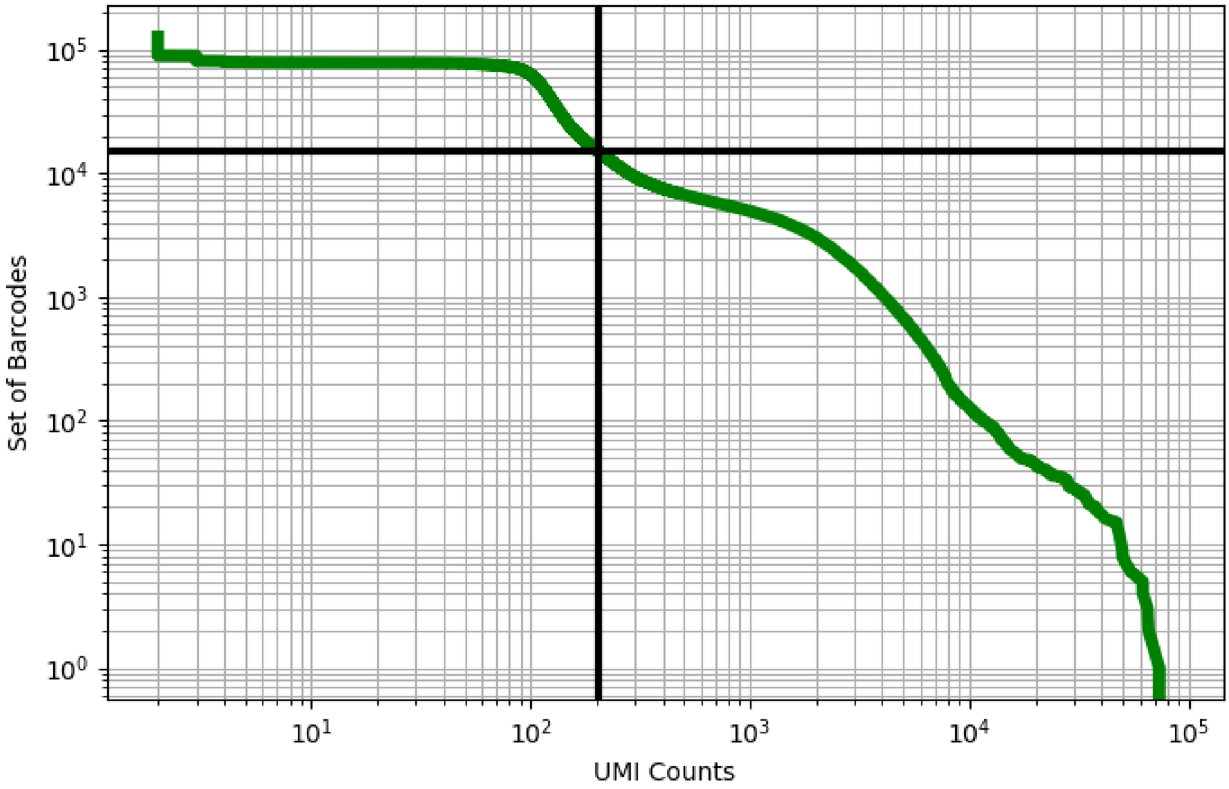


Figure 456: skin/GSM3758115

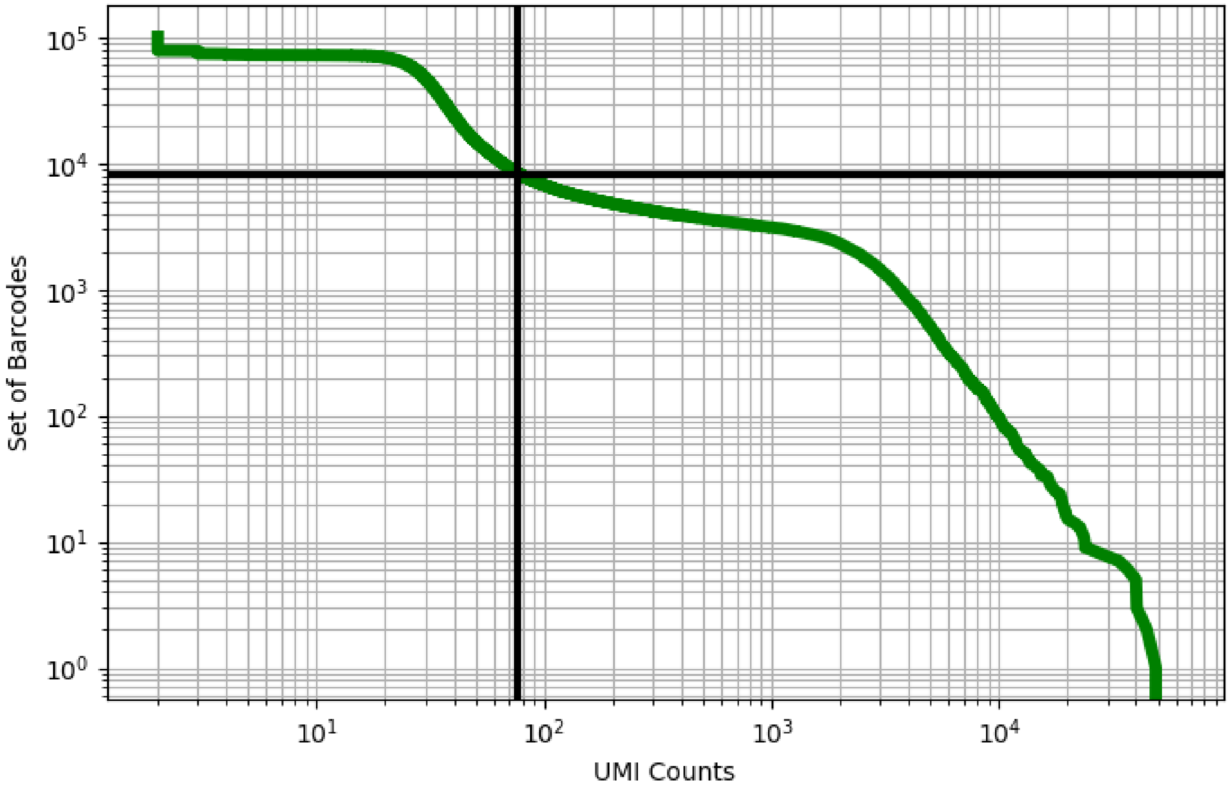


Figure 457: skin/GSM3758116

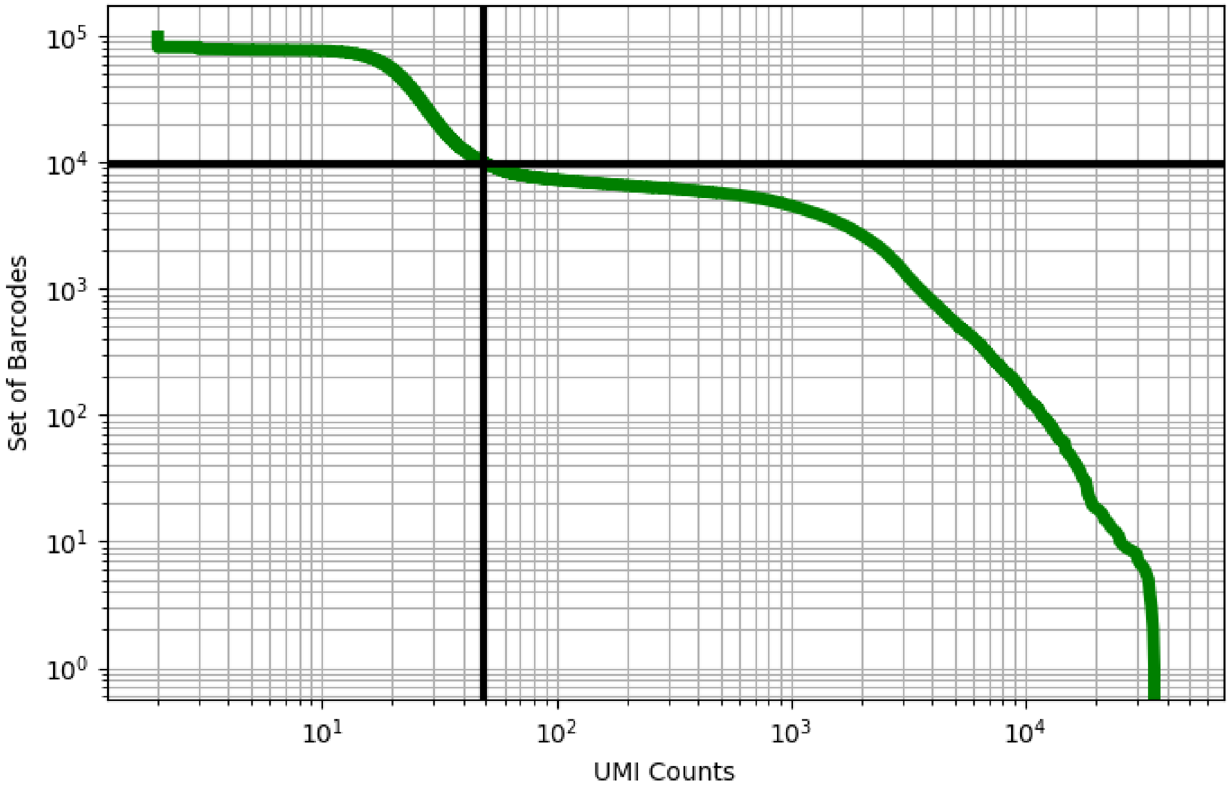


Figure 458: skin/GSM3758117

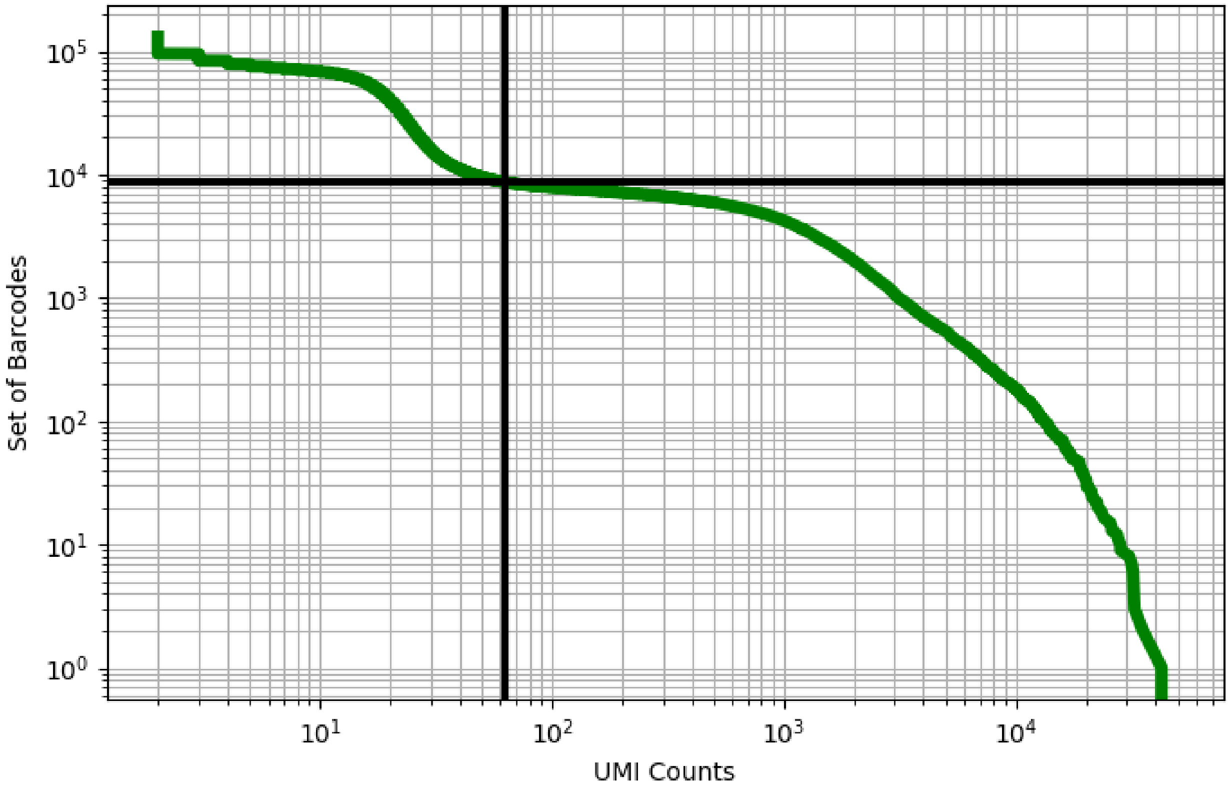


Figure 459: skin/GSM3758118

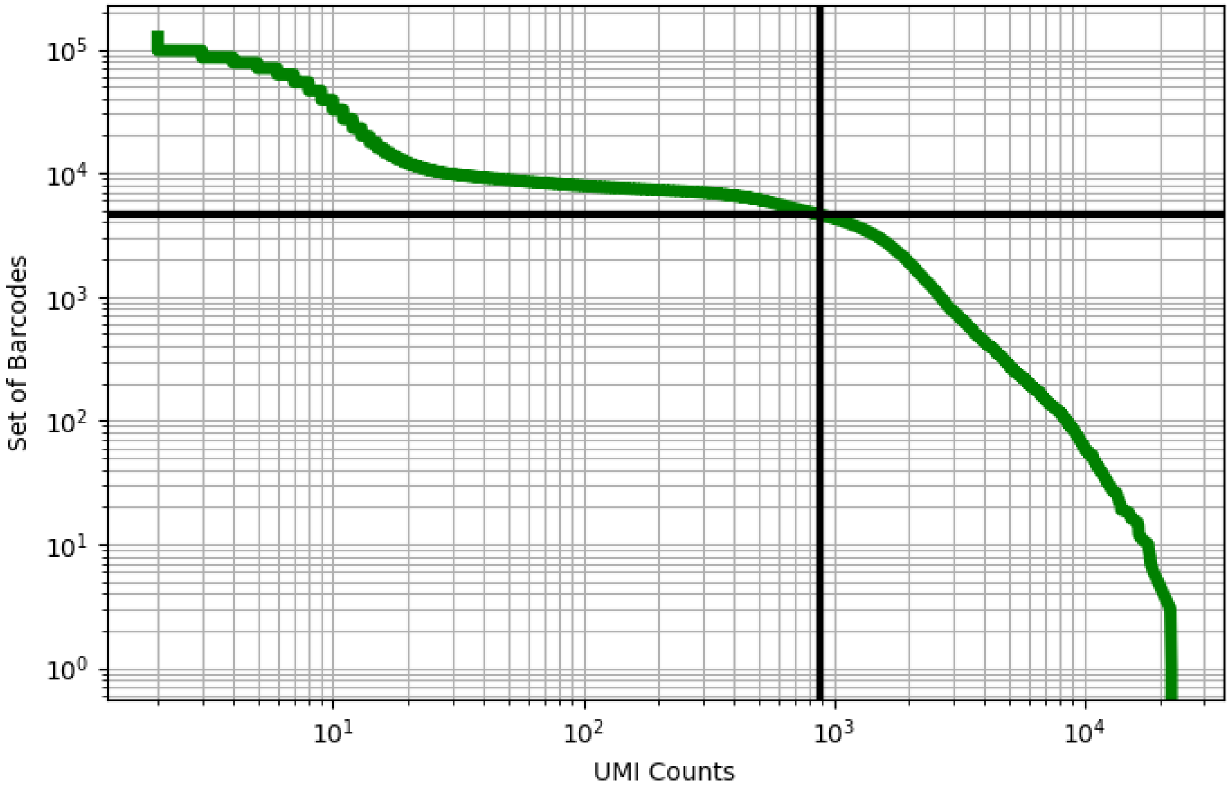


Figure 460: skin/GSM3758119

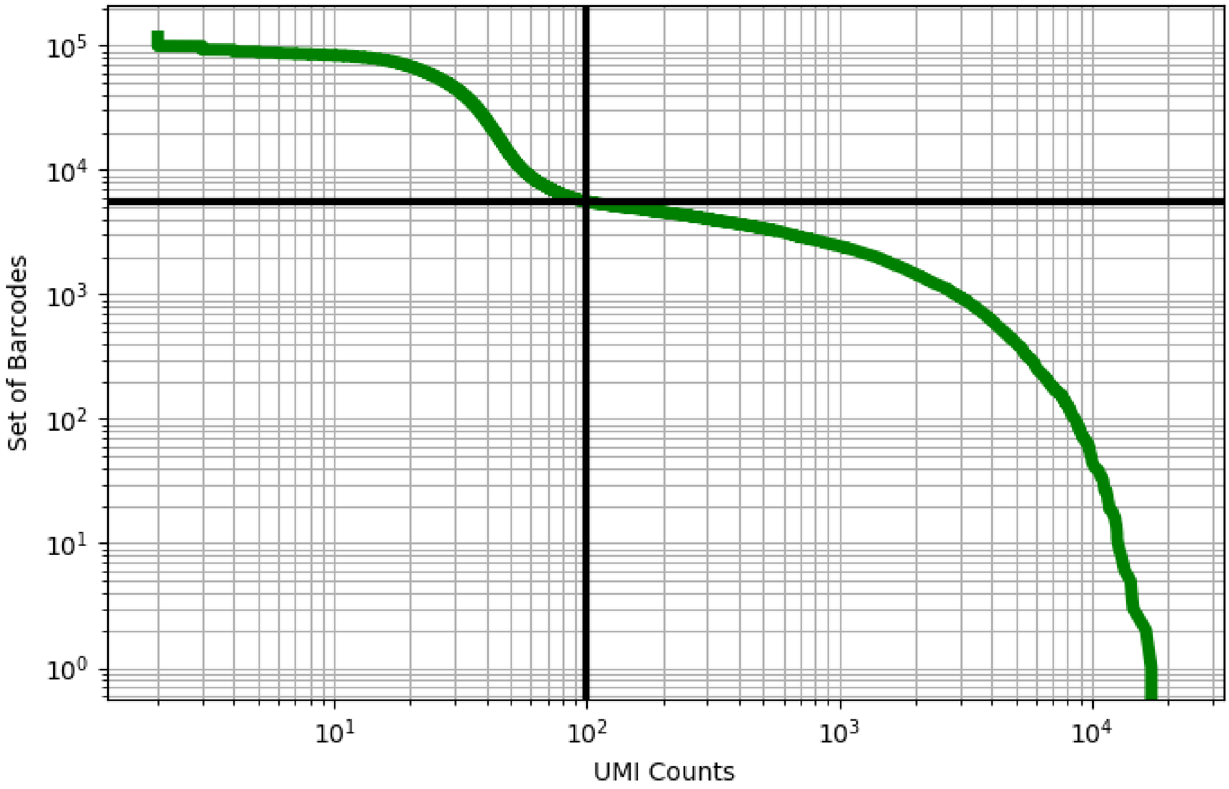


Figure 461: stomach/GSM3954946

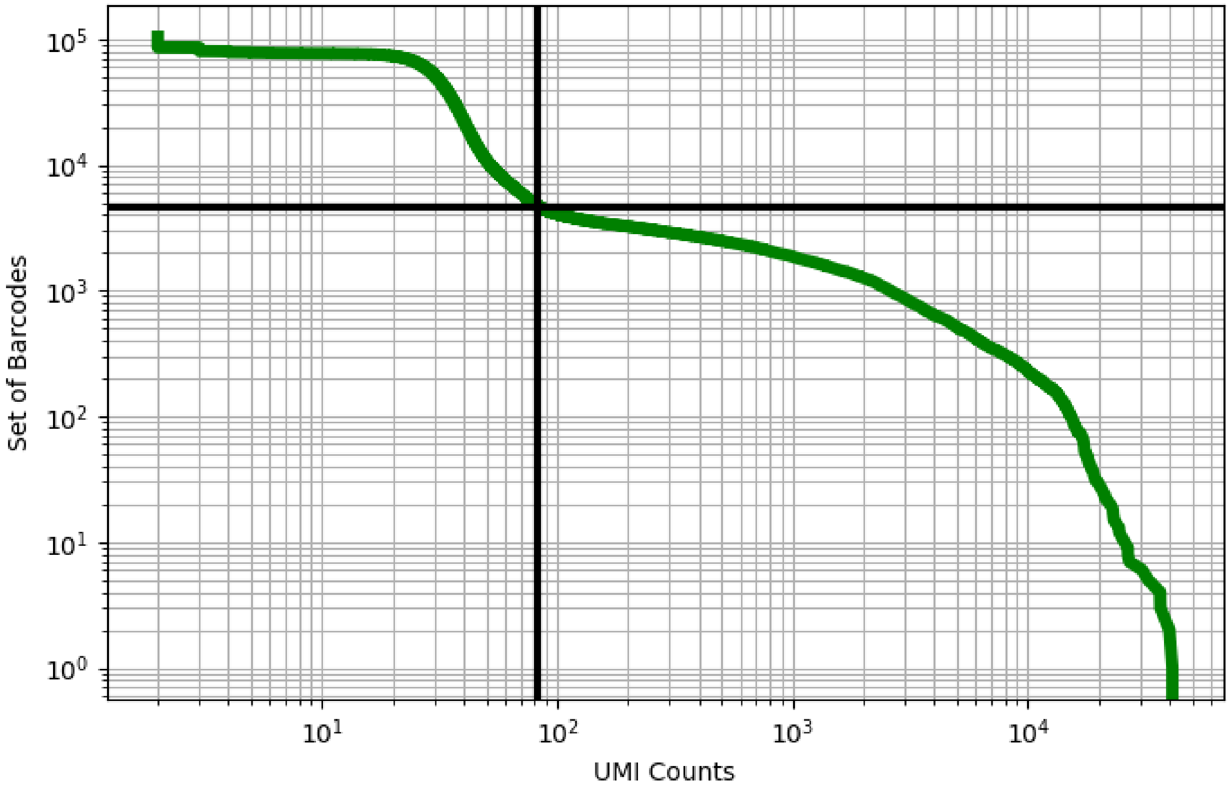


Figure 462: stomach/GSM3954947

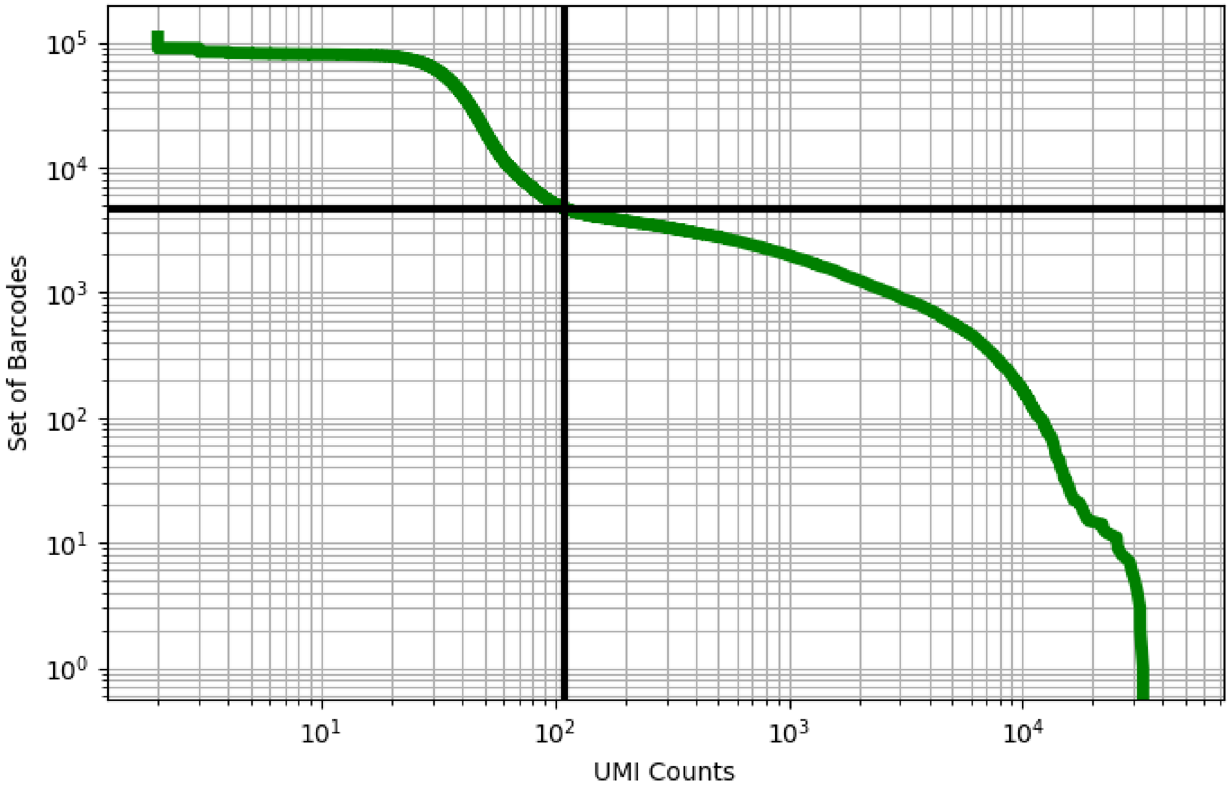


Figure 463: stomach/GSM3954948

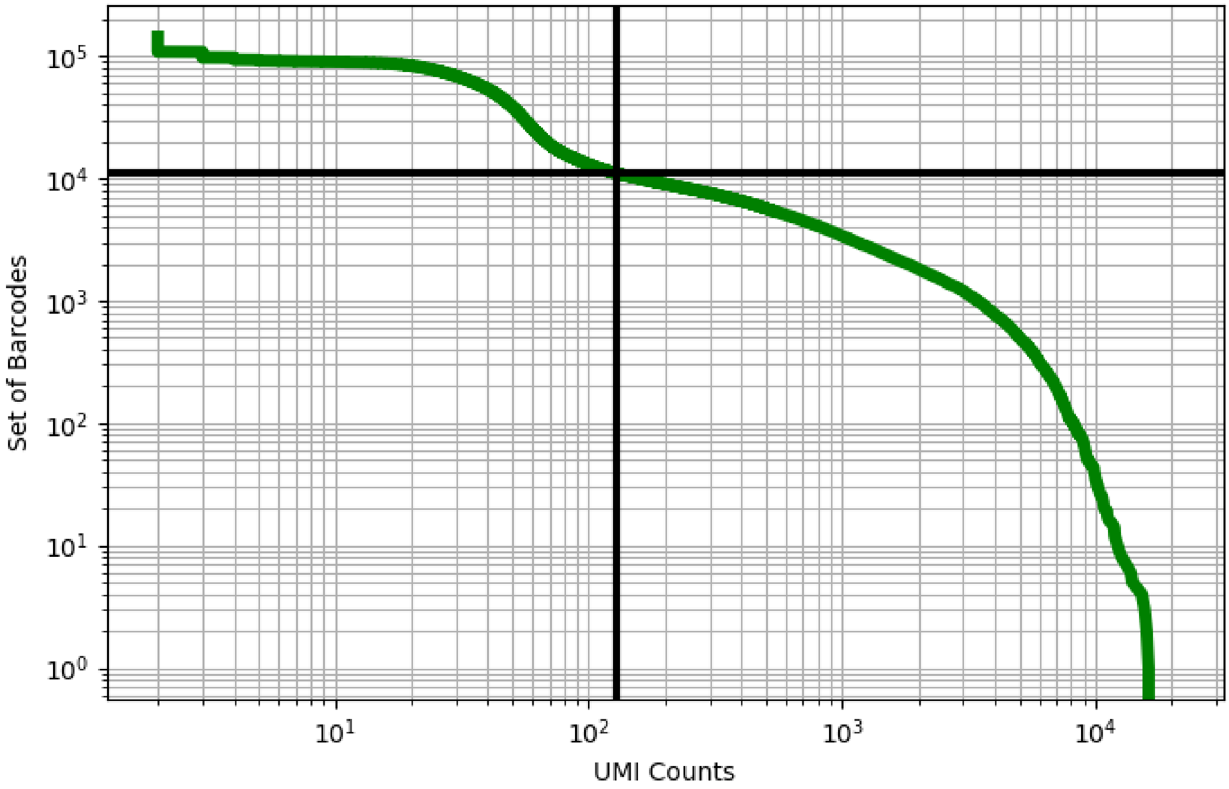


Figure 464: stomach/GSM3954949

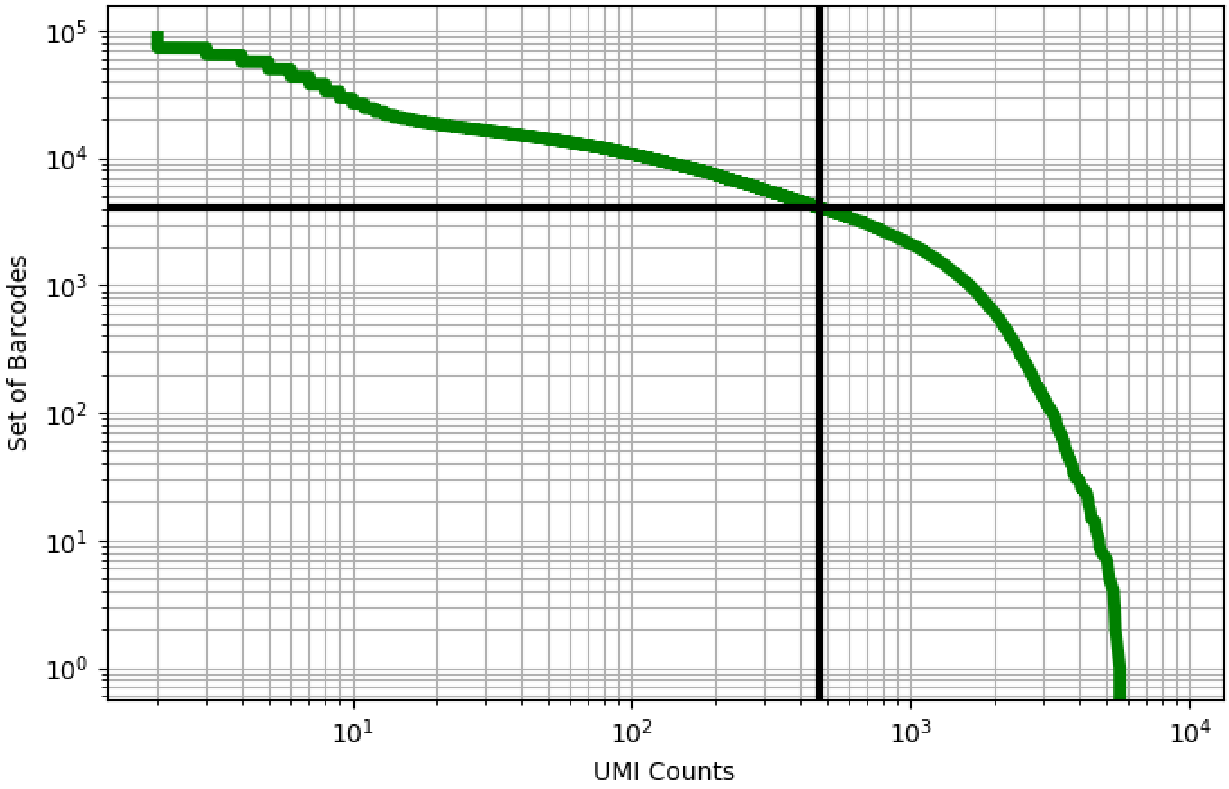


Figure 465: stomach/GSM3954950

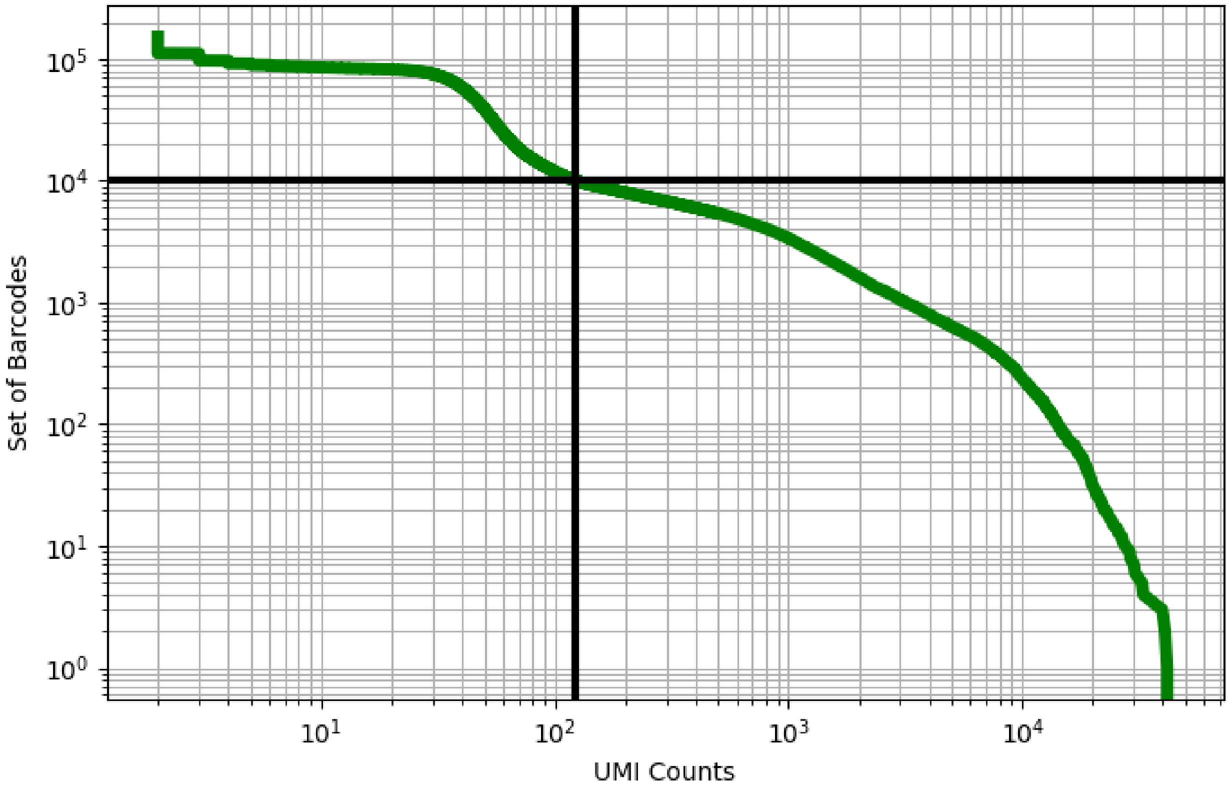


Figure 466: stomach/GSM3954952

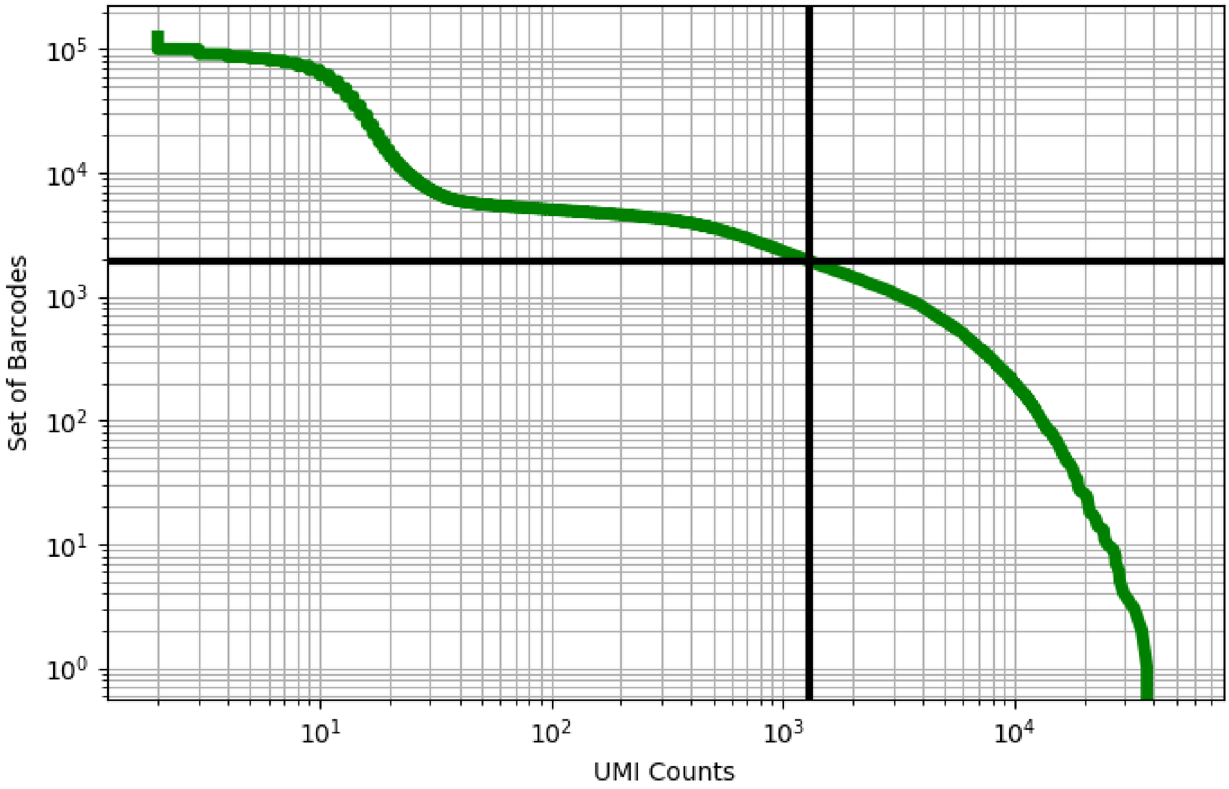


Figure 467: stomach/GSM3954953

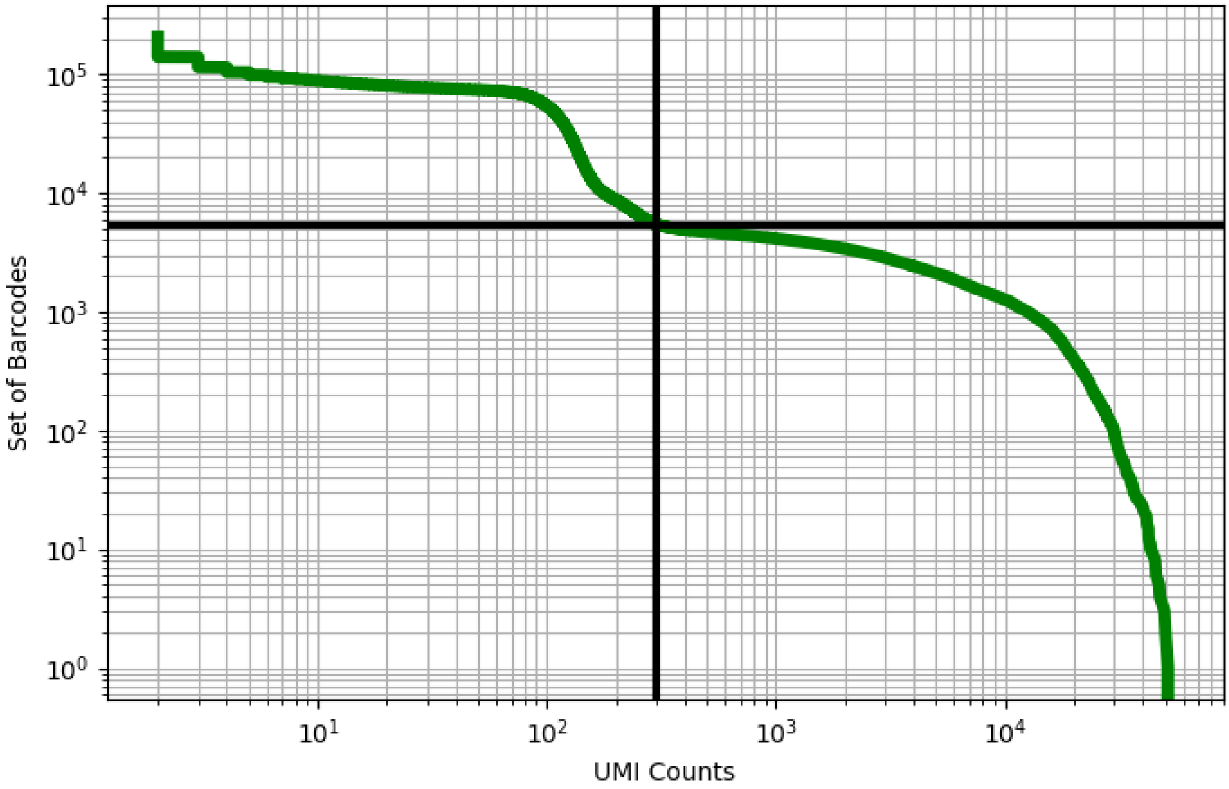


Figure 468: stomach/GSM3954954

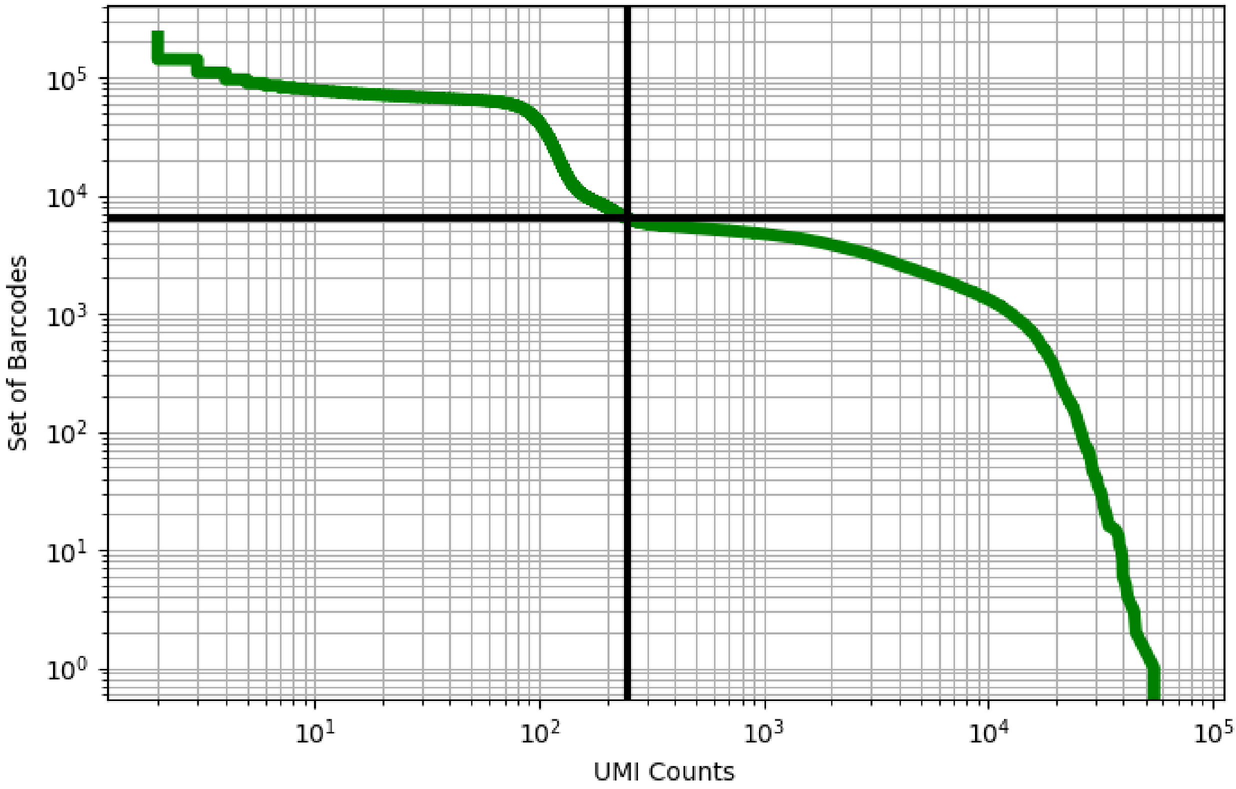


Figure 469: stomach/GSM3954955

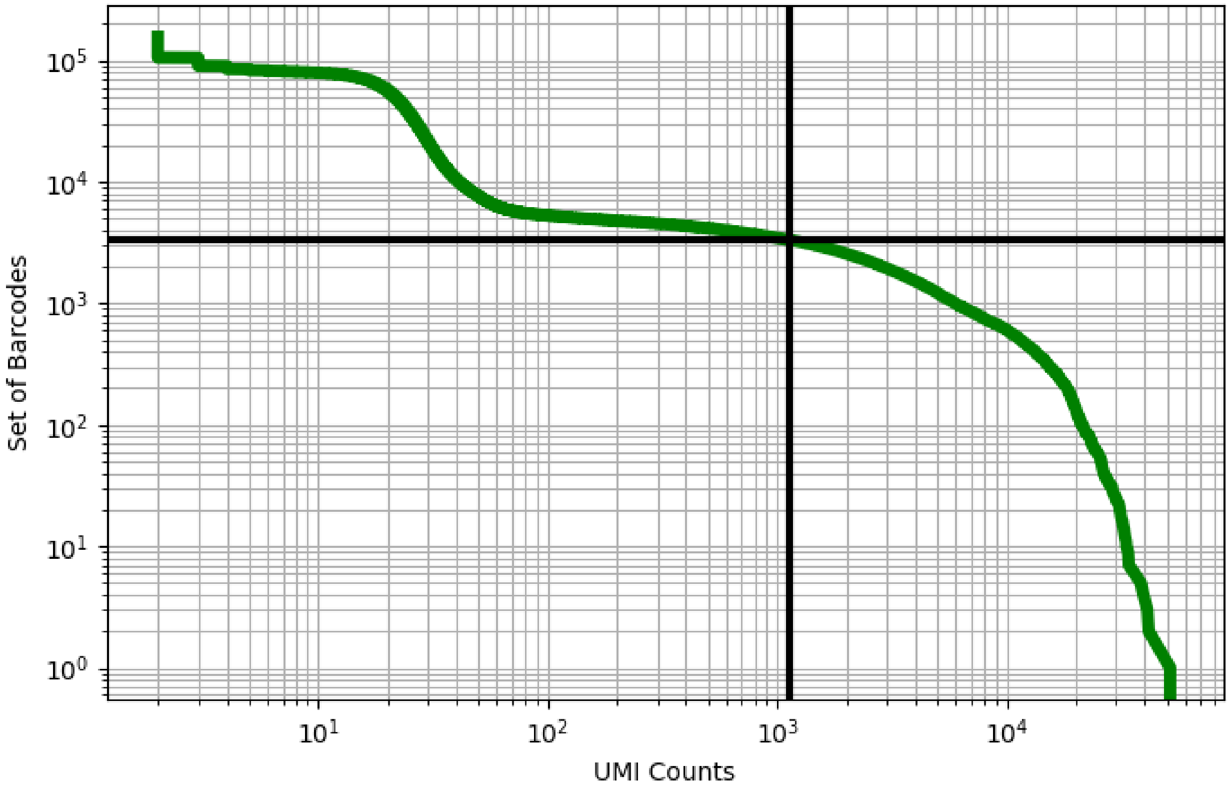


Figure 470: stomach/GSM3954956

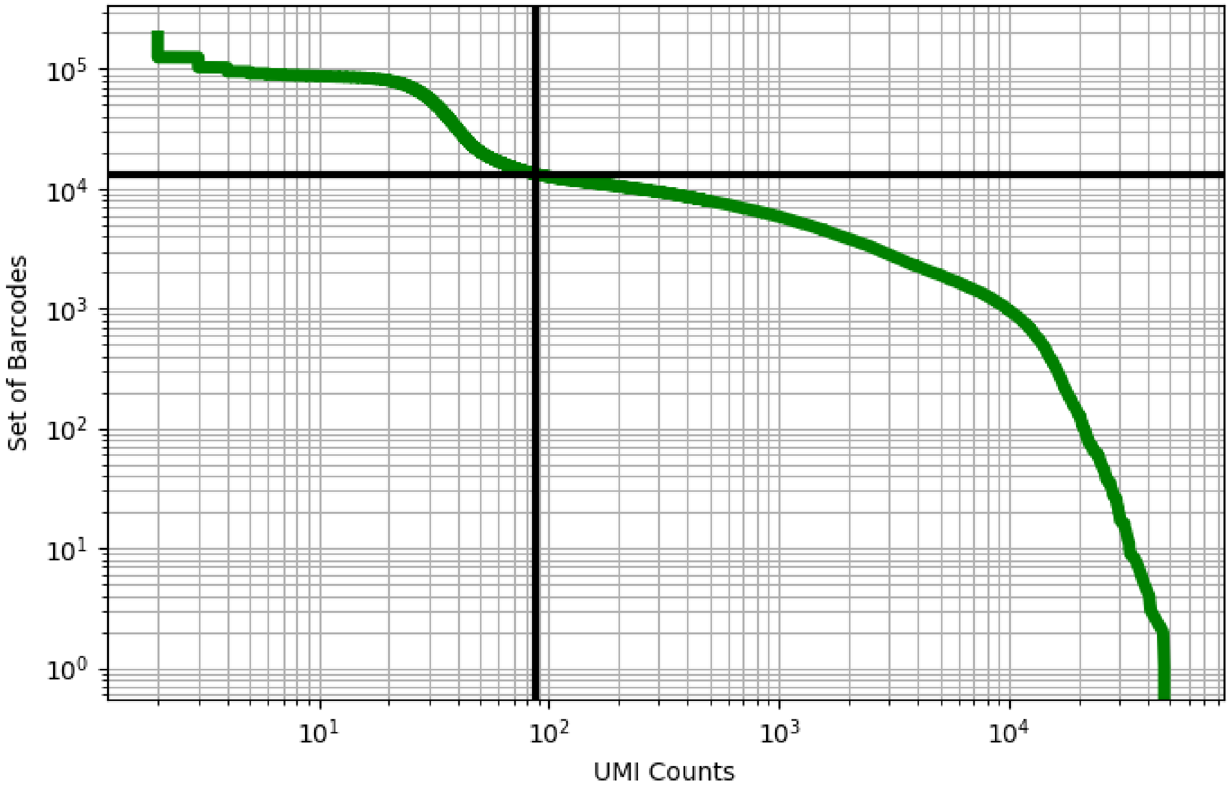


Figure 471: stomach/GSM3954957

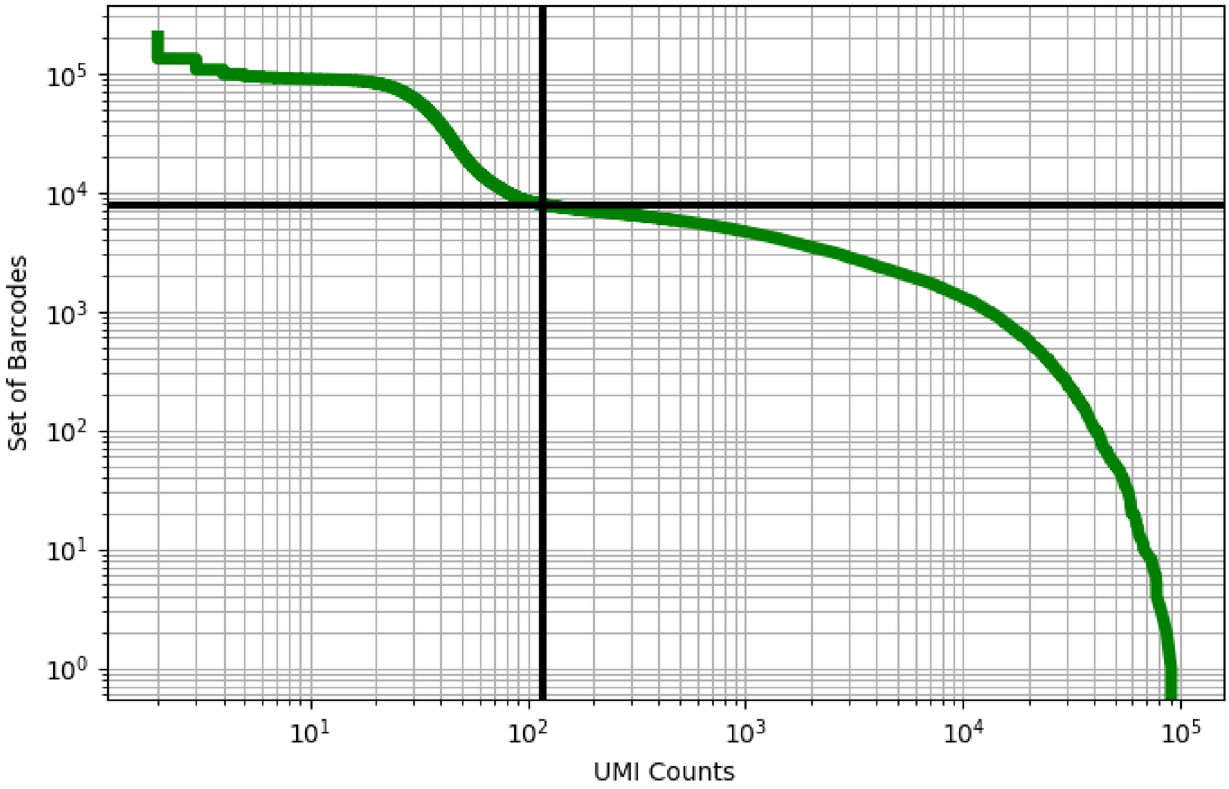


Figure 472: stomach/GSM3954958

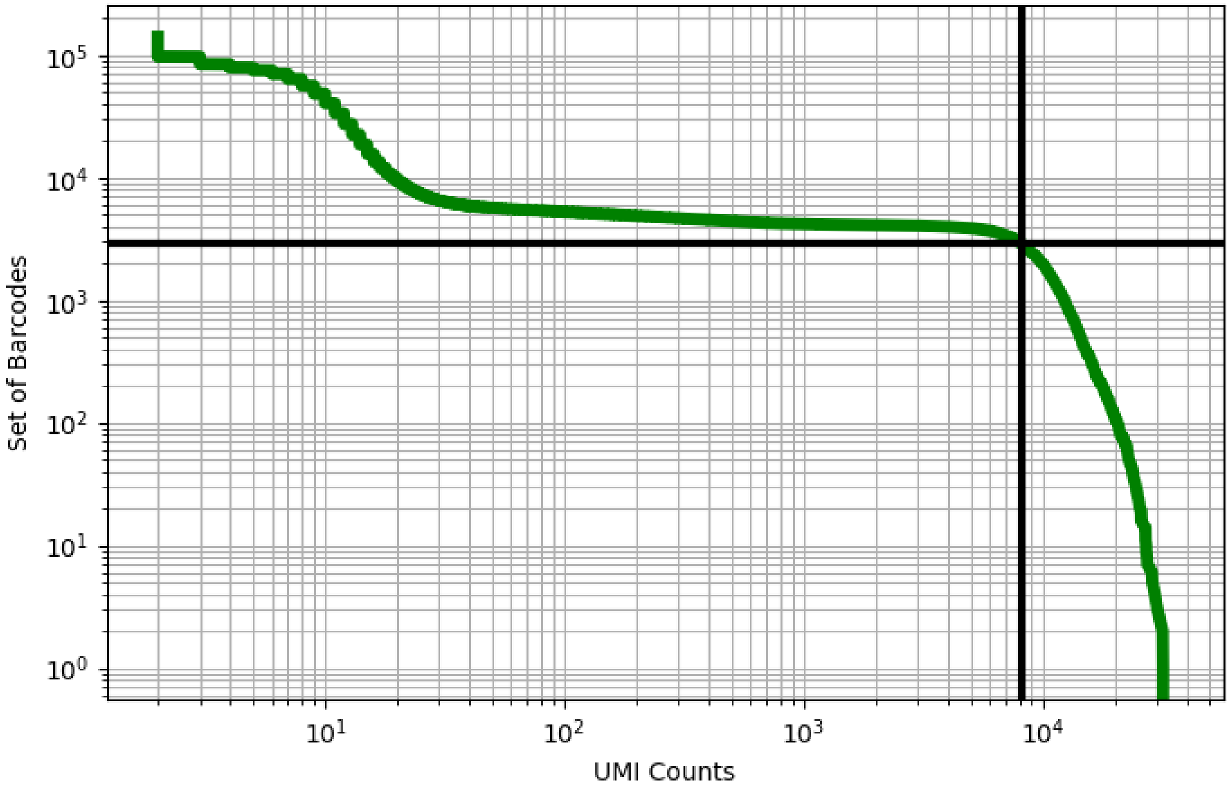


Figure 473: testis/GSM2928377

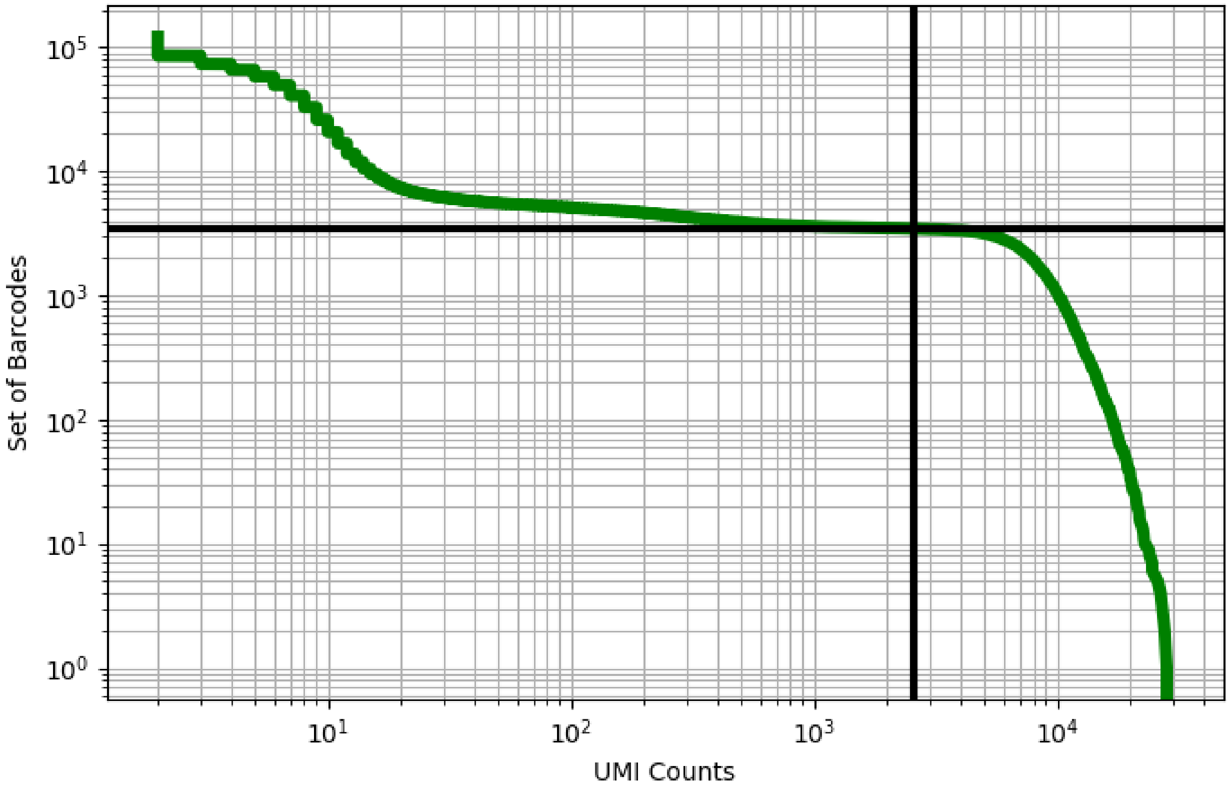


Figure 474: testis/GSM2928378

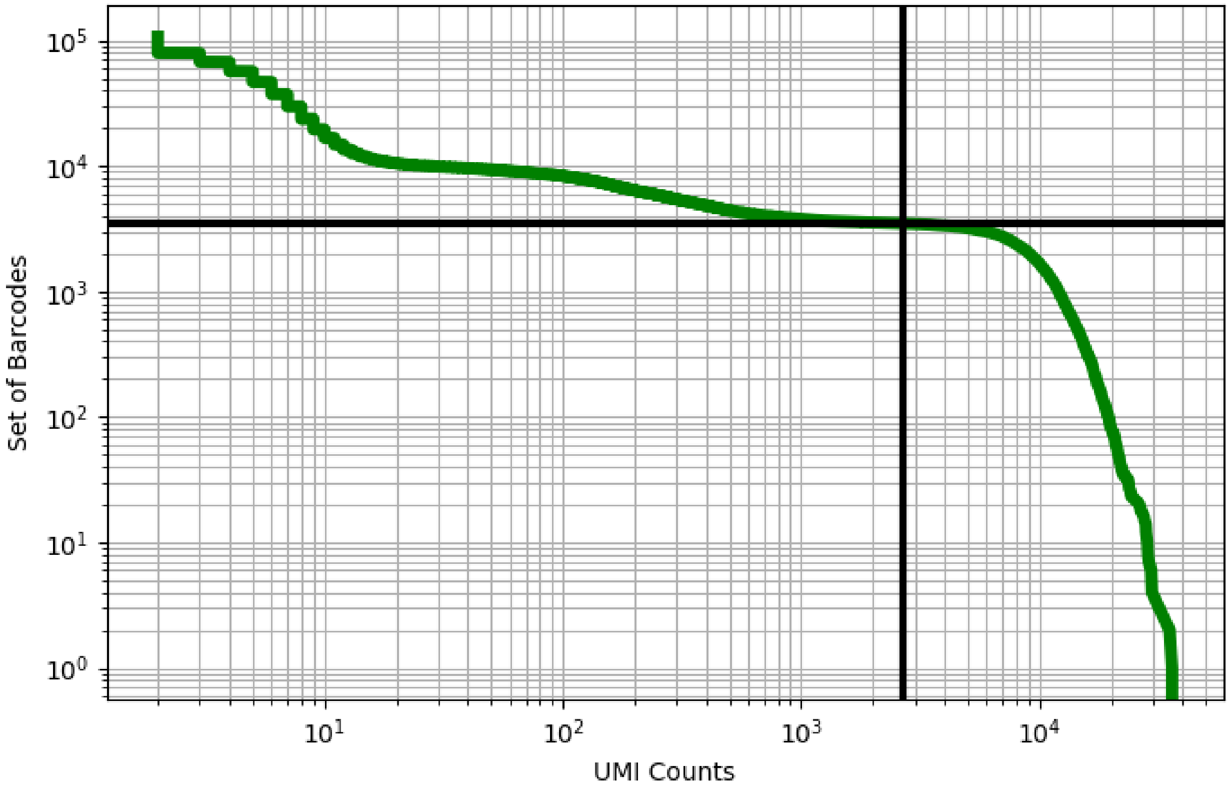


Figure 475: testis/GSM2928379

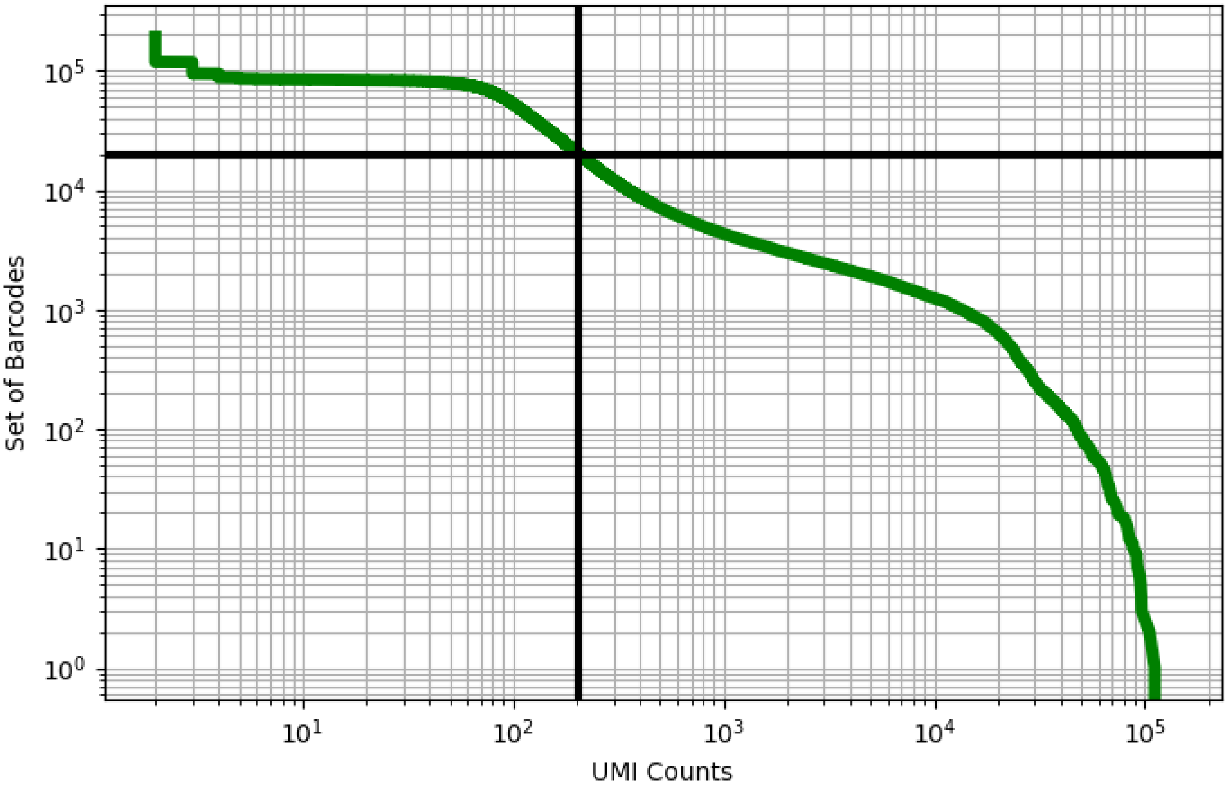


Figure 476: testis/GSM2928380

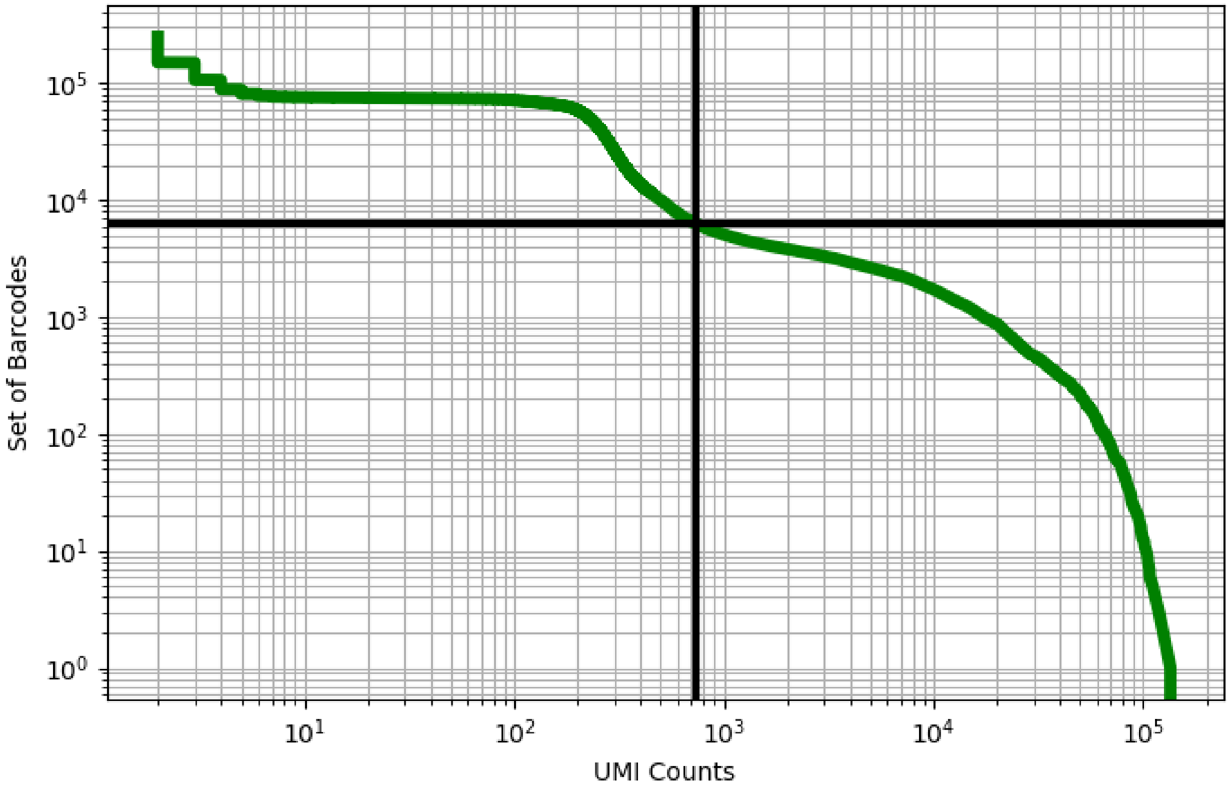


Figure 477: testis/GSM2928381

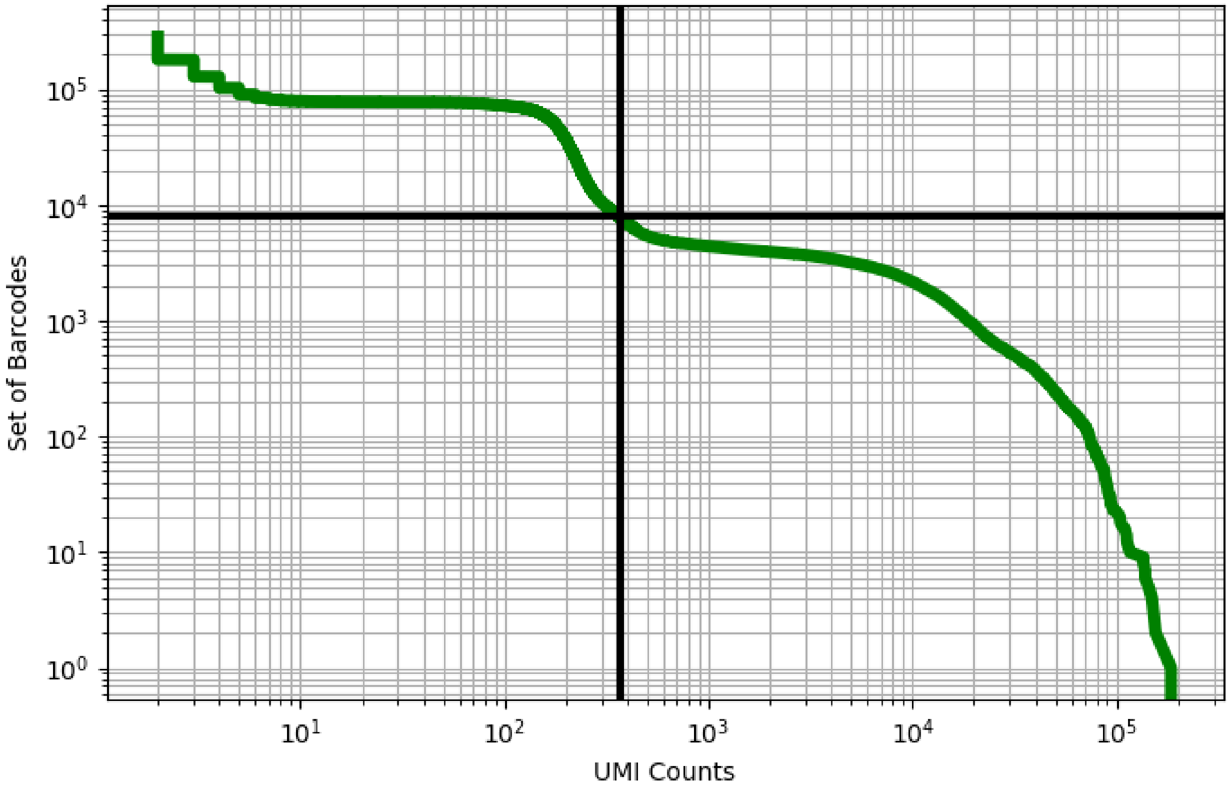


Figure 478: testis/GSM2928382

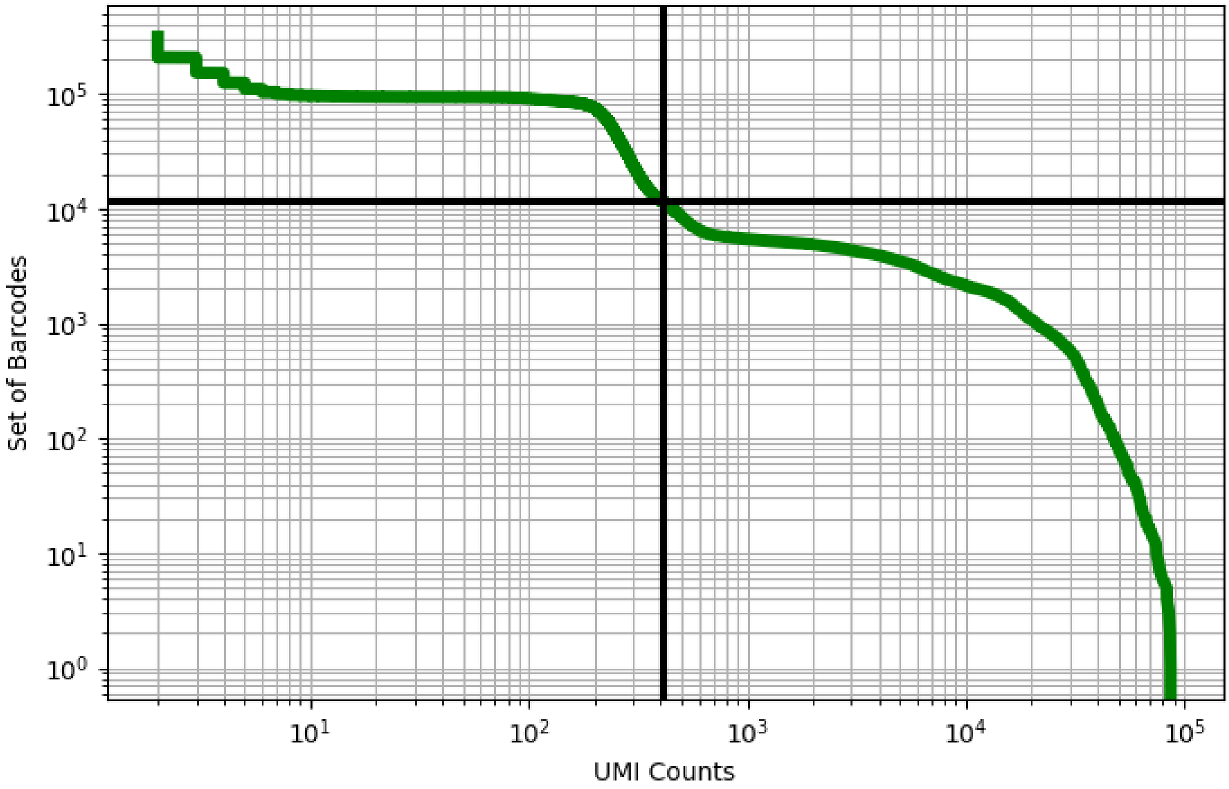


Figure 479: testis/GSM2928384

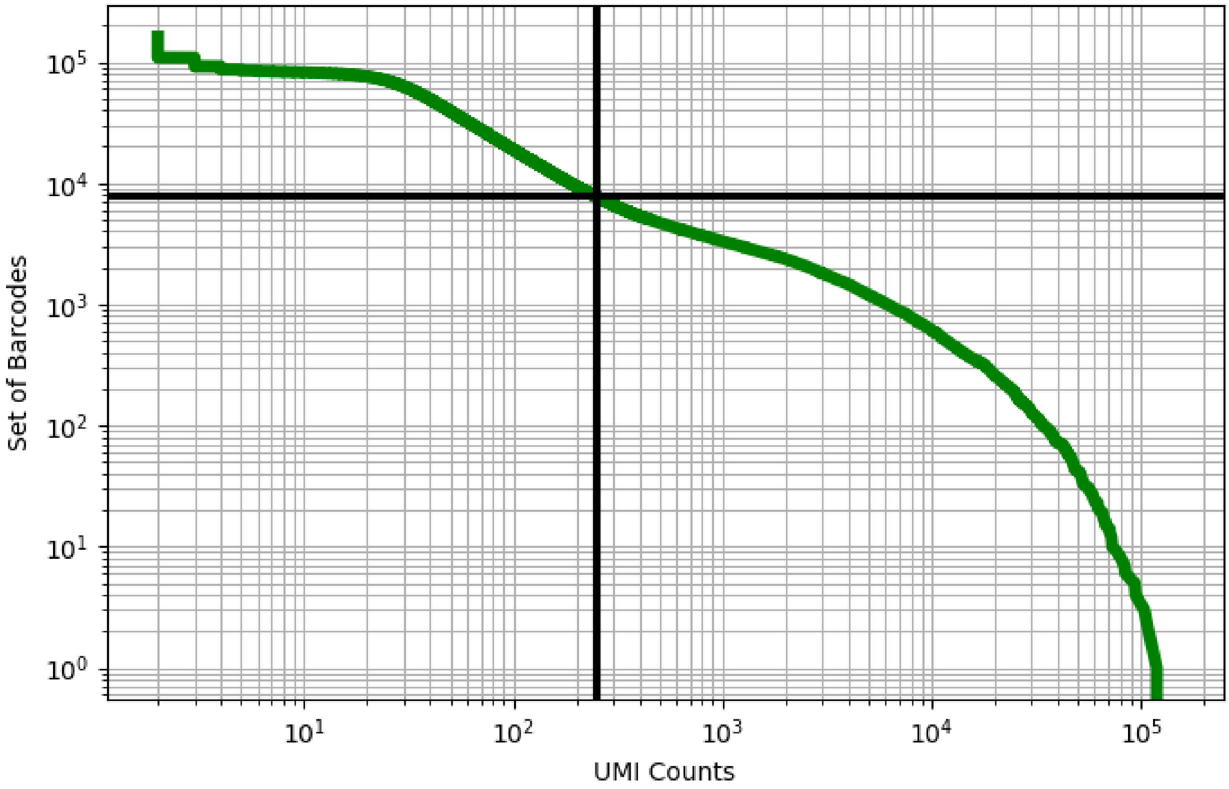


Figure 480: testis/GSM3052919

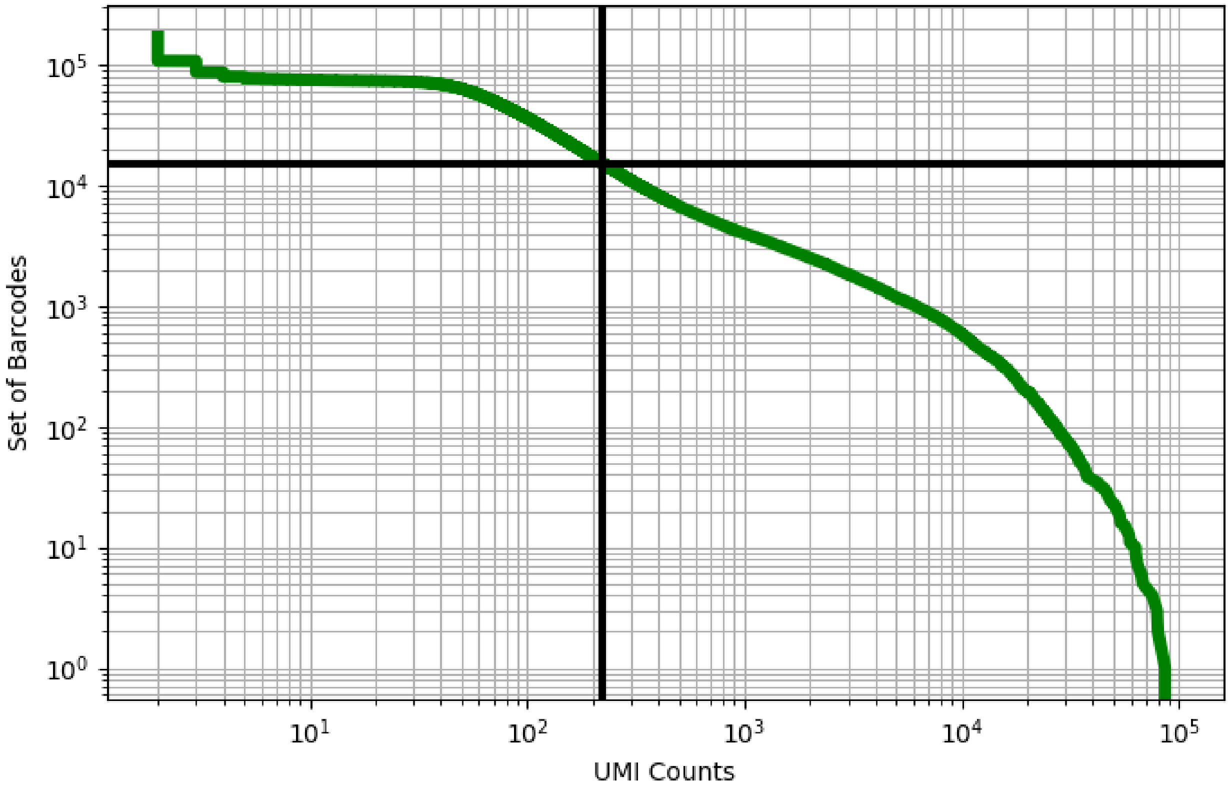


Figure 481: testis/GSM3052921

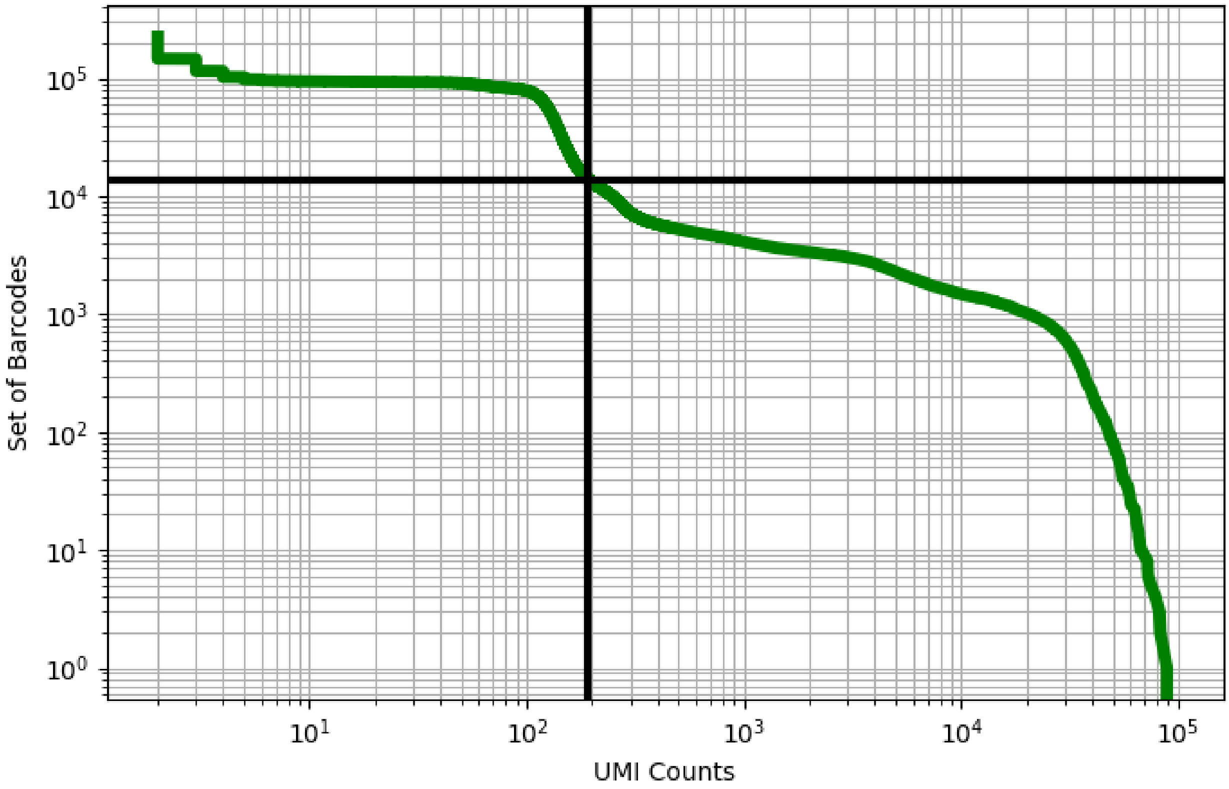


Figure 482: testis/GSM3302524

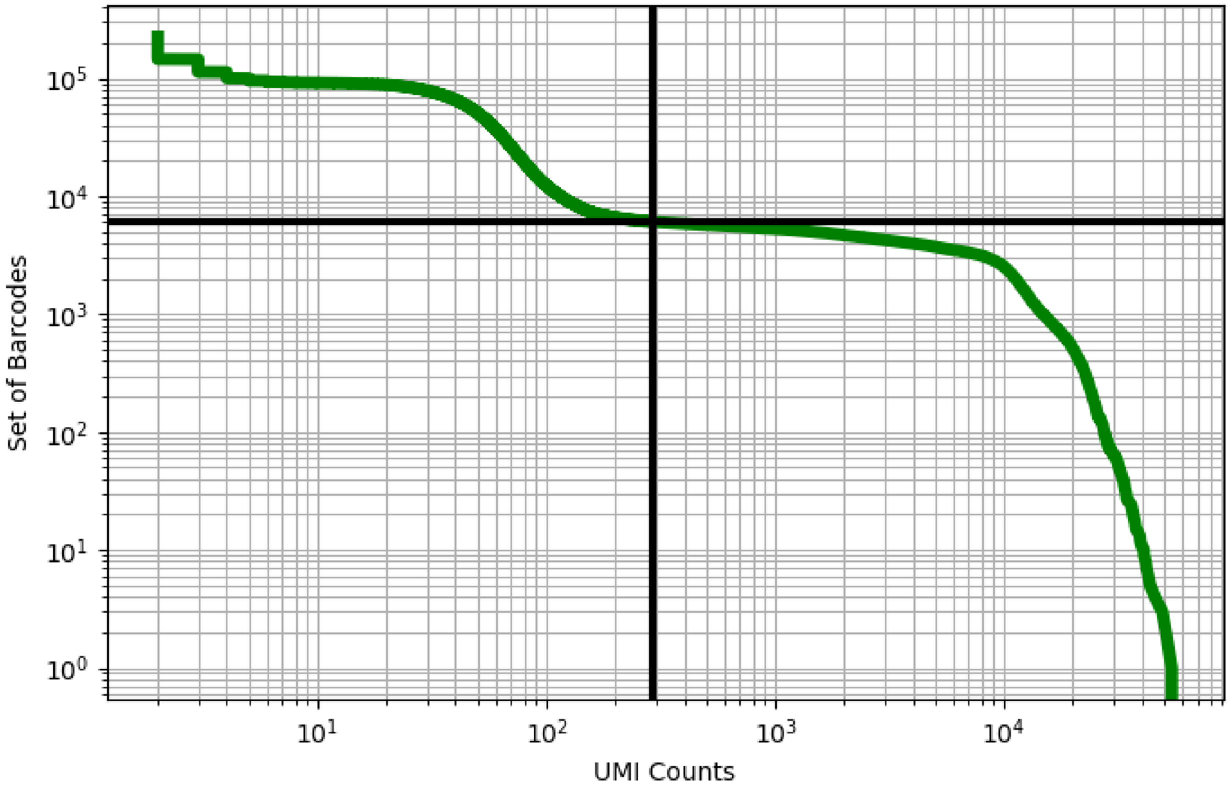


Figure 483: testis/GSM3302525

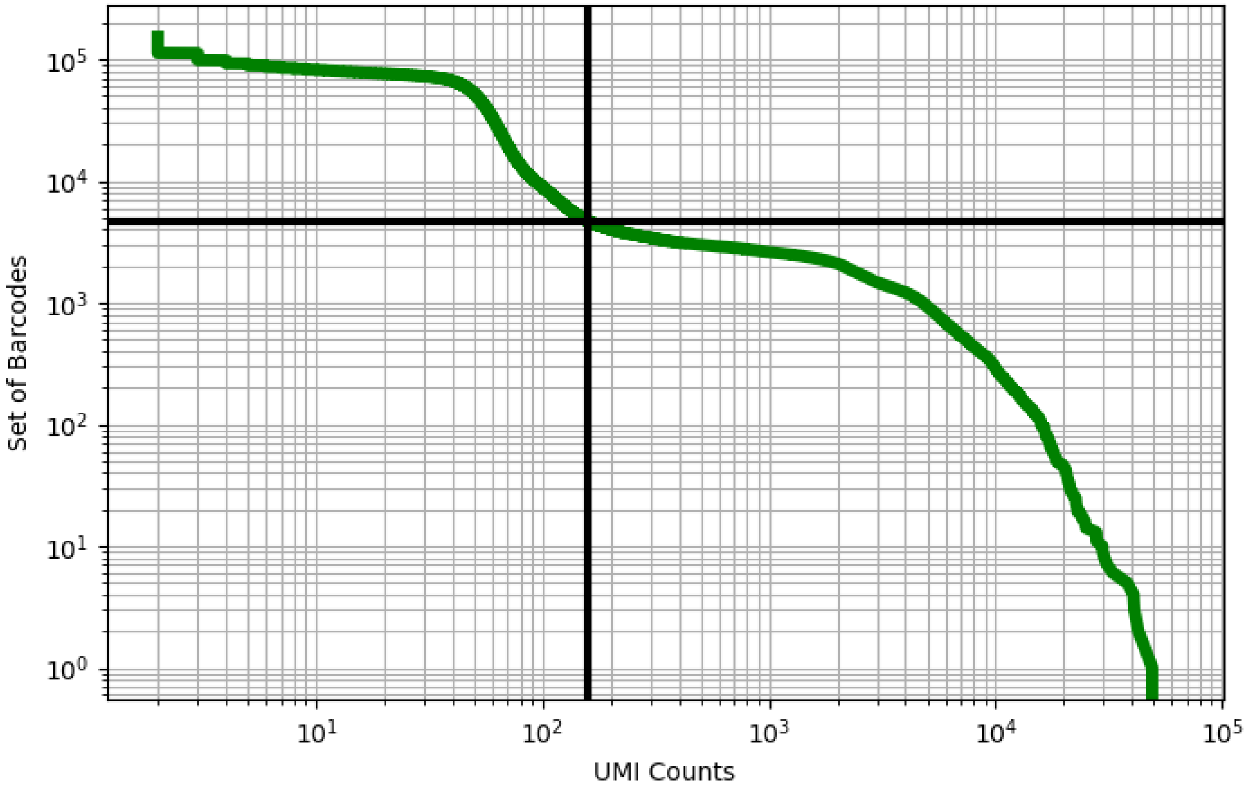


Figure 484: testis/GSM3402078

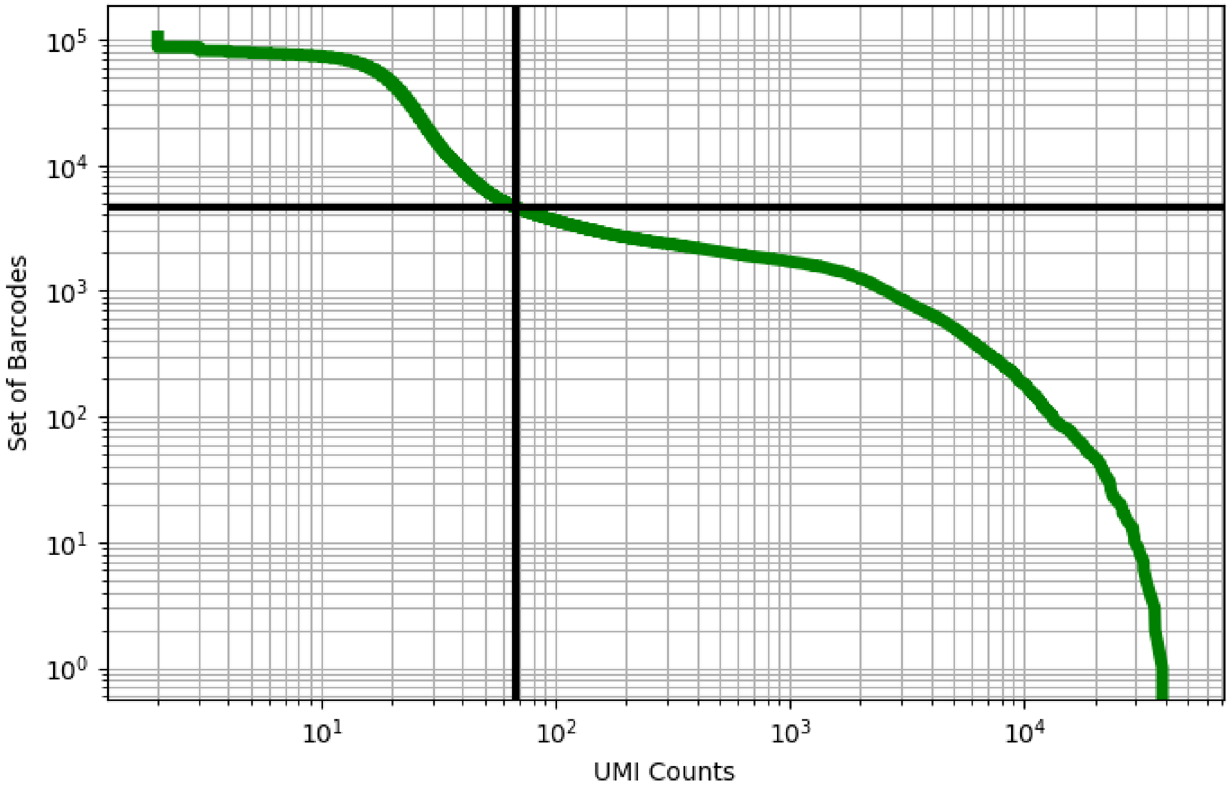


Figure 485: testis/GSM3402080

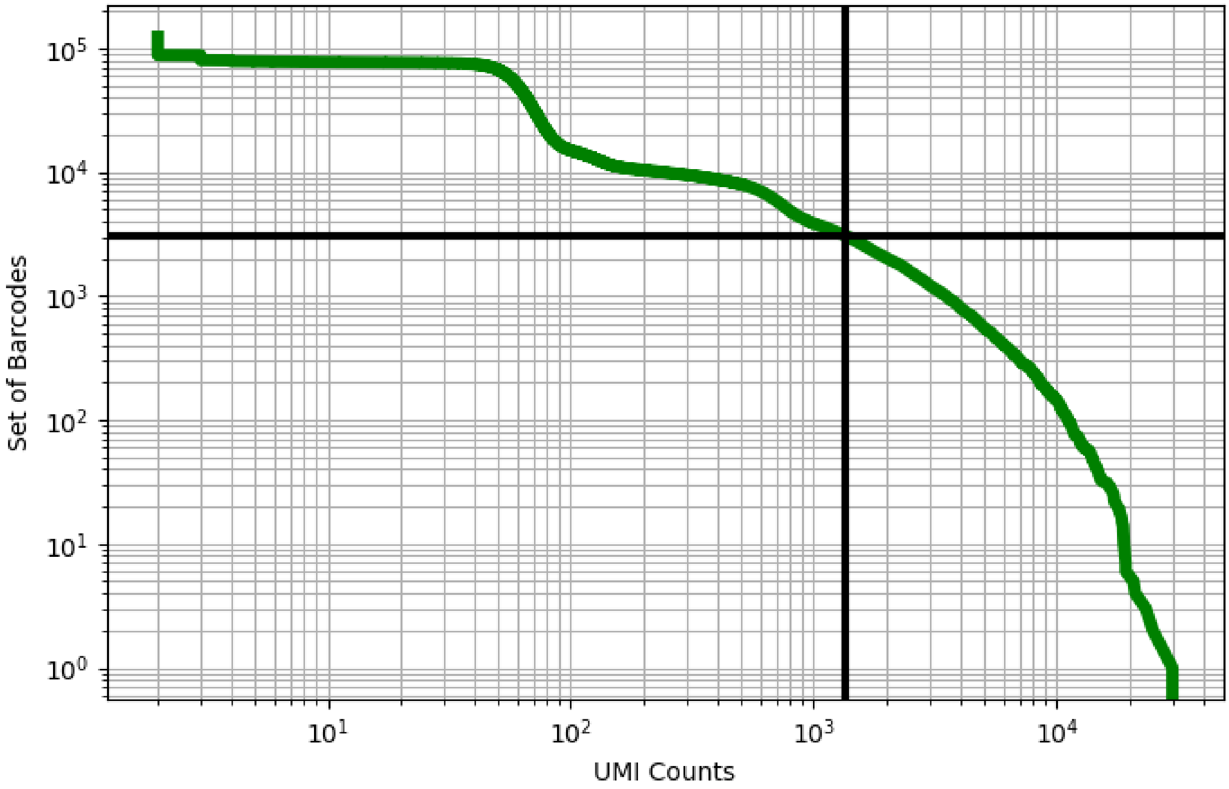


Figure 486: testis/GSM3526583

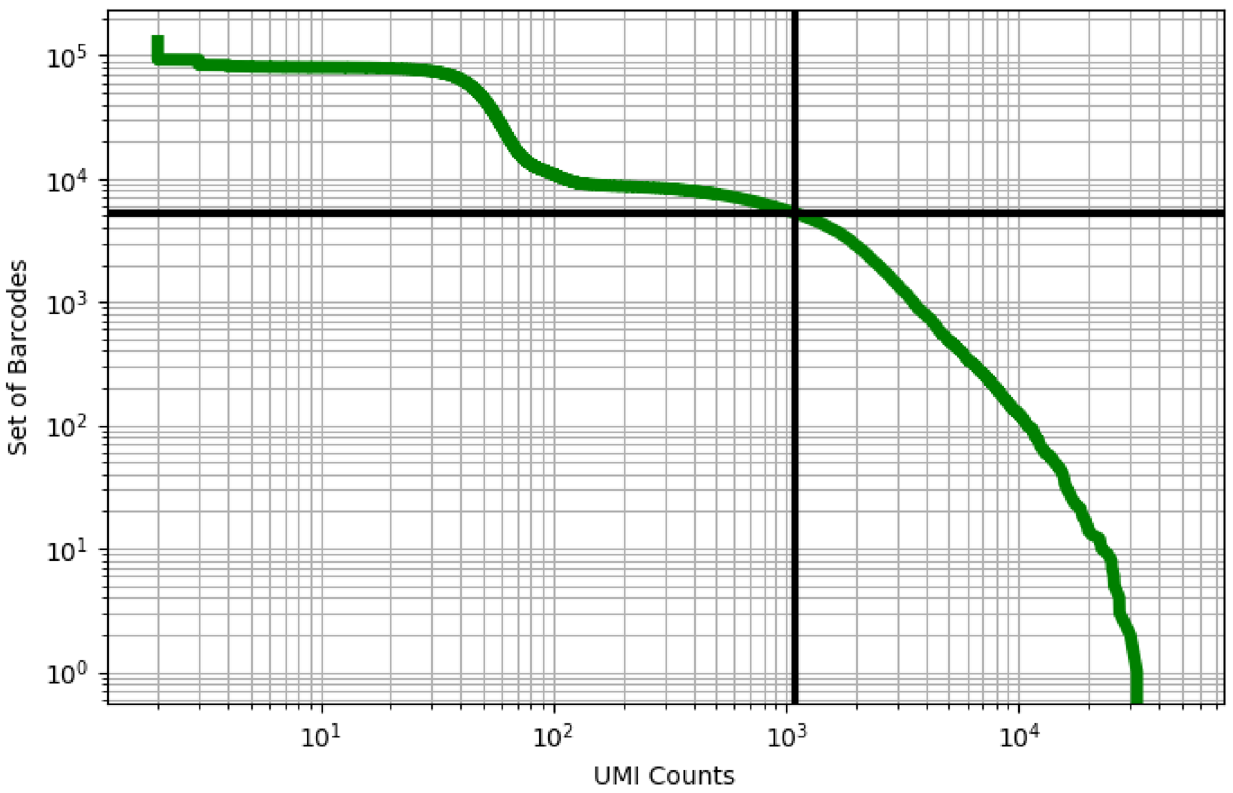


Figure 487: testis/GSM3526584

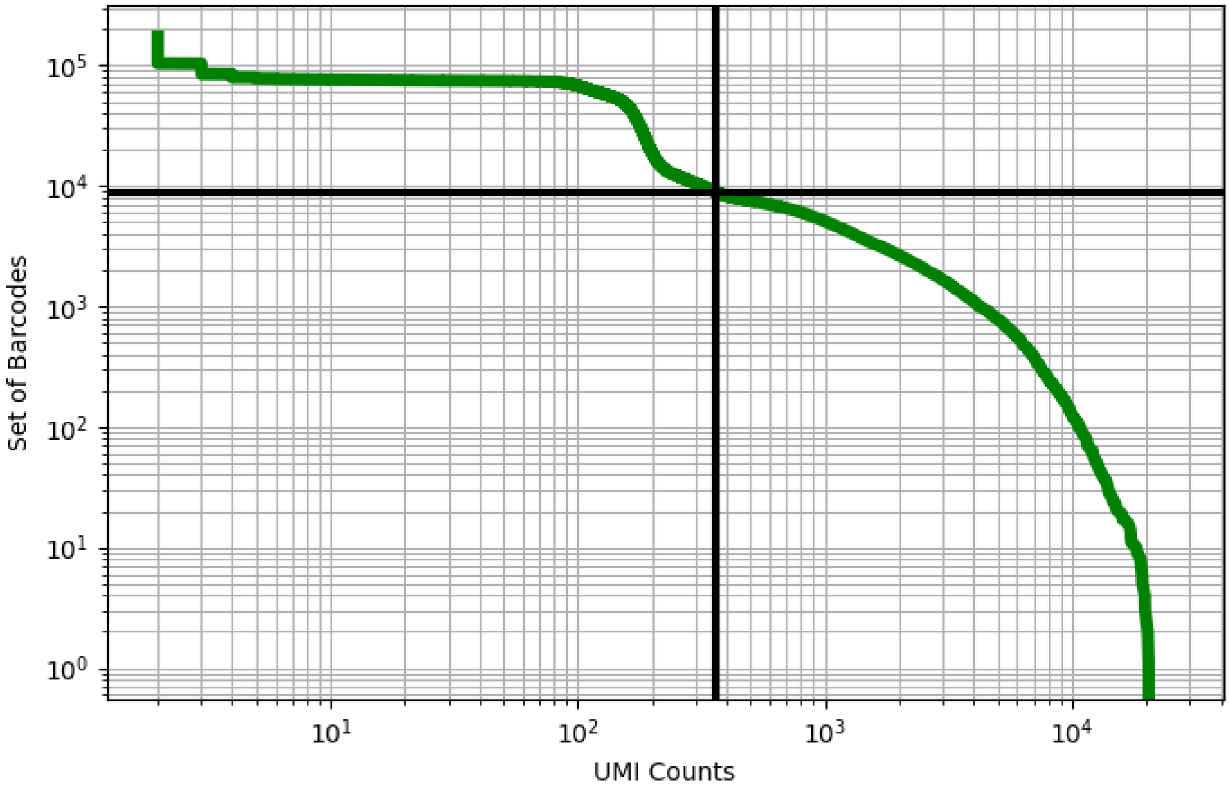


Figure 488: testis/GSM3526585

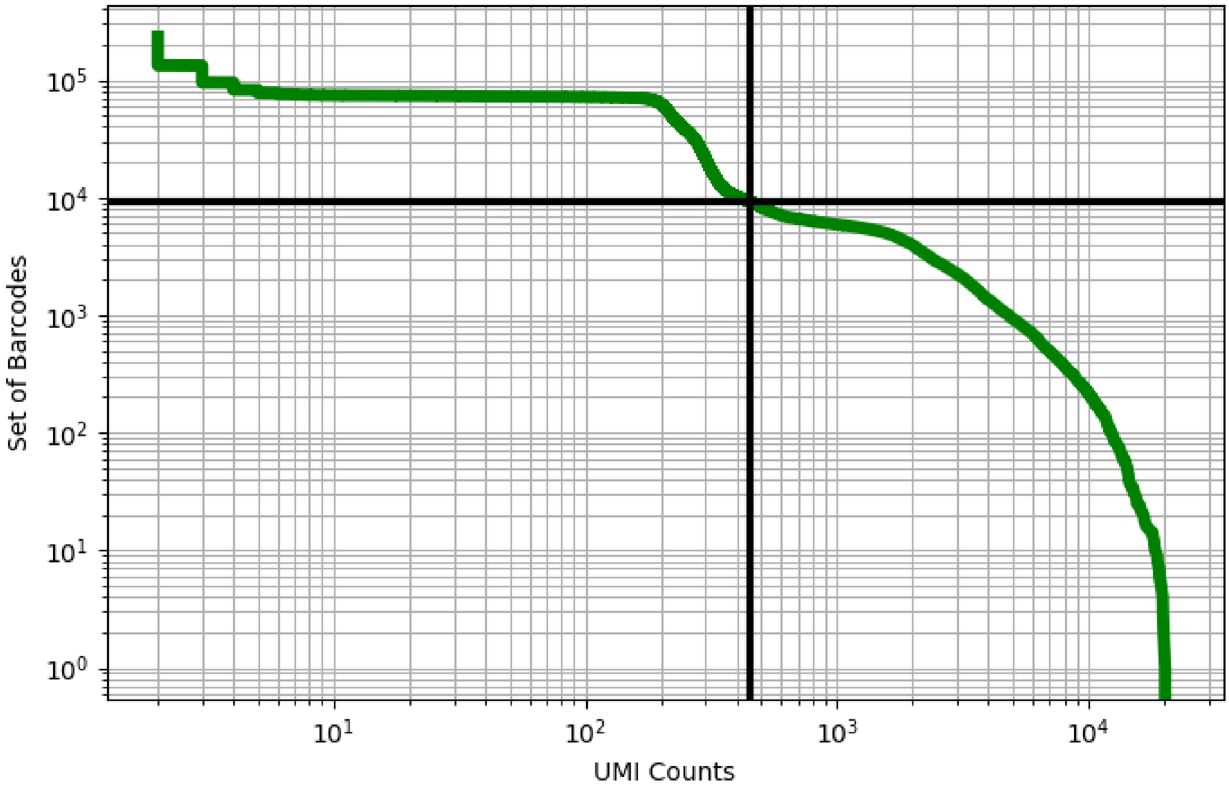


Figure 489: testis/GSM3526586

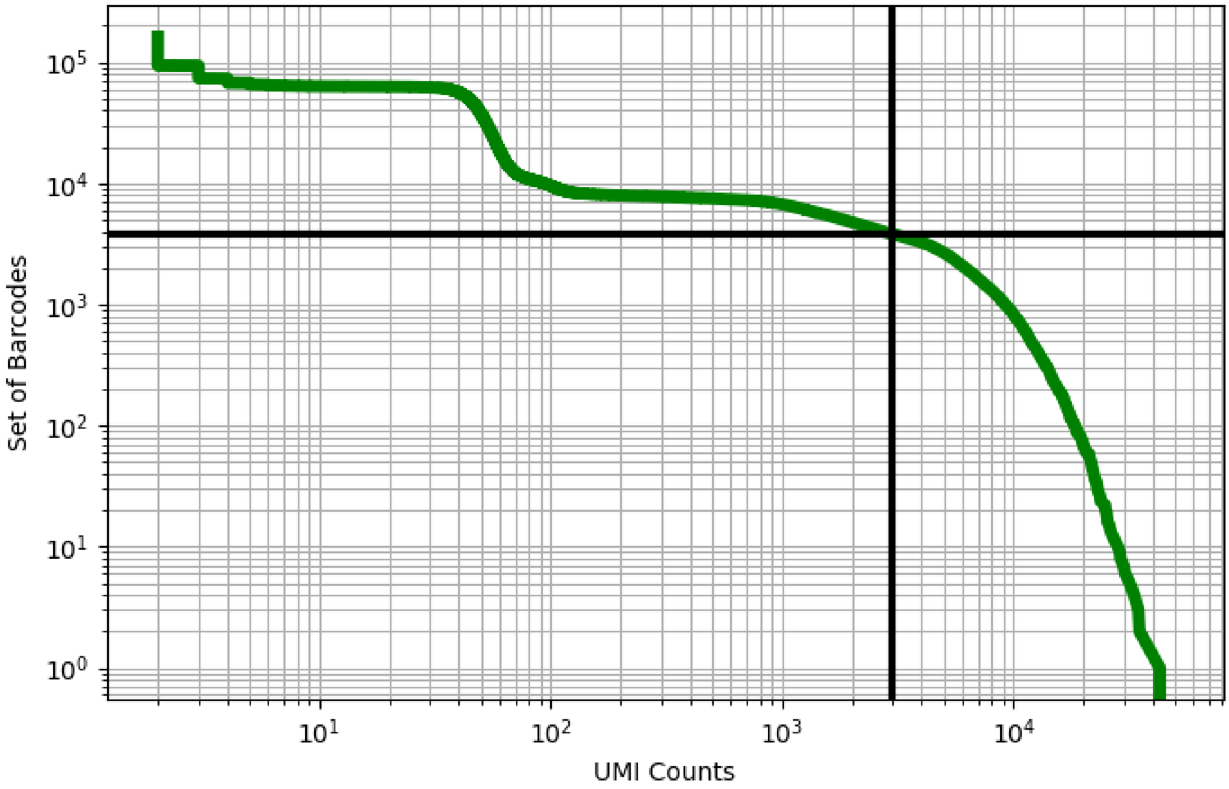


Figure 490: testis/GSM3526587

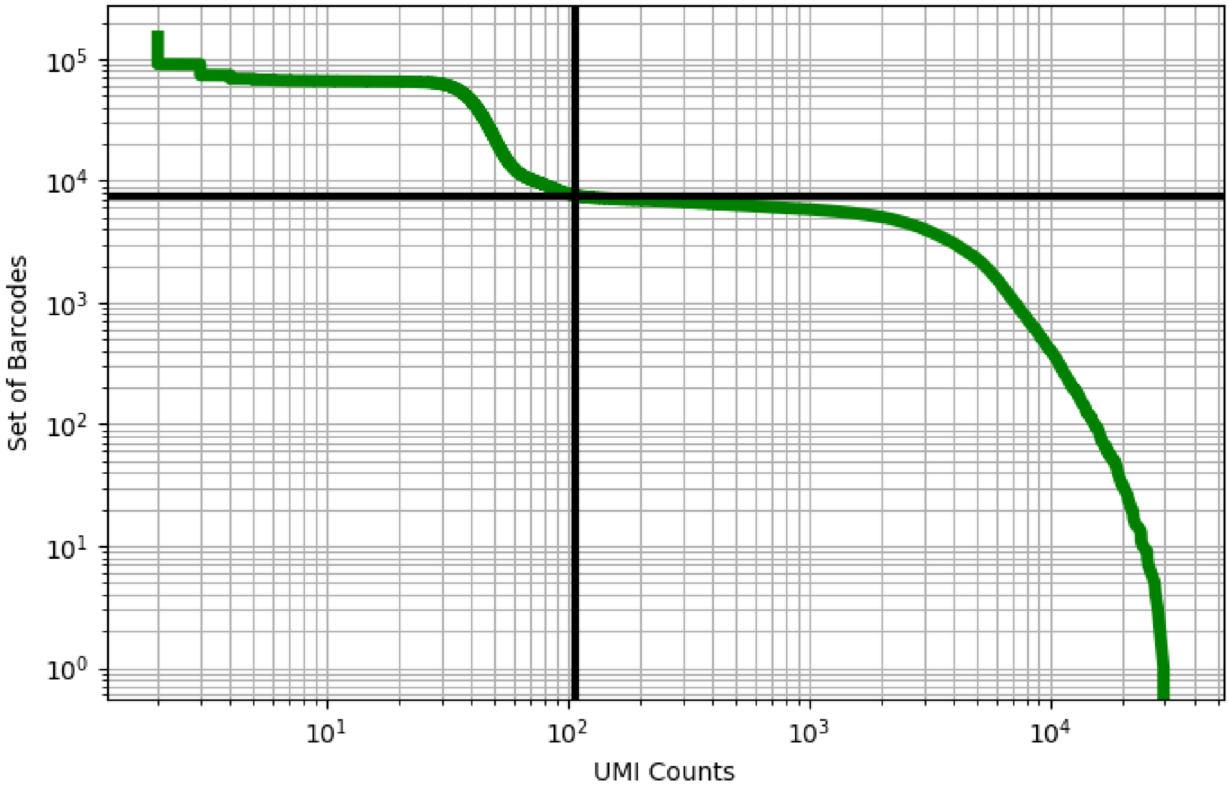


Figure 491: testis/GSM3526588

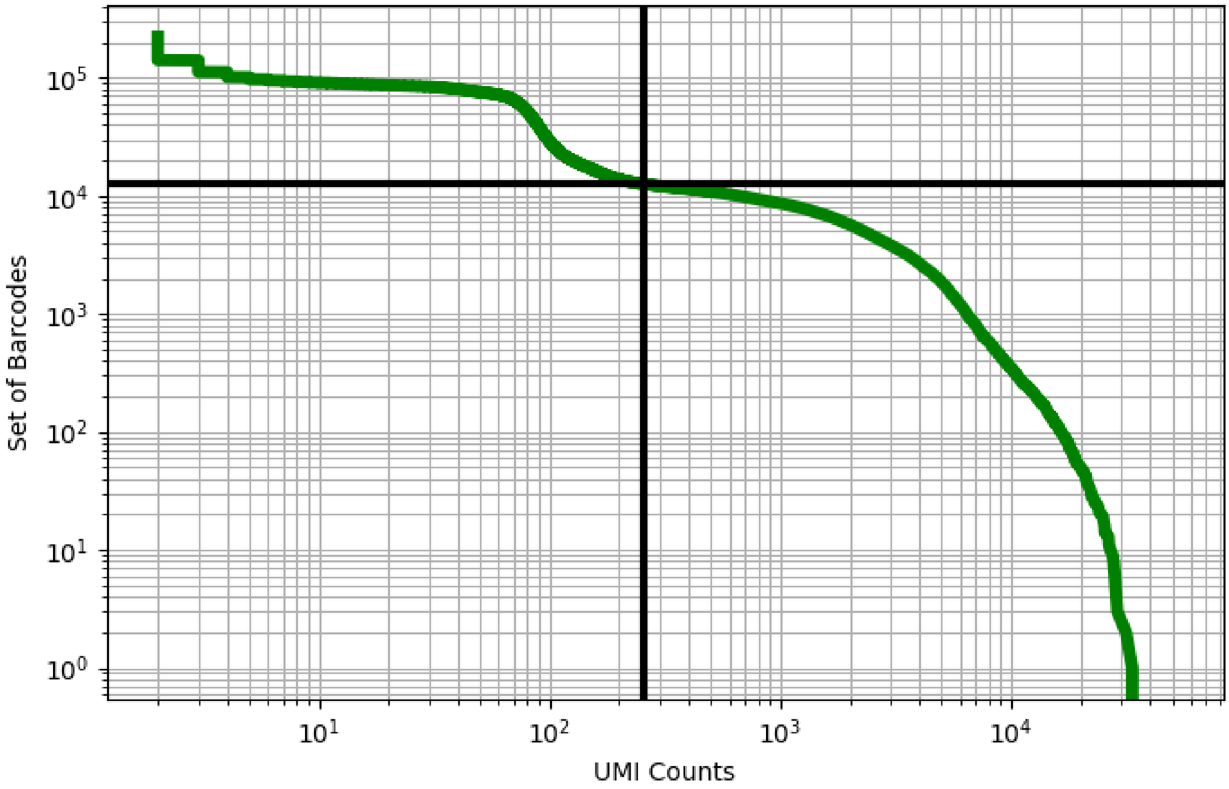


Figure 492: testis/GSM3526589

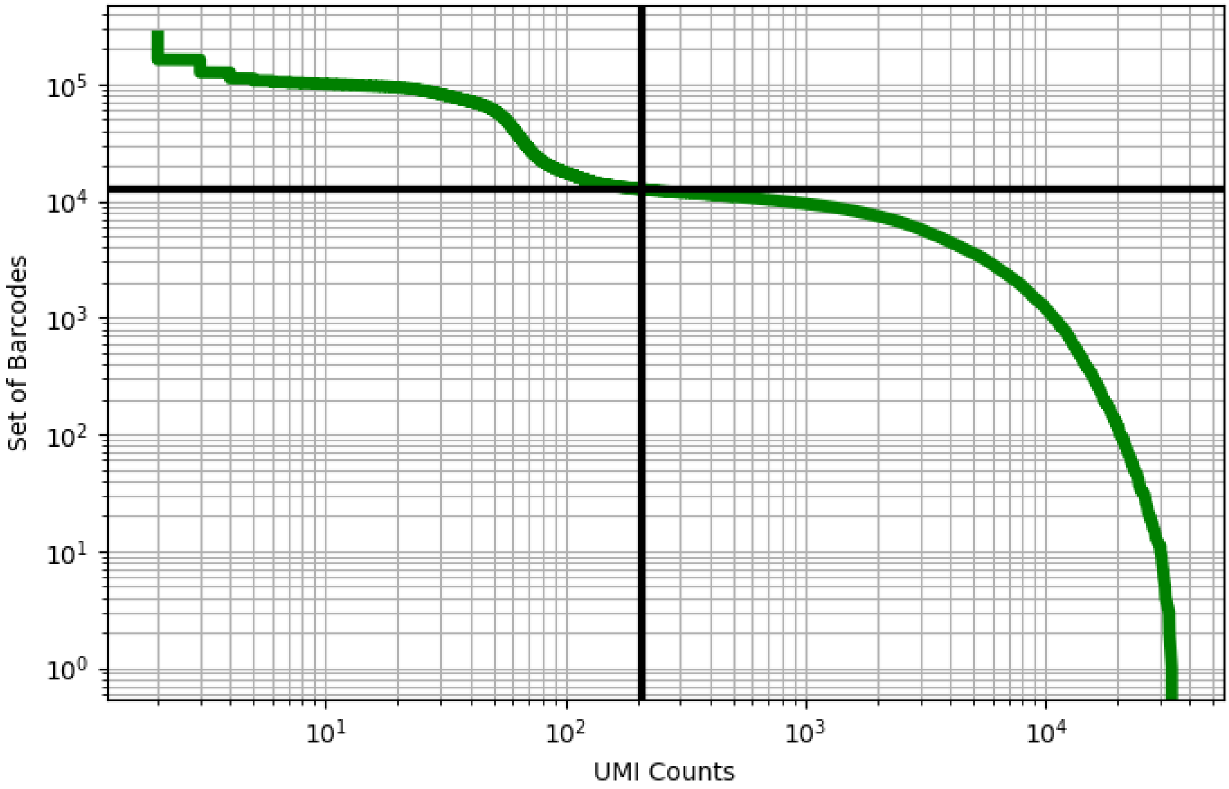


Figure 493: testis/GSM3526590

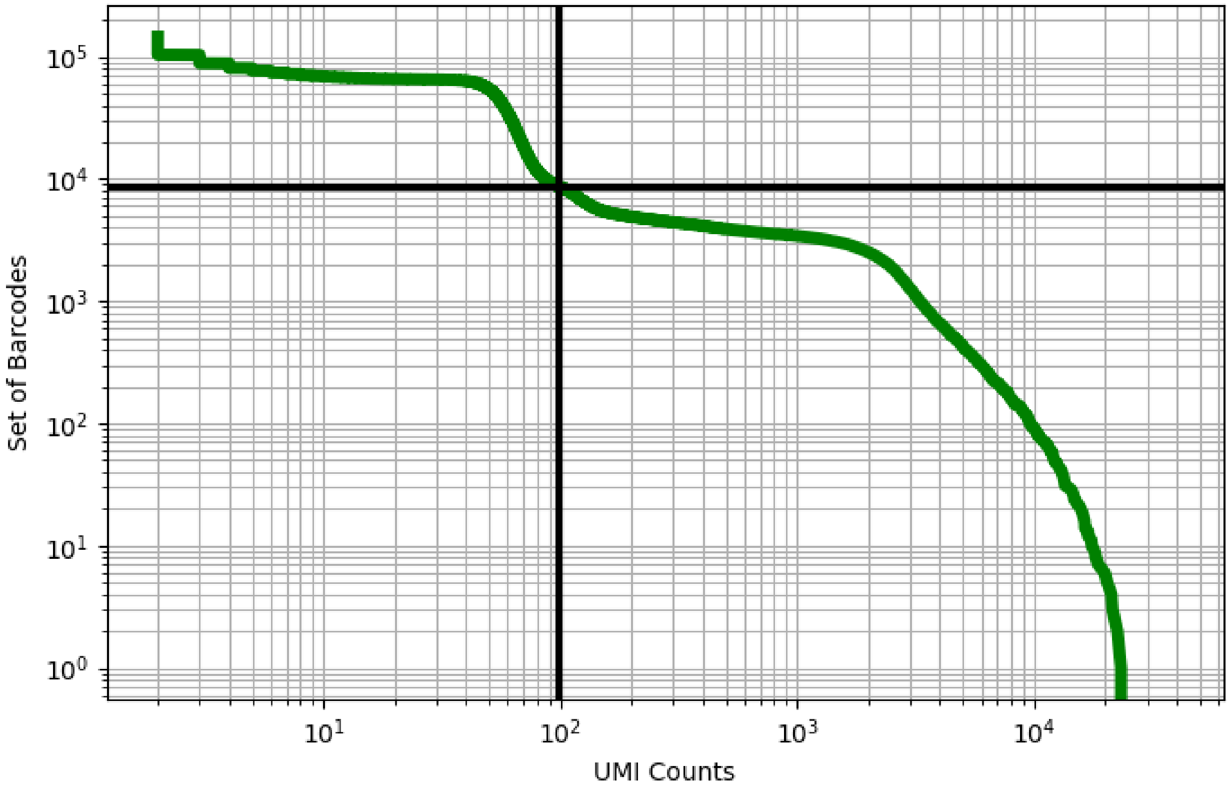


Figure 494: testis/GSM3732871

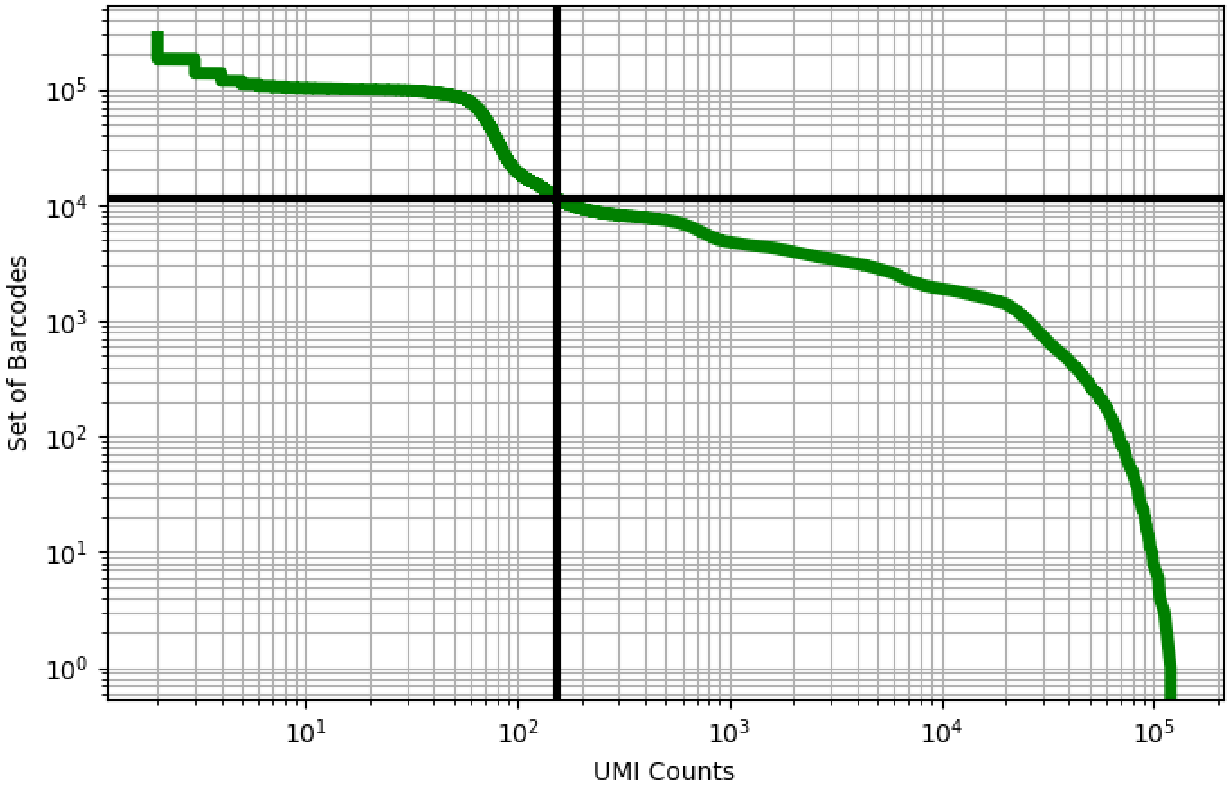


Figure 495: testis/GSM4486714

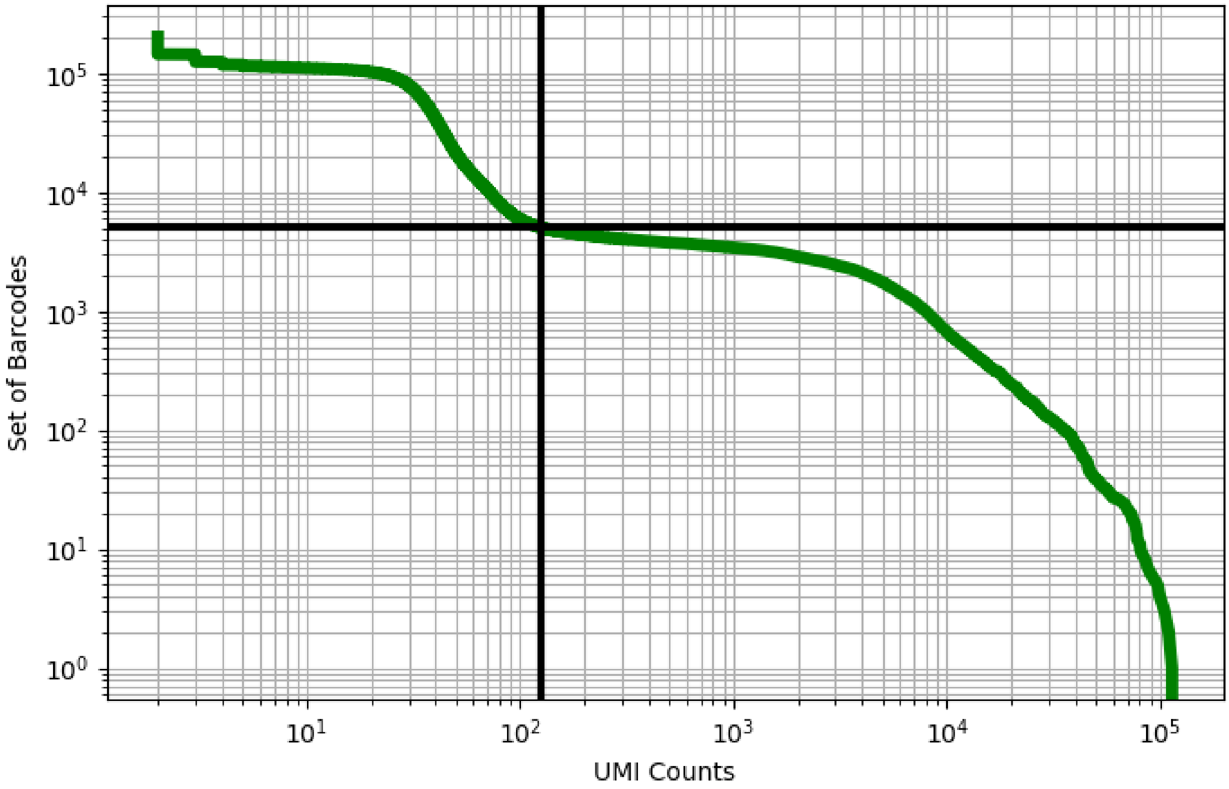


Figure 496: testis/GSM4486715

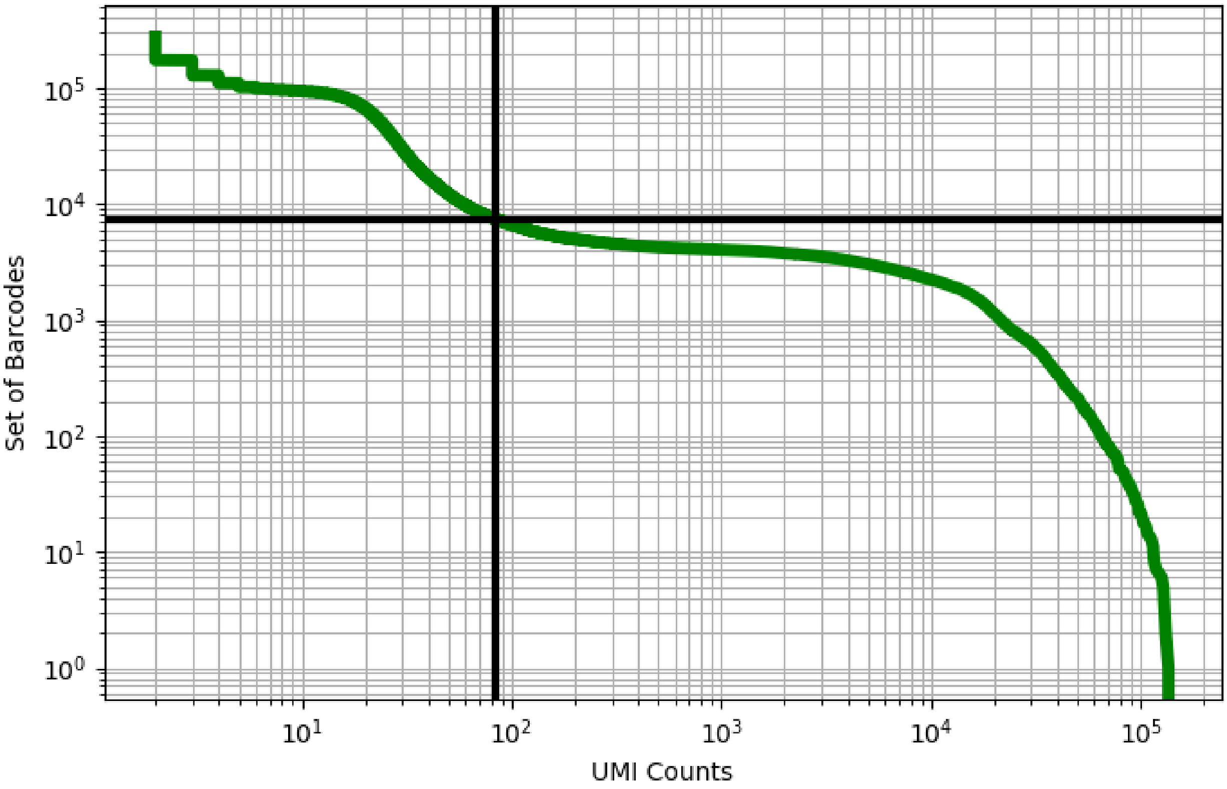


Figure 497: testis/GSM4486716

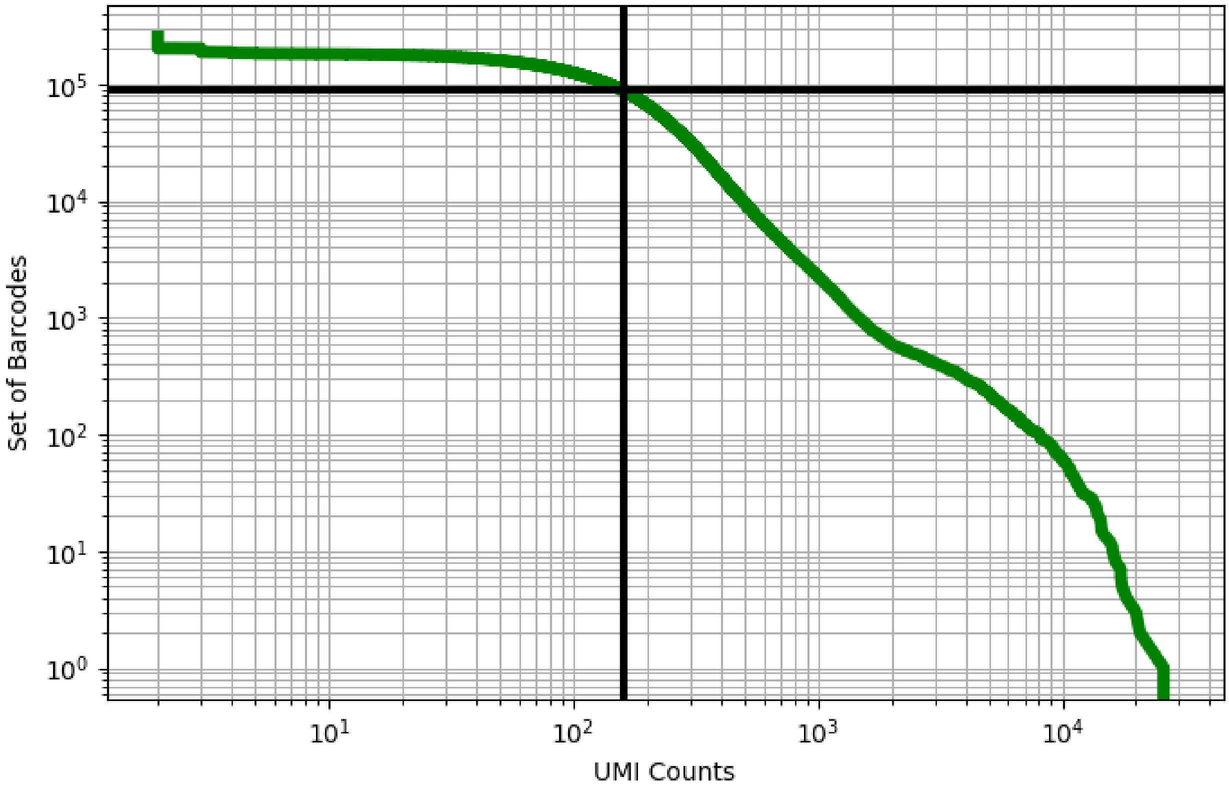


Figure 498: thymus/ERS4228628

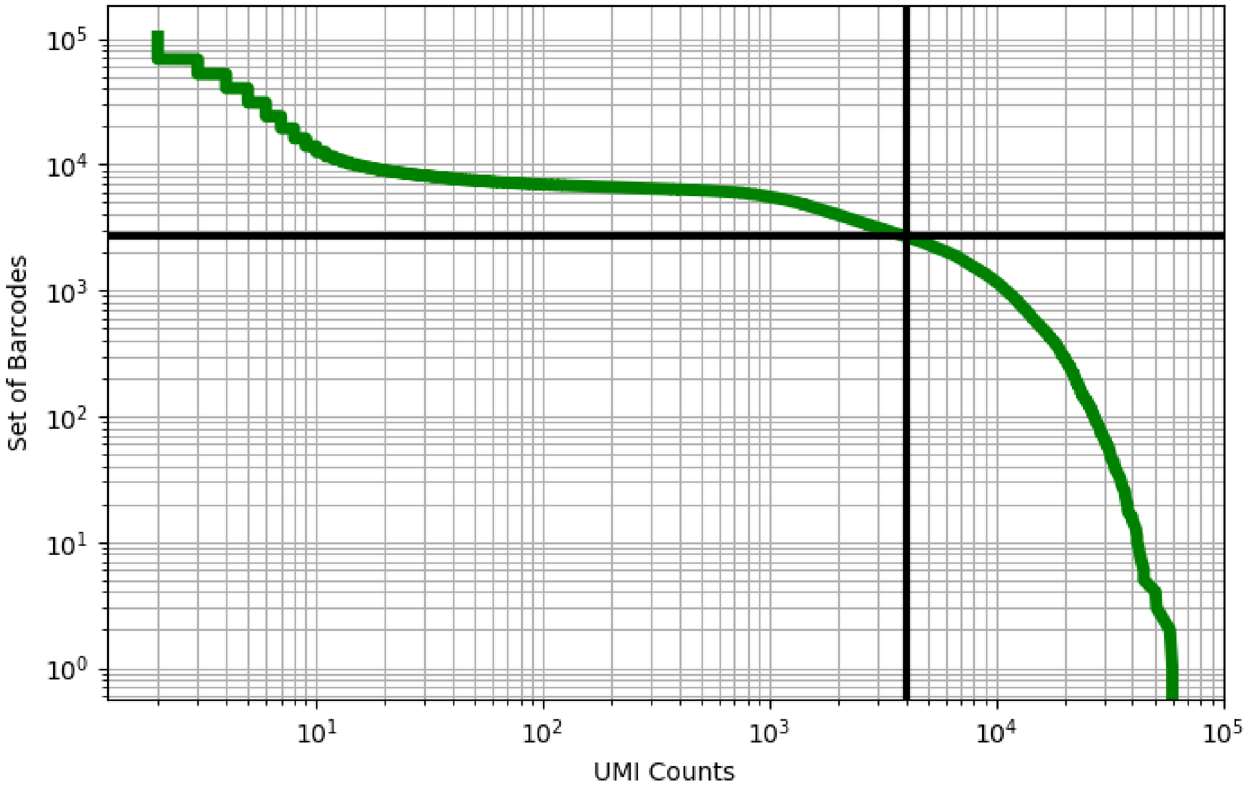


Figure 499: thymus/ERS4228634

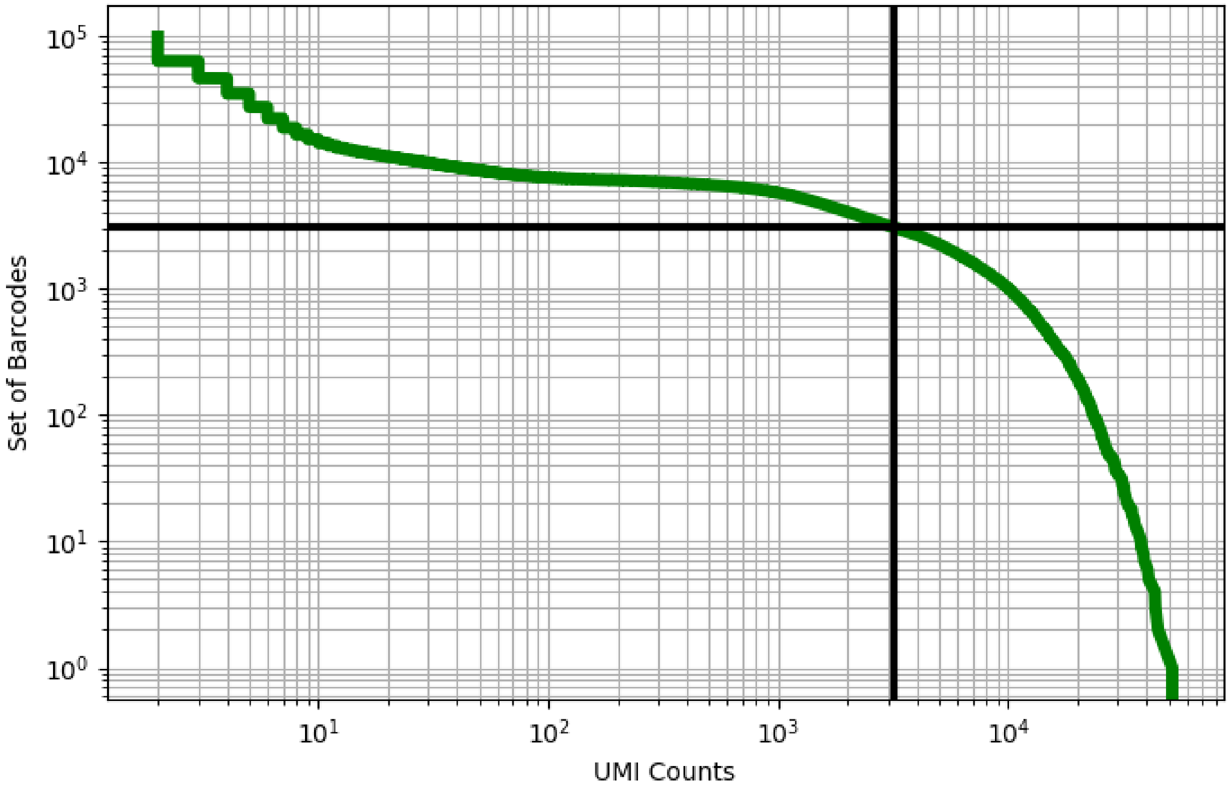


Figure 500: thymus/ERS4228635

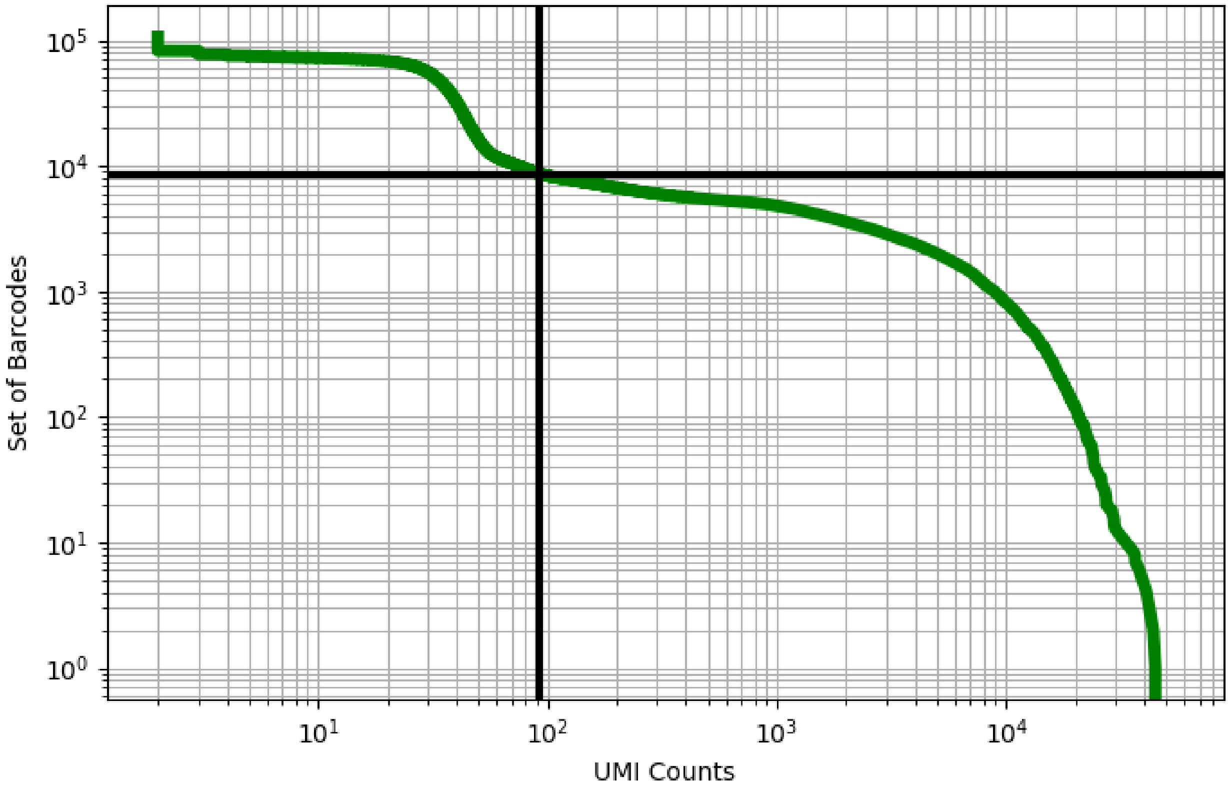


Figure 501: thymus/ERS4228636

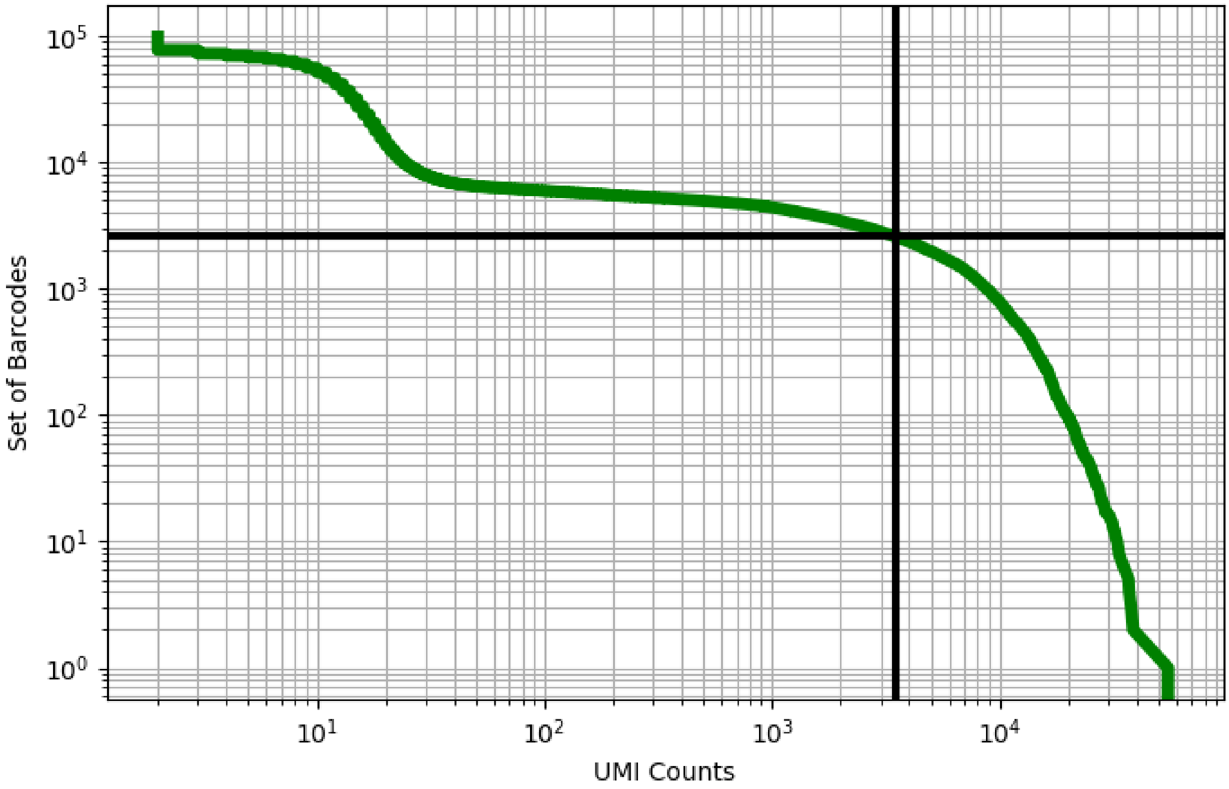


Figure 502: thymus/ERS4228637

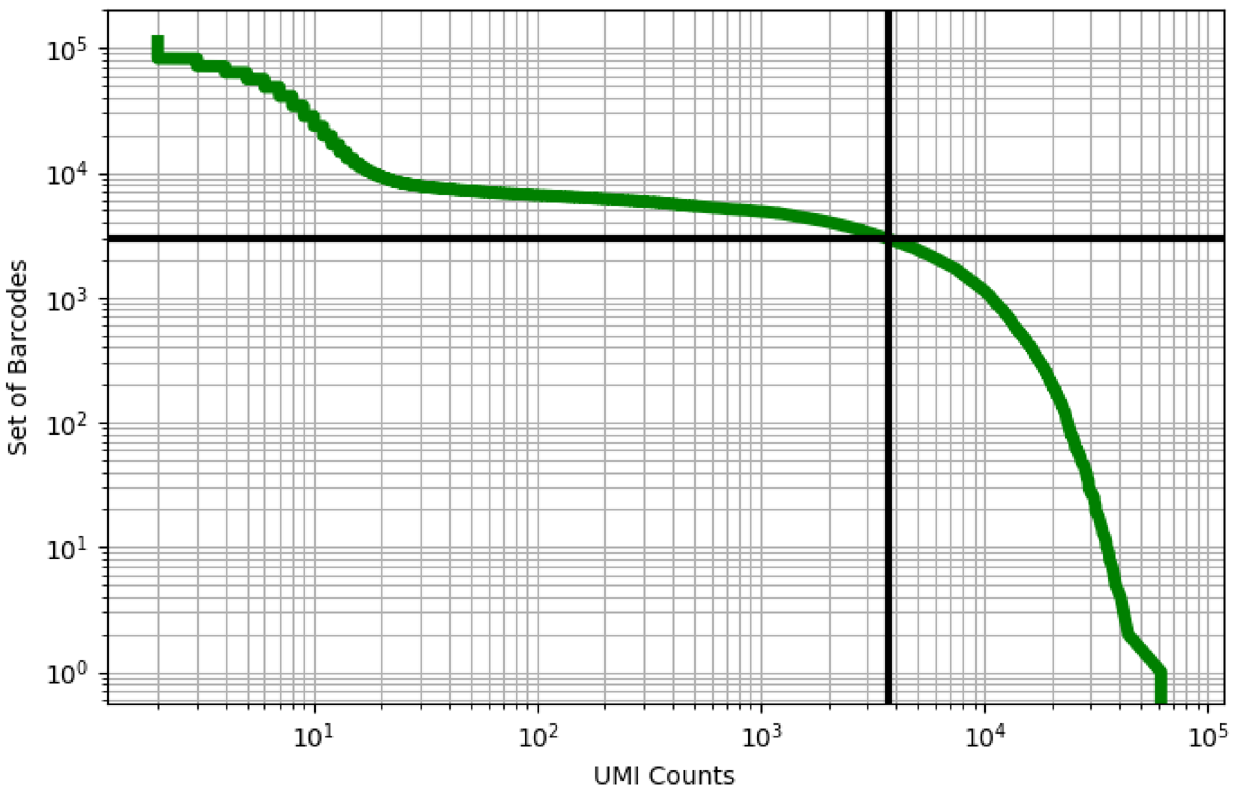


Figure 503: thymus/ERS4228638

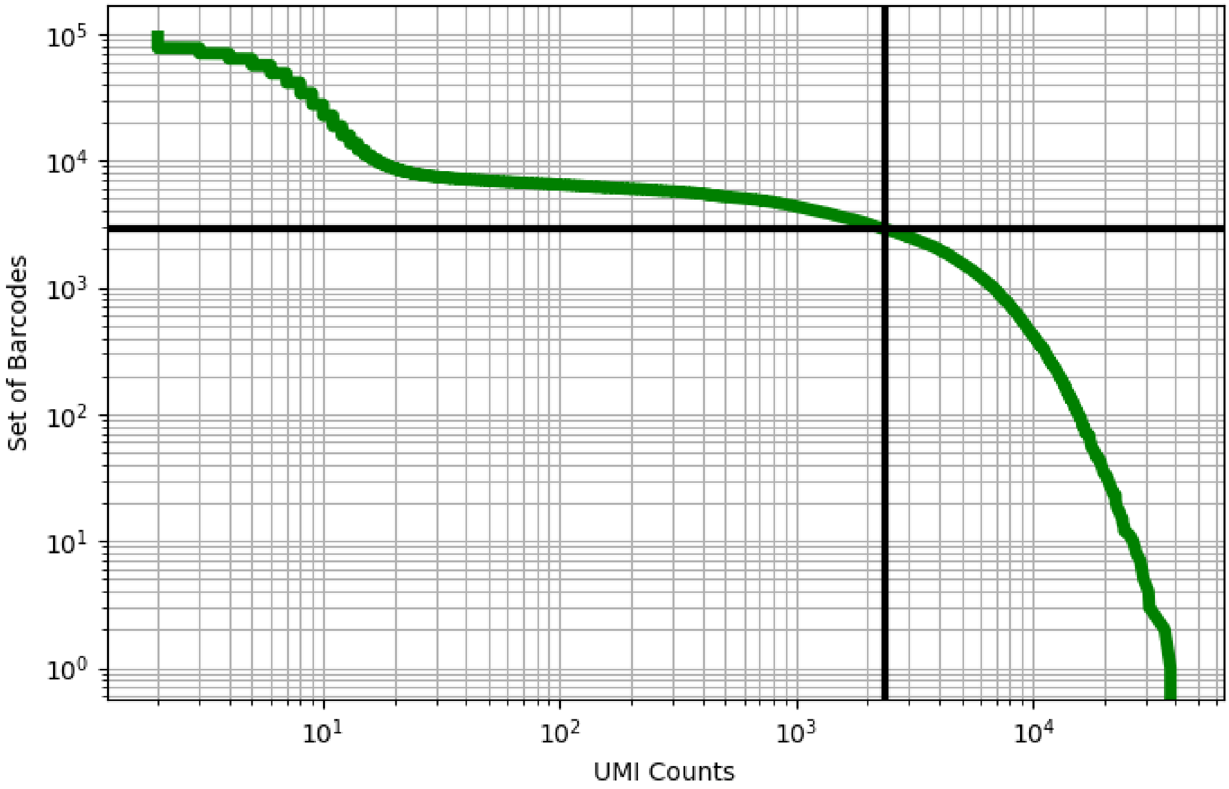


Figure 504: thymus/ERS4228639

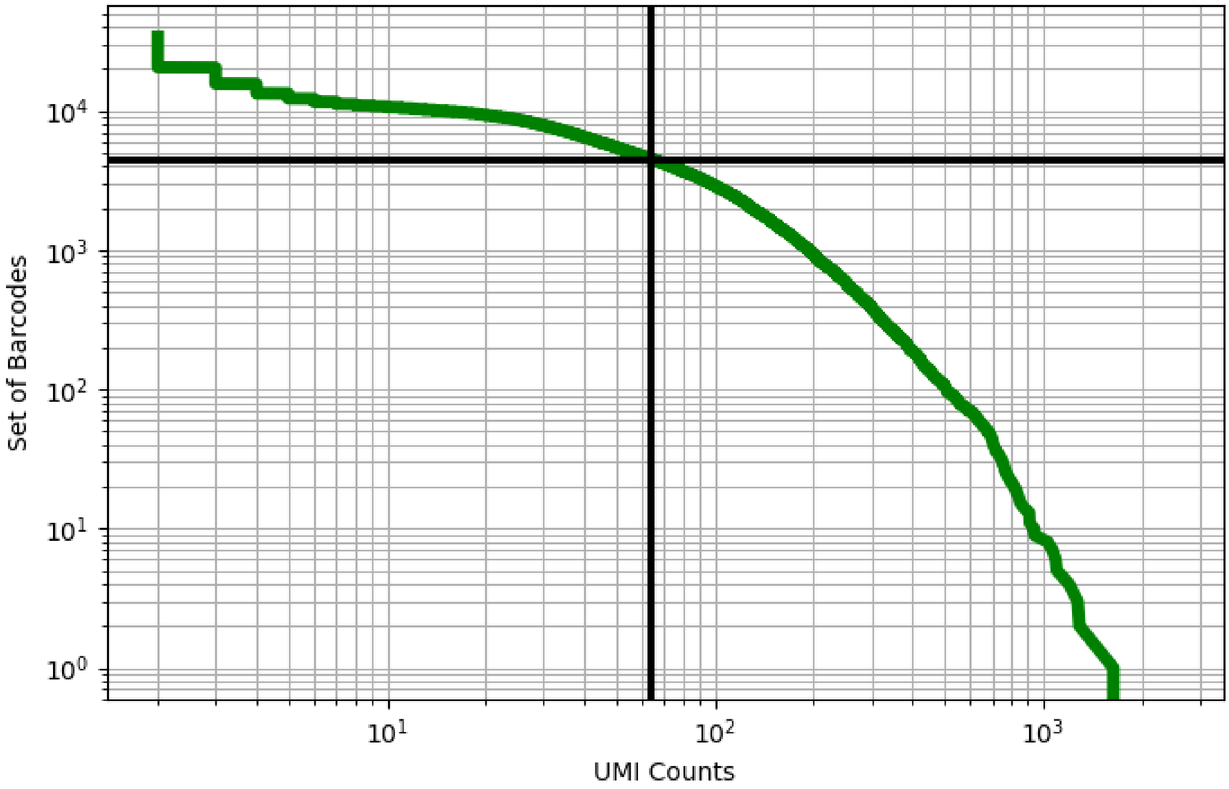


Figure 505: thymus/ERS4228662

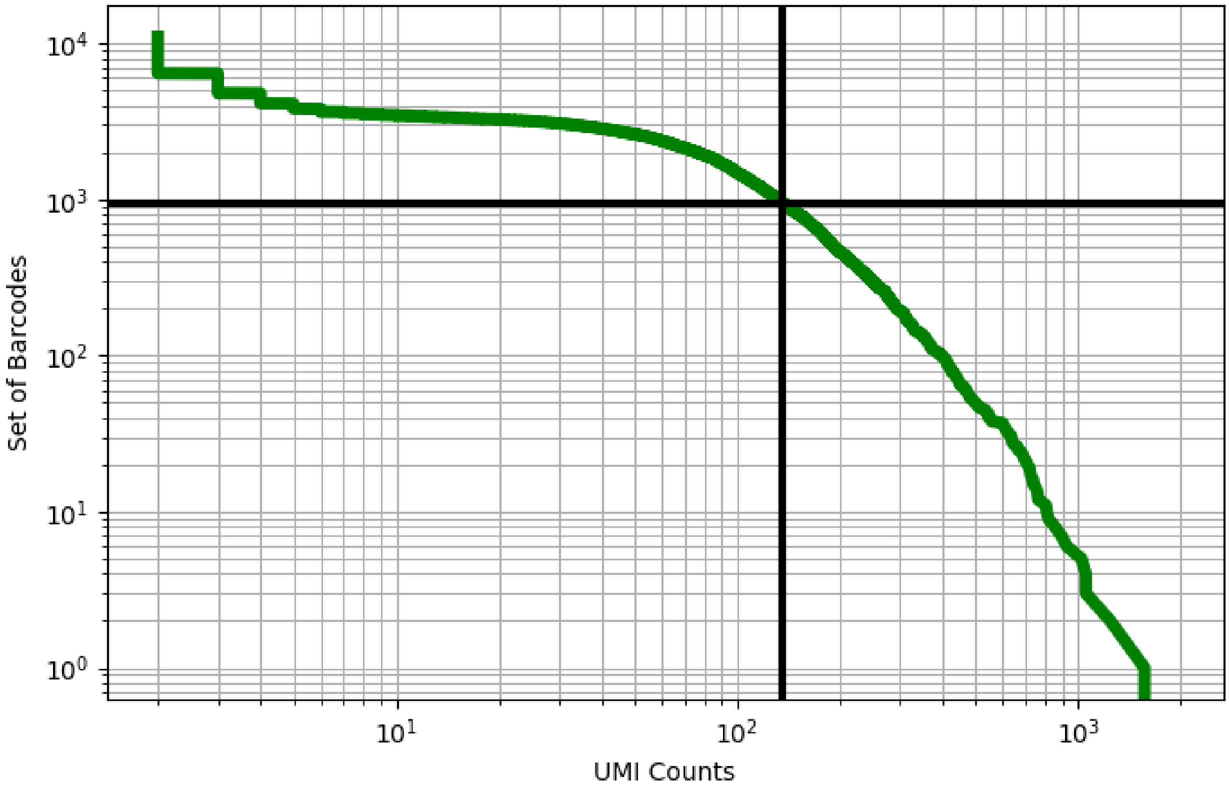


Figure 506: thymus/ERS4228663

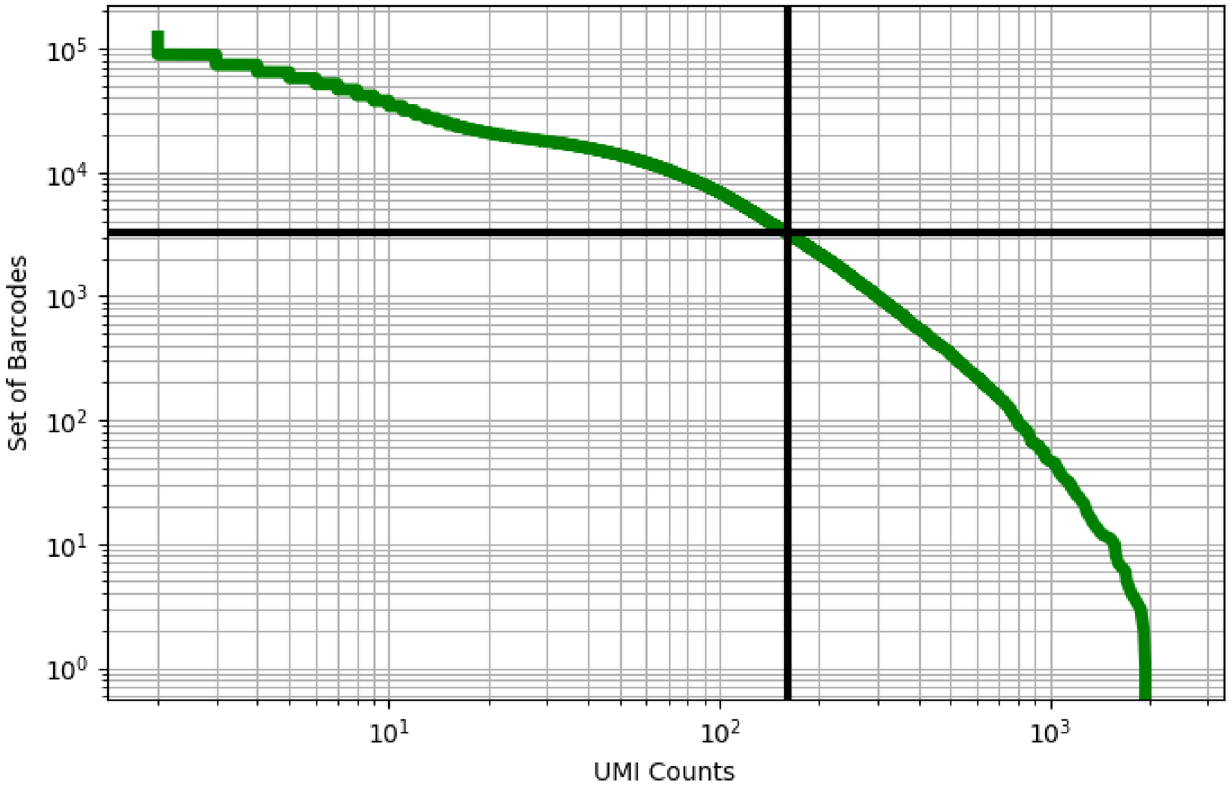


Figure 507: thymus/ERS4228664

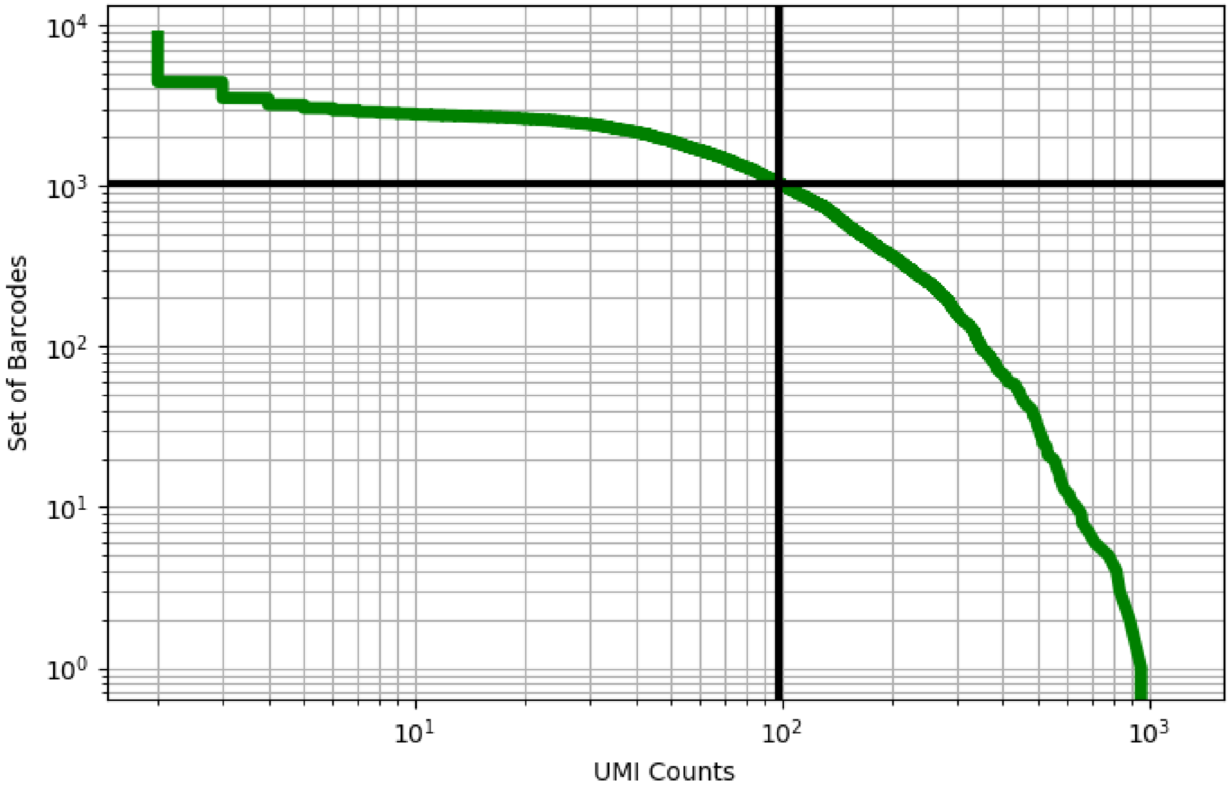


Figure 508: thymus/ERS4228665

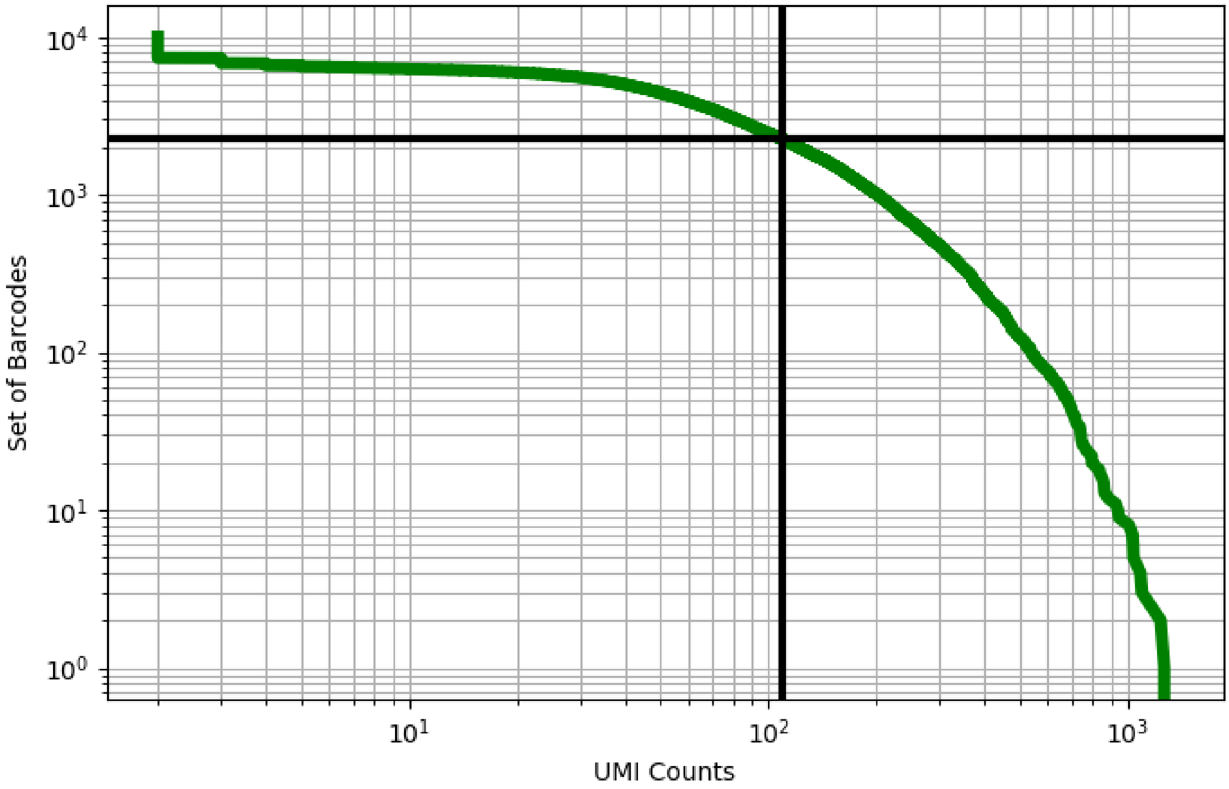


Figure 509: thymus/ERS4228668

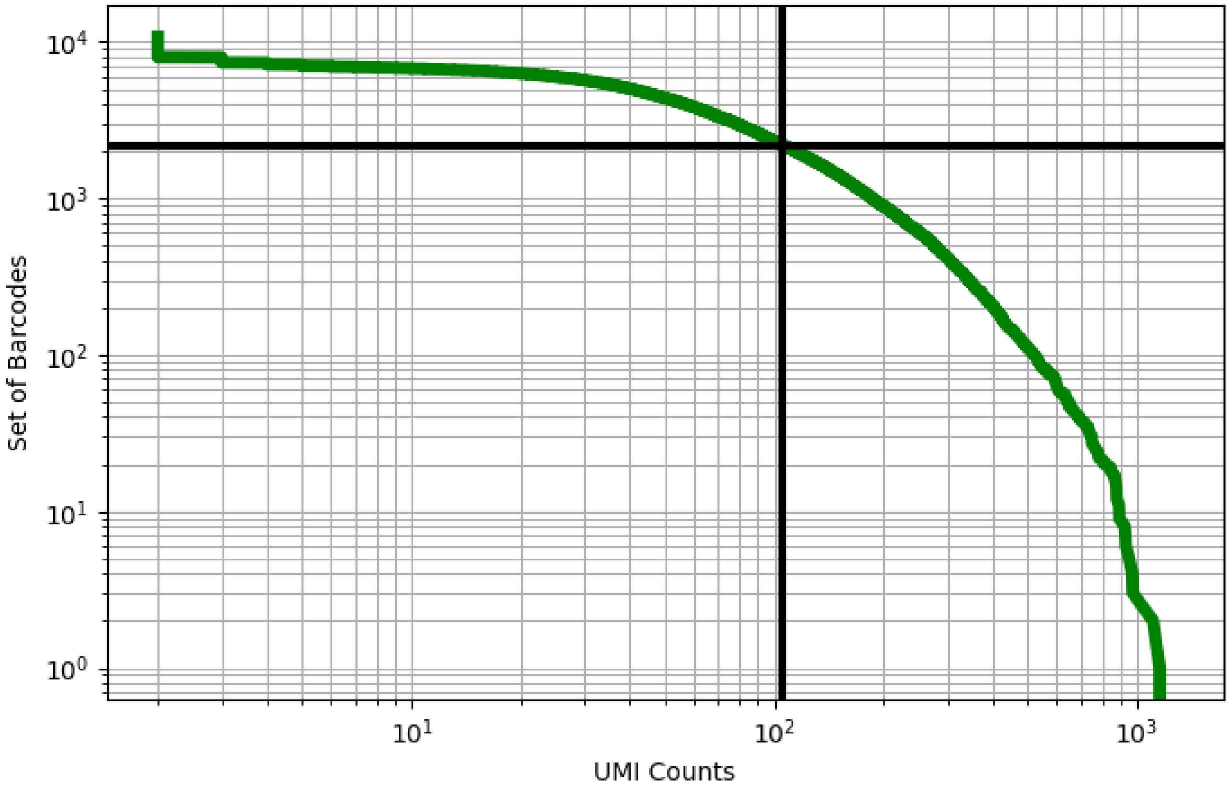


Figure 510: thymus/ERS4228669

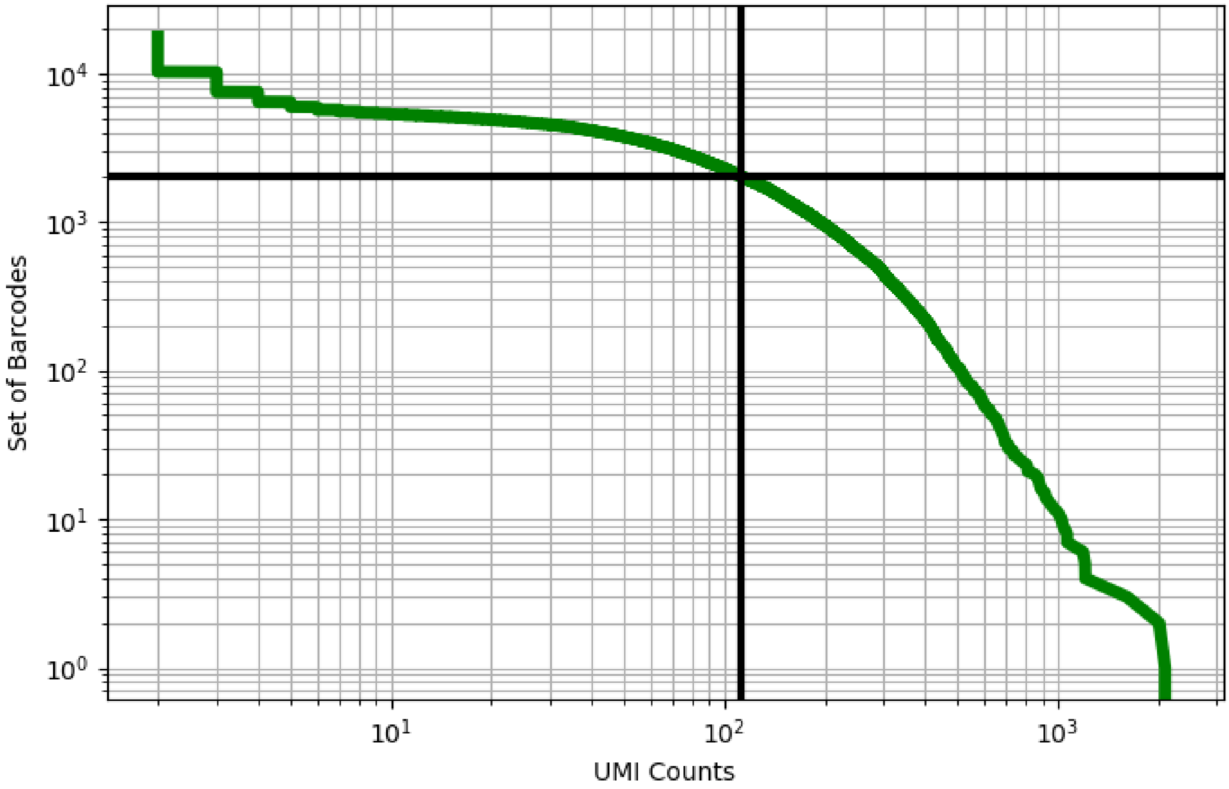


Figure 511: thymus/ERS4228670

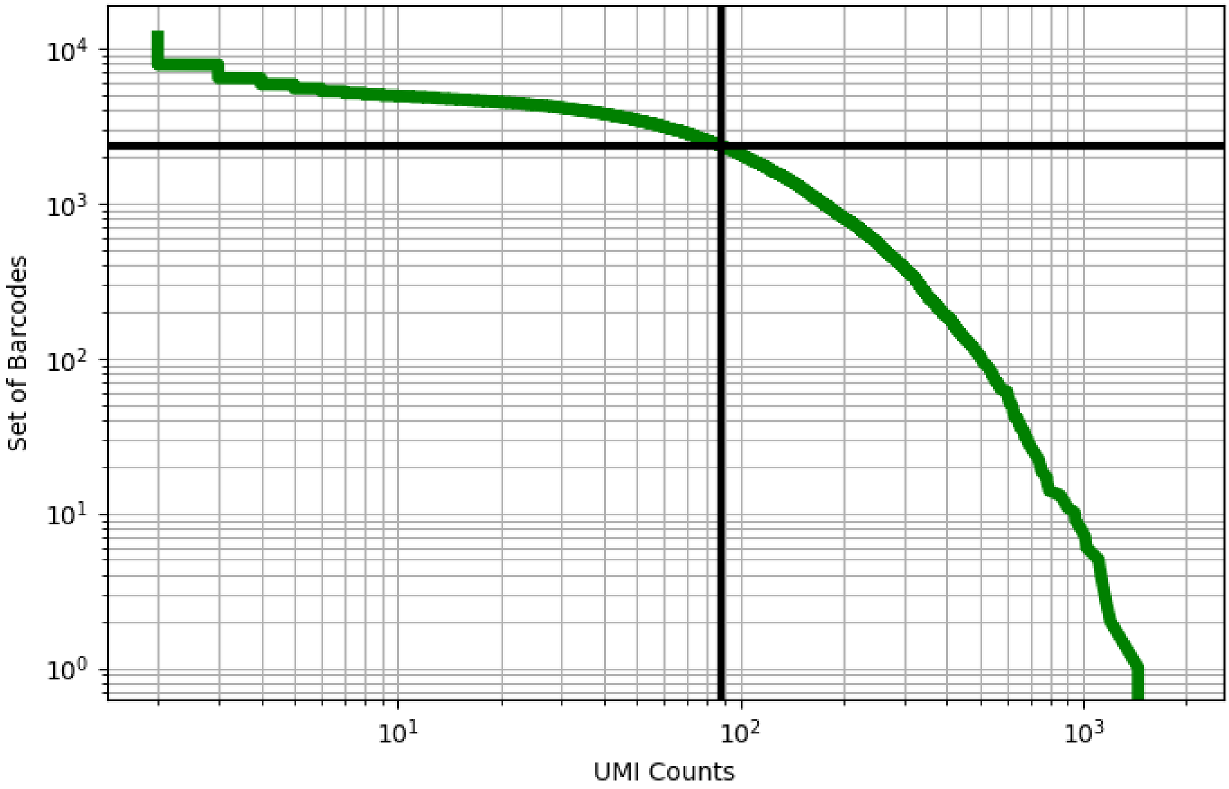


Figure 512: thymus/ERS4228671

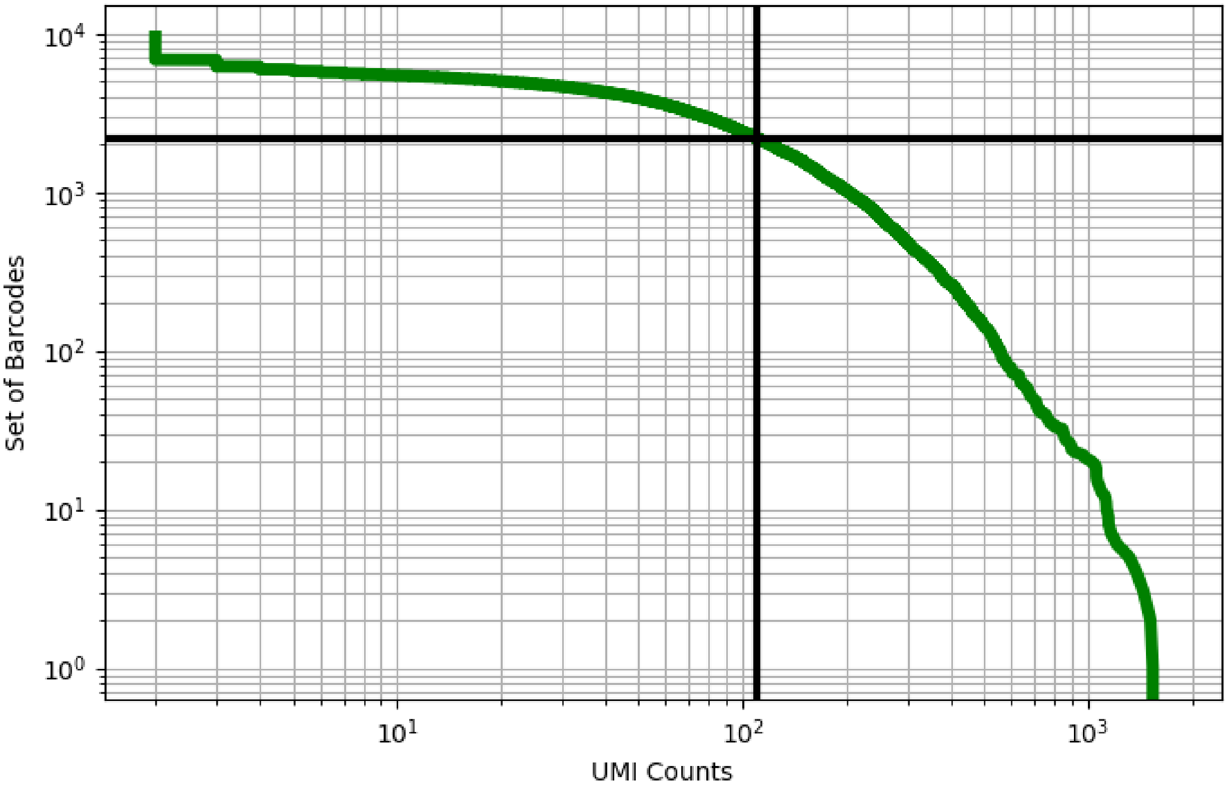


Figure 513: thymus/ERS4228672

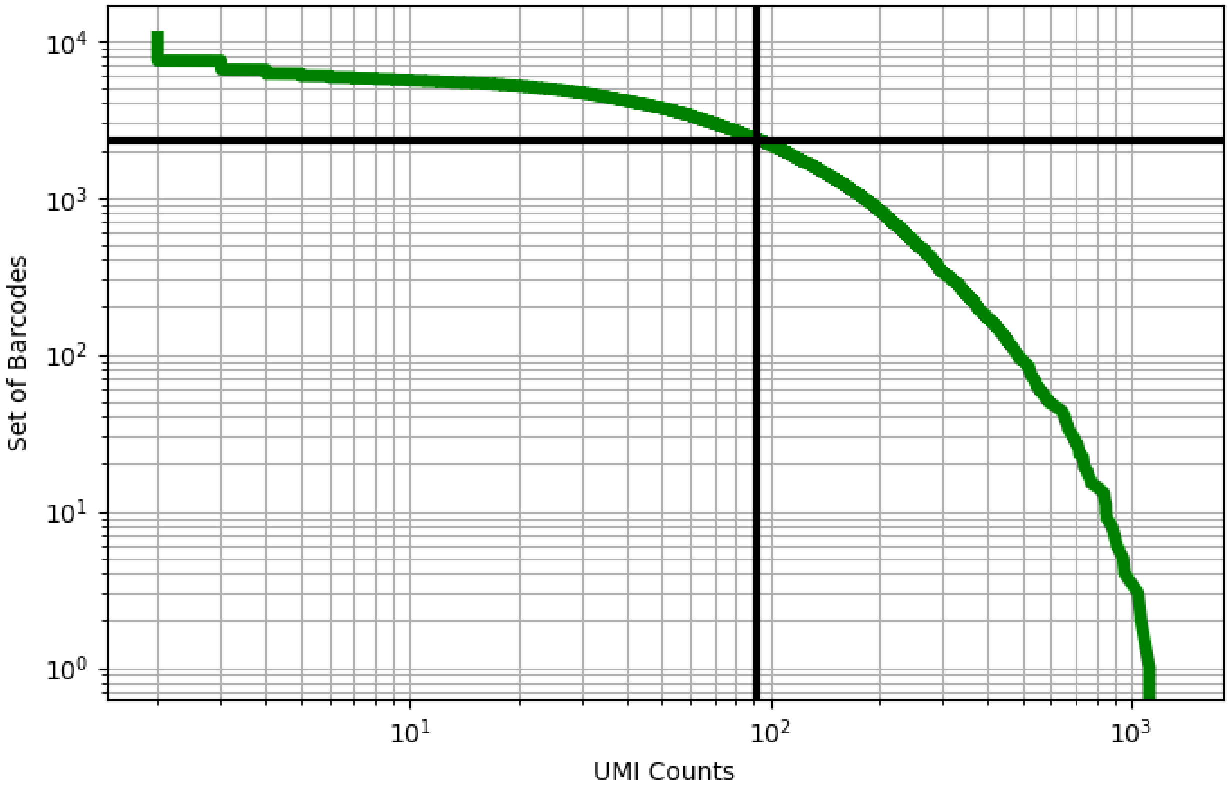


Figure 514: thymus/ERS4228673

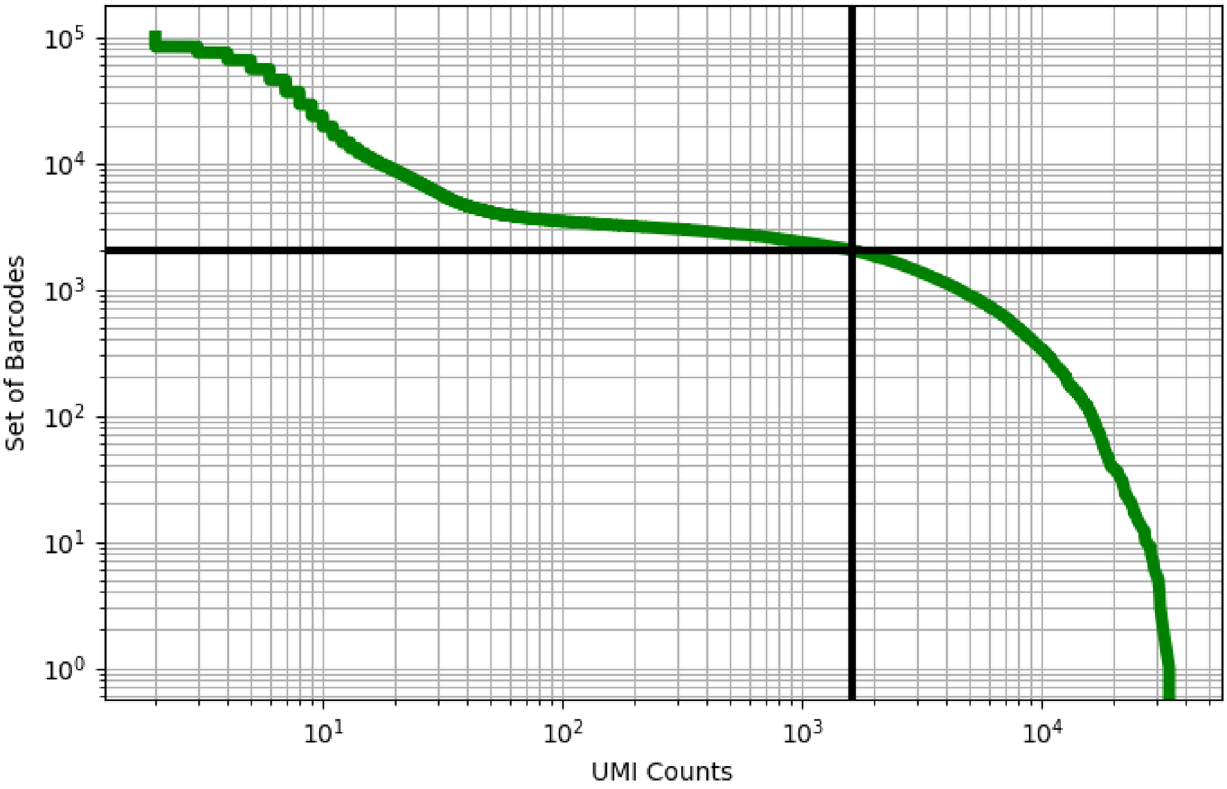


Figure 515: tonsil/GSM3375767

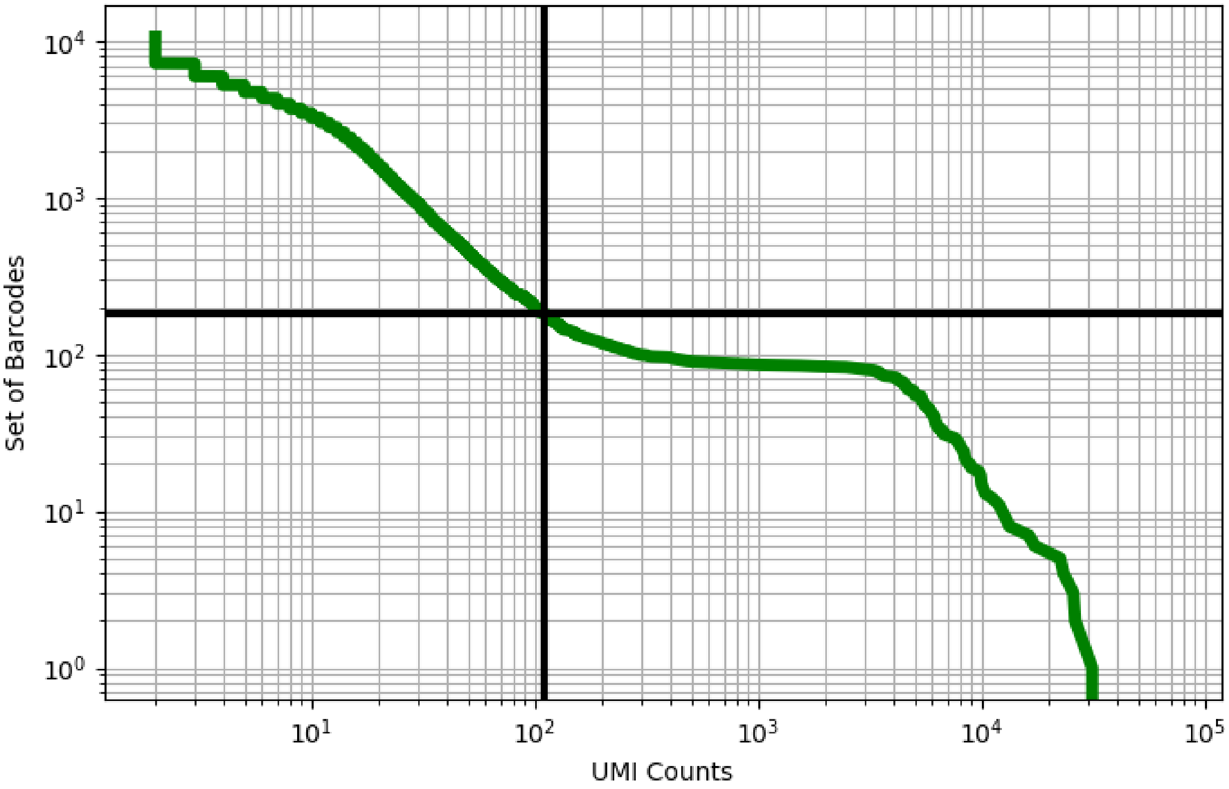


Figure 516: tonsil/GSM5051494

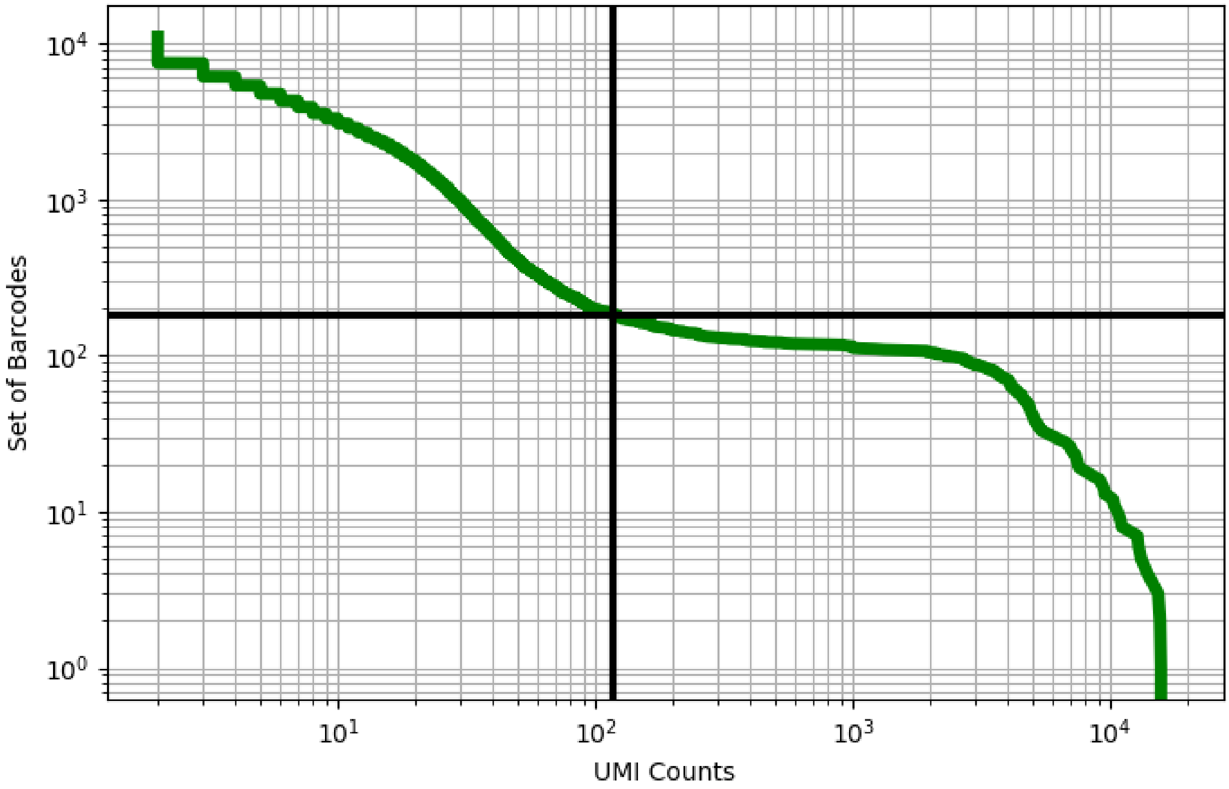


Figure 517: tonsil/GSM5051495

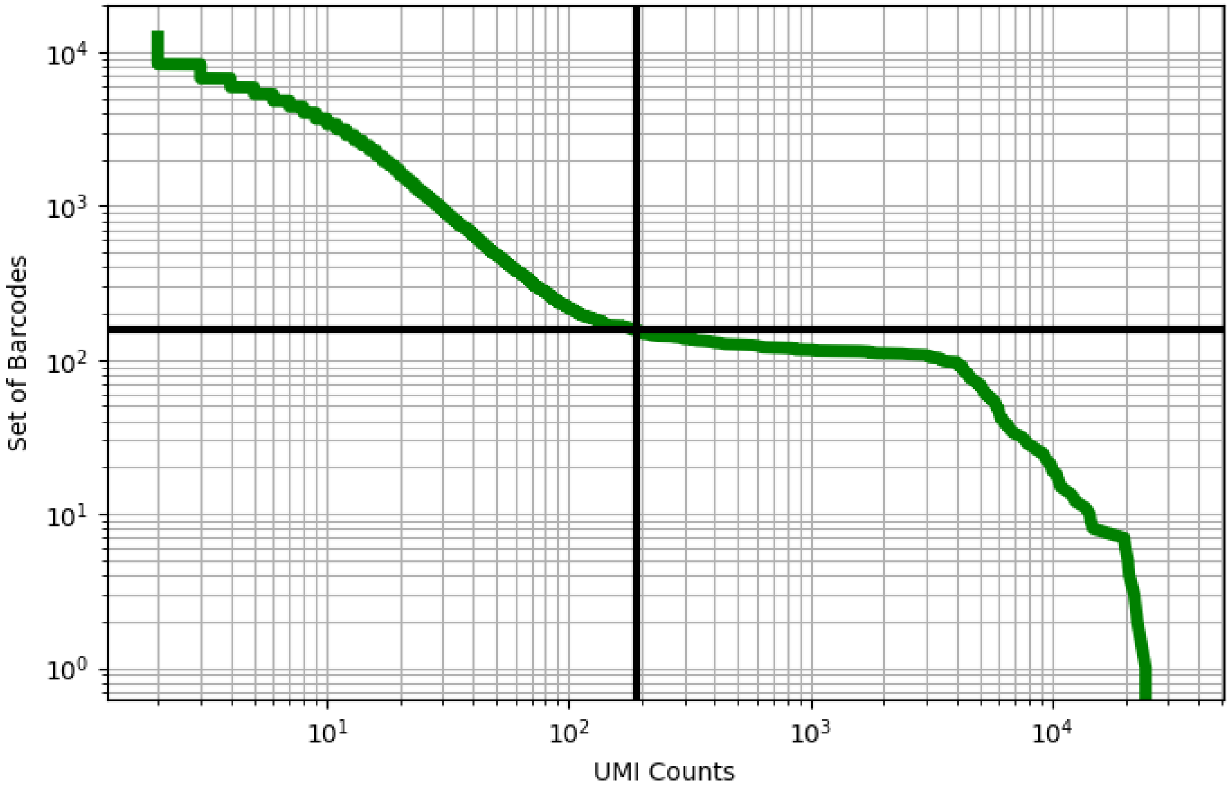


Figure 518: tonsil/GSM5051497

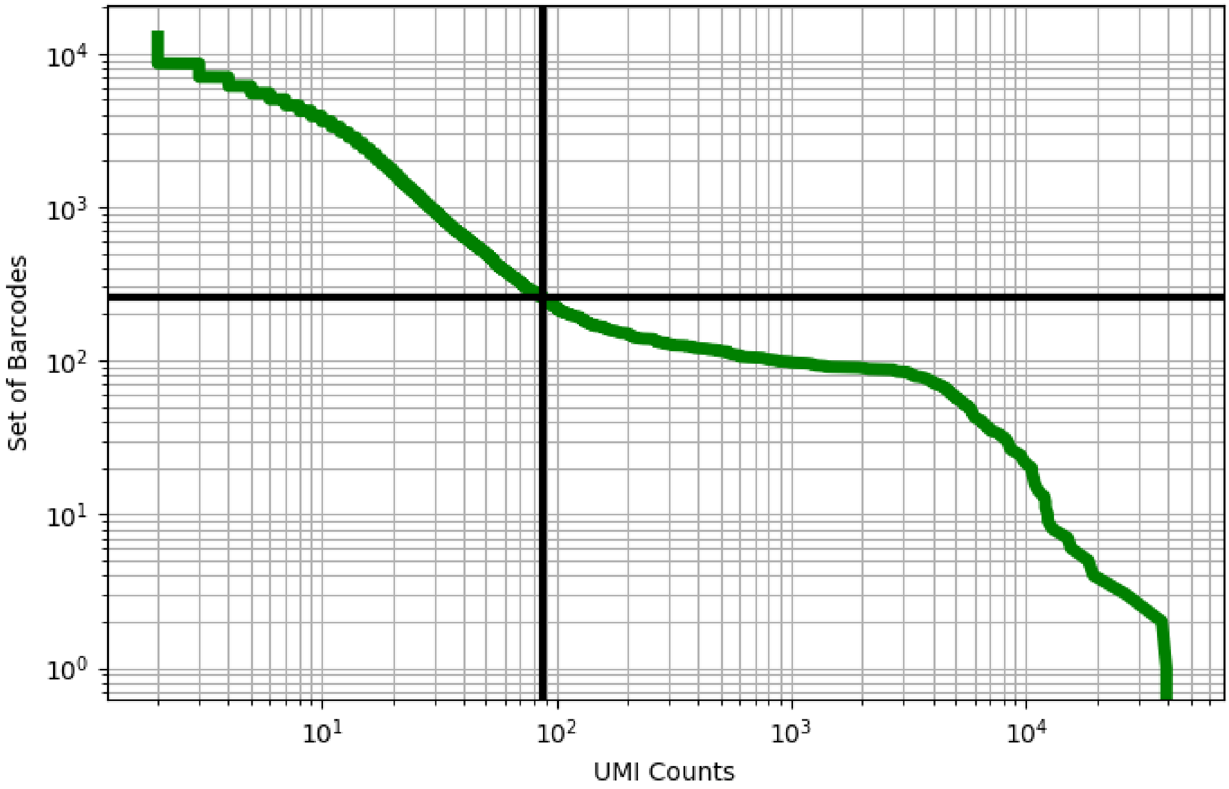


Figure 519: tonsil/GSM5051498

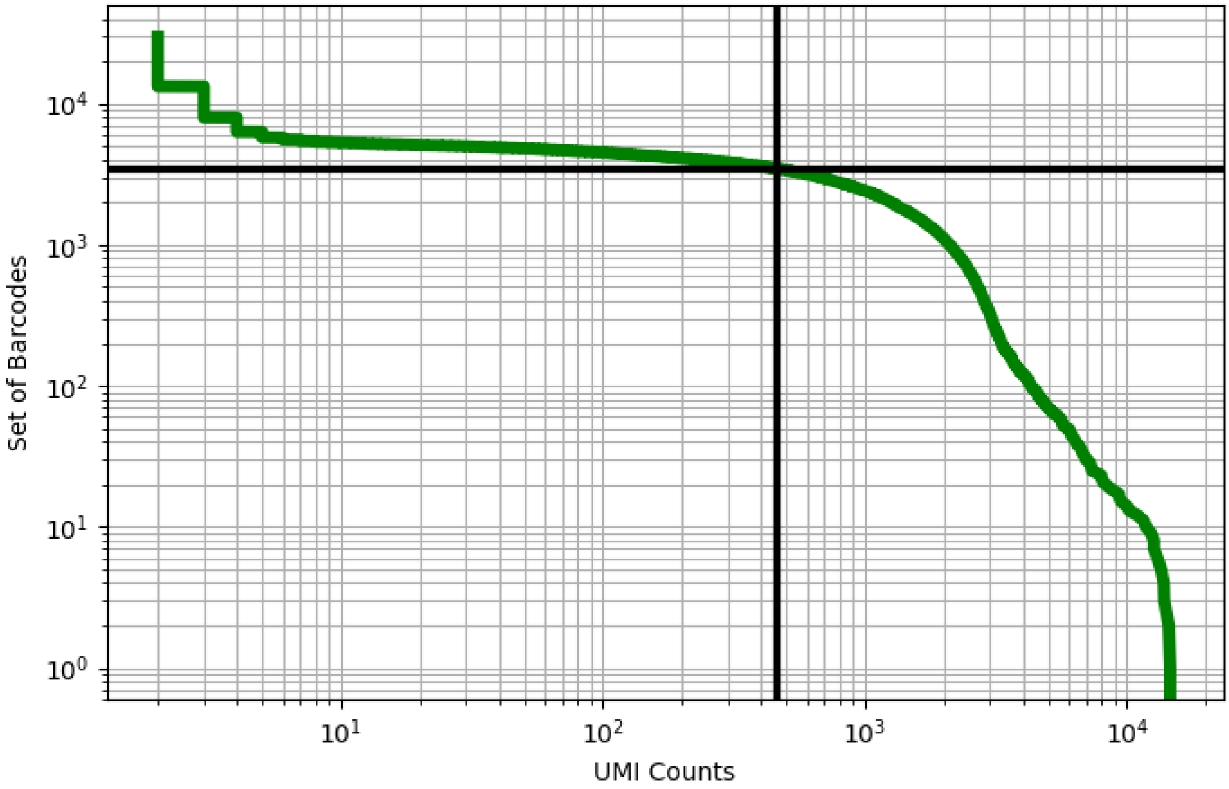


Figure 520: yolk_sac/ERS3861813

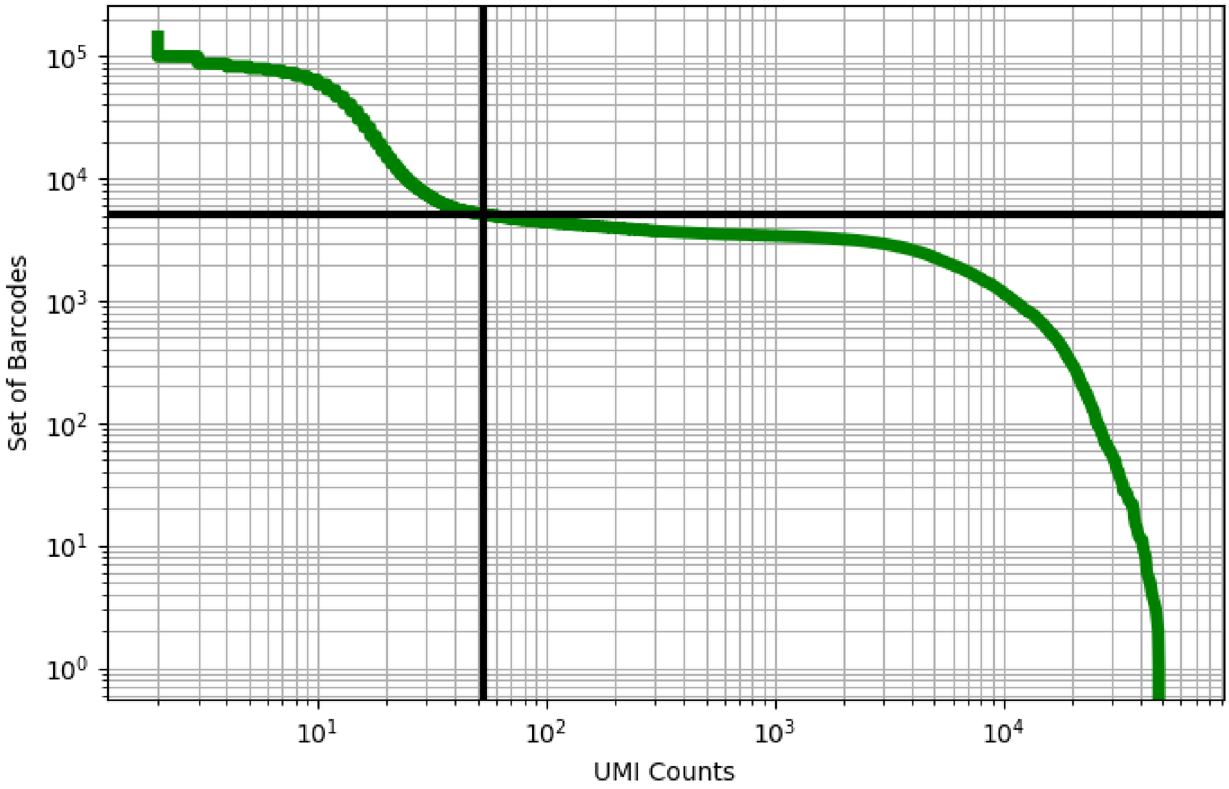


Figure 521: yolk_sac/ERS3861831

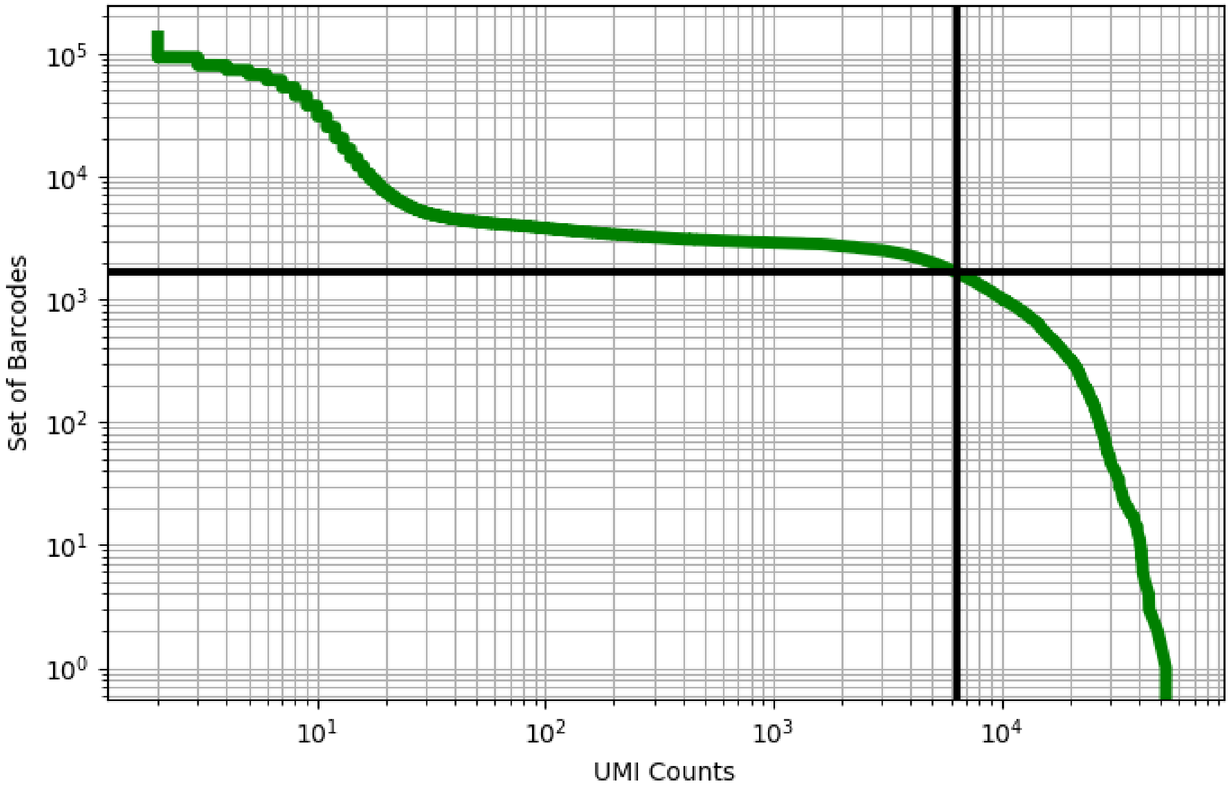


Figure 522: yolk_sac/ERS3861832

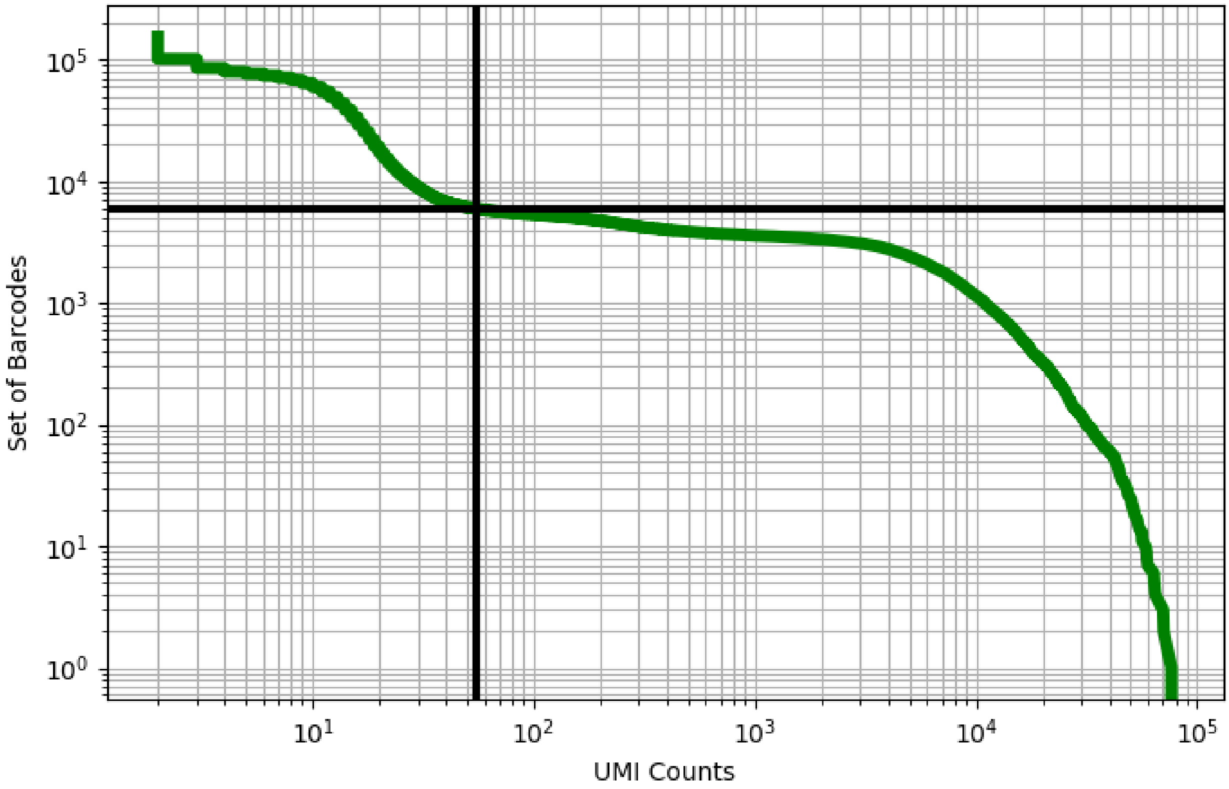


Figure 523: yolk_sac/ERS3861833

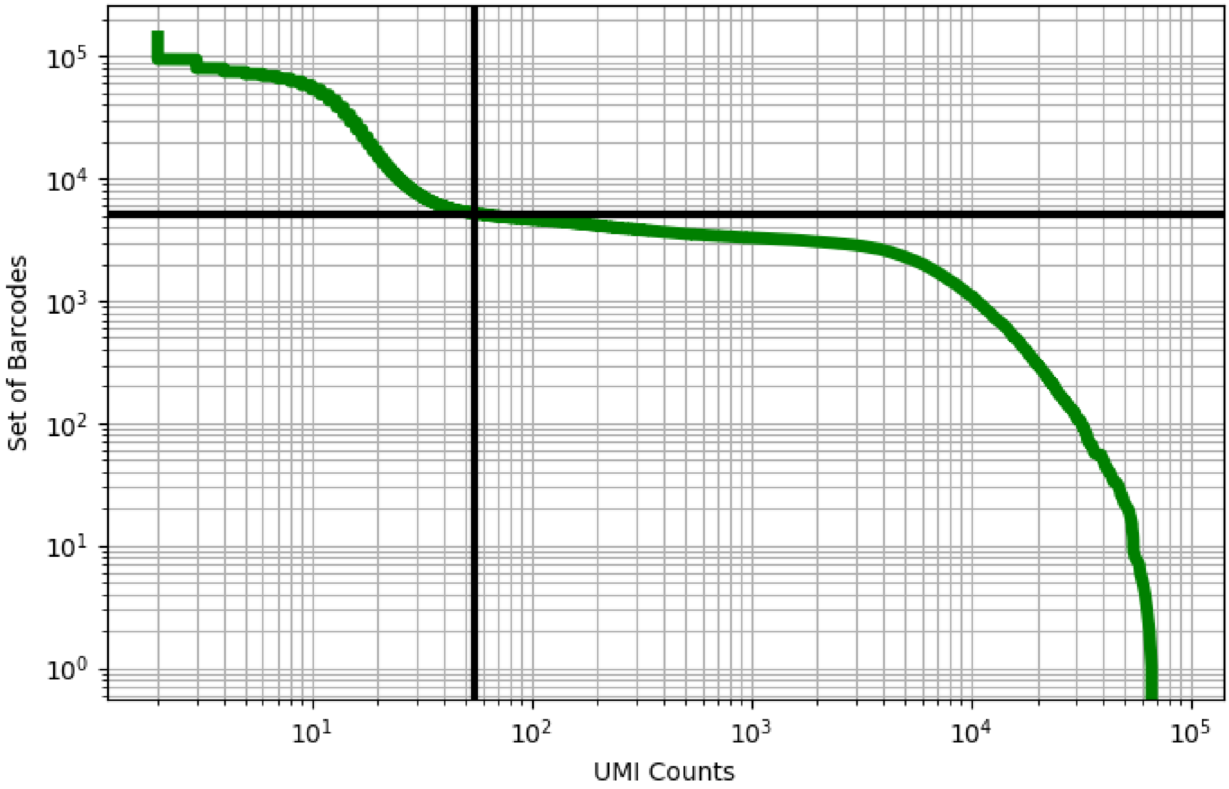


Figure 524: yolk_sac/ERS3861834

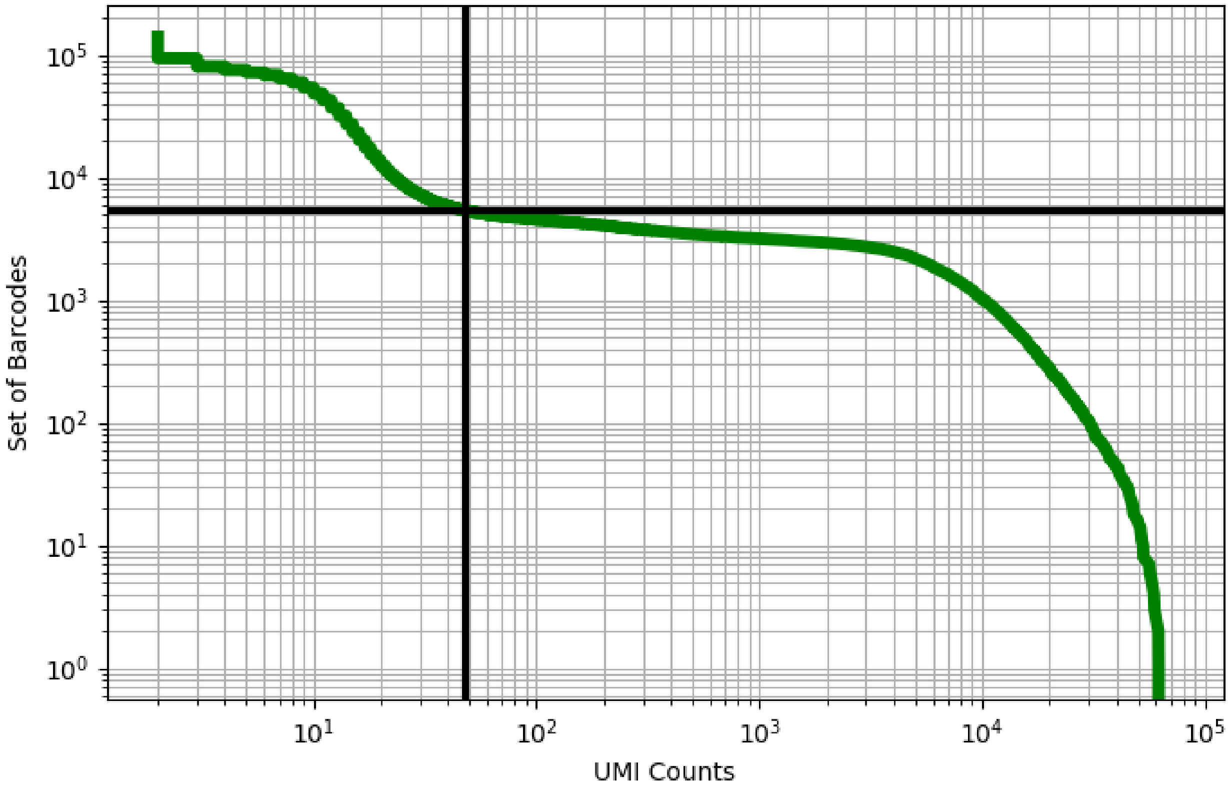


Figure 525: yolk_sac/ERS3861835
